# Loss of the RNA helicase *Dhx15* impairs endothelial energy metabolism, lymphatic drainage and tumor metastasis in mice

**DOI:** 10.1101/2020.11.03.366286

**Authors:** Jordi Ribera, Irene Portolés, Bernat Córdoba, Juan Rodríguez-Vita, Gregori Casals, Bernardino González de la Presa, Mariona Graupera, Guadalupe Soria, Raúl Tudela, Anna Esteve-Codina, Guadalupe Espadas, Eduard Sabidó, Wladimiro Jiménez, William C. Sessa, Manuel Morales-Ruiz

## Abstract

DHX15 is an ATP-dependent RNA helicase involved in pre-mRNA splicing and a downstream substrate for Akt1, which plays a significant role in vascular biology. The aim of this study was to explore the regulatory function of DHX15 over the vasculature and endothelial cell biology. **Results:** DHX15^-/-^ was lethal in mouse and zebrafish embryos. DHX15^-/-^ zebrafish also showed an undeveloped parachordal line, which leads to the formation of lymphatic structures. DHX15^+/-^ mice triggered lower vascular network density and impaired lymphatic function postnatally. Transcriptome and proteome analysis of DHX15 silenced LEC revealed alterations in the glycolysis and gluconeogenesis pathways. The validation of these results demonstrated an uncoupling of the glycolysis with the oxidation of pyruvate in the mitochondria and a lower activity of the Complex I, resulting in lower cellular ATP production. Noteworthy, DHX15^+/-^ mice partially inhibited primary tumor growth and reduced lung metastasis after injection of LLC1 tumor cells.

## INTRODUCTION

RNA helicases are highly conserved and widespread enzymes that play a fundamental role in RNA metabolism through the control of basic RNA processes such as ribosome formation, pre-mRNA maturation, nuclear transportation, RNA translation, transcription initiation, degradation and folding of RNA (Patel and Donmez, 2006). They also play an essential role in the detection of viral RNAs (Jankowsky et al., 2011). RNA helicases are classified into two different subfamilies: the DEAD-box family (DDX), and the DEAH-box (DHX) family (Jankowsky et al., 2011; Umate et al., 2011). Specifically, the DHX family consists of 16 members, which have been identified based on their homology within the amino acid sequences of the helicase domain (Suthar et al., 2016; Umate et al., 2011).

RNA helicases act by remodelling RNA structures through ATP hydrolysis, which exerts a mechanical force resulting in the alteration of the RNA configuration that is fundamental for many cellular processes. This RNA reconfiguration is due to the translocation of the helicase along the RNA which unwinds RNA duplexes and dissociates bound proteins (Bourgeois et al., 2016; Chen et al., 2001; Jankowsky et al., 2001). It has been recently discovered that since DHX helicases lack target selectivity they require the action of adapter proteins, such as G-patch proteins, to aid in the recruitment of RNA targets to the functional site of the helicase. These kind of DHX activators stabilize a functional conformation with high RNA affinity enhancing the catalytic activity of the helicase (Studer et al., 2020).

In recent years, RNA helicases have gained notoriety due to their role in cell maintenance, controlling many biological processes, including cell differentiation and apoptosis (Jiang and Wu, 1999). Also, several groups have linked the defects in helicase functioning with cancer, infectious diseases, and neurodegenerative disorders (Jankowsky et al., 2011). For instance, recent studies have shown that the deregulated expression of an increasing number of these enzymes usually appears in many types of tumors (Abdelhaleem, 2004; Robert and Pelletier, 2013), hence being related to carcinogenesis and cancer progression (Abdelhaleem et al., 2003; Fuller-Pace, 2013, 2006; Heerma van Voss et al., 2017; Steimer and Klostermeier, 2012). Despite these initial studies, the understanding of the pathological mechanisms driven by the RNA helicases and their individual specificity or redundancy over their molecular targets is still limited.

DHX15 is a newly identified member of the DEAH-box RNA helicase family, located in the cell nucleus that regulates pre-mRNA maturation (Fouraux et al., 2002; Niu et al., 2012). DHX15 is known to contribute to ribosome biogenesis by participating in some steps of the small subunit maturation, and in splicing by dissociating the spliceosome modules after completion of its function (Arenas and Abelson, 1997; Combs et al., 2006; Martin et al., 2002; Tsai et al., 2005). DHX15 has ubiquitous variable expression in healthy tissues and organs (Imamura et al., 1997). This helicase is also present in retinal endothelial cells that line the arborizing microvasculature in the human retina (Bharadwaj et al., 2013). In pathological situations, some studies have shown that DHX15 expression can be dysregulated due to exacerbated autoimmune response (Mosallanejad et al., 2014; Wang et al., 2016) and different types of cancer (Albrecht et al., 2004; Lin et al., 2009; Nakagawa et al., 2006; Pan et al., 2017). The serine/threonine kinase Akt1 plays an essential role in vascular biology as a central signaling node that coordinates major cellular processes (Chen et al., 2008; Lee et al., 2014; Pauta et al., 2016). Its primary signaling function relays on the fact that Akt1-dependent phosphorylation leads to the regulation of critical mediators that control different cellular processes, including cell death, cell growth, and chemotaxis. In a previous study, we demonstrated that only the Akt1 isoform can phosphorylate DHX15 in mouse lung endothelial cells (Lee et al., 2014). This observation led us to hypothesize that DHX15 contributes to some extent to the vascular functions of Akt. Therefore, the goal of the present study was to characterize the vascular phenotypes and the pathological mechanism associated with the DHX15 gene deficiency generated by gene editing of this enzyme in mice and zebrafish.

## RESULTS

### Homozygous loss of DHX15 gene is associated with embryonic lethality in mice

The physiological and pathophysiological effect of perturbations in the DHX15 gene has not previously been assessed due to the lack of genetically modified experimental models for this gene. Therefore, we generated a global knockout mouse for DHX15 as described under the Materials and Methods section. Exon 2 of the *DHX15* gene was selected as TALEN target site for knockout mouse production, resulting in two different DHX15-deficient mouse clones: Mouse-ID#35, and Mouse-ID#39. Both clones were used interchangeably in the subsequent experiments without detecting significant differences in the phenotypes or the results obtained (Fig. 1A).

**Figure 1.**
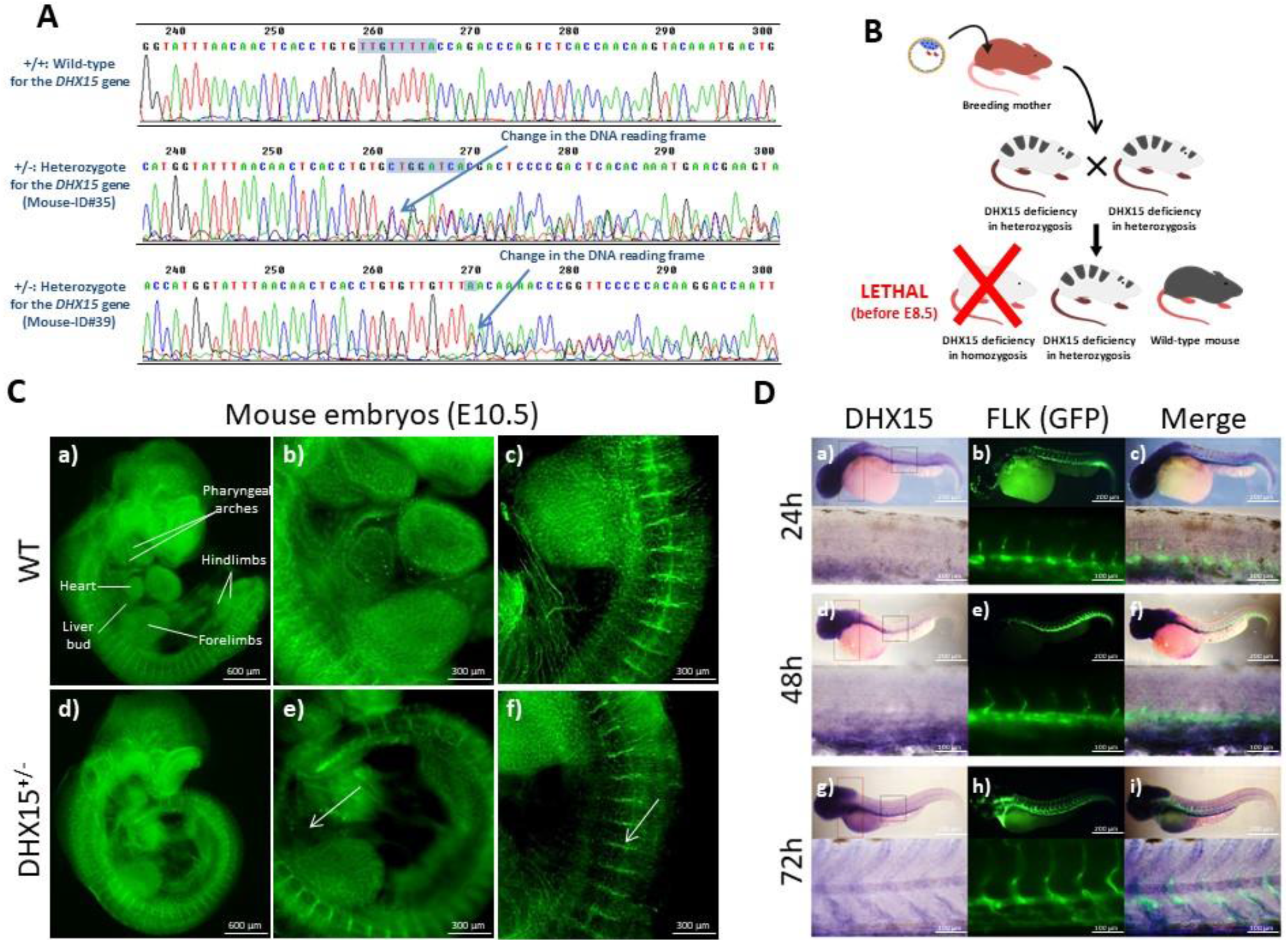
Embryonic characterization of DHX15 gene deficiency and expression in the gene-edited mouse and zebrafish models. (A) Comparison of sequencing chromatograms from wild-type and the heterozygote mouse of the *DHX15* gene obtained from the clones Mouse-ID#35 and Mouse-ID#39. The transcription activator-like effector nuclease technology (TALEN) target site is highlighted within the box. The arrow shows the first base from which the DNA reading frame undergoes nonsense-mediated decay. (B) Scheme showing the transgenic mouse production, from TALEN RNA injection in pronucleated oocytes. (C) Representative immunostaining of the vasculature with endomucin (green) from mouse embryos at the stage E10.5 of embryonic development. The white arrows denote areas of decreased vascular density (n=6). Maximal projection and 3D rendering from the microscope are showed for each genotype. Original magnification: 20X and 40X. (D) Representative results obtained from in situ hybridization using a labelled complementary RNA strand to localize the specific DHX15 sequence on whole-mount zebrafish embryos. DHX15 (blue) and vasculature (FLK1:EGFP; green) in zebrafish embryos at 24, 48 and 72h of post-natal development. Merged panels show DHX15 and vasculature (green) colocalization (n=15).

Heterozygous DHX15 (DHX15^+/-^) mice were viable without any apparent phenotypic abnormalities but intercrosses between DHX15^+/-^ showed no viable homozygous mice (DHX15^-/-^). To establish the embryonic lethality period, timed pregnancies of DHX15^+/-^ breeding were examined at embryonic day (E) 8.5. No DHX15^-/-^ embryos were obtained at this time point suggesting post-implantation embryonic lethality before E8.5 (Fig. 1B).

### DHX15 gene deficiency causes blood and lymphatic vascular defects during embryonic stages

Previous studies demonstrated the role of Akt in vascular development (Chen et al., 2008; Lee et al., 2014). To investigate whether DHX15, as a downstream target of Akt, contributes to these vascular defects we performed whole-mount blood vessel staining followed by 3D visualization in E10.5 mouse embryos to quantify potential vascular abnormalities. DHX15^+/-^ embryos do not exhibit significant defects in segmentation. However, heterozygous embryos showed lower vascular density compared with wild-type (Fig. 1C). This deficiency is mainly evident in the heart region and the intersomitic arteries that sprout out or are located between the somites.

Early embryonic lethality caused in mice by DHX15 deficiency limits the characterization of embryonic vascular anomalies motivated by this genetic disturbance in homozygosis. In order to overcome this limitation, we generated a DHX15 gene deficient zebrafish mutant by Crispr/Cas9 editing in a Tg(*flk1*:EGFP) background, as described in the material and methods section. Wild-type zebrafish embryos showed DHX15 mRNA expressed broadly across the larvae on 5 day post fertilization (dpf) (Fig. 1D, panels a, d and g) with an enriched expression in the vascular system, specifically in the dorsal aorta and the intersegmental vessels (Fig. 1D panels c, f and i), as post-natal development progresses (24, 48 and 72 hours). DHX15^-/-^ zebrafish embryos at the stage 5 dpf were also screened for the expression of GFP in the vasculature. DHX15 deficiency caused vascular development impairment in primary arteries and veins, compared with the wild-type embryos. This impairment was characterized by generalized dilatation of the vasculature, especially in the cardinal vein and the intersegmental vessels (ISV) (Fig 2A). These defects were extended to developing lymphatic structures, such as the parachordal line (Jung et al., 2017). In control embryos, the parachordal line was detected in nearly every somite segment (Figure 3A, arrows). By contrast, the number of parachordal lines was 63% lower in the DHX15^-/-^ embryos.

**Figure 2.**
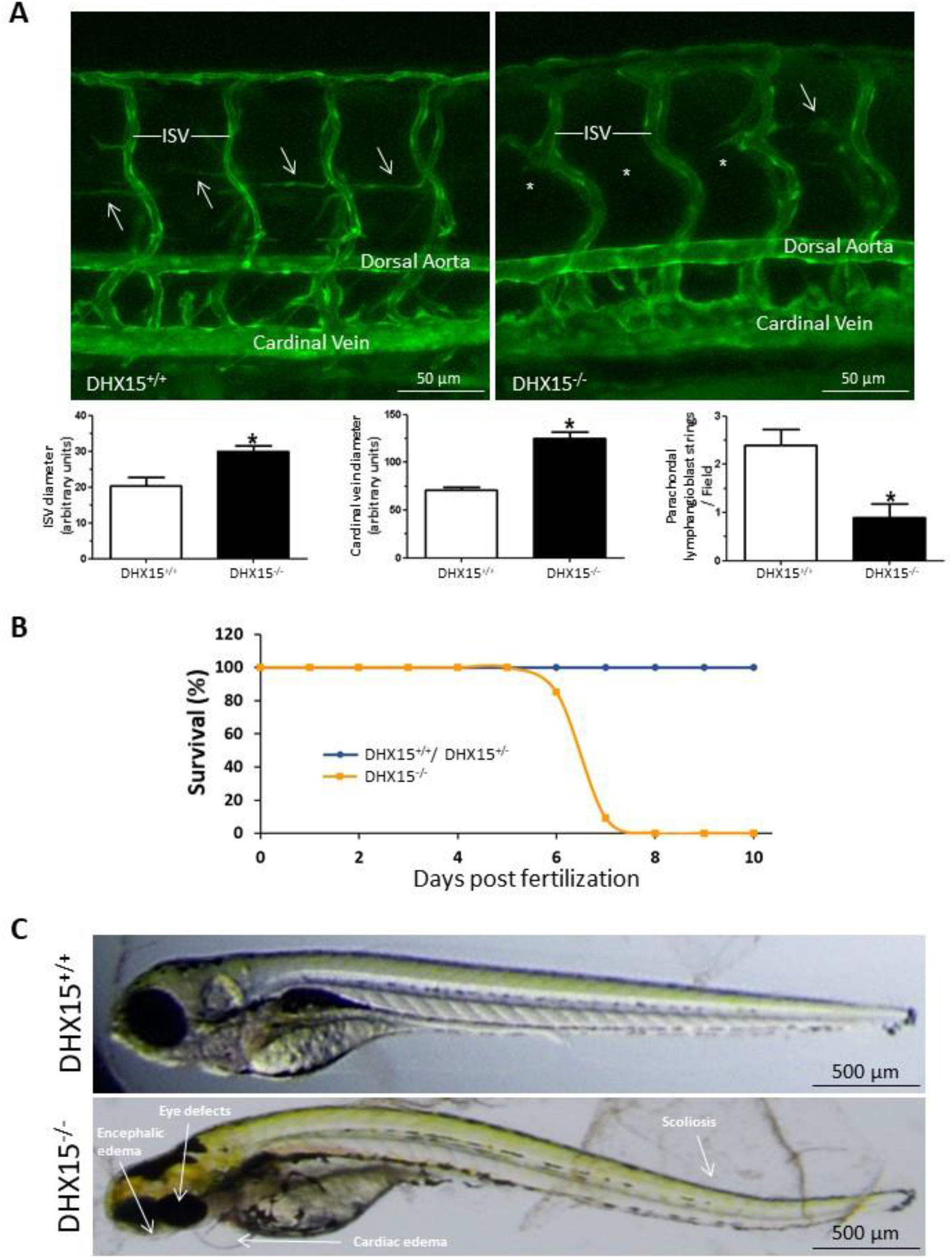
Characterization of embryonic vascular anomalies motivated by DHX15 gene deficient in zebrafish. (A) Representative vascular images of DHX15^+/+^ and DHX15^-/-^ larvae at 5 day post fertilization (dpf) revealing a reduced formation of the parachordal line (arrows). Asterisks denote the absence of these vascular structures in DHX15^-/-^ animals. Quantifications of cardinal vein diameter, intersegmental vessels (ISV), and number of parachordal structures are shown in the graphs; \**p*<0.01 vs. wild-type zebrafish. (B) Survival assessment assay. The graph shows the larvae survival rate through the first 10 dpf according to their different genotype (n=15). (C) Representative images comparing wilt-type and DHX15^-/-^ larvae at 7 dpf where morphological defects including encephalic and cardiac edema, scoliosis, and impaired neural/eye growth are evident.

**Figure 3.**
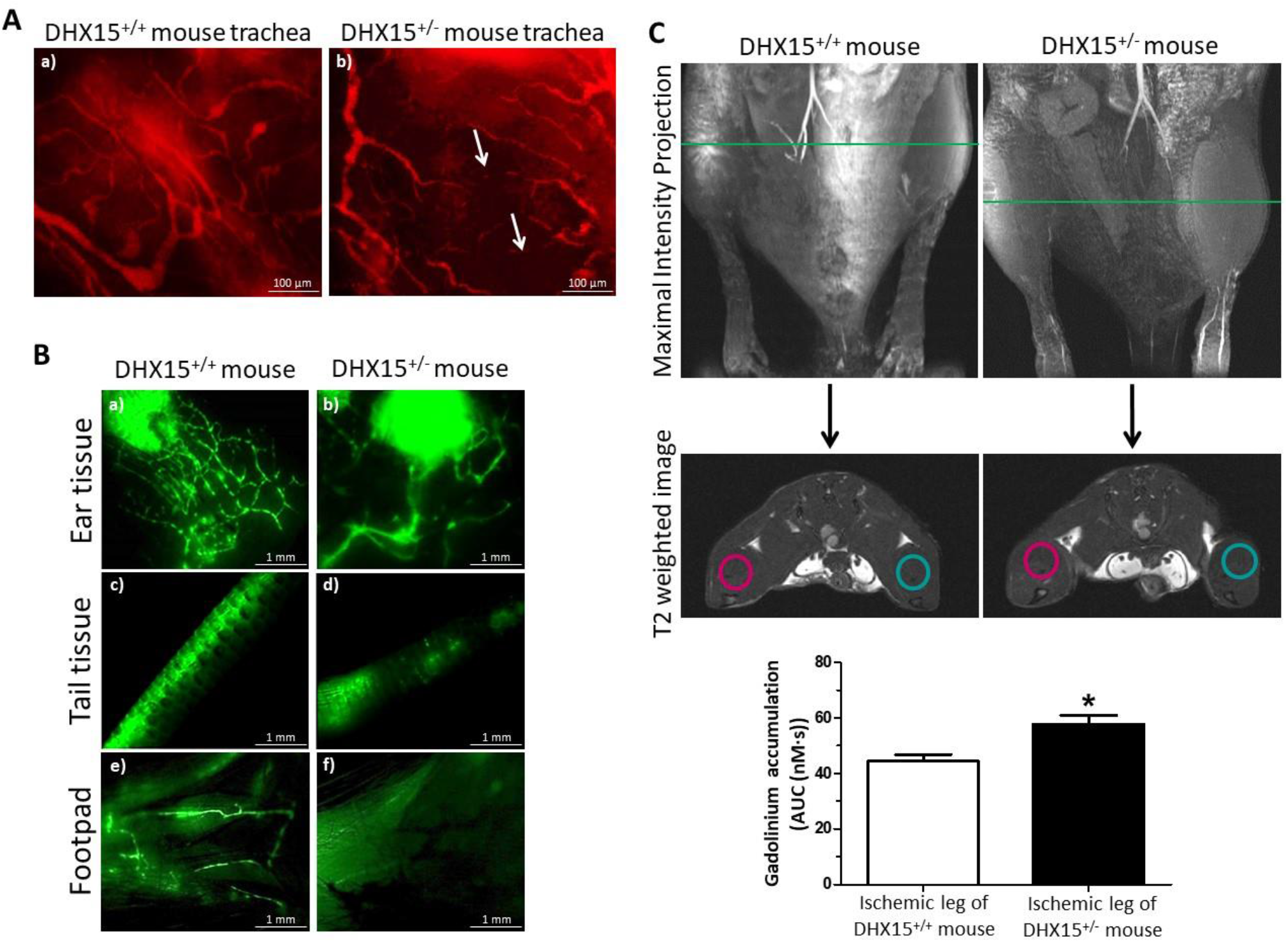
DHX15^+/-^ mice showed cardiovascular and lymphatic vasculature alterations. (A) Representative immunofluorescent images (red CD31 staining) of mouse trachea vessels. White arrowhead evidences lack of connectivity between large vessels (n=5). (B) Lymphatic drainage of 2000 KDa FITC-dextran analyzed by lymphangiography. Fluorescent dye was injected intradermally in the ear (panels a and b), in the interstitium of the tail-tip (panels c and d) and in the footpad (panels e and f) to assess lymphatic uptake (n=5). (C) Representative magnetic resonance images (MRI) for both strains of mice. First row shows the maximal intensity projection of the time of flight (TOF) angiography. The green line indicates the position of the coronal image (second row: T2 weighted image) where the regions of interest (ROIs) for the analysis of the dynamic contrast enhanced-MRI experiment where positioned. In blue, ROIs for the control leg, in red, ROIs for the ischemic leg. The lower graph shows the area under the concentration curve (AUC) calculated for the ischemic leg in WT and DHX15^+/-^ mice; \**p*<0.05 vs. wild-type mouse (n=8).

Similar to mice, completed DHX15 deficiency was lethal in zebrafish. To establish the embryonic lethality period, we monitored the larvae mortality during the first 10 dpf (Fig. 2B). The DHX15^-/-^ larvae started to die at 6 dpf and reached a 100% mortality at the stage 8 dpf. These animals developed morphological defects including encephalic and cardiac edema, scoliosis, and impaired neural/eye growth; compared to wild-type embryos (Fig. 2C) and the onset of these morphological defects was at 3 dpf. In contrast, DHX15^+/+^ and DHX15^+/-^ did not show significant differences in survival throughout embryonic development or postnatally.

### Vascular density and lymphatic functionality are impaired postnatally in DHX15^+/-^ mice

Due to the mortality associated with the lack of DHX15 in homozygotes, it was only possible to characterize the vascular role of DHX15 in viable adult heterozygotic mice (DHX15^+/-^). DHX15^+/-^ mice showed significant vascular malformations compared with wild-type mice. Whole-mount preparations of trachea tissue for the quantification of vessel pruning demonstrated that DHX15^+/-^ mice exhibited reduced vascular densities and an impaired connectivity between large vessels (arrows), compared with littermate WT mice (Fig. 3A).

We also evaluated lymphatic function in adult DHX15^+/-^ mice by two different methodologies. First, fluorescent lymphangiographies were performed in peripheral regions using FITC-dextran. The lymphatic vasculature of DHX15^+/-^ mice depicted diminished fluid drainage, compared with WT mice in three different peripheral tissues: the ear, the tail and the foodpad (Fig. 3B). Second, we quantified lymphatic drainage of the contrast agent gadolinium in the hindlimbs after femoral artery ligation. This model is associated with increased vascular permeability and the consequent accumulation of fluid in interstitial spaces. For this purpose, we performed MRI in WT and DHX15^+/-^ mice, 4 weeks after the femoral artery ligation. In the ischemic legs (red circles Fig. 3C), all the DHX15^+/-^ mice showed impaired lymphatic drainage with the consequent significant increase in gadolinium accumulation, as observed by the increase area under the concentration curve calculated in DHX15^+/-^ mice, compared to control animals (57.72±3.19 vs. 44.45±2.35 nM·s, respectively; *p*<0.05).

### Mechanistic insights of DHX15 deficiency in mouse endothelial cells. Role of DHX15 in carbohydrate metabolism

The *in vivo* experiments documented that reduced levels of DHX15 caused cardiovascular and lymphatic abnormalities. To obtain information on the mechanisms responsible for these phenotypes in both vascular systems, we performed RNAseq and proteomics in endothelial cells with or without DHX15 gene silencing using a LEC cell line engineered to express the Tet-On^®^ induction system for silencing DHX15 (siL-DHX15-LEC, see material and methods). The advantage of using this cell line is that these cells maintain both cardiovascular and lymphatic endothelial cell characteristics, enabling the potential extrapolation of the mechanistic findings into both vascular systems (supplemental figure 1). For instance, LEC displayed endothelial cell markers (CD31, eNOS and uptake of oxidized low-density lipoprotein; supplemental figure 1A and C) but also classical lymphatic cell markers (LYVE-1 and podoplanin; supplemental figure 1B and C, respectively). Supplemental figure 1D documents the total DHX15 levels in the starting cells with or without DHX15 silencing that were used for genomic and proteo-genomics experiments. The RNA-seq experiment allowed us generating 122.968 paired-end reads that were successfully mapped to the mouse reference GRCm38 genome assembly. After quality control analysis, four samples per experimental condition were included in the analysis. A total of 5,408 isoforms were differentially expressed when considering a stringent threshold of FDR <0.001 (Supplementary Table S1). For the proteomics experiment, protein extracts (n=5 for each experimental condition) were digested into peptides and they were quantified by mass spectrometry analysis, as described in material and methods. We prioritized those protein variations that showed a p-value lower than 0.05 or when the protein was present in one condition, and undetectable in all the cases of the other experimental condition. The list of proteins with relevant changes in their abundance due to the silencing of DHX15 are detailed in Supplementary Table S2.

In an effort to understand the biological relevance of the -omis results, we combined the results of the RNAseq and proteomic experiments and subjected the final list to pathway analysis using the Ingenuity Pathway Analysis (IPA; Ingenuity) database. The results were compared against global molecular networks to identify associated diseases and canonical pathways affected by the perturbation of DHX15. The DHX15 silencing significantly affects two networks: 1) endocrine system disorders, organismal injury and abnormalities and cancer, and 2) gastrointestinal disease, organismal injury and abnormalities and carbohydrate metabolism (Supplemental figure 2, A and B). Also, the reduction of DHX15, was associated with differential expression in nine additional signalling pathways (Supplemental figure 3A). Among them, the glycolysis and gluconeogenesis pathways.

To functionally validate these bioinformatic results, we measured the glycolytic activity and ATP generation in LEC cells. In accordance to our findings, siL-DHX15-LEC cells showed higher levels of glycolytic activity (measured as nM of L-Lactate) compared to cells with the DHX15 wild-type gene (0.41±0.04 vs. 0.19±0.05 nM L-lactate, respectively; *p*<0.05) (Fig. 4A). The alterations in glucose metabolism was linked to lower ATP production compared to WT LEC cells (417.7±31.37 vs. 524.2±20.40 nM ATP, respectively; *p*<0.05) (Fig. 4B), suggesting uncoupling between glycolysis and oxidative phosphorylation. In this context, the rMATS analysis of alternative 5’ splice site from the RNAseq data identified 220 significant splicing event modifications (with FDR<5%, absolute difference >5% and >70 number of reads) caused by DHX15 silencing (Supplementary Table S3). Among these genes, NADH ubiquinone oxidoreductase core subunit S1 (NDUFS1) showed a differential alternative splicing characterized by the presence of an alternative 5’ splice site, giving rise to a longer exon 1 for NDUFS1, when DHX15 was silenced in LECs (Fig. 4C). This gene encodes a subunit of the mitochondrial respiratory chain complex I. In agreement with this observation, the in-gel activity measurement of complex I was reduced significantly by ~50% in the siL-DHX15-LEC condition, compared with non-silenced LEC cells (0.50±0.04 vs. 1.00±0.03 relative units, respectively; *p*<0.01) (Fig. 4D).

**Figure 4.**
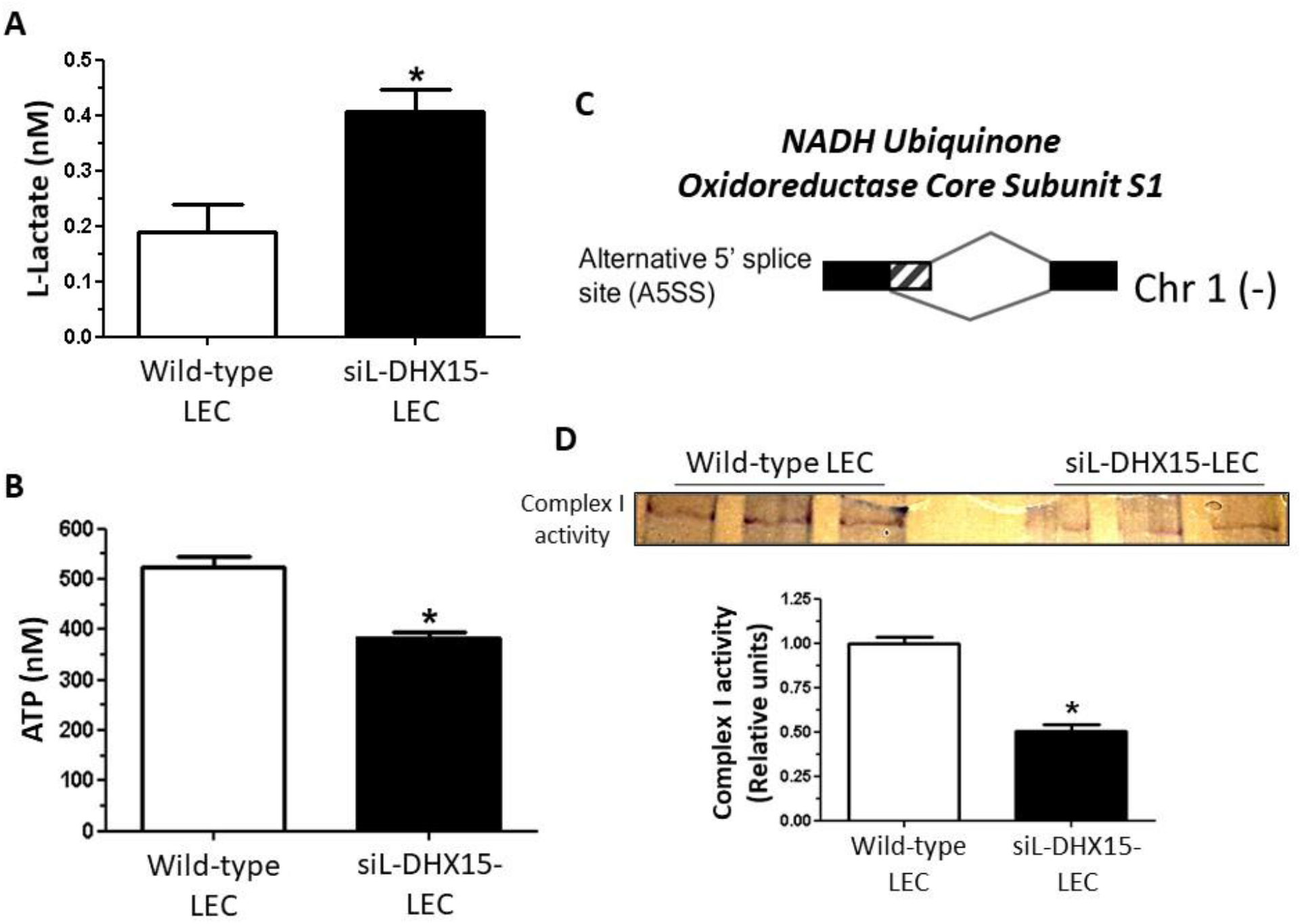
Uncoupling of aerobic glucose metabolism in siL-DHX15-LEC. (A) Glycolysis levels were evaluated by a colorimetric enzymatic reaction in wild-type and silenced DHX15 liver endothelial cells (siL-DHX15-LEC) as nM of L-Lactate; \**p*<0.01 *vs*. wild-type LEC (n=6). (B) ATP production was evaluated by a luminescence assay in wild-type and siL-DHX15-LEC; \**p*<0.05 *vs*. wild-type LEC (n=6). (C) Alternative splicing was quantified as described in Material and Methods. The diagram shows the significant splicing event occurring on the gene NDUFS1 with an inclusion level of 6% and a FDR=0.02. The striped bar represents the lengthening of the alternative 5’ limit size of exon 1 caused by the DHX15 silencing. (D), *In-gel* activity staining on clear-native page (CN-PAGE) of the respiratory complex I from wild-type and siL-DHX15-LEC’s mitochondria. The quantification of the relative band intensities of complex I activity is shown in the graph below; \**p*<0.01 vs. wild-type LEC (n=6).

Next, to assess the impact of reduced energy biosynthesis in siL-DHX15-LEC, we quantified cell proliferation and migration. Silencing of *DHX15* reduced BrdU uptake compared to control LEC (55.53±0.49 vs. 66.43±0.46 % of cells BrdU positives, respectively; *p*<0.01) (Fig. 5A). Accordingly, the reduction of DHX15 resulted in an impaired cell migration. Silenced DHX15-LECs presented delayed wound-healing after 24 hours compared with control LEC, as measured in a scratch-induced directional wound-healing assay (Fig. 5B).

**Figure 5.**
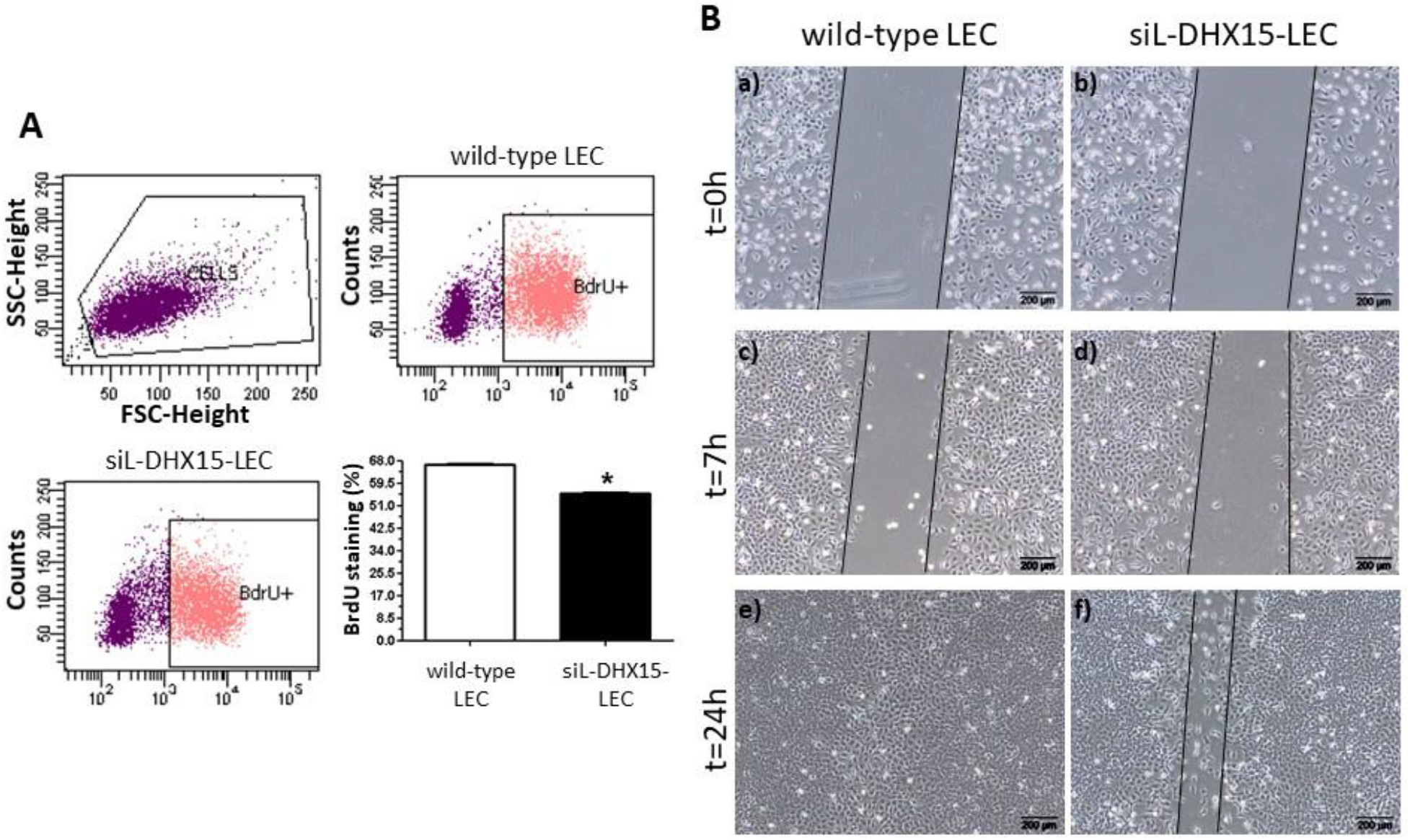
siL-DHX15-LEC presented less cell migration and proliferation. (A) Representative figures of the proliferation assay performed in wild-type and silenced DHX15 liver endothelial cells (siL-DHX15-LEC). Bromodeoxyuridine (BrdU) incorporation was quantified by flow cytometry. First panel shows a dot-blot graph of the cell population. Cells within the oval scatter gate were analyzed. The negative control population was chosen from cells cultured in the absence of BrdU. The percentage of cells that stained positively for BrdU for each experimental condition is depicted in the bar graph; \**p*<0.01 *vs*. wild-type LEC (n=6). (B) Cell migration was quantified after performing a scratch wound in confluent non-silenced and silenced LECs cells that were cultured in 6-well plates. Then images of wound healing were acquired after 0, 7 and 24 hours (n=6).

### Heterozygous *DHX15* gene deficiency reduced tumor growth and metastases

Abnormal function of blood and lymphatic vessels plays a pathological role in multiple pathological conditions including inflammation and cancer. Also, there is a strong link between the lymphatic vasculature and tumor spread, as lymphatic vessels constitute one of the main routes of metastasis in most cancers (Achen and Stacker, 2008; Mumprecht and Detmar, 2009; Sleeman and Thiele, 2009). Considering also that one of the pathways modified by the reduction of DHX15 was “endocrine system disorders, organismal injury and abnormalities and cancer”, we aimed at evaluating the role of DHX15 in tumor development.

First, we explored how the loss of DHX15 affected growth of tumors implanted into WT and DHX15^+/-^ mice by injecting syngeneic LLC1 cells (1×10^5^) into the flanks of both strains. We measured the volume of the primary tumors implanted in these mice 21 days post-injection of the LLC1 cells. Primary tumors were significantly smaller in DHX15^+/-^ mice compared with controls (0.54±0.07 vs. 1.06±0.17 cm^3^, respectively; *p*<0.01) (Fig. 6, panels a and d). Tumors from DHX15^+/-^ mice also showed an impaired vascular network characterized by smaller vascular perimeter, compared with WT littermates (199.1±9.81 vs. 312.5±17.16 μm perimeter, respectively; *p*<0.01) (Fig. 6, panels b and e). Since growth of the primary tumor is rate-limiting and precludes analyses of metastasis in this model, we established a postsurgical metastasis model. Three weeks after subcutaneous LLC1 cells implantation, primary tumors were completely resected and postsurgical lung metastasis were evaluated 2 weeks later. Fewer primary tumors from DHX15^+/-^ mice metastasized the lungs compared with wild-type mice (*p<0.01*) (Supplemental figure 3B). Noteworthy, and considering the mice with lung metastasis, the overall area of metastases was strongly decreased in DHX15^+/-^ mice, compared with the wild-type group (0.88±0.32 vs. 2.53±0.69 % lung metastases, respectively; *p*<0.05) (Fig. 6, panels c and f).

**Figure 6.**
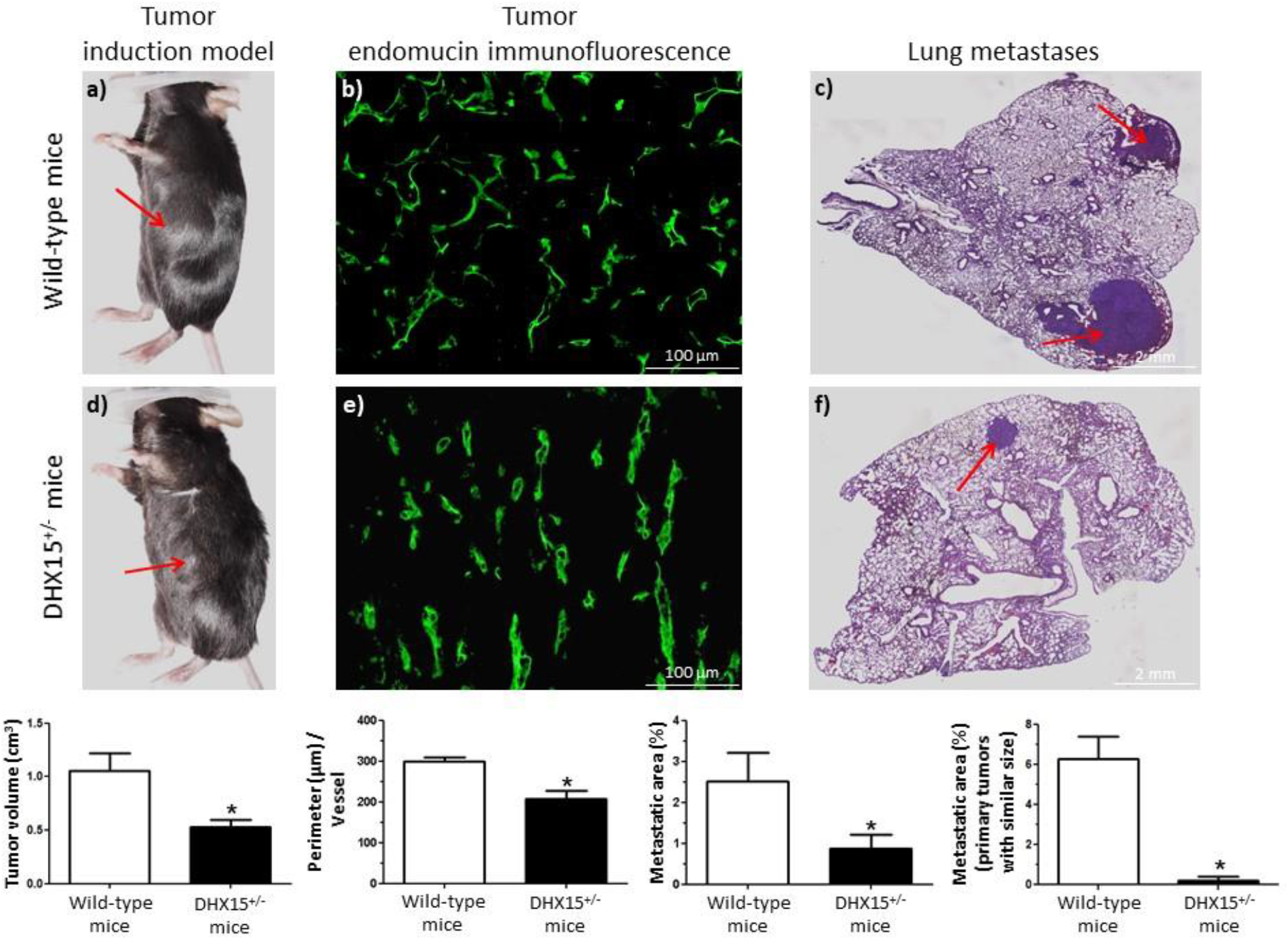
Tumor growth and metastases in DHX15^+/-^ mouse. (A) Macroscopic images of tumor size in wild-type and DHX15^+/-^ mice 3 weeks after mouse Lewis lung cancer cells (LLC1) implantation. The arrows indicate the primary tumor. The quantification of tumor volume (cm^3^) is shown on the lower graph; \**p*<0.01 vs. wild-type mice (n=15). (B) Endomucin immunostaining of intratumoral blood vessels in wild-type and DHX15^+/-^ mice. Quantification of vessel perimeter is shown in the graph below; \**p*<0.01 *vs*. wild-type mice (n=15). Original magnification: 200X. (C) Representative lung sections and quantification of lung metastatic area after haematoxylin-eosin staining (H&E) (FIJI software analysis) are shown. The arrows indicate the metastatic areas. Quantifications of the percentage of lung metastases are shown in the graphs below; all tumors \**p<0.05 vs*. wild-type mice (n=15) and primary tumors with similar size, \**p*<0.01 *vs*. wild-type mice (n=3). Original magnification: 10X.

To investigate further whether the differences in the size of lung metastasis were due to a potential lower LLC1 cell sedding in the primary tumors, we followed over time a group of mice with similar primary large-size tumors. In this experimental group (size of primary tumors ranging from 0.81 to 1.13 cm^3^), we also found significant differences when we compared the areas of lung metastasis between wild-type and DHX15^+/-^ mice, being significantly lower in the DHX15^+/-^ mice (6.27±1.12 vs. 0.20±0.20 vs. % lung metastases, respectively; p<0.01) (Fig. 6, lower right graph).

## DISCUSSION

In previous studies, we identified DHX15 as a downstream target of Akt1, which has the greatest influence on vascular regulation compared with the other Akt isoforms (Chen et al., 2008; Lee et al., 2014; Pauta et al., 2016). Considering this observation, we hypothesized as a starting point for this study that DHX15 may also play a role in the vascular function. Here, we show that homozygous DHX15 deficiency was associated with embryonic mortality in mice. To determine when lethality occurred, timed pregnancies of DHX15^+/-^ breeding were studied at E8.5. We did not detect any DHX15^-/-^ embryos at this time point, suggesting early lethality post-implantation. On the other hand, the loss of just one DHX15 allele resulted in impaired lymphatic drainage and decreased vascular blood density. These results are in agreement with the reduced cell proliferation and migration that we observed in LEC after silencing the expression of DHX15. Why vasculature is affected by the partial loss of DHX15 in adult mice remains unclear if we only consider the results of this experimental model. Under this experimental setting, we cannot discard the possibility that adult mice developed these vascular defects because of accumulated damage occurring during the postnatal stage in the DHX15 deficiency background. However, and by studying the role of this RNA helicase in DHX15 gene-deficient zebrafish, we showed that the DHX15-related vascular defects are also occurring during development. In this experimental model, we found that DHX15^-/-^ larvae were not viable. Also, these unviable larvae presented blood vascular alterations and reduced formation of developing lymphatic structures, which were associated with cardiac and encephalic edema in the embryos.

Some unaddressed questions are whether each member of the RNA helicases family has unique physiological functions and whether they are used redundantly by the cell. The lethal phenotype of the DHX15^-/-^ mouse and zebrafish embryo supports the concept that RNA helicases may present unique functions and specificity for molecular targets, likely in combination with adapter proteins as demonstrated by Studer et al. (Studer et al., 2020).

Vascular growth and lymphangiogenesis are crucial in tumor growth and metastasis (He et al., 2015; Nishida et al., 2006). Therefore, we assessed the role played by DHX15 deficiency in tumor formation and expansion. After performing syngeneic LLC1 cells allotransplantation in mice, we found that heterozygous DHX15 deficiency partially inhibits primary tumor growth and reduced lung metastases. This effect was associated with an intratumoral reduction in the length of the blood vessel capillaries. Our results agree with several clinical studies that underline a role of DHX15 in tumor progression in several types of cancer, such as acute myeloid leukemia (Pan et al., 2017), hepatocellular carcinoma (Xie et al., 2019), malignant peripheral nerve sheath tumors (Nakagawa et al., 2006), prostate cancer (Jing et al., 2018) and non-small-cell lung cancer (Yao et al., 2019). In all these situations, modifications of DHX15 activity and/or its overexpression favors tumor growth. Several mechanisms have been proposed in these studies to explain the tumorigenicity effect of DHX15, such as the co-activation of the androgen receptor in prostate cancer and the transcriptional activation of NF-kB in leukemia cells. In our study, we provided an additional and a more general mechanism of DHX15 that is relevant for tumor progression: the regulation of the growth and the function of the vasculature.

The use of RNAseq and proteomics, combined with bioinformatics down-stream analysis, is one of the most potent approaches to investigate the role played by RNA helicases, as changes in differential splicing of a gene may or may not be associated with changes in its protein abundance; therefore we need the combined result of both high-throughput approaches. Adopting this strategy, we were able to identify in the DHX15-silenced LEC significant changes in key pathways that metabolize carbohydrates. Some of the genes products that varied were UDP-glucose:glycoprotein glucosyltransferase 1, glyceraldehyde-3-phosphate dehydrogenase, fructose-bisphosphate aldolase A, and pyruvate kinase. This differential expression was also associated with a significant increase in the lactate levels and a reduction in the intracellular concentration of ATP. These metabolic changes suggest a regulatory compensatory effect of the glycolytic enzymes due to the uncoupling of glycolysis with the oxidation of pyruvate into the mitochondria. Supporting this possibility, we observed by RNAseq significant expression changes in members of the family of the NADH dehydrogenase [ubiquinone] 1 alpha subcomplex assembly factor (members from NDUFS1, NDUFS5 and NDUFS7). These members are accessory subunits of the mitochondrial membrane respiratory chain NADH dehydrogenase (Complex I), that transfer electrons from NADH to ubiquinone in the mitochondrial respiratory chain. Among them, we observed a significant alteration in the splicing of the NDUFS1 gene. This alteration in gene splicing was linked with a 50% reduction of the activity of the complex I in the mitochondria of DHX15 silenced cells that may explain the decrease of ATP intracellular levels and, as a consequence, the impairment in endothelial cell proliferation and vascular growth.

Endothelial cells are mainly considered a glycolytic cell type that maintain their energy demands by exploiting the glycolytic pathway preferentially without need of coupling to the tricarboxilic acid cylce and the oxidative phosphorylation (Martinive et al., 2006; Rath et al., 2012). However, some studies support the notion that mitochondria electron transport chain activity is also playing a role in driving endothelium angiogenesis and function. For example, low concentration of oxygen results in the generation of mitochondrial ROS through the Complex III of the electron transport chain (Bell et al., 2007). This increase in mitochondrial ROS regulates angiogenic factors such as HIF-1a, promoting its stabilization and allowing the transcription of the VEGFa gene (Jung et al., 2013; Xia et al., 2007). Also, HUVEC and HDMEC depend on mitochondrial oxidative phosphorylation to maintain energy supplies for proliferation and growth, as demonstrated by measuring oxygen consumption and ATP production in the presence of the mitochondrial uncoupler embelin (Coutelle et al., 2014). Another transport electron chain inhibitor, rotenone that blocks Complex I activity, reduced the angiogenic capacity of vasa vasorum endothelial cells (Lapel et al., 2017). Our study is in line with these observations and supports the idea that for optimal angiogenic response, endothelial cells require dynamic crosstalk between glycolysis and mitochondria activity, driven likely by regulators that promote a metabolic switch, as we have seen when we modified the levels of DHX15.

In conclusion, our study establishes for the first time a vascular and metabolic regulatory role for one RNA helicase member, DHX15, that has a repercussion in pathological processes such as impaired lymphatic drainage and tumor growth. Our results highlight the therapeutic potential of the modulation of DHX15 expression in the context of these diseases. However, our study also raised challenging and exciting questions concerning the function specificity and the redundancy of the RNA-helicase family that need clarification before moving forward in the selection of potential therapeutic targets from this family of enzymes.

## MATERIALS AND METHODS

### DHX15 transgenic mice

*DHX15* gene deficient C57BL/6 mice were generated by genomic editing by microinjecting TALEN (transcription activator-like effector nuclease technology) RNA in pronucleated oocytes (Cyagen). The mouse *DHX15* gene (GenBank accession number: NM_007839.3; Ensembl: ENSMUSG00000029169) is located on chromosome 5. Fourteen exons have been identified for this gene, with the ATG start codon in exon 1 and the TGA stop codon in exon 14. Exon 2 was selected as TALEN target sites. TALENs were constructed using the Golden Gate Assembly method (Cermak et al., 2011) and confirmed by sequencing. The amplicons were then purified and sent for DNA sequencing analysis. TALEN mRNAs generated by in vitro transcription were injected into fertilized eggs for knockout mouse production (cDNA sequence: TGTTGGTGAGACTGGGTC). The pups were genotyped by PCR followed by sequence analysis. The positive founders were breeding to the next generation, which was genotyped by PCR and DNA sequencing analysis. DNA sequencing revealed two different DHX15-deficient mouse clones: Mouse-ID#35 that was missing 8 bases in one strand, and Mouse-ID#39 that was missing 1 base in one strand. Wild-type DNA was used as a negative control for sequencing in parallel. The mRNA transcribed from targeted allele with frameshift undergoes nonsense-mediated decay (NMD).

All animals were kept under constant temperature and humidity in a 12 hours controlled dark/light cycle, and they were fed *ad libitum* on a standard pellet diet. We performed the study following the guidelines of the Investigation and Ethics Committees of the Hospital Clínic and the University of Barcelona.

### Mouse genotyping

Mouse genomic DNA was isolated from tail biopsies using a specific kit (Extract-N-Amp™ Tissue PCR Kit; Sigma-Aldrich, Darmstadt, Germany). PCR was performed using the primer pairs to amplify the *DHX15* gene (primer forward: 5’CACCAACCTGCCCCATACTCCT-3’ and primer reverse: 5’-TGTATTGTCCCAGGGTAAAATGTGTTG-3’). PCR conditions were as follows: 35 cycles at 94°C for 30 s, 59.3°C for 30 s, and 72°C for 60 s. PCR product was sequenced by sanger sequencing to distinguish the *DHX15* wild type mice and *DHX15* transgenic mice.

### Immunological staining of mouse embryo whole-mounts

Embryos were harvested at different points between E8.5 to E10.5. Embryos at E10.5 were fixed in 4% paraformaldehyde overnight at 4°C. For immunostaining of whole mount embryos, after paraformaldehyde fixation, the embryos were sequentially dehydrated in methanol and then incubated in the permeabilization buffer (PBlec) (PBS pH6.8, 1% Tween 20, 1mM CaCl_2_, 1mM MgCl_2_, 0.1 mM MnCl_2_) for 20 minutes at room temperature. After permeabilization, the embryos were incubated with primary antibody rat anti-endomucin (Abcam, Cambridge, UK) (1:20 dilution) in PBlec buffer overnight at 4°C. To remove residual primary antibody, embryos were washed with PBT (PBS pH6.8, 0.1% Tween) for 5×10 minutes. Next, the embryos were incubated with secondary antibody Alexa Fluor 488-conjugated goat anti-rat (Thermo Fisher, Waltham, MA, USA) (1:500 dilution) in the dark overnight at 4°C, and then washed with PBT for 3×10 minutes and postfix in 4% paraformaldehyde. Images were acquired using fluorescence stereomicroscope (Leica Microsystems, Heerbrugg, Switzerland) and immunofluorescence microscope (Nikon Eclipse E600, Kanagawa, Japan) systems.

### Generation of the Zebrafish animal model

Adults wild-type zebrafish (*Danio rerio*), in a Tg(*flk1*:EGFP);Tg(*fabp10*:RFP) background, purchased from KIT - European Zebrafish Resource Center (EZRC), were maintained at 28–29 °C on a light cycle of 14 hours light/10 hours dark. The Crispr/cas9 design for gene knock-out was performed as follows: Gene sequences were retrieved from http://www.ncbi.nlm.nih.gov/gene and http://www.ensembl.org/Danio_rerio/Info/Index. The sgRNA was designed using the online tool http://crispor.tefor.net, based on exon site and high efficacy and off-target published algorithms. Microinjection was performed at 1-cell stage embryos. Fertilized zebrafish embryos were collected in E3 medium. Then, embryos were grown at 28.5 °C.

### Zebrafish whole mount in situ hybridization (ISH)

cDNAs were amplified by PCR from a custom zebrafish cDNAs library obtained by RT-PCR from an mRNA pool coming from 5 days post fertilization zebrafish larvae. We included a SP6 sequence linker in reverse primers to directly use the synthesized PCR products as templates to amplify the reverse riboprobe to be used for ISH. For the ISH, embryos were fixed overnight with 4% PFA and washed twice with PBT. Then, embryos were incubated with hybridation buffer (50% formamide, 5X SSC buffer pH 6 (0.75M NaCl, 0.075M sodium citrate), 0.1% triton, 50 μg/mL yeast RNA, 50 μg/mL heparin) at least 1 hour. Next, embryos were incubated with hybridation buffer containing the reverse riboprobe overnight. Finally, embryos were washed with washing solution (50% formamide, 1X SSC buffer, 0.1% Tween) 30 minutes twice and with MABT (100 mM maleic acid pH 7.5, 150 mM NaCl, 0.1% Tween) 10 minutes five times. Stained embryos were processed for imaging with bright field stereoscope to determine the overall expression pattern.

### Zebrafish vascular characterization

Five-day old larvae obtained by pairwise mating of adult Tg(*flk1*:EGFP;*fab10*:RFP; *DHX15*^+/-^) were sorted in two groups depending on two criteria: a) curly larvae with abnormal development or b) normal developed larvae. Larvae were flat-mounted and analyzed by confocal imaging (Zeiss AxioObserver Z1) to evaluate putative phenotypical defects in the trunk angiogenesis caused by the gene knockout. Genomic DNA extracted from the whole embryos (using Extract-N-Amp™ Tissue PCR Kit, Sigma) was used for the genotyping after vasculature imaging.

### Mouse femoral artery ligation model and magnetic resonance imaging (MRI)

Mice were anesthetized with a mixture of 4% isoflurane and 100% oxygen. The femoral artery was isolated, and 5-0 suture was tied tightly around artery at a ~3 mm distance to the inguinal ligament. Mice were allowed 4 weeks to recover following the surgical procedure.

MRI experiments were conducted on a 7T BioSpec 70/30 horizontal animal scanner (Bruker BioSpin, Ettlingen, Germany), equipped with a 12 cm inner diameter actively shielded gradient system (400 mT/m) using a surface coil dedicated to abdominal imaging. Animals were first anesthetized (1.5% isoflurane in a mixture of 30% O_2_ and 70% CO_2_) and the tail vein was cannulated for administration of contrast agent. Then, animals were transferred under the same anesthesia regime to a Plexiglas holder in supine position with a nose cone for administering anesthetic gases and fixed by a tooth bar, ear bars and adhesive tape. 3D-localizer scans were used to ensure accurate position of the animal’s midline at the level of the posterior limbs in the isocenter of the magnet. T2-weighted images were acquired by a RARE (rapid acquisition with relaxation enhancement) sequence with an effective echo time (TE) of 24 ms, repetition time (TR) 1201 ms and RARE factor 8. Matrix size was 256×256 with an in-plane voxel size of 0.156×0.156 mm^2^, 15 slices, slice thickness 1mm, resulting in a field of view (FOV) of 40×40×15 mm^3^. Time of flight 3D angiography was acquired a FLASH (Fast Low Angle Shot) protocol with TE=2.4 ms, TR=14000 ms, flip angle 20°, 3 averages, matrix size: 448×256×128, voxel size 0.078×0.078×0.234 mm^3^, resulting in a FOV of 35×20×30 mm^3^. The shortening of the T1-relaxation time by the contrast agent enhanced the tissue signal. T1 map was acquired using RARE-VTR (rapid acquisition with relaxation enhancement and variable repetition time) sequence, with TE=7 ms, 6 TR=200, 400, 800, 1500, 3000, 5500 ms, matrix size: 96×96×3, voxel size 0.417×0.417×1 mm^3^, resulting in a FOV of 40×40×3 mm^3^.

To estimate the T1-relaxation rate and to measure the contrast agent relative concentration DCE (dynamic contrast enhanced)-MR imaging was used with a T1-weighted gradient-echo sequence. FLASH protocol was used with TE=1.5 ms, TR=12500 ms, flip angle 15°, 600 repetitions and identical matrix, resolution and FOV than T1 map. 0.025 mM/kg of gadoteridol was administered after the first 100 repetitions were acquired as baseline. Altogether, the MRI session lasted for 50 minutes approximately. After that, mice were returned to their home cage under close supervision until they were recovered from the anesthesia.

T1 maps were calculated in Paravision 6.0 software (Bruker BioSpin, Ettlingen, Germany). Later these maps were processed with custom-made algorithms programmed in Matlab (The MathWorks, Inc, Natick, MA, USA). A binary mask was manually drawn over the T1 map in order to segment the muscle in both legs. The first 40 volumes of the DCE acquistion were removed to assure the signal stabilization. Also, the last 50 volumes were discarded to avoid second pass effects. The slices of the temporal acquisition were spatially smoothed with a Gaussian filter (standard deviation = 0.5) and temporal smoothed with a moving average of 25 neighbors. The baseline of the signal was considered using the 20 volumes after the 10^th^. The signal intensities of the temporal acquisitions were converted to gadolinium concentrations using the method described in (Barboriak et al., 2008; Li et al., 2000; Ortuño et al., 2013). The T1 map acquired before the DCE was used as reference values for the magnetization and the Gadoteridol relaxivity was considered to be 3.35 s^-1^mM^-1^ (Shen et al., 2014). Finally, the obtained concentration curves were also smoothed with a moving average of 9 neighbors. From the concentration curves, different parameters were estimated, such as the time to peak (TTP), the bolus arrival time (BAT) (using the method described in (Cheong et al., 2003), relative time to peak (rTTP) (considering the bolus arrival time as starting point), the wash in and wash out slopes, and the area under the gadolinium concentration curve (AUC).

### Mouse-induced tumor model

LLC1 (Mouse Lewis lung cancer cells) (ATCC, Manassas, VA, USA) were cultured in Dulbecco’s Modified Eagle Medium (DMEM) supplemented with 10% fetal calf serum, 50 U/ml penicillin and 50 μg/ml streptomycin in humified atmosphere at 37°C and 5% CO_2_. Syngeneic LLC1 tumor cells (1×10^5^) were subcutaneously injected into the flank of DHX15^+/-^ and wild-type mice. Primary tumor growth was controlled during the first 3 weeks. Primary tumors were surgically removed 21 days after seeding. Tumor volume was calculated by following formula: V = 4/3 x π x [length x depth x width]. Primary tumors were fixed in 4% PFA and cryoconserved in tissue-tek O.C.T. compound (Sakura, Flemingweg, Netherlands). The Post-surgical metastasis model was performed as follows: Two weeks after primary tumor removal, LLC1 injected mice showed distant metastasis formed in the lungs. Tile scan images of haematoxylin-eosin (H&E) stained paraffin lung sections were visualized using a microscope system (Nikon Eclipse E600, Kanagawa, Japan) and the percentage of pulmonary metastatic area as percent of total lung area was measured with Image J software (ImageJ version 1.52b; National Institutes of Health, Bethesda, MD, USA).

### DHX15 silencing in liver endothelial cells

The silencing of *DHX15* was carried out in mouse primary hepatic endothelial cells immortalized with the SV40 virus LEC; abmGood, Richmond, Canada), through shRNA by lentiviral infection (Dharmacon, Lafayette, Colorado, USA). The SMARTvector incorporated the bipartite 3G Tet-On^®^ induction system, an inducible system with minimal basal expression and potent activation after induction with doxycycline. Cells were cultured in Microvascular Endothelial Cell Growth Complete Medium (Pelobiotech, Planegg, Germany) in humified atmosphere at 37°C and 5% CO_2_.

### Proteome and transcriptome analysis of siL-DHX15-LECs

For proteomics, proteins from non-silenced and DHX15-silenced LECs (1 mg/mL) were extracted in 100 mM NH4HCO3, 8 M urea, 2.5 mM sodium pyrophosphate, 1 mM sodium orthovanadate and 1 mM β-glycerol phosphate buffer. Samples were reduced with dithiothreitol (37 °C, 60 min) and alkylated in the dark with iodoacetamide (25 °C, 30 min). The resulting protein extract was first diluted to 2M urea with 200 mM ammonium bicarbonate for digestion with endoproteinase LysC (1:10 w:w, 37°C, o/n, Wako, cat # 129-02541), and then diluted 2-fold with 200 mM ammonium bicarbonate for trypsin digestion (1:10 w:w, 37°C, 8h, Promega cat # V5113). After digestion, peptide mix was acidified with formic acid and desalted with a MicroSpin C18 column (The Nest Group, Inc) prior to LC-MS/MS analysis. Samples were analyzed using a LTQ-Orbitrap Fusion Lumos mass spectrometer (Thermo Fisher Scientific, San Jose, CA, USA) coupled to an EASY-nLC 1000 (Thermo Fisher Scientific (Proxeon), Odense, Denmark). Peptides were loaded directly onto the analytical column and were separated reversed-phase chromatography with a 90-min gradient (0-35% ACN) in a 50-cm column with an inner diameter of 75 μm, packed with 2 μm C18 particles spectrometer (Thermo Scientific, San Jose, CA, USA). Acquired spectra were analyzed with ProteomeDiscoverer software (v2.0, Swiss-Prot mouse database as in November 2016, 16831 entries). Protein abundances were estimated with the average of the area corresponding to the three most intense peptides. Protein abundance estimates were then log-transformed, normalized by the median, and a fold change, and an adjusted *p-value* was calculated with Perseus 1.5.6.0 (Supplementary Material). The raw proteomics data have been deposited to the PRIDE repository with the dataset identifier PXD018104.

The transcriptome analysis was carried out using RNAseq (HiSeq, Illumina). Total RNA from *Mus musculus* was quantified by Qubit^®^ RNA BR Assay kit (Thermo Fisher Scientific) and the RNA integrity was estimated by using RNA 6000 Nano Bioanalyzer 2100 Assay (Agilent). The RNASeq libraries were prepared with KAPA Stranded mRNA-Seq Illumina^®^ Platforms Kit (Roche-Kapa Biosystems) following the manufacturer’s recommendations. Briefly, 500ng of total RNA was used as the input material, the poly-A fraction was enriched with oligo-dT magnetic beads and the mRNA was fragmented. The strand specificity was achieved during the second strand synthesis performed in the presence of dUTP instead of dTTP. The blunt-ended double stranded cDNA was 3’adenylated and Illumina indexed adapters (Illumina) were ligated. The ligation product was enriched with 15 PCR cycles and the final library was validated on an Agilent 2100 Bioanalyzer with the DNA 7500 assay. The libraries were sequenced on HiSeq 4000 (Illumina, Inc) in paired-end mode with a read length of 2×76bp using HiSeq 4000 SBS kit in a fraction of a HiSeq 4000 PE Cluster kit sequencing flow cell lane, following the manufacturer’s protocol. Image analysis, base calling and quality scoring of the run were processed using the manufacturer’s software Real Time Analysis (RTA 2.7.6) and followed by generation of FASTQ sequence files by CASAVA. RNA-seq reads were mapped against the mouse reference genome (GRCm38) using STAR version 2.5.2a (Dobin et al., 2013) with ENCODE parameters for long RNA. Genes were quantified with RSEM version 1.2.28 (Li and Dewey, 2011) using the gencode M12 version. Differential expression analysis was performed with DESeq2 version 1.18 (Love et al., 2014) with default parameters. Differential alternative splicing was performed with rMATS (Shen et al., 2014). Significant splicing events with FDR<5%, absolute inclusion difference >5% and >70 number of reads were considered significant.

Signaling pathways altered by DHX15 deficiency were modeled using Ingenuity Pathway Software (Ingenuity^®^Systems, Inc., Redwood City, USA). The resulting *p-values* obtained by the Ingenuity Pathways Knowledge Base were adjusted for multiple comparisons using Benjamini and Hochberg’s method.

### Separation of respiratory complexes and supercomplexes by Clear-Native Page (CN-PAGE) and in-gel activity of Complex I

Solubilization of mitochondrial membranes by detergents, CN-PAGE, and staining was performed as described by Jha et al. with minor modifications (Jha et al., 2017). For this, after the mitochondria isolation from wild-type and siL-DHX15-LEC, 150μg of mitochondrial protein were suspended in a low-salt buffer (50 mM NaCl, 50 mM imidazole, pH 7.0) and solubilized with digitonin (8 g/g protein, for solubilization of respiratory chain supercomplexes). Immediately after the electrophoretic run (4–13% gradient polyacrylamide gels), enzymatic colorimetric reaction was performed. Complex I activity was determined by incubating the gel with 2 mM Tris-HCl pH=7.4, 0.1 mg/mL NADH, and 2.5 mg/mL nitro blue tetrazolium (NTB) at room temperature. The original colour of the complex I was preserved by fixing the gels in 50% methanol and 10% acetic acid. After gel scanning, the intensity of each band was quantified by Image J software (ImageJ version 1.52b; National Institutes of Health, Bethesda, MD, USA).

### Statistical analysis

In the case of homoscedasticity and normally distributed data (assessed by Shapiro-Wilk test), groups were compared using a two-sided Student t test or analysis of variance for independent samples. For other types of data, Mann-Whitney U test, or Kruskal-Wallis test was used. Tukey’s test (with analysis of variance) or Dunn’s test (with Kruskal-Wallis) was used as a post hoc test to perform pairwise comparisons. The statistical analysis of contingency tables for proportions was performed using the Fisher’s exact test. Differences were considered to be significant at a *p-value* less than 0.05. The data are presented as the mean±standard error of the mean.

## ACKNOWLEDGEMENTS

This study was supported by grants from the Ministerio de Economia y Competitividad (SAF2016-75358-R and PDI2019-105502RB-100 to MM-R), co-financed by FEDER. CIBERehd, CIBERonc and CIBERbbn are financed by the Instituto de Salud Carlos III. AE-C is funded by ISCIII of the MINECO (reference PT17/0009/0019) and co-financed by FEDER. JR-V was a recipient of a BIOTRACK Postdoctoral Fellowship supported by the European Community’s Seventh Framework Programme and the MINECO (Contract 229673). The CRG/UPF Proteomics Unit is part of the Spanish Infrastructure for Omics Technologies (ICTS OmicsTech) and it is a member of the ProteoRed PRB3 consortium which is supported by grant PT17/0019 of the PE I+D+i 2013-2016 from the Instituto de Salud Carlos III (ISCIII) and ERDF. We acknowledge support from the Spanish Ministry of Science, Innovation and Universities, “Centro de Excelencia Severo Ochoa 2013-2017”, SEV-2012-0208, and “Secretaria d’Universitats i Recerca del Departament d’Economia i Coneixement de la Generalitat de Catalunya” (2017SGR595). The authors would like to thank Drs. Simone Calzolari, Rafael Miñana and Javier Terriente from ZeClinics for their helpful collaboration with the zebrafish model generation and experimentation.

