## Supplemental material and figures for "Loss of the RNA helicase *Dhx15* impairs endothelial energy metabolism, lymphatic drainage and tumor metastasis in mice"

### EXPANDED MATERIAL AND METHODS

***Mouse trachea whole-mount immunofluorescence staining:*** Trachea tissue was fixed overnight in a solution containing 80% methanol and 20% DMSO at -20ºC, and permeabilized in Tris-buffered saline (TBS) containing 2% BSA and 0.1% Triton for 5 h. Tissue was incubated with rat anti-mouse CD31 antibody (BD Pharmigen, Franklin Lakes, New Jersey, USA) at a 1:100 dilution overnight, washed 4 times with TBS and blocked again with TBS containing 2% BSA and 0.1% Triton for 3 h. Finally, the tissue was incubated with Alexa Fluor 555-conjugated rat anti-mouse antibody (Thermo Fisher, Waltham, MA, USA) in a 1:500 dilution overnight and washed 8 times with TBS. The immunofluorescence signal was visualized using an immunofluorescence microscope system (Nikon Eclipse E600, Kanagawa, Japan).

***Mouse fluorescence lymphangiography:*** To evaluate the lymphatic drainage in peripheral areas, 0.05 mL of FITC-Dextran 2000 KDa (Sigma-Aldrich, Darmstadt, Germany) was subcutaneously injected in ear, tail and footpad of wild-type and transgenic mice, with a 30-gauge needle. Immediately after injection, fluorescence images of subcutaneous lymphatic drainage were visualized on a fluorescence stereomicroscope system (Leica Microsystems, Heerbrugg, Switzerland).

***Western Blot experiments.*** Cell lysates were prepared in a lysis buffer (Tris–HCl 20 mM pH=7.4 containing 1% Triton X-100, 0.1% SDS, 50 mM NaCl, 2.5 mM EDTA, 1 mM $Na_4P_2O_7 \cdot 10H_2O$, 20 mM NaF, 1 mM $Na_3VO_4$, 2 mM Pefabloc and Complete® from Roche). Proteins were separated on a 10% SDSpolyacrylamide gel (Mini Protean III; Biorad, Hercules, CA, USA) and transferred for 2 hours at 4ºC to nitrocellulose membranes (Transblot Transfer Medium; Biorad, Hercules, CA, USA) that were stained with Ponceau-S red as a control for protein loading. Then, the membranes were incubated at 4ºC with the following antibodies: mouse monoclonal anti-mouse DHX15 (Santa Cruz Biotechnology, Dallas, Texas, USA), rabbit polyclonal anti-mouse CD31, rabbit anti-mouse β-actin HRP Conjugate (Cell Signaling, Danvers, MA, USA) or mouse monoclonal anti-mouse eNOS (BD Bioscience, San Jose, CA, USA) overnight in a 1:1000 dilution. Next, the membranes were incubated with goat anti-rabbit peroxidase-conjugated secondary antibody or goat anti-mouse peroxidase-conjugated secondary antibody at a 1:2000 dilution (Cell Signaling, Danvers, MA, USA) for 1 hour at room temperature. The bands were visualized by chemiluminescence (Clarity Western ECLS substrate; Biorad, Hercules, CA, USA). The intensity of each band was quantified by Image J software (ImageJ version 1.52b; National Institutes of Health, Bethesda, MD, USA). Band intensities were measured and normalized to the indicated sample (shown as 1.00) on the same membrane.

***Immunocytofluorescence and uptake of oxiLDL.*** For *immunocytofluorescence*, LECs were fixed with 4% paraformaldehyde and blocked with 5% normal goat serum. Next, cells were incubated with rabbit anti-mouse podoplanin (Sigma Chemical, St Louis, Missouri, USA) for 1 hour in a 1:200 dilution. The respective secondary antibody was Alexa Fluor 488-conjugated goat anti-mouse (Molecular Probes, Invitrogen, San Diego, California, USA). In addition, 4',6-diamidino-2-phenylindole (DAPI, Vectashield, Vector laboratories, Burlingame, Ca) was used to counterstain cell nuclei.

Immunofluorescence was visualized with an immunofluorescence microscope system (Nikon Eclipse E600, Kawasaki, Kanagawa, Japan).

For the *uptake of oxiLDL*, cells were incubated with Alexa Fluor 488-conjugated AcLDL (Thermo Fisher, Waltham, MA, USA) at a concentration of 10 µg/mL for 3 hours. Then, cells were washed 3 times with PBS and fixed with 4% paraformaldehyde. Fluorescence was visualized with an immunofluorescence microscope system (Nikon Eclipse E600, Kawasaki, Kanagawa, Japan).

***Real-Time PCR.*** Total RNA was extracted from cultured LECs using the Trizol reagent (Life Technologies, Carlsbad, CA, USA). One microgram of total RNA was reverse transcribed using First Strand cDNA Synthesis Kit (Roche, Mannheim, Germany). Subsequently, complementary DNA samples were amplified for 30-35 cycles (94 for 30 seconds, 55-60°C for 30 seconds, and 72ºC during 1 minute; LigthCycler 480-Roche Diagnostics). To normalize the results, HPRT gene was used as reference. Specific primers for amplification of the complementary DNA were:

Podoplanin, 5'-TGAATCTACTGGCAAGGCACCTCT-3' (forward primer) and 5'-TGCTGAGGTGGACAGTTCCTCTAA-3' (reverse primer);

VEGFR3, 5’-AACAAAGTGGGCCAGGATGAGAGA-3' (forward primer) and 5'-AGCGCAGATGTTCGTACGTGTAGT-3' (reverse primer);

HPRT, 5’-AGTCCCAGCGTCGTGATTAG-3' (forward primer) and 5'-TGATGGCCTCCCATCTCCTT-3' (reverse primer).

***Cell proliferation assay.*** The bromodeoxyuridine (BrdU) cell proliferation assay kit (BrdU Flow kit; BD Pharmigen, Franklin Lakes, New Jersey, USA) was used to measure the incorporation of BrdU during DNA synthesis following the manufacturer’s protocols. Briefly, when cells reached about 60-70% of confluence, BrdU (10 μM) was added to the culture medium for 1h. Then, the BrdU-labeled cells were fixed and the DNA was denatured in fixative solution for 1h at 37°C. Next, the cells were incubated with Alexa Fluor 555-conjugated anti-BrdU antibody for 1h at room temperature. Immunofluorescence was detected by flow cytometry (LSRFortessa; BD Bioscience, San Jose, CA, USA).

***Wound healing assay.*** The wound healing assay was performed in wild-type and siL-DHX15-LEC. Cells were cultured in 6-well plates and incubated at 37°C. When cells were confluent, the straight scratch wound was performed with a 100 μL sterile pipette tip. The scratched areas were photographed using an inverted phase contrast microscope (Olympus IX51, Tokyo, Japan) at 0, 7 and 24 hours, respectively.

***Levels of glycolytic activity and intracellular ATP production.*** The glycolytic activity was determined in non-silenced and DHX15-silenced endothelial cells by an assay coupled to a colorimetric enzymatic reaction (Cayman Chem, Ann Arbor, Michigan, USA) according to the manufacturer's instructions. The cellular production of ATP was determined with a luminescence assay (Cayman Chem, Ann Arbor, MI, USA) according to the manufacturer's instructions.

***Immunofluorescence*****.** Tissues were fixed in 4% paraformaldehyde, cryoprotected overnight in a 30% sucrose solution and embedded in optimal cutting temperature medium. Next, 2-μm frozen sections were rehydrated, blocked with 5% normal goat serum and incubated with rat anti-endomucin (Abcam, Cambridge, UK) as a primary antibody overnight at 4ºC. The binding sites of the primary antibodies were revealed with Alexa Fluor 488-conjugated goat anti-rat (Thermo Fisher, Waltham, MA, USA). Tissues for which immunostaining was performed without primary antibodies were used as negative controls. Slides were mounted using Vectashield (Vector Laboratories, Burlingame, CA, USA) and samples were visualized with a fluorescence microscope (Nikon Eclipse E600, Kanagawa, Japan).

### SUPPLEMENTAL FIGURE LEGENDS

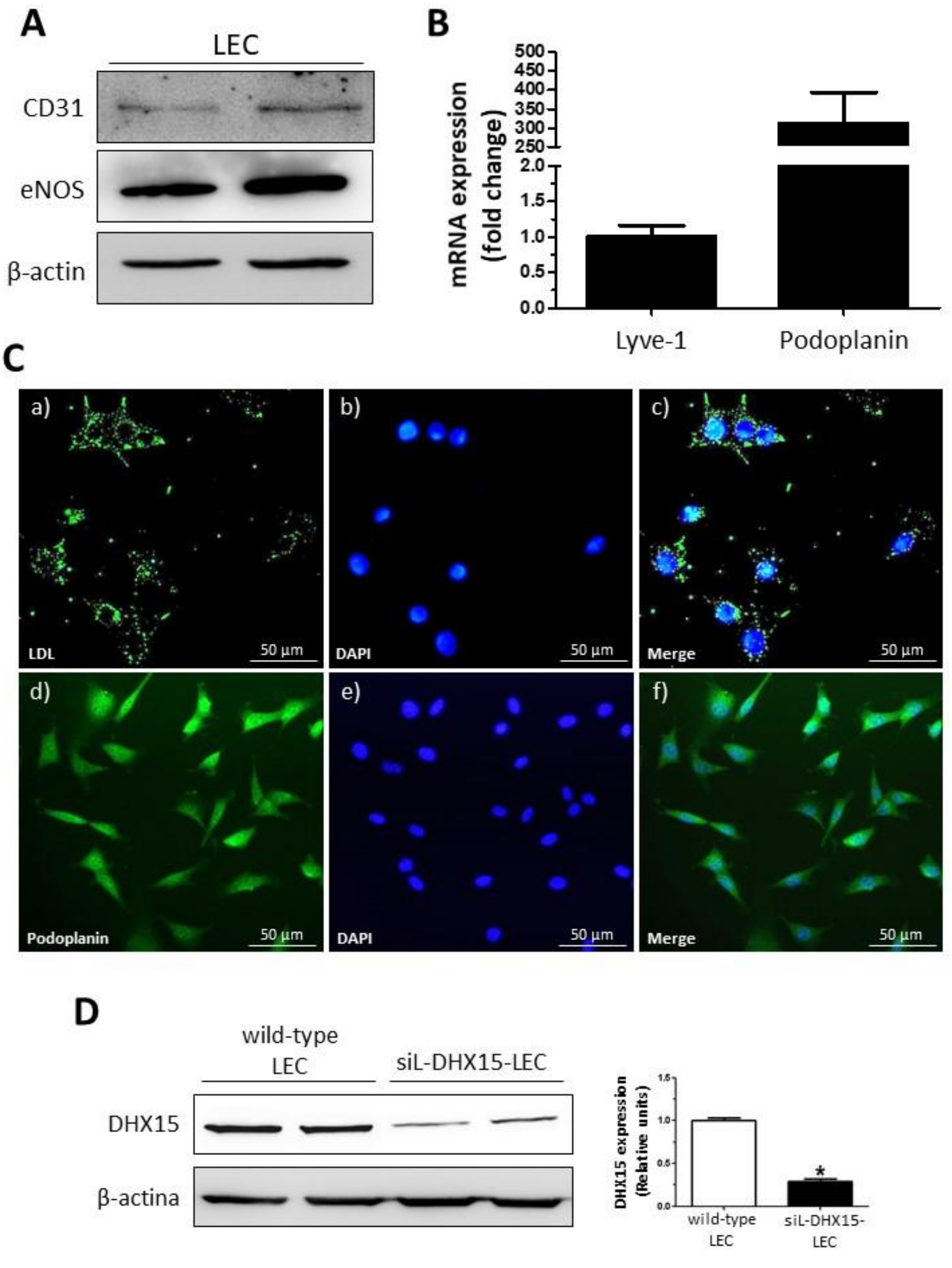


**Supplemental Figure 1. LEC display cardiovascular and lymphatic endothelial markers.** (A) The expression of CD31 and eNOS protein was evaluated by western blot using cell lysates from liver endothelial cells (LEC). β-actin was used as a loading control. (B) LEC were lysed in trizol and their mRNA expression was analyzed by real-time PCR, as described in materials and methods. Graph show the expression levels for the genes Lyve-1 and podoplanin. mRNA levels are illustrated as fold change relative to HPRT mRNA levels (n=5). (C) Representative images of oxidized low-density lipoprotein (oxLDL) uptake (upper panels a and c; green) and immunostaining of the lymphatic endothelial marker podoplanin (lower panels d and f; green) are shown for LEC. Cell nuclei were stained with DAPI (blue). Epifluorescence microscope, original magnification: 200X (n=3). (D) The expression of DHX15 protein was evaluated by western blot using cell lysates from wild-type and silenced DHX15 liver endothelial cells (siL-DHX15-LEC). β-actin was used as a loading control. The densitometric analysis of the protein expression is shown on the right bar graph; $^*p<0.01$ *vs*. wild-type LEC (n=10).

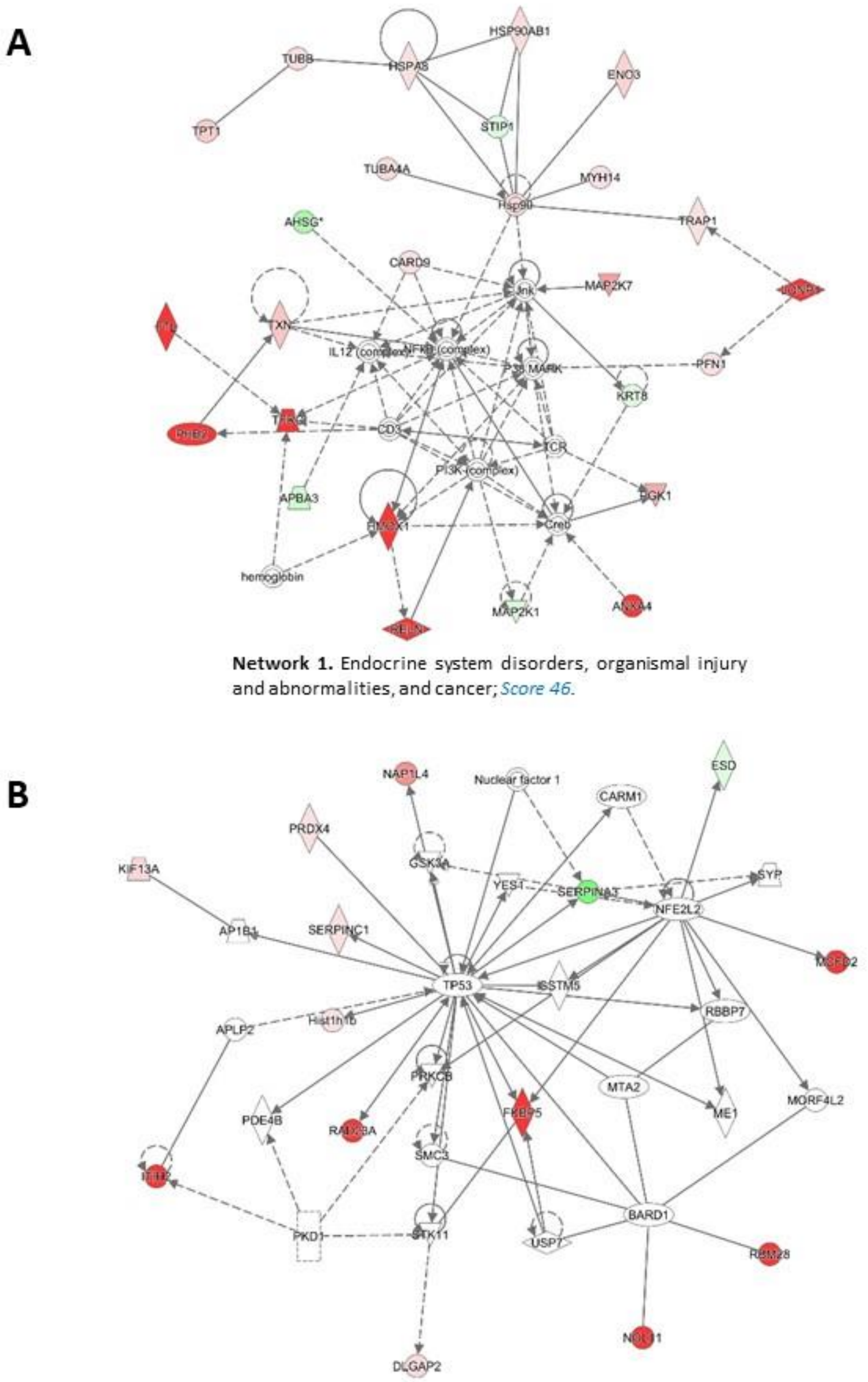


**Network 1.** Endocrine system disorders, organismal injury and abnormalities, and cancer; *Score 46.*

**Network 2.** Gastrointestinal disease, organismal injury and abnormalities, and carbohydrate metabolism; *Score 21.*

**Supplemental Figure 2. Network analysis of molecular changes derived from DHX15 deficiency in LE**C. Networks of pathways obtained from the proteogenomic analysis of wild-type and silenced DHX15 liver endothelial cells (siL-DHX15-LEC) were algorithmically generated based on their connectivity.

The proteins are represented as nodes, and the biological relationship between two nodes is represented as an edge (line). A colored node indicates a protein that was detected by the proteogenomic screening (red: overexpressed and green: reduced expression in silenced-LEC). Nodes are displayed using various shapes that represent the functional class of the proteins. Edges with dashed lines show indirect interaction.

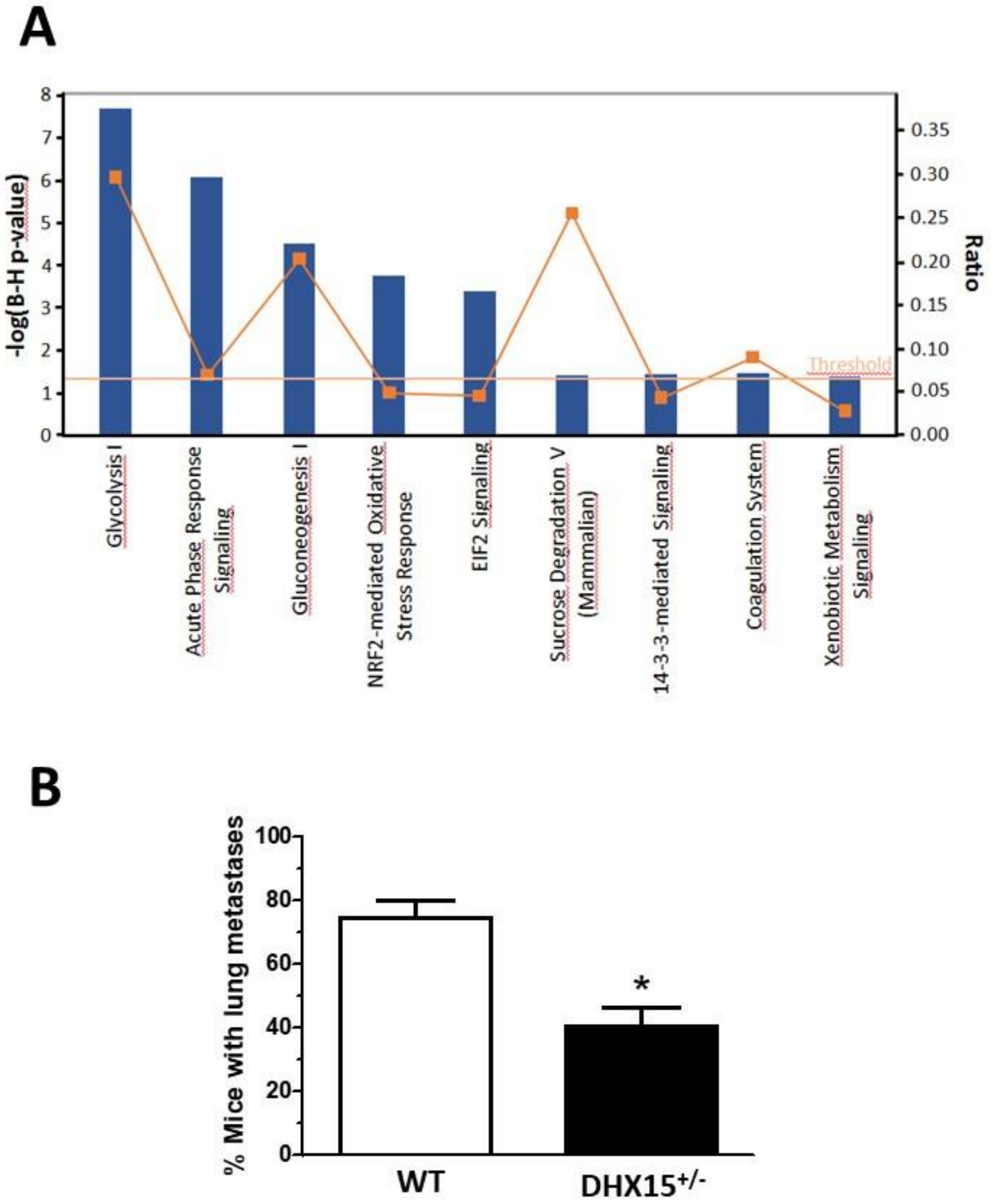


**Supplemental Figure 3.** (A) Pathway analysis of molecular changes derived from DHX15 deficiency in liver endothelial cells (LEC). Signaling pathways affected by the loss DHX15 in LECs were modeled using using the Ingenuity Pathways Knowledge Base (IPA). The significant proteogenomics results obtained from wild-type or silenced DHX15 liver endothelial cells (siL-DHX15-LECs) were compared with global molecular networks using the Fisher's Exact Test. The resulting P-values were adjusted for multiple comparisons using the Benjamini and Hochberg’s method to control the false discovery rate. After multitest adjustment, differences were considered to be significant at a P value less than 0.05 (left axis). The red line across the graph indicates the point where the significance value equals 0.05. In addition, the significance of the association was measured considering the ratio of “the number of targets from the data set that map to the pathway” divided by “the total number of targets that are included in the canonical pathway” (yellow line and right axis). (B) Quantification of lung metastases occurrence in wild-type (WT) and DHX15$^{+/-}$ mice. The graph shows the percentage of mice with distant metastases formed in the lungs after primary tumor removal induced by subcutaneous injection of mouse Lewis lung cancer cells (LLC1); *$p$<0.01 vs. WT mice (n=15).
