## Supplementary Table 1 for "Loss of the RNA helicase *Dhx15* impairs endothelial energy metabolism, lymphatic drainage and tumor metastasis in mice"

| Accession number and ID | Base mean | log2-FoldChange | *p-value* | adjusted *p-value* |
|---|---|---|---|---|
| ENSMUST00000031314.9,Alb-001,protein_coding,ENSMUSG00000029368.10,protein_coding | 280.0184115 | -4.681715526 | 3.96E-173 | 8.51E-169 |
| ENSMUST00000109262.7,Zmynd8-010,protein_coding,ENSMUSG00000039671.18,protein_coding | 317.9776733 | -4.65931549 | 6.48E-165 | 9.30E-161 |
| ENSMUST00000043058.4,Serpina3k-001,protein_coding,ENSMUSG00000058207.6,protein_coding | 75.0993299 | -3.084451115 | 2.84E-64 | 1.22E-60 |
| ENSMUST00000079697.9,Mup7-001,protein_coding,ENSMUSG00000073842.10,protein_coding | 73.55693354 | -2.754991418 | 7.72E-50 | 2.22E-46 |
| ENSMUST00000074018.3,Mup20-001,protein_coding,ENSMUSG00000078672.2,protein_coding | 39.21030999 | -2.295920828 | 8.82E-35 | 9.04E-32 |
| ENSMUST00000032779.11,Ctsc-001,protein_coding,ENSMUSG00000030560.17,protein_coding | 5619.993225 | 0.690874383 | 8.41E-80 | 6.03E-76 |
| ENSMUST00000106112.1,Bnip3-001,protein_coding,ENSMUSG00000078566.8,protein_coding | 10000.77233 | 0.428358432 | 3.36E-78 | 2.07E-74 |
| ENSMUST00000066509.9,Etl4-006,protein_coding,ENSMUSG00000036617.16,protein_coding | 108.4159399 | -2.26303611 | 5.74E-34 | 5.37E-31 |
| ENSMUST00000173631.7,Txn2-008,protein_coding,ENSMUSG00000005354.15,protein_coding | 1174.706303 | 1.278974105 | 1.85E-65 | 8.85E-62 |
| ENSMUST00000085054.4,Serpina1e-002,protein_coding,ENSMUSG00000072849.10,protein_coding | 38.00800991 | -2.227535371 | 7.54E-33 | 6.01E-30 |
| ENSMUST00000161567.7,Pam-002,protein_coding,ENSMUSG00000026335.16,protein_coding | 3451.869374 | -0.837684864 | 1.27E-58 | 4.98E-55 |
| ENSMUST00000177163.7,Rpl41-002,protein_coding,ENSMUSG00000093674.7,protein_coding | 22848.63483 | 0.914577052 | 1.29E-56 | 4.62E-53 |
| ENSMUST00000115124.7,Tnk2-001,protein_coding,ENSMUSG00000022791.16,protein_coding | 2938.721772 | 0.912177814 | 1.50E-55 | 4.95E-52 |
| ENSMUST00000029630.14,Fga-001,protein_coding,ENSMUSG00000028001.16,protein_coding | 30.36285958 | -2.085278112 | 5.46E-29 | 3.09E-26 |
| ENSMUST00000099309.4,Bhmt-001,protein_coding,ENSMUSG00000074768.5,protein_coding | 28.92150001 | -2.022850579 | 2.06E-27 | 1.01E-24 |
| ENSMUST00000187117.1,Gm29216-001,unprocessed_pseudogene,ENSMUSG00000101249.1,unprocessed_pseudogene | 29634.47389 | 0.67185837 | 1.89E-49 | 5.08E-46 |
| ENSMUST00000163694.3,Pkm-004,protein_coding,ENSMUSG00000032294.17,protein_coding | 23570.32749 | -0.847148418 | 1.80E-48 | 4.55E-45 |
| ENSMUST00000150571.1,Ddx27-002,protein_coding,ENSMUSG00000017999.16,protein_coding | 810.0581375 | 0.905021411 | 1.29E-45 | 3.10E-42 |
| ENSMUST00000023583.5,Ahsg-201,protein_coding,ENSMUSG00000022868.5,protein_coding | 41.99520433 | -1.981897863 | 1.58E-26 | 7.57E-24 |
| ENSMUST00000113487.7,Sf1-001,protein_coding,ENSMUSG00000024949.17,protein_coding | 27527.11891 | 0.458778252 | 3.15E-45 | 6.78E-42 |
| ENSMUST00000032228.8,Mug1-001,protein_coding,ENSMUSG00000059908.9,protein_coding | 27.11142362 | -1.947586225 | 1.35E-25 | 5.80E-23 |
| ENSMUST00000100164.4,Hoxc4-002,protein_coding,ENSMUSG00000075394.4,protein_coding | 1672.337209 | -0.595515353 | 9.34E-45 | 1.83E-41 |
| ENSMUST00000134661.7,Knstrn-001,protein_coding,ENSMUSG00000027331.15,protein_coding | 4883.451928 | -0.54856005 | 2.28E-44 | 4.26E-41 |
| ENSMUST00000122129.7,Map7d1-006,protein_coding,ENSMUSG00000028849.17,protein_coding | 1963.019538 | -0.964814653 | 1.04E-42 | 1.86E-39 |
| ENSMUST00000115246.8,Csnk1a1-003,protein_coding,ENSMUSG00000024576.14,protein_coding | 866.1090638 | 1.178583418 | 1.75E-42 | 3.01E-39 |
| ENSMUST00000074051.5,Serpina1c-001,protein_coding,ENSMUSG00000079015.2,protein_coding | 27.88913039 | -1.928278246 | 3.67E-25 | 1.45E-22 |
| ENSMUST00000005862.7,Tfap4-201,protein_coding,ENSMUSG00000005718.7,protein_coding | 4276.614831 | 0.514961121 | 8.82E-41 | 1.37E-37 |
| ENSMUST00000166526.7,U2af1-202,protein_coding,ENSMUSG00000061613.12,protein_coding | 5674.858046 | 0.694525777 | 8.94E-41 | 1.37E-37 |
| ENSMUST00000079158.12,Abcc5-001,protein_coding,ENSMUSG00000022822.15,protein_coding | 9266.193466 | 0.434968021 | 2.33E-39 | 3.45E-36 |
| ENSMUST00000001547.7,Col1a1-001,protein_coding,ENSMUSG00000001506.10,protein_coding | 37472.12452 | -0.255788156 | 3.17E-39 | 4.54E-36 |
| ENSMUST00000041401.10,Herc3-001,protein_coding,ENSMUSG00000029804.16,protein_coding | 1280.469954 | -0.779625478 | 1.24E-38 | 1.72E-35 |
| ENSMUST00000114476.7,Ncam1-005,protein_coding,ENSMUSG00000039542.16,protein_coding | 916.010467 | 1.188009178 | 3.49E-38 | 4.70E-35 |
| ENSMUST00000165860.7,Ubr4-201,protein_coding,ENSMUSG00000066036.14,protein_coding | 16070.89281 | 0.400207995 | 6.39E-38 | 8.33E-35 |
| ENSMUST00000075451.11,Cr1l-001,protein_coding,ENSMUSG00000016481.15,protein_coding | 2576.910526 | 0.744855325 | 2.06E-37 | 2.60E-34 |
| ENSMUST00000048246.4,Fgb-001,protein_coding,ENSMUSG00000033831.5,protein_coding | 27.88170721 | -1.873407018 | 6.24E-24 | 2.18E-21 |
| ENSMUST00000129847.7,Rps19-004,nonsense_mediated_decay,ENSMUSG00000040952.16,protein_coding | 6384.882503 | 0.668809888 | 4.73E-37 | 5.66E-34 |
| ENSMUST00000211875.1,RP24-346O11.2-001,protein_coding,ENSMUSG00000110439.1,protein_coding | 29.90187957 | -1.841067987 | 2.91E-23 | 9.20E-21 |
| ENSMUST00000185899.1,Slc35a4-005,protein_coding,ENSMUSG00000033272.12,protein_coding | 100.7632842 | -1.799697742 | 1.21E-22 | 3.60E-20 |
| ENSMUST00000094499.10,Kif1c-002,protein_coding,ENSMUSG00000020821.17,protein_coding | 32626.4903 | -0.33417557 | 3.07E-36 | 3.39E-33 |
| ENSMUST00000062606.7,Upk3b-001,protein_coding,ENSMUSG00000042985.7,protein_coding | 5897.341854 | -0.582533625 | 1.55E-35 | 1.66E-32 |
| ENSMUST00000095450.10,Serpina1b-001,protein_coding,ENSMUSG00000071178.11,protein_coding | 25.06623486 | -1.785018739 | 5.34E-22 | 1.52E-19 |
| ENSMUST00000027144.7,Cps1-001,protein_coding,ENSMUSG00000025991.8,protein_coding | 37.15580097 | -1.69470213 | 1.10E-19 | 2.45E-17 |
| ENSMUST00000097788.10,Mthfr-003,protein_coding,ENSMUSG00000029009.17,protein_coding | 620.9207629 | 1.203495122 | 9.50E-35 | 9.51E-32 |
| ENSMUST00000031281.13,Antxr2-001,protein_coding,ENSMUSG00000029338.13,protein_coding | 8013.278957 | 0.417315704 | 2.64E-34 | 2.58E-31 |
| ENSMUST00000034740.14,Nedd4-001,protein_coding,ENSMUSG00000032216.14,protein_coding | 110006.986 | -0.395080785 | 3.01E-34 | 2.88E-31 |
| ENSMUST00000049209.12,Gc-001,protein_coding,ENSMUSG00000035540.12,protein_coding | 21.39226668 | -1.629421089 | 7.06E-19 | 1.40E-16 |
| ENSMUST00000121334.7,Sept8-004,protein_coding,ENSMUSG00000018398.18,protein_coding | 720.0319193 | 1.256969984 | 6.67E-34 | 6.11E-31 |
| ENSMUST00000029553.15,Ubap2l-001,protein_coding,ENSMUSG00000042520.16,protein_coding | 3859.688904 | 0.669915317 | 7.77E-34 | 6.97E-31 |
| ENSMUST00000117243.7,Dctn6-004,protein_coding,ENSMUSG00000031516.11,protein_coding | 676.4403138 | 0.900423586 | 2.32E-33 | 2.04E-30 |
| ENSMUST00000050372.9,Crb2-001,protein_coding,ENSMUSG00000035403.10,protein_coding | 1165.262877 | -0.659711966 | 2.59E-33 | 2.23E-30 |
| ENSMUST00000028522.9,Itga6-001,protein_coding,ENSMUSG00000027111.15,protein_coding | 14327.55848 | 0.483284047 | 3.15E-33 | 2.66E-30 |
| ENSMUST00000020381.3,Frs2-201,protein_coding,ENSMUSG00000020170.3,protein_coding | 6601.163991 | -0.486469218 | 3.74E-33 | 3.10E-30 |
| ENSMUST00000031447.11,Anxa3-001,protein_coding,ENSMUSG00000029484.12,protein_coding | 27649.35347 | -0.340913689 | 3.84E-33 | 3.12E-30 |
| ENSMUST00000112132.7,Pzp-001,protein_coding,ENSMUSG00000030359.14,protein_coding | 27.90752696 | -1.595648877 | 3.67E-18 | 6.50E-16 |
| ENSMUST00000088169.5,Rtn3-203,protein_coding,ENSMUSG00000024758.14,protein_coding | 3506.337776 | 0.719051207 | 2.19E-32 | 1.72E-29 |
| ENSMUST00000103172.3,Dstn-001,protein_coding,ENSMUSG00000015932.8,protein_coding | 31234.40563 | -0.529385636 | 3.06E-32 | 2.35E-29 |
| ENSMUST00000020702.10,Igfbp3-001,protein_coding,ENSMUSG00000020427.11,protein_coding | 8073.86065 | 0.738032704 | 6.22E-32 | 4.70E-29 |
| ENSMUST00000031061.11,Dhx15-001,protein_coding,ENSMUSG00000029169.11,protein_coding | 14321.98347 | -1.593451737 | 5.90E-144 | 6.35E-140 |
| ENSMUST00000099856.4,Mtr-201,protein_coding,ENSMUSG00000021311.8,protein_coding | 4932.338354 | 0.502665882 | 1.21E-31 | 8.84E-29 |
| ENSMUST00000065086.4,Gas1-001,protein_coding,ENSMUSG00000052957.7,protein_coding | 2748.393226 | -0.757896783 | 1.23E-31 | 8.84E-29 |
| ENSMUST00000120276.7,Brd4-005,protein_coding,ENSMUSG00000024002.16,protein_coding | 2414.227047 | 0.495878569 | 1.49E-31 | 1.05E-28 |
| ENSMUST00000119499.7,Osbpl5-001,protein_coding,ENSMUSG00000037606.18,protein_coding | 2514.747123 | -0.635667555 | 2.02E-31 | 1.40E-28 |
| ENSMUST00000043765.13,Hnrnpul1-001,protein_coding,ENSMUSG00000040725.13,protein_coding | 10389.88413 | 0.33653699 | 3.28E-31 | 2.24E-28 |
| ENSMUST00000143791.7,Bicc1-001,protein_coding,ENSMUSG00000014329.14,protein_coding | 3719.753037 | -0.384519286 | 3.82E-31 | 2.57E-28 |
| ENSMUST00000215660.1,Pkm-006,protein_coding,ENSMUSG00000032294.17,protein_coding | 306.0107839 | 1.102058256 | 4.41E-31 | 2.92E-28 |
| ENSMUST00000118599.7,Hmga1-005,protein_coding,ENSMUSG00000046711.15,protein_coding | 3069.134965 | 0.422046008 | 5.25E-31 | 3.43E-28 |
| ENSMUST00000143043.7,Wt1-001,protein_coding,ENSMUSG00000016458.14,protein_coding | 3008.92913 | -0.475897158 | 6.40E-31 | 4.11E-28 |
| ENSMUST00000149090.1,Spata5l1-001,processed_transcript,ENSMUSG00000074876.5,processed_transcript | 1724.357367 | 0.653039252 | 7.96E-31 | 5.04E-28 |
| ENSMUST00000025541.5,Gnaq-001,protein_coding,ENSMUSG00000024639.5,protein_coding | 5349.182676 | -0.516286878 | 1.39E-30 | 8.69E-28 |
| ENSMUST00000018625.9,Appbp2-001,protein_coding,ENSMUSG00000018481.9,protein_coding | 10702.4734 | -0.448262206 | 1.56E-30 | 9.60E-28 |
| ENSMUST00000041297.14,Zbed4-001,protein_coding,ENSMUSG00000034333.15,protein_coding | 3114.707783 | -0.554743309 | 1.68E-30 | 1.02E-27 |
| ENSMUST00000138502.1,Rps20-001,protein_coding,ENSMUSG00000028234.6,protein_coding | 33953.96029 | 0.293103647 | 3.74E-30 | 2.23E-27 |
| ENSMUST00000093271.7,Adgrg1-001,protein_coding,ENSMUSG00000031785.16,protein_coding | 8742.00882 | 0.404049551 | 5.41E-30 | 3.19E-27 |
| ENSMUST00000150290.7,Lgals3-002,protein_coding,ENSMUSG00000050335.16,protein_coding | 31312.33473 | 0.550224112 | 1.82E-29 | 1.06E-26 |
| ENSMUST00000004057.7,Fam162a-201,protein_coding,ENSMUSG00000003955.7,protein_coding | 9954.213271 | 0.277647719 | 1.90E-29 | 1.09E-26 |
| ENSMUST00000107472.7,Mup3-002,protein_coding,ENSMUSG00000066154.11,protein_coding | 22.60249006 | -1.590630956 | 3.32E-18 | 5.95E-16 |
| ENSMUST00000114229.2,Cxadr-002,protein_coding,ENSMUSG00000022865.13,protein_coding | 1314.364464 | 1.079138415 | 5.63E-29 | 3.15E-26 |
| ENSMUST00000037141.8,Setd7-001,protein_coding,ENSMUSG00000037111.9,protein_coding | 13210.78301 | -0.446789115 | 6.62E-29 | 3.65E-26 |
| ENSMUST00000201143.1,Sdad1-002,protein_coding,ENSMUSG00000029415.4,protein_coding | 4227.725123 | 0.561718241 | 1.68E-28 | 9.13E-26 |
| ENSMUST00000085056.7,Serpina1a-002,protein_coding,ENSMUSG00000066366.14,protein_coding | 21.69542682 | -1.577230159 | 6.11E-18 | 1.05E-15 |
| ENSMUST00000031325.5,Areg-001,protein_coding,ENSMUSG00000029378.5,protein_coding | 1959.400793 | 0.601054793 | 2.47E-28 | 1.31E-25 |
| ENSMUST00000057816.14,Uckl1-001,protein_coding,ENSMUSG00000089917.7,protein_coding | 2261.33783 | 0.713055578 | 2.80E-28 | 1.47E-25 |
| ENSMUST00000212654.1,Dync1li2-002,nonsense_mediated_decay,ENSMUSG00000035770.8,protein_coding | 1543.996697 | 0.903577241 | 3.34E-28 | 1.73E-25 |
| ENSMUST00000111632.3,Ndrg2-202,protein_coding,ENSMUSG00000004558.14,protein_coding | 1780.06564 | -0.524854013 | 3.52E-28 | 1.81E-25 |
| ENSMUST00000037811.12,Apob-001,protein_coding,ENSMUSG00000020609.14,protein_coding | 328.9065519 | -1.365486913 | 9.00E-28 | 4.56E-25 |
| ENSMUST00000006579.4,Pfdn2-002,protein_coding,ENSMUSG00000006412.10,protein_coding | 623.5746553 | 1.3063709 | 1.06E-27 | 5.32E-25 |
| ENSMUST00000174924.2,Rn7s1-201,misc_RNA,ENSMUSG00000099021.1,misc_RNA | 506.9716949 | -1.571852082 | 2.13E-36 | 2.42E-33 |
| ENSMUST00000175032.2,Rn7s2-201,misc_RNA,ENSMUSG00000099250.1,misc_RNA | 506.9716949 | -1.571852082 | 2.13E-36 | 2.42E-33 |
| ENSMUST00000081654.12,Col18a1-001,protein_coding,ENSMUSG00000001435.14,protein_coding | 5361.004016 | 0.526297246 | 3.38E-27 | 1.64E-24 |
| ENSMUST00000075206.11,Mup14-001,protein_coding,ENSMUSG00000073830.11,protein_coding | 19.83895191 | -1.436707806 | 1.60E-15 | 1.98E-13 |
| ENSMUST00000118566.7,Tpm3-004,protein_coding,ENSMUSG00000027940.18,protein_coding | 5864.370374 | 0.660780934 | 1.62E-26 | 7.64E-24 |

| | | | | |
|---|---|---|---|---|
| ENSMUST00000132620.7,Mprip-006,protein_coding,ENSMUSG00000005417.17,protein_coding | 17767.09835 | 0.454099668 | 1.95E-26 | 9.14E-24 |
| ENSMUST00000038757.7,Csdc2-001,protein_coding,ENSMUSG00000042109.7,protein_coding | 1082.918654 | -0.583411103 | 2.31E-26 | 1.07E-23 |
| ENSMUST00000140713.2,Raly-004,protein_coding,ENSMUSG00000027593.15,protein_coding | 5739.550981 | 0.527900472 | 9.41E-26 | 4.26E-23 |
| ENSMUST00000105348.7,Mbd3-003,protein_coding,ENSMUSG00000035478.14,protein_coding | 2153.352884 | 0.868260555 | 9.32E-26 | 4.26E-23 |
| ENSMUST00000060500.8,Eif5a2-001,protein_coding,ENSMUSG00000050192.8,protein_coding | 1348.253056 | -1.42500014 | 2.69E-37 | 3.31E-34 |
| ENSMUST00000012161.3,Scarf2-201,protein_coding,ENSMUSG00000012017.3,protein_coding | 9915.057836 | -0.381735816 | 9.63E-26 | 4.28E-23 |
| ENSMUST00000055738.11,Tsc22d3-001,protein_coding,ENSMUSG00000031431.13,protein_coding | 3907.515055 | 0.828778061 | 1.17E-25 | 5.13E-23 |
| ENSMUST00000077088.10,Rad18-001,protein_coding,ENSMUSG00000030254.16,protein_coding | 4465.710183 | 0.477520036 | 1.30E-25 | 5.64E-23 |
| ENSMUST00000200186.4,Dhx15-004,nonsense_mediated_decay,ENSMUSG00000029169.11,protein_coding | 3233.119048 | -1.404533537 | 6.48E-25 | 2.49E-22 |
| ENSMUST00000154339.7,Ide-003,processed_transcript,ENSMUSG00000056999.14,protein_coding | 540.394558 | 1.027472785 | 1.39E-25 | 5.94E-23 |
| ENSMUST00000051129.9,Lrrc75b-001,protein_coding,ENSMUSG00000046807.9,protein_coding | 2345.580012 | -0.525699352 | 1.52E-25 | 6.41E-23 |
| ENSMUST00000107911.7,Nr1h2-002,protein_coding,ENSMUSG00000060601.13,protein_coding | 653.4644045 | -1.374531883 | 2.09E-25 | 8.74E-23 |
| ENSMUST00000021018.10,Taf15-001,protein_coding,ENSMUSG00000020680.15,protein_coding | 10694.93375 | 0.613396271 | 2.58E-25 | 1.07E-22 |
| ENSMUST00000111663.8,Fgfr1op2-002,protein_coding,ENSMUSG00000040242.14,protein_coding | 3653.307613 | -0.628626658 | 2.75E-25 | 1.13E-22 |
| ENSMUST00000069600.12,Ndrg3-002,protein_coding,ENSMUSG00000027634.14,protein_coding | 611.6457165 | 0.948484302 | 2.86E-25 | 1.16E-22 |
| ENSMUST00000100335.9,Mrc2-001,protein_coding,ENSMUSG00000020695.14,protein_coding | 2223.071685 | -0.487317596 | 3.02E-25 | 1.22E-22 |
| ENSMUST00000026500.11,Avil-001,protein_coding,ENSMUSG00000025432.11,protein_coding | 3755.001006 | 1.238306512 | 3.13E-25 | 1.25E-22 |
| ENSMUST00000100544.10,Agfg2-002,protein_coding,ENSMUSG00000029722.15,protein_coding | 780.8084561 | 1.435795211 | 1.50E-20 | 3.69E-18 |
| ENSMUST00000039571.13,2410004B18Rik-001,protein_coding,ENSMUSG00000036873.13,protein_coding | 1335.96471 | -0.603867741 | 4.55E-25 | 1.78E-22 |
| ENSMUST00000112516.7,Ltbp1-002,protein_coding,ENSMUSG00000001870.15,protein_coding | 18053.73524 | 0.433850708 | 6.03E-25 | 2.34E-22 |
| ENSMUST00000110999.7,Map2k7-004,protein_coding,ENSMUSG00000002948.19,protein_coding | 853.7925311 | 1.445950659 | 4.74E-45 | 9.72E-42 |
| ENSMUST00000029451.11,Tspan2-001,protein_coding,ENSMUSG00000027858.13,protein_coding | 487.8037367 | -0.938492991 | 1.01E-24 | 3.83E-22 |
| ENSMUST00000013931.11,Ehmt2-001,protein_coding,ENSMUSG00000013787.15,protein_coding | 9263.346964 | 0.363816069 | 1.31E-24 | 4.96E-22 |
| ENSMUST00000091504.4,Nmt2-002,protein_coding,ENSMUSG00000026643.16,protein_coding | 3271.500216 | 0.450074094 | 2.90E-24 | 1.08E-21 |
| ENSMUST00000056153.7,Fads6-001,protein_coding,ENSMUSG00000044788.10,protein_coding | 1383.199895 | -0.510447571 | 3.28E-24 | 1.22E-21 |
| ENSMUST00000213770.1,Thyn1-001,protein_coding,ENSMUSG00000035443.10,protein_coding | 2549.250181 | 0.488267125 | 4.04E-24 | 1.49E-21 |
| ENSMUST00000153499.1,Sars-003,protein_coding,ENSMUSG00000068739.13,protein_coding | 4585.106689 | 0.505669772 | 4.09E-24 | 1.49E-21 |
| ENSMUST00000143278.7,Fam131b-001,protein_coding,ENSMUSG00000029861.17,protein_coding | 3009.244464 | -0.560211596 | 4.21E-24 | 1.52E-21 |
| ENSMUST00000009789.14,P4ha1-001,protein_coding,ENSMUSG00000019916.14,protein_coding | 3260.873488 | 0.413504431 | 4.50E-24 | 1.61E-21 |
| ENSMUST00000056924.13,Plagl2-001,protein_coding,ENSMUSG00000051413.17,protein_coding | 5056.476251 | 0.28998883 | 4.79E-24 | 1.71E-21 |
| ENSMUST00000073043.4,Cxcl12-003,protein_coding,ENSMUSG00000061353.11,protein_coding | 2426.496933 | -1.03637569 | 5.94E-24 | 2.10E-21 |
| ENSMUST00000046476.12,Dusp12-003,nonsense_mediated_decay,ENSMUSG00000026659.13,protein_coding | 182.0334535 | 1.471250914 | 1.79E-23 | 5.76E-21 |
| ENSMUST00000053819.5,Srsf4-001,protein_coding,ENSMUSG00000028911.16,protein_coding | 12436.45402 | 0.282258269 | 6.90E-24 | 2.39E-21 |
| ENSMUST00000042012.6,Sf3b3-001,protein_coding,ENSMUSG00000033732.10,protein_coding | 60158.85043 | 0.413275075 | 6.98E-24 | 2.40E-21 |
| ENSMUST00000100500.8,Med24-010,protein_coding,ENSMUSG00000017210.14,protein_coding | 1006.166918 | 0.950842167 | 7.81E-24 | 2.67E-21 |
| ENSMUST00000159345.7,Naaa-002,protein_coding,ENSMUSG00000029413.14,protein_coding | 1650.662436 | 0.579654777 | 8.85E-24 | 3.00E-21 |
| ENSMUST00000207628.1,Zfp787-002,protein_coding,ENSMUSG00000046792.8,protein_coding | 314.476666 | 1.063605827 | 9.60E-24 | 3.23E-21 |
| ENSMUST00000172306.2,Dusp7-001,protein_coding,ENSMUSG00000053716.9,protein_coding | 3361.442684 | -0.395506614 | 1.08E-23 | 3.62E-21 |
| ENSMUST00000126787.1,Itga6-005,processed_transcript,ENSMUSG00000027111.15,protein_coding | 6054.113058 | 0.441742052 | 1.31E-23 | 4.33E-21 |
| ENSMUST00000126243.8,Mbd6-007,nonsense_mediated_decay,ENSMUSG00000025409.14,protein_coding | 445.3596516 | 0.799173811 | 1.50E-23 | 4.91E-21 |
| ENSMUST00000029371.2,Smad9-001,protein_coding,ENSMUSG00000027796.2,protein_coding | 4979.448763 | -0.3198399 | 1.58E-23 | 5.14E-21 |
| ENSMUST00000107756.3,Coro2a-001,protein_coding,ENSMUSG00000028337.14,protein_coding | 2468.856347 | 0.651457807 | 1.76E-23 | 5.70E-21 |
| ENSMUST00000004673.13,Ndrg2-201,protein_coding,ENSMUSG00000004558.14,protein_coding | 668.3464841 | 1.539054571 | 2.26E-28 | 1.22E-25 |
| ENSMUST00000113488.7,Sf1-003,protein_coding,ENSMUSG00000024949.17,protein_coding | 5420.142529 | 0.390967041 | 1.90E-23 | 6.07E-21 |
| ENSMUST00000155415.7,Ppfibp1-009,protein_coding,ENSMUSG00000016487.15,protein_coding | 338.4536415 | 1.572995917 | 1.73E-45 | 3.91E-42 |
| ENSMUST00000057767.4,Upk1b-201,protein_coding,ENSMUSG00000049436.4,protein_coding | 2258.046001 | -0.573641982 | 4.07E-23 | 1.27E-20 |
| ENSMUST00000113337.8,Ubr2-202,protein_coding,ENSMUSG00000023977.14,protein_coding | 9721.445482 | 0.528745314 | 4.08E-23 | 1.27E-20 |
| ENSMUST00000156793.1,Pabpc1-006,processed_transcript,ENSMUSG00000022283.14,protein_coding | 7882.429961 | 0.533551988 | 6.18E-23 | 1.91E-20 |
| ENSMUST00000020085.6,Ube2d1-001,protein_coding,ENSMUSG00000019927.6,protein_coding | 2659.691167 | -0.476864649 | 6.92E-23 | 2.13E-20 |
| ENSMUST00000081593.12,Pgk1-001,protein_coding,ENSMUSG00000062070.12,protein_coding | 216236.2783 | 0.471066313 | 7.49E-23 | 2.29E-20 |
| ENSMUST00000028005.2,Mgst3-001,protein_coding,ENSMUSG00000026688.5,protein_coding | 4855.297553 | 0.633095979 | 7.90E-23 | 2.38E-20 |
| ENSMUST00000079413.9,Fkbp5-001,protein_coding,ENSMUSG00000024222.16,protein_coding | 31493.57049 | 0.374852517 | 7.86E-23 | 2.38E-20 |
| ENSMUST00000216139.1,Akap12-002,processed_transcript,ENSMUSG00000038587.9,protein_coding | 17822.61441 | -0.687736329 | 1.08E-22 | 3.22E-20 |
| ENSMUST00000087566.10,Aldoa-003,protein_coding,ENSMUSG00000030695.14,protein_coding | 12034.42891 | 1.622014889 | 0 | 0 |
| ENSMUST00000046257.13,Sptan1-003,protein_coding,ENSMUSG00000057738.13,protein_coding | 46387.50858 | -0.288096384 | 1.44E-22 | 4.20E-20 |
| ENSMUST00000112996.8,Tsc22d3-006,protein_coding,ENSMUSG00000031431.13,protein_coding | 528.7148307 | 0.799729309 | 1.43E-22 | 4.20E-20 |
| ENSMUST00000033880.6,Eif4ebp1-001,protein_coding,ENSMUSG00000031490.6,protein_coding | 18942.45656 | 0.501714444 | 1.60E-22 | 4.64E-20 |
| ENSMUST00000106746.7,Helz-002,protein_coding,ENSMUSG00000020721.16,protein_coding | 3921.498307 | -0.547858973 | 2.63E-22 | 7.61E-20 |
| ENSMUST00000152804.8,Cystm1-005,protein_coding,ENSMUSG00000046727.12,protein_coding | 246.9245233 | 1.051954064 | 3.80E-22 | 1.09E-19 |
| ENSMUST00000072727.6,Nap1l4-001,protein_coding,ENSMUSG00000059119.9,protein_coding | 2371.156328 | 1.63385349 | 5.81E-66 | 3.13E-62 |
| ENSMUST00000021866.8,Riok1-201,protein_coding,ENSMUSG00000021428.8,protein_coding | 4836.069413 | 0.341899895 | 6.42E-22 | 1.82E-19 |
| ENSMUST00000166397.2,Suds3-002,protein_coding,ENSMUSG00000066900.11,protein_coding | 5703.559693 | 0.476347655 | 8.12E-22 | 2.29E-19 |
| ENSMUST00000016315.15,Lamb3-001,protein_coding,ENSMUSG00000026639.18,protein_coding | 1870.558874 | 0.451234924 | 9.25E-22 | 2.58E-19 |
| ENSMUST00000037547.8,Disp2-001,protein_coding,ENSMUSG00000040035.14,protein_coding | 1613.453095 | 0.593482452 | 1.12E-21 | 3.11E-19 |
| ENSMUST00000173166.1,Uggt1-007,nonsense_mediated_decay,ENSMUSG00000037470.14,protein_coding | 746.2067811 | 0.850736503 | 1.17E-21 | 3.23E-19 |
| ENSMUST00000047611.2,Nthl1-201,protein_coding,ENSMUSG00000041429.2,protein_coding | 1262.701751 | 0.485441584 | 1.20E-21 | 3.29E-19 |
| ENSMUST00000116456.9,Cyth3-001,protein_coding,ENSMUSG00000018001.18,protein_coding | 4462.735177 | -0.519704329 | 1.96E-21 | 5.33E-19 |
| ENSMUST00000092167.5,Ptprb-201,protein_coding,ENSMUSG00000020154.9,protein_coding | 1567.333132 | -0.48847923 | 2.02E-21 | 5.46E-19 |
| ENSMUST00000044355.9,Selenow-001,protein_coding,ENSMUSG00000041571.9,protein_coding | 7875.968639 | 0.579018657 | 2.83E-21 | 7.62E-19 |
| ENSMUST00000060655.14,Nod1-001,protein_coding,ENSMUSG00000038058.14,protein_coding | 1763.043181 | -0.516779372 | 3.16E-21 | 8.44E-19 |
| ENSMUST00000041403.11,Becn1-002,nonsense_mediated_decay,ENSMUSG00000035086.13,protein_coding | 5172.793816 | -0.348850163 | 4.06E-21 | 1.08E-18 |
| ENSMUST00000126610.1,Rpl12-001,protein_coding,ENSMUSG00000038900.17,protein_coding | 119116.8514 | 0.406866458 | 4.56E-21 | 1.21E-18 |
| ENSMUST00000021532.4,Snapc1-201,protein_coding,ENSMUSG00000021113.4,protein_coding | 1461.560185 | -0.678387251 | 4.87E-21 | 1.28E-18 |
| ENSMUST00000048099.4,Nemp1-001,protein_coding,ENSMUSG00000040195.11,protein_coding | 3356.986719 | 0.354284852 | 4.98E-21 | 1.30E-18 |
| ENSMUST00000017610.9,Timp2-001,protein_coding,ENSMUSG00000017466.9,protein_coding | 42939.06297 | -0.334641392 | 6.25E-21 | 1.62E-18 |
| ENSMUST00000022345.6,Dnajc9-201,protein_coding,ENSMUSG00000021811.6,protein_coding | 6395.331512 | -0.452856637 | 6.73E-21 | 1.73E-18 |
| ENSMUST00000049948.5,Asrgl1-001,protein_coding,ENSMUSG00000024654.8,protein_coding | 1886.735479 | 0.484179533 | 7.73E-21 | 1.98E-18 |
| ENSMUST00000005256.12,Ndrg1-001,protein_coding,ENSMUSG00000005125.12,protein_coding | 11855.04079 | 1.181696736 | 8.21E-21 | 2.09E-18 |
| ENSMUST00000022875.6,Ank-001,protein_coding,ENSMUSG00000022265.6,protein_coding | 8425.186474 | 0.370208559 | 8.81E-21 | 2.23E-18 |
| ENSMUST00000029382.12,Ppid-001,protein_coding,ENSMUSG00000027804.13,protein_coding | 12849.05 | -0.670342055 | 1.04E-20 | 2.61E-18 |
| ENSMUST00000110049.7,Eif4enif1-003,protein_coding,ENSMUSG00000020454.17,protein_coding | 8503.583985 | 0.31953327 | 1.16E-20 | 2.91E-18 |
| ENSMUST00000094761.10,Stk40-001,protein_coding,ENSMUSG00000042608.15,protein_coding | 2576.885635 | 0.444334789 | 1.23E-20 | 3.05E-18 |
| ENSMUST00000027726.13,Cyb5r1-001,protein_coding,ENSMUSG00000026456.18,protein_coding | 15571.97403 | 0.806672002 | 1.40E-20 | 3.47E-18 |
| ENSMUST00000002603.11,Scrib-001,protein_coding,ENSMUSG00000022568.16,protein_coding | 1299.974256 | 1.700342759 | 9.65E-26 | 4.28E-23 |
| ENSMUST00000017637.12,Igfbp4-001,protein_coding,ENSMUSG00000017493.12,protein_coding | 38285.84209 | 0.338371588 | 1.60E-20 | 3.90E-18 |
| ENSMUST00000065353.12,Yap1-001,protein_coding,ENSMUSG00000053110.13,protein_coding | 7647.474998 | 0.437961508 | 1.70E-20 | 4.14E-18 |
| ENSMUST00000037370.13,Sorcs2-001,protein_coding,ENSMUSG00000029093.14,protein_coding | 629.0986785 | 0.656528186 | 1.71E-20 | 4.14E-18 |
| ENSMUST00000196466.1,Snhg8-001,lincRNA,ENSMUSG00000104960.1,processed_transcript | 2507.225101 | 0.322378287 | 1.94E-20 | 4.67E-18 |
| ENSMUST00000185827.6,Ndufs1-005,retained_intron,ENSMUSG00000025968.16,protein_coding | 562.5818813 | 0.817351479 | 2.02E-20 | 4.83E-18 |
| ENSMUST00000050149.11,Mical2-001,protein_coding,ENSMUSG00000038244.13,protein_coding | 29093.32503 | -0.24662421 | 2.03E-20 | 4.83E-18 |
| ENSMUST00000069799.2,Irs1-001,protein_coding,ENSMUSG00000055980.2,protein_coding | 4547.540789 | -0.589552196 | 2.51E-20 | 5.94E-18 |
| ENSMUST00000103009.4,Rps20-002,retained_intron,ENSMUSG00000028234.6,protein_coding | 47387.19222 | 0.303475506 | 3.22E-20 | 7.56E-18 |

| | | | | |
|---|---|---|---|---|
| ENSMUST00000007317.7,Krt19-001,protein_coding,ENSMUSG00000020911.14,protein_coding | 34105.44165 | 0.566012589 | 3.63E-20 | 8.48E-18 |
| ENSMUST00000114888.9,Hmga1-002,protein_coding,ENSMUSG00000046711.15,protein_coding | 79856.93345 | 0.349390492 | 5.02E-20 | 1.17E-17 |
| ENSMUST00000020629.4,Gfpt2-001,protein_coding,ENSMUSG00000020363.6,protein_coding | 2298.522553 | 0.461735287 | 5.72E-20 | 1.32E-17 |
| ENSMUST00000097541.9,Fam129a-002,nonsense_mediated_decay,ENSMUSG00000026483.13,protein_coding | 5883.30524 | 0.767496186 | 6.18E-20 | 1.42E-17 |
| ENSMUST00000112866.7,Cxcl12-001,protein_coding,ENSMUSG00000061353.11,protein_coding | 5205.226223 | -0.610307567 | 6.94E-20 | 1.59E-17 |
| ENSMUST00000114496.1,Cd47-002,protein_coding,ENSMUSG00000055447.18,protein_coding | 4153.385653 | 0.466555446 | 7.13E-20 | 1.62E-17 |
| ENSMUST00000018992.3,Rars-001,protein_coding,ENSMUSG00000018848.4,protein_coding | 27744.06076 | 0.383461605 | 8.71E-20 | 1.97E-17 |
| ENSMUST00000024894.1,Cyp1b1-201,protein_coding,ENSMUSG00000024087.3,protein_coding | 7276.869454 | -0.958428621 | 9.51E-20 | 2.14E-17 |
| ENSMUST00000076352.6,Serpinb1a-201,protein_coding,ENSMUSG00000044734.15,protein_coding | 518.4857423 | 0.814758449 | 1.02E-19 | 2.29E-17 |
| ENSMUST00000040202.14,Atxn2l-001,protein_coding,ENSMUSG00000032637.15,protein_coding | 7384.582156 | 0.303657283 | 1.05E-19 | 2.34E-17 |
| ENSMUST00000170633.7,Trappc9-204,protein_coding,ENSMUSG00000047921.15,protein_coding | 112.4010938 | 1.727555954 | 7.42E-32 | 5.51E-29 |
| ENSMUST00000107545.8,Med1-001,protein_coding,ENSMUSG00000018160.15,protein_coding | 11140.26652 | -0.247242052 | 1.13E-19 | 2.50E-17 |
| ENSMUST00000165307.7,Anp32e-001,protein_coding,ENSMUSG00000015749.12,protein_coding | 18445.37209 | -0.487706557 | 1.16E-19 | 2.54E-17 |
| ENSMUST00000106289.8,Cant1-004,protein_coding,ENSMUSG00000025575.14,protein_coding | 224.561432 | 1.01373847 | 1.31E-19 | 2.87E-17 |
| ENSMUST00000103208.1,Tmem201-002,protein_coding,ENSMUSG00000044700.15,protein_coding | 3296.077702 | -0.403315452 | 1.61E-19 | 3.48E-17 |
| ENSMUST00000038867.12,Opa1-003,protein_coding,ENSMUSG00000038084.16,protein_coding | 3300.764446 | 0.29327115 | 1.61E-19 | 3.48E-17 |
| ENSMUST00000110087.8,Fgd3-001,protein_coding,ENSMUSG00000037946.11,protein_coding | 1971.355103 | -0.449891949 | 1.65E-19 | 3.56E-17 |
| ENSMUST00000020123.6,Tmpo-004,protein_coding,ENSMUSG00000019961.17,protein_coding | 10813.43912 | 0.318382929 | 1.68E-19 | 3.59E-17 |
| ENSMUST00000033828.6,Gas6-001,protein_coding,ENSMUSG00000031451.6,protein_coding | 12712.09944 | -0.366481947 | 1.94E-19 | 4.14E-17 |
| ENSMUST00000093769.7,Magi1-004,protein_coding,ENSMUSG00000045095.16,protein_coding | 1892.012519 | 0.417064354 | 2.00E-19 | 4.25E-17 |
| ENSMUST00000040261.5,Macrod1-201,protein_coding,ENSMUSG00000036278.6,protein_coding | 621.9011197 | 0.710690583 | 2.05E-19 | 4.32E-17 |
| ENSMUST00000154483.7,Csnk1d-006,protein_coding,ENSMUSG00000025162.17,protein_coding | 412.4563282 | 1.028798302 | 2.34E-19 | 4.91E-17 |
| ENSMUST00000134778.2,Mars-004,protein_coding,ENSMUSG00000040354.14,protein_coding | 240.561763 | 1.372716486 | 2.42E-19 | 5.05E-17 |
| ENSMUST00000215233.1,Clk3-005,protein_coding,ENSMUSG00000032316.8,protein_coding | 1488.960312 | 0.678336805 | 2.81E-19 | 5.85E-17 |
| ENSMUST00000204890.2,Cnbp-006,protein_coding,ENSMUSG00000030057.15,protein_coding | 12941.90237 | 0.459516268 | 2.99E-19 | 6.19E-17 |
| ENSMUST00000114468.8,Osbpl3-003,protein_coding,ENSMUSG00000029822.15,protein_coding | 1473.293192 | -0.763770826 | 3.29E-19 | 6.77E-17 |
| ENSMUST00000105366.1,Atp5d-003,protein_coding,ENSMUSG00000003072.15,protein_coding | 717.2564125 | 1.019291664 | 3.65E-19 | 7.49E-17 |
| ENSMUST00000046290.2,Lpcat2-001,protein_coding,ENSMUSG00000033192.5,protein_coding | 701.6484527 | 0.816656053 | 3.98E-19 | 8.13E-17 |
| ENSMUST00000170288.1,Slc35a4-003,processed_transcript,ENSMUSG00000033272.12,protein_coding | 338.3156945 | 0.879636561 | 5.12E-19 | 1.04E-16 |
| ENSMUST00000106435.8,Fbf1-201,protein_coding,ENSMUSG00000020776.18,protein_coding | 4756.657963 | 0.619000522 | 5.20E-19 | 1.05E-16 |
| ENSMUST00000112514.1,Ltbp1-007,protein_coding,ENSMUSG00000001870.15,protein_coding | 18349.10883 | -0.348943925 | 5.44E-19 | 1.09E-16 |
| ENSMUST00000022340.3,Nid2-201,protein_coding,ENSMUSG00000021806.3,protein_coding | 1349.122511 | -0.48111942 | 5.51E-19 | 1.10E-16 |
| ENSMUST00000095794.3,Tbpl1-002,protein_coding,ENSMUSG00000071359.13,protein_coding | 2012.729359 | 0.379790936 | 6.56E-19 | 1.31E-16 |
| ENSMUST00000179784.7,Emc1-202,protein_coding,ENSMUSG00000078517.11,protein_coding | 744.6456169 | 1.74096937 | 1.95E-35 | 2.05E-32 |
| ENSMUST00000026415.7,Cdk2-201,protein_coding,ENSMUSG00000025358.15,protein_coding | 2435.419479 | -0.602653974 | 7.07E-19 | 1.40E-16 |
| ENSMUST00000173094.7,Dap3-010,protein_coding,ENSMUSG00000068921.14,protein_coding | 1006.451503 | 0.775929474 | 8.20E-19 | 1.61E-16 |
| ENSMUST00000009885.13,Ubp1-201,protein_coding,ENSMUSG00000009741.15,protein_coding | 2267.51087 | 0.692541815 | 8.42E-19 | 1.65E-16 |
| ENSMUST00000168902.7,Serpinf1-007,nonsense_mediated_decay,ENSMUSG00000000753.15,protein_coding | 2225.931343 | 0.506271708 | 8.55E-19 | 1.67E-16 |
| ENSMUST00000174287.1,Neat1-002,lincRNA,ENSMUSG00000092274.2,lincRNA | 7812.782252 | -0.707028495 | 9.38E-19 | 1.82E-16 |
| ENSMUST00000199369.4,Aldh2-005,protein_coding,ENSMUSG00000029455.14,protein_coding | 7195.128929 | 0.398421307 | 9.46E-19 | 1.83E-16 |
| ENSMUST00000160770.8,Amotl1-005,processed_transcript,ENSMUSG00000013076.16,protein_coding | 1095.330209 | 0.632050422 | 1.01E-18 | 1.93E-16 |
| ENSMUST00000007236.4,Syngr3-001,protein_coding,ENSMUSG00000007021.4,protein_coding | 311.3117876 | -0.841347649 | 1.31E-18 | 2.50E-16 |
| ENSMUST00000064062.12,Immt-005,protein_coding,ENSMUSG00000052337.15,protein_coding | 23080.10151 | 0.38647364 | 1.45E-18 | 2.77E-16 |
| ENSMUST00000072289.11,Flot2-001,protein_coding,ENSMUSG00000061981.14,protein_coding | 8291.140597 | -0.314116701 | 1.47E-18 | 2.79E-16 |
| ENSMUST00000165649.2,Ghitm-202,protein_coding,ENSMUSG00000041028.15,protein_coding | 43329.50846 | 0.530911669 | 1.49E-18 | 2.81E-16 |
| ENSMUST00000186302.6,Agfg1-002,protein_coding,ENSMUSG00000026159.13,protein_coding | 6745.986768 | 0.306700663 | 1.56E-18 | 2.93E-16 |
| ENSMUST00000119486.7,Hmga1-006,protein_coding,ENSMUSG00000046711.15,protein_coding | 10666.48564 | 0.36497582 | 1.88E-18 | 3.52E-16 |
| ENSMUST00000212129.1,Rpl18a-003,nonsense_mediated_decay,ENSMUSG00000045128.9,protein_coding | 58223.54862 | 0.352164902 | 1.91E-18 | 3.56E-16 |
| ENSMUST00000199050.1,Ubap2l-023,protein_coding,ENSMUSG00000042520.16,protein_coding | 253.5421068 | 0.905516739 | 2.07E-18 | 3.84E-16 |
| ENSMUST00000021614.13,Pfkp-005,protein_coding,ENSMUSG00000021196.13,protein_coding | 8567.575399 | 0.464532283 | 2.16E-18 | 3.99E-16 |
| ENSMUST00000079085.10,Rpl34-002,protein_coding,ENSMUSG00000062006.12,protein_coding | 43476.8255 | 0.311268119 | 2.28E-18 | 4.19E-16 |
| ENSMUST00000042646.7,Atoh8-001,protein_coding,ENSMUSG00000037621.8,protein_coding | 13575.00148 | -0.472445021 | 2.55E-18 | 4.68E-16 |
| ENSMUST00000173938.7,Psmd11-004,protein_coding,ENSMUSG00000017428.16,protein_coding | 6438.565447 | 0.329678059 | 2.58E-18 | 4.70E-16 |
| ENSMUST00000133861.7,Mprip-004,protein_coding,ENSMUSG00000005417.17,protein_coding | 14899.57291 | -0.406150899 | 3.00E-18 | 5.44E-16 |
| ENSMUST00000103191.10,Rpl22-001,protein_coding,ENSMUSG00000028936.15,protein_coding | 32949.29059 | 0.339730739 | 3.11E-18 | 5.62E-16 |
| ENSMUST00000111024.9,Enah-009,protein_coding,ENSMUSG00000022995.16,protein_coding | 1979.028565 | -0.473453406 | 3.28E-18 | 5.90E-16 |
| ENSMUST00000209847.1,Pih1d1-006,protein_coding,ENSMUSG00000003423.15,protein_coding | 107.7214589 | 1.771919582 | 1.14E-27 | 5.64E-25 |
| ENSMUST00000113127.6,G3bp2-001,protein_coding,ENSMUSG00000029405.16,protein_coding | 14156.55003 | -0.426010499 | 3.39E-18 | 6.05E-16 |
| ENSMUST00000113697.7,Tpm1-006,protein_coding,ENSMUSG00000032366.15,protein_coding | 13080.58904 | -0.656987477 | 3.50E-18 | 6.23E-16 |
| ENSMUST00000100215.10,Sh3glb2-002,protein_coding,ENSMUSG00000026860.16,protein_coding | 1009.673782 | 1.806048995 | 1.51E-50 | 4.63E-47 |
| ENSMUST00000003964.16,Gys1-001,protein_coding,ENSMUSG00000003865.16,protein_coding | 20161.05642 | 0.547852081 | 4.01E-18 | 7.08E-16 |
| ENSMUST00000015877.13,Capza2-001,protein_coding,ENSMUSG00000015733.13,protein_coding | 18482.59486 | -0.482542489 | 4.30E-18 | 7.56E-16 |
| ENSMUST00000117583.7,Nfu1-001,protein_coding,ENSMUSG00000029993.16,protein_coding | 7668.966202 | 0.503292168 | 4.36E-18 | 7.62E-16 |
| ENSMUST00000109848.9,Wars-001,protein_coding,ENSMUSG00000021266.16,protein_coding | 21626.37044 | 0.592006433 | 4.60E-18 | 8.01E-16 |
| ENSMUST00000054098.2,Slc35g1-201,protein_coding,ENSMUSG00000044026.2,protein_coding | 1175.049252 | -0.491832612 | 4.65E-18 | 8.07E-16 |
| ENSMUST00000077037.11,Pcbp2-201,protein_coding,ENSMUSG00000056851.13,protein_coding | 59375.13357 | 0.3321075 | 5.92E-18 | 1.02E-15 |
| ENSMUST00000186864.6,Reep6-201,protein_coding,ENSMUSG00000035504.16,protein_coding | 1641.296521 | 1.84329656 | 1.88E-41 | 3.11E-38 |
| ENSMUST00000123548.1,Gm13205-001,antisense,ENSMUSG00000086606.1,antisense | 487.9025012 | 0.750070677 | 6.63E-18 | 1.13E-15 |
| ENSMUST00000023829.6,Cdkn1a-001,protein_coding,ENSMUSG00000023067.13,protein_coding | 4343.729309 | 0.438348971 | 6.61E-18 | 1.13E-15 |
| ENSMUST00000120834.7,Pld1-001,protein_coding,ENSMUSG00000027695.16,protein_coding | 1914.666877 | 0.427821133 | 6.78E-18 | 1.15E-15 |
| ENSMUST00000202601.3,Ube2k-004,nonsense_mediated_decay,ENSMUSG00000029203.16,protein_coding | 4985.357859 | 0.254356151 | 7.19E-18 | 1.22E-15 |
| ENSMUST00000100052.10,Igf2bp2-001,protein_coding,ENSMUSG00000033581.16,protein_coding | 10066.90428 | 0.360010442 | 7.53E-18 | 1.27E-15 |
| ENSMUST00000058790.11,Ldlrad3-001,protein_coding,ENSMUSG00000048058.17,protein_coding | 6740.005977 | 0.513570624 | 7.60E-18 | 1.28E-15 |
| ENSMUST00000019998.8,Perp-001,protein_coding,ENSMUSG00000019851.8,protein_coding | 2979.138563 | 0.400305463 | 8.19E-18 | 1.37E-15 |
| ENSMUST00000162003.7,Mff-005,protein_coding,ENSMUSG00000026150.14,protein_coding | 393.7105968 | 1.206896826 | 9.30E-18 | 1.55E-15 |
| ENSMUST00000196739.4,Fubp1-004,protein_coding,ENSMUSG00000028034.15,protein_coding | 2246.084154 | 0.564680087 | 9.39E-18 | 1.56E-15 |
| ENSMUST00000172272.7,Ddx46-202,protein_coding,ENSMUSG00000021500.15,protein_coding | 9739.024747 | -0.67272269 | 9.97E-18 | 1.65E-15 |
| ENSMUST00000019382.16,Tecr-002,protein_coding,ENSMUSG00000031708.16,protein_coding | 19826.57536 | 0.356862489 | 1.13E-17 | 1.86E-15 |
| ENSMUST00000069949.12,Hnrnpa2b1-001,protein_coding,ENSMUSG00000004980.16,protein_coding | 20049.79389 | -0.485444087 | 1.24E-17 | 2.04E-15 |
| ENSMUST00000153290.7,Htra1-002,processed_transcript,ENSMUSG00000006205.13,protein_coding | 1019.962571 | 1.190293611 | 1.26E-17 | 2.07E-15 |
| ENSMUST00000059619.1,Cdc42ep1-201,protein_coding,ENSMUSG00000049521.1,protein_coding | 13826.99181 | -0.300719468 | 1.27E-17 | 2.07E-15 |
| ENSMUST00000044804.7,Cdsn-001,protein_coding,ENSMUSG00000039518.7,protein_coding | 8471.609634 | 1.040892341 | 1.34E-17 | 2.17E-15 |
| ENSMUST00000080953.11,Lrp2-001,protein_coding,ENSMUSG00000027070.14,protein_coding | 3648.608313 | -0.407606799 | 1.42E-17 | 2.29E-15 |
| ENSMUST00000099673.8,Cd44-002,protein_coding,ENSMUSG00000005087.17,protein_coding | 30187.80574 | 0.261893736 | 1.59E-17 | 2.56E-15 |
| ENSMUST00000020220.14,Nuak1-001,protein_coding,ENSMUSG00000020032.14,protein_coding | 1346.565642 | -0.605336474 | 1.61E-17 | 2.59E-15 |
| ENSMUST00000135988.7,Desi1-002,nonsense_mediated_decay,ENSMUSG00000022472.16,protein_coding | 3985.029101 | 0.387995348 | 1.64E-17 | 2.62E-15 |
| ENSMUST00000102811.9,Rpl12-002,nonsense_mediated_decay,ENSMUSG00000038900.17,protein_coding | 86394.18597 | 0.439065648 | 1.68E-17 | 2.68E-15 |
| ENSMUST00000175934.7,Tcof1-005,protein_coding,ENSMUSG00000024613.16,protein_coding | 33994.88783 | 0.386096833 | 1.95E-17 | 3.09E-15 |
| ENSMUST00000132790.1,Cul7-004,retained_intron,ENSMUSG00000038545.13,protein_coding | 627.4793351 | -0.653450853 | 2.07E-17 | 3.28E-15 |
| ENSMUST00000006367.7,Htra1-001,protein_coding,ENSMUSG00000006205.13,protein_coding | 9999.797555 | 0.645074229 | 2.16E-17 | 3.40E-15 |
| ENSMUST00000107974.2,Iglon5-001,protein_coding,ENSMUSG00000013367.5,protein_coding | 382.8094796 | -0.694852973 | 2.51E-17 | 3.92E-15 |
| ENSMUST00000171252.7,Serpinb6a-011,protein_coding,ENSMUSG00000060147.15,protein_coding | 1087.291754 | 0.709222136 | 2.50E-17 | 3.92E-15 |

| | | | | |
|---|---|---|---|---|
| ENSMUST00000118539.1,Cth-001,protein_coding,ENSMUSG00000028179.12,protein_coding | 4961.569752 | 0.882720253 | 2.65E-17 | 4.13E-15 |
| ENSMUST00000067512.7,Smpd3-001,protein_coding,ENSMUSG00000031906.9,protein_coding | 5173.186965 | -0.472930183 | 2.88E-17 | 4.48E-15 |
| ENSMUST00000051594.11,Depdc1b-001,protein_coding,ENSMUSG00000021697.12,protein_coding | 1299.019529 | -0.509711251 | 3.03E-17 | 4.70E-15 |
| ENSMUST00000072031.12,Mprip-201,protein_coding,ENSMUSG00000005417.17,protein_coding | 3490.271435 | -0.454544035 | 3.62E-17 | 5.58E-15 |
| ENSMUST00000004994.15,Pax3-002,protein_coding,ENSMUSG00000004872.15,protein_coding | 2931.109548 | 0.48967516 | 3.70E-17 | 5.68E-15 |
| ENSMUST00000167163.7,Serpinb6a-202,protein_coding,ENSMUSG00000060147.15,protein_coding | 1895.955534 | 0.847026831 | 3.96E-17 | 6.06E-15 |
| ENSMUST00000198811.1,Bri3bp-002,protein_coding,ENSMUSG00000037905.13,protein_coding | 2031.928432 | 0.400140846 | 4.35E-17 | 6.64E-15 |
| ENSMUST00000190390.6,Srcap-013,protein_coding,ENSMUSG00000053877.12,protein_coding | 16424.32805 | -0.222732816 | 4.98E-17 | 7.57E-15 |
| ENSMUST00000033486.5,Plp2-001,protein_coding,ENSMUSG00000031146.6,protein_coding | 24412.59038 | 0.302520624 | 5.02E-17 | 7.60E-15 |
| ENSMUST00000190096.6,Agap1-002,protein_coding,ENSMUSG00000055013.15,protein_coding | 5419.661575 | 0.60525355 | 5.92E-17 | 8.94E-15 |
| ENSMUST00000073488.11,Nr1h2-001,protein_coding,ENSMUSG00000060601.13,protein_coding | 2152.137981 | 0.585105142 | 6.95E-17 | 1.05E-14 |
| ENSMUST00000044524.15,Dap-001,protein_coding,ENSMUSG00000039168.15,protein_coding | 21203.73632 | 0.329820629 | 7.61E-17 | 1.14E-14 |
| ENSMUST00000068947.13,Acox3-001,protein_coding,ENSMUSG00000029098.16,protein_coding | 1608.550037 | -0.419180369 | 8.53E-17 | 1.27E-14 |
| ENSMUST00000196709.4,Fdps-002,protein_coding,ENSMUSG00000059743.12,protein_coding | 2051.066909 | 0.852052762 | 8.56E-17 | 1.28E-14 |
| ENSMUST00000076813.6,Iah1-201,protein_coding,ENSMUSG00000062054.6,protein_coding | 2643.935812 | 0.56724155 | 8.88E-17 | 1.32E-14 |
| ENSMUST00000199420.4,Csde1-005,protein_coding,ENSMUSG00000068823.10,protein_coding | 1521.017217 | -1.116232582 | 9.08E-17 | 1.34E-14 |
| ENSMUST00000071539.9,Slc39a11-001,protein_coding,ENSMUSG00000041654.15,protein_coding | 3095.617593 | 0.345308749 | 1.00E-16 | 1.48E-14 |
| ENSMUST00000165485.7,Efemp2-001,protein_coding,ENSMUSG00000024909.14,protein_coding | 14001.88427 | -0.380689474 | 1.01E-16 | 1.48E-14 |
| ENSMUST00000032071.12,Dusp11-001,protein_coding,ENSMUSG00000030002.15,protein_coding | 22448.20612 | -0.286511994 | 1.04E-16 | 1.53E-14 |
| ENSMUST00000167070.1,Ergic1-007,protein_coding,ENSMUSG00000001576.14,protein_coding | 782.5031577 | 0.56762942 | 1.18E-16 | 1.73E-14 |
| ENSMUST00000034140.8,Itfg1-001,protein_coding,ENSMUSG00000031703.8,protein_coding | 8866.677851 | 0.277568503 | 1.22E-16 | 1.77E-14 |
| ENSMUST00000051955.7,Rps23-201,protein_coding,ENSMUSG00000049517.7,protein_coding | 82521.80113 | 0.268847192 | 1.32E-16 | 1.91E-14 |
| ENSMUST00000089119.11,Rnaset2b-201,protein_coding,ENSMUSG00000094724.7,protein_coding | 7092.316595 | 0.409960341 | 1.32E-16 | 1.91E-14 |
| ENSMUST00000186129.6,Fn1-002,protein_coding,ENSMUSG00000026193.15,protein_coding | 56745.7091 | 0.212898859 | 1.33E-16 | 1.92E-14 |
| ENSMUST00000029629.14,Pla2g12a-001,protein_coding,ENSMUSG00000027999.15,protein_coding | 10152.28348 | 0.5631462 | 1.36E-16 | 1.94E-14 |
| ENSMUST00000211121.1,Snrnp70-008,protein_coding,ENSMUSG00000063511.11,protein_coding | 627.0457073 | 0.59884504 | 1.36E-16 | 1.94E-14 |
| ENSMUST00000125824.7,Set-005,processed_transcript,ENSMUSG00000054766.13,protein_coding | 12366.6941 | 0.46610673 | 1.37E-16 | 1.95E-14 |
| ENSMUST00000190240.6,Fam134a-002,protein_coding,ENSMUSG00000049339.16,protein_coding | 1168.023374 | 0.945245598 | 1.56E-16 | 2.22E-14 |
| ENSMUST00000074127.13,Aldh3a2-002,protein_coding,ENSMUSG00000010025.19,protein_coding | 6713.243468 | 0.412394273 | 1.59E-16 | 2.26E-14 |
| ENSMUST00000121493.8,Palld-004,protein_coding,ENSMUSG00000058056.16,protein_coding | 2643.073638 | -0.658318832 | 1.66E-16 | 2.34E-14 |
| ENSMUST00000001561.11,Nrip2-001,protein_coding,ENSMUSG00000001520.12,protein_coding | 1271.93696 | 1.163310366 | 1.74E-16 | 2.45E-14 |
| ENSMUST00000144265.5,Sri-004,retained_intron,ENSMUSG00000003161.15,protein_coding | 622.1542361 | 0.673869093 | 1.84E-16 | 2.58E-14 |
| ENSMUST00000041133.8,Fam110c-201,protein_coding,ENSMUSG00000036136.8,protein_coding | 1179.386263 | 0.651742166 | 1.87E-16 | 2.61E-14 |
| ENSMUST00000020188.12,Ifngr1-001,protein_coding,ENSMUSG00000020009.12,protein_coding | 3699.241325 | 0.299513572 | 1.88E-16 | 2.62E-14 |
| ENSMUST00000007131.15,Acly-001,protein_coding,ENSMUSG00000020917.17,protein_coding | 10476.0138 | 0.718059248 | 1.94E-16 | 2.69E-14 |
| ENSMUST00000100680.9,Stx2-002,protein_coding,ENSMUSG00000029428.13,protein_coding | 1584.173319 | 0.474369514 | 1.94E-16 | 2.69E-14 |
| ENSMUST00000041364.12,Nop14-001,protein_coding,ENSMUSG00000036693.12,protein_coding | 17521.55645 | 0.433739171 | 2.00E-16 | 2.76E-14 |
| ENSMUST00000210959.1,Eif4ebp1-002,processed_transcript,ENSMUSG00000031490.6,protein_coding | 12749.21261 | 0.478649373 | 2.05E-16 | 2.82E-14 |
| ENSMUST00000041475.14,Myof-201,protein_coding,ENSMUSG00000048612.15,protein_coding | 71767.85509 | 0.22378093 | 2.06E-16 | 2.83E-14 |
| ENSMUST00000000844.14,Tpd52l2-002,protein_coding,ENSMUSG00000000827.18,protein_coding | 2353.149805 | -0.39805088 | 2.26E-16 | 3.08E-14 |
| ENSMUST00000151394.7,Dusp11-002,processed_transcript,ENSMUSG00000030002.15,protein_coding | 287.8013184 | 0.916295294 | 2.36E-16 | 3.22E-14 |
| ENSMUST00000049053.8,Fam168a-002,protein_coding,ENSMUSG00000029461.17,protein_coding | 2472.922232 | -0.60484714 | 2.44E-16 | 3.31E-14 |
| ENSMUST00000034469.6,Egln1-001,protein_coding,ENSMUSG00000031987.6,protein_coding | 11257.42641 | 0.265244705 | 2.45E-16 | 3.31E-14 |
| ENSMUST00000026737.11,Shisa5-001,protein_coding,ENSMUSG00000025647.16,protein_coding | 5623.193745 | 0.351130424 | 2.46E-16 | 3.31E-14 |
| ENSMUST00000025095.7,Cox7a2l-201,protein_coding,ENSMUSG00000024248.13,protein_coding | 13883.59781 | 0.320977331 | 2.50E-16 | 3.36E-14 |
| ENSMUST00000186932.1,Ptma-004,processed_transcript,ENSMUSG00000026238.14,protein_coding | 109.4484971 | 1.131522257 | 2.73E-16 | 3.67E-14 |
| ENSMUST00000080067.12,Slc4a2-001,protein_coding,ENSMUSG00000028962.14,protein_coding | 13201.82688 | -0.329682148 | 2.84E-16 | 3.79E-14 |
| ENSMUST00000047226.8,Lonp1-201,protein_coding,ENSMUSG00000041168.9,protein_coding | 61854.84059 | 0.523794468 | 3.01E-16 | 4.01E-14 |
| ENSMUST00000138045.7,Capzb-011,protein_coding,ENSMUSG00000028745.18,protein_coding | 859.5826009 | 0.68051934 | 3.03E-16 | 4.03E-14 |
| ENSMUST00000173908.7,Cyb5r1-009,protein_coding,ENSMUSG00000026456.18,protein_coding | 939.9560258 | 1.011608172 | 3.15E-16 | 4.17E-14 |
| ENSMUST00000077983.12,Sdccag3-003,protein_coding,ENSMUSG00000026927.17,protein_coding | 4609.380524 | 0.342061783 | 3.58E-16 | 4.72E-14 |
| ENSMUST00000126460.7,Krt19-002,processed_transcript,ENSMUSG00000020911.14,protein_coding | 7278.106737 | 0.563707644 | 4.32E-16 | 5.69E-14 |
| ENSMUST00000053230.6,Ulk3-001,protein_coding,ENSMUSG00000032308.9,protein_coding | 1290.784996 | 0.56345571 | 4.97E-16 | 6.52E-14 |
| ENSMUST00000100302.3,Prkca-004,protein_coding,ENSMUSG00000050965.14,protein_coding | 2006.697874 | 0.417594115 | 5.14E-16 | 6.72E-14 |
| ENSMUST00000107619.2,Klf4-001,protein_coding,ENSMUSG00000003032.8,protein_coding | 9808.405337 | 0.3511227 | 5.40E-16 | 7.04E-14 |
| ENSMUST00000117836.7,Ifnar2-002,protein_coding,ENSMUSG00000022971.18,protein_coding | 2798.968204 | 0.412955121 | 5.51E-16 | 7.16E-14 |
| ENSMUST00000121799.7,Arhgap23-003,protein_coding,ENSMUSG00000049807.16,protein_coding | 7916.604134 | -0.268191097 | 6.16E-16 | 7.99E-14 |
| ENSMUST00000028214.14,Sh3glb2-001,protein_coding,ENSMUSG00000026860.16,protein_coding | 7934.903443 | -0.410247067 | 6.38E-16 | 8.25E-14 |
| ENSMUST00000021271.13,Per1-001,protein_coding,ENSMUSG00000020893.17,protein_coding | 9696.044895 | 0.31694379 | 6.45E-16 | 8.31E-14 |
| ENSMUST00000105465.7,Psap-001,protein_coding,ENSMUSG00000004207.14,protein_coding | 64895.17291 | 0.253634018 | 6.60E-16 | 8.48E-14 |
| ENSMUST00000003442.7,Cacnb3-201,protein_coding,ENSMUSG00000003352.13,protein_coding | 8827.469778 | 0.458231762 | 6.79E-16 | 8.70E-14 |
| ENSMUST00000031766.11,Asns-001,protein_coding,ENSMUSG00000029752.12,protein_coding | 89626.02569 | 0.420261078 | 7.45E-16 | 9.51E-14 |
| ENSMUST00000197984.4,Smarcc1-002,protein_coding,ENSMUSG00000032481.16,protein_coding | 8336.40111 | 0.256629308 | 7.70E-16 | 9.81E-14 |
| ENSMUST00000102691.10,Cdca7-001,protein_coding,ENSMUSG00000055612.15,protein_coding | 16690.65182 | 0.421376891 | 8.33E-16 | 1.06E-13 |
| ENSMUST00000030677.6,Map3k6-001,protein_coding,ENSMUSG00000028862.6,protein_coding | 2839.02461 | 0.43616594 | 9.08E-16 | 1.15E-13 |
| ENSMUST00000179748.7,Nudt5-201,protein_coding,ENSMUSG00000025817.12,protein_coding | 5919.218268 | 0.367365953 | 9.55E-16 | 1.21E-13 |
| ENSMUST00000052602.5,C77080-005,protein_coding,ENSMUSG00000050390.12,protein_coding | 5368.019574 | -0.333790432 | 9.91E-16 | 1.24E-13 |
| ENSMUST00000143829.4,Slain2-002,protein_coding,ENSMUSG00000036087.18,protein_coding | 4261.573883 | -0.638067388 | 9.89E-16 | 1.24E-13 |
| ENSMUST00000102930.9,Ewsr1-009,protein_coding,ENSMUSG00000009079.16,protein_coding | 9458.937374 | 0.458688617 | 1.05E-15 | 1.32E-13 |
| ENSMUST00000106226.8,Tial1-001,protein_coding,ENSMUSG00000030846.15,protein_coding | 682.4264393 | -0.77126715 | 1.11E-15 | 1.39E-13 |
| ENSMUST00000176308.1,Gnb1-007,retained_intron,ENSMUSG00000029064.15,protein_coding | 9955.215389 | 0.34067795 | 1.13E-15 | 1.41E-13 |
| ENSMUST00000163419.7,Scaf4-202,protein_coding,ENSMUSG00000022983.15,protein_coding | 4099.166532 | 0.400146124 | 1.22E-15 | 1.52E-13 |
| ENSMUST00000014065.15,Clip3-001,protein_coding,ENSMUSG00000013921.15,protein_coding | 4121.98075 | -0.362337448 | 1.35E-15 | 1.67E-13 |
| ENSMUST00000179238.7,Psap-204,protein_coding,ENSMUSG00000004207.14,protein_coding | 3472.545093 | 2.885555142 | 4.16E-139 | 3.58E-135 |
| ENSMUST00000099747.3,B3galnt2-201,protein_coding,ENSMUSG00000039242.9,protein_coding | 4487.477369 | 0.351354803 | 1.61E-15 | 1.98E-13 |
| ENSMUST00000024657.11,Phf10-001,protein_coding,ENSMUSG00000023883.13,protein_coding | 26052.07957 | 0.549434503 | 1.69E-15 | 2.07E-13 |
| ENSMUST00000117192.7,Klhl22-002,protein_coding,ENSMUSG00000022750.17,protein_coding | 1996.530335 | -0.45177572 | 1.79E-15 | 2.19E-13 |
| ENSMUST00000003572.9,Gars-001,protein_coding,ENSMUSG00000029777.10,protein_coding | 135186.558 | 0.433484821 | 1.81E-15 | 2.21E-13 |
| ENSMUST00000161779.7,Hipk2-004,protein_coding,ENSMUSG00000061436.15,protein_coding | 11882.14142 | -0.25680602 | 1.87E-15 | 2.28E-13 |
| ENSMUST00000203828.2,Eif4g3-013,protein_coding,ENSMUSG00000028760.16,protein_coding | 6021.468011 | 0.250286457 | 1.98E-15 | 2.40E-13 |
| ENSMUST00000027863.12,Atp1b1-001,protein_coding,ENSMUSG00000026576.12,protein_coding | 1198.674666 | -0.53528877 | 2.23E-15 | 2.70E-13 |
| ENSMUST00000162114.7,4933434E20Rik-005,protein_coding,ENSMUSG00000027942.16,protein_coding | 1118.745622 | 0.488358727 | 2.48E-15 | 2.99E-13 |
| ENSMUST00000159772.7,Kif2a-007,protein_coding,ENSMUSG00000021693.19,protein_coding | 2407.602535 | 0.557254114 | 2.55E-15 | 3.07E-13 |
| ENSMUST00000105687.8,Tmem201-001,protein_coding,ENSMUSG00000044700.15,protein_coding | 3780.397449 | 0.342084183 | 2.58E-15 | 3.10E-13 |
| ENSMUST00000058906.6,Chchd10-201,protein_coding,ENSMUSG00000049422.6,protein_coding | 5417.812124 | 0.657931475 | 2.79E-15 | 3.34E-13 |
| ENSMUST00000032265.12,Olr1-001,protein_coding,ENSMUSG00000030162.14,protein_coding | 1016.594233 | -0.832825779 | 2.85E-15 | 3.40E-13 |
| ENSMUST00000097836.9,Hp1bp3-202,protein_coding,ENSMUSG00000028759.13,protein_coding | 3387.534966 | 0.613088703 | 2.88E-15 | 3.42E-13 |
| ENSMUST00000115456.5,Cav1-008,protein_coding,ENSMUSG00000007655.16,protein_coding | 738.4129353 | 0.848987231 | 2.97E-15 | 3.52E-13 |
| ENSMUST00000046520.12,Fkbp14-001,protein_coding,ENSMUSG00000038074.16,protein_coding | 1459.315358 | -0.462910673 | 3.29E-15 | 3.89E-13 |
| ENSMUST00000017458.10,Mpp2-001,protein_coding,ENSMUSG00000017314.12,protein_coding | 4435.563865 | -0.4908807 | 3.30E-15 | 3.90E-13 |
| ENSMUST00000023432.8,Nit2-201,protein_coding,ENSMUSG00000022751.8,protein_coding | 2456.83529 | 0.463089497 | 3.38E-15 | 3.98E-13 |
| ENSMUST00000131787.1,2410006H16Rik-001,processed_transcript,ENSMUSG00000086841.1,processed_transcript | 10232.29249 | 0.571960146 | 3.48E-15 | 4.08E-13 |

| | | | | |
|---|---|---|---|---|
| ENSMUST00000023226.12,Plec-004,protein_coding,ENSMUSG00000022565.15,protein_coding | 89668.59363 | 0.314090712 | 3.66E-15 | 4.28E-13 |
| ENSMUST00000063199.12,Ralgps2-008,protein_coding,ENSMUSG00000026594.14,protein_coding | 2557.195401 | 0.415913823 | 3.78E-15 | 4.40E-13 |
| ENSMUST00000106787.7,Rhoc-002,protein_coding,ENSMUSG00000002233.13,protein_coding | 5839.827536 | 0.398034351 | 3.82E-15 | 4.44E-13 |
| ENSMUST00000078482.12,Dchs1-001,protein_coding,ENSMUSG00000036862.15,protein_coding | 297.468538 | -0.815553807 | 3.85E-15 | 4.47E-13 |
| ENSMUST00000102625.10,Sars-002,protein_coding,ENSMUSG00000068739.13,protein_coding | 60818.1569 | 0.399233793 | 3.95E-15 | 4.57E-13 |
| ENSMUST00000125184.7,Uba52-001,protein_coding,ENSMUSG00000090137.8,protein_coding | 42579.98644 | 0.285746117 | 4.58E-15 | 5.28E-13 |
| ENSMUST00000037960.10,Zdhhc9-001,protein_coding,ENSMUSG00000036985.12,protein_coding | 3657.372765 | 0.376975025 | 4.74E-15 | 5.45E-13 |
| ENSMUST00000049230.10,Kcnab1-001,protein_coding,ENSMUSG00000027827.16,protein_coding | 2397.419788 | -0.498167257 | 4.82E-15 | 5.53E-13 |
| ENSMUST00000110333.1,Anapc1-005,protein_coding,ENSMUSG00000014355.10,protein_coding | 14172.25992 | 0.283802765 | 5.12E-15 | 5.86E-13 |
| ENSMUST00000020308.3,Ddit4-201,protein_coding,ENSMUSG00000020108.3,protein_coding | 14902.67658 | 0.575821288 | 5.50E-15 | 6.28E-13 |
| ENSMUST00000042163.14,Naa25-001,protein_coding,ENSMUSG00000042719.17,protein_coding | 10410.07649 | -0.364423013 | 5.84E-15 | 6.66E-13 |
| ENSMUST00000166192.6,Copg2-001,protein_coding,ENSMUSG00000025607.15,protein_coding | 7851.879958 | 0.408879566 | 6.26E-15 | 7.11E-13 |
| ENSMUST00000111484.8,Acin1-003,protein_coding,ENSMUSG00000022185.18,protein_coding | 5349.045287 | -0.621152315 | 6.40E-15 | 7.26E-13 |
| ENSMUST00000164470.2,Kars-003,protein_coding,ENSMUSG00000031948.14,protein_coding | 2246.279812 | 0.829140839 | 6.57E-15 | 7.42E-13 |
| ENSMUST00000109252.7,Ncoa3-001,protein_coding,ENSMUSG00000027678.17,protein_coding | 7508.03711 | 0.392469485 | 6.64E-15 | 7.48E-13 |
| ENSMUST00000021512.9,Dhrs7-201,protein_coding,ENSMUSG00000021094.9,protein_coding | 2935.229786 | 0.633558144 | 6.77E-15 | 7.61E-13 |
| ENSMUST00000102749.10,Arnt-001,protein_coding,ENSMUSG00000015522.18,protein_coding | 4124.574979 | 0.293985029 | 6.80E-15 | 7.63E-13 |
| ENSMUST00000090945.4,Syt11-201,protein_coding,ENSMUSG00000068923.14,protein_coding | 1600.567847 | -0.560724437 | 7.34E-15 | 8.21E-13 |
| ENSMUST00000211995.1,Tk2-002,nonsense_mediated_decay,ENSMUSG00000035824.7,protein_coding | 875.6088681 | 0.482923387 | 7.48E-15 | 8.34E-13 |
| ENSMUST00000023550.7,Pdia5-201,protein_coding,ENSMUSG00000022844.7,protein_coding | 2757.734739 | 0.437705148 | 7.57E-15 | 8.42E-13 |
| ENSMUST00000187005.6,Wdfy1-013,nonsense_mediated_decay,ENSMUSG00000073643.11,protein_coding | 1493.434527 | 0.36534858 | 8.76E-15 | 9.70E-13 |
| ENSMUST00000098645.10,Dda1-009,retained_intron,ENSMUSG00000074247.10,protein_coding | 1391.744885 | 0.562663679 | 8.76E-15 | 9.70E-13 |
| ENSMUST00000126717.7,Rnf44-006,protein_coding,ENSMUSG00000034928.16,protein_coding | 242.8529233 | 1.171744118 | 9.20E-15 | 1.02E-12 |
| ENSMUST00000067072.3,Cthrc1-001,protein_coding,ENSMUSG00000054196.6,protein_coding | 3167.152625 | -0.485442264 | 9.60E-15 | 1.06E-12 |
| ENSMUST00000061618.7,Patl1-201,protein_coding,ENSMUSG00000046139.7,protein_coding | 11216.27308 | 0.26139572 | 1.01E-14 | 1.11E-12 |
| ENSMUST00000028378.3,Galnt3-001,protein_coding,ENSMUSG00000026994.9,protein_coding | 521.0004123 | 0.635716434 | 1.05E-14 | 1.15E-12 |
| ENSMUST00000052999.12,Echdc2-001,protein_coding,ENSMUSG00000028601.18,protein_coding | 1933.04821 | -0.37685772 | 1.08E-14 | 1.18E-12 |
| ENSMUST00000075281.6,Mark3-201,protein_coding,ENSMUSG00000007411.15,protein_coding | 4681.968695 | 0.314712044 | 1.15E-14 | 1.26E-12 |
| ENSMUST00000047762.7,Gata6-201,protein_coding,ENSMUSG00000005836.8,protein_coding | 9051.829431 | -0.296981526 | 1.17E-14 | 1.27E-12 |
| ENSMUST00000196506.1,Tm4sf1-003,protein_coding,ENSMUSG00000027800.14,protein_coding | 10645.08749 | -0.506341112 | 1.23E-14 | 1.33E-12 |
| ENSMUST00000118317.7,Hipk1-001,protein_coding,ENSMUSG00000008730.17,protein_coding | 10715.62782 | -0.386940981 | 1.27E-14 | 1.37E-12 |
| ENSMUST00000086861.11,Fam132b-001,protein_coding,ENSMUSG00000047443.14,protein_coding | 1858.762794 | 0.548877385 | 1.28E-14 | 1.38E-12 |
| ENSMUST00000047431.10,AU040320-003,protein_coding,ENSMUSG00000028830.14,protein_coding | 6928.585711 | 0.470981053 | 1.28E-14 | 1.38E-12 |
| ENSMUST00000049355.10,Mapk6-001,protein_coding,ENSMUSG00000042688.16,protein_coding | 10055.69317 | -0.392904512 | 1.32E-14 | 1.42E-12 |
| ENSMUST00000024742.7,Nfkbie-201,protein_coding,ENSMUSG00000023947.7,protein_coding | 252.9571481 | -0.744837265 | 1.37E-14 | 1.46E-12 |
| ENSMUST00000029650.8,Ints12-001,protein_coding,ENSMUSG00000028016.9,protein_coding | 4119.775499 | 0.336024442 | 1.38E-14 | 1.47E-12 |
| ENSMUST00000028241.6,Stom-001,protein_coding,ENSMUSG00000026880.11,protein_coding | 7827.629389 | 0.31657853 | 1.39E-14 | 1.48E-12 |
| ENSMUST00000034570.6,Pts-001,protein_coding,ENSMUSG00000032067.6,protein_coding | 691.7502335 | 0.527461325 | 1.45E-14 | 1.54E-12 |
| ENSMUST00000060404.4,Tor1aip2-001,protein_coding,ENSMUSG00000050565.16,protein_coding | 6613.776309 | 0.34689065 | 1.60E-14 | 1.70E-12 |
| ENSMUST00000120595.7,Eml2-002,protein_coding,ENSMUSG00000040811.15,protein_coding | 3528.39378 | 0.41760721 | 1.62E-14 | 1.71E-12 |
| ENSMUST00000031779.16,Calu-001,protein_coding,ENSMUSG00000029767.16,protein_coding | 63084.24713 | 0.254355487 | 1.66E-14 | 1.75E-12 |
| ENSMUST00000075994.10,Pdzrn3-001,protein_coding,ENSMUSG00000035357.16,protein_coding | 3666.691788 | -0.551580354 | 1.69E-14 | 1.77E-12 |
| ENSMUST00000020546.2,Stc2-001,protein_coding,ENSMUSG00000020303.2,protein_coding | 1097.017873 | 0.642641429 | 1.69E-14 | 1.77E-12 |
| ENSMUST00000025802.9,Nudt8-001,protein_coding,ENSMUSG00000110949.1,protein_coding | 2290.202134 | 0.393131289 | 1.74E-14 | 1.82E-12 |
| ENSMUST00000112439.8,Cops7a-002,protein_coding,ENSMUSG00000030127.15,protein_coding | 7266.808949 | 0.397246137 | 1.78E-14 | 1.86E-12 |
| ENSMUST00000073792.9,Mon2-202,protein_coding,ENSMUSG00000034602.13,protein_coding | 555.3255488 | 1.28571928 | 1.86E-14 | 1.94E-12 |
| ENSMUST00000054963.9,Fdft1-201,protein_coding,ENSMUSG00000021273.9,protein_coding | 20641.48075 | 0.462797421 | 1.90E-14 | 1.98E-12 |
| ENSMUST00000114225.7,Gng12-006,protein_coding,ENSMUSG00000036402.13,protein_coding | 792.5437614 | -0.560486843 | 2.08E-14 | 2.16E-12 |
| ENSMUST00000117291.7,Mthfd1l-001,protein_coding,ENSMUSG00000040675.17,protein_coding | 19935.21995 | 0.364609641 | 2.09E-14 | 2.16E-12 |
| ENSMUST00000165126.8,Uba52-201,protein_coding,ENSMUSG00000090137.8,protein_coding | 27664.39939 | 0.343313849 | 2.10E-14 | 2.17E-12 |
| ENSMUST00000109784.1,Nol4l-006,protein_coding,ENSMUSG00000061411.12,protein_coding | 4409.535486 | -0.364033043 | 2.11E-14 | 2.17E-12 |
| ENSMUST00000067853.5,Tmem119-001,protein_coding,ENSMUSG00000054675.5,protein_coding | 3205.010827 | -0.576019133 | 2.18E-14 | 2.24E-12 |
| ENSMUST00000102672.4,Nfe2l2-001,protein_coding,ENSMUSG00000015839.6,protein_coding | 14839.86236 | 0.240591038 | 2.30E-14 | 2.36E-12 |
| ENSMUST00000182500.7,Xpnpep1-004,nonsense_mediated_decay,ENSMUSG00000025027.17,protein_coding | 1771.587145 | 0.365760988 | 2.33E-14 | 2.38E-12 |
| ENSMUST00000063690.3,Dhrs9-001,protein_coding,ENSMUSG00000027068.6,protein_coding | 3144.095499 | 0.496362726 | 2.36E-14 | 2.40E-12 |
| ENSMUST00000027322.13,Rhbdd1-001,protein_coding,ENSMUSG00000026142.15,protein_coding | 10199.9171 | 0.596552624 | 2.42E-14 | 2.46E-12 |
| ENSMUST00000026475.14,Ddit3-001,protein_coding,ENSMUSG00000025408.15,protein_coding | 12249.76893 | 0.836935404 | 2.49E-14 | 2.53E-12 |
| ENSMUST00000032597.11,Rpl28-001,protein_coding,ENSMUSG00000030432.12,protein_coding | 47076.39676 | 0.284178338 | 2.51E-14 | 2.55E-12 |
| ENSMUST00000094154.5,Serpinh1-001,protein_coding,ENSMUSG00000070436.12,protein_coding | 64529.76715 | 0.349704652 | 2.55E-14 | 2.57E-12 |
| ENSMUST00000065330.7,Clk3-001,protein_coding,ENSMUSG00000032316.8,protein_coding | 3871.035943 | 0.513487444 | 2.64E-14 | 2.66E-12 |
| ENSMUST00000006614.2,Epha2-001,protein_coding,ENSMUSG00000006445.3,protein_coding | 54874.79623 | 0.318529295 | 2.73E-14 | 2.75E-12 |
| ENSMUST00000102911.9,Slc44a1-001,protein_coding,ENSMUSG00000028412.17,protein_coding | 2298.925983 | 0.374684587 | 2.85E-14 | 2.86E-12 |
| ENSMUST00000079300.12,Arhgef26-001,protein_coding,ENSMUSG00000036885.14,protein_coding | 304.156451 | -0.729272386 | 3.02E-14 | 3.03E-12 |
| ENSMUST00000172785.7,H2-D1-001,protein_coding,ENSMUSG00000073411.11,protein_coding | 11044.15357 | 0.34130884 | 3.05E-14 | 3.05E-12 |
| ENSMUST00000041023.13,Ipo9-001,protein_coding,ENSMUSG00000041879.13,protein_coding | 11913.52914 | -0.357766739 | 3.06E-14 | 3.05E-12 |
| ENSMUST00000016771.12,Myh9-001,protein_coding,ENSMUSG00000022443.16,protein_coding | 318494.312 | 0.204689477 | 3.14E-14 | 3.12E-12 |
| ENSMUST00000050630.13,Supt3-001,protein_coding,ENSMUSG00000038954.14,protein_coding | 2777.417895 | 0.493929009 | 3.14E-14 | 3.12E-12 |
| ENSMUST00000006618.8,Arhgef19-001,protein_coding,ENSMUSG00000028919.11,protein_coding | 1693.360195 | -0.404142567 | 3.22E-14 | 3.19E-12 |
| ENSMUST00000110778.1,Wipi2-003,protein_coding,ENSMUSG00000029578.15,protein_coding | 4302.092656 | 0.392682557 | 3.76E-14 | 3.71E-12 |
| ENSMUST00000134674.7,Mtmr2-003,nonsense_mediated_decay,ENSMUSG00000031918.16,protein_coding | 6034.237987 | 0.401252482 | 3.81E-14 | 3.75E-12 |
| ENSMUST00000120683.7,Hdac7-004,protein_coding,ENSMUSG00000022475.18,protein_coding | 353.2992937 | 1.080205558 | 3.81E-14 | 3.75E-12 |
| ENSMUST00000129025.7,Ulk2-002,processed_transcript,ENSMUSG00000004798.14,protein_coding | 1164.083481 | 0.438800613 | 3.92E-14 | 3.84E-12 |
| ENSMUST00000159814.1,Arl4c-001,protein_coding,ENSMUSG00000049866.12,protein_coding | 7184.343712 | -0.332503146 | 3.96E-14 | 3.87E-12 |
| ENSMUST00000031399.12,Psph-001,protein_coding,ENSMUSG00000029446.14,protein_coding | 22122.71731 | 0.486900679 | 4.02E-14 | 3.93E-12 |
| ENSMUST00000106282.7,Bcl7c-002,nonsense_mediated_decay,ENSMUSG00000030814.17,protein_coding | 755.1623246 | -0.525958079 | 4.27E-14 | 4.15E-12 |
| ENSMUST00000026243.3,Mgea5-201,protein_coding,ENSMUSG00000025220.7,protein_coding | 12214.69402 | 0.289620056 | 4.29E-14 | 4.17E-12 |
| ENSMUST00000107704.1,Msantd3-003,protein_coding,ENSMUSG00000039693.11,protein_coding | 2845.945815 | 0.339445138 | 4.31E-14 | 4.18E-12 |
| ENSMUST00000156800.1,Dpm1-004,retained_intron,ENSMUSG00000078919.10,protein_coding | 594.5250202 | 0.857202601 | 4.43E-14 | 4.29E-12 |
| ENSMUST00000027297.10,Plekhb2-001,protein_coding,ENSMUSG00000026123.11,protein_coding | 14127.13931 | 0.285505145 | 4.48E-14 | 4.32E-12 |
| ENSMUST00000053265.6,Lyrm4-201,protein_coding,ENSMUSG00000046573.6,protein_coding | 1221.610252 | 0.513644219 | 4.62E-14 | 4.45E-12 |
| ENSMUST00000085404.9,Dtd2-002,protein_coding,ENSMUSG00000020956.14,protein_coding | 424.9866523 | 0.649050279 | 4.75E-14 | 4.56E-12 |
| ENSMUST00000112862.6,Arpc5l-001,protein_coding,ENSMUSG00000026755.14,protein_coding | 8476.612853 | 0.395466753 | 4.78E-14 | 4.57E-12 |
| ENSMUST00000106393.7,Hyi-001,nonsense_mediated_decay,ENSMUSG00000006395.16,polymorphic_pseudogene | 520.6407129 | 0.63767487 | 4.78E-14 | 4.57E-12 |
| ENSMUST00000192355.5,Map4k4-006,protein_coding,ENSMUSG00000026074.14,protein_coding | 1111.549972 | 0.414493609 | 5.42E-14 | 5.17E-12 |
| ENSMUST00000203220.2,Hnrnpa2b1-007,protein_coding,ENSMUSG00000004980.16,protein_coding | 34626.90221 | -0.544172772 | 5.46E-14 | 5.20E-12 |
| ENSMUST00000107808.2,Tomm5-003,protein_coding,ENSMUSG00000078713.8,protein_coding | 983.94616 | 0.583910872 | 5.63E-14 | 5.35E-12 |
| ENSMUST00000032157.8,Gxylt2-001,protein_coding,ENSMUSG00000030074.9,protein_coding | 1799.819176 | -0.512916163 | 5.76E-14 | 5.46E-12 |
| ENSMUST00000036418.9,Ciart-001,protein_coding,ENSMUSG00000038550.10,protein_coding | 1819.775206 | 0.758971293 | 5.87E-14 | 5.56E-12 |
| ENSMUST00000203653.2,Wdr19-001,protein_coding,ENSMUSG00000037890.13,protein_coding | 1183.930717 | -0.50268889 | 5.98E-14 | 5.65E-12 |
| ENSMUST00000107771.11,Ruvbl2-002,protein_coding,ENSMUSG00000003868.14,protein_coding | 26275.23378 | 0.321531564 | 6.32E-14 | 5.96E-12 |
| ENSMUST00000168747.2,Atp10a-001,protein_coding,ENSMUSG00000025324.8,protein_coding | 3487.715349 | 0.37506978 | 6.52E-14 | 6.13E-12 |
| ENSMUST00000107032.2,Arhgef17-001,protein_coding,ENSMUSG00000032875.8,protein_coding | 9069.984479 | -0.252201267 | 6.65E-14 | 6.23E-12 |

| | | | | |
|---|---|---|---|---|
| ENSMUST00000162049.1,Slc4a8-002,protein_coding,ENSMUSG00000023032.12,protein_coding | 674.1593461 | -0.465172475 | 6.77E-14 | 6.33E-12 |
| ENSMUST00000117179.8,Fgfr1-002,protein_coding,ENSMUSG00000031565.18,protein_coding | 5902.241431 | 0.369668953 | 6.88E-14 | 6.42E-12 |
| ENSMUST00000015796.8,Steap1-001,protein_coding,ENSMUSG00000015652.9,protein_coding | 6799.501221 | 0.594405862 | 7.10E-14 | 6.61E-12 |
| ENSMUST00000093157.12,Ap1g1-001,protein_coding,ENSMUSG00000031731.16,protein_coding | 10148.82739 | -0.367144934 | 7.47E-14 | 6.94E-12 |
| ENSMUST00000114151.9,Immt-004,protein_coding,ENSMUSG00000052337.15,protein_coding | 12949.81282 | -0.423573315 | 7.74E-14 | 7.18E-12 |
| ENSMUST00000110840.7,Dcun1d2-003,protein_coding,ENSMUSG00000038506.13,protein_coding | 611.6227862 | -0.614472818 | 7.75E-14 | 7.18E-12 |
| ENSMUST00000113535.8,Fip1l1-004,protein_coding,ENSMUSG00000029227.15,protein_coding | 11767.99735 | 0.348214898 | 7.96E-14 | 7.34E-12 |
| ENSMUST00000045356.7,Rpl37-201,protein_coding,ENSMUSG00000041841.7,protein_coding | 58414.98259 | 0.195724728 | 7.95E-14 | 7.34E-12 |
| ENSMUST00000146021.7,Epb41-004,processed_transcript,ENSMUSG00000028906.16,protein_coding | 8778.593207 | 0.561945855 | 8.68E-14 | 7.99E-12 |
| ENSMUST00000108838.3,Pcbp2-203,protein_coding,ENSMUSG00000056851.13,protein_coding | 39916.46207 | 0.314365798 | 8.71E-14 | 7.99E-12 |
| ENSMUST00000210576.1,Tnpo2-005,retained_intron,ENSMUSG00000031691.14,protein_coding | 4128.742011 | 0.336829925 | 8.84E-14 | 8.10E-12 |
| ENSMUST00000017981.9,Nkiras2-001,protein_coding,ENSMUSG00000017837.12,protein_coding | 4525.15008 | 0.342599107 | 8.86E-14 | 8.10E-12 |
| ENSMUST00000025906.10,Esrra-001,protein_coding,ENSMUSG00000024955.13,protein_coding | 3513.387302 | -0.277855441 | 8.92E-14 | 8.13E-12 |
| ENSMUST00000028004.10,Aldh9a1-001,protein_coding,ENSMUSG00000026687.14,protein_coding | 9782.568964 | 0.296135253 | 8.95E-14 | 8.15E-12 |
| ENSMUST00000078945.11,Grsf1-001,protein_coding,ENSMUSG00000044221.14,protein_coding | 22915.43852 | 0.193882515 | 9.00E-14 | 8.18E-12 |
| ENSMUST00000008733.14,Dnajb6-001,protein_coding,ENSMUSG00000029131.14,protein_coding | 10554.99839 | 0.326144286 | 9.21E-14 | 8.35E-12 |
| ENSMUST00000031255.14,Agpat9-002,protein_coding,ENSMUSG00000029314.14,protein_coding | 1275.667248 | 0.760783463 | 9.32E-14 | 8.42E-12 |
| ENSMUST00000154546.7,Aars-003,retained_intron,ENSMUSG00000031960.14,protein_coding | 2558.356678 | 0.336813568 | 9.38E-14 | 8.47E-12 |
| ENSMUST00000065504.15,Anxa7-201,protein_coding,ENSMUSG00000021814.16,protein_coding | 11316.51409 | 0.361425771 | 9.55E-14 | 8.60E-12 |
| ENSMUST00000110648.7,Trp53bp1-001,protein_coding,ENSMUSG00000043909.16,protein_coding | 3186.869208 | -0.382001295 | 9.60E-14 | 8.62E-12 |
| ENSMUST00000114434.8,Hoxa3-001,protein_coding,ENSMUSG00000079560.13,protein_coding | 1196.786149 | -0.446339816 | 9.63E-14 | 8.62E-12 |
| ENSMUST00000075995.6,Cand2-001,protein_coding,ENSMUSG00000030319.8,protein_coding | 4353.146257 | -0.474661999 | 9.63E-14 | 8.62E-12 |
| ENSMUST00000020171.11,Ctgf-001,protein_coding,ENSMUSG00000019997.11,protein_coding | 16690.51836 | -0.343924481 | 9.81E-14 | 8.76E-12 |
| ENSMUST00000069817.14,Prrc2b-003,protein_coding,ENSMUSG00000039262.16,protein_coding | 29337.25403 | -0.177660024 | 1.04E-13 | 9.26E-12 |
| ENSMUST00000058943.7,Ankrd34a-001,protein_coding,ENSMUSG00000049097.9,protein_coding | 497.3897543 | -0.604185423 | 1.05E-13 | 9.33E-12 |
| ENSMUST00000167377.2,Iqgap1-001,protein_coding,ENSMUSG00000030536.10,protein_coding | 84606.80861 | 0.275496226 | 1.08E-13 | 9.60E-12 |
| ENSMUST00000113755.7,Odf2-008,protein_coding,ENSMUSG00000026790.19,protein_coding | 826.7222787 | 0.579308558 | 1.09E-13 | 9.64E-12 |
| ENSMUST00000113444.7,Agfg1-004,protein_coding,ENSMUSG00000026159.13,protein_coding | 5402.605708 | 0.238435041 | 1.10E-13 | 9.74E-12 |
| ENSMUST00000211733.1,2900026A02Rik-003,protein_coding,ENSMUSG00000051339.10,protein_coding | 7389.0879 | -0.318948723 | 1.10E-13 | 9.74E-12 |
| ENSMUST00000070257.13,Ascc2-001,protein_coding,ENSMUSG00000020412.16,protein_coding | 10078.75856 | 0.494526955 | 1.12E-13 | 9.83E-12 |
| ENSMUST00000040381.14,Ncoa5-001,protein_coding,ENSMUSG00000039804.15,protein_coding | 5442.04244 | 0.41528757 | 1.14E-13 | 9.99E-12 |
| ENSMUST00000123781.7,Stk39-002,retained_intron,ENSMUSG00000027030.15,protein_coding | 1003.16508 | 1.020264337 | 1.15E-13 | 1.01E-11 |
| ENSMUST00000017188.13,Serpinb6a-002,protein_coding,ENSMUSG00000060147.15,protein_coding | 2987.701231 | 0.625814595 | 1.15E-13 | 1.01E-11 |
| ENSMUST00000204598.2,Kcmf1-002,protein_coding,ENSMUSG00000055239.10,protein_coding | 13387.64775 | 0.39522445 | 1.21E-13 | 1.05E-11 |
| ENSMUST00000167415.7,Gt(ROSA)26Sor-004,antisense,ENSMUSG00000086429.9,antisense | 202.6910035 | 0.831709806 | 1.27E-13 | 1.11E-11 |
| ENSMUST00000189896.1,Cdv3-002,protein_coding,ENSMUSG00000032803.15,protein_coding | 3085.397454 | 0.309406484 | 1.27E-13 | 1.11E-11 |
| ENSMUST00000040312.6,Trib3-001,protein_coding,ENSMUSG00000032715.9,protein_coding | 30427.70984 | 0.748810821 | 1.30E-13 | 1.12E-11 |
| ENSMUST00000000312.11,Cdh1-001,protein_coding,ENSMUSG00000000303.12,protein_coding | 2739.533173 | 0.517888366 | 1.37E-13 | 1.19E-11 |
| ENSMUST00000027752.14,Lamc1-001,protein_coding,ENSMUSG00000026478.14,protein_coding | 43663.80479 | 0.343899609 | 1.43E-13 | 1.23E-11 |
| ENSMUST00000107734.9,Kat7-001,protein_coding,ENSMUSG00000038909.16,protein_coding | 17799.13086 | 0.316149997 | 1.44E-13 | 1.24E-11 |
| ENSMUST00000153655.7,Zmynd8-013,protein_coding,ENSMUSG00000039671.18,protein_coding | 1064.720076 | 0.724014814 | 1.44E-13 | 1.24E-11 |
| ENSMUST00000001920.12,Aif1l-001,protein_coding,ENSMUSG00000001864.13,protein_coding | 2682.637003 | -0.443555703 | 1.45E-13 | 1.25E-11 |
| ENSMUST00000093107.11,P4ha2-001,protein_coding,ENSMUSG00000018906.14,protein_coding | 3712.894402 | 0.600305816 | 1.48E-13 | 1.27E-11 |
| ENSMUST00000047144.12,5031439G07Rik-001,protein_coding,ENSMUSG00000036046.14,protein_coding | 4459.291181 | -0.35014953 | 1.60E-13 | 1.37E-11 |
| ENSMUST00000113865.4,Rbpj-002,protein_coding,ENSMUSG00000039191.12,protein_coding | 15009.11243 | 0.508607402 | 1.64E-13 | 1.40E-11 |
| ENSMUST00000030357.14,Ndc1-003,nonsense_mediated_decay,ENSMUSG00000028614.14,protein_coding | 3287.472822 | 0.59774388 | 1.65E-13 | 1.40E-11 |
| ENSMUST00000045743.12,Prkab2-001,protein_coding,ENSMUSG00000038205.12,protein_coding | 4519.711881 | -0.291393909 | 1.67E-13 | 1.42E-11 |
| ENSMUST00000032949.13,Coro1a-001,protein_coding,ENSMUSG00000030707.15,protein_coding | 223.0178387 | -0.759028005 | 1.67E-13 | 1.42E-11 |
| ENSMUST00000072119.14,Ccnb1-001,protein_coding,ENSMUSG00000041431.16,protein_coding | 28600.32256 | -0.152387258 | 1.80E-13 | 1.53E-11 |
| ENSMUST00000109087.7,Gnas-201,protein_coding,ENSMUSG00000027523.19,protein_coding | 126066.1154 | 0.223919206 | 1.84E-13 | 1.55E-11 |
| ENSMUST00000062677.11,Tmem11-001,protein_coding,ENSMUSG00000043284.12,protein_coding | 6019.133312 | 0.356315343 | 1.85E-13 | 1.56E-11 |
| ENSMUST00000182944.7,Birc6-003,protein_coding,ENSMUSG00000024073.14,protein_coding | 8663.902402 | -0.425596009 | 1.86E-13 | 1.57E-11 |
| ENSMUST00000113820.8,Arpc2-001,protein_coding,ENSMUSG00000006304.14,protein_coding | 24578.38656 | 0.272445681 | 1.88E-13 | 1.58E-11 |
| ENSMUST00000111861.3,Pdcd6ip-002,protein_coding,ENSMUSG00000032504.15,protein_coding | 15823.44683 | 0.258004996 | 1.89E-13 | 1.59E-11 |
| ENSMUST00000023362.14,Ntan1-001,protein_coding,ENSMUSG00000022681.14,protein_coding | 2940.279266 | 0.304627865 | 1.93E-13 | 1.61E-11 |
| ENSMUST00000077220.13,Gtpbp6-001,protein_coding,ENSMUSG00000033434.15,protein_coding | 3303.358011 | 0.429653677 | 1.94E-13 | 1.62E-11 |
| ENSMUST00000197859.4,Ube2d3-001,protein_coding,ENSMUSG00000078578.9,protein_coding | 1936.295578 | -0.407036467 | 1.98E-13 | 1.65E-11 |
| ENSMUST00000051209.10,Peg3-001,protein_coding,ENSMUSG00000002265.16,protein_coding | 1798.105918 | -0.620322318 | 2.02E-13 | 1.68E-11 |
| ENSMUST00000177083.7,Asap1-005,protein_coding,ENSMUSG00000022377.16,protein_coding | 4971.118508 | 0.601931628 | 2.10E-13 | 1.74E-11 |
| ENSMUST00000024988.14,C3-001,protein_coding,ENSMUSG00000024164.15,protein_coding | 168.2900973 | -0.945075191 | 2.10E-13 | 1.74E-11 |
| ENSMUST00000142734.7,Lgals3-001,protein_coding,ENSMUSG00000050335.16,protein_coding | 1894.383551 | 0.512600869 | 2.11E-13 | 1.75E-11 |
| ENSMUST00000109605.3,Atf4-201,protein_coding,ENSMUSG00000042406.7,protein_coding | 117277.2596 | 0.384021193 | 2.25E-13 | 1.86E-11 |
| ENSMUST00000047040.3,Prkcdbp-001,protein_coding,ENSMUSG00000037060.3,protein_coding | 4446.24746 | 0.386585509 | 2.29E-13 | 1.89E-11 |
| ENSMUST00000046206.3,Rprd1a-201,protein_coding,ENSMUSG00000040446.3,protein_coding | 4294.733785 | -0.373403028 | 2.32E-13 | 1.91E-11 |
| ENSMUST00000162022.7,Glis3-001,protein_coding,ENSMUSG00000052942.13,protein_coding | 2094.303754 | -0.325794694 | 2.34E-13 | 1.92E-11 |
| ENSMUST00000042105.10,Ppp1r15a-001,protein_coding,ENSMUSG00000040435.12,protein_coding | 13470.73224 | 0.90566209 | 2.38E-13 | 1.95E-11 |
| ENSMUST00000134244.1,Cxcl12-004,retained_intron,ENSMUSG00000061353.11,protein_coding | 597.3692821 | -0.816889085 | 2.38E-13 | 1.95E-11 |
| ENSMUST00000053177.13,Wdfy3-001,protein_coding,ENSMUSG00000043940.15,protein_coding | 10873.26396 | -0.247499986 | 2.50E-13 | 2.04E-11 |
| ENSMUST00000195382.1,Gm38357-001,TEC,ENSMUSG00000103313.1,TEC | 487.815348 | -0.642999814 | 2.74E-13 | 2.23E-11 |
| ENSMUST00000105680.8,Rere-003,protein_coding,ENSMUSG00000039852.16,protein_coding | 2902.727727 | -0.753559639 | 2.84E-13 | 2.31E-11 |
| ENSMUST00000071698.12,Erlin1-201,protein_coding,ENSMUSG00000025198.16,protein_coding | 5305.840302 | 0.888723964 | 2.97E-13 | 2.41E-11 |
| ENSMUST00000061767.4,Crebzf-001,protein_coding,ENSMUSG00000051451.6,protein_coding | 3933.604686 | -0.541427379 | 2.98E-13 | 2.42E-11 |
| ENSMUST00000070949.4,Prelid2-201,protein_coding,ENSMUSG00000056671.6,protein_coding | 816.657665 | 0.527306883 | 3.18E-13 | 2.58E-11 |
| ENSMUST00000102727.2,Anxa6-002,protein_coding,ENSMUSG00000018340.13,protein_coding | 8978.000373 | -0.341111833 | 3.20E-13 | 2.58E-11 |
| ENSMUST00000113485.8,Sf1-012,nonsense_mediated_decay,ENSMUSG00000024949.17,protein_coding | 18798.07538 | 0.248911313 | 3.51E-13 | 2.83E-11 |
| ENSMUST00000117597.1,Fbxo7-002,protein_coding,ENSMUSG00000001786.14,protein_coding | 1507.519625 | 0.423694028 | 3.74E-13 | 3.01E-11 |
| ENSMUST00000093357.11,Wdr83-001,protein_coding,ENSMUSG00000005150.16,protein_coding | 1755.488397 | 0.374560037 | 3.79E-13 | 3.04E-11 |
| ENSMUST00000001809.14,Pabpc1-001,protein_coding,ENSMUSG00000022283.14,protein_coding | 388166.7916 | 0.245144591 | 3.81E-13 | 3.05E-11 |
| ENSMUST00000063339.13,Rarg-002,protein_coding,ENSMUSG00000001288.14,protein_coding | 7205.613086 | 0.295132079 | 4.15E-13 | 3.32E-11 |
| ENSMUST00000072357.13,Hk1-201,protein_coding,ENSMUSG00000037012.18,protein_coding | 14622.11591 | 0.272988396 | 4.23E-13 | 3.38E-11 |
| ENSMUST00000138703.7,Pfkp-001,protein_coding,ENSMUSG00000021196.13,protein_coding | 4371.572649 | 0.407682329 | 4.24E-13 | 3.38E-11 |
| ENSMUST00000155579.7,Sertad4-001,protein_coding,ENSMUSG00000016262.14,protein_coding | 965.6434961 | -0.512187191 | 4.27E-13 | 3.40E-11 |
| ENSMUST00000043726.7,Angptl6-001,protein_coding,ENSMUSG00000038742.7,protein_coding | 1164.842423 | 0.948859364 | 4.29E-13 | 3.40E-11 |
| ENSMUST00000152441.1,Rpl35-003,processed_transcript,ENSMUSG00000062997.6,protein_coding | 32475.40753 | 0.322776099 | 4.50E-13 | 3.57E-11 |
| ENSMUST00000036126.5,Parp2-201,protein_coding,ENSMUSG00000036023.5,protein_coding | 4834.530535 | 0.244482983 | 4.56E-13 | 3.61E-11 |
| ENSMUST00000072837.6,Foxk1-001,protein_coding,ENSMUSG00000056493.9,protein_coding | 11719.04276 | 0.286247545 | 4.62E-13 | 3.65E-11 |
| ENSMUST00000028062.7,Vim-001,protein_coding,ENSMUSG00000026728.9,protein_coding | 262746.4287 | 0.368894609 | 4.86E-13 | 3.83E-11 |
| ENSMUST00000182432.1,Pdlim1-003,protein_coding,ENSMUSG00000055044.12,protein_coding | 8266.556398 | 0.332990784 | 4.90E-13 | 3.86E-11 |
| ENSMUST00000030296.8,Txndc12-001,protein_coding,ENSMUSG00000028567.8,protein_coding | 6833.758837 | 0.33013978 | 4.98E-13 | 3.92E-11 |
| ENSMUST00000084325.9,B4galt2-004,nonsense_mediated_decay,ENSMUSG00000028541.14,protein_coding | 665.4014866 | -0.565555544 | 5.01E-13 | 3.92E-11 |
| ENSMUST00000058997.13,Zbtb7c-201,protein_coding,ENSMUSG00000044646.14,protein_coding | 1401.512018 | 0.422256559 | 5.01E-13 | 3.92E-11 |
| ENSMUST00000200373.4,Gatad2b-005,retained_intron,ENSMUSG00000042390.9,protein_coding | 133.6131594 | 1.078173789 | 5.39E-13 | 4.21E-11 |

| | | | | |
|---|---|---|---|---|
| ENSMUST00000128326.7,Phldb1-005,protein_coding,ENSMUSG00000048537.16,protein_coding | 2821.795555 | 0.377173659 | 5.51E-13 | 4.29E-11 |
| ENSMUST00000058762.14,Pam-001,protein_coding,ENSMUSG00000026335.16,protein_coding | 2736.984266 | -0.624696469 | 5.77E-13 | 4.49E-11 |
| ENSMUST00000161712.7,Alkbh1-002,nonsense_mediated_decay,ENSMUSG00000079036.10,protein_coding | 824.2030689 | 0.774450779 | 5.91E-13 | 4.60E-11 |
| ENSMUST00000023779.6,Nr4a1-201,protein_coding,ENSMUSG00000023034.6,protein_coding | 2674.073997 | -0.335815717 | 6.00E-13 | 4.65E-11 |
| ENSMUST00000068439.12,Pdlim1-001,protein_coding,ENSMUSG00000055044.12,protein_coding | 8350.701037 | 0.299278402 | 6.38E-13 | 4.94E-11 |
| ENSMUST00000167498.1,Gm6548-001,processed_pseudogene,ENSMUSG00000091549.1,processed_pseudogene | 499.8437501 | -0.527410902 | 6.79E-13 | 5.25E-11 |
| ENSMUST00000160272.7,Pard3-002,protein_coding,ENSMUSG00000025812.17,protein_coding | 3316.319646 | 0.393646346 | 7.24E-13 | 5.59E-11 |
| ENSMUST00000042334.14,Rpl10a-001,protein_coding,ENSMUSG00000037805.14,protein_coding | 74060.01064 | 0.353056938 | 7.26E-13 | 5.59E-11 |
| ENSMUST00000133514.7,Aldoa-006,protein_coding,ENSMUSG00000030695.14,protein_coding | 240.9246867 | 1.259201818 | 7.48E-13 | 5.75E-11 |
| ENSMUST00000026625.5,Clybl-201,protein_coding,ENSMUSG00000025545.9,protein_coding | 1297.587443 | 0.423118129 | 7.94E-13 | 6.09E-11 |
| ENSMUST00000079949.12,Ewsr1-002,protein_coding,ENSMUSG00000009079.16,protein_coding | 9396.908134 | 0.452477048 | 8.54E-13 | 6.54E-11 |
| ENSMUST00000127939.1,Rnf114-003,nonsense_mediated_decay,ENSMUSG00000006418.17,protein_coding | 4627.208277 | 0.375313016 | 8.57E-13 | 6.55E-11 |
| ENSMUST00000149835.1,Arl6ip4-002,protein_coding,ENSMUSG00000029404.15,protein_coding | 251.9296816 | 0.74043636 | 8.65E-13 | 6.60E-11 |
| ENSMUST00000085614.5,Usp12-001,protein_coding,ENSMUSG00000029640.17,protein_coding | 8655.660127 | -0.401671568 | 8.85E-13 | 6.74E-11 |
| ENSMUST00000034206.5,Bbs2-001,protein_coding,ENSMUSG00000031755.6,protein_coding | 1238.589979 | 0.387584898 | 8.90E-13 | 6.77E-11 |
| ENSMUST00000100461.6,Smurf1-003,protein_coding,ENSMUSG00000038780.14,protein_coding | 3996.750397 | 0.547174219 | 8.94E-13 | 6.79E-11 |
| ENSMUST00000214037.1,RP23-320D23.6-001,TEC,ENSMUSG00000111394.1,TEC | 4808.16719 | -0.294897799 | 9.00E-13 | 6.80E-11 |
| ENSMUST00000116403.8,Hnrnpk-001,protein_coding,ENSMUSG00000021546.17,protein_coding | 4706.747329 | -0.459048505 | 8.99E-13 | 6.80E-11 |
| ENSMUST00000102774.10,Sqstm1-001,protein_coding,ENSMUSG00000015837.15,protein_coding | 76978.87272 | 0.519123482 | 9.02E-13 | 6.81E-11 |
| ENSMUST00000085708.2,Stat2-201,protein_coding,ENSMUSG00000040033.15,protein_coding | 3323.477562 | 0.368457472 | 9.20E-13 | 6.94E-11 |
| ENSMUST00000163382.1,St13-003,protein_coding,ENSMUSG00000022403.14,protein_coding | 1119.818382 | 0.446153832 | 9.24E-13 | 6.95E-11 |
| ENSMUST00000054254.11,Proser2-002,protein_coding,ENSMUSG00000045319.13,protein_coding | 596.8917157 | 0.545852141 | 9.35E-13 | 7.02E-11 |
| ENSMUST00000105683.8,Car6-002,protein_coding,ENSMUSG00000028972.11,protein_coding | 673.3413897 | 1.184152401 | 9.37E-13 | 7.02E-11 |
| ENSMUST00000039694.11,Stard3nl-001,protein_coding,ENSMUSG00000003062.12,protein_coding | 3809.385822 | 0.450063068 | 9.72E-13 | 7.28E-11 |
| ENSMUST00000020927.8,Hs1bp3-201,protein_coding,ENSMUSG00000020605.8,protein_coding | 4446.879342 | 0.24721398 | 1.00E-12 | 7.50E-11 |
| ENSMUST00000110615.7,Serf2-003,protein_coding,ENSMUSG00000074884.12,protein_coding | 405.6450223 | 0.813352239 | 1.03E-12 | 7.69E-11 |
| ENSMUST00000151790.1,G3bp1-002,processed_transcript,ENSMUSG00000018583.5,protein_coding | 721.2020052 | 0.556172492 | 1.04E-12 | 7.78E-11 |
| ENSMUST00000021738.8,Gpr137b-201,protein_coding,ENSMUSG00000021306.10,protein_coding | 4358.314793 | 0.339058133 | 1.05E-12 | 7.81E-11 |
| ENSMUST00000076859.11,Plk3-001,protein_coding,ENSMUSG00000028680.14,protein_coding | 5108.419599 | 0.468511956 | 1.09E-12 | 8.11E-11 |
| ENSMUST00000174387.1,Usp3-011,protein_coding,ENSMUSG00000032376.12,protein_coding | 1431.481005 | 0.540412335 | 1.09E-12 | 8.11E-11 |
| ENSMUST00000058550.14,Ccni-001,protein_coding,ENSMUSG00000063015.12,protein_coding | 29192.20667 | 0.351074087 | 1.12E-12 | 8.27E-11 |
| ENSMUST00000204188.2,Hnrnpa2b1-005,protein_coding,ENSMUSG00000004980.16,protein_coding | 13735.82054 | 0.452089423 | 1.12E-12 | 8.27E-11 |
| ENSMUST00000133808.1,C330006A16Rik-001,lincRNA,ENSMUSG00000087679.2,lincRNA | 6387.56363 | 0.295798076 | 1.13E-12 | 8.31E-11 |
| ENSMUST00000107042.8,Fam168a-001,protein_coding,ENSMUSG00000029461.17,protein_coding | 6281.848991 | -0.449157546 | 1.15E-12 | 8.45E-11 |
| ENSMUST00000161576.7,Herpud1-001,protein_coding,ENSMUSG00000031770.16,protein_coding | 9872.487543 | 0.538154802 | 1.16E-12 | 8.49E-11 |
| ENSMUST00000169681.1,Eif4b-201,protein_coding,ENSMUSG00000058655.8,protein_coding | 163227.6187 | 0.276322494 | 1.22E-12 | 8.97E-11 |
| ENSMUST00000185641.6,Atp5g2-001,protein_coding,ENSMUSG00000062683.11,protein_coding | 31362.48271 | 0.292350809 | 1.26E-12 | 9.19E-11 |
| ENSMUST00000021120.5,Trim47-002,protein_coding,ENSMUSG00000020773.11,protein_coding | 2017.633983 | 0.488335199 | 1.26E-12 | 9.24E-11 |
| ENSMUST00000212981.1,Nudt21-003,protein_coding,ENSMUSG00000031754.10,protein_coding | 8838.100042 | 0.299076616 | 1.40E-12 | 1.02E-10 |
| ENSMUST00000154269.6,Pter-003,protein_coding,ENSMUSG00000026730.12,protein_coding | 598.4087857 | 0.478550958 | 1.41E-12 | 1.03E-10 |
| ENSMUST00000166811.8,Ncam1-002,protein_coding,ENSMUSG00000039542.16,protein_coding | 1712.192663 | 0.489384406 | 1.43E-12 | 1.04E-10 |
| ENSMUST00000049389.10,Zdhhc2-001,protein_coding,ENSMUSG00000039470.15,protein_coding | 1723.439027 | 0.343271057 | 1.44E-12 | 1.04E-10 |
| ENSMUST00000112028.9,Erlin1-001,protein_coding,ENSMUSG00000025198.16,protein_coding | 6867.26645 | 0.564965612 | 1.45E-12 | 1.05E-10 |
| ENSMUST00000058096.13,Whsc1-002,protein_coding,ENSMUSG00000057406.16,protein_coding | 13425.84976 | 0.413159308 | 1.49E-12 | 1.08E-10 |
| ENSMUST00000042498.13,Hdlbp-003,protein_coding,ENSMUSG00000034088.15,protein_coding | 89205.20816 | 0.233206315 | 1.53E-12 | 1.11E-10 |
| ENSMUST00000000287.8,Scpep1-001,protein_coding,ENSMUSG00000000278.10,protein_coding | 30367.20016 | 0.545436468 | 1.59E-12 | 1.15E-10 |
| ENSMUST00000171654.7,Thsd4-201,protein_coding,ENSMUSG00000032289.15,protein_coding | 395.3148595 | 0.85311872 | 1.63E-12 | 1.17E-10 |
| ENSMUST00000124780.7,Depdc5-005,nonsense_mediated_decay,ENSMUSG00000037426.17,protein_coding | 36.44091379 | 1.21422927 | 1.68E-12 | 1.21E-10 |
| ENSMUST00000114513.8,Gls-002,protein_coding,ENSMUSG00000026103.14,protein_coding | 7997.132648 | 0.361373525 | 1.68E-12 | 1.21E-10 |
| ENSMUST00000026649.13,Syngr2-001,protein_coding,ENSMUSG00000048277.15,protein_coding | 12884.92733 | -0.301450725 | 1.69E-12 | 1.21E-10 |
| ENSMUST00000038377.7,Btg1-201,protein_coding,ENSMUSG00000036478.7,protein_coding | 11048.56713 | 0.340515451 | 1.71E-12 | 1.22E-10 |
| ENSMUST00000022849.5,Tars-201,protein_coding,ENSMUSG00000022241.5,protein_coding | 54623.31573 | 0.342940879 | 1.73E-12 | 1.23E-10 |
| ENSMUST00000071044.12,Capg-003,protein_coding,ENSMUSG00000056737.14,protein_coding | 5511.369845 | 0.442899951 | 1.77E-12 | 1.26E-10 |
| ENSMUST00000049461.6,Cars2-001,protein_coding,ENSMUSG00000056228.10,protein_coding | 787.0661506 | -0.660514461 | 1.78E-12 | 1.27E-10 |
| ENSMUST00000122044.7,Csnk1e-002,protein_coding,ENSMUSG00000022433.18,protein_coding | 1723.200998 | -0.399469622 | 1.82E-12 | 1.29E-10 |
| ENSMUST00000147884.1,Txndc11-004,nonsense_mediated_decay,ENSMUSG00000022498.16,protein_coding | 278.9014556 | 0.742900263 | 1.91E-12 | 1.35E-10 |
| ENSMUST00000030269.13,Slc6a9-001,protein_coding,ENSMUSG00000028542.17,protein_coding | 4429.53233 | 0.646249181 | 1.92E-12 | 1.36E-10 |
| ENSMUST00000020957.12,Adi1-001,protein_coding,ENSMUSG00000020629.12,protein_coding | 7283.97931 | 0.326315464 | 1.99E-12 | 1.41E-10 |
| ENSMUST00000075418.14,Stard5-001,protein_coding,ENSMUSG00000046027.17,protein_coding | 2877.015419 | 0.883882623 | 2.01E-12 | 1.42E-10 |
| ENSMUST00000030398.9,Slc2a1-001,protein_coding,ENSMUSG00000028645.11,protein_coding | 11525.70214 | 0.396910245 | 2.11E-12 | 1.48E-10 |
| ENSMUST00000156314.7,Rnf20-002,protein_coding,ENSMUSG00000028309.14,protein_coding | 641.8878448 | -0.614598852 | 2.14E-12 | 1.50E-10 |
| ENSMUST00000090381.10,Exoc4-004,protein_coding,ENSMUSG00000029763.17,protein_coding | 470.0398045 | 0.640898635 | 2.21E-12 | 1.55E-10 |
| ENSMUST00000034245.15,Usb1-001,protein_coding,ENSMUSG00000031792.15,protein_coding | 4136.732068 | 0.307395886 | 2.27E-12 | 1.59E-10 |
| ENSMUST00000043517.7,Pvr-001,protein_coding,ENSMUSG00000040511.14,protein_coding | 5955.124104 | 0.410769416 | 2.30E-12 | 1.61E-10 |
| ENSMUST00000034375.10,Dus2-001,protein_coding,ENSMUSG00000031901.12,protein_coding | 1579.336823 | 0.509016957 | 2.34E-12 | 1.63E-10 |
| ENSMUST00000196695.4,Fubp1-008,protein_coding,ENSMUSG00000028034.15,protein_coding | 13340.95487 | 0.350005066 | 2.38E-12 | 1.66E-10 |
| ENSMUST00000045288.13,Tgfb2-001,protein_coding,ENSMUSG00000039239.14,protein_coding | 570.5306728 | -0.60096662 | 2.47E-12 | 1.72E-10 |
| ENSMUST00000087865.3,Ybx3-001,protein_coding,ENSMUSG00000030189.15,protein_coding | 99258.46664 | 0.311920627 | 2.50E-12 | 1.74E-10 |
| ENSMUST00000033939.12,Ikbkb-001,protein_coding,ENSMUSG00000031537.15,protein_coding | 4871.303873 | -0.26914823 | 2.53E-12 | 1.76E-10 |
| ENSMUST00000110388.8,Gphn-202,protein_coding,ENSMUSG00000047454.11,protein_coding | 3016.497069 | 0.273541243 | 2.61E-12 | 1.81E-10 |
| ENSMUST00000109161.2,Zfp64-003,protein_coding,ENSMUSG00000027551.15,protein_coding | 1025.652982 | 0.564598363 | 2.65E-12 | 1.83E-10 |
| ENSMUST00000108588.7,Dlg4-001,protein_coding,ENSMUSG00000020886.16,protein_coding | 4311.22224 | 0.377731557 | 2.70E-12 | 1.86E-10 |
| ENSMUST00000070191.9,Cebpg-002,protein_coding,ENSMUSG00000056216.9,protein_coding | 746.203709 | 0.645797245 | 2.73E-12 | 1.89E-10 |
| ENSMUST00000073466.12,Cryzl1-001,protein_coding,ENSMUSG00000058240.13,protein_coding | 4162.558316 | 0.653733902 | 2.91E-12 | 2.00E-10 |
| ENSMUST00000212276.1,2900026A02Rik-004,protein_coding,ENSMUSG00000051339.10,protein_coding | 136.5066815 | -1.212346427 | 2.91E-12 | 2.00E-10 |
| ENSMUST00000019220.15,Strn4-001,protein_coding,ENSMUSG00000030374.15,protein_coding | 1788.975874 | 0.484462896 | 2.93E-12 | 2.01E-10 |
| ENSMUST00000045004.9,Dopey2-201,protein_coding,ENSMUSG00000022946.8,protein_coding | 2987.343306 | 0.295802672 | 2.94E-12 | 2.01E-10 |
| ENSMUST00000034588.8,Apoa1-001,protein_coding,ENSMUSG00000032083.8,protein_coding | 49.59292027 | -1.204904433 | 2.96E-12 | 2.02E-10 |
| ENSMUST00000161250.7,Tmbim6-003,protein_coding,ENSMUSG00000023010.14,protein_coding | 1533.924939 | 0.815467983 | 2.98E-12 | 2.04E-10 |
| ENSMUST00000034215.7,Mt1-001,protein_coding,ENSMUSG00000031765.8,protein_coding | 4342.951534 | 0.308866929 | 3.03E-12 | 2.07E-10 |
| ENSMUST00000015712.14,Lpl-001,protein_coding,ENSMUSG00000015568.16,protein_coding | 11811.74663 | 0.365227881 | 3.24E-12 | 2.21E-10 |
| ENSMUST00000042729.15,Npnt-002,protein_coding,ENSMUSG00000040998.18,protein_coding | 3508.573766 | -0.636551178 | 3.25E-12 | 2.21E-10 |
| ENSMUST00000120539.7,Lmo4-001,protein_coding,ENSMUSG00000028266.17,protein_coding | 12732.16553 | 0.393984715 | 3.28E-12 | 2.22E-10 |
| ENSMUST00000077159.7,Mical3-001,protein_coding,ENSMUSG00000051586.16,protein_coding | 1288.96743 | 0.432246383 | 3.28E-12 | 2.22E-10 |
| ENSMUST00000111230.7,Tagln2-001,protein_coding,ENSMUSG00000026547.15,protein_coding | 84450.05528 | 0.311166422 | 3.34E-12 | 2.26E-10 |
| ENSMUST00000212270.1,Slc38a7-004,protein_coding,ENSMUSG00000036534.6,protein_coding | 6003.141061 | 0.566290823 | 3.62E-12 | 2.45E-10 |
| ENSMUST00000105410.9,Slc19a1-003,protein_coding,ENSMUSG00000001436.15,protein_coding | 5600.440432 | 0.328042007 | 3.80E-12 | 2.56E-10 |
| ENSMUST00000116376.8,Sh3bp5l-201,protein_coding,ENSMUSG00000013646.17,protein_coding | 969.5057439 | -0.406067254 | 3.81E-12 | 2.57E-10 |
| ENSMUST00000112198.2,Kdm5b-003,protein_coding,ENSMUSG00000042207.17,protein_coding | 4174.517104 | 0.394992467 | 3.82E-12 | 2.57E-10 |
| ENSMUST00000000394.13,Tbrg4-007,protein_coding,ENSMUSG00000000384.15,protein_coding | 10097.55248 | 0.273958879 | 3.91E-12 | 2.62E-10 |
| ENSMUST00000026991.15,Faf2-006,protein_coding,ENSMUSG00000025873.15,protein_coding | 3722.024256 | 0.376646218 | 3.93E-12 | 2.64E-10 |
| ENSMUST00000124954.1,Got1-003,processed_transcript,ENSMUSG00000025190.13,protein_coding | 73.61023978 | 1.068634239 | 4.13E-12 | 2.76E-10 |

| | | | | |
|---|---|---|---|---|
| ENSMUST00000029663.10,Aimp1-001,protein_coding,ENSMUSG00000028029.10,protein_coding | 9026.340242 | 0.216327036 | 4.30E-12 | 2.88E-10 |
| ENSMUST00000035724.4,Akna-001,protein_coding,ENSMUSG00000039158.11,protein_coding | 2326.373697 | 0.821561824 | 4.59E-12 | 3.06E-10 |
| ENSMUST00000025866.13,Vldlr-005,nonsense_mediated_decay,ENSMUSG00000024924.14,protein_coding | 2601.453852 | 0.775654991 | 4.70E-12 | 3.13E-10 |
| ENSMUST00000087328.3,Hspa1a-001,protein_coding,ENSMUSG00000091971.3,protein_coding | 1847.065807 | -0.312792774 | 4.81E-12 | 3.20E-10 |
| ENSMUST00000028780.3,Chac1-001,protein_coding,ENSMUSG00000027313.3,protein_coding | 14342.45082 | 0.557243649 | 4.85E-12 | 3.22E-10 |
| ENSMUST00000109767.8,Trmt1-002,protein_coding,ENSMUSG00000001909.15,protein_coding | 8233.449259 | 0.384507756 | 4.85E-12 | 3.22E-10 |
| ENSMUST00000138858.7,Aldh1l2-003,retained_intron,ENSMUSG00000020256.14,protein_coding | 243.6984631 | 1.225962794 | 4.91E-12 | 3.25E-10 |
| ENSMUST00000206577.1,Atxn2l-005,protein_coding,ENSMUSG00000032637.15,protein_coding | 17888.89871 | 0.390564278 | 4.97E-12 | 3.29E-10 |
| ENSMUST00000107229.1,Pip5k1a-005,processed_transcript,ENSMUSG00000028126.16,protein_coding | 507.6717102 | 0.661233974 | 5.10E-12 | 3.37E-10 |
| ENSMUST00000070681.6,Gid4-001,protein_coding,ENSMUSG00000018415.14,protein_coding | 6207.096916 | 0.33208248 | 5.17E-12 | 3.41E-10 |
| ENSMUST00000054865.12,Cfap69-001,protein_coding,ENSMUSG00000040473.15,protein_coding | 32.3436134 | -1.287281333 | 5.23E-12 | 3.44E-10 |
| ENSMUST00000028949.15,Nsfl1c-002,protein_coding,ENSMUSG00000027455.16,protein_coding | 6347.615555 | 0.340759798 | 5.29E-12 | 3.48E-10 |
| ENSMUST00000202622.3,Eif4h-001,protein_coding,ENSMUSG00000040731.13,protein_coding | 23669.57942 | -0.34764953 | 5.41E-12 | 3.55E-10 |
| ENSMUST00000016530.13,Mpped2-002,protein_coding,ENSMUSG00000016386.15,protein_coding | 585.2716124 | 0.5165046 | 5.49E-12 | 3.60E-10 |
| ENSMUST00000123194.7,Nrf1-008,protein_coding,ENSMUSG00000058440.14,protein_coding | 271.3434287 | 0.931445461 | 5.62E-12 | 3.68E-10 |
| ENSMUST00000167110.7,Nfe2l1-202,protein_coding,ENSMUSG00000038615.17,protein_coding | 13259.17824 | 0.741518647 | 5.67E-12 | 3.70E-10 |
| ENSMUST00000105682.8,Rere-001,protein_coding,ENSMUSG00000039852.16,protein_coding | 18011.79946 | 0.371908471 | 5.74E-12 | 3.74E-10 |
| ENSMUST00000102741.7,Btf3l4-002,protein_coding,ENSMUSG00000028568.16,protein_coding | 769.3294494 | 0.501646257 | 5.83E-12 | 3.80E-10 |
| ENSMUST00000173078.7,Otud4-001,protein_coding,ENSMUSG00000036990.13,protein_coding | 7006.431767 | -0.539517227 | 6.05E-12 | 3.94E-10 |
| ENSMUST00000190679.6,Cnppd1-003,protein_coding,ENSMUSG00000033159.14,protein_coding | 2513.384315 | 0.424711295 | 6.07E-12 | 3.94E-10 |
| ENSMUST00000102922.9,Pold2-001,protein_coding,ENSMUSG00000020471.11,protein_coding | 19662.00143 | 0.279179001 | 6.13E-12 | 3.98E-10 |
| ENSMUST00000040532.11,Smim14-007,nonsense_mediated_decay,ENSMUSG00000037822.12,protein_coding | 1302.007224 | 0.540814394 | 6.17E-12 | 3.99E-10 |
| ENSMUST00000006103.8,Ctns-001,protein_coding,ENSMUSG00000005949.9,protein_coding | 899.0726055 | 0.422987651 | 6.20E-12 | 4.01E-10 |
| ENSMUST00000054450.6,Pdcd2-001,protein_coding,ENSMUSG00000014771.12,protein_coding | 4279.086079 | 0.324158233 | 6.27E-12 | 4.04E-10 |
| ENSMUST00000016168.8,Lbp-001,protein_coding,ENSMUSG00000016024.9,protein_coding | 285.5023359 | 0.78683615 | 6.36E-12 | 4.10E-10 |
| ENSMUST00000092736.10,Cacnb1-002,protein_coding,ENSMUSG00000020882.17,protein_coding | 180.6272212 | 0.921751112 | 6.51E-12 | 4.19E-10 |
| ENSMUST00000081840.5,Rpl32-001,protein_coding,ENSMUSG00000057841.5,protein_coding | 159248.5298 | 0.305007392 | 6.60E-12 | 4.24E-10 |
| ENSMUST00000108661.7,Chd3-001,protein_coding,ENSMUSG00000018474.17,protein_coding | 18814.35064 | -0.188984711 | 6.62E-12 | 4.24E-10 |
| ENSMUST00000177094.7,Gnb1-003,protein_coding,ENSMUSG00000029064.15,protein_coding | 9079.546868 | 0.401657321 | 6.84E-12 | 4.38E-10 |
| ENSMUST00000205870.1,Gpi1-011,nonsense_mediated_decay,ENSMUSG00000036427.5,protein_coding | 9060.390958 | 0.307921874 | 7.01E-12 | 4.47E-10 |
| ENSMUST00000034756.14,Anxa2-001,protein_coding,ENSMUSG00000032231.14,protein_coding | 168353.9982 | 0.337533517 | 7.00E-12 | 4.47E-10 |
| ENSMUST00000078804.11,Rpl35a-001,protein_coding,ENSMUSG00000060636.13,protein_coding | 44009.78882 | 0.265770907 | 7.06E-12 | 4.50E-10 |
| ENSMUST00000113498.8,Fbln2-001,protein_coding,ENSMUSG00000064080.12,protein_coding | 143755.2434 | 0.230538258 | 7.08E-12 | 4.51E-10 |
| ENSMUST00000027127.13,Sf3b1-001,protein_coding,ENSMUSG00000025982.13,protein_coding | 33832.84668 | -0.335410639 | 7.14E-12 | 4.53E-10 |
| ENSMUST00000114608.2,Etl4-014,protein_coding,ENSMUSG00000036617.16,protein_coding | 1010.579389 | 0.640510778 | 7.14E-12 | 4.53E-10 |
| ENSMUST00000091263.11,Qsox2-002,protein_coding,ENSMUSG00000036327.18,protein_coding | 2645.25576 | 0.3314954 | 7.22E-12 | 4.58E-10 |
| ENSMUST00000189909.6,Zfas1-001,antisense,ENSMUSG00000074578.13,processed_transcript | 3672.697338 | 0.429441544 | 7.30E-12 | 4.62E-10 |
| ENSMUST00000091288.12,Prnp-001,protein_coding,ENSMUSG00000079037.9,protein_coding | 6518.752241 | 0.356646734 | 7.32E-12 | 4.63E-10 |
| ENSMUST00000034776.12,Cln6-001,protein_coding,ENSMUSG00000032245.12,protein_coding | 2470.417002 | 0.433909212 | 7.39E-12 | 4.66E-10 |
| ENSMUST00000097590.3,Lars-201,protein_coding,ENSMUSG00000024493.8,protein_coding | 43404.501 | 0.338405441 | 7.41E-12 | 4.67E-10 |
| ENSMUST00000057793.10,Adnp-002,protein_coding,ENSMUSG00000051149.15,protein_coding | 3533.556976 | -0.42703738 | 7.56E-12 | 4.76E-10 |
| ENSMUST00000172753.1,Hspa1b-001,protein_coding,ENSMUSG00000090877.3,protein_coding | 6295.361106 | -0.357019187 | 7.61E-12 | 4.78E-10 |
| ENSMUST00000168900.2,Krtcap2-004,protein_coding,ENSMUSG00000042747.12,protein_coding | 4760.451957 | 0.287080634 | 7.92E-12 | 4.97E-10 |
| ENSMUST00000091752.3,Hist1h3c-001,protein_coding,ENSMUSG00000069310.3,protein_coding | 798.5411288 | 0.47106801 | 8.20E-12 | 5.14E-10 |
| ENSMUST00000071982.6,Bmper-001,protein_coding,ENSMUSG00000031963.8,protein_coding | 1245.722244 | -0.492475541 | 8.23E-12 | 5.15E-10 |
| ENSMUST00000050214.7,Akap8l-201,protein_coding,ENSMUSG00000002625.9,protein_coding | 2857.830654 | 0.394703289 | 8.38E-12 | 5.24E-10 |
| ENSMUST00000174070.7,Snx21-004,protein_coding,ENSMUSG00000050373.13,protein_coding | 463.1865187 | 0.675660152 | 8.66E-12 | 5.40E-10 |
| ENSMUST00000082223.12,Rpl5-001,protein_coding,ENSMUSG00000058558.12,protein_coding | 77334.24089 | 0.215739129 | 9.01E-12 | 5.61E-10 |
| ENSMUST00000032427.14,Asun-001,protein_coding,ENSMUSG00000040250.14,protein_coding | 8263.152254 | 0.268757456 | 9.10E-12 | 5.66E-10 |
| ENSMUST00000026408.6,Gdf11-001,protein_coding,ENSMUSG00000025352.6,protein_coding | 1535.682724 | -0.534500658 | 9.18E-12 | 5.70E-10 |
| ENSMUST00000054080.14,Fam60a-001,protein_coding,ENSMUSG00000039985.14,protein_coding | 10801.16333 | 0.306197317 | 9.38E-12 | 5.81E-10 |
| ENSMUST00000023807.6,Igfbp6-001,protein_coding,ENSMUSG00000023046.6,protein_coding | 8742.15386 | 0.338901975 | 9.37E-12 | 5.81E-10 |
| ENSMUST00000178691.1,Ubl4a-201,protein_coding,ENSMUSG00000015290.15,protein_coding | 1789.846091 | 0.402245194 | 9.67E-12 | 5.98E-10 |
| ENSMUST00000154131.1,Cdk9-004,nonsense_mediated_decay,ENSMUSG00000009555.16,protein_coding | 81.5461371 | 1.105292999 | 9.69E-12 | 5.98E-10 |
| ENSMUST00000125113.7,Samd4-004,protein_coding,ENSMUSG00000021838.16,protein_coding | 345.7685133 | 0.888215523 | 9.78E-12 | 6.03E-10 |
| ENSMUST00000056919.8,Csnk2a2-001,protein_coding,ENSMUSG00000046707.9,protein_coding | 3291.930451 | 0.419911635 | 9.81E-12 | 6.04E-10 |
| ENSMUST00000111930.8,Osbpl6-002,protein_coding,ENSMUSG00000042359.18,protein_coding | 806.2997472 | 0.528214084 | 1.03E-11 | 6.36E-10 |
| ENSMUST00000087689.4,Prune2-201,protein_coding,ENSMUSG00000039126.9,protein_coding | 13289.49892 | 0.552742083 | 1.04E-11 | 6.40E-10 |
| ENSMUST00000167624.1,Glo1-202,protein_coding,ENSMUSG00000024026.12,protein_coding | 1481.37792 | 0.581251268 | 1.08E-11 | 6.60E-10 |
| ENSMUST00000041124.12,Zfp704-001,protein_coding,ENSMUSG00000040209.12,protein_coding | 2573.519218 | -0.504843859 | 1.11E-11 | 6.81E-10 |
| ENSMUST00000165766.8,Dnm2-001,protein_coding,ENSMUSG00000033335.17,protein_coding | 2846.557346 | 0.44033088 | 1.11E-11 | 6.82E-10 |
| ENSMUST00000025979.12,Aldh18a1-001,protein_coding,ENSMUSG00000025007.13,protein_coding | 65640.44843 | 0.401970727 | 1.13E-11 | 6.87E-10 |
| ENSMUST00000001412.16,Vps50-001,protein_coding,ENSMUSG00000001376.17,protein_coding | 5000.667097 | -0.575960226 | 1.17E-11 | 7.10E-10 |
| ENSMUST00000066994.6,Zfp395-201,protein_coding,ENSMUSG00000034522.9,protein_coding | 9444.944391 | 0.277804962 | 1.19E-11 | 7.27E-10 |
| ENSMUST00000148752.7,Wdr11-004,nonsense_mediated_decay,ENSMUSG00000042055.13,protein_coding | 28.93770598 | 1.156100216 | 1.20E-11 | 7.31E-10 |
| ENSMUST00000026846.10,Tyms-001,protein_coding,ENSMUSG00000025747.12,protein_coding | 3273.986346 | -0.413521713 | 1.21E-11 | 7.33E-10 |
| ENSMUST00000099084.8,Zmynd8-015,protein_coding,ENSMUSG00000039671.18,protein_coding | 3420.892996 | 0.586084344 | 1.22E-11 | 7.42E-10 |
| ENSMUST00000057676.6,Ubald2-001,protein_coding,ENSMUSG00000050628.6,protein_coding | 12087.38034 | 0.314294347 | 1.24E-11 | 7.50E-10 |
| ENSMUST00000171530.7,Ifrd1-007,nonsense_mediated_decay,ENSMUSG00000001627.11,protein_coding | 5936.67431 | 0.209561387 | 1.24E-11 | 7.51E-10 |
| ENSMUST00000114657.8,Fmr1-006,protein_coding,ENSMUSG00000000838.17,protein_coding | 2100.853422 | -0.854373399 | 1.25E-11 | 7.54E-10 |
| ENSMUST00000078432.4,Rpl28-002,protein_coding,ENSMUSG00000030432.12,protein_coding | 26190.58621 | 0.327193601 | 1.26E-11 | 7.59E-10 |
| ENSMUST00000021668.8,Npc2-201,protein_coding,ENSMUSG00000021242.8,protein_coding | 25048.84398 | 0.303675152 | 1.26E-11 | 7.59E-10 |
| ENSMUST00000107392.7,Anpep-001,protein_coding,ENSMUSG00000039062.15,protein_coding | 6190.564087 | 0.303286036 | 1.27E-11 | 7.62E-10 |
| ENSMUST00000017841.3,Ada-001,protein_coding,ENSMUSG00000017697.3,protein_coding | 1117.964296 | -0.4199322 | 1.28E-11 | 7.66E-10 |
| ENSMUST00000107559.2,Mef2d-004,protein_coding,ENSMUSG00000001419.17,protein_coding | 397.8951111 | 0.922408555 | 1.30E-11 | 7.78E-10 |
| ENSMUST00000176849.7,Hnrnpk-012,protein_coding,ENSMUSG00000021546.17,protein_coding | 9513.246764 | 0.300518212 | 1.32E-11 | 7.88E-10 |
| ENSMUST00000061568.8,Slc36a4-001,protein_coding,ENSMUSG00000043885.14,protein_coding | 1910.472105 | -0.42742335 | 1.34E-11 | 7.99E-10 |
| ENSMUST00000107913.9,Tpm2-201,protein_coding,ENSMUSG00000028464.16,protein_coding | 5917.898813 | 0.362356514 | 1.34E-11 | 8.02E-10 |
| ENSMUST00000114195.7,Jam2-001,protein_coding,ENSMUSG00000053062.15,protein_coding | 383.5164513 | -0.622070195 | 1.39E-11 | 8.30E-10 |
| ENSMUST00000018429.11,Pld2-001,protein_coding,ENSMUSG00000020828.13,protein_coding | 2521.171531 | 0.378136972 | 1.41E-11 | 8.41E-10 |
| ENSMUST00000023343.3,Atg3-001,protein_coding,ENSMUSG00000022663.3,protein_coding | 3887.404285 | 0.370171343 | 1.43E-11 | 8.49E-10 |
| ENSMUST00000020307.10,Anapc16-002,protein_coding,ENSMUSG00000020107.10,protein_coding | 2797.963384 | 0.403009579 | 1.44E-11 | 8.55E-10 |
| ENSMUST00000143840.1,Nadk-005,protein_coding,ENSMUSG00000029063.16,protein_coding | 164.644429 | 0.763553375 | 1.44E-11 | 8.55E-10 |
| ENSMUST00000022629.8,Dpysl2-001,protein_coding,ENSMUSG00000022048.8,protein_coding | 8561.467348 | 0.358453599 | 1.45E-11 | 8.60E-10 |
| ENSMUST00000136507.3,Psph-002,protein_coding,ENSMUSG00000029446.14,protein_coding | 177.0414916 | 0.895774843 | 1.45E-11 | 8.60E-10 |
| ENSMUST00000005077.6,Hspb1-001,protein_coding,ENSMUSG00000004951.10,protein_coding | 14783.10251 | -0.375026489 | 1.48E-11 | 8.71E-10 |
| ENSMUST00000152918.7,Rps16-002,retained_intron,ENSMUSG00000037563.15,protein_coding | 27937.31092 | 0.293352764 | 1.49E-11 | 8.80E-10 |
| ENSMUST00000022691.13,Hr-001,protein_coding,ENSMUSG00000022096.14,protein_coding | 6760.79395 | 0.258688556 | 1.55E-11 | 9.13E-10 |
| ENSMUST00000118623.1,Dpf2-002,protein_coding,ENSMUSG00000024826.14,protein_coding | 12004.35827 | 0.44565189 | 1.58E-11 | 9.31E-10 |
| ENSMUST00000156081.1,Gm12840-001,lincRNA,ENSMUSG00000086320.1,lincRNA | 5080.703853 | -0.594095154 | 1.66E-11 | 9.72E-10 |
| ENSMUST00000042661.7,Ttyh3-001,protein_coding,ENSMUSG00000036565.7,protein_coding | 10221.91833 | -0.240735235 | 1.66E-11 | 9.73E-10 |
| ENSMUST00000112143.3,Oasl1-002,protein_coding,ENSMUSG00000041827.15,protein_coding | 6265.349999 | 0.420017624 | 1.66E-11 | 9.73E-10 |

| | | | | |
|---|---|---|---|---|
| ENSMUST00000068700.6,Wdr6-001,protein_coding,ENSMUSG00000066357.6,protein_coding | 11469.87999 | -0.403626269 | 1.73E-11 | 1.01E-09 |
| ENSMUST00000042990.6,Serbp1-002,protein_coding,ENSMUSG00000036371.6,protein_coding | 6338.061488 | -0.617413054 | 1.81E-11 | 1.06E-09 |
| ENSMUST00000196736.1,Pla2g12a-003,retained_intron,ENSMUSG00000027999.15,protein_coding | 555.6460776 | 0.635354144 | 1.90E-11 | 1.11E-09 |
| ENSMUST00000112935.7,Nmnat3-001,protein_coding,ENSMUSG00000032456.13,protein_coding | 680.4339955 | 0.50081739 | 1.97E-11 | 1.15E-09 |
| ENSMUST00000033910.8,Leprotl1-001,protein_coding,ENSMUSG00000031513.8,protein_coding | 10988.06806 | 0.406384331 | 1.97E-11 | 1.15E-09 |
| ENSMUST00000067404.12,Fgfr1op2-001,protein_coding,ENSMUSG00000040242.14,protein_coding | 2812.63635 | 0.629748687 | 2.12E-11 | 1.23E-09 |
| ENSMUST00000024866.4,Xdh-201,protein_coding,ENSMUSG00000024066.8,protein_coding | 5993.386402 | 0.54369477 | 2.20E-11 | 1.28E-09 |
| ENSMUST00000215568.1,Rpsa-005,retained_intron,ENSMUSG00000032518.6,protein_coding | 10975.60642 | 0.417049224 | 2.22E-11 | 1.29E-09 |
| ENSMUST00000049057.4,Fam171a2-001,protein_coding,ENSMUSG00000034685.4,protein_coding | 1185.311258 | -0.385549206 | 2.38E-11 | 1.38E-09 |
| ENSMUST00000022507.11,Pspc1-001,protein_coding,ENSMUSG00000021938.11,protein_coding | 4678.719016 | -0.49487022 | 2.38E-11 | 1.38E-09 |
| ENSMUST00000105466.2,P4ha1-002,protein_coding,ENSMUSG00000019916.14,protein_coding | 14183.13192 | 0.3666449 | 2.43E-11 | 1.40E-09 |
| ENSMUST00000067476.8,Spcs3-001,protein_coding,ENSMUSG00000054408.9,protein_coding | 11128.03685 | -0.345142572 | 2.46E-11 | 1.42E-09 |
| ENSMUST00000151289.7,Sgms1-004,protein_coding,ENSMUSG00000040451.17,protein_coding | 751.3298706 | 0.461740775 | 2.48E-11 | 1.43E-09 |
| ENSMUST00000142698.7,Avil-003,protein_coding,ENSMUSG00000025432.11,protein_coding | 83.87154193 | 1.163282956 | 2.51E-11 | 1.44E-09 |
| ENSMUST00000196981.4,Lrrfip2-003,protein_coding,ENSMUSG00000032497.16,protein_coding | 4458.477168 | 0.297962874 | 2.53E-11 | 1.45E-09 |
| ENSMUST00000047257.14,Katnal1-001,protein_coding,ENSMUSG00000041298.15,protein_coding | 3039.241801 | 0.358889224 | 2.54E-11 | 1.46E-09 |
| ENSMUST00000203442.1,Gm44275-001,lincRNA,ENSMUSG00000107768.1,lincRNA | 448.7356528 | -0.690081739 | 2.56E-11 | 1.46E-09 |
| ENSMUST00000015107.12,Tlk2-008,protein_coding,ENSMUSG00000020694.16,protein_coding | 647.2696367 | -0.776087045 | 2.56E-11 | 1.46E-09 |
| ENSMUST00000129589.7,Azin1-003,nonsense_mediated_decay,ENSMUSG00000037458.14,protein_coding | 1929.684858 | -0.504367606 | 2.56E-11 | 1.46E-09 |
| ENSMUST00000161409.1,Zcrb1-003,nonsense_mediated_decay,ENSMUSG00000022635.9,protein_coding | 438.7003762 | 0.714678801 | 2.57E-11 | 1.47E-09 |
| ENSMUST00000181645.7,Raet1e-001,protein_coding,ENSMUSG00000053219.15,protein_coding | 1982.857631 | 0.506293188 | 2.57E-11 | 1.47E-09 |
| ENSMUST00000087225.5,Pnkd-004,protein_coding,ENSMUSG00000026179.14,protein_coding | 2048.56712 | -0.323642194 | 2.61E-11 | 1.49E-09 |
| ENSMUST00000023112.11,Pmm1-001,protein_coding,ENSMUSG00000022474.14,protein_coding | 7523.162801 | 0.356294896 | 2.65E-11 | 1.51E-09 |
| ENSMUST00000064883.13,Plekha2-001,protein_coding,ENSMUSG00000031557.16,protein_coding | 2075.196063 | -0.358747632 | 2.72E-11 | 1.55E-09 |
| ENSMUST00000105839.7,Usp48-003,protein_coding,ENSMUSG00000043411.15,protein_coding | 778.1488891 | 0.425452384 | 2.76E-11 | 1.56E-09 |
| ENSMUST00000030464.13,Pik3r3-001,protein_coding,ENSMUSG00000028698.13,protein_coding | 1262.032524 | -0.601661803 | 2.80E-11 | 1.58E-09 |
| ENSMUST00000034136.11,Gpt2-001,protein_coding,ENSMUSG00000031700.11,protein_coding | 22728.02316 | 0.460069557 | 2.89E-11 | 1.63E-09 |
| ENSMUST00000181873.7,5430416N02Rik-001,lincRNA,ENSMUSG00000097772.7,lincRNA | 1050.848249 | 0.658786689 | 2.92E-11 | 1.65E-09 |
| ENSMUST00000073192.13,Erlec1-004,protein_coding,ENSMUSG00000020311.17,protein_coding | 2822.288935 | 0.34357433 | 2.94E-11 | 1.66E-09 |
| ENSMUST00000048184.3,Pdcd7-001,protein_coding,ENSMUSG00000041837.4,protein_coding | 2495.876135 | 0.329947302 | 3.03E-11 | 1.70E-09 |
| ENSMUST00000130530.7,Igfbp4-004,processed_transcript,ENSMUSG00000017493.12,protein_coding | 9250.28713 | 0.270100094 | 3.08E-11 | 1.73E-09 |
| ENSMUST00000176745.7,Cux1-006,protein_coding,ENSMUSG00000029705.17,protein_coding | 14923.81326 | 0.189631683 | 3.09E-11 | 1.73E-09 |
| ENSMUST00000035861.5,Pex5-002,protein_coding,ENSMUSG00000005069.12,protein_coding | 4051.405368 | 0.360035241 | 3.14E-11 | 1.76E-09 |
| ENSMUST00000010239.5,Slc3a2-002,protein_coding,ENSMUSG00000010095.13,protein_coding | 40013.77095 | 0.446206513 | 3.16E-11 | 1.77E-09 |
| ENSMUST00000194649.1,Rpl22l1-002,protein_coding,ENSMUSG00000039221.10,protein_coding | 2927.333611 | -0.329242445 | 3.22E-11 | 1.80E-09 |
| ENSMUST00000182115.1,Gapdh-016,processed_transcript,ENSMUSG00000057666.18,protein_coding | 13140.99006 | 0.507455073 | 3.27E-11 | 1.83E-09 |
| ENSMUST00000062069.5,Ptgs1-001,protein_coding,ENSMUSG00000047250.13,protein_coding | 17669.83981 | 0.31738458 | 3.40E-11 | 1.90E-09 |
| ENSMUST00000050785.13,Lcn2-001,protein_coding,ENSMUSG00000026822.14,protein_coding | 84.52788202 | 1.103212816 | 3.49E-11 | 1.94E-09 |
| ENSMUST00000047883.9,Tspan15-201,protein_coding,ENSMUSG00000037031.9,protein_coding | 3958.719169 | 0.314894168 | 3.58E-11 | 1.99E-09 |
| ENSMUST00000191436.6,Ly6e-002,protein_coding,ENSMUSG00000022587.14,protein_coding | 15236.53554 | 0.259741078 | 3.58E-11 | 1.99E-09 |
| ENSMUST00000116584.1,Mrpl39-001,protein_coding,ENSMUSG00000022889.7,protein_coding | 6385.319766 | 0.207129271 | 3.71E-11 | 2.06E-09 |
| ENSMUST00000117600.7,Hmga1-001,protein_coding,ENSMUSG00000046711.15,protein_coding | 48290.98364 | 0.291443202 | 3.86E-11 | 2.14E-09 |
| ENSMUST00000110272.8,Entpd5-001,protein_coding,ENSMUSG00000021236.16,protein_coding | 3100.591964 | 0.433956572 | 3.99E-11 | 2.20E-09 |
| ENSMUST00000079716.5,Rpl17-201,protein_coding,ENSMUSG00000062328.7,protein_coding | 80408.27306 | 0.254811073 | 3.98E-11 | 2.20E-09 |
| ENSMUST00000063042.10,Glb1-001,protein_coding,ENSMUSG00000045594.10,protein_coding | 4277.189245 | 0.299156243 | 4.16E-11 | 2.30E-09 |
| ENSMUST00000144236.7,Mcfd2-002,protein_coding,ENSMUSG00000024150.11,protein_coding | 1007.291504 | 0.604843147 | 4.25E-11 | 2.34E-09 |
| ENSMUST00000024748.13,Gtpbp2-001,protein_coding,ENSMUSG00000023952.13,protein_coding | 5671.905513 | 0.653104636 | 4.30E-11 | 2.37E-09 |
| ENSMUST00000036158.6,Slc35a4-001,protein_coding,ENSMUSG00000033272.12,protein_coding | 7297.295252 | -0.364910187 | 4.41E-11 | 2.42E-09 |
| ENSMUST00000100023.1,Mb21d2-201,protein_coding,ENSMUSG00000051065.8,protein_coding | 1834.397838 | 0.290334959 | 4.59E-11 | 2.52E-09 |
| ENSMUST00000032934.11,Aldoa-001,protein_coding,ENSMUSG00000030695.14,protein_coding | 174012.7888 | 0.280818625 | 4.60E-11 | 2.52E-09 |
| ENSMUST00000032185.8,Slc6a6-001,protein_coding,ENSMUSG00000030096.8,protein_coding | 52033.37054 | -0.344264627 | 4.66E-11 | 2.55E-09 |
| ENSMUST00000190840.7,Tenm3-005,protein_coding,ENSMUSG00000031561.16,protein_coding | 5344.581521 | -0.366252343 | 4.96E-11 | 2.71E-09 |
| ENSMUST00000127559.1,Slc2a4rg-ps-001,processed_transcript,ENSMUSG00000085028.2,unitary_pseudogene | 168.9007668 | -0.832319748 | 5.02E-11 | 2.74E-09 |
| ENSMUST00000204398.1,Anxa4-008,protein_coding,ENSMUSG00000029994.17,protein_coding | 809.7829645 | 0.557991361 | 5.05E-11 | 2.75E-09 |
| ENSMUST00000173945.1,Snx21-006,retained_intron,ENSMUSG00000050373.13,protein_coding | 309.9748064 | 0.719165456 | 5.21E-11 | 2.84E-09 |
| ENSMUST00000022529.6,Tkt-001,protein_coding,ENSMUSG00000021957.6,protein_coding | 63434.7763 | 0.270052228 | 5.35E-11 | 2.91E-09 |
| ENSMUST00000057792.8,Pon2-001,protein_coding,ENSMUSG00000032667.17,protein_coding | 7196.908543 | 0.263576003 | 5.38E-11 | 2.92E-09 |
| ENSMUST00000108987.7,Sept8-001,protein_coding,ENSMUSG00000018398.18,protein_coding | 11930.05037 | 0.31701008 | 5.38E-11 | 2.92E-09 |
| ENSMUST00000056427.8,Tubb2a-201,protein_coding,ENSMUSG00000058672.6,protein_coding | 6743.786289 | -0.439267981 | 5.38E-11 | 2.92E-09 |
| ENSMUST00000147210.7,Ptpn1-006,processed_transcript,ENSMUSG00000027540.13,protein_coding | 303.7681845 | 0.752670125 | 5.41E-11 | 2.93E-09 |
| ENSMUST00000138567.8,Mms22l-005,protein_coding,ENSMUSG00000045751.16,protein_coding | 1137.762226 | 0.412381672 | 5.44E-11 | 2.94E-09 |
| ENSMUST00000174806.7,Snx14-006,protein_coding,ENSMUSG00000032422.17,protein_coding | 268.2938174 | 1.2202032 | 5.58E-11 | 3.02E-09 |
| ENSMUST00000202305.3,Tmem248-005,protein_coding,ENSMUSG00000053094.15,protein_coding | 277.295512 | 0.950601636 | 5.75E-11 | 3.10E-09 |
| ENSMUST00000085254.6,Rdh11-002,protein_coding,ENSMUSG00000066441.14,protein_coding | 1682.053239 | 0.375975475 | 5.75E-11 | 3.10E-09 |
| ENSMUST00000087916.10,Hdac8-001,protein_coding,ENSMUSG00000067567.12,protein_coding | 1441.808741 | 0.36300634 | 5.80E-11 | 3.12E-09 |
| ENSMUST00000106008.2,Pddc1-001,protein_coding,ENSMUSG00000051007.15,protein_coding | 1388.718911 | 0.376415991 | 5.89E-11 | 3.17E-09 |
| ENSMUST00000089519.12,Aak1-001,protein_coding,ENSMUSG00000057230.13,protein_coding | 13560.65172 | -0.190241261 | 5.91E-11 | 3.17E-09 |
| ENSMUST00000050952.3,Stbd1-001,protein_coding,ENSMUSG00000047963.7,protein_coding | 7963.020147 | 0.670280023 | 6.14E-11 | 3.29E-09 |
| ENSMUST00000172407.7,Psma4-002,protein_coding,ENSMUSG00000032301.13,protein_coding | 276.9972952 | 0.942110408 | 6.21E-11 | 3.32E-09 |
| ENSMUST00000047368.7,Mnd1-001,protein_coding,ENSMUSG00000033752.7,protein_coding | 834.7504754 | 0.466008858 | 6.51E-11 | 3.48E-09 |
| ENSMUST00000049583.7,Zbtb40-001,protein_coding,ENSMUSG00000060862.9,protein_coding | 2432.204239 | -0.32205406 | 6.52E-11 | 3.48E-09 |
| ENSMUST00000025418.3,Psmg2-201,protein_coding,ENSMUSG00000024537.4,protein_coding | 3639.12936 | 0.229167453 | 6.51E-11 | 3.48E-09 |
| ENSMUST00000105365.8,Cirbp-001,protein_coding,ENSMUSG00000045193.13,protein_coding | 1633.844418 | 0.383659612 | 6.65E-11 | 3.54E-09 |
| ENSMUST00000021933.7,Ctsl-201,protein_coding,ENSMUSG00000021477.8,protein_coding | 25588.09539 | 0.303801809 | 7.02E-11 | 3.74E-09 |
| ENSMUST00000074989.5,Bcl9l-201,protein_coding,ENSMUSG00000063382.5,protein_coding | 7026.67772 | -0.197855336 | 7.06E-11 | 3.75E-09 |
| ENSMUST00000155493.1,Pafah1b1-001,protein_coding,ENSMUSG00000020745.15,protein_coding | 1571.936746 | 0.509065949 | 7.14E-11 | 3.79E-09 |
| ENSMUST00000080861.5,Rpl35-001,protein_coding,ENSMUSG00000062997.6,protein_coding | 48637.14275 | 0.26516518 | 7.19E-11 | 3.81E-09 |
| ENSMUST00000023918.12,Ivns1abp-001,protein_coding,ENSMUSG00000023150.14,protein_coding | 11240.56285 | -0.286437093 | 7.20E-11 | 3.81E-09 |
| ENSMUST00000114357.9,St3gal6-002,protein_coding,ENSMUSG00000022747.16,protein_coding | 1826.14688 | 0.378117298 | 7.36E-11 | 3.89E-09 |
| ENSMUST00000032992.6,Eif3c-001,protein_coding,ENSMUSG00000030738.11,protein_coding | 90370.4328 | 0.302270981 | 7.43E-11 | 3.92E-09 |
| ENSMUST00000059472.9,Mat2a-001,protein_coding,ENSMUSG00000053907.10,protein_coding | 16093.75925 | -0.582846849 | 7.46E-11 | 3.94E-09 |
| ENSMUST00000003870.13,Mxi1-201,protein_coding,ENSMUSG00000025025.13,protein_coding | 3103.948832 | 0.298455765 | 7.55E-11 | 3.98E-09 |
| ENSMUST00000061326.4,Uqcrq-001,protein_coding,ENSMUSG00000044894.14,protein_coding | 6879.259349 | 0.315089597 | 7.68E-11 | 4.04E-09 |
| ENSMUST00000142482.7,Dpm1-006,retained_intron,ENSMUSG00000078919.10,protein_coding | 194.5038967 | 1.0788826 | 7.81E-11 | 4.11E-09 |
| ENSMUST00000026699.14,Mkln1-001,protein_coding,ENSMUSG00000025609.15,protein_coding | 1896.970187 | -0.707011615 | 8.00E-11 | 4.20E-09 |
| ENSMUST00000025724.8,Il33-002,protein_coding,ENSMUSG00000024810.16,protein_coding | 826.6922041 | -0.659295446 | 8.16E-11 | 4.28E-09 |
| ENSMUST00000211840.1,Golga4-004,protein_coding,ENSMUSG00000038708.10,protein_coding | 500.4596658 | -0.5370792 | 8.34E-11 | 4.37E-09 |
| ENSMUST00000059562.13,Lhfp-001,protein_coding,ENSMUSG00000048332.13,protein_coding | 5823.904197 | -0.208923763 | 8.71E-11 | 4.56E-09 |
| ENSMUST00000115854.3,Dnaja3-002,protein_coding,ENSMUSG00000004069.16,protein_coding | 11959.12813 | 0.348547159 | 8.73E-11 | 4.56E-09 |
| ENSMUST00000046274.11,Ralgapb-201,protein_coding,ENSMUSG00000027652.15,protein_coding | 5481.917412 | -0.505806855 | 9.01E-11 | 4.70E-09 |
| ENSMUST00000144070.2,Mrps23-004,protein_coding,ENSMUSG00000023723.17,protein_coding | 1074.40381 | 0.349312815 | 9.18E-11 | 4.79E-09 |
| ENSMUST00000128187.7,Hnrnpdl-001,protein_coding,ENSMUSG00000029328.15,protein_coding | 5678.007955 | -0.471451931 | 9.24E-11 | 4.81E-09 |

| | | | | |
|---|---|---|---|---|
| ENSMUST00000060215.11,Srgap3-003,retained_intron,ENSMUSG00000030257.16,protein_coding | 412.9121 | 0.665610614 | 9.27E-11 | 4.82E-09 |
| ENSMUST00000090726.5,Slc43a3-001,protein_coding,ENSMUSG00000027074.14,protein_coding | 18007.8721 | 0.382760994 | 9.36E-11 | 4.86E-09 |
| ENSMUST00000128788.7,Ift20-001,protein_coding,ENSMUSG00000001105.15,protein_coding | 1576.240425 | 0.381249595 | 9.54E-11 | 4.95E-09 |
| ENSMUST00000026196.13,Got1-001,protein_coding,ENSMUSG00000025190.13,protein_coding | 23491.68344 | 0.466470655 | 9.67E-11 | 5.01E-09 |
| ENSMUST00000020397.9,Snrpd3-001,protein_coding,ENSMUSG00000020180.9,protein_coding | 9459.67626 | -0.355606392 | 9.77E-11 | 5.05E-09 |
| ENSMUST00000166931.1,Erh-002,protein_coding,ENSMUSG00000021131.13,protein_coding | 3885.036051 | 0.35719722 | 9.77E-11 | 5.05E-09 |
| ENSMUST00000053273.14,Itpr2-002,protein_coding,ENSMUSG00000030287.15,protein_coding | 1445.187903 | 0.691430122 | 9.91E-11 | 5.11E-09 |
| ENSMUST00000008179.6,Mid1ip1-002,protein_coding,ENSMUSG00000008035.12,protein_coding | 5213.256085 | 0.442360694 | 1.01E-10 | 5.23E-09 |
| ENSMUST00000101638.3,Limk2-001,protein_coding,ENSMUSG00000020451.17,protein_coding | 4114.435715 | 0.360426459 | 1.02E-10 | 5.25E-09 |
| ENSMUST00000025631.6,Ostf1-001,protein_coding,ENSMUSG00000024725.12,protein_coding | 7350.025067 | 0.338407765 | 1.11E-10 | 5.70E-09 |
| ENSMUST00000146453.1,Slc39a14-009,retained_intron,ENSMUSG00000022094.15,protein_coding | 233.2479615 | -0.668371649 | 1.11E-10 | 5.71E-09 |
| ENSMUST00000032057.7,Gfpt1-002,protein_coding,ENSMUSG00000029992.14,protein_coding | 1363.786865 | 0.611078104 | 1.13E-10 | 5.80E-09 |
| ENSMUST00000115248.9,C330007P06Rik-201,protein_coding,ENSMUSG00000006423.15,protein_coding | 4035.817147 | -0.532990389 | 1.17E-10 | 6.02E-09 |
| ENSMUST00000193547.5,Ncam1-001,protein_coding,ENSMUSG00000039542.16,protein_coding | 5331.379359 | 0.263739898 | 1.18E-10 | 6.06E-09 |
| ENSMUST00000078357.4,Emp2-001,protein_coding,ENSMUSG00000022505.8,protein_coding | 8398.961792 | 0.510387075 | 1.21E-10 | 6.17E-09 |
| ENSMUST00000109426.1,Arhgef28-201,protein_coding,ENSMUSG00000021662.9,protein_coding | 1489.351856 | 0.490968386 | 1.22E-10 | 6.20E-09 |
| ENSMUST00000098300.5,Alg8-001,protein_coding,ENSMUSG00000035704.17,protein_coding | 4079.131447 | 0.249231212 | 1.22E-10 | 6.23E-09 |
| ENSMUST00000022378.7,Ero1l-201,protein_coding,ENSMUSG00000021831.8,protein_coding | 48981.85997 | 0.64668583 | 1.24E-10 | 6.33E-09 |
| ENSMUST00000113811.7,Yipf6-002,protein_coding,ENSMUSG00000047694.12,protein_coding | 784.5363088 | 0.410607606 | 1.27E-10 | 6.48E-09 |
| ENSMUST00000109021.3,Uqcrq-003,protein_coding,ENSMUSG00000044894.14,protein_coding | 9942.938759 | 0.308654328 | 1.30E-10 | 6.60E-09 |
| ENSMUST00000134719.2,Ppp1cc-008,nonsense_mediated_decay,ENSMUSG00000004455.16,protein_coding | 283.0158114 | 1.068692512 | 1.31E-10 | 6.65E-09 |
| ENSMUST00000107162.7,Tenm4-001,protein_coding,ENSMUSG00000048078.16,protein_coding | 349.5219767 | 0.980578518 | 1.32E-10 | 6.67E-09 |
| ENSMUST00000108800.7,Tpd52l2-003,protein_coding,ENSMUSG00000000827.18,protein_coding | 563.9158426 | -0.721666726 | 1.32E-10 | 6.69E-09 |
| ENSMUST00000105686.2,Slc25a33-001,protein_coding,ENSMUSG00000028982.9,protein_coding | 3816.085956 | 0.420093066 | 1.32E-10 | 6.70E-09 |
| ENSMUST00000035086.12,Pdcd6ip-001,protein_coding,ENSMUSG00000032504.15,protein_coding | 9167.539881 | -0.250528166 | 1.36E-10 | 6.88E-09 |
| ENSMUST00000113365.7,Dnm1-001,protein_coding,ENSMUSG00000026825.17,protein_coding | 688.3819727 | 0.741516616 | 1.37E-10 | 6.90E-09 |
| ENSMUST00000066134.5,Rpia-001,protein_coding,ENSMUSG00000053604.5,protein_coding | 4140.091881 | 0.313936322 | 1.37E-10 | 6.90E-09 |
| ENSMUST00000046770.9,Nenf-001,protein_coding,ENSMUSG00000037499.9,protein_coding | 4366.666062 | 0.309122286 | 1.38E-10 | 6.94E-09 |
| ENSMUST00000141251.1,Emg1-002,retained_intron,ENSMUSG00000004268.11,protein_coding | 588.0622082 | 0.590325645 | 1.39E-10 | 6.99E-09 |
| ENSMUST00000128482.7,Rps15a-003,protein_coding,ENSMUSG00000008683.16,protein_coding | 5865.002928 | 0.296726625 | 1.39E-10 | 7.00E-09 |
| ENSMUST00000071095.13,Setd3-001,protein_coding,ENSMUSG00000056770.15,protein_coding | 10861.65541 | 0.375749913 | 1.43E-10 | 7.16E-09 |
| ENSMUST00000006760.2,Cdt1-001,protein_coding,ENSMUSG00000006585.3,protein_coding | 20449.78775 | 0.332172822 | 1.44E-10 | 7.23E-09 |
| ENSMUST00000174545.7,Xpo4-001,protein_coding,ENSMUSG00000021952.15,protein_coding | 7676.042125 | -0.422306687 | 1.45E-10 | 7.28E-09 |
| ENSMUST00000172501.7,1110038B12Rik-001,lincRNA,ENSMUSG00000092203.7,processed_transcript | 2579.345576 | 0.452089503 | 1.47E-10 | 7.34E-09 |
| ENSMUST00000166214.7,Wdr36-202,protein_coding,ENSMUSG00000038299.15,protein_coding | 10784.34531 | 0.311799008 | 1.49E-10 | 7.43E-09 |
| ENSMUST00000105909.3,Cars-002,protein_coding,ENSMUSG00000010755.17,protein_coding | 23380.27189 | 0.311068998 | 1.49E-10 | 7.44E-09 |
| ENSMUST00000080666.7,Ndrg4-002,protein_coding,ENSMUSG00000036564.17,protein_coding | 879.7638351 | -0.35221283 | 1.51E-10 | 7.52E-09 |
| ENSMUST00000105887.7,Dhdds-003,protein_coding,ENSMUSG00000012117.13,protein_coding | 1849.724968 | 0.361283512 | 1.52E-10 | 7.57E-09 |
| ENSMUST00000092512.10,P4ha1-003,protein_coding,ENSMUSG00000019916.14,protein_coding | 4793.101514 | 0.493259914 | 1.54E-10 | 7.65E-09 |
| ENSMUST00000065134.3,Ier5l-001,protein_coding,ENSMUSG00000089762.3,protein_coding | 2333.86359 | -0.536202109 | 1.57E-10 | 7.77E-09 |
| ENSMUST00000148042.2,Zmynd19-002,protein_coding,ENSMUSG00000026974.11,protein_coding | 3589.849183 | 0.245698032 | 1.60E-10 | 7.92E-09 |
| ENSMUST00000151085.1,Rnf10-005,processed_transcript,ENSMUSG00000041740.16,protein_coding | 83.06851995 | 1.124349097 | 1.62E-10 | 8.04E-09 |
| ENSMUST00000148248.2,Vim-004,retained_intron,ENSMUSG00000026728.9,protein_coding | 22690.92847 | 0.292354839 | 1.63E-10 | 8.08E-09 |
| ENSMUST00000032961.3,Nupr1-001,protein_coding,ENSMUSG00000030717.9,protein_coding | 3687.3515 | 0.611822644 | 1.65E-10 | 8.15E-09 |
| ENSMUST00000032492.8,Cd9-001,protein_coding,ENSMUSG00000030342.8,protein_coding | 17317.7764 | 0.261179952 | 1.66E-10 | 8.21E-09 |
| ENSMUST00000028280.13,Col5a1-001,protein_coding,ENSMUSG00000026837.15,protein_coding | 21056.87964 | 0.253956756 | 1.68E-10 | 8.26E-09 |
| ENSMUST00000021283.7,Pik3r5-001,protein_coding,ENSMUSG00000020901.13,protein_coding | 1619.157501 | 0.270734736 | 1.68E-10 | 8.26E-09 |
| ENSMUST00000065692.13,Ap2b1-005,protein_coding,ENSMUSG00000035152.14,protein_coding | 8928.905858 | 0.223315056 | 1.68E-10 | 8.26E-09 |
| ENSMUST00000009707.13,Tor2a-001,protein_coding,ENSMUSG00000009563.16,protein_coding | 2950.81856 | 0.225527277 | 1.69E-10 | 8.30E-09 |
| ENSMUST00000004955.13,Prpsap2-001,protein_coding,ENSMUSG00000020528.14,protein_coding | 1511.847477 | 0.405621839 | 1.72E-10 | 8.45E-09 |
| ENSMUST00000189262.6,Clasp1-007,protein_coding,ENSMUSG00000064302.13,protein_coding | 162.4927514 | 1.057742602 | 1.73E-10 | 8.46E-09 |
| ENSMUST00000046506.6,Clcf1-001,protein_coding,ENSMUSG00000040663.8,protein_coding | 4549.54546 | -0.219007276 | 1.74E-10 | 8.52E-09 |
| ENSMUST00000146080.7,Nadk-003,protein_coding,ENSMUSG00000029063.16,protein_coding | 279.038221 | 0.646380488 | 1.78E-10 | 8.72E-09 |
| ENSMUST00000161154.1,Wars-004,protein_coding,ENSMUSG00000021266.16,protein_coding | 1062.697706 | 0.760517997 | 1.79E-10 | 8.73E-09 |
| ENSMUST00000021649.7,Acot2-001,protein_coding,ENSMUSG00000021226.7,protein_coding | 5709.944562 | 0.652671241 | 1.82E-10 | 8.87E-09 |
| ENSMUST00000095083.10,Sptan1-002,protein_coding,ENSMUSG00000057738.13,protein_coding | 44296.55919 | 0.383626082 | 1.83E-10 | 8.93E-09 |
| ENSMUST00000037875.5,Rrbp1-001,protein_coding,ENSMUSG00000027422.15,protein_coding | 627.1824144 | -0.561791975 | 1.84E-10 | 8.96E-09 |
| ENSMUST00000016678.13,Lamp2-001,protein_coding,ENSMUSG00000016534.15,protein_coding | 5404.371107 | 0.297796659 | 1.97E-10 | 9.57E-09 |
| ENSMUST00000114444.8,Pcnp-001,protein_coding,ENSMUSG00000071533.11,protein_coding | 8069.868891 | -0.417206362 | 2.03E-10 | 9.84E-09 |
| ENSMUST00000048994.6,Nfkbil1-001,protein_coding,ENSMUSG00000042419.8,protein_coding | 2901.555183 | 0.364193519 | 2.05E-10 | 9.96E-09 |
| ENSMUST00000069712.8,Tpd52l2-004,protein_coding,ENSMUSG00000000827.18,protein_coding | 2703.085273 | 0.438128648 | 2.06E-10 | 9.99E-09 |
| ENSMUST00000040967.8,Vps37b-001,protein_coding,ENSMUSG00000066278.6,protein_coding | 7690.987234 | 0.221389514 | 2.06E-10 | 9.99E-09 |
| ENSMUST00000165582.1,St13-005,protein_coding,ENSMUSG00000022403.14,protein_coding | 1900.716741 | 0.499761963 | 2.09E-10 | 1.01E-08 |
| ENSMUST00000090219.12,Bmf-001,protein_coding,ENSMUSG00000040093.15,protein_coding | 129.9670647 | -0.845117148 | 2.13E-10 | 1.03E-08 |
| ENSMUST00000114233.2,Htra3-002,protein_coding,ENSMUSG00000029096.15,protein_coding | 851.9015002 | 0.651089565 | 2.14E-10 | 1.03E-08 |
| ENSMUST00000092891.5,Cacng7-001,protein_coding,ENSMUSG00000069806.5,protein_coding | 1586.367546 | -0.371853544 | 2.16E-10 | 1.04E-08 |
| ENSMUST00000027974.6,Atf6-001,protein_coding,ENSMUSG00000026663.6,protein_coding | 21023.60943 | 0.439788935 | 2.18E-10 | 1.05E-08 |
| ENSMUST00000018821.8,Slc25a39-001,protein_coding,ENSMUSG00000018677.9,protein_coding | 23122.97632 | 0.303969873 | 2.17E-10 | 1.05E-08 |
| ENSMUST00000084434.10,Rplp2-001,protein_coding,ENSMUSG00000025508.13,protein_coding | 49960.80205 | 0.272159269 | 2.23E-10 | 1.07E-08 |
| ENSMUST00000022169.7,Hexb-201,protein_coding,ENSMUSG00000021665.7,protein_coding | 3668.255414 | 0.293681379 | 2.23E-10 | 1.07E-08 |
| ENSMUST00000073926.6,Rps12-201,protein_coding,ENSMUSG00000061983.6,protein_coding | 123097.8589 | 0.235962869 | 2.26E-10 | 1.09E-08 |
| ENSMUST00000144826.1,Noct-002,protein_coding,ENSMUSG00000023087.15,protein_coding | 1353.989862 | -0.527814292 | 2.30E-10 | 1.10E-08 |
| ENSMUST00000035560.8,Acbd6-001,protein_coding,ENSMUSG00000033701.13,protein_coding | 628.4741845 | -0.451121616 | 2.36E-10 | 1.13E-08 |
| ENSMUST00000111350.9,Uap1-002,protein_coding,ENSMUSG00000026670.15,protein_coding | 758.1720158 | 0.593437318 | 2.39E-10 | 1.14E-08 |
| ENSMUST00000056937.11,Hscb-001,protein_coding,ENSMUSG00000043510.12,protein_coding | 961.0992735 | 0.431780203 | 2.40E-10 | 1.15E-08 |
| ENSMUST00000199798.4,Kmt5a-011,nonsense_mediated_decay,ENSMUSG00000049327.17,protein_coding | 4630.947998 | 0.472531044 | 2.41E-10 | 1.15E-08 |
| ENSMUST00000165532.2,Rpl14-001,protein_coding,ENSMUSG00000025794.9,protein_coding | 75083.70696 | 0.304407493 | 2.41E-10 | 1.15E-08 |
| ENSMUST00000032330.15,Emp1-201,protein_coding,ENSMUSG00000030208.15,protein_coding | 113587.3896 | 0.226826009 | 2.42E-10 | 1.15E-08 |
| ENSMUST00000170122.3,Rps27-001,protein_coding,ENSMUSG00000090733.6,protein_coding | 39345.05686 | 0.263066532 | 2.44E-10 | 1.16E-08 |
| ENSMUST00000066160.2,Chek2-002,protein_coding,ENSMUSG00000029521.7,protein_coding | 3408.573277 | 0.405774914 | 2.47E-10 | 1.17E-08 |
| ENSMUST00000161816.7,Rad52-003,retained_intron,ENSMUSG00000030166.14,protein_coding | 270.1256485 | 0.775872166 | 2.48E-10 | 1.17E-08 |
| ENSMUST00000146640.7,Aldh1l2-002,protein_coding,ENSMUSG00000020256.14,protein_coding | 4596.49682 | 0.729762964 | 2.49E-10 | 1.18E-08 |
| ENSMUST00000053063.6,Hexim1-001,protein_coding,ENSMUSG00000048878.6,protein_coding | 3814.72911 | -0.502924674 | 2.53E-10 | 1.19E-08 |
| ENSMUST00000008812.7,Rps18-001,protein_coding,ENSMUSG00000008668.14,protein_coding | 123931.8243 | 0.272474346 | 2.53E-10 | 1.19E-08 |
| ENSMUST00000020454.10,Hmg20b-001,protein_coding,ENSMUSG00000020232.17,protein_coding | 3027.737449 | -0.406922972 | 2.65E-10 | 1.25E-08 |
| ENSMUST00000032884.15,Anapc15-005,retained_intron,ENSMUSG00000030649.18,protein_coding | 706.7789709 | 0.429764084 | 2.66E-10 | 1.25E-08 |
| ENSMUST00000019169.7,Use1-001,protein_coding,ENSMUSG00000002395.14,protein_coding | 3019.185592 | 0.291036551 | 2.68E-10 | 1.26E-08 |
| ENSMUST00000138693.7,Brd3-003,protein_coding,ENSMUSG00000026918.16,protein_coding | 3643.901268 | 0.416560755 | 2.76E-10 | 1.30E-08 |
| ENSMUST00000168360.1,P4hb-002,protein_coding,ENSMUSG00000025130.11,protein_coding | 4147.885367 | 0.382472151 | 2.85E-10 | 1.34E-08 |
| ENSMUST00000030336.10,Tnfrsf1b-001,protein_coding,ENSMUSG00000028599.10,protein_coding | 3773.449103 | 0.311195443 | 2.86E-10 | 1.34E-08 |
| ENSMUST00000032916.5,Maz-001,protein_coding,ENSMUSG00000030678.7,protein_coding | 6163.302945 | 0.448154607 | 2.87E-10 | 1.35E-08 |
| ENSMUST00000099576.8,Fmn1-201,protein_coding,ENSMUSG00000044042.18,protein_coding | 5345.859882 | -0.583712707 | 2.89E-10 | 1.35E-08 |

| | | | | |
|---|---|---|---|---|
| ENSMUST00000113134.7,Ugt1a6a-002,protein_coding,ENSMUSG00000054545.17,protein_coding | 4596.919468 | 0.63220988 | 2.96E-10 | 1.39E-08 |
| ENSMUST00000129935.7,Rpl10a-002,nonsense_mediated_decay,ENSMUSG00000037805.14,protein_coding | 7167.869252 | 0.43200894 | 2.99E-10 | 1.40E-08 |
| ENSMUST00000183247.1,Sec31a-010,protein_coding,ENSMUSG00000035325.16,protein_coding | 406.2403289 | 0.807289801 | 3.03E-10 | 1.41E-08 |
| ENSMUST00000034746.9,Mns1-001,protein_coding,ENSMUSG00000032221.14,protein_coding | 916.2318163 | -0.590267859 | 3.03E-10 | 1.41E-08 |
| ENSMUST00000152672.1,Tulp4-005,processed_transcript,ENSMUSG00000034377.10,protein_coding | 1207.657365 | -0.394135192 | 3.12E-10 | 1.45E-08 |
| ENSMUST00000212494.1,Rpl18a-008,protein_coding,ENSMUSG00000045128.9,protein_coding | 31873.55598 | 0.263512183 | 3.13E-10 | 1.46E-08 |
| ENSMUST00000142395.7,Whsc1l1-005,protein_coding,ENSMUSG00000054823.16,protein_coding | 5044.678643 | -0.391050114 | 3.18E-10 | 1.48E-08 |
| ENSMUST00000151936.1,Gas2l1-007,processed_transcript,ENSMUSG00000034201.13,protein_coding | 778.8773447 | -0.444756012 | 3.20E-10 | 1.49E-08 |
| ENSMUST00000025718.8,Ankrd1-201,protein_coding,ENSMUSG00000024803.8,protein_coding | 1947.870078 | -0.387991037 | 3.23E-10 | 1.50E-08 |
| ENSMUST00000040005.12,Alkbh3-201,protein_coding,ENSMUSG00000040174.14,protein_coding | 4088.278017 | 0.330203591 | 3.26E-10 | 1.51E-08 |
| ENSMUST00000117676.7,Rgs17-001,protein_coding,ENSMUSG00000019775.17,protein_coding | 682.5148379 | 0.502007559 | 3.26E-10 | 1.51E-08 |
| ENSMUST00000107213.7,Nbr1-003,protein_coding,ENSMUSG00000017119.19,protein_coding | 8618.857196 | 0.302577267 | 3.26E-10 | 1.51E-08 |
| ENSMUST00000050687.12,Jdp2-201,protein_coding,ENSMUSG00000034271.15,protein_coding | 10209.81383 | 0.355518695 | 3.30E-10 | 1.52E-08 |
| ENSMUST00000210369.1,Ssbp4-002,protein_coding,ENSMUSG00000070003.7,protein_coding | 2858.180672 | -0.32795798 | 3.30E-10 | 1.52E-08 |
| ENSMUST00000103061.1,Amz2-002,protein_coding,ENSMUSG00000020610.16,protein_coding | 3575.781316 | 0.357029861 | 3.33E-10 | 1.53E-08 |
| ENSMUST00000138843.7,Slc11a2-002,protein_coding,ENSMUSG00000023030.16,protein_coding | 3797.808677 | -0.381797022 | 3.33E-10 | 1.53E-08 |
| ENSMUST00000110186.8,Smox-008,protein_coding,ENSMUSG00000027333.18,protein_coding | 508.2902793 | 0.746342959 | 3.40E-10 | 1.56E-08 |
| ENSMUST00000033049.8,Cox6a2-001,protein_coding,ENSMUSG00000030785.8,protein_coding | 165.0057165 | 0.898988688 | 3.48E-10 | 1.60E-08 |
| ENSMUST00000174834.1,Abcf1-002,retained_intron,ENSMUSG00000038762.15,protein_coding | 346.8459326 | 0.559569475 | 3.60E-10 | 1.65E-08 |
| ENSMUST00000099858.3,Prep-001,protein_coding,ENSMUSG00000019849.11,protein_coding | 29603.9173 | 0.290223565 | 3.71E-10 | 1.70E-08 |
| ENSMUST00000035648.4,Atg12-201,protein_coding,ENSMUSG00000032905.4,protein_coding | 4003.167231 | 0.317251165 | 3.75E-10 | 1.72E-08 |
| ENSMUST00000129093.1,Myl12a-007,retained_intron,ENSMUSG00000024048.14,protein_coding | 1689.865709 | 0.694265235 | 3.78E-10 | 1.73E-08 |
| ENSMUST00000201637.1,Ccnd2-004,protein_coding,ENSMUSG00000000184.12,protein_coding | 1583.417329 | 0.564676792 | 3.83E-10 | 1.75E-08 |
| ENSMUST00000030724.8,Sesn2-001,protein_coding,ENSMUSG00000028893.8,protein_coding | 11099.145 | 0.477778874 | 3.91E-10 | 1.79E-08 |
| ENSMUST00000092498.11,Sgpl1-001,protein_coding,ENSMUSG00000020097.14,protein_coding | 6257.970361 | 0.344352264 | 3.97E-10 | 1.81E-08 |
| ENSMUST00000107691.7,Trim2-003,protein_coding,ENSMUSG00000027993.16,protein_coding | 1426.44337 | -0.568256055 | 4.00E-10 | 1.82E-08 |
| ENSMUST00000153590.1,Rpl5-002,protein_coding,ENSMUSG00000058558.12,protein_coding | 26286.15797 | 0.179248497 | 4.03E-10 | 1.83E-08 |
| ENSMUST00000042390.4,Man1b1-001,protein_coding,ENSMUSG00000036646.13,protein_coding | 14522.52837 | 0.224180172 | 4.03E-10 | 1.83E-08 |
| ENSMUST00000074267.3,Rps7-201,protein_coding,ENSMUSG00000061477.3,protein_coding | 99933.78058 | 0.318601418 | 4.08E-10 | 1.85E-08 |
| ENSMUST00000109940.1,Hmgxb4-003,protein_coding,ENSMUSG00000034518.14,protein_coding | 125.6158772 | -0.907659555 | 4.13E-10 | 1.87E-08 |
| ENSMUST00000030030.14,Tex10-003,protein_coding,ENSMUSG00000028345.15,protein_coding | 9793.073359 | 0.295543983 | 4.37E-10 | 1.98E-08 |
| ENSMUST00000189529.6,Farsb-004,nonsense_mediated_decay,ENSMUSG00000026245.16,protein_coding | 1601.489993 | 0.620684754 | 4.38E-10 | 1.98E-08 |
| ENSMUST00000023612.14,Ets2-001,protein_coding,ENSMUSG00000022895.14,protein_coding | 9253.078343 | 0.484748527 | 4.38E-10 | 1.98E-08 |
| ENSMUST00000030202.13,Glipr2-001,protein_coding,ENSMUSG00000028480.14,protein_coding | 4556.157344 | 0.338159109 | 4.43E-10 | 2.00E-08 |
| ENSMUST00000118487.7,Fam234a-002,protein_coding,ENSMUSG00000024187.14,protein_coding | 4742.132883 | 0.301807631 | 4.47E-10 | 2.02E-08 |
| ENSMUST00000169744.7,Adipor2-002,protein_coding,ENSMUSG00000030168.13,protein_coding | 8537.771301 | 0.248758509 | 4.52E-10 | 2.04E-08 |
| ENSMUST00000048139.11,Wrap53-001,protein_coding,ENSMUSG00000041346.11,protein_coding | 2296.903643 | -0.341876322 | 4.69E-10 | 2.11E-08 |
| ENSMUST00000018113.7,Ptgis-001,protein_coding,ENSMUSG00000017969.13,protein_coding | 6187.48765 | -0.390777578 | 4.97E-10 | 2.23E-08 |
| ENSMUST00000167818.7,Becn1-017,nonsense_mediated_decay,ENSMUSG00000035086.13,protein_coding | 940.0441914 | -0.405241304 | 5.06E-10 | 2.27E-08 |
| ENSMUST00000043098.8,Gadd45a-001,protein_coding,ENSMUSG00000036390.8,protein_coding | 4842.187725 | 0.886010542 | 5.09E-10 | 2.28E-08 |
| ENSMUST00000034796.13,Elovl4-001,protein_coding,ENSMUSG00000032262.13,protein_coding | 703.7560728 | -0.498196808 | 5.09E-10 | 2.28E-08 |
| ENSMUST00000041369.6,Socs5-201,protein_coding,ENSMUSG00000037104.8,protein_coding | 8111.351174 | -0.348529188 | 5.27E-10 | 2.36E-08 |
| ENSMUST00000048249.6,Ndufa7-001,protein_coding,ENSMUSG00000041881.12,protein_coding | 12542.81114 | 0.275402801 | 5.35E-10 | 2.39E-08 |
| ENSMUST00000000193.5,Ccl2-001,protein_coding,ENSMUSG00000035385.5,protein_coding | 283.8463506 | -0.626328063 | 5.37E-10 | 2.40E-08 |
| ENSMUST00000103040.10,Nat9-001,protein_coding,ENSMUSG00000015542.17,protein_coding | 765.106276 | 0.461136024 | 5.37E-10 | 2.40E-08 |
| ENSMUST00000058905.7,Pars2-001,protein_coding,ENSMUSG00000043572.7,protein_coding | 912.0846315 | 0.396892982 | 5.40E-10 | 2.41E-08 |
| ENSMUST00000067857.10,Aifm2-002,protein_coding,ENSMUSG00000020085.15,protein_coding | 623.0439674 | 0.505365149 | 5.53E-10 | 2.46E-08 |
| ENSMUST00000000769.13,Serpinf1-001,protein_coding,ENSMUSG00000000753.15,protein_coding | 2089.733938 | 0.562219686 | 5.55E-10 | 2.47E-08 |
| ENSMUST00000021676.11,0610007P14Rik-001,protein_coding,ENSMUSG00000021252.11,protein_coding | 4846.92833 | 0.355913027 | 5.63E-10 | 2.50E-08 |
| ENSMUST00000024575.6,Rps6ka2-201,protein_coding,ENSMUSG00000023809.9,protein_coding | 12915.70745 | 0.406896001 | 5.67E-10 | 2.52E-08 |
| ENSMUST00000047864.9,Eef2-201,protein_coding,ENSMUSG00000034994.9,protein_coding | 663555.5375 | 0.265684762 | 5.72E-10 | 2.54E-08 |
| ENSMUST00000117757.8,Gapdh-003,protein_coding,ENSMUSG00000057666.18,protein_coding | 313454.4345 | 0.32659424 | 5.92E-10 | 2.63E-08 |
| ENSMUST00000073350.12,Ctse-001,protein_coding,ENSMUSG00000004552.16,protein_coding | 150.0711143 | 0.830487578 | 5.98E-10 | 2.64E-08 |
| ENSMUST00000160870.7,Apbb2-004,protein_coding,ENSMUSG00000029207.16,protein_coding | 6593.172962 | -0.272007603 | 5.98E-10 | 2.64E-08 |
| ENSMUST00000025483.9,Nars-201,protein_coding,ENSMUSG00000024587.9,protein_coding | 76049.87231 | 0.333168782 | 5.98E-10 | 2.64E-08 |
| ENSMUST00000119865.7,Mtfr1-003,protein_coding,ENSMUSG00000027601.13,protein_coding | 3478.937854 | 0.331870572 | 6.04E-10 | 2.66E-08 |
| ENSMUST00000075510.11,Safb2-001,protein_coding,ENSMUSG00000042625.16,protein_coding | 5248.343626 | 0.21189443 | 6.08E-10 | 2.68E-08 |
| ENSMUST00000027975.7,Phyh-001,protein_coding,ENSMUSG00000026664.7,protein_coding | 1181.111317 | 0.331462715 | 6.11E-10 | 2.69E-08 |
| ENSMUST00000060501.4,Cdkn2a-001,protein_coding,ENSMUSG00000044303.6,protein_coding | 13029.48301 | 0.262709634 | 6.26E-10 | 2.75E-08 |
| ENSMUST00000056484.9,Nlgn2-002,protein_coding,ENSMUSG00000051790.15,protein_coding | 1237.528226 | -0.683362114 | 6.29E-10 | 2.77E-08 |
| ENSMUST00000028045.3,Mrc1-001,protein_coding,ENSMUSG00000026712.3,protein_coding | 56.22569216 | -1.045715823 | 6.31E-10 | 2.77E-08 |
| ENSMUST00000041500.7,Srxn1-001,protein_coding,ENSMUSG00000032802.8,protein_coding | 9626.506551 | -0.258439942 | 6.32E-10 | 2.77E-08 |
| ENSMUST00000172435.7,Fndc4-001,protein_coding,ENSMUSG00000038552.14,protein_coding | 1245.081154 | -0.368698154 | 6.42E-10 | 2.81E-08 |
| ENSMUST00000048545.9,Arglu1-001,protein_coding,ENSMUSG00000040459.11,protein_coding | 1411.807862 | -0.378434607 | 6.43E-10 | 2.82E-08 |
| ENSMUST00000006828.8,Aplp1-001,protein_coding,ENSMUSG00000006651.8,protein_coding | 1053.228621 | -0.472695349 | 6.57E-10 | 2.87E-08 |
| ENSMUST00000034834.15,Pkm-001,protein_coding,ENSMUSG00000032294.17,protein_coding | 386716.5398 | 0.277649804 | 6.63E-10 | 2.90E-08 |
| ENSMUST00000148810.7,Fam129a-001,protein_coding,ENSMUSG00000026483.13,protein_coding | 26893.74898 | 0.410402846 | 6.72E-10 | 2.93E-08 |
| ENSMUST00000207505.1,Gm19410-001,protein_coding,ENSMUSG00000109372.1,protein_coding | 361.2544739 | 0.997766798 | 6.73E-10 | 2.94E-08 |
| ENSMUST00000005810.8,Mthfd2-001,protein_coding,ENSMUSG00000005667.8,protein_coding | 54773.82825 | 0.331555938 | 6.77E-10 | 2.95E-08 |
| ENSMUST00000177053.7,Evi5l-005,protein_coding,ENSMUSG00000011832.15,protein_coding | 854.4927281 | 0.515060749 | 6.82E-10 | 2.97E-08 |
| ENSMUST00000033965.13,Tenm3-001,protein_coding,ENSMUSG00000031561.16,protein_coding | 2404.277806 | -0.314983227 | 6.83E-10 | 2.97E-08 |
| ENSMUST00000021339.7,Dtd2-001,protein_coding,ENSMUSG00000020956.14,protein_coding | 1131.515096 | 0.321726767 | 7.01E-10 | 3.04E-08 |
| ENSMUST00000044415.15,Npepl1-001,protein_coding,ENSMUSG00000039263.16,protein_coding | 8299.91609 | 0.295215304 | 7.08E-10 | 3.07E-08 |
| ENSMUST00000038739.4,Rfwd3-001,protein_coding,ENSMUSG00000033596.5,protein_coding | 15873.12407 | -0.274677929 | 7.10E-10 | 3.07E-08 |
| ENSMUST00000164259.7,Os9-202,protein_coding,ENSMUSG00000040462.12,protein_coding | 2748.969377 | 0.511192158 | 7.09E-10 | 3.07E-08 |
| ENSMUST00000202899.3,Gsn-007,protein_coding,ENSMUSG00000026879.14,protein_coding | 2020.03639 | 0.532134945 | 7.19E-10 | 3.11E-08 |
| ENSMUST00000060710.7,Cdc25c-201,protein_coding,ENSMUSG00000044201.9,protein_coding | 3724.736687 | -0.250547608 | 7.27E-10 | 3.14E-08 |
| ENSMUST00000030872.11,Orc5-001,protein_coding,ENSMUSG00000029012.11,protein_coding | 3526.266078 | 0.281045933 | 7.30E-10 | 3.15E-08 |
| ENSMUST00000188482.7,Myc-201,protein_coding,ENSMUSG00000022346.15,protein_coding | 3478.12448 | 0.445280587 | 7.70E-10 | 3.32E-08 |
| ENSMUST00000084250.10,Rcc1-001,protein_coding,ENSMUSG00000028896.13,protein_coding | 14958.62923 | 0.281173012 | 7.74E-10 | 3.33E-08 |
| ENSMUST00000027422.6,Slc16a14-001,protein_coding,ENSMUSG00000026220.6,protein_coding | 148.096828 | 0.868825673 | 7.79E-10 | 3.36E-08 |
| ENSMUST00000059287.13,Ephb2-002,protein_coding,ENSMUSG00000028664.14,protein_coding | 1222.087488 | -0.360626687 | 7.81E-10 | 3.36E-08 |
| ENSMUST00000059775.14,Tgif1-001,protein_coding,ENSMUSG00000047407.17,protein_coding | 4503.538523 | 0.368095367 | 7.82E-10 | 3.36E-08 |
| ENSMUST00000031091.12,D5Ertd579e-001,protein_coding,ENSMUSG00000029190.19,protein_coding | 5661.926719 | -0.318536178 | 7.87E-10 | 3.38E-08 |
| ENSMUST00000210967.1,Rpl13a-009,nonsense_mediated_decay,ENSMUSG00000074129.13,protein_coding | 13189.43612 | 0.3543365 | 7.87E-10 | 3.38E-08 |
| ENSMUST00000042235.14,Eef1a1-001,protein_coding,ENSMUSG00000037742.14,protein_coding | 966374.5731 | 0.288975104 | 7.88E-10 | 3.38E-08 |
| ENSMUST00000088898.10,Zfp280c-004,protein_coding,ENSMUSG00000036916.13,protein_coding | 819.5731765 | 1.05136345 | 7.92E-10 | 3.39E-08 |
| ENSMUST00000089651.5,Dok1-001,protein_coding,ENSMUSG00000068335.6,protein_coding | 7490.986644 | 0.257319859 | 8.01E-10 | 3.42E-08 |
| ENSMUST00000212468.1,Hsbp1-003,protein_coding,ENSMUSG00000031839.7,protein_coding | 1041.997265 | 0.590502093 | 8.01E-10 | 3.42E-08 |
| ENSMUST00000108530.1,Nup88-002,protein_coding,ENSMUSG00000040667.11,protein_coding | 16130.70911 | 0.264494991 | 8.06E-10 | 3.44E-08 |
| ENSMUST00000043839.4,F11r-001,protein_coding,ENSMUSG00000038235.4,protein_coding | 14914.42976 | 0.206042991 | 8.19E-10 | 3.49E-08 |
| ENSMUST00000027760.13,Rgl1-001,protein_coding,ENSMUSG00000026482.13,protein_coding | 762.7543247 | -0.474701392 | 8.26E-10 | 3.52E-08 |

| | | | | |
|---|---|---|---|---|
| ENSMUST00000085425.4,Isg15-201,protein_coding,ENSMUSG00000035692.6,protein_coding | 1729.602595 | 0.420592853 | 8.29E-10 | 3.53E-08 |
| ENSMUST00000131853.1,Rmnd1-004,protein_coding,ENSMUSG00000019763.11,protein_coding | 35.92125577 | 1.111935598 | 8.38E-10 | 3.56E-08 |
| ENSMUST00000085635.5,Mfsd7b-001,protein_coding,ENSMUSG00000066595.9,protein_coding | 1556.246406 | -0.530123735 | 8.49E-10 | 3.60E-08 |
| ENSMUST00000022634.8,Bnip3l-002,protein_coding,ENSMUSG00000022051.15,protein_coding | 11304.15775 | 0.217751774 | 8.54E-10 | 3.62E-08 |
| ENSMUST00000171368.7,Anp32e-004,protein_coding,ENSMUSG00000015749.12,protein_coding | 727.2269307 | 0.91368485 | 8.60E-10 | 3.64E-08 |
| ENSMUST00000020562.4,Cct4-004,protein_coding,ENSMUSG00000007739.10,protein_coding | 23280.25302 | 0.233695837 | 8.64E-10 | 3.66E-08 |
| ENSMUST00000044532.10,Dock3-001,protein_coding,ENSMUSG00000039716.12,protein_coding | 155.9876345 | -0.9517304 | 8.74E-10 | 3.69E-08 |
| ENSMUST00000174757.7,Ddx39b-002,protein_coding,ENSMUSG00000019432.15,protein_coding | 284.359443 | 0.769267027 | 8.76E-10 | 3.70E-08 |
| ENSMUST00000112384.9,Rps24-201,protein_coding,ENSMUSG00000025290.16,protein_coding | 52077.97274 | 0.20994749 | 8.83E-10 | 3.73E-08 |
| ENSMUST00000201700.3,Sept11-002,protein_coding,ENSMUSG00000058013.11,protein_coding | 1160.029808 | -0.363729276 | 9.31E-10 | 3.92E-08 |
| ENSMUST00000034220.7,Herpud1-002,protein_coding,ENSMUSG00000031770.16,protein_coding | 1583.183852 | 0.700945249 | 9.32E-10 | 3.92E-08 |
| ENSMUST00000001806.9,Coasy-001,protein_coding,ENSMUSG00000001755.12,protein_coding | 2505.657906 | 0.465746134 | 1.01E-09 | 4.23E-08 |
| ENSMUST00000090180.2,Sema3g-001,protein_coding,ENSMUSG00000021904.5,protein_coding | 656.7096495 | 0.554350204 | 1.02E-09 | 4.27E-08 |
| ENSMUST00000029771.12,F3-001,protein_coding,ENSMUSG00000028128.13,protein_coding | 3351.355974 | 0.311147988 | 1.02E-09 | 4.29E-08 |
| ENSMUST00000191837.5,Pcdh7-001,protein_coding,ENSMUSG00000029108.14,protein_coding | 2424.203826 | -0.708149505 | 1.02E-09 | 4.29E-08 |
| ENSMUST00000026613.13,Gusb-001,protein_coding,ENSMUSG00000025534.17,protein_coding | 8413.596414 | -0.252477969 | 1.03E-09 | 4.30E-08 |
| ENSMUST00000029540.12,Npr1-001,protein_coding,ENSMUSG00000027931.12,protein_coding | 521.6545658 | -0.471706008 | 1.03E-09 | 4.32E-08 |
| ENSMUST00000000579.2,Sox9-001,protein_coding,ENSMUSG00000000567.5,protein_coding | 5431.018206 | -0.247454339 | 1.04E-09 | 4.35E-08 |
| ENSMUST00000015934.12,Surf1-001,protein_coding,ENSMUSG00000015790.16,protein_coding | 2927.30723 | 0.35858943 | 1.04E-09 | 4.36E-08 |
| ENSMUST00000026267.15,Parp6-001,protein_coding,ENSMUSG00000025237.16,protein_coding | 387.3579916 | 0.949633661 | 1.05E-09 | 4.38E-08 |
| ENSMUST00000112101.7,Itga6-002,protein_coding,ENSMUSG00000027111.15,protein_coding | 1158.220212 | 0.960195931 | 1.05E-09 | 4.39E-08 |
| ENSMUST00000003493.8,Prkcsh-002,protein_coding,ENSMUSG00000003402.14,protein_coding | 10904.85612 | 0.220710497 | 1.08E-09 | 4.49E-08 |
| ENSMUST00000167327.1,Actn1-202,protein_coding,ENSMUSG00000015143.14,protein_coding | 5900.188029 | 0.251976275 | 1.09E-09 | 4.53E-08 |
| ENSMUST00000029476.8,Sec22b-001,protein_coding,ENSMUSG00000027879.9,protein_coding | 2562.286516 | -0.306191598 | 1.13E-09 | 4.68E-08 |
| ENSMUST00000095576.3,Scel-201,protein_coding,ENSMUSG00000022123.8,protein_coding | 910.8065006 | -0.603322997 | 1.13E-09 | 4.68E-08 |
| ENSMUST00000082122.13,Dut-001,protein_coding,ENSMUSG00000027203.15,protein_coding | 16079.99902 | 0.264994134 | 1.14E-09 | 4.73E-08 |
| ENSMUST00000106113.1,Foxk2-001,protein_coding,ENSMUSG00000039275.9,protein_coding | 20784.39329 | 0.208745121 | 1.17E-09 | 4.84E-08 |
| ENSMUST00000100757.5,Adam1a-001,protein_coding,ENSMUSG00000072647.5,protein_coding | 123.2991623 | -0.812003566 | 1.18E-09 | 4.87E-08 |
| ENSMUST00000165740.8,Tecr-001,protein_coding,ENSMUSG00000031708.16,protein_coding | 3258.733864 | -0.378278199 | 1.21E-09 | 5.00E-08 |
| ENSMUST00000080394.12,Sh3kbp1-002,protein_coding,ENSMUSG00000040990.17,protein_coding | 2660.735505 | 0.358886536 | 1.23E-09 | 5.07E-08 |
| ENSMUST00000123104.1,Knstrn-006,nonsense_mediated_decay,ENSMUSG00000027331.15,protein_coding | 192.9883321 | 0.646053701 | 1.24E-09 | 5.13E-08 |
| ENSMUST00000021135.4,Ncbp3-001,protein_coding,ENSMUSG00000020783.14,protein_coding | 6448.094306 | 0.225389783 | 1.24E-09 | 5.13E-08 |
| ENSMUST00000156036.7,Pdk1-002,processed_transcript,ENSMUSG00000006494.11,protein_coding | 7017.581156 | 0.390537255 | 1.25E-09 | 5.13E-08 |
| ENSMUST00000102823.9,Ttc39b-001,protein_coding,ENSMUSG00000038172.14,protein_coding | 7395.068955 | -0.319619161 | 1.25E-09 | 5.15E-08 |
| ENSMUST00000032924.5,Kctd13-001,protein_coding,ENSMUSG00000030685.5,protein_coding | 2948.379709 | 0.424276859 | 1.29E-09 | 5.29E-08 |
| ENSMUST00000198241.4,Asic3-003,retained_intron,ENSMUSG00000038276.12,protein_coding | 92.99889187 | -0.95436828 | 1.31E-09 | 5.37E-08 |
| ENSMUST00000021166.5,Cygb-001,protein_coding,ENSMUSG00000020810.5,protein_coding | 122.5397899 | -0.880201966 | 1.31E-09 | 5.37E-08 |
| ENSMUST00000113590.7,Speg-002,protein_coding,ENSMUSG00000026207.16,protein_coding | 1192.196271 | -0.380657319 | 1.31E-09 | 5.39E-08 |
| ENSMUST00000103122.9,Tgm2-001,protein_coding,ENSMUSG00000037820.15,protein_coding | 668.4835954 | 0.434340445 | 1.37E-09 | 5.61E-08 |
| ENSMUST00000004554.13,Rps5-001,protein_coding,ENSMUSG00000012848.15,protein_coding | 86319.97218 | 0.280743159 | 1.39E-09 | 5.70E-08 |
| ENSMUST00000173070.7,1110038B12Rik-002,lincRNA,ENSMUSG00000092203.7,processed_transcript | 1471.56467 | 0.431139611 | 1.43E-09 | 5.86E-08 |
| ENSMUST00000201022.1,Psph-008,retained_intron,ENSMUSG00000029446.14,protein_coding | 1601.870836 | 0.495575051 | 1.44E-09 | 5.89E-08 |
| ENSMUST00000212520.1,Capn15-001,protein_coding,ENSMUSG00000037326.10,protein_coding | 1270.4916 | 0.419025171 | 1.44E-09 | 5.90E-08 |
| ENSMUST00000106602.9,Rpl38-001,protein_coding,ENSMUSG00000057322.12,protein_coding | 26142.32481 | 0.206086419 | 1.47E-09 | 6.02E-08 |
| ENSMUST00000033770.12,Mecp2-002,protein_coding,ENSMUSG00000031393.16,protein_coding | 1891.308471 | 0.384691993 | 1.49E-09 | 6.08E-08 |
| ENSMUST00000106653.3,Usp47-004,protein_coding,ENSMUSG00000059263.9,protein_coding | 8050.37801 | -0.449004996 | 1.50E-09 | 6.11E-08 |
| ENSMUST00000117399.1,Msn-001,protein_coding,ENSMUSG00000031207.16,protein_coding | 68509.51967 | 0.215486537 | 1.51E-09 | 6.16E-08 |
| ENSMUST00000064174.11,Cemip-001,protein_coding,ENSMUSG00000052353.13,protein_coding | 490.9065138 | -0.65591677 | 1.53E-09 | 6.21E-08 |
| ENSMUST00000022993.5,Derl1-201,protein_coding,ENSMUSG00000022365.5,protein_coding | 23353.42058 | 0.327834173 | 1.55E-09 | 6.30E-08 |
| ENSMUST00000114071.7,Capg-001,protein_coding,ENSMUSG00000056737.14,protein_coding | 771.0171209 | 0.817262466 | 1.56E-09 | 6.35E-08 |
| ENSMUST00000032998.12,Rps3-001,protein_coding,ENSMUSG00000030744.13,protein_coding | 98965.0303 | 0.273020426 | 1.57E-09 | 6.35E-08 |
| ENSMUST00000151958.1,Gadd45a-004,retained_intron,ENSMUSG00000036390.8,protein_coding | 729.0544093 | 0.872768104 | 1.61E-09 | 6.53E-08 |
| ENSMUST00000153258.7,Dpm1-008,retained_intron,ENSMUSG00000078919.10,protein_coding | 44.20800525 | 1.106297597 | 1.62E-09 | 6.54E-08 |
| ENSMUST00000156359.1,Rplp0-004,protein_coding,ENSMUSG00000067274.10,protein_coding | 46865.76366 | 0.452574951 | 1.63E-09 | 6.59E-08 |
| ENSMUST00000090553.11,Sars-001,protein_coding,ENSMUSG00000068739.13,protein_coding | 11057.16233 | 0.387569165 | 1.66E-09 | 6.71E-08 |
| ENSMUST00000046831.10,Tmem205-001,protein_coding,ENSMUSG00000040883.16,protein_coding | 687.1504096 | 0.469638308 | 1.68E-09 | 6.76E-08 |
| ENSMUST00000041213.11,Cnppd1-002,protein_coding,ENSMUSG00000033159.14,protein_coding | 2057.218809 | 0.394526039 | 1.69E-09 | 6.80E-08 |
| ENSMUST00000105349.7,Mbd3-002,protein_coding,ENSMUSG00000035478.14,protein_coding | 10945.33641 | -0.342970627 | 1.71E-09 | 6.89E-08 |
| ENSMUST00000107333.7,S100a16-006,protein_coding,ENSMUSG00000074457.10,protein_coding | 1063.013927 | 0.553878658 | 1.72E-09 | 6.91E-08 |
| ENSMUST00000049822.9,Them4-001,protein_coding,ENSMUSG00000028145.9,protein_coding | 1068.233538 | 0.524797836 | 1.72E-09 | 6.93E-08 |
| ENSMUST00000045970.7,Gpc1-001,protein_coding,ENSMUSG00000034220.7,protein_coding | 33463.93889 | 0.246809533 | 1.73E-09 | 6.94E-08 |
| ENSMUST00000030010.3,Abca1-001,protein_coding,ENSMUSG00000015243.4,protein_coding | 238.205176 | -0.666912516 | 1.79E-09 | 7.17E-08 |
| ENSMUST00000019854.12,Mrpl24-001,protein_coding,ENSMUSG00000019710.13,protein_coding | 1785.105046 | 0.467319595 | 1.80E-09 | 7.21E-08 |
| ENSMUST00000176500.7,Arhgef2-003,protein_coding,ENSMUSG00000028059.15,protein_coding | 22669.21372 | 0.616128717 | 1.80E-09 | 7.21E-08 |
| ENSMUST00000113867.8,Tsc1-007,protein_coding,ENSMUSG00000026812.16,protein_coding | 1768.206811 | -0.475416927 | 1.81E-09 | 7.25E-08 |
| ENSMUST00000041055.7,Atg2b-201,protein_coding,ENSMUSG00000041341.9,protein_coding | 11571.70566 | 0.209377855 | 1.82E-09 | 7.29E-08 |
| ENSMUST00000107812.7,Ift80-001,protein_coding,ENSMUSG00000027778.15,protein_coding | 2333.008049 | -0.538958275 | 1.85E-09 | 7.38E-08 |
| ENSMUST00000116629.8,Arid5a-005,nonsense_mediated_decay,ENSMUSG00000037447.16,protein_coding | 2168.204183 | 0.57397759 | 1.85E-09 | 7.38E-08 |
| ENSMUST00000017620.9,Cant1-002,protein_coding,ENSMUSG00000025575.14,protein_coding | 3105.798566 | -0.207719919 | 1.87E-09 | 7.44E-08 |
| ENSMUST00000025350.9,Dcp2-001,protein_coding,ENSMUSG00000024472.9,protein_coding | 5528.981018 | -0.566034889 | 1.90E-09 | 7.57E-08 |
| ENSMUST00000049357.9,Pnrc1-001,protein_coding,ENSMUSG00000040128.9,protein_coding | 3385.351426 | 0.301228876 | 1.94E-09 | 7.72E-08 |
| ENSMUST00000099475.11,Serf2-004,protein_coding,ENSMUSG00000074884.12,protein_coding | 10399.44923 | 0.234196325 | 1.95E-09 | 7.75E-08 |
| ENSMUST00000027727.14,Adipor1-001,protein_coding,ENSMUSG00000026457.14,protein_coding | 23522.29454 | 0.241003036 | 2.00E-09 | 7.93E-08 |
| ENSMUST00000041029.5,Smad6-001,protein_coding,ENSMUSG00000036867.7,protein_coding | 6149.324162 | -0.34317003 | 2.00E-09 | 7.95E-08 |
| ENSMUST00000138160.7,Bnip3l-007,processed_transcript,ENSMUSG00000022051.15,protein_coding | 2007.968539 | 0.423394498 | 2.01E-09 | 7.95E-08 |
| ENSMUST00000151839.1,Agfg2-003,protein_coding,ENSMUSG00000029722.15,protein_coding | 226.2026108 | 0.756587362 | 2.01E-09 | 7.96E-08 |
| ENSMUST00000047511.13,Shtn1-201,protein_coding,ENSMUSG00000041362.16,protein_coding | 696.2225288 | 0.404479447 | 2.02E-09 | 8.01E-08 |
| ENSMUST00000125036.7,Shc1-005,protein_coding,ENSMUSG00000042626.13,protein_coding | 161.7515223 | 0.882035271 | 2.03E-09 | 8.02E-08 |
| ENSMUST00000109816.7,Tpx2-002,protein_coding,ENSMUSG00000027469.16,protein_coding | 21656.05901 | 0.242416771 | 2.04E-09 | 8.04E-08 |
| ENSMUST00000027985.7,Ddr2-201,protein_coding,ENSMUSG00000026674.9,protein_coding | 1486.788847 | 0.834651182 | 2.08E-09 | 8.20E-08 |
| ENSMUST00000022704.7,Itm2b-201,protein_coding,ENSMUSG00000022108.7,protein_coding | 13206.15492 | 0.221234502 | 2.10E-09 | 8.29E-08 |
| ENSMUST00000196213.1,Galnt11-005,processed_transcript,ENSMUSG00000038072.14,protein_coding | 679.7342667 | -0.542429742 | 2.12E-09 | 8.35E-08 |
| ENSMUST00000115457.7,Eif4g1-003,protein_coding,ENSMUSG00000045983.15,protein_coding | 5287.222823 | -0.791818444 | 2.13E-09 | 8.40E-08 |
| ENSMUST00000040656.6,Bphl-201,protein_coding,ENSMUSG00000038286.10,protein_coding | 1279.34219 | 0.3350752 | 2.16E-09 | 8.46E-08 |
| ENSMUST00000082243.6,Spire1-202,protein_coding,ENSMUSG00000024533.15,protein_coding | 3522.328497 | 0.35362443 | 2.15E-09 | 8.46E-08 |
| ENSMUST00000029815.7,Cisd2-001,protein_coding,ENSMUSG00000028165.9,protein_coding | 5259.955586 | -0.238306585 | 2.19E-09 | 8.61E-08 |
| ENSMUST00000108275.1,Ift20-002,protein_coding,ENSMUSG00000001105.15,protein_coding | 580.1090949 | 0.485173781 | 2.20E-09 | 8.64E-08 |
| ENSMUST00000057893.6,Fzd2-001,protein_coding,ENSMUSG00000050288.6,protein_coding | 5743.264492 | -0.251079148 | 2.22E-09 | 8.69E-08 |
| ENSMUST00000031331.12,Ccng2-001,protein_coding,ENSMUSG00000029385.14,protein_coding | 1699.591985 | 0.577335946 | 2.22E-09 | 8.69E-08 |
| ENSMUST00000019939.11,Snx3-001,protein_coding,ENSMUSG00000019804.12,protein_coding | 18281.29067 | 0.161730703 | 2.25E-09 | 8.81E-08 |
| ENSMUST00000095224.10,Safb-001,protein_coding,ENSMUSG00000071054.12,protein_coding | 9025.188201 | 0.265027354 | 2.27E-09 | 8.88E-08 |
| ENSMUST00000075774.3,Tubb2b-201,protein_coding,ENSMUSG00000045136.5,protein_coding | 3643.0933 | -0.380577573 | 2.29E-09 | 8.94E-08 |

| | | | | |
|---|---|---|---|---|
| ENSMUST00000020497.13,Aldh1l2-001,protein_coding,ENSMUSG00000020256.14,protein_coding | 28986.00317 | 0.49028261 | 2.33E-09 | 9.08E-08 |
| ENSMUST00000033899.13,Col4a2-001,protein_coding,ENSMUSG00000031503.13,protein_coding | 39050.65295 | 0.183177702 | 2.34E-09 | 9.10E-08 |
| ENSMUST00000045740.7,Zxdc-004,protein_coding,ENSMUSG00000034430.16,protein_coding | 669.5405469 | 0.677246554 | 2.35E-09 | 9.13E-08 |
| ENSMUST00000019677.11,Mknk1-001,protein_coding,ENSMUSG00000028708.16,protein_coding | 2490.228845 | 0.401848136 | 2.39E-09 | 9.29E-08 |
| ENSMUST00000048781.3,Pck2-201,protein_coding,ENSMUSG00000040618.6,protein_coding | 45284.52699 | 0.413633674 | 2.43E-09 | 9.46E-08 |
| ENSMUST00000177056.7,Taf6l-005,protein_coding,ENSMUSG00000003680.15,protein_coding | 552.0807909 | -0.917325179 | 2.46E-09 | 9.55E-08 |
| ENSMUST00000199202.4,Fubp1-005,protein_coding,ENSMUSG00000028034.15,protein_coding | 1808.651653 | 0.40218027 | 2.47E-09 | 9.60E-08 |
| ENSMUST00000103031.7,Fbf1-001,protein_coding,ENSMUSG00000020776.18,protein_coding | 4718.390618 | -0.490078553 | 2.49E-09 | 9.66E-08 |
| ENSMUST00000150757.7,Mrgbp-002,retained_intron,ENSMUSG00000027569.15,protein_coding | 809.5960348 | 0.350758207 | 2.51E-09 | 9.71E-08 |
| ENSMUST00000091002.7,Fhdc1-001,protein_coding,ENSMUSG00000041842.15,protein_coding | 7462.132566 | 0.221518765 | 2.61E-09 | 1.01E-07 |
| ENSMUST00000160298.1,Gm8113-001,protein_coding,ENSMUSG00000089901.1,protein_coding | 519.1812577 | -0.517261642 | 2.61E-09 | 1.01E-07 |
| ENSMUST00000202148.1,Gm20186-002,lincRNA,ENSMUSG00000106874.1,lincRNA | 895.1771199 | -0.634594896 | 2.62E-09 | 1.01E-07 |
| ENSMUST00000020647.9,3010026O09Rik-001,protein_coding,ENSMUSG00000020381.10,protein_coding | 1060.304449 | 0.404062536 | 2.64E-09 | 1.02E-07 |
| ENSMUST00000049460.10,Grn-001,protein_coding,ENSMUSG00000034708.11,protein_coding | 11349.17789 | 0.281231639 | 2.68E-09 | 1.03E-07 |
| ENSMUST00000102678.4,H2-T23-001,protein_coding,ENSMUSG00000067212.8,protein_coding | 544.5588608 | 0.398830667 | 2.70E-09 | 1.04E-07 |
| ENSMUST00000050096.13,Add3-202,protein_coding,ENSMUSG00000025026.14,protein_coding | 6918.50706 | 0.259390651 | 2.71E-09 | 1.04E-07 |
| ENSMUST00000187771.6,Morf4l1-013,processed_transcript,ENSMUSG00000062270.13,protein_coding | 3232.524493 | 0.364683052 | 2.72E-09 | 1.04E-07 |
| ENSMUST00000075447.13,Nectin2-001,protein_coding,ENSMUSG00000062300.14,protein_coding | 2820.064248 | 0.318619292 | 2.72E-09 | 1.05E-07 |
| ENSMUST00000037930.12,Msl1-003,protein_coding,ENSMUSG00000052915.14,protein_coding | 333.9697255 | 0.584366781 | 2.75E-09 | 1.06E-07 |
| ENSMUST00000165331.7,Immt-003,protein_coding,ENSMUSG00000052337.15,protein_coding | 177.7596003 | -0.8485738 | 2.77E-09 | 1.06E-07 |
| ENSMUST00000129753.7,Aldh2-002,protein_coding,ENSMUSG00000029455.14,protein_coding | 28152.33037 | 0.260135375 | 2.78E-09 | 1.06E-07 |
| ENSMUST00000163600.7,Ccnh-003,protein_coding,ENSMUSG00000021548.10,protein_coding | 1253.993709 | 0.402207465 | 2.79E-09 | 1.07E-07 |
| ENSMUST00000126486.7,Fyn-003,protein_coding,ENSMUSG00000019843.14,protein_coding | 1157.444468 | 0.709110169 | 2.86E-09 | 1.10E-07 |
| ENSMUST00000047857.15,Fbxl5-001,protein_coding,ENSMUSG00000039753.16,protein_coding | 5380.325055 | 0.241549581 | 2.91E-09 | 1.11E-07 |
| ENSMUST00000001187.14,Anxa4-007,protein_coding,ENSMUSG00000029994.17,protein_coding | 12269.9154 | 0.210453159 | 2.90E-09 | 1.11E-07 |
| ENSMUST00000021322.6,Fmnl1-006,retained_intron,ENSMUSG00000055805.14,protein_coding | 82.58455294 | -0.982115063 | 2.90E-09 | 1.11E-07 |
| ENSMUST00000127906.7,Dsp-002,protein_coding,ENSMUSG00000054889.9,protein_coding | 1035.310449 | 0.584956297 | 2.91E-09 | 1.11E-07 |
| ENSMUST00000112969.9,Paqr3-004,protein_coding,ENSMUSG00000055725.11,protein_coding | 913.9570855 | 0.678835403 | 2.91E-09 | 1.11E-07 |
| ENSMUST00000113045.8,Mum1l1-203,protein_coding,ENSMUSG00000042515.13,protein_coding | 765.5506332 | -0.864290243 | 2.93E-09 | 1.12E-07 |
| ENSMUST00000187504.6,Lbhd1-001,protein_coding,ENSMUSG00000096740.7,protein_coding | 834.9343556 | -0.51591269 | 2.96E-09 | 1.12E-07 |
| ENSMUST00000120542.7,Mettl1-002,protein_coding,ENSMUSG00000006732.14,protein_coding | 831.8202627 | 0.635852524 | 3.00E-09 | 1.14E-07 |
| ENSMUST00000174031.7,Ube2i-010,protein_coding,ENSMUSG00000015120.15,protein_coding | 12621.32032 | 0.188357764 | 3.01E-09 | 1.14E-07 |
| ENSMUST00000007959.13,Rhoa-001,protein_coding,ENSMUSG00000007815.13,protein_coding | 40074.11746 | 0.252819669 | 3.01E-09 | 1.14E-07 |
| ENSMUST00000189664.1,Ndufs1-009,protein_coding,ENSMUSG00000025968.16,protein_coding | 61.64280811 | 0.978564693 | 3.04E-09 | 1.15E-07 |
| ENSMUST00000130256.2,B930095G15Rik-002,sense_intronic,ENSMUSG00000085133.2,sense_intronic | 847.550428 | -0.334843566 | 3.04E-09 | 1.15E-07 |
| ENSMUST00000087401.11,Morc4-001,protein_coding,ENSMUSG00000031434.15,protein_coding | 2321.273929 | 0.450712643 | 3.07E-09 | 1.16E-07 |
| ENSMUST00000034187.8,Mmp2-001,protein_coding,ENSMUSG00000031740.8,protein_coding | 2681.224241 | -0.198591983 | 3.09E-09 | 1.17E-07 |
| ENSMUST00000151843.7,Mib2-002,processed_transcript,ENSMUSG00000029060.17,protein_coding | 1080.752643 | 0.664990886 | 3.11E-09 | 1.17E-07 |
| ENSMUST00000022459.3,Phf7-201,protein_coding,ENSMUSG00000021902.6,protein_coding | 1548.024464 | 0.311041457 | 3.12E-09 | 1.18E-07 |
| ENSMUST00000109626.3,Cyld-203,protein_coding,ENSMUSG00000036712.15,protein_coding | 700.8247536 | -0.849527579 | 3.18E-09 | 1.20E-07 |
| ENSMUST00000120878.8,Sept8-007,protein_coding,ENSMUSG00000018398.18,protein_coding | 1689.108586 | 0.354249713 | 3.28E-09 | 1.24E-07 |
| ENSMUST00000165536.7,Klf3-001,protein_coding,ENSMUSG00000029178.14,protein_coding | 4699.738815 | -0.422216159 | 3.30E-09 | 1.24E-07 |
| ENSMUST00000139560.7,Ndc1-001,protein_coding,ENSMUSG00000028614.14,protein_coding | 10534.50926 | -0.2718983 | 3.34E-09 | 1.25E-07 |
| ENSMUST00000134040.1,Tpt1-005,processed_transcript,ENSMUSG00000060126.14,protein_coding | 86891.31041 | 0.200294071 | 3.34E-09 | 1.25E-07 |
| ENSMUST00000165721.7,Csnk1a1-002,protein_coding,ENSMUSG00000024576.14,protein_coding | 16546.18338 | 0.213124934 | 3.37E-09 | 1.27E-07 |
| ENSMUST00000028619.4,Hsd17b12-001,protein_coding,ENSMUSG00000027195.10,protein_coding | 19500.41033 | 0.242764361 | 3.39E-09 | 1.27E-07 |
| ENSMUST00000161309.7,Pole4-004,processed_transcript,ENSMUSG00000030042.15,protein_coding | 192.9224575 | 0.675784501 | 3.42E-09 | 1.28E-07 |
| ENSMUST00000105324.8,Hmg20b-003,protein_coding,ENSMUSG00000020232.17,protein_coding | 1820.891056 | 0.367661176 | 3.42E-09 | 1.28E-07 |
| ENSMUST00000025217.8,Hspa9-001,protein_coding,ENSMUSG00000024359.8,protein_coding | 121189.7208 | 0.263838148 | 3.42E-09 | 1.28E-07 |
| ENSMUST00000205865.1,Gpi1-010,nonsense_mediated_decay,ENSMUSG00000036427.5,protein_coding | 4126.955724 | 0.253040343 | 3.43E-09 | 1.28E-07 |
| ENSMUST00000032410.13,Tada3-001,protein_coding,ENSMUSG00000048930.12,protein_coding | 3967.861065 | 0.249254795 | 3.45E-09 | 1.29E-07 |
| ENSMUST00000031597.6,Plbd2-001,protein_coding,ENSMUSG00000029598.12,protein_coding | 6811.624156 | 0.205574239 | 3.46E-09 | 1.29E-07 |
| ENSMUST00000211699.1,Trak1-002,retained_intron,ENSMUSG00000032536.11,protein_coding | 187.5957016 | 0.967265899 | 3.47E-09 | 1.29E-07 |
| ENSMUST00000021359.5,Pole2-201,protein_coding,ENSMUSG00000020974.5,protein_coding | 2959.834651 | 0.230571851 | 3.50E-09 | 1.30E-07 |
| ENSMUST00000181283.2,Ssh2-002,protein_coding,ENSMUSG00000037926.15,protein_coding | 541.2090477 | 0.579558557 | 3.52E-09 | 1.31E-07 |
| ENSMUST00000022977.13,Sqle-001,protein_coding,ENSMUSG00000022351.14,protein_coding | 22831.06782 | 0.319178705 | 3.56E-09 | 1.32E-07 |
| ENSMUST00000143254.7,Svil-002,protein_coding,ENSMUSG00000024236.18,protein_coding | 6062.097394 | 0.269318878 | 3.56E-09 | 1.32E-07 |
| ENSMUST00000090973.11,Nes-001,protein_coding,ENSMUSG00000004891.16,protein_coding | 1637.427847 | -0.41586949 | 3.58E-09 | 1.33E-07 |
| ENSMUST00000078434.6,Ppp3cc-201,protein_coding,ENSMUSG00000022092.10,protein_coding | 1623.975357 | 0.41660527 | 3.58E-09 | 1.33E-07 |
| ENSMUST00000027738.13,Tor1aip1-001,protein_coding,ENSMUSG00000026466.16,protein_coding | 1193.854838 | 0.480293898 | 3.59E-09 | 1.33E-07 |
| ENSMUST00000174372.2,Slc2a1-004,nonsense_mediated_decay,ENSMUSG00000028645.11,protein_coding | 6555.667061 | 0.336172722 | 3.61E-09 | 1.34E-07 |
| ENSMUST00000093943.9,Cbx1-001,protein_coding,ENSMUSG00000018666.13,protein_coding | 3012.526978 | 0.360241737 | 3.62E-09 | 1.34E-07 |
| ENSMUST00000106588.7,Rps15a-004,protein_coding,ENSMUSG00000008683.16,protein_coding | 29245.59147 | 0.205380925 | 3.67E-09 | 1.36E-07 |
| ENSMUST00000023039.14,St13-004,protein_coding,ENSMUSG00000022403.14,protein_coding | 15442.08898 | 0.201305927 | 3.67E-09 | 1.36E-07 |
| ENSMUST00000035301.6,Atp6v1g1-001,protein_coding,ENSMUSG00000039105.6,protein_coding | 15235.06397 | 0.226990288 | 3.70E-09 | 1.36E-07 |
| ENSMUST00000047034.8,Ttbk1-201,protein_coding,ENSMUSG00000015599.8,protein_coding | 137.840328 | -0.784472237 | 3.70E-09 | 1.36E-07 |
| ENSMUST00000114471.1,Cblb-201,protein_coding,ENSMUSG00000022637.10,protein_coding | 5568.639423 | 0.192797739 | 3.73E-09 | 1.37E-07 |
| ENSMUST00000193168.1,Rpl22l1-006,nonsense_mediated_decay,ENSMUSG00000039221.10,protein_coding | 397.3521714 | 0.559542211 | 3.74E-09 | 1.37E-07 |
| ENSMUST00000063814.14,Gnpda1-001,protein_coding,ENSMUSG00000052102.14,protein_coding | 2961.034659 | 0.307254375 | 3.75E-09 | 1.38E-07 |
| ENSMUST00000025746.13,Dpf2-009,nonsense_mediated_decay,ENSMUSG00000024826.14,protein_coding | 7436.801419 | 0.420298164 | 3.78E-09 | 1.39E-07 |
| ENSMUST00000046093.5,Hmx3-001,protein_coding,ENSMUSG00000040148.5,protein_coding | 74.9642312 | 1.011166535 | 3.80E-09 | 1.39E-07 |
| ENSMUST00000176305.7,Hnrnpk-014,protein_coding,ENSMUSG00000021546.17,protein_coding | 12813.47555 | 0.261799387 | 3.84E-09 | 1.41E-07 |
| ENSMUST00000032386.10,Bhlhe41-001,protein_coding,ENSMUSG00000030256.11,protein_coding | 1077.274499 | 0.312427889 | 3.91E-09 | 1.43E-07 |
| ENSMUST00000090559.11,Cry2-001,protein_coding,ENSMUSG00000068742.11,protein_coding | 3079.16279 | 0.400954459 | 3.94E-09 | 1.44E-07 |
| ENSMUST00000191775.1,Cr1l-007,protein_coding,ENSMUSG00000016481.15,protein_coding | 928.3965545 | 0.544089651 | 3.98E-09 | 1.45E-07 |
| ENSMUST00000195943.1,Smarcd3-009,protein_coding,ENSMUSG00000028949.13,protein_coding | 901.7077436 | -0.485661567 | 3.98E-09 | 1.45E-07 |
| ENSMUST00000212230.1,Dync1li2-007,protein_coding,ENSMUSG00000035770.8,protein_coding | 821.760156 | 0.500353608 | 4.01E-09 | 1.46E-07 |
| ENSMUST00000087638.3,Aldh1a1-001,protein_coding,ENSMUSG00000053279.6,protein_coding | 3341.994744 | -0.359360461 | 4.03E-09 | 1.47E-07 |
| ENSMUST00000018700.12,Dlg4-003,protein_coding,ENSMUSG00000020886.16,protein_coding | 3499.827505 | 0.353057821 | 4.07E-09 | 1.48E-07 |
| ENSMUST00000165223.7,Clasp1-003,protein_coding,ENSMUSG00000064302.13,protein_coding | 8346.373489 | -0.246243467 | 4.10E-09 | 1.49E-07 |
| ENSMUST00000140164.7,Abi1-001,protein_coding,ENSMUSG00000058835.14,protein_coding | 2639.524368 | -0.720533268 | 4.19E-09 | 1.52E-07 |
| ENSMUST00000023774.11,Slc11a2-001,protein_coding,ENSMUSG00000023030.16,protein_coding | 2723.60445 | 0.365090973 | 4.21E-09 | 1.53E-07 |
| ENSMUST00000003612.12,Dusp3-001,protein_coding,ENSMUSG00000003518.13,protein_coding | 1951.639128 | -0.351773357 | 4.23E-09 | 1.53E-07 |
| ENSMUST00000000619.7,Clcn4-001,protein_coding,ENSMUSG00000000605.9,protein_coding | 2229.908187 | -0.366490659 | 4.26E-09 | 1.54E-07 |
| ENSMUST00000130818.7,Pcnp-002,nonsense_mediated_decay,ENSMUSG00000071533.11,protein_coding | 1053.384111 | 0.377129423 | 4.28E-09 | 1.55E-07 |
| ENSMUST00000216867.1,Rps25-002,protein_coding,ENSMUSG00000009927.9,protein_coding | 47837.60275 | 0.268214728 | 4.29E-09 | 1.55E-07 |
| ENSMUST00000052622.5,1810011O10Rik-001,protein_coding,ENSMUSG00000056313.5,protein_coding | 838.3621995 | 0.39337678 | 4.32E-09 | 1.56E-07 |
| ENSMUST00000021620.12,Otub2-001,protein_coding,ENSMUSG00000021203.15,protein_coding | 2298.115619 | 0.547144425 | 4.33E-09 | 1.57E-07 |
| ENSMUST00000140322.1,Ssu72-003,retained_intron,ENSMUSG00000029038.9,protein_coding | 1035.758724 | 0.386298135 | 4.39E-09 | 1.59E-07 |
| ENSMUST00000001079.14,Sec24b-001,protein_coding,ENSMUSG00000001052.15,protein_coding | 13402.05881 | 0.154675611 | 4.40E-09 | 1.59E-07 |
| ENSMUST00000103217.10,Pex14-001,protein_coding,ENSMUSG00000028975.16,protein_coding | 10907.44233 | 0.255265543 | 4.43E-09 | 1.60E-07 |
| ENSMUST00000025490.8,Prrc1-201,protein_coding,ENSMUSG00000024594.8,protein_coding | 8247.733216 | 0.307721628 | 4.47E-09 | 1.61E-07 |

| | | | | |
|---|---|---|---|---|
| ENSMUST00000069180.7,Zcchc24-001,protein_coding,ENSMUSG00000055538.7,protein_coding | 6724.20613 | 0.353688312 | 4.49E-09 | 1.62E-07 |
| ENSMUST00000165316.7,Iars-001,protein_coding,ENSMUSG00000037851.13,protein_coding | 63134.81319 | 0.283983683 | 4.70E-09 | 1.69E-07 |
| ENSMUST00000206378.1,Kcmf1-006,protein_coding,ENSMUSG00000055239.10,protein_coding | 431.8425934 | 0.904284441 | 4.76E-09 | 1.71E-07 |
| ENSMUST00000097333.9,Ddr1-002,protein_coding,ENSMUSG00000003534.17,protein_coding | 2345.479005 | -0.404088686 | 4.82E-09 | 1.73E-07 |
| ENSMUST00000123555.7,Parp3-004,protein_coding,ENSMUSG00000023249.15,protein_coding | 1191.580102 | -0.508032222 | 4.88E-09 | 1.75E-07 |
| ENSMUST00000211769.1,Glrx3-005,nonsense_mediated_decay,ENSMUSG00000031068.8,protein_coding | 142.0800334 | 0.865182936 | 4.88E-09 | 1.75E-07 |
| ENSMUST00000119848.7,Eme2-003,protein_coding,ENSMUSG00000073436.11,protein_coding | 491.9471236 | -0.556126302 | 4.89E-09 | 1.75E-07 |
| ENSMUST00000188750.6,Tfdp2-013,protein_coding,ENSMUSG00000032411.15,protein_coding | 622.8812145 | 0.611307118 | 4.97E-09 | 1.78E-07 |
| ENSMUST00000020974.6,Id2-201,protein_coding,ENSMUSG00000020644.8,protein_coding | 6053.472873 | -0.613636959 | 5.03E-09 | 1.80E-07 |
| ENSMUST00000096367.4,Spin4-001,protein_coding,ENSMUSG00000071722.4,protein_coding | 324.73146 | -0.786111588 | 5.03E-09 | 1.80E-07 |
| ENSMUST00000021297.5,Lsm12-001,protein_coding,ENSMUSG00000020922.11,protein_coding | 17958.3736 | 0.174298161 | 5.12E-09 | 1.83E-07 |
| ENSMUST00000197767.4,Hax1-004,protein_coding,ENSMUSG00000027944.13,protein_coding | 1790.305054 | 0.359635194 | 5.17E-09 | 1.84E-07 |
| ENSMUST00000100799.8,Rpl15-002,protein_coding,ENSMUSG00000012405.15,protein_coding | 87135.78926 | 0.286136958 | 5.18E-09 | 1.85E-07 |
| ENSMUST00000022580.6,Slc25a30-201,protein_coding,ENSMUSG00000022003.6,protein_coding | 4525.539543 | 0.279366102 | 5.24E-09 | 1.86E-07 |
| ENSMUST00000086580.11,Yap1-002,protein_coding,ENSMUSG00000053110.13,protein_coding | 9272.75402 | -0.260662545 | 5.34E-09 | 1.90E-07 |
| ENSMUST00000040514.7,Irs2-001,protein_coding,ENSMUSG00000038894.7,protein_coding | 14653.455 | -0.266608252 | 5.38E-09 | 1.91E-07 |
| ENSMUST00000111639.7,Arntl2-001,protein_coding,ENSMUSG00000040187.15,protein_coding | 712.1764923 | 0.423917041 | 5.39E-09 | 1.91E-07 |
| ENSMUST00000025511.9,Rps14-001,protein_coding,ENSMUSG00000024608.10,protein_coding | 61738.9796 | 0.301397201 | 5.39E-09 | 1.91E-07 |
| ENSMUST00000026133.14,Pycr1-001,protein_coding,ENSMUSG00000025140.15,protein_coding | 6736.397508 | 0.46856454 | 5.43E-09 | 1.93E-07 |
| ENSMUST00000033898.9,Col4a1-001,protein_coding,ENSMUSG00000031502.11,protein_coding | 78653.24731 | 0.202826792 | 5.50E-09 | 1.95E-07 |
| ENSMUST00000102923.9,Aebp1-001,protein_coding,ENSMUSG00000020473.13,protein_coding | 22153.26728 | -0.212465149 | 5.50E-09 | 1.95E-07 |
| ENSMUST00000086523.6,Pxn-002,protein_coding,ENSMUSG00000029528.18,protein_coding | 3705.582728 | 0.335427903 | 5.54E-09 | 1.96E-07 |
| ENSMUST00000088217.11,Tbl1x-001,protein_coding,ENSMUSG00000025246.13,protein_coding | 24564.15009 | 0.228495514 | 5.57E-09 | 1.97E-07 |
| ENSMUST00000089559.10,Ddrgk1-001,protein_coding,ENSMUSG00000068290.11,protein_coding | 6447.348409 | 0.288771577 | 5.60E-09 | 1.98E-07 |
| ENSMUST00000130871.1,Gm28043-001,nonsense_mediated_decay,ENSMUSG00000098374.1,protein_coding | 1852.004088 | 0.44619528 | 5.69E-09 | 2.01E-07 |
| ENSMUST00000206265.1,Atxn2l-006,protein_coding,ENSMUSG00000032637.15,protein_coding | 202.5451652 | 0.873209555 | 5.73E-09 | 2.02E-07 |
| ENSMUST00000110086.1,Fgd3-003,protein_coding,ENSMUSG00000037946.11,protein_coding | 505.5378056 | -0.568097359 | 5.82E-09 | 2.05E-07 |
| ENSMUST00000084947.9,Trmt61a-201,protein_coding,ENSMUSG00000060950.11,protein_coding | 1469.327353 | -0.463417452 | 5.83E-09 | 2.05E-07 |
| ENSMUST00000194462.5,Slc7a11-004,protein_coding,ENSMUSG00000027737.10,protein_coding | 296.1479363 | 1.060669149 | 5.86E-09 | 2.06E-07 |
| ENSMUST00000094801.4,Ttc17-002,protein_coding,ENSMUSG00000027194.16,protein_coding | 1616.41015 | 0.533177278 | 5.89E-09 | 2.07E-07 |
| ENSMUST00000025215.8,Sil1-201,protein_coding,ENSMUSG00000024357.9,protein_coding | 3432.54396 | 0.370490713 | 5.95E-09 | 2.09E-07 |
| ENSMUST00000115239.9,Sept6-001,protein_coding,ENSMUSG00000050379.15,protein_coding | 1642.510121 | 0.337332453 | 5.98E-09 | 2.10E-07 |
| ENSMUST00000132404.7,Selenoi-003,nonsense_mediated_decay,ENSMUSG00000075703.15,protein_coding | 3197.485239 | 0.255434011 | 6.06E-09 | 2.12E-07 |
| ENSMUST00000004565.14,Ralb-001,protein_coding,ENSMUSG00000004451.14,protein_coding | 8089.124966 | 0.216180784 | 6.11E-09 | 2.14E-07 |
| ENSMUST00000112709.7,Tro-201,protein_coding,ENSMUSG00000025272.16,protein_coding | 263.19125 | -1.070752619 | 6.20E-09 | 2.17E-07 |
| ENSMUST00000150427.7,Ddb2-003,processed_transcript,ENSMUSG00000002109.14,protein_coding | 119.8450555 | 1.024920881 | 6.29E-09 | 2.20E-07 |
| ENSMUST00000176364.7,Retsat-004,protein_coding,ENSMUSG00000056666.13,protein_coding | 2590.628037 | 0.358522315 | 6.28E-09 | 2.20E-07 |
| ENSMUST00000128791.1,Ctsc-004,protein_coding,ENSMUSG00000030560.17,protein_coding | 403.2529533 | 0.77891579 | 6.33E-09 | 2.21E-07 |
| ENSMUST00000141108.1,Mib2-005,protein_coding,ENSMUSG00000029060.17,protein_coding | 611.6975499 | 0.593697736 | 6.35E-09 | 2.22E-07 |
| ENSMUST00000025803.8,Dkk1-001,protein_coding,ENSMUSG00000024868.8,protein_coding | 183.2159484 | -0.7031611 | 6.37E-09 | 2.22E-07 |
| ENSMUST00000025563.6,Fth1-201,protein_coding,ENSMUSG00000024661.6,protein_coding | 186555.5532 | 0.260905688 | 6.45E-09 | 2.25E-07 |
| ENSMUST00000173844.7,Rps28-001,protein_coding,ENSMUSG00000067288.12,protein_coding | 49009.62009 | 0.217729321 | 6.46E-09 | 2.25E-07 |
| ENSMUST00000042194.9,Trpm4-001,protein_coding,ENSMUSG00000038260.10,protein_coding | 880.314352 | 0.383443366 | 6.50E-09 | 2.26E-07 |
| ENSMUST00000134962.1,Abcc5-003,retained_intron,ENSMUSG00000022822.15,protein_coding | 788.3681702 | -0.569664411 | 6.64E-09 | 2.31E-07 |
| ENSMUST00000008826.13,Rpl10-001,protein_coding,ENSMUSG00000008682.13,protein_coding | 72522.48493 | 0.250025746 | 6.95E-09 | 2.41E-07 |
| ENSMUST00000054477.7,AI837181-002,processed_transcript,ENSMUSG00000047423.9,protein_coding | 1236.481681 | 0.818562522 | 7.00E-09 | 2.43E-07 |
| ENSMUST00000022616.12,Clu-001,protein_coding,ENSMUSG00000022037.14,protein_coding | 1217.579859 | -0.371321305 | 7.02E-09 | 2.43E-07 |
| ENSMUST00000080514.8,Rpp25-001,protein_coding,ENSMUSG00000062309.8,protein_coding | 530.4749947 | -0.471255435 | 7.07E-09 | 2.45E-07 |
| ENSMUST00000163031.1,Vti1b-003,nonsense_mediated_decay,ENSMUSG00000021124.13,protein_coding | 38.98962957 | 1.051906327 | 7.09E-09 | 2.45E-07 |
| ENSMUST00000031726.14,Gnb2-001,protein_coding,ENSMUSG00000029713.15,protein_coding | 35789.20518 | 0.223819524 | 7.11E-09 | 2.46E-07 |
| ENSMUST00000119787.8,Top3b-004,protein_coding,ENSMUSG00000022779.14,protein_coding | 1767.676202 | 0.548649447 | 7.15E-09 | 2.47E-07 |
| ENSMUST00000004134.10,Gstm5-001,protein_coding,ENSMUSG00000004032.10,protein_coding | 4023.129485 | 0.262062277 | 7.16E-09 | 2.47E-07 |
| ENSMUST00000166005.7,Zfp426-201,protein_coding,ENSMUSG00000059475.13,protein_coding | 52.55129601 | 0.948074943 | 7.16E-09 | 2.47E-07 |
| ENSMUST00000070118.13,Efemp2-002,protein_coding,ENSMUSG00000024909.14,protein_coding | 1384.718047 | -0.479308304 | 7.19E-09 | 2.48E-07 |
| ENSMUST00000128490.7,Dcaf11-015,protein_coding,ENSMUSG00000022214.14,protein_coding | 89.6598681 | -1.037326378 | 7.23E-09 | 2.49E-07 |
| ENSMUST00000142351.8,Vegfa-003,protein_coding,ENSMUSG00000023951.17,protein_coding | 1666.787409 | 0.466102289 | 7.25E-09 | 2.49E-07 |
| ENSMUST00000034377.7,Pla2g15-001,protein_coding,ENSMUSG00000031903.7,protein_coding | 2070.450708 | 0.334945696 | 7.26E-09 | 2.49E-07 |
| ENSMUST00000113215.9,Sh3bgrl2-001,protein_coding,ENSMUSG00000032261.16,protein_coding | 3052.954078 | 0.295714326 | 7.29E-09 | 2.51E-07 |
| ENSMUST00000170991.7,Serpinb6a-012,protein_coding,ENSMUSG00000060147.15,protein_coding | 190.7217613 | 0.72167596 | 7.37E-09 | 2.53E-07 |
| ENSMUST00000153474.8,Snhg12-002,lincRNA,ENSMUSG00000086290.8,lincRNA | 1361.19829 | 0.667852645 | 7.44E-09 | 2.55E-07 |
| ENSMUST00000027186.11,Trak2-001,protein_coding,ENSMUSG00000026028.12,protein_coding | 5305.996338 | -0.304706325 | 7.50E-09 | 2.57E-07 |
| ENSMUST00000201883.3,Rbpj-007,protein_coding,ENSMUSG00000039191.12,protein_coding | 2280.996439 | 0.565160647 | 7.53E-09 | 2.58E-07 |
| ENSMUST00000122328.7,Pxdn-001,protein_coding,ENSMUSG00000020674.16,protein_coding | 13506.49717 | -0.205128829 | 7.68E-09 | 2.63E-07 |
| ENSMUST00000114900.7,Zc3hav1-004,protein_coding,ENSMUSG00000029826.14,protein_coding | 13708.22461 | 0.299641764 | 7.72E-09 | 2.64E-07 |
| ENSMUST00000025503.8,Isoc1-201,protein_coding,ENSMUSG00000024601.8,protein_coding | 3476.804071 | 0.279654661 | 7.74E-09 | 2.65E-07 |
| ENSMUST00000044111.9,Rras-001,protein_coding,ENSMUSG00000038387.9,protein_coding | 2883.060599 | 0.25161386 | 7.76E-09 | 2.65E-07 |
| ENSMUST00000123355.7,Hspb6-003,processed_transcript,ENSMUSG00000036854.14,protein_coding | 1166.698462 | 0.503832005 | 7.93E-09 | 2.71E-07 |
| ENSMUST00000185198.6,Ralgps2-005,protein_coding,ENSMUSG00000026594.14,protein_coding | 916.2293382 | 0.706621099 | 7.96E-09 | 2.71E-07 |
| ENSMUST00000024884.4,Eif2ak2-001,protein_coding,ENSMUSG00000024079.4,protein_coding | 2607.11942 | -0.59309764 | 8.13E-09 | 2.77E-07 |
| ENSMUST00000178839.7,Ubtf-201,protein_coding,ENSMUSG00000020923.17,protein_coding | 3537.763291 | 0.217152509 | 8.17E-09 | 2.78E-07 |
| ENSMUST00000089482.11,Xpo4-201,protein_coding,ENSMUSG00000021952.15,protein_coding | 1554.484215 | 0.714097918 | 8.23E-09 | 2.80E-07 |
| ENSMUST00000060077.5,Cpox-201,protein_coding,ENSMUSG00000022742.5,protein_coding | 11250.3046 | 0.493611103 | 8.39E-09 | 2.85E-07 |
| ENSMUST00000009777.3,G0s2-001,protein_coding,ENSMUSG00000009633.3,protein_coding | 339.277069 | 0.722087977 | 8.48E-09 | 2.88E-07 |
| ENSMUST00000151358.7,Pdk1-003,processed_transcript,ENSMUSG00000006494.11,protein_coding | 1250.551959 | 0.491778698 | 8.53E-09 | 2.89E-07 |
| ENSMUST00000143107.1,Rpl27a-001,protein_coding,ENSMUSG00000046364.14,protein_coding | 85067.05967 | 0.230540631 | 8.61E-09 | 2.92E-07 |
| ENSMUST00000184105.7,Pank2-003,nonsense_mediated_decay,ENSMUSG00000037514.17,protein_coding | 1876.980256 | 0.330734758 | 8.65E-09 | 2.93E-07 |
| ENSMUST00000095439.9,Syne3-202,protein_coding,ENSMUSG00000054150.11,protein_coding | 7138.929846 | 0.237913841 | 8.68E-09 | 2.94E-07 |
| ENSMUST00000033342.6,Eif3f-001,protein_coding,ENSMUSG00000031029.6,protein_coding | 32074.28384 | 0.243453223 | 8.78E-09 | 2.97E-07 |
| ENSMUST00000021911.14,2010111I01Rik-201,protein_coding,ENSMUSG00000021458.16,protein_coding | 1266.555625 | -0.481300491 | 8.95E-09 | 3.02E-07 |
| ENSMUST00000043938.6,Plcl2-201,protein_coding,ENSMUSG00000038910.6,protein_coding | 833.778733 | 0.635277923 | 9.08E-09 | 3.07E-07 |
| ENSMUST00000119597.7,Slc18b1-001,protein_coding,ENSMUSG00000037455.16,protein_coding | 1586.214508 | 0.405452774 | 9.10E-09 | 3.07E-07 |
| ENSMUST00000037551.14,Ppp1r16a-001,protein_coding,ENSMUSG00000033819.15,protein_coding | 3641.545081 | -0.219589257 | 9.11E-09 | 3.07E-07 |
| ENSMUST00000030538.4,Ddost-001,protein_coding,ENSMUSG00000028757.4,protein_coding | 34670.0455 | 0.214882064 | 9.13E-09 | 3.07E-07 |
| ENSMUST00000145477.7,Sms-005,nonsense_mediated_decay,ENSMUSG00000071708.11,protein_coding | 6007.894869 | 0.267961164 | 9.20E-09 | 3.10E-07 |
| ENSMUST00000119299.7,Ogt-002,protein_coding,ENSMUSG00000034160.13,protein_coding | 1169.002894 | -0.452696191 | 9.28E-09 | 3.12E-07 |
| ENSMUST00000201352.3,Bre-001,protein_coding,ENSMUSG00000052139.18,protein_coding | 4025.827056 | 0.298888216 | 9.31E-09 | 3.13E-07 |
| ENSMUST00000048400.3,Tnks1bp1-003,protein_coding,ENSMUSG00000033955.12,protein_coding | 8702.118768 | 0.414510728 | 9.36E-09 | 3.14E-07 |
| ENSMUST00000057311.3,Sfn-001,protein_coding,ENSMUSG00000047281.3,protein_coding | 511.5881875 | 0.558587811 | 9.46E-09 | 3.17E-07 |
| ENSMUST00000209787.1,Primpol-006,protein_coding,ENSMUSG00000038225.15,protein_coding | 466.3302944 | 0.448777407 | 9.51E-09 | 3.19E-07 |
| ENSMUST00000041544.7,Fbln2-002,protein_coding,ENSMUSG00000064080.12,protein_coding | 8457.200674 | -0.354834913 | 9.53E-09 | 3.19E-07 |
| ENSMUST00000099224.9,Csnk2a1-001,protein_coding,ENSMUSG00000074698.10,protein_coding | 28920.48591 | 0.135082455 | 9.66E-09 | 3.23E-07 |
| ENSMUST00000057897.7,Zfand3-201,protein_coding,ENSMUSG00000044477.10,protein_coding | 11117.62569 | 0.282287092 | 9.81E-09 | 3.28E-07 |

| | | | | |
|---|---|---|---|---|
| ENSMUST00000163272.1,Suz12-201,protein_coding,ENSMUSG00000017548.15,protein_coding | 1334.83134 | -0.686653991 | 9.83E-09 | 3.28E-07 |
| ENSMUST00000023087.12,Twf1-001,protein_coding,ENSMUSG00000022451.12,protein_coding | 16812.9612 | -0.237322359 | 1.01E-08 | 3.36E-07 |
| ENSMUST00000172314.8,Hbp1-001,protein_coding,ENSMUSG00000002996.17,protein_coding | 2383.248818 | 0.481319956 | 1.01E-08 | 3.38E-07 |
| ENSMUST00000152813.1,Spata5l1-002,processed_transcript,ENSMUSG00000074876.5,processed_transcript | 592.3521732 | 0.524796521 | 1.04E-08 | 3.47E-07 |
| ENSMUST00000026420.5,Rps26-201,protein_coding,ENSMUSG00000025362.5,protein_coding | 84710.33744 | 0.190195884 | 1.04E-08 | 3.47E-07 |
| ENSMUST00000096246.3,Ganab-201,protein_coding,ENSMUSG00000071650.5,protein_coding | 44487.24064 | 0.193278347 | 1.05E-08 | 3.48E-07 |
| ENSMUST00000013262.14,Zkscan17-001,protein_coding,ENSMUSG00000020472.14,protein_coding | 10353.6137 | 0.330954686 | 1.05E-08 | 3.50E-07 |
| ENSMUST00000167911.1,Sptlc2-004,processed_transcript,ENSMUSG00000021036.10,protein_coding | 1504.080295 | 0.317214368 | 1.06E-08 | 3.51E-07 |
| ENSMUST00000170010.1,Banf1-202,protein_coding,ENSMUSG00000024844.14,protein_coding | 1711.859059 | 0.404754303 | 1.06E-08 | 3.51E-07 |
| ENSMUST00000117146.7,Ube2w-001,protein_coding,ENSMUSG00000025939.17,protein_coding | 3918.407228 | 0.346856132 | 1.07E-08 | 3.55E-07 |
| ENSMUST00000040718.5,Ostm1-004,protein_coding,ENSMUSG00000038280.12,protein_coding | 2280.211704 | -0.240703958 | 1.08E-08 | 3.57E-07 |
| ENSMUST00000108796.1,Dnajc5-004,protein_coding,ENSMUSG00000000826.16,protein_coding | 692.1597397 | 0.61570271 | 1.08E-08 | 3.59E-07 |
| ENSMUST00000106004.7,Rplp2-004,protein_coding,ENSMUSG00000025508.13,protein_coding | 25976.75943 | 0.238426618 | 1.10E-08 | 3.64E-07 |
| ENSMUST00000073308.10,Ewsr1-007,protein_coding,ENSMUSG00000009079.16,protein_coding | 16353.83868 | 0.296440205 | 1.10E-08 | 3.65E-07 |
| ENSMUST00000030993.7,Nelfa-001,protein_coding,ENSMUSG00000029111.7,protein_coding | 8475.177046 | 0.194050805 | 1.11E-08 | 3.68E-07 |
| ENSMUST00000005953.10,Sqrdl-001,protein_coding,ENSMUSG00000005803.14,protein_coding | 2614.965335 | 0.343875668 | 1.12E-08 | 3.71E-07 |
| ENSMUST00000021938.9,Aaed1-201,protein_coding,ENSMUSG00000021482.9,protein_coding | 1256.467607 | 0.428875781 | 1.12E-08 | 3.71E-07 |
| ENSMUST00000023760.12,Gpd1-001,protein_coding,ENSMUSG00000023019.12,protein_coding | 1120.604899 | -0.787084528 | 1.13E-08 | 3.71E-07 |
| ENSMUST00000042290.13,Itih2-001,protein_coding,ENSMUSG00000037254.18,protein_coding | 974.4599433 | 0.467237972 | 1.17E-08 | 3.86E-07 |
| ENSMUST00000079724.8,Hax1-001,protein_coding,ENSMUSG00000027944.13,protein_coding | 15695.22416 | 0.312104927 | 1.17E-08 | 3.87E-07 |
| ENSMUST00000033878.13,Rab11fip1-001,protein_coding,ENSMUSG00000031488.14,protein_coding | 1479.195202 | 0.286360727 | 1.19E-08 | 3.90E-07 |
| ENSMUST00000145593.7,Paip2b-003,retained_intron,ENSMUSG00000045896.14,protein_coding | 255.2497367 | 0.641160408 | 1.19E-08 | 3.92E-07 |
| ENSMUST00000038514.14,Nup54-001,protein_coding,ENSMUSG00000034826.14,protein_coding | 9538.57192 | 0.222696331 | 1.20E-08 | 3.94E-07 |
| ENSMUST00000025077.6,Mtpap-201,protein_coding,ENSMUSG00000024234.6,protein_coding | 3472.379481 | 0.202729299 | 1.21E-08 | 3.96E-07 |
| ENSMUST00000093138.12,Tbc1d9b-001,protein_coding,ENSMUSG00000036644.15,protein_coding | 9151.882345 | 0.307508319 | 1.22E-08 | 4.01E-07 |
| ENSMUST00000111742.7,Bcat1-002,protein_coding,ENSMUSG00000030268.17,protein_coding | 41386.73698 | 0.294524143 | 1.22E-08 | 4.01E-07 |
| ENSMUST00000214547.1,Neo1-002,protein_coding,ENSMUSG00000032340.8,protein_coding | 16559.911 | 0.173214116 | 1.25E-08 | 4.08E-07 |
| ENSMUST00000114871.1,Hs6st2-002,protein_coding,ENSMUSG00000062184.11,protein_coding | 1997.312983 | -0.521638903 | 1.25E-08 | 4.10E-07 |
| ENSMUST00000167496.7,Rnf20-201,protein_coding,ENSMUSG00000028309.14,protein_coding | 177.9089994 | -0.908686152 | 1.26E-08 | 4.12E-07 |
| ENSMUST00000025208.5,Dnajc18-201,protein_coding,ENSMUSG00000024350.5,protein_coding | 2357.395407 | -0.248864325 | 1.29E-08 | 4.20E-07 |
| ENSMUST00000109354.9,Npm1-002,protein_coding,ENSMUSG00000057113.13,protein_coding | 47086.21284 | 0.460951521 | 1.29E-08 | 4.22E-07 |
| ENSMUST00000031727.9,Gigyf1-001,protein_coding,ENSMUSG00000029714.11,protein_coding | 4261.394548 | 0.235441403 | 1.30E-08 | 4.26E-07 |
| ENSMUST00000118365.2,Card6-001,protein_coding,ENSMUSG00000041849.7,protein_coding | 377.3352856 | 0.641723363 | 1.31E-08 | 4.26E-07 |
| ENSMUST00000055458.4,Cdc42ep2-201,protein_coding,ENSMUSG00000045664.4,protein_coding | 1432.552871 | -0.568540107 | 1.31E-08 | 4.26E-07 |
| ENSMUST00000092990.3,Agpat9-003,protein_coding,ENSMUSG00000029314.14,protein_coding | 683.3028474 | 0.61390617 | 1.31E-08 | 4.27E-07 |
| ENSMUST00000133182.1,Mkln1-008,retained_intron,ENSMUSG00000025609.15,protein_coding | 37.36949429 | 1.031806095 | 1.31E-08 | 4.27E-07 |
| ENSMUST00000181974.7,Ank3-024,protein_coding,ENSMUSG00000069601.13,protein_coding | 818.7775052 | -0.781978584 | 1.32E-08 | 4.28E-07 |
| ENSMUST00000026262.7,Hexa-001,protein_coding,ENSMUSG00000025232.8,protein_coding | 7450.986134 | 0.321712929 | 1.37E-08 | 4.44E-07 |
| ENSMUST00000030709.8,Smpdl3b-001,protein_coding,ENSMUSG00000028885.8,protein_coding | 1304.363066 | -0.342432433 | 1.39E-08 | 4.50E-07 |
| ENSMUST00000088295.8,Chpf2-001,protein_coding,ENSMUSG00000038181.16,protein_coding | 453.483354 | -0.474453777 | 1.40E-08 | 4.54E-07 |
| ENSMUST00000023694.9,1110004E09Rik-001,protein_coding,ENSMUSG00000022972.9,protein_coding | 4548.967993 | 0.218333728 | 1.40E-08 | 4.54E-07 |
| ENSMUST00000095350.10,Aebp2-003,protein_coding,ENSMUSG00000030232.17,protein_coding | 310.4081373 | 0.627329854 | 1.40E-08 | 4.55E-07 |
| ENSMUST00000023462.12,Mapk1-002,protein_coding,ENSMUSG00000063358.15,protein_coding | 13816.46213 | 0.159380885 | 1.42E-08 | 4.58E-07 |
| ENSMUST00000028815.14,Slc23a2-001,protein_coding,ENSMUSG00000027340.15,protein_coding | 6938.21943 | 0.272480543 | 1.42E-08 | 4.61E-07 |
| ENSMUST00000089157.9,Cenpm-202,protein_coding,ENSMUSG00000068101.10,protein_coding | 3231.512711 | 0.260672975 | 1.43E-08 | 4.61E-07 |
| ENSMUST00000192702.5,Uck2-002,nonsense_mediated_decay,ENSMUSG00000026558.13,protein_coding | 3215.778593 | 0.343501111 | 1.43E-08 | 4.63E-07 |
| ENSMUST00000200069.4,Nras-006,protein_coding,ENSMUSG00000027852.14,protein_coding | 1765.384704 | 0.425874474 | 1.48E-08 | 4.78E-07 |
| ENSMUST00000170217.7,Farsb-002,protein_coding,ENSMUSG00000026245.16,protein_coding | 555.1571461 | 0.910136974 | 1.50E-08 | 4.82E-07 |
| ENSMUST00000155551.7,Dcaf10-001,protein_coding,ENSMUSG00000035572.16,protein_coding | 2321.675212 | -0.333843879 | 1.50E-08 | 4.83E-07 |
| ENSMUST00000107868.7,Tom1l1-002,protein_coding,ENSMUSG00000020541.12,protein_coding | 1138.301284 | 0.322565491 | 1.51E-08 | 4.87E-07 |
| ENSMUST00000114036.8,Son-002,protein_coding,ENSMUSG00000022961.17,protein_coding | 10019.35433 | 0.226985894 | 1.51E-08 | 4.87E-07 |
| ENSMUST00000059193.5,F2r-201,protein_coding,ENSMUSG00000048376.5,protein_coding | 31911.25119 | -0.187199152 | 1.52E-08 | 4.88E-07 |
| ENSMUST00000169397.7,Slc39a7-201,protein_coding,ENSMUSG00000024327.15,protein_coding | 4062.068281 | 0.554665222 | 1.53E-08 | 4.93E-07 |
| ENSMUST00000176257.7,Hnrnpk-020,retained_intron,ENSMUSG00000021546.17,protein_coding | 2249.25078 | 0.349587482 | 1.56E-08 | 5.00E-07 |
| ENSMUST00000008573.8,Herpud2-001,protein_coding,ENSMUSG00000008429.8,protein_coding | 2573.76395 | 0.298963865 | 1.61E-08 | 5.16E-07 |
| ENSMUST00000022082.7,Glrx-001,protein_coding,ENSMUSG00000021591.7,protein_coding | 757.9876632 | -0.438658142 | 1.62E-08 | 5.20E-07 |
| ENSMUST00000000828.12,Txnrd3-001,protein_coding,ENSMUSG00000000811.13,protein_coding | 3326.738045 | 0.316555845 | 1.67E-08 | 5.34E-07 |
| ENSMUST00000061052.11,Atat1-001,protein_coding,ENSMUSG00000024426.17,protein_coding | 327.2285044 | 0.671181576 | 1.68E-08 | 5.39E-07 |
| ENSMUST00000151854.1,Hspa4-002,processed_transcript,ENSMUSG00000020361.13,protein_coding | 11288.07942 | 0.364651614 | 1.70E-08 | 5.44E-07 |
| ENSMUST00000204145.2,Plekha5-004,nonsense_mediated_decay,ENSMUSG00000030231.11,protein_coding | 1512.159865 | 0.424803597 | 1.71E-08 | 5.46E-07 |
| ENSMUST00000029135.14,Acss2-001,protein_coding,ENSMUSG00000027605.18,protein_coding | 1442.318596 | 0.536109546 | 1.72E-08 | 5.49E-07 |
| ENSMUST00000097649.9,Lrrfip1-003,protein_coding,ENSMUSG00000026305.15,protein_coding | 7814.84827 | -0.414330365 | 1.72E-08 | 5.49E-07 |
| ENSMUST00000048260.13,Lman1-001,protein_coding,ENSMUSG00000041891.15,protein_coding | 13904.13086 | 0.268142653 | 1.72E-08 | 5.49E-07 |
| ENSMUST00000035938.2,Ccl5-001,protein_coding,ENSMUSG00000035042.2,protein_coding | 754.4648592 | 0.473424625 | 1.73E-08 | 5.50E-07 |
| ENSMUST00000022992.12,Tbc1d31-001,protein_coding,ENSMUSG00000022364.14,protein_coding | 10414.27404 | 0.322418923 | 1.74E-08 | 5.54E-07 |
| ENSMUST00000080926.12,Eno1-001,protein_coding,ENSMUSG00000063524.13,protein_coding | 392938.5075 | 0.257667866 | 1.74E-08 | 5.55E-07 |
| ENSMUST00000071370.5,Pcdh17-201,protein_coding,ENSMUSG00000035566.6,protein_coding | 248.1728691 | -0.657472825 | 1.78E-08 | 5.66E-07 |
| ENSMUST00000060722.6,Cxxc5-201,protein_coding,ENSMUSG00000046668.8,protein_coding | 2674.131262 | -0.348076908 | 1.80E-08 | 5.72E-07 |
| ENSMUST00000032768.14,Nr2f2-001,protein_coding,ENSMUSG00000030551.14,protein_coding | 5910.507535 | 0.445164853 | 1.81E-08 | 5.75E-07 |
| ENSMUST00000193682.5,Map4k4-005,protein_coding,ENSMUSG00000026074.14,protein_coding | 22711.22617 | 0.18880272 | 1.82E-08 | 5.78E-07 |
| ENSMUST00000060837.9,Rab11fip5-002,protein_coding,ENSMUSG00000051343.11,protein_coding | 6055.199409 | 0.381618656 | 1.83E-08 | 5.80E-07 |
| ENSMUST00000033699.12,Flna-003,protein_coding,ENSMUSG00000031328.15,protein_coding | 215450.6663 | 0.203565092 | 1.86E-08 | 5.89E-07 |
| ENSMUST00000105336.8,Dot1l-001,protein_coding,ENSMUSG00000061589.14,protein_coding | 9002.29757 | -0.203948565 | 1.86E-08 | 5.89E-07 |
| ENSMUST00000003808.7,Gdf15-001,protein_coding,ENSMUSG00000038508.7,protein_coding | 917.260826 | 0.581639555 | 1.89E-08 | 5.97E-07 |
| ENSMUST00000131810.7,Dnajb12-003,nonsense_mediated_decay,ENSMUSG00000020109.13,protein_coding | 1139.730209 | 0.51626482 | 1.94E-08 | 6.12E-07 |
| ENSMUST00000098106.8,Rusc2-003,protein_coding,ENSMUSG00000035969.15,protein_coding | 1195.298266 | -0.703598453 | 1.99E-08 | 6.28E-07 |
| ENSMUST00000026470.4,Shmt2-201,protein_coding,ENSMUSG00000025403.4,protein_coding | 86761.23558 | 0.27077178 | 1.99E-08 | 6.29E-07 |
| ENSMUST00000071750.12,Col12a1-001,protein_coding,ENSMUSG00000032332.17,protein_coding | 4184.568851 | -0.307882945 | 1.99E-08 | 6.29E-07 |
| ENSMUST00000116605.7,Mad2l1-001,protein_coding,ENSMUSG00000029910.14,protein_coding | 1608.27781 | -0.483010915 | 2.00E-08 | 6.32E-07 |
| ENSMUST00000100857.9,Dusp16-001,protein_coding,ENSMUSG00000030203.17,protein_coding | 3677.20135 | -0.352781148 | 2.02E-08 | 6.35E-07 |
| ENSMUST00000148894.1,Luc7l-001,nonsense_mediated_decay,ENSMUSG00000024188.15,protein_coding | 823.2787107 | 0.464115084 | 2.02E-08 | 6.36E-07 |
| ENSMUST00000037607.10,Map3k7-001,protein_coding,ENSMUSG00000028284.13,protein_coding | 2818.506019 | -0.582556255 | 2.04E-08 | 6.42E-07 |
| ENSMUST00000203626.1,Zyx-011,processed_transcript,ENSMUSG00000029860.16,protein_coding | 772.5288791 | 0.456817066 | 2.04E-08 | 6.42E-07 |
| ENSMUST00000027941.13,Atf3-001,protein_coding,ENSMUSG00000026628.13,protein_coding | 1641.872057 | 0.640443994 | 2.08E-08 | 6.52E-07 |
| ENSMUST00000073080.5,Gm10053-201,protein_coding,ENSMUSG00000058927.5,protein_coding | 2030.050405 | -0.359796824 | 2.10E-08 | 6.60E-07 |
| ENSMUST00000115628.8,Tango2-001,protein_coding,ENSMUSG00000013539.13,protein_coding | 900.4158794 | -0.378348318 | 2.12E-08 | 6.63E-07 |
| ENSMUST00000026234.4,Kazald1-201,protein_coding,ENSMUSG00000025213.11,protein_coding | 1255.065702 | -0.465209722 | 2.17E-08 | 6.79E-07 |
| ENSMUST00000020851.14,Cox11-001,protein_coding,ENSMUSG00000020544.14,protein_coding | 800.5454334 | -0.392263049 | 2.17E-08 | 6.80E-07 |
| ENSMUST00000029769.13,Gclm-001,protein_coding,ENSMUSG00000028124.15,protein_coding | 4606.230095 | -0.433542384 | 2.20E-08 | 6.88E-07 |
| ENSMUST00000052740.13,Tcaim-001,protein_coding,ENSMUSG00000046603.16,protein_coding | 589.8838864 | 0.599397068 | 2.21E-08 | 6.90E-07 |
| ENSMUST00000086199.11,Glul-001,protein_coding,ENSMUSG00000026473.15,protein_coding | 5692.195074 | 0.26229678 | 2.21E-08 | 6.91E-07 |
| ENSMUST00000186247.6,Ank3-038,protein_coding,ENSMUSG00000069601.13,protein_coding | 211.38893 | -0.757315801 | 2.23E-08 | 6.95E-07 |

| | | | | |
|---|---|---|---|---|
| ENSMUST00000037941.9,Cd81-001,protein_coding,ENSMUSG00000037706.17,protein_coding | 11203.05084 | 0.222735196 | 2.24E-08 | 6.99E-07 |
| ENSMUST00000047218.3,Reep4-201,protein_coding,ENSMUSG00000033589.3,protein_coding | 9059.762031 | 0.195175896 | 2.25E-08 | 7.03E-07 |
| ENSMUST00000110104.8,Zc3h14-203,protein_coding,ENSMUSG00000021012.15,protein_coding | 3047.501825 | 0.290948498 | 2.26E-08 | 7.03E-07 |
| ENSMUST00000048774.12,Copg2-201,protein_coding,ENSMUSG00000025607.15,protein_coding | 4121.620489 | 0.352001389 | 2.26E-08 | 7.05E-07 |
| ENSMUST00000189413.6,Kif14-001,protein_coding,ENSMUSG00000041498.13,protein_coding | 2465.348726 | -0.239362877 | 2.27E-08 | 7.07E-07 |
| ENSMUST00000038361.10,Mgat5-001,protein_coding,ENSMUSG00000036155.13,protein_coding | 5655.417104 | -0.173626637 | 2.29E-08 | 7.11E-07 |
| ENSMUST00000112350.7,Mta3-002,protein_coding,ENSMUSG00000055817.17,protein_coding | 2854.615817 | 0.410311726 | 2.29E-08 | 7.11E-07 |
| ENSMUST00000146511.7,Gtf3a-001,protein_coding,ENSMUSG00000016503.16,protein_coding | 9590.897768 | 0.218429367 | 2.29E-08 | 7.11E-07 |
| ENSMUST00000022239.6,Parp8-201,protein_coding,ENSMUSG00000021725.8,protein_coding | 2095.12215 | 0.337014166 | 2.29E-08 | 7.11E-07 |
| ENSMUST00000035672.3,Ppl-201,protein_coding,ENSMUSG00000039457.3,protein_coding | 6021.102933 | -0.511473803 | 2.30E-08 | 7.13E-07 |
| ENSMUST00000140772.1,Igfbp4-002,protein_coding,ENSMUSG00000017493.12,protein_coding | 453.9203339 | 0.951683517 | 2.36E-08 | 7.30E-07 |
| ENSMUST00000135179.7,Cdk4-010,retained_intron,ENSMUSG00000006728.12,protein_coding | 1342.995013 | 0.5248851 | 2.42E-08 | 7.49E-07 |
| ENSMUST00000045756.13,S100a10-001,protein_coding,ENSMUSG00000041959.14,protein_coding | 50655.28291 | 0.292156439 | 2.44E-08 | 7.54E-07 |
| ENSMUST00000115028.9,Itgb5-001,protein_coding,ENSMUSG00000022817.14,protein_coding | 10163.81369 | 0.306302349 | 2.45E-08 | 7.56E-07 |
| ENSMUST00000017868.6,Etv4-002,protein_coding,ENSMUSG00000017724.14,protein_coding | 7561.793195 | 0.301767758 | 2.45E-08 | 7.57E-07 |
| ENSMUST00000014614.3,Rnf166-001,protein_coding,ENSMUSG00000014470.4,protein_coding | 4655.785683 | 0.378728003 | 2.49E-08 | 7.67E-07 |
| ENSMUST00000078508.5,Sp7-201,protein_coding,ENSMUSG00000060284.6,protein_coding | 1592.437382 | -0.555528553 | 2.53E-08 | 7.80E-07 |
| ENSMUST00000111018.1,Srp9-002,protein_coding,ENSMUSG00000026511.7,protein_coding | 937.228662 | 0.513710157 | 2.55E-08 | 7.87E-07 |
| ENSMUST00000118875.7,Gapdh-001,protein_coding,ENSMUSG00000057666.18,protein_coding | 431927.1415 | 0.247497809 | 2.56E-08 | 7.89E-07 |
| ENSMUST00000021062.11,Ddx5-001,protein_coding,ENSMUSG00000020719.14,protein_coding | 37033.20585 | -0.317432629 | 2.58E-08 | 7.93E-07 |
| ENSMUST00000155873.2,Rpl11-002,protein_coding,ENSMUSG00000059291.15,protein_coding | 41333.39588 | 0.270694496 | 2.59E-08 | 7.95E-07 |
| ENSMUST00000081554.12,P2rx4-002,protein_coding,ENSMUSG00000029470.15,protein_coding | 2206.01709 | 0.443020829 | 2.62E-08 | 8.04E-07 |
| ENSMUST00000183461.7,Camsap1-202,protein_coding,ENSMUSG00000026933.17,protein_coding | 3122.056788 | 0.244390605 | 2.64E-08 | 8.10E-07 |
| ENSMUST00000048248.8,Brsk1-001,protein_coding,ENSMUSG00000035390.16,protein_coding | 2748.176918 | -0.247681965 | 2.66E-08 | 8.16E-07 |
| ENSMUST00000057343.15,Ptbp1-008,nonsense_mediated_decay,ENSMUSG00000006498.16,protein_coding | 10944.54187 | 0.262408386 | 2.67E-08 | 8.17E-07 |
| ENSMUST00000138509.7,Pank2-005,processed_transcript,ENSMUSG00000037514.17,protein_coding | 726.1612678 | 0.705453168 | 2.67E-08 | 8.17E-07 |
| ENSMUST00000050331.12,Hnrnph2-004,protein_coding,ENSMUSG00000045427.13,protein_coding | 2224.556938 | -0.391008389 | 2.71E-08 | 8.29E-07 |
| ENSMUST00000200097.4,Ptbp2-001,protein_coding,ENSMUSG00000028134.11,protein_coding | 705.9966418 | -0.555561923 | 2.72E-08 | 8.32E-07 |
| ENSMUST00000172361.7,Hnrnpd-001,protein_coding,ENSMUSG00000000568.15,protein_coding | 924.3799645 | -0.748146819 | 2.73E-08 | 8.35E-07 |
| ENSMUST00000100527.10,Puf60-202,protein_coding,ENSMUSG00000002524.15,protein_coding | 10188.35973 | -0.297143086 | 2.73E-08 | 8.35E-07 |
| ENSMUST00000147891.7,Son-004,nonsense_mediated_decay,ENSMUSG00000022961.17,protein_coding | 1417.921672 | -0.769980619 | 2.74E-08 | 8.35E-07 |
| ENSMUST00000054220.9,Rpl18a-001,protein_coding,ENSMUSG00000045128.9,protein_coding | 1161.009665 | 0.352431887 | 2.74E-08 | 8.36E-07 |
| ENSMUST00000059658.5,Gm867-001,antisense,ENSMUSG00000050157.5,antisense | 157.2789995 | 0.819481335 | 2.76E-08 | 8.40E-07 |
| ENSMUST00000204206.1,Serbp1-011,retained_intron,ENSMUSG00000036371.6,protein_coding | 10888.00257 | 0.331652661 | 2.76E-08 | 8.41E-07 |
| ENSMUST00000154995.1,Anapc1-009,processed_transcript,ENSMUSG00000014355.10,protein_coding | 1152.690682 | 0.406751815 | 2.76E-08 | 8.41E-07 |
| ENSMUST00000147632.1,Psmg4-001,protein_coding,ENSMUSG00000071451.9,protein_coding | 1373.359641 | 0.344484627 | 2.77E-08 | 8.43E-07 |
| ENSMUST00000059997.8,A430105I19Rik-001,protein_coding,ENSMUSG00000045838.8,protein_coding | 2748.735393 | -0.379277465 | 2.79E-08 | 8.48E-07 |
| ENSMUST00000169810.7,Racgap1-011,protein_coding,ENSMUSG00000023015.13,protein_coding | 1074.315077 | 0.508179106 | 2.81E-08 | 8.55E-07 |
| ENSMUST00000028610.9,Cat-001,protein_coding,ENSMUSG00000027187.10,protein_coding | 9874.084383 | 0.167911139 | 2.82E-08 | 8.57E-07 |
| ENSMUST00000110894.8,Tpt1-001,protein_coding,ENSMUSG00000060126.14,protein_coding | 160260.2241 | 0.191518758 | 2.84E-08 | 8.63E-07 |
| ENSMUST00000034441.7,Aars-001,protein_coding,ENSMUSG00000031960.14,protein_coding | 105236.9859 | 0.244173353 | 2.87E-08 | 8.69E-07 |
| ENSMUST00000044828.13,Lgr6-001,protein_coding,ENSMUSG00000042793.13,protein_coding | 4577.923145 | -0.484817344 | 2.88E-08 | 8.71E-07 |
| ENSMUST00000013759.5,Fnbp4-001,protein_coding,ENSMUSG00000008200.14,protein_coding | 7412.528555 | 0.185476033 | 2.95E-08 | 8.92E-07 |
| ENSMUST00000038644.4,Rangrf-001,protein_coding,ENSMUSG00000032892.14,protein_coding | 2909.250564 | -0.364128765 | 3.06E-08 | 9.26E-07 |
| ENSMUST00000022341.5,2700060E02Rik-201,protein_coding,ENSMUSG00000021807.5,protein_coding | 17085.38001 | 0.231867506 | 3.08E-08 | 9.32E-07 |
| ENSMUST00000023119.14,Vdr-001,protein_coding,ENSMUSG00000022479.15,protein_coding | 984.367185 | -0.383801634 | 3.17E-08 | 9.59E-07 |
| ENSMUST00000120524.1,Tmem132a-002,protein_coding,ENSMUSG00000024736.14,protein_coding | 1328.784976 | -0.563737666 | 3.25E-08 | 9.82E-07 |
| ENSMUST00000105346.9,Tcf3-001,protein_coding,ENSMUSG00000020167.14,protein_coding | 7001.829631 | 0.237548192 | 3.28E-08 | 9.90E-07 |
| ENSMUST00000173801.7,Syncrip-002,protein_coding,ENSMUSG00000032423.12,protein_coding | 2885.762272 | -0.47161623 | 3.32E-08 | 1.00E-06 |
| ENSMUST00000143899.1,Ndel1-004,processed_transcript,ENSMUSG00000018736.14,protein_coding | 381.9234313 | -0.571834987 | 3.34E-08 | 1.01E-06 |
| ENSMUST00000010597.9,Kansl3-001,protein_coding,ENSMUSG00000010453.12,protein_coding | 10761.44134 | 0.25376265 | 3.35E-08 | 1.01E-06 |
| ENSMUST00000055576.11,Pcsk6-001,protein_coding,ENSMUSG00000030513.14,protein_coding | 1469.960748 | 0.709961634 | 3.36E-08 | 1.01E-06 |
| ENSMUST00000156160.1,Smarce1-006,processed_transcript,ENSMUSG00000037935.16,protein_coding | 2072.991408 | 0.308242011 | 3.39E-08 | 1.02E-06 |
| ENSMUST00000027839.13,Uck2-008,protein_coding,ENSMUSG00000026558.13,protein_coding | 3098.630259 | 0.282724578 | 3.42E-08 | 1.03E-06 |
| ENSMUST00000034904.13,Elovl5-001,protein_coding,ENSMUSG00000032349.13,protein_coding | 12266.69897 | 0.178665928 | 3.43E-08 | 1.03E-06 |
| ENSMUST00000041048.5,Orai2-001,protein_coding,ENSMUSG00000039747.11,protein_coding | 5966.466585 | -0.305088824 | 3.52E-08 | 1.06E-06 |
| ENSMUST00000162244.7,Cpsf4-003,protein_coding,ENSMUSG00000029625.16,protein_coding | 1425.731453 | 0.434917637 | 3.52E-08 | 1.06E-06 |
| ENSMUST00000190997.6,Rbbp5-002,protein_coding,ENSMUSG00000026439.14,protein_coding | 3942.390018 | 0.2839868 | 3.54E-08 | 1.06E-06 |
| ENSMUST00000106644.8,Nrd1-002,protein_coding,ENSMUSG00000053510.12,protein_coding | 10426.36322 | 0.221325622 | 3.57E-08 | 1.07E-06 |
| ENSMUST00000072838.5,Hsf1-201,protein_coding,ENSMUSG00000022556.9,protein_coding | 11041.82635 | -0.148135142 | 3.57E-08 | 1.07E-06 |
| ENSMUST00000035007.9,Cmtm6-001,protein_coding,ENSMUSG00000032434.9,protein_coding | 3518.694552 | -0.294866636 | 3.59E-08 | 1.07E-06 |
| ENSMUST00000040668.8,Osbpl2-001,protein_coding,ENSMUSG00000039050.8,protein_coding | 4616.52894 | 0.356514531 | 3.59E-08 | 1.07E-06 |
| ENSMUST00000115094.7,Xiap-003,protein_coding,ENSMUSG00000025860.14,protein_coding | 3122.135999 | -0.583431563 | 3.59E-08 | 1.07E-06 |
| ENSMUST00000036248.12,Pmepa1-001,protein_coding,ENSMUSG00000038400.15,protein_coding | 10375.63389 | -0.475421819 | 3.62E-08 | 1.08E-06 |
| ENSMUST00000140205.7,Adam10-002,nonsense_mediated_decay,ENSMUSG00000054693.14,protein_coding | 2312.794067 | -0.593596163 | 3.62E-08 | 1.08E-06 |
| ENSMUST00000028102.13,Kif5c-001,protein_coding,ENSMUSG00000026764.15,protein_coding | 332.1441817 | -0.63325052 | 3.65E-08 | 1.09E-06 |
| ENSMUST00000032441.13,Ccdc91-001,protein_coding,ENSMUSG00000030301.17,protein_coding | 3989.7594 | 0.272228289 | 3.65E-08 | 1.09E-06 |
| ENSMUST00000060738.8,S100a1-001,protein_coding,ENSMUSG00000044080.9,protein_coding | 4519.798414 | 0.34898394 | 3.68E-08 | 1.10E-06 |
| ENSMUST00000033827.7,Grk1-201,protein_coding,ENSMUSG00000031450.8,protein_coding | 27.14442036 | 1.026250408 | 3.69E-08 | 1.10E-06 |
| ENSMUST00000208532.1,Rps3-005,protein_coding,ENSMUSG00000030744.13,protein_coding | 21243.50509 | 0.295536865 | 3.71E-08 | 1.10E-06 |
| ENSMUST00000109895.1,Gas2l1-002,protein_coding,ENSMUSG00000034201.13,protein_coding | 2663.58553 | -0.456572634 | 3.71E-08 | 1.10E-06 |
| ENSMUST00000029944.12,Ube2j1-003,protein_coding,ENSMUSG00000028277.13,protein_coding | 1219.17428 | 0.474119718 | 3.73E-08 | 1.11E-06 |
| ENSMUST00000150320.2,Ankrd16-003,retained_intron,ENSMUSG00000047909.11,protein_coding | 130.8582107 | -0.759284595 | 3.78E-08 | 1.12E-06 |
| ENSMUST00000173581.7,Tnfaip2-008,nonsense_mediated_decay,ENSMUSG00000021281.15,protein_coding | 245.1228292 | 0.71091454 | 3.81E-08 | 1.13E-06 |
| ENSMUST00000000291.8,Mnt-001,protein_coding,ENSMUSG00000000282.12,protein_coding | 4708.641458 | 0.346326947 | 3.83E-08 | 1.13E-06 |
| ENSMUST00000015456.8,Gadd45b-201,protein_coding,ENSMUSG00000015312.8,protein_coding | 1477.170816 | -0.436313414 | 3.85E-08 | 1.14E-06 |
| ENSMUST00000109579.8,Mkl1-002,protein_coding,ENSMUSG00000042292.16,protein_coding | 1319.055516 | -0.476370613 | 3.86E-08 | 1.14E-06 |
| ENSMUST00000136448.7,Ift80-002,retained_intron,ENSMUSG00000027778.15,protein_coding | 184.2066435 | -0.679198798 | 3.87E-08 | 1.14E-06 |
| ENSMUST00000107214.8,Psip1-003,protein_coding,ENSMUSG00000028484.16,protein_coding | 991.1327202 | 0.750926402 | 3.93E-08 | 1.16E-06 |
| ENSMUST00000068044.13,Slc39a14-001,protein_coding,ENSMUSG00000022094.15,protein_coding | 11636.91686 | 0.272439156 | 3.94E-08 | 1.16E-06 |
| ENSMUST00000139897.7,Ybx1-003,processed_transcript,ENSMUSG00000028639.14,protein_coding | 9371.675232 | 0.35729606 | 3.94E-08 | 1.16E-06 |
| ENSMUST00000163516.7,Nedd4l-202,protein_coding,ENSMUSG00000024589.15,protein_coding | 7983.250255 | 0.191832454 | 3.95E-08 | 1.16E-06 |
| ENSMUST00000065793.11,Phgdh-001,protein_coding,ENSMUSG00000053398.11,protein_coding | 115713.7556 | 0.270002542 | 3.99E-08 | 1.17E-06 |
| ENSMUST00000153837.7,Inpp5b-004,protein_coding,ENSMUSG00000028894.18,protein_coding | 88.35544948 | 0.881552588 | 4.00E-08 | 1.18E-06 |
| ENSMUST00000152704.7,Btf3-002,protein_coding,ENSMUSG00000021660.14,protein_coding | 51159.91672 | 0.212812932 | 4.03E-08 | 1.19E-06 |
| ENSMUST00000160420.7,9930104L06Rik-002,retained_intron,ENSMUSG00000044730.15,processed_transcript | 268.36091 | -0.545899614 | 4.11E-08 | 1.21E-06 |
| ENSMUST00000101055.4,Ccnk-201,protein_coding,ENSMUSG00000021258.9,protein_coding | 11481.0846 | 0.178650146 | 4.19E-08 | 1.23E-06 |
| ENSMUST00000156503.1,Uqcrq-004,processed_transcript,ENSMUSG00000044894.14,protein_coding | 1338.724806 | 0.351435751 | 4.23E-08 | 1.24E-06 |
| ENSMUST00000000500.6,Pdgfb-201,protein_coding,ENSMUSG00000000489.6,protein_coding | 314.0100238 | -0.588166985 | 4.28E-08 | 1.26E-06 |
| ENSMUST00000168977.7,Ylpm1-004,protein_coding,ENSMUSG00000021244.15,protein_coding | 1560.212673 | 0.373802093 | 4.39E-08 | 1.29E-06 |
| ENSMUST00000045898.3,Pcyt1b-001,protein_coding,ENSMUSG00000035246.16,protein_coding | 2146.306357 | -0.302088801 | 4.39E-08 | 1.29E-06 |
| ENSMUST00000058011.7,Mcm2-001,protein_coding,ENSMUSG00000002870.8,protein_coding | 72364.50284 | 0.256627394 | 4.39E-08 | 1.29E-06 |

| | | | | |
|---|---|---|---|---|
| ENSMUST00000154032.1,Spryd3-001,protein_coding,ENSMUSG00000036966.15,protein_coding | 6675.944994 | 0.265582793 | 4.41E-08 | 1.29E-06 |
| ENSMUST00000060311.11,Hip1-001,protein_coding,ENSMUSG00000039959.13,protein_coding | 19145.52679 | -0.192712431 | 4.42E-08 | 1.29E-06 |
| ENSMUST00000025818.6,Rin1-201,protein_coding,ENSMUSG00000024883.6,protein_coding | 12290.85953 | 0.230540827 | 4.43E-08 | 1.29E-06 |
| ENSMUST00000031081.10,Pi4k2b-001,protein_coding,ENSMUSG00000029186.12,protein_coding | 8930.263231 | 0.299508212 | 4.44E-08 | 1.30E-06 |
| ENSMUST00000199406.4,Anapc5-007,protein_coding,ENSMUSG00000029472.13,protein_coding | 17886.46541 | 0.256196033 | 4.52E-08 | 1.32E-06 |
| ENSMUST00000092912.11,Mettl16-004,nonsense_mediated_decay,ENSMUSG00000010554.14,protein_coding | 876.2104317 | 0.542790109 | 4.52E-08 | 1.32E-06 |
| ENSMUST00000023019.11,Trmu-001,protein_coding,ENSMUSG00000022386.12,protein_coding | 1254.616722 | 0.428800789 | 4.53E-08 | 1.32E-06 |
| ENSMUST00000027940.5,Tmem206-001,protein_coding,ENSMUSG00000026627.7,protein_coding | 1995.98221 | 0.303981536 | 4.53E-08 | 1.32E-06 |
| ENSMUST00000074628.12,Wbp2-001,protein_coding,ENSMUSG00000034341.17,protein_coding | 16538.79739 | 0.208552766 | 4.55E-08 | 1.32E-06 |
| ENSMUST00000138243.1,Map3k10-002,protein_coding,ENSMUSG00000040390.13,protein_coding | 1941.093715 | 0.281365965 | 4.61E-08 | 1.34E-06 |
| ENSMUST00000001304.7,Ckb-201,protein_coding,ENSMUSG00000001270.8,protein_coding | 2625.269508 | -0.500129501 | 4.71E-08 | 1.37E-06 |
| ENSMUST00000029845.14,Ddah1-001,protein_coding,ENSMUSG00000028194.15,protein_coding | 3731.402997 | -0.304410198 | 4.72E-08 | 1.37E-06 |
| ENSMUST00000114754.7,Mapk14-003,protein_coding,ENSMUSG00000053436.14,protein_coding | 161.3624993 | 1.016368706 | 4.72E-08 | 1.37E-06 |
| ENSMUST00000199854.4,Sh3glb1-005,protein_coding,ENSMUSG00000037062.13,protein_coding | 16052.94597 | 0.188363374 | 4.73E-08 | 1.37E-06 |
| ENSMUST00000079464.12,Tuba4a-002,protein_coding,ENSMUSG00000026202.13,protein_coding | 2964.994467 | -0.351264984 | 4.74E-08 | 1.37E-06 |
| ENSMUST00000041110.11,Ttc7-001,protein_coding,ENSMUSG00000036918.15,protein_coding | 3518.649649 | 0.298687457 | 4.76E-08 | 1.38E-06 |
| ENSMUST00000020099.11,Cdk1-001,protein_coding,ENSMUSG00000019942.12,protein_coding | 1122.708964 | -0.386907434 | 4.79E-08 | 1.39E-06 |
| ENSMUST00000102814.4,Rps6-001,protein_coding,ENSMUSG00000028495.14,protein_coding | 130936.7206 | 0.248313745 | 4.81E-08 | 1.39E-06 |
| ENSMUST00000021379.7,Gemin2-001,protein_coding,ENSMUSG00000060121.15,protein_coding | 3140.761907 | 0.277767065 | 4.81E-08 | 1.39E-06 |
| ENSMUST00000030742.10,Mecr-001,protein_coding,ENSMUSG00000028910.12,protein_coding | 3752.369329 | 0.303820168 | 4.83E-08 | 1.39E-06 |
| ENSMUST00000202604.1,Sdad1-003,retained_intron,ENSMUSG00000029415.4,protein_coding | 442.5491574 | 0.736031881 | 4.83E-08 | 1.39E-06 |
| ENSMUST00000039318.14,Cdc14b-201,protein_coding,ENSMUSG00000033102.14,protein_coding | 3189.559114 | -0.411667712 | 4.89E-08 | 1.41E-06 |
| ENSMUST00000115026.7,Cald1-015,protein_coding,ENSMUSG00000029761.16,protein_coding | 37009.13435 | -0.359775961 | 4.92E-08 | 1.42E-06 |
| ENSMUST00000026011.6,Sfxn2-201,protein_coding,ENSMUSG00000025036.6,protein_coding | 1486.497982 | -0.36296984 | 4.93E-08 | 1.42E-06 |
| ENSMUST00000022693.7,Bmp1-201,protein_coding,ENSMUSG00000022098.9,protein_coding | 7851.880133 | -0.230476033 | 4.94E-08 | 1.42E-06 |
| ENSMUST00000030183.9,Car9-001,protein_coding,ENSMUSG00000028463.14,protein_coding | 237.6449211 | 0.657043354 | 4.97E-08 | 1.43E-06 |
| ENSMUST00000102992.9,Zc3hc1-001,protein_coding,ENSMUSG00000039130.18,protein_coding | 138.1109003 | 0.76721959 | 5.01E-08 | 1.44E-06 |
| ENSMUST00000079817.7,Stk24-201,protein_coding,ENSMUSG00000063410.7,protein_coding | 17102.44591 | 0.217063912 | 5.03E-08 | 1.44E-06 |
| ENSMUST00000036540.11,Fam117b-001,protein_coding,ENSMUSG00000041040.12,protein_coding | 7374.848873 | 0.195187813 | 5.07E-08 | 1.46E-06 |
| ENSMUST00000002080.11,Pdzd4-001,protein_coding,ENSMUSG00000002006.12,protein_coding | 2084.368884 | -0.28215486 | 5.09E-08 | 1.46E-06 |
| ENSMUST00000047356.10,Atf5-001,protein_coding,ENSMUSG00000038539.15,protein_coding | 15577.93903 | -0.235984408 | 5.12E-08 | 1.47E-06 |
| ENSMUST00000058237.13,Ugt1a7c-001,protein_coding,ENSMUSG00000090124.7,protein_coding | 6938.646962 | 0.368523053 | 5.15E-08 | 1.47E-06 |
| ENSMUST00000192638.5,Ildr2-007,protein_coding,ENSMUSG00000040612.13,protein_coding | 287.1688083 | -0.743686526 | 5.15E-08 | 1.47E-06 |
| ENSMUST00000063857.10,Slc6a9-003,protein_coding,ENSMUSG00000028542.17,protein_coding | 6344.501034 | 0.679185954 | 5.20E-08 | 1.49E-06 |
| ENSMUST00000074963.8,Bbs5-001,protein_coding,ENSMUSG00000063145.10,protein_coding | 713.7004127 | -0.49980685 | 5.30E-08 | 1.52E-06 |
| ENSMUST00000051396.12,Soat1-003,protein_coding,ENSMUSG00000026600.12,protein_coding | 19806.82915 | 0.216830404 | 5.32E-08 | 1.52E-06 |
| ENSMUST00000071430.5,2310039H08Rik-201,protein_coding,ENSMUSG00000062619.5,protein_coding | 1995.231089 | 0.278914727 | 5.35E-08 | 1.53E-06 |
| ENSMUST00000036453.13,Map3k10-001,protein_coding,ENSMUSG00000040390.13,protein_coding | 1046.910214 | -0.368212742 | 5.38E-08 | 1.53E-06 |
| ENSMUST00000119730.7,Ncald-003,protein_coding,ENSMUSG00000051359.14,protein_coding | 195.8451098 | -0.854114406 | 5.39E-08 | 1.54E-06 |
| ENSMUST00000124828.7,Fbf1-003,protein_coding,ENSMUSG00000020776.18,protein_coding | 428.7135619 | 0.727409806 | 5.40E-08 | 1.54E-06 |
| ENSMUST00000135941.7,Pfdn2-001,protein_coding,ENSMUSG00000006412.10,protein_coding | 4917.155575 | -0.203493446 | 5.42E-08 | 1.54E-06 |
| ENSMUST00000017851.3,Serinc3-001,protein_coding,ENSMUSG00000017707.9,protein_coding | 14785.57828 | 0.23107698 | 5.43E-08 | 1.55E-06 |
| ENSMUST00000079041.5,Slc35e3-201,protein_coding,ENSMUSG00000060181.5,protein_coding | 2215.650397 | -0.313787327 | 5.50E-08 | 1.56E-06 |
| ENSMUST00000036004.15,Hnrnpa1-201,protein_coding,ENSMUSG00000046434.15,protein_coding | 10735.07051 | -0.402818838 | 5.52E-08 | 1.57E-06 |
| ENSMUST00000105352.1,Adamtsl5-201,protein_coding,ENSMUSG00000043822.18,protein_coding | 422.4300621 | 0.753770241 | 5.56E-08 | 1.58E-06 |
| ENSMUST00000136579.2,Drap1-010,protein_coding,ENSMUSG00000024914.16,protein_coding | 7776.4293 | 0.260608656 | 5.60E-08 | 1.59E-06 |
| ENSMUST00000008036.8,Rplp1-001,protein_coding,ENSMUSG00000007892.8,protein_coding | 142314.7798 | 0.262316425 | 5.61E-08 | 1.59E-06 |
| ENSMUST00000005600.5,Rfx1-201,protein_coding,ENSMUSG00000031706.7,protein_coding | 4264.182262 | 0.184672389 | 5.68E-08 | 1.61E-06 |
| ENSMUST00000130606.7,Capza2-007,nonsense_mediated_decay,ENSMUSG00000015733.13,protein_coding | 1891.767134 | -0.39927601 | 5.88E-08 | 1.67E-06 |
| ENSMUST00000102845.10,Rps27a-003,protein_coding,ENSMUSG00000020460.15,protein_coding | 47115.73104 | 0.226541607 | 5.89E-08 | 1.67E-06 |
| ENSMUST00000205428.1,Kcnn4-003,protein_coding,ENSMUSG00000054342.8,protein_coding | 2310.190275 | 0.30180453 | 5.91E-08 | 1.67E-06 |
| ENSMUST00000172254.2,Iars-002,protein_coding,ENSMUSG00000037851.13,protein_coding | 243.4622569 | 0.593382001 | 5.92E-08 | 1.67E-06 |
| ENSMUST00000045557.9,Slc7a5-001,protein_coding,ENSMUSG00000040010.10,protein_coding | 65679.0612 | 0.363746592 | 5.95E-08 | 1.68E-06 |
| ENSMUST00000148460.7,Bloc1s6-002,processed_transcript,ENSMUSG00000005804.14,protein_coding | 2877.946943 | 0.217964087 | 6.02E-08 | 1.70E-06 |
| ENSMUST00000110115.8,Asap1-203,protein_coding,ENSMUSG00000022377.16,protein_coding | 17960.91449 | -0.281353773 | 6.04E-08 | 1.70E-06 |
| ENSMUST00000050125.8,2900026A02Rik-001,protein_coding,ENSMUSG00000051339.10,protein_coding | 962.5636841 | -0.549671283 | 6.06E-08 | 1.71E-06 |
| ENSMUST00000034094.10,Gins3-001,protein_coding,ENSMUSG00000031669.10,protein_coding | 1215.399965 | -0.347644171 | 6.06E-08 | 1.71E-06 |
| ENSMUST00000105361.9,Dazap1-002,protein_coding,ENSMUSG00000069565.12,protein_coding | 7747.942606 | 0.336728503 | 6.07E-08 | 1.71E-06 |
| ENSMUST00000095263.9,Yipf3-001,protein_coding,ENSMUSG00000071074.9,protein_coding | 4230.25132 | 0.294436757 | 6.12E-08 | 1.72E-06 |
| ENSMUST00000070597.12,Retsat-001,protein_coding,ENSMUSG00000056666.13,protein_coding | 4358.070533 | 0.269799606 | 6.14E-08 | 1.73E-06 |
| ENSMUST00000198834.1,Fbrsl1-003,protein_coding,ENSMUSG00000043323.16,protein_coding | 612.4338934 | 0.533860994 | 6.22E-08 | 1.75E-06 |
| ENSMUST00000179543.7,Noc2l-201,protein_coding,ENSMUSG00000095567.7,protein_coding | 42163.19724 | 0.231054105 | 6.22E-08 | 1.75E-06 |
| ENSMUST00000182961.1,Arhgef40-009,retained_intron,ENSMUSG00000004562.16,protein_coding | 105.0377585 | -0.836678435 | 6.29E-08 | 1.77E-06 |
| ENSMUST00000038169.6,Mif-201,protein_coding,ENSMUSG00000033307.6,protein_coding | 73090.9384 | 0.238458782 | 6.30E-08 | 1.77E-06 |
| ENSMUST00000122088.1,S1pr5-001,protein_coding,ENSMUSG00000045087.8,protein_coding | 194.7592225 | -0.609072892 | 6.31E-08 | 1.77E-06 |
| ENSMUST00000030028.4,Erp44-001,protein_coding,ENSMUSG00000028343.10,protein_coding | 5643.504449 | 0.255213915 | 6.42E-08 | 1.80E-06 |
| ENSMUST00000114884.7,Rbm33-003,protein_coding,ENSMUSG00000048271.14,protein_coding | 9134.467859 | 0.260040423 | 6.43E-08 | 1.80E-06 |
| ENSMUST00000160853.7,Maf1-002,protein_coding,ENSMUSG00000022553.15,protein_coding | 1867.141325 | 0.363200724 | 6.51E-08 | 1.82E-06 |
| ENSMUST00000106488.2,Eef2k-003,protein_coding,ENSMUSG00000035064.17,protein_coding | 1116.184424 | 0.641324674 | 6.61E-08 | 1.85E-06 |
| ENSMUST00000119222.7,Mtx1-002,protein_coding,ENSMUSG00000064068.13,protein_coding | 475.506897 | -0.448127434 | 6.62E-08 | 1.85E-06 |
| ENSMUST00000057208.12,Mrps9-002,protein_coding,ENSMUSG00000060679.14,protein_coding | 6693.440466 | 0.210671316 | 6.64E-08 | 1.85E-06 |
| ENSMUST00000072503.12,Rpl18-201,protein_coding,ENSMUSG00000059070.16,protein_coding | 85676.29783 | 0.25166867 | 6.67E-08 | 1.86E-06 |
| ENSMUST00000117646.7,Dixdc1-002,protein_coding,ENSMUSG00000032064.17,protein_coding | 611.8135719 | -0.51102016 | 6.69E-08 | 1.86E-06 |
| ENSMUST00000153073.7,Rad51c-004,nonsense_mediated_decay,ENSMUSG00000007646.13,protein_coding | 1632.551443 | 0.33210881 | 6.68E-08 | 1.86E-06 |
| ENSMUST00000095476.5,Lig4-001,protein_coding,ENSMUSG00000049717.9,protein_coding | 681.582104 | -0.465228915 | 6.77E-08 | 1.89E-06 |
| ENSMUST00000108883.9,Anxa6-001,protein_coding,ENSMUSG00000018340.13,protein_coding | 9736.033619 | 0.390712458 | 6.80E-08 | 1.89E-06 |
| ENSMUST00000020831.12,Fam114a2-001,protein_coding,ENSMUSG00000020523.14,protein_coding | 5991.230424 | 0.246982429 | 6.80E-08 | 1.89E-06 |
| ENSMUST00000145846.1,Vmp1-002,processed_transcript,ENSMUSG00000018171.9,protein_coding | 1774.04721 | -0.294725011 | 6.82E-08 | 1.89E-06 |
| ENSMUST00000031530.8,Sppl3-001,protein_coding,ENSMUSG00000029550.11,protein_coding | 8235.688405 | 0.155150385 | 6.89E-08 | 1.91E-06 |
| ENSMUST00000170935.7,Mon2-203,protein_coding,ENSMUSG00000034602.13,protein_coding | 3066.611782 | 0.402219131 | 6.91E-08 | 1.92E-06 |
| ENSMUST00000114389.9,Naa10-002,protein_coding,ENSMUSG00000031388.14,protein_coding | 4242.322271 | 0.429212073 | 6.93E-08 | 1.92E-06 |
| ENSMUST00000097420.5,Rnaset2a-201,protein_coding,ENSMUSG00000095687.1,protein_coding | 3490.059692 | 0.295214279 | 7.03E-08 | 1.95E-06 |
| ENSMUST00000200271.4,Ttc14-002,protein_coding,ENSMUSG00000027677.17,protein_coding | 150.3974514 | -0.834451305 | 7.08E-08 | 1.96E-06 |
| ENSMUST00000154550.7,Use1-003,retained_intron,ENSMUSG00000002395.14,protein_coding | 598.4648595 | 0.466954437 | 7.27E-08 | 2.01E-06 |
| ENSMUST00000032151.2,Eif4e3-001,protein_coding,ENSMUSG00000093661.1,protein_coding | 304.6003885 | 0.506963074 | 7.28E-08 | 2.01E-06 |
| ENSMUST00000101270.4,Tbc1d9b-002,protein_coding,ENSMUSG00000036644.15,protein_coding | 2055.920642 | 0.384725822 | 7.32E-08 | 2.02E-06 |
| ENSMUST00000052832.5,2410015M20Rik-201,protein_coding,ENSMUSG00000049760.5,protein_coding | 4490.188138 | 0.25398998 | 7.33E-08 | 2.02E-06 |
| ENSMUST00000033683.7,Rps4x-001,protein_coding,ENSMUSG00000031320.9,protein_coding | 119386.3878 | 0.19745427 | 7.37E-08 | 2.04E-06 |
| ENSMUST00000023100.6,Srebf2-201,protein_coding,ENSMUSG00000022463.7,protein_coding | 66578.30526 | 0.243364587 | 7.40E-08 | 2.04E-06 |
| ENSMUST00000020504.5,Hint1-001,protein_coding,ENSMUSG00000020267.6,protein_coding | 40337.30565 | 0.264063722 | 7.43E-08 | 2.05E-06 |
| ENSMUST00000136983.7,Dpf2-001,protein_coding,ENSMUSG00000024826.14,protein_coding | 3240.551265 | 0.373456111 | 7.45E-08 | 2.05E-06 |
| ENSMUST00000036456.6,Cep72-201,protein_coding,ENSMUSG00000021572.8,protein_coding | 1108.456933 | 0.304352629 | 7.48E-08 | 2.06E-06 |

| | | | | |
|---|---|---|---|---|
| ENSMUST00000033099.5,Fgf21-001,protein_coding,ENSMUSG00000030827.5,protein_coding | 332.3016011 | 0.745935638 | 7.56E-08 | 2.08E-06 |
| ENSMUST00000043074.11,Fau-201,protein_coding,ENSMUSG00000038274.11,protein_coding | 63288.25998 | 0.245358316 | 7.58E-08 | 2.09E-06 |
| ENSMUST00000111468.9,Mtch2-004,nonsense_mediated_decay,ENSMUSG00000027282.17,protein_coding | 9051.10711 | 0.235567467 | 7.61E-08 | 2.09E-06 |
| ENSMUST00000168665.2,2300009A05Rik-001,protein_coding,ENSMUSG00000032403.8,protein_coding | 541.6629469 | 0.399907964 | 7.62E-08 | 2.09E-06 |
| ENSMUST00000035092.6,Myd88-001,protein_coding,ENSMUSG00000032508.8,protein_coding | 2444.567436 | 0.314990073 | 7.62E-08 | 2.09E-06 |
| ENSMUST00000022834.6,Cma1-201,protein_coding,ENSMUSG00000022225.6,protein_coding | 154.4504947 | 0.671346839 | 7.67E-08 | 2.10E-06 |
| ENSMUST00000076354.12,Tspan7-001,protein_coding,ENSMUSG00000058254.12,protein_coding | 1265.507952 | 0.286655757 | 7.75E-08 | 2.12E-06 |
| ENSMUST00000108834.2,4930438A08Rik-001,protein_coding,ENSMUSG00000069873.4,protein_coding | 65.80995433 | 0.956615126 | 7.87E-08 | 2.16E-06 |
| ENSMUST00000028835.12,Crls1-001,protein_coding,ENSMUSG00000027357.16,protein_coding | 5655.892744 | 0.302721298 | 7.91E-08 | 2.16E-06 |
| ENSMUST00000194746.5,Rpl31-002,protein_coding,ENSMUSG00000073702.11,protein_coding | 11370.98074 | 0.185584731 | 7.98E-08 | 2.19E-06 |
| ENSMUST00000102995.8,Ahcyl2-002,protein_coding,ENSMUSG00000029772.17,protein_coding | 5637.240019 | 0.292935753 | 8.00E-08 | 2.19E-06 |
| ENSMUST00000043409.7,Zfp503-001,protein_coding,ENSMUSG00000039081.9,protein_coding | 1672.538904 | -0.473253453 | 8.04E-08 | 2.20E-06 |
| ENSMUST00000030931.10,Pank4-001,protein_coding,ENSMUSG00000029056.13,protein_coding | 3127.349873 | -0.257647295 | 8.24E-08 | 2.25E-06 |
| ENSMUST00000045747.4,Wnt4-001,protein_coding,ENSMUSG00000036856.4,protein_coding | 322.5696031 | -0.626326864 | 8.26E-08 | 2.26E-06 |
| ENSMUST00000108840.3,Tcf25-003,protein_coding,ENSMUSG00000001472.17,protein_coding | 14869.87409 | 0.270895239 | 8.33E-08 | 2.27E-06 |
| ENSMUST00000143987.7,Ndufv2-001,protein_coding,ENSMUSG00000024099.14,protein_coding | 4283.485278 | 0.247955844 | 8.40E-08 | 2.29E-06 |
| ENSMUST00000179818.2,Atxn2l-004,protein_coding,ENSMUSG00000032637.15,protein_coding | 723.4593131 | 0.525868607 | 8.59E-08 | 2.34E-06 |
| ENSMUST00000113827.1,Brwd1-003,protein_coding,ENSMUSG00000022914.15,protein_coding | 672.6602214 | -0.364681827 | 8.63E-08 | 2.35E-06 |
| ENSMUST00000034234.14,Coq9-001,protein_coding,ENSMUSG00000031782.15,protein_coding | 5669.950755 | 0.25776392 | 8.65E-08 | 2.35E-06 |
| ENSMUST00000049863.11,Pofut1-001,protein_coding,ENSMUSG00000046020.13,protein_coding | 6321.335692 | 0.218757242 | 8.72E-08 | 2.37E-06 |
| ENSMUST00000040440.6,Calm2-001,protein_coding,ENSMUSG00000036438.12,protein_coding | 34076.27405 | -0.298359864 | 8.73E-08 | 2.37E-06 |
| ENSMUST00000216325.1,Mrpl4-008,processed_transcript,ENSMUSG00000003299.10,protein_coding | 67.57720553 | 0.922931013 | 8.75E-08 | 2.38E-06 |
| ENSMUST00000127637.7,Napa-002,nonsense_mediated_decay,ENSMUSG00000006024.13,protein_coding | 308.2642487 | 0.658469232 | 8.79E-08 | 2.39E-06 |
| ENSMUST00000179343.2,Purb-001,protein_coding,ENSMUSG00000094483.2,protein_coding | 16700.14563 | -0.468430844 | 9.04E-08 | 2.45E-06 |
| ENSMUST00000112498.2,Crim1-201,protein_coding,ENSMUSG00000024074.7,protein_coding | 11181.80802 | -0.338626098 | 9.04E-08 | 2.45E-06 |
| ENSMUST00000000449.8,Mkrn2-001,protein_coding,ENSMUSG00000000439.9,protein_coding | 7557.054913 | 0.282281324 | 9.09E-08 | 2.46E-06 |
| ENSMUST00000099270.4,Thbd-001,protein_coding,ENSMUSG00000074743.4,protein_coding | 6651.519842 | 0.434171101 | 9.14E-08 | 2.47E-06 |
| ENSMUST00000038343.6,B4galnt2-001,protein_coding,ENSMUSG00000013418.8,protein_coding | 163.0993618 | 0.812093101 | 9.16E-08 | 2.48E-06 |
| ENSMUST00000102724.4,Faf1-001,protein_coding,ENSMUSG00000010517.7,protein_coding | 13200.23687 | 0.297676101 | 9.25E-08 | 2.50E-06 |
| ENSMUST00000036074.14,Gmip-001,protein_coding,ENSMUSG00000036246.14,protein_coding | 2853.878648 | -0.362818475 | 9.27E-08 | 2.50E-06 |
| ENSMUST00000194192.2,Dst-020,protein_coding,ENSMUSG00000026131.18,protein_coding | 2536.575352 | -0.311701719 | 9.31E-08 | 2.51E-06 |
| ENSMUST00000099582.2,Fars2-202,protein_coding,ENSMUSG00000021420.12,protein_coding | 55.3149459 | 0.886913652 | 9.33E-08 | 2.52E-06 |
| ENSMUST00000110143.7,Mef2b-003,protein_coding,ENSMUSG00000079033.9,protein_coding | 294.738121 | 0.708107589 | 9.45E-08 | 2.55E-06 |
| ENSMUST00000162529.7,Sidt2-007,protein_coding,ENSMUSG00000034908.16,protein_coding | 102.7129214 | 0.789543559 | 9.51E-08 | 2.56E-06 |
| ENSMUST00000134362.7,Pdk1-007,processed_transcript,ENSMUSG00000006494.11,protein_coding | 779.0353392 | 0.619420501 | 9.57E-08 | 2.58E-06 |
| ENSMUST00000210449.1,Nucb1-005,protein_coding,ENSMUSG00000030824.17,protein_coding | 2184.910451 | 0.354132977 | 9.58E-08 | 2.58E-06 |
| ENSMUST00000154308.1,Ide-002,retained_intron,ENSMUSG00000056999.14,protein_coding | 1682.233473 | 0.42404698 | 9.59E-08 | 2.58E-06 |
| ENSMUST00000044116.13,Spg20-001,protein_coding,ENSMUSG00000036580.15,protein_coding | 7746.57941 | 0.396257321 | 9.61E-08 | 2.58E-06 |
| ENSMUST00000114227.7,Gng12-003,protein_coding,ENSMUSG00000036402.13,protein_coding | 289.5654269 | 0.738712997 | 9.63E-08 | 2.59E-06 |
| ENSMUST00000141365.2,Vim-003,protein_coding,ENSMUSG00000026728.9,protein_coding | 4004.784546 | 0.346673718 | 9.69E-08 | 2.60E-06 |
| ENSMUST00000051395.8,Prr11-001,protein_coding,ENSMUSG00000020493.8,protein_coding | 15247.0051 | -0.173138896 | 9.70E-08 | 2.60E-06 |
| ENSMUST00000089874.8,Mcc-001,protein_coding,ENSMUSG00000071856.10,protein_coding | 6755.158478 | -0.186711497 | 9.76E-08 | 2.62E-06 |
| ENSMUST00000086458.9,Fbxl12-003,protein_coding,ENSMUSG00000066892.14,protein_coding | 546.9169694 | 0.511694356 | 9.78E-08 | 2.62E-06 |
| ENSMUST00000145058.1,Cbx4-002,processed_transcript,ENSMUSG00000039989.7,protein_coding | 2345.841571 | 0.361586977 | 9.77E-08 | 2.62E-06 |
| ENSMUST00000056177.6,Pacsin2-201,protein_coding,ENSMUSG00000016664.14,protein_coding | 7014.802909 | 0.27026752 | 9.80E-08 | 2.62E-06 |
| ENSMUST00000070951.6,Plpp1-202,protein_coding,ENSMUSG00000021759.13,protein_coding | 1533.008432 | -0.384517866 | 9.93E-08 | 2.66E-06 |
| ENSMUST00000070368.7,Siah2-001,protein_coding,ENSMUSG00000036432.8,protein_coding | 4690.551519 | 0.408517091 | 9.94E-08 | 2.66E-06 |
| ENSMUST00000035657.7,Tssc1-201,protein_coding,ENSMUSG00000036613.7,protein_coding | 1529.529863 | -0.360086576 | 9.97E-08 | 2.66E-06 |
| ENSMUST00000162036.7,Rsrc1-001,protein_coding,ENSMUSG00000034544.17,protein_coding | 978.0874667 | 0.343937289 | 1.00E-07 | 2.67E-06 |
| ENSMUST00000033578.6,Magee1-001,protein_coding,ENSMUSG00000031227.8,protein_coding | 1923.347375 | -0.32986509 | 1.00E-07 | 2.67E-06 |
| ENSMUST00000060918.10,Amz1-001,protein_coding,ENSMUSG00000050022.16,protein_coding | 129.5285682 | 0.779319434 | 1.00E-07 | 2.68E-06 |
| ENSMUST00000050544.7,Has2-001,protein_coding,ENSMUSG00000022367.7,protein_coding | 1496.119151 | -0.591683186 | 1.00E-07 | 2.68E-06 |
| ENSMUST00000139129.8,Srsf1-001,protein_coding,ENSMUSG00000018379.17,protein_coding | 6704.862787 | -0.592901552 | 1.01E-07 | 2.70E-06 |
| ENSMUST00000169377.7,Map3k12-001,protein_coding,ENSMUSG00000023050.8,protein_coding | 3315.299259 | -0.298714135 | 1.02E-07 | 2.71E-06 |
| ENSMUST00000101375.4,Npm1-003,protein_coding,ENSMUSG00000057113.13,protein_coding | 11317.05973 | -0.955429383 | 1.02E-07 | 2.71E-06 |
| ENSMUST00000033995.13,Rbpms-003,protein_coding,ENSMUSG00000031586.16,protein_coding | 1253.980356 | 0.352407298 | 1.02E-07 | 2.72E-06 |
| ENSMUST00000028257.2,Gca-001,protein_coding,ENSMUSG00000026893.4,protein_coding | 105.526128 | -0.704197762 | 1.02E-07 | 2.72E-06 |
| ENSMUST00000107485.7,Msl1-004,protein_coding,ENSMUSG00000052915.14,protein_coding | 2443.412782 | -0.261211315 | 1.02E-07 | 2.72E-06 |
| ENSMUST00000027271.8,Inpp1-001,protein_coding,ENSMUSG00000026102.9,protein_coding | 1350.735832 | -0.413033625 | 1.03E-07 | 2.73E-06 |
| ENSMUST00000028831.14,Mcm8-001,protein_coding,ENSMUSG00000027353.14,protein_coding | 3476.645906 | 0.295245793 | 1.04E-07 | 2.75E-06 |
| ENSMUST00000108814.7,Rtel1-007,protein_coding,ENSMUSG00000038685.18,protein_coding | 1411.626066 | -0.54545477 | 1.05E-07 | 2.78E-06 |
| ENSMUST00000072055.12,Chka-001,protein_coding,ENSMUSG00000024843.15,protein_coding | 1298.923368 | 0.398868766 | 1.11E-07 | 2.93E-06 |
| ENSMUST00000093456.11,Kxd1-001,protein_coding,ENSMUSG00000055553.16,protein_coding | 3781.778014 | 0.272119569 | 1.13E-07 | 3.00E-06 |
| ENSMUST00000140653.1,D5Ertd579e-005,protein_coding,ENSMUSG00000029190.19,protein_coding | 3159.389809 | 0.253192276 | 1.16E-07 | 3.06E-06 |
| ENSMUST00000106640.1,Parva-004,protein_coding,ENSMUSG00000030770.15,protein_coding | 1423.972345 | 0.513289733 | 1.16E-07 | 3.06E-06 |
| ENSMUST00000027961.11,Hspa14-001,protein_coding,ENSMUSG00000109865.1,protein_coding | 11593.92621 | 0.236913182 | 1.16E-07 | 3.06E-06 |
| ENSMUST00000070877.5,Amacr-201,protein_coding,ENSMUSG00000022244.6,protein_coding | 2204.454929 | 0.342750454 | 1.16E-07 | 3.06E-06 |
| ENSMUST00000040888.11,Krtcap2-001,protein_coding,ENSMUSG00000042747.12,protein_coding | 4870.104981 | 0.287237569 | 1.16E-07 | 3.07E-06 |
| ENSMUST00000063445.12,Klhdc1-001,protein_coding,ENSMUSG00000051890.13,protein_coding | 434.2911387 | 0.503700259 | 1.16E-07 | 3.07E-06 |
| ENSMUST00000038027.5,Gpi1-001,protein_coding,ENSMUSG00000036427.5,protein_coding | 59968.38112 | 0.228981123 | 1.17E-07 | 3.08E-06 |
| ENSMUST00000051777.14,C2cd3-001,protein_coding,ENSMUSG00000047248.20,protein_coding | 5000.856008 | -0.186715939 | 1.17E-07 | 3.08E-06 |
| ENSMUST00000032888.9,Arl6ip1-001,protein_coding,ENSMUSG00000030654.9,protein_coding | 24441.1344 | 0.179765802 | 1.17E-07 | 3.09E-06 |
| ENSMUST00000159333.1,Stk19-007,protein_coding,ENSMUSG00000061207.11,protein_coding | 117.6642895 | -0.795547594 | 1.18E-07 | 3.10E-06 |
| ENSMUST00000045174.5,Ypel5-201,protein_coding,ENSMUSG00000039770.5,protein_coding | 2270.223628 | 0.471767402 | 1.19E-07 | 3.13E-06 |
| ENSMUST00000047783.13,Rpgrip1l-001,protein_coding,ENSMUSG00000033282.14,protein_coding | 872.0689947 | -0.531631014 | 1.19E-07 | 3.13E-06 |
| ENSMUST00000089959.6,Gch1-201,protein_coding,ENSMUSG00000037580.9,protein_coding | 2252.780047 | 0.399093431 | 1.19E-07 | 3.13E-06 |
| ENSMUST00000113695.7,Tpm1-016,protein_coding,ENSMUSG00000032366.15,protein_coding | 24426.87936 | 0.234343753 | 1.20E-07 | 3.14E-06 |
| ENSMUST00000109381.7,Brd1-001,protein_coding,ENSMUSG00000022387.16,protein_coding | 12122.3255 | 0.250054517 | 1.20E-07 | 3.14E-06 |
| ENSMUST00000174819.7,Tmem234-014,nonsense_mediated_decay,ENSMUSG00000028797.20,protein_coding | 385.637227 | 0.583609845 | 1.20E-07 | 3.15E-06 |
| ENSMUST00000023776.12,Slc4a8-001,protein_coding,ENSMUSG00000023032.12,protein_coding | 165.7796081 | -0.659767186 | 1.20E-07 | 3.15E-06 |
| ENSMUST00000063905.11,Clip1-003,protein_coding,ENSMUSG00000049550.17,protein_coding | 125.1257558 | 0.98711394 | 1.21E-07 | 3.17E-06 |
| ENSMUST00000176274.1,Cyhr1-003,protein_coding,ENSMUSG00000053929.16,protein_coding | 9740.530634 | -0.239749847 | 1.21E-07 | 3.17E-06 |
| ENSMUST00000031411.14,Aldh2-001,protein_coding,ENSMUSG00000029455.14,protein_coding | 7096.368145 | 0.287784536 | 1.21E-07 | 3.18E-06 |
| ENSMUST00000109764.7,Nfix-001,protein_coding,ENSMUSG00000001911.16,protein_coding | 18453.40928 | -0.099057125 | 1.22E-07 | 3.20E-06 |
| ENSMUST00000076894.10,Rfwd2-001,protein_coding,ENSMUSG00000040782.12,protein_coding | 5474.061299 | 0.183952958 | 1.22E-07 | 3.20E-06 |
| ENSMUST00000089402.9,Porcn-004,protein_coding,ENSMUSG00000031169.13,protein_coding | 835.5101134 | 0.451953248 | 1.23E-07 | 3.21E-06 |
| ENSMUST00000184142.7,Plac9b-001,protein_coding,ENSMUSG00000072674.10,protein_coding | 197.5102653 | 0.610604412 | 1.24E-07 | 3.24E-06 |
| ENSMUST00000095017.10,Dmtf1-002,protein_coding,ENSMUSG00000042508.16,protein_coding | 1792.301694 | -0.438876802 | 1.24E-07 | 3.24E-06 |
| ENSMUST00000119870.8,Cngb1-001,protein_coding,ENSMUSG00000031789.17,protein_coding | 292.907257 | 0.567832696 | 1.24E-07 | 3.24E-06 |
| ENSMUST00000127996.7,Foxred1-001,protein_coding,ENSMUSG00000039048.15,protein_coding | 2253.635999 | 0.25654547 | 1.25E-07 | 3.27E-06 |
| ENSMUST00000034463.3,Arv1-001,protein_coding,ENSMUSG00000031982.5,protein_coding | 537.2510534 | -0.384123484 | 1.26E-07 | 3.27E-06 |
| ENSMUST00000076698.11,Sypl-001,protein_coding,ENSMUSG00000020570.14,protein_coding | 2066.961932 | -0.410739185 | 1.27E-07 | 3.30E-06 |

| | | | | |
|---|---|---|---|---|
| ENSMUST00000053288.5,Cdr2l-001,protein_coding,ENSMUSG00000050910.5,protein_coding | 21623.53122 | 0.236220022 | 1.28E-07 | 3.32E-06 |
| ENSMUST00000155707.2,Timp2-002,protein_coding,ENSMUSG00000017466.9,protein_coding | 4349.624178 | 0.270602819 | 1.28E-07 | 3.32E-06 |
| ENSMUST00000010506.9,Rdm1-001,protein_coding,ENSMUSG00000010362.9,protein_coding | 1362.066001 | 0.463878096 | 1.28E-07 | 3.33E-06 |
| ENSMUST00000103070.2,Tex2-002,protein_coding,ENSMUSG00000040548.16,protein_coding | 249.1355359 | 0.69085383 | 1.29E-07 | 3.34E-06 |
| ENSMUST00000022087.6,Nsun2-002,protein_coding,ENSMUSG00000021595.18,protein_coding | 470.2160118 | -0.497039522 | 1.29E-07 | 3.34E-06 |
| ENSMUST00000206598.1,Slc3a2-008,protein_coding,ENSMUSG00000010095.13,protein_coding | 9683.30368 | 0.349540287 | 1.29E-07 | 3.34E-06 |
| ENSMUST00000071103.8,Ddx1-201,protein_coding,ENSMUSG00000037149.9,protein_coding | 24195.46241 | 0.137331842 | 1.31E-07 | 3.38E-06 |
| ENSMUST00000068408.13,Rps15-001,protein_coding,ENSMUSG00000063457.14,protein_coding | 110302.4145 | 0.22578461 | 1.31E-07 | 3.39E-06 |
| ENSMUST00000044255.12,Ggta1-201,protein_coding,ENSMUSG00000035778.17,protein_coding | 119.8752413 | 0.943572258 | 1.32E-07 | 3.41E-06 |
| ENSMUST00000048824.8,Gm5617-001,protein_coding,ENSMUSG00000042293.8,protein_coding | 307.9079174 | 0.49996542 | 1.33E-07 | 3.43E-06 |
| ENSMUST00000105290.8,Nr2c1-001,protein_coding,ENSMUSG00000005897.14,protein_coding | 1606.800016 | 0.448556396 | 1.35E-07 | 3.50E-06 |
| ENSMUST00000127010.1,Gm12758-001,antisense,ENSMUSG00000085105.2,antisense | 279.9852109 | 0.564180916 | 1.36E-07 | 3.51E-06 |
| ENSMUST00000132253.4,Ost4-001,protein_coding,ENSMUSG00000038803.7,protein_coding | 2534.469972 | -0.289024398 | 1.36E-07 | 3.52E-06 |
| ENSMUST00000002855.13,Kdelr1-001,protein_coding,ENSMUSG00000002778.14,protein_coding | 12115.37869 | 0.20534625 | 1.36E-07 | 3.52E-06 |
| ENSMUST00000054607.15,Ahcy-001,protein_coding,ENSMUSG00000027597.15,protein_coding | 60751.41597 | 0.236797771 | 1.37E-07 | 3.52E-06 |
| ENSMUST00000162007.7,Stau2-004,protein_coding,ENSMUSG00000025920.19,protein_coding | 1105.90597 | 0.410372978 | 1.38E-07 | 3.55E-06 |
| ENSMUST00000047973.3,Dhcr24-001,protein_coding,ENSMUSG00000034926.3,protein_coding | 24501.29489 | 0.245099774 | 1.38E-07 | 3.55E-06 |
| ENSMUST00000107583.2,Cisd3-001,protein_coding,ENSMUSG00000078695.8,protein_coding | 3494.467402 | -0.389087638 | 1.39E-07 | 3.57E-06 |
| ENSMUST00000005964.6,Adh5-001,protein_coding,ENSMUSG00000028138.8,protein_coding | 15357.67763 | 0.291987034 | 1.39E-07 | 3.58E-06 |
| ENSMUST00000034590.3,Tagln-001,protein_coding,ENSMUSG00000032085.5,protein_coding | 2464.352251 | -0.377075035 | 1.40E-07 | 3.60E-06 |
| ENSMUST00000062686.10,Map2k7-002,protein_coding,ENSMUSG00000002948.19,protein_coding | 5854.155576 | 0.317036937 | 1.42E-07 | 3.65E-06 |
| ENSMUST00000110103.1,Gdf15-002,protein_coding,ENSMUSG00000038508.7,protein_coding | 2541.938964 | 0.702879392 | 1.43E-07 | 3.67E-06 |
| ENSMUST00000005592.6,Siglecg-001,protein_coding,ENSMUSG00000030468.12,protein_coding | 530.5610666 | -0.551333738 | 1.43E-07 | 3.67E-06 |
| ENSMUST00000169609.1,Sc5d-201,protein_coding,ENSMUSG00000032018.14,protein_coding | 4002.487451 | 0.275014621 | 1.44E-07 | 3.70E-06 |
| ENSMUST00000134910.1,Prkca-003,processed_transcript,ENSMUSG00000050965.14,protein_coding | 569.6199485 | 0.40122718 | 1.45E-07 | 3.72E-06 |
| ENSMUST00000115098.6,Kcnh2-002,protein_coding,ENSMUSG00000038319.14,protein_coding | 1906.313541 | -0.274627397 | 1.46E-07 | 3.73E-06 |
| ENSMUST00000102483.4,Rpl23a-001,protein_coding,ENSMUSG00000058546.8,protein_coding | 137763.3589 | 0.173443311 | 1.46E-07 | 3.75E-06 |
| ENSMUST00000166409.5,Klf3-201,protein_coding,ENSMUSG00000029178.14,protein_coding | 1084.695489 | 0.543407095 | 1.48E-07 | 3.78E-06 |
| ENSMUST00000099194.3,Tspyl3-001,protein_coding,ENSMUSG00000074671.6,protein_coding | 796.651651 | -0.336676303 | 1.48E-07 | 3.78E-06 |
| ENSMUST00000140129.1,Srm-003,processed_transcript,ENSMUSG00000006442.10,protein_coding | 66.48002289 | 0.947287108 | 1.50E-07 | 3.83E-06 |
| ENSMUST00000109627.7,Cbx6-001,protein_coding,ENSMUSG00000089715.11,protein_coding | 14875.73052 | -0.189312022 | 1.50E-07 | 3.84E-06 |
| ENSMUST00000049470.10,Tmem183a-001,protein_coding,ENSMUSG00000042305.12,protein_coding | 7728.832675 | 0.280526508 | 1.52E-07 | 3.87E-06 |
| ENSMUST00000050467.7,Tob2-201,protein_coding,ENSMUSG00000048546.7,protein_coding | 11636.68189 | 0.191799265 | 1.52E-07 | 3.89E-06 |
| ENSMUST00000114776.2,Pias2-201,protein_coding,ENSMUSG00000025423.15,protein_coding | 2691.165509 | 0.246190175 | 1.54E-07 | 3.91E-06 |
| ENSMUST00000074085.11,Rpl10-002,protein_coding,ENSMUSG00000008682.13,protein_coding | 11740.98811 | -0.210631993 | 1.54E-07 | 3.91E-06 |
| ENSMUST00000028728.5,Ubr1-001,protein_coding,ENSMUSG00000027272.5,protein_coding | 6707.303241 | -0.312892118 | 1.54E-07 | 3.91E-06 |
| ENSMUST00000037618.12,Rbpj-001,protein_coding,ENSMUSG00000039191.12,protein_coding | 1926.226245 | 0.510294394 | 1.56E-07 | 3.98E-06 |
| ENSMUST00000053024.6,Pgp-201,protein_coding,ENSMUSG00000043445.6,protein_coding | 4997.718037 | -0.273057837 | 1.56E-07 | 3.98E-06 |
| ENSMUST00000094549.10,Szrd1-002,processed_transcript,ENSMUSG00000040842.17,protein_coding | 7610.124824 | 0.261141858 | 1.57E-07 | 3.98E-06 |
| ENSMUST00000005066.8,Map2k1-001,protein_coding,ENSMUSG00000004936.8,protein_coding | 12106.9335 | 0.229251765 | 1.57E-07 | 3.98E-06 |
| ENSMUST00000001724.11,Ddx18-001,protein_coding,ENSMUSG00000001674.11,protein_coding | 22448.91536 | 0.248123414 | 1.58E-07 | 4.02E-06 |
| ENSMUST00000113102.9,Naaa-001,protein_coding,ENSMUSG00000029413.14,protein_coding | 1376.152863 | 0.419515883 | 1.58E-07 | 4.02E-06 |
| ENSMUST00000172699.1,Mex3a-001,protein_coding,ENSMUSG00000074480.4,protein_coding | 9703.477822 | -0.171828708 | 1.59E-07 | 4.04E-06 |
| ENSMUST00000069025.13,Srsf11-010,protein_coding,ENSMUSG00000055436.18,protein_coding | 2132.946615 | -0.265247441 | 1.61E-07 | 4.08E-06 |
| ENSMUST00000107131.1,Cdkn2a-002,protein_coding,ENSMUSG00000044303.6,protein_coding | 52256.61514 | 0.226389777 | 1.62E-07 | 4.10E-06 |
| ENSMUST00000034913.4,Fam195b-001,protein_coding,ENSMUSG00000061111.8,protein_coding | 6172.257515 | 0.261975436 | 1.63E-07 | 4.12E-06 |
| ENSMUST00000136510.2,Cstf2-006,protein_coding,ENSMUSG00000031256.11,protein_coding | 77.41260091 | 0.935576847 | 1.63E-07 | 4.12E-06 |
| ENSMUST00000055327.7,Aqp3-001,protein_coding,ENSMUSG00000028435.8,protein_coding | 609.6072252 | 0.54328755 | 1.64E-07 | 4.14E-06 |
| ENSMUST00000081848.12,Fdps-001,protein_coding,ENSMUSG00000059743.12,protein_coding | 24236.22287 | 0.29734812 | 1.65E-07 | 4.18E-06 |
| ENSMUST00000109921.8,Dbn1-009,protein_coding,ENSMUSG00000034675.17,protein_coding | 7086.742878 | -0.248084169 | 1.66E-07 | 4.18E-06 |
| ENSMUST00000102758.7,Vdac1-001,protein_coding,ENSMUSG00000020402.11,protein_coding | 58403.85798 | 0.24355875 | 1.66E-07 | 4.19E-06 |
| ENSMUST00000063761.7,Cpt1c-001,protein_coding,ENSMUSG00000007783.10,protein_coding | 1932.168282 | -0.27788602 | 1.66E-07 | 4.20E-06 |
| ENSMUST00000172912.7,H2-K1-003,protein_coding,ENSMUSG00000061232.15,protein_coding | 890.7854727 | 0.743894091 | 1.67E-07 | 4.22E-06 |
| ENSMUST00000046212.1,Slc16a1-001,protein_coding,ENSMUSG00000032902.1,protein_coding | 20205.8977 | 0.289453413 | 1.68E-07 | 4.23E-06 |
| ENSMUST00000109614.8,Chd9-201,protein_coding,ENSMUSG00000056608.14,protein_coding | 1748.339617 | -0.610480465 | 1.72E-07 | 4.32E-06 |
| ENSMUST00000079510.5,Cherp-003,protein_coding,ENSMUSG00000052488.7,protein_coding | 10246.35267 | 0.353509739 | 1.74E-07 | 4.37E-06 |
| ENSMUST00000217537.1,Gtf3c6-001,protein_coding,ENSMUSG00000019837.8,protein_coding | 1306.446154 | -0.45342829 | 1.75E-07 | 4.40E-06 |
| ENSMUST00000198254.4,Sh3glb1-003,protein_coding,ENSMUSG00000037062.13,protein_coding | 9666.77239 | -0.423097755 | 1.77E-07 | 4.45E-06 |
| ENSMUST00000113051.8,Shroom3-005,protein_coding,ENSMUSG00000029381.15,protein_coding | 1611.103464 | -0.422471703 | 1.78E-07 | 4.48E-06 |
| ENSMUST00000100467.9,Trrap-001,protein_coding,ENSMUSG00000045482.16,protein_coding | 34056.89145 | -0.131431201 | 1.79E-07 | 4.50E-06 |
| ENSMUST00000217352.1,Rpsa-006,nonsense_mediated_decay,ENSMUSG00000032518.6,protein_coding | 1693.9503 | 0.394585109 | 1.80E-07 | 4.53E-06 |
| ENSMUST00000002979.14,Lamb1-001,protein_coding,ENSMUSG00000002900.15,protein_coding | 32338.8489 | 0.161521731 | 1.83E-07 | 4.60E-06 |
| ENSMUST00000028232.6,Phf19-001,protein_coding,ENSMUSG00000026873.9,protein_coding | 2069.266346 | 0.273736496 | 1.84E-07 | 4.60E-06 |
| ENSMUST00000112173.7,Tspan9-001,protein_coding,ENSMUSG00000030352.14,protein_coding | 7160.366782 | 0.223104117 | 1.84E-07 | 4.62E-06 |
| ENSMUST00000205070.2,Isy1-007,nonsense_mediated_decay,ENSMUSG00000030056.8,protein_coding | 568.8602343 | 0.584873584 | 1.86E-07 | 4.66E-06 |
| ENSMUST00000036125.9,Eif4h-002,protein_coding,ENSMUSG00000040731.13,protein_coding | 53000.33162 | 0.230471009 | 1.87E-07 | 4.68E-06 |
| ENSMUST00000025760.12,Chka-002,protein_coding,ENSMUSG00000024843.15,protein_coding | 1569.091464 | 0.4274028 | 1.89E-07 | 4.72E-06 |
| ENSMUST00000162503.7,Adck5-006,protein_coding,ENSMUSG00000022550.17,protein_coding | 2193.951285 | 0.247569353 | 1.89E-07 | 4.73E-06 |
| ENSMUST00000031817.9,Herc6-001,protein_coding,ENSMUSG00000029798.11,protein_coding | 844.2830225 | -0.372631897 | 1.90E-07 | 4.76E-06 |
| ENSMUST00000032169.7,Abtb1-001,protein_coding,ENSMUSG00000030083.11,protein_coding | 924.976001 | 0.393015617 | 1.91E-07 | 4.78E-06 |
| ENSMUST00000059295.9,Syt12-001,protein_coding,ENSMUSG00000049303.10,protein_coding | 1534.586487 | -0.349526321 | 1.92E-07 | 4.78E-06 |
| ENSMUST00000033096.15,Nucb1-012,protein_coding,ENSMUSG00000030824.17,protein_coding | 15872.58676 | 0.244671747 | 1.93E-07 | 4.81E-06 |
| ENSMUST00000049020.8,Irgq-001,protein_coding,ENSMUSG00000041037.8,protein_coding | 1300.740597 | -0.300702748 | 1.94E-07 | 4.82E-06 |
| ENSMUST00000027517.13,Dgkd-001,protein_coding,ENSMUSG00000070738.10,protein_coding | 16091.04075 | 0.377528876 | 1.94E-07 | 4.84E-06 |
| ENSMUST00000118316.7,N4bp2l2-001,protein_coding,ENSMUSG00000029655.17,protein_coding | 2311.46493 | -0.454261598 | 1.94E-07 | 4.84E-06 |
| ENSMUST00000153867.7,Mcm7-004,protein_coding,ENSMUSG00000029730.16,protein_coding | 3904.810396 | 0.373667842 | 1.95E-07 | 4.86E-06 |
| ENSMUST00000001675.13,Stk38l-001,protein_coding,ENSMUSG00000001630.13,protein_coding | 1415.472512 | -0.31930755 | 1.96E-07 | 4.88E-06 |
| ENSMUST00000173015.7,March2-006,protein_coding,ENSMUSG00000079557.10,protein_coding | 331.8333752 | 0.773545254 | 1.99E-07 | 4.93E-06 |
| ENSMUST00000114882.7,Rps10-001,protein_coding,ENSMUSG00000052146.15,protein_coding | 97162.38046 | 0.25152997 | 2.00E-07 | 4.98E-06 |
| ENSMUST00000195571.1,Cenpl-010,nonsense_mediated_decay,ENSMUSG00000026708.16,protein_coding | 190.4560505 | -0.752495588 | 2.01E-07 | 4.99E-06 |
| ENSMUST00000044442.8,Ptk7-201,protein_coding,ENSMUSG00000023972.9,protein_coding | 11815.11328 | -0.232937563 | 2.01E-07 | 4.99E-06 |
| ENSMUST00000107894.7,Dgke-001,protein_coding,ENSMUSG00000000276.11,protein_coding | 815.8529819 | -0.360336446 | 2.02E-07 | 4.99E-06 |
| ENSMUST00000136008.7,Ergic2-001,protein_coding,ENSMUSG00000030304.11,protein_coding | 3197.879031 | -0.376828261 | 2.02E-07 | 5.00E-06 |
| ENSMUST00000046869.10,Fbxw4-001,protein_coding,ENSMUSG00000040913.10,protein_coding | 3393.312857 | 0.286402914 | 2.03E-07 | 5.03E-06 |
| ENSMUST00000023426.11,Cldnd1-001,protein_coding,ENSMUSG00000022744.12,protein_coding | 9004.452231 | 0.257776776 | 2.03E-07 | 5.03E-06 |
| ENSMUST00000028250.8,Mrrf-001,protein_coding,ENSMUSG00000026887.9,protein_coding | 3697.763813 | 0.285466261 | 2.04E-07 | 5.05E-06 |
| ENSMUST00000137724.7,Clcn6-001,protein_coding,ENSMUSG00000029016.13,protein_coding | 2181.696757 | 0.289226621 | 2.04E-07 | 5.05E-06 |
| ENSMUST00000114842.7,Anks1-001,protein_coding,ENSMUSG00000024219.15,protein_coding | 3610.285271 | 0.228900341 | 2.05E-07 | 5.07E-06 |
| ENSMUST00000054776.3,Plekhf2-001,protein_coding,ENSMUSG00000049969.3,protein_coding | 2746.979872 | 0.379231202 | 2.10E-07 | 5.19E-06 |
| ENSMUST00000084299.5,Akirin2-001,protein_coding,ENSMUSG00000028291.7,protein_coding | 8382.881759 | 0.210660248 | 2.13E-07 | 5.25E-06 |
| ENSMUST00000096744.7,Myadm-002,protein_coding,ENSMUSG00000068566.12,protein_coding | 10589.84951 | 0.404213664 | 2.14E-07 | 5.29E-06 |
| ENSMUST00000215300.1,Higd1a-005,protein_coding,ENSMUSG00000038412.8,protein_coding | 2612.664031 | 0.284578139 | 2.15E-07 | 5.29E-06 |

| | | | | |
|---|---|---|---|---|
| ENSMUST00000030404.4,Ppie-001,protein_coding,ENSMUSG00000028651.12,protein_coding | 5425.093525 | 0.257584765 | 2.16E-07 | 5.31E-06 |
| ENSMUST00000145409.1,Mapk1-007,processed_transcript,ENSMUSG00000063358.15,protein_coding | 899.4755824 | 0.334912757 | 2.16E-07 | 5.31E-06 |
| ENSMUST00000031262.8,Coq2-001,protein_coding,ENSMUSG00000029319.8,protein_coding | 5673.079221 | 0.251560481 | 2.16E-07 | 5.32E-06 |
| ENSMUST00000048794.6,Hoxa5-001,protein_coding,ENSMUSG00000038253.6,protein_coding | 924.4770199 | -0.421416098 | 2.17E-07 | 5.33E-06 |
| ENSMUST00000051720.5,Sh3tc2-201,protein_coding,ENSMUSG00000045629.7,protein_coding | 6802.405969 | 0.430174552 | 2.17E-07 | 5.34E-06 |
| ENSMUST00000017548.6,Rpl19-001,protein_coding,ENSMUSG00000017404.12,protein_coding | 185392.9097 | 0.257596753 | 2.18E-07 | 5.34E-06 |
| ENSMUST00000020864.8,Pctp-001,protein_coding,ENSMUSG00000020553.8,protein_coding | 1043.153235 | 0.275486492 | 2.19E-07 | 5.38E-06 |
| ENSMUST00000152941.1,Gm4724-001,protein_coding,ENSMUSG00000078897.10,protein_coding | 307.7300652 | -0.857398045 | 2.20E-07 | 5.39E-06 |
| ENSMUST00000163595.2,Cdyl-202,protein_coding,ENSMUSG00000059288.13,protein_coding | 3468.186928 | 0.306735143 | 2.21E-07 | 5.41E-06 |
| ENSMUST00000201359.3,Tbc1d24-005,nonsense_mediated_decay,ENSMUSG00000036473.15,protein_coding | 182.7958037 | -0.966126352 | 2.22E-07 | 5.45E-06 |
| ENSMUST00000133603.1,Nup188-005,retained_intron,ENSMUSG00000052533.14,protein_coding | 59.01178938 | -0.966587533 | 2.24E-07 | 5.48E-06 |
| ENSMUST00000150923.7,Exosc8-003,retained_intron,ENSMUSG00000027752.15,protein_coding | 513.9587694 | -0.47380932 | 2.24E-07 | 5.49E-06 |
| ENSMUST00000154375.7,Tspan9-008,protein_coding,ENSMUSG00000030352.14,protein_coding | 63.69203803 | 0.905797263 | 2.25E-07 | 5.50E-06 |
| ENSMUST00000153269.1,Mboat1-002,processed_transcript,ENSMUSG00000038732.14,protein_coding | 218.0841407 | 0.74605704 | 2.26E-07 | 5.52E-06 |
| ENSMUST00000171797.1,Nap1l1-202,protein_coding,ENSMUSG00000058799.14,protein_coding | 21907.60558 | -0.401408315 | 2.28E-07 | 5.56E-06 |
| ENSMUST00000022925.8,Eif3h-201,protein_coding,ENSMUSG00000022312.10,protein_coding | 71321.93243 | 0.245868413 | 2.28E-07 | 5.56E-06 |
| ENSMUST00000080813.4,Rps17-001,protein_coding,ENSMUSG00000061787.15,protein_coding | 43734.25215 | 0.155518767 | 2.29E-07 | 5.60E-06 |
| ENSMUST00000145976.1,Ybx1-007,processed_transcript,ENSMUSG00000028639.14,protein_coding | 429.5491752 | 0.450151757 | 2.31E-07 | 5.63E-06 |
| ENSMUST00000028332.7,Dpp7-001,protein_coding,ENSMUSG00000026958.13,protein_coding | 727.1614079 | -0.35343874 | 2.31E-07 | 5.64E-06 |
| ENSMUST00000176204.7,Peg10-001,protein_coding,ENSMUSG00000092035.8,protein_coding | 14597.77421 | -0.359542094 | 2.32E-07 | 5.65E-06 |
| ENSMUST00000102824.3,Ifit1-001,protein_coding,ENSMUSG00000034459.8,protein_coding | 1397.623632 | 0.340748779 | 2.33E-07 | 5.67E-06 |
| ENSMUST00000011055.6,Apip-001,protein_coding,ENSMUSG00000010911.12,protein_coding | 2264.366266 | 0.237081842 | 2.34E-07 | 5.69E-06 |
| ENSMUST00000189208.1,Ralgps2-012,retained_intron,ENSMUSG00000026594.14,protein_coding | 457.0831937 | 0.482629199 | 2.35E-07 | 5.71E-06 |
| ENSMUST00000162183.7,Alg12-001,protein_coding,ENSMUSG00000035845.17,protein_coding | 1695.259921 | 0.332173193 | 2.35E-07 | 5.71E-06 |
| ENSMUST00000205191.1,Bet1-006,processed_transcript,ENSMUSG00000032757.6,protein_coding | 82.51339931 | 0.900538914 | 2.39E-07 | 5.81E-06 |
| ENSMUST00000039559.8,Thbs1-001,protein_coding,ENSMUSG00000040152.8,protein_coding | 10791.08205 | -0.44539226 | 2.40E-07 | 5.84E-06 |
| ENSMUST00000153735.7,Pdlim2-002,protein_coding,ENSMUSG00000022090.10,protein_coding | 2062.547645 | 0.262380121 | 2.41E-07 | 5.86E-06 |
| ENSMUST00000119398.9,Fgfr1-003,protein_coding,ENSMUSG00000031565.18,protein_coding | 5531.86085 | 0.301758074 | 2.42E-07 | 5.88E-06 |
| ENSMUST00000026547.8,Tubgcp2-001,protein_coding,ENSMUSG00000025474.9,protein_coding | 4450.984207 | 0.320183465 | 2.43E-07 | 5.89E-06 |
| ENSMUST00000204701.2,Ppp6c-003,protein_coding,ENSMUSG00000026753.5,protein_coding | 2976.268567 | 0.281505693 | 2.43E-07 | 5.90E-06 |
| ENSMUST00000217356.1,Rpsa-002,nonsense_mediated_decay,ENSMUSG00000032518.6,protein_coding | 177662.1616 | 0.23708381 | 2.46E-07 | 5.96E-06 |
| ENSMUST00000003038.11,Ap2a2-001,protein_coding,ENSMUSG00000002957.11,protein_coding | 44922.91352 | 0.196282157 | 2.47E-07 | 5.97E-06 |
| ENSMUST00000054034.6,Polq-002,protein_coding,ENSMUSG00000034206.15,protein_coding | 1470.824931 | -0.715526159 | 2.48E-07 | 6.00E-06 |
| ENSMUST00000069652.7,Rab3gap2-001,protein_coding,ENSMUSG00000039318.12,protein_coding | 1751.922465 | -0.291086362 | 2.49E-07 | 6.01E-06 |
| ENSMUST00000000365.2,Mcts1-001,protein_coding,ENSMUSG00000000355.13,protein_coding | 1103.054148 | -0.404184108 | 2.53E-07 | 6.12E-06 |
| ENSMUST00000109574.7,Epb41l1-002,protein_coding,ENSMUSG00000027624.17,protein_coding | 3157.599697 | -0.244293974 | 2.54E-07 | 6.13E-06 |
| ENSMUST00000139457.1,Aurkb-006,processed_transcript,ENSMUSG00000020897.12,protein_coding | 121.1170486 | 0.716237618 | 2.55E-07 | 6.14E-06 |
| ENSMUST00000101018.9,Cep170b-202,protein_coding,ENSMUSG00000072825.10,protein_coding | 18357.25718 | 0.176860216 | 2.55E-07 | 6.15E-06 |
| ENSMUST00000109415.1,Bod1-002,protein_coding,ENSMUSG00000044502.16,protein_coding | 184.4638758 | 0.702696871 | 2.57E-07 | 6.20E-06 |
| ENSMUST00000124755.1,Slc25a39-004,protein_coding,ENSMUSG00000018677.9,protein_coding | 613.3093086 | 0.738211218 | 2.57E-07 | 6.20E-06 |
| ENSMUST00000052404.12,Epb41l5-003,protein_coding,ENSMUSG00000026383.14,protein_coding | 485.4418409 | -0.377659879 | 2.60E-07 | 6.26E-06 |
| ENSMUST00000113312.8,Shq1-001,protein_coding,ENSMUSG00000035378.17,protein_coding | 462.8486302 | -0.44838889 | 2.62E-07 | 6.31E-06 |
| ENSMUST00000115395.9,Carm1-003,protein_coding,ENSMUSG00000032185.15,protein_coding | 6236.222606 | 0.315682289 | 2.65E-07 | 6.36E-06 |
| ENSMUST00000005017.14,Hdgf-001,protein_coding,ENSMUSG00000004897.16,protein_coding | 96726.98549 | 0.1930383 | 2.65E-07 | 6.38E-06 |
| ENSMUST00000124738.1,Gm12992-002,processed_transcript,ENSMUSG00000085667.1,processed_transcript | 1129.627472 | 0.348802036 | 2.67E-07 | 6.42E-06 |
| ENSMUST00000109412.8,Cpeb4-001,protein_coding,ENSMUSG00000020300.14,protein_coding | 811.2231893 | -0.70038741 | 2.68E-07 | 6.43E-06 |
| ENSMUST00000120339.7,5730559C18Rik-001,protein_coding,ENSMUSG00000041605.16,protein_coding | 1080.614321 | -0.282806435 | 2.70E-07 | 6.47E-06 |
| ENSMUST00000022519.14,Anxa8-001,protein_coding,ENSMUSG00000021950.15,protein_coding | 6016.956005 | 0.329037896 | 2.72E-07 | 6.53E-06 |
| ENSMUST00000106763.7,Bptf-001,protein_coding,ENSMUSG00000040481.17,protein_coding | 6924.67664 | -0.301743551 | 2.73E-07 | 6.54E-06 |
| ENSMUST00000038193.14,Wdr59-001,protein_coding,ENSMUSG00000031959.15,protein_coding | 2977.705334 | -0.308033231 | 2.73E-07 | 6.55E-06 |
| ENSMUST00000106479.7,Ipp-201,protein_coding,ENSMUSG00000028696.12,protein_coding | 1947.793366 | 0.223351818 | 2.77E-07 | 6.64E-06 |
| ENSMUST00000045730.6,Akap12-001,protein_coding,ENSMUSG00000038587.9,protein_coding | 8914.052958 | -0.347647379 | 2.78E-07 | 6.64E-06 |
| ENSMUST00000045562.5,Cox15-001,protein_coding,ENSMUSG00000040018.9,protein_coding | 6018.388593 | -0.154880022 | 2.80E-07 | 6.69E-06 |
| ENSMUST00000110452.1,Hist1h2bj-001,protein_coding,ENSMUSG00000069300.3,protein_coding | 534.3039548 | 0.411169363 | 2.81E-07 | 6.72E-06 |
| ENSMUST00000032373.11,Ldhb-001,protein_coding,ENSMUSG00000030246.11,protein_coding | 1873.756604 | -0.30297816 | 2.82E-07 | 6.73E-06 |
| ENSMUST00000160629.7,Cpsf4-002,protein_coding,ENSMUSG00000029625.16,protein_coding | 2834.527133 | 0.328008523 | 2.85E-07 | 6.80E-06 |
| ENSMUST00000128553.7,Zfp512b-002,protein_coding,ENSMUSG00000000823.16,protein_coding | 2490.287303 | -0.371072251 | 2.87E-07 | 6.85E-06 |
| ENSMUST00000020759.11,Efemp1-001,protein_coding,ENSMUSG00000020467.15,protein_coding | 16089.16046 | 0.217360055 | 2.89E-07 | 6.90E-06 |
| ENSMUST00000016680.13,Cul4a-001,protein_coding,ENSMUSG00000031446.14,protein_coding | 17287.95714 | 0.158013387 | 2.90E-07 | 6.90E-06 |
| ENSMUST00000022623.12,Trim35-001,protein_coding,ENSMUSG00000022043.18,protein_coding | 23615.91598 | 0.286891874 | 2.90E-07 | 6.92E-06 |
| ENSMUST00000040344.6,Gns-201,protein_coding,ENSMUSG00000034707.6,protein_coding | 14586.99099 | 0.24548208 | 2.91E-07 | 6.94E-06 |
| ENSMUST00000058351.15,Pgm2-001,protein_coding,ENSMUSG00000025791.18,protein_coding | 22403.49836 | 0.258056697 | 2.92E-07 | 6.94E-06 |
| ENSMUST00000053558.9,Ankrd37-001,protein_coding,ENSMUSG00000050914.16,protein_coding | 229.852061 | 0.573090383 | 2.96E-07 | 7.03E-06 |
| ENSMUST00000002845.6,Mea1-201,protein_coding,ENSMUSG00000002768.7,protein_coding | 8564.742514 | 0.259474016 | 2.98E-07 | 7.08E-06 |
| ENSMUST00000094783.6,Pcdh7-002,protein_coding,ENSMUSG00000029108.14,protein_coding | 866.5029919 | -0.47676081 | 3.00E-07 | 7.13E-06 |
| ENSMUST00000154256.2,Zfp703-001,protein_coding,ENSMUSG00000085795.8,protein_coding | 4625.015205 | 0.192182594 | 3.01E-07 | 7.14E-06 |
| ENSMUST00000029877.8,Decr1-001,protein_coding,ENSMUSG00000028223.8,protein_coding | 5720.873628 | 0.234614363 | 3.03E-07 | 7.19E-06 |
| ENSMUST00000160197.5,Exoc6b-001,protein_coding,ENSMUSG00000033769.16,protein_coding | 8454.006578 | 0.195211771 | 3.04E-07 | 7.21E-06 |
| ENSMUST00000180667.1,Mirt1-202,lincRNA,ENSMUSG00000097636.7,lincRNA | 189.6084063 | -0.671074795 | 3.08E-07 | 7.29E-06 |
| ENSMUST00000020149.5,Ikbip-001,protein_coding,ENSMUSG00000019975.12,protein_coding | 2368.075267 | -0.359735189 | 3.09E-07 | 7.32E-06 |
| ENSMUST00000040021.11,Ptpn23-001,protein_coding,ENSMUSG00000036057.11,protein_coding | 15879.20562 | 0.185305712 | 3.11E-07 | 7.36E-06 |
| ENSMUST00000137868.7,Speg-003,protein_coding,ENSMUSG00000026207.16,protein_coding | 4726.825917 | -0.181794734 | 3.14E-07 | 7.43E-06 |
| ENSMUST00000025419.7,Ppic-201,protein_coding,ENSMUSG00000024538.7,protein_coding | 6239.787467 | -0.192534301 | 3.14E-07 | 7.43E-06 |
| ENSMUST00000093381.10,Ccdc157-001,protein_coding,ENSMUSG00000051427.14,protein_coding | 1556.759131 | -0.306562672 | 3.15E-07 | 7.44E-06 |
| ENSMUST00000122182.1,Cbx5-003,protein_coding,ENSMUSG00000009575.14,protein_coding | 1684.282203 | 0.417237673 | 3.15E-07 | 7.44E-06 |
| ENSMUST00000123994.2,Slc6a9-011,protein_coding,ENSMUSG00000028542.17,protein_coding | 35.04841412 | 0.95422927 | 3.19E-07 | 7.52E-06 |
| ENSMUST00000132504.1,Got1-004,retained_intron,ENSMUSG00000025190.13,protein_coding | 186.5725561 | 0.775579742 | 3.20E-07 | 7.56E-06 |
| ENSMUST00000145911.3,Bcat1-007,processed_transcript,ENSMUSG00000030268.17,protein_coding | 2859.616692 | 0.267013832 | 3.21E-07 | 7.58E-06 |
| ENSMUST00000066489.12,P3h4-001,protein_coding,ENSMUSG00000006931.15,protein_coding | 11568.46364 | -0.247380933 | 3.23E-07 | 7.63E-06 |
| ENSMUST00000071250.12,Pygl-001,protein_coding,ENSMUSG00000021069.16,protein_coding | 968.3484719 | 0.333223248 | 3.24E-07 | 7.63E-06 |
| ENSMUST00000197451.1,1500002C15Rik-201,antisense,ENSMUSG00000096221.3,antisense | 606.8843154 | 0.662060581 | 3.27E-07 | 7.69E-06 |
| ENSMUST00000201536.1,Gm43268-001,TEC,ENSMUSG00000107182.1,TEC | 49.12436381 | -0.888319169 | 3.30E-07 | 7.76E-06 |
| ENSMUST00000150033.7,Fam53a-003,nonsense_mediated_decay,ENSMUSG00000037339.17,protein_coding | 400.0512555 | 0.634984279 | 3.32E-07 | 7.81E-06 |
| ENSMUST00000169453.7,Nfat5-202,protein_coding,ENSMUSG00000003847.16,protein_coding | 4813.468821 | -0.516189637 | 3.32E-07 | 7.81E-06 |
| ENSMUST00000090031.11,Gng5-001,protein_coding,ENSMUSG00000068523.12,protein_coding | 12286.82836 | 0.180846658 | 3.35E-07 | 7.86E-06 |
| ENSMUST00000028854.14,Mal-001,protein_coding,ENSMUSG00000027375.14,protein_coding | 1230.043874 | 0.487949949 | 3.35E-07 | 7.87E-06 |
| ENSMUST00000162237.1,Sf1-015,protein_coding,ENSMUSG00000024949.17,protein_coding | 1078.563817 | 0.40690764 | 3.35E-07 | 7.87E-06 |
| ENSMUST00000205262.1,Arhgap17-002,protein_coding,ENSMUSG00000030766.15,protein_coding | 255.5617744 | 0.901497364 | 3.40E-07 | 7.98E-06 |
| ENSMUST00000006235.7,Ctsb-201,protein_coding,ENSMUSG00000021939.7,protein_coding | 39272.04108 | 0.183298714 | 3.40E-07 | 7.98E-06 |
| ENSMUST00000177034.1,Mtcl1-011,protein_coding,ENSMUSG00000052105.16,protein_coding | 136.1692428 | 0.719920632 | 3.41E-07 | 7.98E-06 |
| ENSMUST00000208252.1,Arglu1-003,retained_intron,ENSMUSG00000040459.11,protein_coding | 4059.854005 | -0.444604596 | 3.41E-07 | 7.99E-06 |
| ENSMUST00000074077.11,Bmp4-001,protein_coding,ENSMUSG00000021835.14,protein_coding | 384.0990666 | -0.545287235 | 3.42E-07 | 8.00E-06 |

| | | | | |
|---|---|---|---|---|
| ENSMUST00000021728.10,Siva1-201,protein_coding,ENSMUSG00000064326.12,protein_coding | 4898.199022 | 0.252965523 | 3.43E-07 | 8.03E-06 |
| ENSMUST00000108719.3,Llgl1-002,protein_coding,ENSMUSG00000020536.12,protein_coding | 8699.40665 | 0.237641313 | 3.46E-07 | 8.10E-06 |
| ENSMUST00000099145.5,Dlgap4-003,protein_coding,ENSMUSG00000061689.15,protein_coding | 2674.493418 | -0.186439114 | 3.48E-07 | 8.12E-06 |
| ENSMUST00000197357.1,Ube2d3-014,processed_transcript,ENSMUSG00000078578.9,protein_coding | 1122.071029 | 0.319690017 | 3.48E-07 | 8.12E-06 |
| ENSMUST00000038552.12,Coro7-001,protein_coding,ENSMUSG00000039637.14,protein_coding | 2144.587602 | 0.393697621 | 3.48E-07 | 8.12E-06 |
| ENSMUST00000192436.5,Tiprl-004,nonsense_mediated_decay,ENSMUSG00000040843.10,protein_coding | 1745.416547 | 0.266725579 | 3.50E-07 | 8.16E-06 |
| ENSMUST00000058978.7,Nqo2-202,protein_coding,ENSMUSG00000046949.15,protein_coding | 2607.440183 | 0.223445846 | 3.50E-07 | 8.16E-06 |
| ENSMUST00000060366.6,Zfp36l2-001,protein_coding,ENSMUSG00000045817.8,protein_coding | 37641.68748 | 0.142217918 | 3.50E-07 | 8.16E-06 |
| ENSMUST00000044684.13,Helq-001,protein_coding,ENSMUSG00000035266.16,protein_coding | 3323.082984 | 0.258850488 | 3.55E-07 | 8.26E-06 |
| ENSMUST00000084117.12,H6pd-002,protein_coding,ENSMUSG00000028980.14,protein_coding | 6742.669238 | -0.423891804 | 3.55E-07 | 8.26E-06 |
| ENSMUST00000025375.13,Tcerg1-001,protein_coding,ENSMUSG00000024498.15,protein_coding | 15422.81845 | -0.184241156 | 3.56E-07 | 8.28E-06 |
| ENSMUST00000064104.12,Cdc42se2-001,protein_coding,ENSMUSG00000052298.12,protein_coding | 11285.57601 | 0.266964064 | 3.56E-07 | 8.28E-06 |
| ENSMUST00000008542.11,Elk3-001,protein_coding,ENSMUSG00000008398.14,protein_coding | 10339.66768 | -0.157104274 | 3.58E-07 | 8.32E-06 |
| ENSMUST00000036069.6,Mamdc2-201,protein_coding,ENSMUSG00000033207.6,protein_coding | 175.0901449 | 0.804246145 | 3.58E-07 | 8.32E-06 |
| ENSMUST00000196454.4,Orai2-003,protein_coding,ENSMUSG00000039747.11,protein_coding | 143.9319824 | 0.917437314 | 3.59E-07 | 8.34E-06 |
| ENSMUST00000048678.6,Lss-001,protein_coding,ENSMUSG00000033105.12,protein_coding | 15668.48704 | 0.327729754 | 3.60E-07 | 8.36E-06 |
| ENSMUST00000148483.7,Zfp688-003,nonsense_mediated_decay,ENSMUSG00000045251.13,protein_coding | 408.2745576 | 0.705622792 | 3.64E-07 | 8.45E-06 |
| ENSMUST00000023356.6,Snai2-201,protein_coding,ENSMUSG00000022676.6,protein_coding | 5838.495458 | 0.4396923 | 3.64E-07 | 8.45E-06 |
| ENSMUST00000067880.12,Adam10-001,protein_coding,ENSMUSG00000054693.14,protein_coding | 14569.20562 | -0.440559142 | 3.65E-07 | 8.45E-06 |
| ENSMUST00000111678.7,Ctnnd1-010,protein_coding,ENSMUSG00000034101.14,protein_coding | 52.79135004 | 0.779016891 | 3.66E-07 | 8.48E-06 |
| ENSMUST00000204483.1,Arl8b-002,protein_coding,ENSMUSG00000030105.8,protein_coding | 2076.331116 | 0.259699232 | 3.69E-07 | 8.53E-06 |
| ENSMUST00000161976.7,Myc-001,protein_coding,ENSMUSG00000022346.15,protein_coding | 5674.565574 | 0.320014154 | 3.69E-07 | 8.53E-06 |
| ENSMUST00000022971.7,Myc-005,protein_coding,ENSMUSG00000022346.15,protein_coding | 5674.565574 | 0.320014154 | 3.69E-07 | 8.53E-06 |
| ENSMUST00000025853.15,Drap1-001,protein_coding,ENSMUSG00000024914.16,protein_coding | 12384.43996 | 0.228303423 | 3.69E-07 | 8.53E-06 |
| ENSMUST00000123362.7,Mecp2-003,protein_coding,ENSMUSG00000031393.16,protein_coding | 460.2435701 | 0.655248395 | 3.69E-07 | 8.53E-06 |
| ENSMUST00000025786.7,Pacs1-201,protein_coding,ENSMUSG00000024855.9,protein_coding | 14416.05956 | 0.163264749 | 3.75E-07 | 8.66E-06 |
| ENSMUST00000062957.6,Ttc7b-201,protein_coding,ENSMUSG00000033530.7,protein_coding | 2363.717592 | 0.258258357 | 3.83E-07 | 8.83E-06 |
| ENSMUST00000113708.3,Tia1-009,retained_intron,ENSMUSG00000071337.11,protein_coding | 101.3265355 | 0.878955732 | 3.84E-07 | 8.85E-06 |
| ENSMUST00000095457.10,Ptbp1-005,protein_coding,ENSMUSG00000006498.16,protein_coding | 30333.39218 | 0.267729604 | 3.86E-07 | 8.89E-06 |
| ENSMUST00000099362.10,Slc4a11-001,protein_coding,ENSMUSG00000074796.10,protein_coding | 348.2385398 | 0.596256673 | 3.86E-07 | 8.89E-06 |
| ENSMUST00000139167.2,Gatc-001,protein_coding,ENSMUSG00000029536.8,protein_coding | 3934.375544 | 0.256825425 | 3.87E-07 | 8.91E-06 |
| ENSMUST00000071065.7,Nfil3-001,protein_coding,ENSMUSG00000056749.7,protein_coding | 5522.036746 | 0.379460554 | 3.87E-07 | 8.91E-06 |
| ENSMUST00000182980.7,Tmem161a-011,protein_coding,ENSMUSG00000002342.17,protein_coding | 5862.128895 | 0.265710655 | 3.96E-07 | 9.10E-06 |
| ENSMUST00000047488.14,Rap1gap2-005,nonsense_mediated_decay,ENSMUSG00000038807.18,protein_coding | 805.1220287 | 0.639257576 | 3.96E-07 | 9.10E-06 |
| ENSMUST00000058154.14,Tmtc3-001,protein_coding,ENSMUSG00000036676.14,protein_coding | 4879.520802 | -0.590560067 | 3.97E-07 | 9.11E-06 |
| ENSMUST00000114294.7,Abcg2-007,protein_coding,ENSMUSG00000029802.13,protein_coding | 3964.904113 | -0.227938159 | 3.99E-07 | 9.16E-06 |
| ENSMUST00000123183.7,Rpl5-003,retained_intron,ENSMUSG00000058558.12,protein_coding | 18305.64407 | 0.223079177 | 4.00E-07 | 9.17E-06 |
| ENSMUST00000030201.13,Gne-001,protein_coding,ENSMUSG00000028479.18,protein_coding | 237.5495424 | -0.68800932 | 4.05E-07 | 9.27E-06 |
| ENSMUST00000050476.10,Slc35a4-002,protein_coding,ENSMUSG00000033272.12,protein_coding | 2782.023174 | -0.418210024 | 4.04E-07 | 9.27E-06 |
| ENSMUST00000112751.1,Bcl2-001,protein_coding,ENSMUSG00000057329.7,protein_coding | 2900.379715 | -0.374681123 | 4.07E-07 | 9.32E-06 |
| ENSMUST00000202713.1,Suds3-003,retained_intron,ENSMUSG00000066900.11,protein_coding | 420.894471 | -0.449078105 | 4.08E-07 | 9.34E-06 |
| ENSMUST00000076169.3,Mtx3-201,protein_coding,ENSMUSG00000021704.7,protein_coding | 1621.788611 | -0.33036001 | 4.09E-07 | 9.36E-06 |
| ENSMUST00000056045.4,Fam185a-001,protein_coding,ENSMUSG00000047221.5,protein_coding | 1204.717681 | -0.301968429 | 4.09E-07 | 9.36E-06 |
| ENSMUST00000070635.12,Zyx-002,protein_coding,ENSMUSG00000029860.16,protein_coding | 6709.791282 | 0.358189266 | 4.15E-07 | 9.48E-06 |
| ENSMUST00000067284.8,Cpsf3-201,protein_coding,ENSMUSG00000054309.8,protein_coding | 13555.62173 | 0.210607091 | 4.15E-07 | 9.48E-06 |
| ENSMUST00000054096.12,Man1c1-001,protein_coding,ENSMUSG00000037306.13,protein_coding | 1233.391891 | -0.261342218 | 4.16E-07 | 9.51E-06 |
| ENSMUST00000108236.4,Spint2-001,protein_coding,ENSMUSG00000074227.12,protein_coding | 650.8825725 | 0.372255256 | 4.19E-07 | 9.57E-06 |
| ENSMUST00000058444.9,Ppp1r37-001,protein_coding,ENSMUSG00000051403.9,protein_coding | 4181.936686 | -0.33413887 | 4.22E-07 | 9.62E-06 |
| ENSMUST00000065917.14,Nap1l1-201,protein_coding,ENSMUSG00000058799.14,protein_coding | 39085.46161 | 0.245475936 | 4.25E-07 | 9.68E-06 |
| ENSMUST00000168536.7,Nt5c2-202,protein_coding,ENSMUSG00000025041.16,protein_coding | 1745.836669 | -0.41505941 | 4.25E-07 | 9.69E-06 |
| ENSMUST00000034775.9,Fem1b-001,protein_coding,ENSMUSG00000032244.9,protein_coding | 8587.709165 | -0.159633154 | 4.26E-07 | 9.70E-06 |
| ENSMUST00000023818.9,Calcoco1-001,protein_coding,ENSMUSG00000023055.9,protein_coding | 6440.032271 | 0.305318592 | 4.28E-07 | 9.74E-06 |
| ENSMUST00000023083.7,Cyp2d22-201,protein_coding,ENSMUSG00000061740.7,protein_coding | 1174.174042 | 0.435586233 | 4.35E-07 | 9.90E-06 |
| ENSMUST00000032183.5,Tmem43-001,protein_coding,ENSMUSG00000030095.10,protein_coding | 9363.910737 | 0.174754682 | 4.39E-07 | 9.99E-06 |
| ENSMUST00000174266.7,Uggt1-002,nonsense_mediated_decay,ENSMUSG00000037470.14,protein_coding | 17604.97241 | 0.193587603 | 4.41E-07 | 1.00E-05 |
| ENSMUST00000159899.7,Pigk-001,protein_coding,ENSMUSG00000039047.17,protein_coding | 3059.392955 | -0.287236275 | 4.41E-07 | 1.00E-05 |
| ENSMUST00000212206.1,Zfp1-003,nonsense_mediated_decay,ENSMUSG00000055835.10,protein_coding | 54.68008888 | 0.925089558 | 4.42E-07 | 1.00E-05 |
| ENSMUST00000094434.12,Ftl1-001,protein_coding,ENSMUSG00000050708.16,protein_coding | 400431.3302 | 0.195650426 | 4.43E-07 | 1.00E-05 |
| ENSMUST00000019909.7,Rgs17-003,protein_coding,ENSMUSG00000019775.17,protein_coding | 143.695101 | 0.90233867 | 4.43E-07 | 1.00E-05 |
| ENSMUST00000071268.10,Taok2-001,protein_coding,ENSMUSG00000059981.12,protein_coding | 4306.955057 | -0.208349952 | 4.45E-07 | 1.01E-05 |
| ENSMUST00000047348.9,Spag1-201,protein_coding,ENSMUSG00000037617.11,protein_coding | 198.6844251 | -0.719712801 | 4.46E-07 | 1.01E-05 |
| ENSMUST00000033135.8,Tial1-004,protein_coding,ENSMUSG00000030846.15,protein_coding | 4987.599671 | 0.244178667 | 4.52E-07 | 1.02E-05 |
| ENSMUST00000111303.1,Dgkz-003,protein_coding,ENSMUSG00000040479.11,protein_coding | 650.4252399 | -0.466322634 | 4.52E-07 | 1.02E-05 |
| ENSMUST00000090806.4,Rai1-001,protein_coding,ENSMUSG00000062115.15,protein_coding | 3441.468827 | 0.315232046 | 4.53E-07 | 1.02E-05 |
| ENSMUST00000166617.7,Dlk2-002,protein_coding,ENSMUSG00000047428.14,protein_coding | 1624.408721 | -0.389282699 | 4.53E-07 | 1.02E-05 |
| ENSMUST00000040430.7,Vat1-001,protein_coding,ENSMUSG00000034993.7,protein_coding | 29048.48375 | 0.213215351 | 4.58E-07 | 1.03E-05 |
| ENSMUST00000146552.7,Lrp8-009,processed_transcript,ENSMUSG00000028613.14,protein_coding | 494.4819885 | 0.723695389 | 4.59E-07 | 1.04E-05 |
| ENSMUST00000023514.3,Ndufb4-001,protein_coding,ENSMUSG00000022820.3,protein_coding | 3259.185401 | 0.287751264 | 4.59E-07 | 1.04E-05 |
| ENSMUST00000192185.5,Sft2d2-002,nonsense_mediated_decay,ENSMUSG00000040848.11,protein_coding | 1509.822583 | 0.342777879 | 4.61E-07 | 1.04E-05 |
| ENSMUST00000043867.10,Rpl22l1-001,protein_coding,ENSMUSG00000039221.10,protein_coding | 1070.161543 | -0.421155752 | 4.61E-07 | 1.04E-05 |
| ENSMUST00000162461.6,Rad52-001,protein_coding,ENSMUSG00000030166.14,protein_coding | 714.6822377 | 0.575558413 | 4.61E-07 | 1.04E-05 |
| ENSMUST00000171901.7,Gtpbp2-007,processed_transcript,ENSMUSG00000023952.13,protein_coding | 63.92960439 | 0.848809707 | 4.61E-07 | 1.04E-05 |
| ENSMUST00000191189.6,Morf4l1-014,protein_coding,ENSMUSG00000062270.13,protein_coding | 501.6010394 | 0.784186458 | 4.63E-07 | 1.04E-05 |
| ENSMUST00000064272.9,B4galt3-002,protein_coding,ENSMUSG00000052423.14,protein_coding | 4245.407991 | 0.307953584 | 4.70E-07 | 1.06E-05 |
| ENSMUST00000149163.7,Tpd52l2-001,protein_coding,ENSMUSG00000000827.18,protein_coding | 241.9339478 | 0.841324346 | 4.70E-07 | 1.06E-05 |
| ENSMUST00000047616.9,Jmjd6-001,protein_coding,ENSMUSG00000056962.11,protein_coding | 6422.941306 | -0.187200791 | 4.71E-07 | 1.06E-05 |
| ENSMUST00000177752.7,Fkbp2-202,protein_coding,ENSMUSG00000056629.15,protein_coding | 2772.015359 | -0.318520098 | 4.73E-07 | 1.06E-05 |
| ENSMUST00000109869.1,Psmf1-006,protein_coding,ENSMUSG00000032869.15,protein_coding | 4890.366062 | 0.275643292 | 4.73E-07 | 1.06E-05 |
| ENSMUST00000204293.2,Serbp1-004,protein_coding,ENSMUSG00000036371.6,protein_coding | 54025.90215 | 0.161964528 | 4.81E-07 | 1.08E-05 |
| ENSMUST00000163530.7,Tars2-003,protein_coding,ENSMUSG00000028107.14,protein_coding | 1277.11258 | 0.521977651 | 4.81E-07 | 1.08E-05 |
| ENSMUST00000100334.10,Nsmf-001,protein_coding,ENSMUSG00000006476.19,protein_coding | 489.9135407 | 0.872321177 | 4.87E-07 | 1.09E-05 |
| ENSMUST00000043325.8,Hs2st1-001,protein_coding,ENSMUSG00000040151.9,protein_coding | 18664.50931 | -0.147701017 | 4.90E-07 | 1.10E-05 |
| ENSMUST00000106048.9,Sirt3-002,protein_coding,ENSMUSG00000025486.17,protein_coding | 589.2232522 | 0.574064917 | 4.91E-07 | 1.10E-05 |
| ENSMUST00000021142.7,Atp2a3-201,protein_coding,ENSMUSG00000020788.15,protein_coding | 358.6706513 | 0.733425279 | 4.93E-07 | 1.10E-05 |
| ENSMUST00000038696.11,Ppp1r9b-001,protein_coding,ENSMUSG00000038976.12,protein_coding | 13847.46663 | -0.11792025 | 5.01E-07 | 1.12E-05 |
| ENSMUST00000102979.1,Hist1h4n-001,protein_coding,ENSMUSG00000069305.3,protein_coding | 192.9404085 | 0.552675077 | 5.02E-07 | 1.12E-05 |
| ENSMUST00000073791.9,Atp5h-004,protein_coding,ENSMUSG00000034566.10,protein_coding | 241.8500921 | 0.79894854 | 5.05E-07 | 1.13E-05 |
| ENSMUST00000105845.8,Ephb2-001,protein_coding,ENSMUSG00000028664.14,protein_coding | 151.5582978 | -0.758866884 | 5.08E-07 | 1.14E-05 |
| ENSMUST00000126469.1,Eef1b2-007,processed_transcript,ENSMUSG00000025967.16,protein_coding | 4812.059526 | 0.27526739 | 5.09E-07 | 1.14E-05 |
| ENSMUST00000031367.14,Slc15a4-001,protein_coding,ENSMUSG00000029416.17,protein_coding | 2790.046738 | 0.38379573 | 5.12E-07 | 1.14E-05 |
| ENSMUST00000172142.7,Nisch-001,protein_coding,ENSMUSG00000021910.15,protein_coding | 4506.522447 | -0.223053725 | 5.15E-07 | 1.15E-05 |
| ENSMUST00000112025.2,2310009B15Rik-001,protein_coding,ENSMUSG00000079283.2,protein_coding | 968.8744543 | 0.387371581 | 5.19E-07 | 1.16E-05 |

| | | | | |
|---|---|---|---|---|
| ENSMUST00000027521.14,Agap1-001,protein_coding,ENSMUSG00000055013.15,protein_coding | 4830.443031 | 0.451658983 | 5.21E-07 | 1.16E-05 |
| ENSMUST00000065534.9,Def8-001,protein_coding,ENSMUSG00000001482.15,protein_coding | 6223.524346 | -0.270850917 | 5.25E-07 | 1.17E-05 |
| ENSMUST00000101385.2,Plekha8-002,protein_coding,ENSMUSG00000005225.15,protein_coding | 1583.633862 | 0.393184642 | 5.26E-07 | 1.17E-05 |
| ENSMUST00000043616.6,Zyg11b-001,protein_coding,ENSMUSG00000034636.9,protein_coding | 15956.7393 | 0.397064292 | 5.28E-07 | 1.18E-05 |
| ENSMUST00000190495.1,Mcm6-004,protein_coding,ENSMUSG00000026355.11,protein_coding | 723.7599727 | 0.467881771 | 5.33E-07 | 1.19E-05 |
| ENSMUST00000038234.12,Ift122-003,protein_coding,ENSMUSG00000030323.13,protein_coding | 1835.100831 | -0.365941862 | 5.34E-07 | 1.19E-05 |
| ENSMUST00000164766.7,Rem2-001,protein_coding,ENSMUSG00000022176.11,protein_coding | 1088.570874 | -0.445608663 | 5.34E-07 | 1.19E-05 |
| ENSMUST00000042658.4,Echdc3-001,protein_coding,ENSMUSG00000039063.5,protein_coding | 1377.919088 | 0.357037429 | 5.36E-07 | 1.19E-05 |
| ENSMUST00000132139.8,Sorbs2-001,protein_coding,ENSMUSG00000031626.17,protein_coding | 505.5780554 | -0.589973888 | 5.36E-07 | 1.19E-05 |
| ENSMUST00000115510.7,Phf14-001,protein_coding,ENSMUSG00000029629.17,protein_coding | 959.6205418 | 0.615982627 | 5.38E-07 | 1.19E-05 |
| ENSMUST00000029673.9,Efna3-001,protein_coding,ENSMUSG00000028039.11,protein_coding | 195.4712221 | -0.596450234 | 5.42E-07 | 1.20E-05 |
| ENSMUST00000028804.14,Cdc25b-001,protein_coding,ENSMUSG00000027330.16,protein_coding | 8800.060167 | 0.408230361 | 5.43E-07 | 1.20E-05 |
| ENSMUST00000030181.11,Ccdc107-001,protein_coding,ENSMUSG00000028461.12,protein_coding | 3299.041493 | 0.253451784 | 5.43E-07 | 1.20E-05 |
| ENSMUST00000068271.4,Ak1-003,protein_coding,ENSMUSG00000026817.14,protein_coding | 1169.175033 | -0.494849275 | 5.47E-07 | 1.21E-05 |
| ENSMUST00000034966.8,Rpl4-001,protein_coding,ENSMUSG00000032399.8,protein_coding | 207257.6804 | 0.231405888 | 5.55E-07 | 1.23E-05 |
| ENSMUST00000026665.7,Cbx4-001,protein_coding,ENSMUSG00000039989.7,protein_coding | 6129.606438 | 0.378771172 | 5.55E-07 | 1.23E-05 |
| ENSMUST00000026377.8,Padi3-001,protein_coding,ENSMUSG00000025328.9,protein_coding | 324.5581115 | -0.499916993 | 5.58E-07 | 1.23E-05 |
| ENSMUST00000107342.1,Chtop-003,protein_coding,ENSMUSG00000001017.15,protein_coding | 2579.872187 | -0.243685426 | 5.60E-07 | 1.24E-05 |
| ENSMUST00000170356.1,Dpy19l1-201,protein_coding,ENSMUSG00000043067.15,protein_coding | 3597.339136 | -0.318726087 | 5.61E-07 | 1.24E-05 |
| ENSMUST00000042352.10,Zmat4-001,protein_coding,ENSMUSG00000037492.16,protein_coding | 121.1096707 | -0.711385752 | 5.66E-07 | 1.25E-05 |
| ENSMUST00000113193.8,Armcx2-001,protein_coding,ENSMUSG00000033436.13,protein_coding | 4694.408758 | -0.203376768 | 5.68E-07 | 1.25E-05 |
| ENSMUST00000111562.7,Fam60a-007,protein_coding,ENSMUSG00000039985.14,protein_coding | 552.6516424 | 0.589096196 | 5.70E-07 | 1.26E-05 |
| ENSMUST00000105411.8,Pcbp3-001,protein_coding,ENSMUSG00000001120.15,protein_coding | 710.993808 | 0.415083741 | 5.73E-07 | 1.26E-05 |
| ENSMUST00000035325.14,Qsox1-001,protein_coding,ENSMUSG00000033684.14,protein_coding | 2238.391251 | 0.23369483 | 5.76E-07 | 1.27E-05 |
| ENSMUST00000111783.7,Mapkapk5-002,protein_coding,ENSMUSG00000029454.15,protein_coding | 1227.172097 | 0.438406487 | 5.82E-07 | 1.28E-05 |
| ENSMUST00000107209.7,Gabpb2-004,protein_coding,ENSMUSG00000038766.16,protein_coding | 3061.482417 | -0.686395046 | 5.86E-07 | 1.29E-05 |
| ENSMUST00000029203.12,Zfp639-201,protein_coding,ENSMUSG00000027667.13,protein_coding | 2083.446755 | 0.262251257 | 5.87E-07 | 1.29E-05 |
| ENSMUST00000034541.11,Srpr-001,protein_coding,ENSMUSG00000032042.11,protein_coding | 16153.62336 | 0.220424099 | 5.88E-07 | 1.29E-05 |
| ENSMUST00000049699.8,Pskh1-001,protein_coding,ENSMUSG00000048310.8,protein_coding | 4214.505536 | 0.269854808 | 5.93E-07 | 1.30E-05 |
| ENSMUST00000094053.6,Tnrc6a-001,protein_coding,ENSMUSG00000052707.8,protein_coding | 8571.199024 | -0.419413196 | 5.94E-07 | 1.30E-05 |
| ENSMUST00000133890.7,Cdc42bpa-008,protein_coding,ENSMUSG00000026490.18,protein_coding | 24.47473667 | 0.761929108 | 5.98E-07 | 1.31E-05 |
| ENSMUST00000057288.5,Pdia6-001,protein_coding,ENSMUSG00000020571.12,protein_coding | 31004.33763 | 0.193441715 | 5.98E-07 | 1.31E-05 |
| ENSMUST00000002044.9,Map3k3-001,protein_coding,ENSMUSG00000020700.11,protein_coding | 9513.920158 | 0.264096809 | 5.99E-07 | 1.31E-05 |
| ENSMUST00000100387.10,Tmub2-001,protein_coding,ENSMUSG00000034757.15,protein_coding | 559.7997547 | -0.677342433 | 6.01E-07 | 1.32E-05 |
| ENSMUST00000045351.12,Atg2a-001,protein_coding,ENSMUSG00000024773.14,protein_coding | 5837.681211 | -0.354366339 | 6.01E-07 | 1.32E-05 |
| ENSMUST00000148721.7,2310061I04Rik-001,protein_coding,ENSMUSG00000050705.15,protein_coding | 3517.969796 | 0.19096203 | 6.02E-07 | 1.32E-05 |
| ENSMUST00000161189.7,Ipo9-002,nonsense_mediated_decay,ENSMUSG00000041879.13,protein_coding | 20351.4082 | 0.208544325 | 6.03E-07 | 1.32E-05 |
| ENSMUST00000147473.5,Gm42791-001,protein_coding,ENSMUSG00000106892.1,protein_coding | 154.6850652 | 0.818363144 | 6.04E-07 | 1.32E-05 |
| ENSMUST00000071648.11,Vegfa-001,protein_coding,ENSMUSG00000023951.17,protein_coding | 17606.52635 | 0.382660345 | 6.08E-07 | 1.33E-05 |
| ENSMUST00000033935.15,Smim19-002,protein_coding,ENSMUSG00000031534.15,protein_coding | 1254.226454 | 0.258264014 | 6.10E-07 | 1.33E-05 |
| ENSMUST00000031377.8,Scarb2-001,protein_coding,ENSMUSG00000029426.8,protein_coding | 10515.93644 | 0.227638846 | 6.19E-07 | 1.35E-05 |
| ENSMUST00000032926.11,Tmem219-004,protein_coding,ENSMUSG00000060538.14,protein_coding | 730.5472254 | 0.430602793 | 6.21E-07 | 1.35E-05 |
| ENSMUST00000143561.1,Chid1-007,protein_coding,ENSMUSG00000025512.15,protein_coding | 780.3896591 | 0.396159734 | 6.20E-07 | 1.35E-05 |
| ENSMUST00000176896.7,Nptn-010,nonsense_mediated_decay,ENSMUSG00000032336.17,protein_coding | 354.3736442 | 0.426222428 | 6.24E-07 | 1.36E-05 |
| ENSMUST00000128978.1,Abcc5-006,retained_intron,ENSMUSG00000022822.15,protein_coding | 71.75086634 | -0.813669516 | 6.26E-07 | 1.36E-05 |
| ENSMUST00000124331.7,Hspa14-002,protein_coding,ENSMUSG00000051396.15,protein_coding | 960.5906772 | -0.294373698 | 6.27E-07 | 1.36E-05 |
| ENSMUST00000118570.1,Hmga1-003,protein_coding,ENSMUSG00000046711.15,protein_coding | 16218.68444 | 0.253113859 | 6.26E-07 | 1.36E-05 |
| ENSMUST00000171023.7,Krit1-004,protein_coding,ENSMUSG00000000600.15,protein_coding | 127.5566983 | -0.779824179 | 6.28E-07 | 1.37E-05 |
| ENSMUST00000078961.4,Kifc5b-201,protein_coding,ENSMUSG00000024301.9,protein_coding | 3991.127142 | 0.234694675 | 6.33E-07 | 1.38E-05 |
| ENSMUST00000107094.1,Jun-001,protein_coding,ENSMUSG00000052684.4,protein_coding | 2901.511062 | -0.253656042 | 6.34E-07 | 1.38E-05 |
| ENSMUST00000108302.7,Pdp1-005,protein_coding,ENSMUSG00000049225.14,protein_coding | 92.69176727 | 0.896234447 | 6.35E-07 | 1.38E-05 |
| ENSMUST00000142890.7,Tspan5-002,nonsense_mediated_decay,ENSMUSG00000028152.10,protein_coding | 5430.136899 | 0.28656516 | 6.40E-07 | 1.39E-05 |
| ENSMUST00000204368.1,Ppp6c-005,protein_coding,ENSMUSG00000026753.5,protein_coding | 524.8559051 | 0.458771376 | 6.44E-07 | 1.40E-05 |
| ENSMUST00000084838.12,Cd47-001,protein_coding,ENSMUSG00000055447.18,protein_coding | 18614.35468 | 0.226755971 | 6.44E-07 | 1.40E-05 |
| ENSMUST00000037023.8,Rps29-001,protein_coding,ENSMUSG00000034892.8,protein_coding | 34392.81144 | 0.150690879 | 6.49E-07 | 1.41E-05 |
| ENSMUST00000020982.5,Klf11-001,protein_coding,ENSMUSG00000020653.11,protein_coding | 1341.357278 | 0.440700136 | 6.50E-07 | 1.41E-05 |
| ENSMUST00000090840.9,Zrsr2-002,protein_coding,ENSMUSG00000031370.12,protein_coding | 1384.152371 | -0.300767712 | 6.52E-07 | 1.41E-05 |
| ENSMUST00000072740.12,Abr-001,protein_coding,ENSMUSG00000017631.18,protein_coding | 2178.646659 | -0.334660018 | 6.53E-07 | 1.41E-05 |
| ENSMUST00000109563.8,Ehbp1-201,protein_coding,ENSMUSG00000042302.14,protein_coding | 789.3494124 | -0.813371397 | 6.55E-07 | 1.42E-05 |
| ENSMUST00000205958.1,Ino80e-012,nonsense_mediated_decay,ENSMUSG00000030689.15,protein_coding | 388.1624616 | 0.802922778 | 6.60E-07 | 1.43E-05 |
| ENSMUST00000037585.8,Dennd6a-001,protein_coding,ENSMUSG00000040818.9,protein_coding | 2844.895808 | 0.237837519 | 6.62E-07 | 1.43E-05 |
| ENSMUST00000114552.3,Zpr1-004,protein_coding,ENSMUSG00000032078.17,protein_coding | 2529.395583 | 0.235600529 | 6.63E-07 | 1.43E-05 |
| ENSMUST00000109738.9,Rnaseh2a-201,protein_coding,ENSMUSG00000052926.15,protein_coding | 1032.974329 | -0.396365919 | 6.67E-07 | 1.44E-05 |
| ENSMUST00000058230.12,Ensa-002,protein_coding,ENSMUSG00000038619.12,protein_coding | 782.3813008 | 0.442411341 | 6.68E-07 | 1.44E-05 |
| ENSMUST00000009157.3,Dynll1-001,protein_coding,ENSMUSG00000009013.5,protein_coding | 3552.771801 | -0.438092227 | 6.71E-07 | 1.45E-05 |
| ENSMUST00000078218.10,Serhl-001,protein_coding,ENSMUSG00000058586.11,protein_coding | 2911.337307 | 0.328703062 | 6.71E-07 | 1.45E-05 |
| ENSMUST00000165033.1,Egr1-201,protein_coding,ENSMUSG00000038418.7,protein_coding | 4597.528667 | 0.23495597 | 6.73E-07 | 1.45E-05 |
| ENSMUST00000054565.6,Ift43-201,protein_coding,ENSMUSG00000007867.8,protein_coding | 1370.056617 | 0.297092457 | 6.74E-07 | 1.45E-05 |
| ENSMUST00000132417.7,0610037L13Rik-004,nonsense_mediated_decay,ENSMUSG00000028608.15,protein_coding | 394.6817177 | 0.475680133 | 6.77E-07 | 1.46E-05 |
| ENSMUST00000033560.8,Ophn1-001,protein_coding,ENSMUSG00000031214.13,protein_coding | 235.1884452 | -0.570044202 | 6.82E-07 | 1.47E-05 |
| ENSMUST00000198030.1,Tm4sf1-004,processed_transcript,ENSMUSG00000027800.14,protein_coding | 14254.27033 | -0.357947963 | 6.89E-07 | 1.48E-05 |
| ENSMUST00000127464.7,Wdr77-003,retained_intron,ENSMUSG00000000561.14,protein_coding | 1132.639003 | -0.32233153 | 6.90E-07 | 1.48E-05 |
| ENSMUST00000216508.1,Hinfp-001,protein_coding,ENSMUSG00000032119.5,protein_coding | 2601.400396 | 0.182803781 | 6.91E-07 | 1.48E-05 |
| ENSMUST00000039608.7,Dym-201,protein_coding,ENSMUSG00000035765.9,protein_coding | 4559.581651 | 0.23146377 | 6.92E-07 | 1.49E-05 |
| ENSMUST00000041012.8,Ptcra-001,protein_coding,ENSMUSG00000036858.8,protein_coding | 77.39575721 | 0.723845753 | 6.94E-07 | 1.49E-05 |
| ENSMUST00000211741.1,Sae1-002,protein_coding,ENSMUSG00000052833.9,protein_coding | 641.2209742 | 0.628322278 | 7.04E-07 | 1.51E-05 |
| ENSMUST00000145838.7,Ext2-006,protein_coding,ENSMUSG00000027198.16,protein_coding | 932.5487495 | 0.409890705 | 7.04E-07 | 1.51E-05 |
| ENSMUST00000041010.14,Ccdc9-001,protein_coding,ENSMUSG00000041375.18,protein_coding | 1931.066146 | 0.354155538 | 7.19E-07 | 1.54E-05 |
| ENSMUST00000153740.1,Gabre-003,retained_intron,ENSMUSG00000031340.8,protein_coding | 23.85845354 | 0.9126773 | 7.27E-07 | 1.56E-05 |
| ENSMUST00000118830.7,Vps29-002,protein_coding,ENSMUSG00000029462.18,protein_coding | 4250.550997 | 0.152517822 | 7.34E-07 | 1.57E-05 |
| ENSMUST00000092219.13,Tmpo-002,protein_coding,ENSMUSG00000019961.17,protein_coding | 7387.030691 | 0.220586738 | 7.35E-07 | 1.57E-05 |
| ENSMUST00000028915.5,Rbbp9-001,protein_coding,ENSMUSG00000027428.9,protein_coding | 2195.712566 | -0.305205504 | 7.40E-07 | 1.58E-05 |
| ENSMUST00000020493.7,Pofut2-201,protein_coding,ENSMUSG00000020260.8,protein_coding | 10467.40563 | 0.174400174 | 7.40E-07 | 1.58E-05 |
| ENSMUST00000110030.9,Snx5-001,protein_coding,ENSMUSG00000027423.15,protein_coding | 17902.5906 | -0.216463966 | 7.42E-07 | 1.59E-05 |
| ENSMUST00000158210.1,Gm23547-201,snoRNA,ENSMUSG00000088835.1,snoRNA | 752.920653 | 0.414349689 | 7.44E-07 | 1.59E-05 |
| ENSMUST00000161266.1,Cct5-003,protein_coding,ENSMUSG00000022234.14,protein_coding | 1299.745453 | 0.368428894 | 7.46E-07 | 1.59E-05 |
| ENSMUST00000145255.1,Hsp90aa1-004,retained_intron,ENSMUSG00000021270.13,protein_coding | 145.8308731 | -0.72696778 | 7.50E-07 | 1.60E-05 |
| ENSMUST00000025293.3,Ndfip1-201,protein_coding,ENSMUSG00000024425.3,protein_coding | 6221.87268 | 0.169296027 | 7.52E-07 | 1.60E-05 |
| ENSMUST00000170055.7,Ttll7-201,protein_coding,ENSMUSG00000036745.15,protein_coding | 775.0759507 | -0.72882552 | 7.52E-07 | 1.60E-05 |
| ENSMUST00000113676.5,Slc30a9-002,protein_coding,ENSMUSG00000029221.15,protein_coding | 707.5240985 | -0.56161425 | 7.56E-07 | 1.61E-05 |
| ENSMUST00000196908.4,Pdlim5-002,protein_coding,ENSMUSG00000028273.15,protein_coding | 369.4173867 | 0.613557008 | 7.57E-07 | 1.61E-05 |
| ENSMUST00000037260.6,Zfp623-201,protein_coding,ENSMUSG00000050846.8,protein_coding | 3209.552226 | 0.252840519 | 7.59E-07 | 1.61E-05 |

| | | | | |
|---|---|---|---|---|
| ENSMUST00000160916.7,Kctd17-008,retained_intron,ENSMUSG00000033287.15,protein_coding | 4418.606153 | -0.273672277 | 7.76E-07 | 1.65E-05 |
| ENSMUST00000024779.14,Usp49-001,protein_coding,ENSMUSG00000090115.7,protein_coding | 2430.365896 | -0.223717821 | 7.83E-07 | 1.67E-05 |
| ENSMUST00000105520.7,Enpp1-001,protein_coding,ENSMUSG00000037370.13,protein_coding | 460.1663809 | -0.433022807 | 7.85E-07 | 1.67E-05 |
| ENSMUST00000167636.7,Elovl1-201,protein_coding,ENSMUSG00000006390.15,protein_coding | 4389.20007 | 0.238714884 | 7.90E-07 | 1.68E-05 |
| ENSMUST00000151059.7,Bnip3l-004,processed_transcript,ENSMUSG00000022051.15,protein_coding | 115.3003827 | 0.901299141 | 7.91E-07 | 1.68E-05 |
| ENSMUST00000015612.13,Notch4-001,protein_coding,ENSMUSG00000015468.14,protein_coding | 411.8268939 | -0.427311694 | 7.95E-07 | 1.69E-05 |
| ENSMUST00000170905.7,Nfatc1-204,protein_coding,ENSMUSG00000033016.15,protein_coding | 626.2215143 | -0.829092623 | 7.97E-07 | 1.69E-05 |
| ENSMUST00000120512.7,Mdfic-004,protein_coding,ENSMUSG00000041390.18,protein_coding | 2637.260663 | 0.191136717 | 7.98E-07 | 1.69E-05 |
| ENSMUST00000015791.5,Lama5-001,protein_coding,ENSMUSG00000015647.9,protein_coding | 47170.69365 | 0.241594036 | 7.99E-07 | 1.69E-05 |
| ENSMUST00000170075.8,Daxx-001,protein_coding,ENSMUSG00000002307.15,protein_coding | 50.21298104 | -0.913450981 | 8.00E-07 | 1.69E-05 |
| ENSMUST00000102743.9,Rapgef6-001,protein_coding,ENSMUSG00000037533.18,protein_coding | 4428.314973 | -0.469605219 | 8.02E-07 | 1.70E-05 |
| ENSMUST00000063314.6,Gtf2a1-002,protein_coding,ENSMUSG00000020962.13,protein_coding | 705.7529559 | 0.52523212 | 8.08E-07 | 1.71E-05 |
| ENSMUST00000114604.8,Etl4-019,protein_coding,ENSMUSG00000036617.16,protein_coding | 208.6197249 | 0.904046048 | 8.15E-07 | 1.72E-05 |
| ENSMUST00000042135.13,Rasa4-001,protein_coding,ENSMUSG00000004952.13,protein_coding | 1318.691886 | 0.238234769 | 8.27E-07 | 1.75E-05 |
| ENSMUST00000094471.9,Patz1-003,protein_coding,ENSMUSG00000020453.17,protein_coding | 1258.405021 | -0.282612138 | 8.32E-07 | 1.76E-05 |
| ENSMUST00000051241.6,Zfp36-001,protein_coding,ENSMUSG00000044786.6,protein_coding | 1030.968221 | 0.399543405 | 8.41E-07 | 1.77E-05 |
| ENSMUST00000118628.7,Josd2-006,protein_coding,ENSMUSG00000038695.13,protein_coding | 892.6034723 | 0.272106413 | 8.44E-07 | 1.78E-05 |
| ENSMUST00000084881.4,Crtap-001,protein_coding,ENSMUSG00000032431.6,protein_coding | 6916.034986 | 0.182997106 | 8.52E-07 | 1.80E-05 |
| ENSMUST00000039631.8,Acta2-201,protein_coding,ENSMUSG00000035783.8,protein_coding | 8675.237549 | -0.243637973 | 8.53E-07 | 1.80E-05 |
| ENSMUST00000058045.4,Garem2-001,protein_coding,ENSMUSG00000044576.6,protein_coding | 142.1500203 | -0.61605867 | 8.55E-07 | 1.80E-05 |
| ENSMUST00000084125.9,Tardbp-001,protein_coding,ENSMUSG00000041459.15,protein_coding | 3534.34739 | -0.607966714 | 8.55E-07 | 1.80E-05 |
| ENSMUST00000035604.12,Gemin5-001,protein_coding,ENSMUSG00000037275.14,protein_coding | 7272.750289 | 0.304895073 | 8.56E-07 | 1.80E-05 |
| ENSMUST00000029046.8,Fabp5-001,protein_coding,ENSMUSG00000027533.10,protein_coding | 15421.25319 | 0.227394075 | 8.62E-07 | 1.81E-05 |
| ENSMUST00000000642.10,Hk2-001,protein_coding,ENSMUSG00000000628.10,protein_coding | 20467.51578 | 0.309349347 | 8.68E-07 | 1.82E-05 |
| ENSMUST00000210503.1,Tmem143-006,protein_coding,ENSMUSG00000002781.19,protein_coding | 234.7599316 | 0.594251477 | 8.79E-07 | 1.85E-05 |
| ENSMUST00000006311.12,Tead4-001,protein_coding,ENSMUSG00000030353.15,protein_coding | 3070.927828 | 0.241076193 | 8.80E-07 | 1.85E-05 |
| ENSMUST00000020640.7,Rack1-001,protein_coding,ENSMUSG00000020372.15,protein_coding | 202994.2733 | 0.242743174 | 8.83E-07 | 1.85E-05 |
| ENSMUST00000035128.10,Abhd5-002,nonsense_mediated_decay,ENSMUSG00000032540.16,protein_coding | 815.902861 | 0.372715341 | 8.84E-07 | 1.85E-05 |
| ENSMUST00000061937.12,Ctbs-001,protein_coding,ENSMUSG00000028189.12,protein_coding | 1307.831367 | 0.278376327 | 8.90E-07 | 1.87E-05 |
| ENSMUST00000019734.10,Cyb561-001,protein_coding,ENSMUSG00000019590.16,protein_coding | 981.7490869 | 0.319722673 | 8.92E-07 | 1.87E-05 |
| ENSMUST00000164904.1,Ubl3-201,protein_coding,ENSMUSG00000001687.15,protein_coding | 329.1889262 | 0.630545647 | 8.99E-07 | 1.88E-05 |
| ENSMUST00000079601.12,Etv5-001,protein_coding,ENSMUSG00000013089.15,protein_coding | 13939.96456 | 0.21998971 | 9.00E-07 | 1.88E-05 |
| ENSMUST00000191104.6,Tns1-008,protein_coding,ENSMUSG00000055322.15,protein_coding | 22492.04967 | -0.192820144 | 9.05E-07 | 1.90E-05 |
| ENSMUST00000072847.11,Rnf138-202,protein_coding,ENSMUSG00000024317.14,protein_coding | 749.4467488 | 0.442578498 | 9.06E-07 | 1.90E-05 |
| ENSMUST00000073927.4,Slc7a3-201,protein_coding,ENSMUSG00000031297.14,protein_coding | 3771.388007 | 0.476737363 | 9.06E-07 | 1.90E-05 |
| ENSMUST00000005616.15,Pkn1-001,protein_coding,ENSMUSG00000057672.16,protein_coding | 5014.713822 | -0.280806777 | 9.07E-07 | 1.90E-05 |
| ENSMUST00000140706.7,Becn1-005,protein_coding,ENSMUSG00000035086.13,protein_coding | 318.5351667 | -0.495080051 | 9.15E-07 | 1.91E-05 |
| ENSMUST00000063838.10,Fam49b-201,protein_coding,ENSMUSG00000022378.13,protein_coding | 649.9013748 | -0.396165148 | 9.15E-07 | 1.91E-05 |
| ENSMUST00000124355.7,Adgrl1-002,nonsense_mediated_decay,ENSMUSG00000013033.16,protein_coding | 228.5856481 | -0.723382974 | 9.15E-07 | 1.91E-05 |
| ENSMUST00000102518.9,Ece1-001,protein_coding,ENSMUSG00000057530.14,protein_coding | 9109.314687 | 0.211255181 | 9.23E-07 | 1.92E-05 |
| ENSMUST00000024839.4,Sik1-201,protein_coding,ENSMUSG00000024042.6,protein_coding | 2194.639002 | -0.224018684 | 9.23E-07 | 1.92E-05 |
| ENSMUST00000033905.12,Ankrd10-001,protein_coding,ENSMUSG00000031508.14,protein_coding | 860.2591604 | -0.613704548 | 9.26E-07 | 1.93E-05 |
| ENSMUST00000119947.1,Gm15725-001,transcribed_processed_pseudogene,ENSMUSG00000083405.1,transcribed_processed_pseudogene | 485.1687694 | -0.539021441 | 9.28E-07 | 1.93E-05 |
| ENSMUST00000091403.4,Arsb-201,protein_coding,ENSMUSG00000042082.6,protein_coding | 6360.130572 | 0.213587557 | 9.29E-07 | 1.93E-05 |
| ENSMUST00000172908.1,Naa25-004,nonsense_mediated_decay,ENSMUSG00000042719.17,protein_coding | 166.0873488 | 0.829159499 | 9.33E-07 | 1.94E-05 |
| ENSMUST00000175096.1,Rpph1-201,ribozyme,ENSMUSG00000092837.1,ribozyme | 44.82997211 | -0.873512163 | 9.36E-07 | 1.95E-05 |
| ENSMUST00000170263.8,Dctd-001,protein_coding,ENSMUSG00000031562.15,protein_coding | 4048.590883 | 0.288454663 | 9.47E-07 | 1.97E-05 |
| ENSMUST00000077798.12,Ssrp1-001,protein_coding,ENSMUSG00000027067.16,protein_coding | 66652.71697 | 0.205536389 | 9.48E-07 | 1.97E-05 |
| ENSMUST00000030346.12,0610037L13Rik-011,processed_transcript,ENSMUSG00000028608.15,protein_coding | 297.1101693 | -0.790437181 | 9.58E-07 | 1.99E-05 |
| ENSMUST00000063089.11,Nsun3-001,protein_coding,ENSMUSG00000050312.11,protein_coding | 433.3987127 | -0.689088537 | 9.58E-07 | 1.99E-05 |
| ENSMUST00000141722.7,Stxbp5-002,protein_coding,ENSMUSG00000019790.16,protein_coding | 3176.275842 | 0.246480607 | 9.59E-07 | 1.99E-05 |
| ENSMUST00000006105.6,Shpk-002,protein_coding,ENSMUSG00000005951.13,protein_coding | 238.0061347 | 0.513356596 | 9.73E-07 | 2.02E-05 |
| ENSMUST00000022419.6,Ppif-001,protein_coding,ENSMUSG00000021868.6,protein_coding | 4256.893521 | 0.208519371 | 9.75E-07 | 2.02E-05 |
| ENSMUST00000100484.4,Eif3d-201,protein_coding,ENSMUSG00000016554.8,protein_coding | 66239.3234 | 0.20658827 | 9.79E-07 | 2.03E-05 |
| ENSMUST00000110690.8,Oxct1-001,protein_coding,ENSMUSG00000022186.14,protein_coding | 8287.774656 | -0.20635035 | 9.82E-07 | 2.03E-05 |
| ENSMUST00000049346.9,Asic3-001,protein_coding,ENSMUSG00000038276.12,protein_coding | 133.2022646 | -0.873806235 | 9.83E-07 | 2.04E-05 |
| ENSMUST00000030365.5,Mrpl37-001,protein_coding,ENSMUSG00000028622.11,protein_coding | 8138.326867 | -0.18729863 | 9.85E-07 | 2.04E-05 |
| ENSMUST00000103073.8,Cass4-002,protein_coding,ENSMUSG00000074570.13,protein_coding | 176.3537064 | 0.705490815 | 9.90E-07 | 2.05E-05 |
| ENSMUST00000070515.1,Ephx2-201,protein_coding,ENSMUSG00000022040.7,protein_coding | 619.4635177 | 0.431716313 | 9.95E-07 | 2.06E-05 |
| ENSMUST00000048116.14,Slc7a1-001,protein_coding,ENSMUSG00000041313.14,protein_coding | 54935.74929 | 0.287139106 | 9.96E-07 | 2.06E-05 |
| ENSMUST00000023913.8,Wrb-001,protein_coding,ENSMUSG00000023147.15,protein_coding | 3484.272964 | -0.279635346 | 1.00E-06 | 2.07E-05 |
| ENSMUST00000043560.14,Smg7-003,protein_coding,ENSMUSG00000042772.15,protein_coding | 2104.028983 | -0.386607477 | 1.00E-06 | 2.07E-05 |
| ENSMUST00000103143.9,Fbxl20-001,protein_coding,ENSMUSG00000020883.18,protein_coding | 4082.331455 | 0.274617558 | 1.00E-06 | 2.07E-05 |
| ENSMUST00000130675.1,Txnl4a-004,processed_transcript,ENSMUSG00000057130.11,protein_coding | 329.3578107 | 0.557756325 | 1.01E-06 | 2.08E-05 |
| ENSMUST00000209099.1,U2af2-009,protein_coding,ENSMUSG00000030435.16,protein_coding | 11599.3265 | -0.225743071 | 1.01E-06 | 2.08E-05 |
| ENSMUST00000063084.15,Xbp1-001,protein_coding,ENSMUSG00000020484.18,protein_coding | 7810.032599 | 0.280096523 | 1.01E-06 | 2.08E-05 |
| ENSMUST00000112320.7,Spc25-003,protein_coding,ENSMUSG00000005233.16,protein_coding | 456.4009912 | -0.606641449 | 1.02E-06 | 2.09E-05 |
| ENSMUST00000216095.1,Clk3-003,retained_intron,ENSMUSG00000032316.8,protein_coding | 511.5640955 | 0.461499028 | 1.02E-06 | 2.11E-05 |
| ENSMUST00000020361.6,Ndufs7-201,protein_coding,ENSMUSG00000020153.14,protein_coding | 9089.116187 | 0.222913566 | 1.03E-06 | 2.11E-05 |
| ENSMUST00000072187.11,Kif1c-004,protein_coding,ENSMUSG00000020821.17,protein_coding | 3629.708674 | -0.468633689 | 1.04E-06 | 2.14E-05 |
| ENSMUST00000101251.7,Cenpi-002,protein_coding,ENSMUSG00000031262.12,protein_coding | 913.0103117 | -0.782354073 | 1.05E-06 | 2.15E-05 |
| ENSMUST00000139147.1,Rad54l-004,processed_transcript,ENSMUSG00000028702.15,protein_coding | 724.4916479 | -0.333726156 | 1.05E-06 | 2.15E-05 |
| ENSMUST00000124226.2,Armcx4-001,protein_coding,ENSMUSG00000049804.9,protein_coding | 1775.168029 | -0.518945308 | 1.05E-06 | 2.15E-05 |
| ENSMUST00000153128.1,Cnnm4-001,protein_coding,ENSMUSG00000037408.10,protein_coding | 5663.736833 | 0.314128925 | 1.05E-06 | 2.16E-05 |
| ENSMUST00000106442.8,Arhgap17-001,protein_coding,ENSMUSG00000030766.15,protein_coding | 2468.937443 | 0.590527521 | 1.06E-06 | 2.18E-05 |
| ENSMUST00000142407.7,Ube2k-001,protein_coding,ENSMUSG00000029203.16,protein_coding | 3303.258243 | -0.439979721 | 1.06E-06 | 2.18E-05 |
| ENSMUST00000090292.11,Drosha-001,protein_coding,ENSMUSG00000022191.15,protein_coding | 29860.83579 | 0.135020972 | 1.07E-06 | 2.20E-05 |
| ENSMUST00000001619.14,Ergic1-004,processed_transcript,ENSMUSG00000001576.14,protein_coding | 2011.043839 | 0.457197958 | 1.08E-06 | 2.20E-05 |
| ENSMUST00000033824.7,Lamp1-001,protein_coding,ENSMUSG00000031447.7,protein_coding | 66997.36993 | 0.194925969 | 1.08E-06 | 2.21E-05 |
| ENSMUST00000134188.7,Igf2bp2-009,retained_intron,ENSMUSG00000033581.16,protein_coding | 5454.9615 | 0.256945959 | 1.09E-06 | 2.23E-05 |
| ENSMUST00000178310.7,Fau-202,protein_coding,ENSMUSG00000038274.11,protein_coding | 10912.26273 | 0.28970864 | 1.09E-06 | 2.23E-05 |
| ENSMUST00000160852.7,Sssca1-006,nonsense_mediated_decay,ENSMUSG00000079478.8,protein_coding | 983.9163405 | -0.414005084 | 1.10E-06 | 2.25E-05 |
| ENSMUST00000216934.1,Csk-007,protein_coding,ENSMUSG00000032312.7,protein_coding | 232.7083585 | 0.565827191 | 1.12E-06 | 2.29E-05 |
| ENSMUST00000056096.14,Zfp697-001,protein_coding,ENSMUSG00000050064.14,protein_coding | 1081.323076 | -0.386982656 | 1.12E-06 | 2.29E-05 |
| ENSMUST00000153224.1,Exosc8-004,protein_coding,ENSMUSG00000027752.15,protein_coding | 342.937203 | 0.452751444 | 1.12E-06 | 2.29E-05 |
| ENSMUST00000022831.3,Khnyn-201,protein_coding,ENSMUSG00000047153.3,protein_coding | 4147.986293 | 0.329617117 | 1.13E-06 | 2.31E-05 |
| ENSMUST00000125362.7,Socs6-003,protein_coding,ENSMUSG00000056153.14,protein_coding | 2985.726462 | 0.300720694 | 1.14E-06 | 2.32E-05 |
| ENSMUST00000086267.5,Fam129a-003,processed_transcript,ENSMUSG00000026483.13,protein_coding | 5858.70058 | 0.410316397 | 1.15E-06 | 2.34E-05 |
| ENSMUST00000177633.7,Zmynd8-203,protein_coding,ENSMUSG00000039671.18,protein_coding | 4806.650059 | -0.345383323 | 1.15E-06 | 2.35E-05 |
| ENSMUST00000173535.7,Bag6-003,protein_coding,ENSMUSG00000024392.17,protein_coding | 8251.888876 | 0.1780928 | 1.16E-06 | 2.36E-05 |
| ENSMUST00000087475.10,Tmem237-003,protein_coding,ENSMUSG00000038079.14,protein_coding | 2308.269024 | 0.373785727 | 1.18E-06 | 2.41E-05 |
| ENSMUST00000070756.11,Adcyap1r1-002,protein_coding,ENSMUSG00000029778.12,protein_coding | 201.31486 | -0.599218326 | 1.19E-06 | 2.42E-05 |

| | | | | |
|---|---|---|---|---|
| ENSMUST00000110036.8,Ptk2-201,protein_coding,ENSMUSG00000022607.14,protein_coding | 18122.66962 | 0.134027492 | 1.19E-06 | 2.42E-05 |
| ENSMUST00000045866.7,Ddx21-201,protein_coding,ENSMUSG00000020075.7,protein_coding | 47513.20668 | -0.223464948 | 1.20E-06 | 2.43E-05 |
| ENSMUST00000033380.6,Cdk16-001,protein_coding,ENSMUSG00000031065.14,protein_coding | 21747.64067 | 0.211685661 | 1.20E-06 | 2.43E-05 |
| ENSMUST00000208707.1,Eif3k-005,nonsense_mediated_decay,ENSMUSG00000053565.10,protein_coding | 196.1707802 | 0.581007548 | 1.21E-06 | 2.46E-05 |
| ENSMUST00000117364.7,Fam175a-201,protein_coding,ENSMUSG00000035234.18,protein_coding | 1398.412669 | 0.614618566 | 1.21E-06 | 2.46E-05 |
| ENSMUST00000067786.8,Slc20a2-001,protein_coding,ENSMUSG00000037656.9,protein_coding | 11959.23115 | 0.17311496 | 1.22E-06 | 2.47E-05 |
| ENSMUST00000155062.1,Smarce1-005,processed_transcript,ENSMUSG00000037935.16,protein_coding | 1367.000835 | 0.275041934 | 1.22E-06 | 2.48E-05 |
| ENSMUST00000034147.3,Zfp330-001,protein_coding,ENSMUSG00000031711.7,protein_coding | 5030.176353 | 0.19481325 | 1.23E-06 | 2.49E-05 |
| ENSMUST00000047903.9,Colgalt1-001,protein_coding,ENSMUSG00000034807.9,protein_coding | 16272.01259 | 0.194691644 | 1.23E-06 | 2.50E-05 |
| ENSMUST00000045737.13,Galnt11-001,protein_coding,ENSMUSG00000038072.14,protein_coding | 1567.61109 | 0.36513098 | 1.24E-06 | 2.51E-05 |
| ENSMUST00000138005.1,Ngrn-002,retained_intron,ENSMUSG00000047084.8,protein_coding | 846.4977592 | 0.567052863 | 1.24E-06 | 2.52E-05 |
| ENSMUST00000146010.2,Epb41l4aos-001,lincRNA,ENSMUSG00000087590.2,lincRNA | 972.6633834 | 0.252453224 | 1.24E-06 | 2.52E-05 |
| ENSMUST00000001127.10,Poldip2-001,protein_coding,ENSMUSG00000001100.10,protein_coding | 12295.91554 | 0.219812154 | 1.25E-06 | 2.52E-05 |
| ENSMUST00000013294.15,Gfod2-001,protein_coding,ENSMUSG00000013150.15,protein_coding | 1317.847643 | -0.426512574 | 1.25E-06 | 2.52E-05 |
| ENSMUST00000103093.9,Ctsa-011,protein_coding,ENSMUSG00000017760.15,protein_coding | 12446.21795 | 0.196845192 | 1.26E-06 | 2.55E-05 |
| ENSMUST00000032138.14,Cnbp-201,protein_coding,ENSMUSG00000030057.15,protein_coding | 9765.415039 | 0.213555622 | 1.27E-06 | 2.57E-05 |
| ENSMUST00000119223.1,E2f8-001,protein_coding,ENSMUSG00000046179.17,protein_coding | 3166.825889 | 0.216858993 | 1.29E-06 | 2.60E-05 |
| ENSMUST00000172795.1,Cyb5r1-008,nonsense_mediated_decay,ENSMUSG00000026456.18,protein_coding | 424.6470869 | 0.549716179 | 1.30E-06 | 2.62E-05 |
| ENSMUST00000166701.1,Gtpbp2-015,protein_coding,ENSMUSG00000023952.13,protein_coding | 167.5805201 | 0.681769616 | 1.30E-06 | 2.62E-05 |
| ENSMUST00000150206.7,Rabl6-003,processed_transcript,ENSMUSG00000015087.14,protein_coding | 428.5708553 | -0.420564608 | 1.30E-06 | 2.63E-05 |
| ENSMUST00000180432.8,Pvt1-004,lincRNA,ENSMUSG00000097039.8,lincRNA | 449.8329932 | 0.53121772 | 1.30E-06 | 2.63E-05 |
| ENSMUST00000016088.8,Gatsl2-001,protein_coding,ENSMUSG00000015944.9,protein_coding | 2491.986942 | -0.284827984 | 1.31E-06 | 2.63E-05 |
| ENSMUST00000069483.11,Fbrsl1-201,protein_coding,ENSMUSG00000043323.16,protein_coding | 920.707889 | 0.397901936 | 1.31E-06 | 2.63E-05 |
| ENSMUST00000144099.7,Cdk5rap2-001,protein_coding,ENSMUSG00000039298.16,protein_coding | 3605.246309 | -0.172955524 | 1.31E-06 | 2.64E-05 |
| ENSMUST00000170662.1,Sox21-001,protein_coding,ENSMUSG00000061517.8,protein_coding | 2223.063099 | -0.428504126 | 1.32E-06 | 2.66E-05 |
| ENSMUST00000196254.4,Fdps-008,protein_coding,ENSMUSG00000059743.12,protein_coding | 131.325111 | 0.727527818 | 1.32E-06 | 2.66E-05 |
| ENSMUST00000086564.10,Git2-005,protein_coding,ENSMUSG00000041890.17,protein_coding | 4374.898877 | 0.252951156 | 1.32E-06 | 2.66E-05 |
| ENSMUST00000109961.7,Nxt1-002,protein_coding,ENSMUSG00000036992.10,protein_coding | 738.492781 | 0.345785523 | 1.33E-06 | 2.68E-05 |
| ENSMUST00000022663.5,Tnfrsf10b-201,protein_coding,ENSMUSG00000022074.5,protein_coding | 6558.466837 | 0.219371394 | 1.34E-06 | 2.69E-05 |
| ENSMUST00000117509.7,Mapk8ip3-005,protein_coding,ENSMUSG00000024163.17,protein_coding | 498.0843968 | 0.781730997 | 1.34E-06 | 2.70E-05 |
| ENSMUST00000177545.1,Gm20605-001,nonsense_mediated_decay,ENSMUSG00000029720.9,protein_coding | 784.7540146 | -0.520622837 | 1.35E-06 | 2.71E-05 |
| ENSMUST00000001507.4,Cyp51-001,protein_coding,ENSMUSG00000001467.5,protein_coding | 25456.7528 | 0.314576378 | 1.35E-06 | 2.72E-05 |
| ENSMUST00000155916.7,Cobll1-008,protein_coding,ENSMUSG00000034903.18,protein_coding | 208.474583 | 0.725628029 | 1.36E-06 | 2.72E-05 |
| ENSMUST00000032503.11,Tspan9-003,protein_coding,ENSMUSG00000030352.14,protein_coding | 1001.939328 | 0.292566268 | 1.36E-06 | 2.72E-05 |
| ENSMUST00000116463.3,Gatad2a-004,protein_coding,ENSMUSG00000036180.15,protein_coding | 25461.18817 | 0.148877644 | 1.36E-06 | 2.72E-05 |
| ENSMUST00000081510.3,Mgmt-001,protein_coding,ENSMUSG00000054612.4,protein_coding | 758.9249287 | 0.287510297 | 1.36E-06 | 2.73E-05 |
| ENSMUST00000024967.13,Msh2-001,protein_coding,ENSMUSG00000024151.13,protein_coding | 8531.351087 | 0.198793608 | 1.38E-06 | 2.77E-05 |
| ENSMUST00000077874.14,Ptpmt1-201,processed_transcript,ENSMUSG00000063235.16,processed_transcript | 2249.875735 | -0.268224542 | 1.40E-06 | 2.80E-05 |
| ENSMUST00000080492.5,Rpl36-201,protein_coding,ENSMUSG00000057863.5,protein_coding | 54706.26036 | 0.204834972 | 1.40E-06 | 2.80E-05 |
| ENSMUST00000089187.6,Desi1-006,nonsense_mediated_decay,ENSMUSG00000022472.16,protein_coding | 456.0442175 | 0.591017788 | 1.41E-06 | 2.82E-05 |
| ENSMUST00000170715.7,Rps2-201,protein_coding,ENSMUSG00000044533.15,protein_coding | 566456.7895 | 0.249004534 | 1.42E-06 | 2.83E-05 |
| ENSMUST00000029092.12,Arfgap1-001,protein_coding,ENSMUSG00000027575.19,protein_coding | 5173.903927 | 0.201400349 | 1.42E-06 | 2.84E-05 |
| ENSMUST00000089634.11,Bcl2l11-003,protein_coding,ENSMUSG00000027381.16,protein_coding | 139.074623 | -0.714607633 | 1.43E-06 | 2.84E-05 |
| ENSMUST00000113296.7,Frs3-002,protein_coding,ENSMUSG00000023266.11,protein_coding | 927.6114815 | 0.404210303 | 1.43E-06 | 2.85E-05 |
| ENSMUST00000171687.7,Tmem39a-002,protein_coding,ENSMUSG00000002845.14,protein_coding | 618.4913603 | 0.631399872 | 1.44E-06 | 2.87E-05 |
| ENSMUST00000154972.7,Dnajc7-004,processed_transcript,ENSMUSG00000014195.15,protein_coding | 315.925577 | 0.545501198 | 1.45E-06 | 2.88E-05 |
| ENSMUST00000122255.7,Crtc3-001,protein_coding,ENSMUSG00000030527.15,protein_coding | 5386.771734 | 0.209748053 | 1.46E-06 | 2.90E-05 |
| ENSMUST00000140944.7,Idh3g-009,processed_transcript,ENSMUSG00000002010.17,protein_coding | 1101.34754 | -0.344309157 | 1.46E-06 | 2.90E-05 |
| ENSMUST00000049242.7,Lrrc20-201,protein_coding,ENSMUSG00000037151.7,protein_coding | 3125.368884 | 0.213769802 | 1.46E-06 | 2.90E-05 |
| ENSMUST00000034041.8,Irf2-001,protein_coding,ENSMUSG00000031627.9,protein_coding | 3225.188406 | 0.186095601 | 1.47E-06 | 2.92E-05 |
| ENSMUST00000038753.5,Sh3pxd2b-001,protein_coding,ENSMUSG00000040711.7,protein_coding | 10557.19641 | -0.160535367 | 1.47E-06 | 2.92E-05 |
| ENSMUST00000006467.13,Arpc2-002,protein_coding,ENSMUSG00000006304.14,protein_coding | 12165.8528 | -0.154220745 | 1.47E-06 | 2.93E-05 |
| ENSMUST00000071876.12,Cdpf1-001,protein_coding,ENSMUSG00000064284.12,protein_coding | 1968.062823 | 0.338208867 | 1.48E-06 | 2.93E-05 |
| ENSMUST00000101362.7,Slc7a3-002,protein_coding,ENSMUSG00000031297.14,protein_coding | 1158.15527 | 0.592344592 | 1.49E-06 | 2.95E-05 |
| ENSMUST00000074575.10,Snrnp70-001,protein_coding,ENSMUSG00000063511.11,protein_coding | 28387.49548 | 0.185629374 | 1.49E-06 | 2.96E-05 |
| ENSMUST00000046386.4,Zcchc14-001,protein_coding,ENSMUSG00000061410.6,protein_coding | 9135.542815 | -0.153066725 | 1.49E-06 | 2.96E-05 |
| ENSMUST00000142080.1,Whsc1-005,protein_coding,ENSMUSG00000057406.16,protein_coding | 2165.954972 | -0.459473252 | 1.50E-06 | 2.97E-05 |
| ENSMUST00000136390.7,Scrib-004,protein_coding,ENSMUSG00000022568.16,protein_coding | 2306.783339 | 0.350757222 | 1.51E-06 | 2.98E-05 |
| ENSMUST00000121805.7,Dpysl3-001,protein_coding,ENSMUSG00000024501.19,protein_coding | 12393.42377 | 0.288379695 | 1.51E-06 | 2.98E-05 |
| ENSMUST00000100737.9,P2rx7-001,protein_coding,ENSMUSG00000029468.17,protein_coding | 2603.891435 | 0.178406163 | 1.52E-06 | 3.01E-05 |
| ENSMUST00000020215.15,Socs2-001,protein_coding,ENSMUSG00000020027.18,protein_coding | 3431.791486 | 0.56482461 | 1.52E-06 | 3.01E-05 |
| ENSMUST00000134686.1,Ddit3-003,retained_intron,ENSMUSG00000025408.15,protein_coding | 186.4497753 | 0.548858024 | 1.52E-06 | 3.01E-05 |
| ENSMUST00000003687.6,Tgfb3-201,protein_coding,ENSMUSG00000021253.6,protein_coding | 2677.024773 | -0.335560199 | 1.52E-06 | 3.01E-05 |
| ENSMUST00000031055.7,Emilin1-001,protein_coding,ENSMUSG00000029163.9,protein_coding | 477.104835 | -0.433658775 | 1.53E-06 | 3.03E-05 |
| ENSMUST00000088644.12,Wnk1-201,protein_coding,ENSMUSG00000045962.16,protein_coding | 5269.787314 | -0.438649 | 1.54E-06 | 3.04E-05 |
| ENSMUST00000036376.12,Tmub2-002,protein_coding,ENSMUSG00000034757.15,protein_coding | 1194.911212 | 0.455448445 | 1.54E-06 | 3.05E-05 |
| ENSMUST00000019942.5,Aig1-001,protein_coding,ENSMUSG00000019806.13,protein_coding | 182.7332575 | 0.659672285 | 1.55E-06 | 3.05E-05 |
| ENSMUST00000032216.6,Ptms-001,protein_coding,ENSMUSG00000030122.12,protein_coding | 72070.37387 | 0.192152119 | 1.55E-06 | 3.05E-05 |
| ENSMUST00000165111.7,Acly-201,protein_coding,ENSMUSG00000020917.17,protein_coding | 15650.78898 | 0.270956613 | 1.55E-06 | 3.06E-05 |
| ENSMUST00000021333.4,Foxg1-001,protein_coding,ENSMUSG00000020950.10,protein_coding | 9526.009905 | 0.197101863 | 1.56E-06 | 3.07E-05 |
| ENSMUST00000212991.1,Cherp-001,protein_coding,ENSMUSG00000052488.7,protein_coding | 13371.66037 | -0.19713554 | 1.58E-06 | 3.12E-05 |
| ENSMUST00000000505.15,Mcm7-001,protein_coding,ENSMUSG00000029730.16,protein_coding | 47206.75978 | 0.205285407 | 1.59E-06 | 3.13E-05 |
| ENSMUST00000004987.15,Srpk1-002,nonsense_mediated_decay,ENSMUSG00000004865.15,protein_coding | 3491.890861 | 0.315851477 | 1.60E-06 | 3.14E-05 |
| ENSMUST00000126181.7,Cald1-012,protein_coding,ENSMUSG00000029761.16,protein_coding | 570.0327306 | 0.452360526 | 1.60E-06 | 3.15E-05 |
| ENSMUST00000047404.6,Dync1li1-001,protein_coding,ENSMUSG00000032435.9,protein_coding | 13445.65471 | 0.254416877 | 1.63E-06 | 3.19E-05 |
| ENSMUST00000035079.9,Mlh1-001,protein_coding,ENSMUSG00000032498.9,protein_coding | 2660.681051 | 0.188403197 | 1.63E-06 | 3.19E-05 |
| ENSMUST00000169659.7,Pfkfb2-002,protein_coding,ENSMUSG00000026409.14,protein_coding | 501.4194908 | -0.602150238 | 1.64E-06 | 3.21E-05 |
| ENSMUST00000109701.9,Raly-002,protein_coding,ENSMUSG00000027593.15,protein_coding | 4371.905771 | -0.456225358 | 1.64E-06 | 3.23E-05 |
| ENSMUST00000038446.8,Myl12b-001,protein_coding,ENSMUSG00000034868.8,protein_coding | 13265.96195 | 0.218425202 | 1.65E-06 | 3.23E-05 |
| ENSMUST00000146585.1,Sec14l1-002,processed_transcript,ENSMUSG00000020823.16,protein_coding | 98.87612168 | 0.724277685 | 1.66E-06 | 3.25E-05 |
| ENSMUST00000026289.9,Hsd17b10-001,protein_coding,ENSMUSG00000025260.10,protein_coding | 7860.054238 | 0.230623485 | 1.66E-06 | 3.25E-05 |
| ENSMUST00000110918.2,BC052040-001,protein_coding,ENSMUSG00000040282.13,protein_coding | 2026.867659 | 0.279771523 | 1.66E-06 | 3.25E-05 |
| ENSMUST00000124586.1,BC029722-001,protein_coding,ENSMUSG00000074649.6,protein_coding | 4714.127093 | -0.306817584 | 1.67E-06 | 3.27E-05 |
| ENSMUST00000112090.1,Dynll1-003,protein_coding,ENSMUSG00000009013.5,protein_coding | 8160.46618 | -0.342940372 | 1.67E-06 | 3.27E-05 |
| ENSMUST00000035375.13,Mdfi-003,protein_coding,ENSMUSG00000032717.14,protein_coding | 423.106091 | -0.368640393 | 1.67E-06 | 3.27E-05 |
| ENSMUST00000195955.1,Zgrf1-004,protein_coding,ENSMUSG00000051278.12,protein_coding | 144.1966493 | -0.717332902 | 1.67E-06 | 3.27E-05 |
| ENSMUST00000029147.15,Nfs1-001,protein_coding,ENSMUSG00000027618.17,protein_coding | 6524.85645 | 0.203887199 | 1.68E-06 | 3.28E-05 |
| ENSMUST00000163884.7,Tmem39a-003,protein_coding,ENSMUSG00000002845.14,protein_coding | 3456.933033 | 0.226102419 | 1.69E-06 | 3.29E-05 |
| ENSMUST00000193703.5,Enah-003,protein_coding,ENSMUSG00000022995.16,protein_coding | 3132.434243 | -0.359492483 | 1.69E-06 | 3.30E-05 |
| ENSMUST00000160922.4,Luc7l2-008,retained_intron,ENSMUSG00000029823.16,protein_coding | 1279.606737 | -0.691733722 | 1.69E-06 | 3.30E-05 |
| ENSMUST00000149796.1,Rer1-002,processed_transcript,ENSMUSG00000029048.3,protein_coding | 8288.047359 | 0.212945793 | 1.70E-06 | 3.32E-05 |
| ENSMUST00000106355.4,Zfp691-002,protein_coding,ENSMUSG00000045268.13,protein_coding | 366.7353683 | 0.558809345 | 1.71E-06 | 3.34E-05 |

| | | | | |
|---|---|---|---|---|
| ENSMUST00000021726.6,Adssl1-201,protein_coding,ENSMUSG00000011148.13,protein_coding | 1526.033933 | 0.229982828 | 1.73E-06 | 3.37E-05 |
| ENSMUST00000052248.7,Eef1g-001,protein_coding,ENSMUSG00000071644.10,protein_coding | 198670.6558 | 0.229459296 | 1.74E-06 | 3.40E-05 |
| ENSMUST00000080035.10,Cd99l2-006,protein_coding,ENSMUSG00000035776.14,protein_coding | 980.8278882 | -0.555569747 | 1.75E-06 | 3.41E-05 |
| ENSMUST00000032898.7,Ipo5-201,protein_coding,ENSMUSG00000030662.9,protein_coding | 74374.07564 | 0.208775386 | 1.76E-06 | 3.42E-05 |
| ENSMUST00000110031.3,Auh-002,protein_coding,ENSMUSG00000021460.16,protein_coding | 326.5724188 | 0.471330789 | 1.77E-06 | 3.44E-05 |
| ENSMUST00000086829.10,Bicd1-001,protein_coding,ENSMUSG00000003452.15,protein_coding | 724.6216505 | -0.73110109 | 1.77E-06 | 3.44E-05 |
| ENSMUST00000206090.1,Ino80e-010,protein_coding,ENSMUSG00000030689.15,protein_coding | 120.256375 | -0.675074582 | 1.77E-06 | 3.44E-05 |
| ENSMUST00000021077.3,Slc9a3r1-001,protein_coding,ENSMUSG00000020733.3,protein_coding | 13281.13659 | -0.241704917 | 1.77E-06 | 3.44E-05 |
| ENSMUST00000107166.7,Tenm4-009,protein_coding,ENSMUSG00000048078.16,protein_coding | 223.204948 | 0.787631317 | 1.78E-06 | 3.45E-05 |
| ENSMUST00000103226.9,Gabpb1-008,protein_coding,ENSMUSG00000027361.15,protein_coding | 1769.529152 | 0.324227832 | 1.78E-06 | 3.46E-05 |
| ENSMUST00000114976.8,Luc7l-003,protein_coding,ENSMUSG00000024188.15,protein_coding | 4661.216305 | 0.194553555 | 1.78E-06 | 3.46E-05 |
| ENSMUST00000050909.6,Amigo1-201,protein_coding,ENSMUSG00000050947.9,protein_coding | 592.260497 | -0.326427793 | 1.79E-06 | 3.47E-05 |
| ENSMUST00000031333.3,Gtf2h3-001,protein_coding,ENSMUSG00000029387.10,protein_coding | 2579.232141 | 0.270360055 | 1.79E-06 | 3.47E-05 |
| ENSMUST00000113355.8,Frmd4b-001,protein_coding,ENSMUSG00000030064.16,protein_coding | 2604.522506 | -0.34667877 | 1.81E-06 | 3.51E-05 |
| ENSMUST00000087956.5,Med12-003,protein_coding,ENSMUSG00000079487.11,protein_coding | 968.1029205 | 0.607956496 | 1.81E-06 | 3.51E-05 |
| ENSMUST00000100251.8,Rxra-002,protein_coding,ENSMUSG00000015846.14,protein_coding | 636.7480166 | 0.506051371 | 1.83E-06 | 3.54E-05 |
| ENSMUST00000139477.7,Gid4-002,nonsense_mediated_decay,ENSMUSG00000018415.14,protein_coding | 63.31629622 | 0.85535297 | 1.84E-06 | 3.57E-05 |
| ENSMUST00000122284.7,Dyrk1a-002,protein_coding,ENSMUSG00000022897.14,protein_coding | 2794.241324 | 0.246299369 | 1.85E-06 | 3.58E-05 |
| ENSMUST00000214571.1,Pkm-002,retained_intron,ENSMUSG00000032294.17,protein_coding | 2866.09359 | -0.277708346 | 1.86E-06 | 3.60E-05 |
| ENSMUST00000131668.2,Rusc2-004,protein_coding,ENSMUSG00000035969.15,protein_coding | 4996.741293 | 0.265402579 | 1.86E-06 | 3.60E-05 |
| ENSMUST00000182886.7,Sec31a-002,protein_coding,ENSMUSG00000035325.16,protein_coding | 1731.354642 | 0.740807067 | 1.86E-06 | 3.60E-05 |
| ENSMUST00000167288.7,Lck-201,protein_coding,ENSMUSG00000000409.14,protein_coding | 576.5343966 | 0.560347327 | 1.87E-06 | 3.61E-05 |
| ENSMUST00000039810.7,Xpot-201,protein_coding,ENSMUSG00000034667.7,protein_coding | 46454.71125 | 0.251106861 | 1.88E-06 | 3.63E-05 |
| ENSMUST00000096053.3,Fam178a-201,protein_coding,ENSMUSG00000036097.7,protein_coding | 7416.068409 | -0.438943226 | 1.90E-06 | 3.67E-05 |
| ENSMUST00000032748.14,Unc45a-201,protein_coding,ENSMUSG00000030533.15,protein_coding | 5397.904111 | -0.293130248 | 1.90E-06 | 3.67E-05 |
| ENSMUST00000130716.7,Mtmr3-001,nonsense_mediated_decay,ENSMUSG00000034354.17,protein_coding | 2853.90004 | 0.465133692 | 1.90E-06 | 3.67E-05 |
| ENSMUST00000135243.2,Gpi1-002,retained_intron,ENSMUSG00000036427.5,protein_coding | 1011.145829 | 0.342456232 | 1.92E-06 | 3.70E-05 |
| ENSMUST00000025841.3,Mus81-002,nonsense_mediated_decay,ENSMUSG00000024906.9,protein_coding | 101.1646151 | -0.882907389 | 1.92E-06 | 3.70E-05 |
| ENSMUST00000140975.7,Cenpo-001,protein_coding,ENSMUSG00000020652.17,protein_coding | 343.9749338 | -0.487707407 | 1.93E-06 | 3.71E-05 |
| ENSMUST00000070532.7,Fabp3-001,protein_coding,ENSMUSG00000028773.8,protein_coding | 256.8978354 | -0.480992931 | 1.95E-06 | 3.76E-05 |
| ENSMUST00000034650.14,Mcam-001,protein_coding,ENSMUSG00000032135.15,protein_coding | 733.5616455 | -0.507370385 | 1.95E-06 | 3.76E-05 |
| ENSMUST00000108375.8,Myo18a-002,protein_coding,ENSMUSG00000000631.20,protein_coding | 13314.30337 | -0.161756562 | 1.98E-06 | 3.80E-05 |
| ENSMUST00000100171.2,Hspa5-001,protein_coding,ENSMUSG00000026864.13,protein_coding | 91086.81515 | 0.319038208 | 1.99E-06 | 3.82E-05 |
| ENSMUST00000170296.1,Syce2-201,protein_coding,ENSMUSG00000003824.12,protein_coding | 1870.754649 | 0.255747136 | 1.99E-06 | 3.82E-05 |
| ENSMUST00000176363.8,Atp2c1-018,protein_coding,ENSMUSG00000032570.17,protein_coding | 97.80268427 | 0.885423463 | 1.99E-06 | 3.82E-05 |
| ENSMUST00000041623.7,Enc1-201,protein_coding,ENSMUSG00000041773.7,protein_coding | 11259.55822 | -0.132126478 | 2.00E-06 | 3.83E-05 |
| ENSMUST00000121312.1,Gata4-003,protein_coding,ENSMUSG00000021944.15,protein_coding | 154.7297712 | 0.847308451 | 2.01E-06 | 3.85E-05 |
| ENSMUST00000097653.10,Col6a3-001,protein_coding,ENSMUSG00000048126.16,protein_coding | 58.21442226 | 0.761101279 | 2.01E-06 | 3.86E-05 |
| ENSMUST00000131222.1,Gm4285-001,antisense,ENSMUSG00000085829.1,antisense | 95.24910665 | 0.688284138 | 2.02E-06 | 3.87E-05 |
| ENSMUST00000150576.7,Rpl29-001,protein_coding,ENSMUSG00000048758.14,protein_coding | 41399.12755 | 0.168839135 | 2.02E-06 | 3.87E-05 |
| ENSMUST00000126215.1,Ncbp2-004,processed_transcript,ENSMUSG00000022774.12,protein_coding | 184.4336157 | 0.7643052 | 2.03E-06 | 3.88E-05 |
| ENSMUST00000156473.7,Snx12-003,protein_coding,ENSMUSG00000046032.16,protein_coding | 976.8776263 | 0.316341775 | 2.03E-06 | 3.89E-05 |
| ENSMUST00000010807.4,Fads1-201,protein_coding,ENSMUSG00000010663.4,protein_coding | 39190.56633 | 0.228205229 | 2.03E-06 | 3.89E-05 |
| ENSMUST00000173432.2,Hoxb6-001,protein_coding,ENSMUSG00000000690.5,protein_coding | 1224.942885 | -0.323735445 | 2.04E-06 | 3.89E-05 |
| ENSMUST00000021714.7,Zfyve21-201,protein_coding,ENSMUSG00000021286.7,protein_coding | 1429.640318 | 0.283359796 | 2.04E-06 | 3.90E-05 |
| ENSMUST00000021046.5,Ddx42-001,protein_coding,ENSMUSG00000020705.6,protein_coding | 14000.27994 | 0.167739201 | 2.05E-06 | 3.92E-05 |
| ENSMUST00000119991.7,Stard4-003,protein_coding,ENSMUSG00000024378.9,protein_coding | 434.9316974 | 0.499334082 | 2.07E-06 | 3.96E-05 |
| ENSMUST00000016309.15,Tmbim1-001,protein_coding,ENSMUSG00000006301.17,protein_coding | 7724.128224 | 0.245819391 | 2.08E-06 | 3.96E-05 |
| ENSMUST00000114897.8,Atp5c1-001,protein_coding,ENSMUSG00000025781.14,protein_coding | 37018.32636 | 0.232817257 | 2.08E-06 | 3.97E-05 |
| ENSMUST00000162053.7,Psat1-002,protein_coding,ENSMUSG00000024640.9,protein_coding | 19431.92718 | 0.346731762 | 2.08E-06 | 3.97E-05 |
| ENSMUST00000160439.7,Zfand2b-003,protein_coding,ENSMUSG00000026197.12,protein_coding | 1552.264845 | 0.361344081 | 2.09E-06 | 3.98E-05 |
| ENSMUST00000065625.11,Trmt1l-001,protein_coding,ENSMUSG00000053286.11,protein_coding | 2238.67352 | -0.257763649 | 2.09E-06 | 3.98E-05 |
| ENSMUST00000182149.7,Prrc2c-001,protein_coding,ENSMUSG00000040225.15,protein_coding | 11258.14869 | -0.434299242 | 2.09E-06 | 3.99E-05 |
| ENSMUST00000121693.7,Rab27b-001,protein_coding,ENSMUSG00000024511.15,protein_coding | 203.1374856 | -0.791825853 | 2.11E-06 | 4.02E-05 |
| ENSMUST00000103176.9,Mib2-001,protein_coding,ENSMUSG00000029060.17,protein_coding | 3335.194644 | 0.470382611 | 2.14E-06 | 4.08E-05 |
| ENSMUST00000045487.3,Rhou-001,protein_coding,ENSMUSG00000039960.5,protein_coding | 4595.369486 | -0.259152037 | 2.15E-06 | 4.09E-05 |
| ENSMUST00000079063.6,Sox11-001,protein_coding,ENSMUSG00000063632.6,protein_coding | 2622.273786 | -0.37301626 | 2.15E-06 | 4.09E-05 |
| ENSMUST00000018304.6,Med1-002,protein_coding,ENSMUSG00000018160.15,protein_coding | 1554.27601 | 0.513546216 | 2.16E-06 | 4.10E-05 |
| ENSMUST00000155520.7,Bcas2-005,nonsense_mediated_decay,ENSMUSG00000005687.14,protein_coding | 1136.152677 | 0.318412997 | 2.16E-06 | 4.10E-05 |
| ENSMUST00000069763.2,Lancl3-001,protein_coding,ENSMUSG00000047344.9,protein_coding | 194.4083914 | -0.636871012 | 2.17E-06 | 4.13E-05 |
| ENSMUST00000134465.7,Phldb1-007,protein_coding,ENSMUSG00000048537.16,protein_coding | 11164.45079 | -0.227529396 | 2.17E-06 | 4.13E-05 |
| ENSMUST00000128764.7,Sap18-001,protein_coding,ENSMUSG00000021963.16,protein_coding | 4598.161492 | -0.205692995 | 2.19E-06 | 4.15E-05 |
| ENSMUST00000041003.6,Tfb1m-201,protein_coding,ENSMUSG00000036983.6,protein_coding | 860.6526359 | 0.340123913 | 2.20E-06 | 4.17E-05 |
| ENSMUST00000033325.8,Swap70-001,protein_coding,ENSMUSG00000031015.8,protein_coding | 4742.316567 | 0.30229922 | 2.22E-06 | 4.20E-05 |
| ENSMUST00000026859.10,Mfsd8-001,protein_coding,ENSMUSG00000025759.11,protein_coding | 517.8476595 | -0.352693455 | 2.22E-06 | 4.20E-05 |
| ENSMUST00000166750.8,Cmip-001,protein_coding,ENSMUSG00000034390.16,protein_coding | 5642.003466 | 0.190047167 | 2.25E-06 | 4.25E-05 |
| ENSMUST00000040308.13,Sh2b3-002,protein_coding,ENSMUSG00000042594.16,protein_coding | 2601.559493 | 0.292621387 | 2.26E-06 | 4.28E-05 |
| ENSMUST00000077376.2,2810006K23Rik-001,protein_coding,ENSMUSG00000047635.6,protein_coding | 2299.470996 | 0.311739577 | 2.27E-06 | 4.29E-05 |
| ENSMUST00000031731.13,Pcolce-001,protein_coding,ENSMUSG00000029718.14,protein_coding | 22581.53276 | -0.180568105 | 2.28E-06 | 4.32E-05 |
| ENSMUST00000054344.10,Eogt-001,protein_coding,ENSMUSG00000035245.12,protein_coding | 3935.576102 | -0.229442668 | 2.30E-06 | 4.35E-05 |
| ENSMUST00000115231.3,Rpl39-001,protein_coding,ENSMUSG00000079641.3,protein_coding | 40816.53296 | 0.207767774 | 2.31E-06 | 4.37E-05 |
| ENSMUST00000102617.4,Adprhl2-001,protein_coding,ENSMUSG00000042558.16,protein_coding | 3494.70934 | 0.248576565 | 2.34E-06 | 4.43E-05 |
| ENSMUST00000027499.12,Bok-001,protein_coding,ENSMUSG00000026278.14,protein_coding | 8718.101097 | 0.194833194 | 2.36E-06 | 4.45E-05 |
| ENSMUST00000041725.13,Mgat4b-001,protein_coding,ENSMUSG00000036620.14,protein_coding | 14543.74345 | 0.205515046 | 2.37E-06 | 4.47E-05 |
| ENSMUST00000172719.7,Dnmt3a-017,protein_coding,ENSMUSG00000020661.15,protein_coding | 114.5608368 | 0.689970537 | 2.38E-06 | 4.50E-05 |
| ENSMUST00000112832.7,Rorb-005,protein_coding,ENSMUSG00000036192.15,protein_coding | 431.1318452 | -0.589675986 | 2.40E-06 | 4.53E-05 |
| ENSMUST00000038188.13,Limch1-001,protein_coding,ENSMUSG00000037736.18,protein_coding | 364.1192759 | -0.50598179 | 2.41E-06 | 4.54E-05 |
| ENSMUST00000171737.1,Odc1-201,protein_coding,ENSMUSG00000011179.7,protein_coding | 90212.88642 | 0.212405356 | 2.43E-06 | 4.57E-05 |
| ENSMUST00000109923.8,Dbn1-002,protein_coding,ENSMUSG00000034675.17,protein_coding | 7218.518572 | -0.300389098 | 2.45E-06 | 4.61E-05 |
| ENSMUST00000190378.1,Prmt6-201,protein_coding,ENSMUSG00000049300.11,protein_coding | 786.0587973 | 0.343652781 | 2.46E-06 | 4.63E-05 |
| ENSMUST00000002007.4,Siae-005,protein_coding,ENSMUSG00000001942.8,protein_coding | 457.9242861 | 0.363261669 | 2.46E-06 | 4.63E-05 |
| ENSMUST00000042988.6,Atg14-201,protein_coding,ENSMUSG00000037526.6,protein_coding | 1855.392687 | 0.34893597 | 2.48E-06 | 4.66E-05 |
| ENSMUST00000106100.8,Zzz3-001,protein_coding,ENSMUSG00000039068.16,protein_coding | 3403.810025 | -0.461898124 | 2.48E-06 | 4.67E-05 |
| ENSMUST00000125959.1,Dynll1-002,retained_intron,ENSMUSG00000009013.5,protein_coding | 3302.260772 | -0.381519326 | 2.48E-06 | 4.67E-05 |
| ENSMUST00000030791.11,Smarcd3-001,protein_coding,ENSMUSG00000028949.13,protein_coding | 2098.973128 | -0.301599827 | 2.50E-06 | 4.69E-05 |
| ENSMUST00000131260.2,1810032O08Rik-002,nonsense_mediated_decay,ENSMUSG00000020812.17,protein_coding | 163.4067203 | 0.699874907 | 2.51E-06 | 4.72E-05 |
| ENSMUST00000088737.10,Grk2-001,protein_coding,ENSMUSG00000024858.13,protein_coding | 22377.83649 | 0.181176876 | 2.52E-06 | 4.73E-05 |
| ENSMUST00000111325.3,Sdr39u1-201,protein_coding,ENSMUSG00000022223.8,protein_coding | 3486.804372 | -0.247442542 | 2.52E-06 | 4.73E-05 |
| ENSMUST00000160429.7,Gas5-014,retained_intron,ENSMUSG00000053332.13,processed_transcript | 509.4859805 | -0.615624253 | 2.53E-06 | 4.74E-05 |
| ENSMUST00000030457.11,Nasp-002,protein_coding,ENSMUSG00000028693.15,protein_coding | 3657.418239 | -0.35798407 | 2.54E-06 | 4.75E-05 |
| ENSMUST00000119368.7,Son-003,protein_coding,ENSMUSG00000022961.17,protein_coding | 5943.720736 | -0.216493739 | 2.54E-06 | 4.76E-05 |
| ENSMUST00000112493.7,Rps6ka3-002,protein_coding,ENSMUSG00000031309.15,protein_coding | 1412.332548 | 0.400329705 | 2.54E-06 | 4.76E-05 |

| | | | | |
|---|---|---|---|---|
| ENSMUST00000211440.1,Ruvbl2-008,nonsense_mediated_decay,ENSMUSG00000003868.14,protein_coding | 183.7205409 | 0.69106179 | 2.55E-06 | 4.77E-05 |
| ENSMUST00000054487.8,Ajuba-201,protein_coding,ENSMUSG00000022178.10,protein_coding | 9674.194377 | -0.220439787 | 2.56E-06 | 4.80E-05 |
| ENSMUST00000191150.1,2310035C23Rik-008,processed_transcript,ENSMUSG00000026319.13,protein_coding | 187.0764291 | -0.589615819 | 2.58E-06 | 4.82E-05 |
| ENSMUST00000045909.7,Zbed3-201,protein_coding,ENSMUSG00000041995.7,protein_coding | 7292.299806 | 0.15304103 | 2.61E-06 | 4.88E-05 |
| ENSMUST00000009550.13,Elk1-001,protein_coding,ENSMUSG00000009406.13,protein_coding | 4653.293615 | 0.213057033 | 2.62E-06 | 4.89E-05 |
| ENSMUST00000108612.7,Eif5a-005,protein_coding,ENSMUSG00000078812.10,protein_coding | 5528.312081 | 0.360445376 | 2.63E-06 | 4.92E-05 |
| ENSMUST00000022280.10,Smg9-001,protein_coding,ENSMUSG00000022210.11,protein_coding | 3712.158992 | 0.186493762 | 2.64E-06 | 4.92E-05 |
| ENSMUST00000004072.8,Rpl8-201,protein_coding,ENSMUSG00000003970.8,protein_coding | 288566.1936 | 0.224691855 | 2.63E-06 | 4.92E-05 |
| ENSMUST00000018186.15,Cyb5r3-001,protein_coding,ENSMUSG00000018042.17,protein_coding | 39524.29563 | 0.168421738 | 2.64E-06 | 4.92E-05 |
| ENSMUST00000120346.7,BC004004-002,protein_coding,ENSMUSG00000052712.16,protein_coding | 2472.924599 | 0.341774986 | 2.64E-06 | 4.92E-05 |
| ENSMUST00000014220.14,Tctex1d2-001,protein_coding,ENSMUSG00000014075.15,protein_coding | 564.48677 | -0.520188522 | 2.64E-06 | 4.92E-05 |
| ENSMUST00000139585.7,Wt1-004,protein_coding,ENSMUSG00000016458.14,protein_coding | 1060.966587 | -0.299134625 | 2.65E-06 | 4.94E-05 |
| ENSMUST00000184914.1,Slc25a37-003,nonsense_mediated_decay,ENSMUSG00000034248.7,protein_coding | 870.2549206 | 0.477179741 | 2.66E-06 | 4.95E-05 |
| ENSMUST00000188299.1,Gm28438-001,unprocessed_pseudogene,ENSMUSG00000101939.1,unprocessed_pseudogene | 27100.72704 | 0.399872962 | 2.66E-06 | 4.95E-05 |
| ENSMUST00000161668.7,Lmo7-012,processed_transcript,ENSMUSG00000033060.15,protein_coding | 394.285335 | -0.388210335 | 2.67E-06 | 4.97E-05 |
| ENSMUST00000110455.3,Hist1h2bk-001,protein_coding,ENSMUSG00000062727.4,protein_coding | 493.3314083 | 0.391163418 | 2.68E-06 | 4.98E-05 |
| ENSMUST00000069476.4,Ldoc1l-201,protein_coding,ENSMUSG00000055745.4,protein_coding | 659.6137798 | -0.342620966 | 2.68E-06 | 4.98E-05 |
| ENSMUST00000106441.7,Trim47-001,protein_coding,ENSMUSG00000020773.11,protein_coding | 3481.431961 | 0.323725415 | 2.70E-06 | 5.01E-05 |
| ENSMUST00000161204.7,Rdh11-001,protein_coding,ENSMUSG00000066441.14,protein_coding | 4825.156257 | 0.294258401 | 2.70E-06 | 5.02E-05 |
| ENSMUST00000135887.2,Igfbp3-002,protein_coding,ENSMUSG00000020427.11,protein_coding | 433.3665676 | 0.687733807 | 2.71E-06 | 5.03E-05 |
| ENSMUST00000108531.7,Nup88-001,protein_coding,ENSMUSG00000040667.11,protein_coding | 7222.060567 | -0.169179838 | 2.71E-06 | 5.03E-05 |
| ENSMUST00000192757.5,Mbnl1-025,protein_coding,ENSMUSG00000027763.13,protein_coding | 77.03439732 | -0.783906539 | 2.72E-06 | 5.04E-05 |
| ENSMUST00000121326.7,Srsf11-001,protein_coding,ENSMUSG00000055436.18,protein_coding | 1404.165525 | -0.471297562 | 2.72E-06 | 5.04E-05 |
| ENSMUST00000039280.7,Scaf4-201,protein_coding,ENSMUSG00000022983.15,protein_coding | 6377.650784 | 0.156897651 | 2.72E-06 | 5.04E-05 |
| ENSMUST00000025915.11,Dnajc4-201,protein_coding,ENSMUSG00000024963.11,protein_coding | 973.8002248 | 0.308519837 | 2.73E-06 | 5.06E-05 |
| ENSMUST00000110492.1,D8Ertd82e-001,protein_coding,ENSMUSG00000050271.12,protein_coding | 2898.588 | 0.199273456 | 2.75E-06 | 5.09E-05 |
| ENSMUST00000144124.1,Tulp4-011,processed_transcript,ENSMUSG00000034377.10,protein_coding | 753.7686449 | 0.431243419 | 2.76E-06 | 5.10E-05 |
| ENSMUST00000049744.3,Mplkip-201,protein_coding,ENSMUSG00000012429.8,protein_coding | 1784.804008 | 0.222092322 | 2.76E-06 | 5.11E-05 |
| ENSMUST00000178400.7,Ccdc191-201,protein_coding,ENSMUSG00000022701.17,protein_coding | 736.7755945 | 0.468225243 | 2.77E-06 | 5.12E-05 |
| ENSMUST00000086075.12,Scarb1-001,protein_coding,ENSMUSG00000037936.15,protein_coding | 9141.746295 | 0.192797681 | 2.79E-06 | 5.16E-05 |
| ENSMUST00000022894.13,Ywhaz-002,protein_coding,ENSMUSG00000022285.16,protein_coding | 75340.5684 | 0.125425492 | 2.82E-06 | 5.21E-05 |
| ENSMUST00000117360.7,Litaf-002,protein_coding,ENSMUSG00000022500.14,protein_coding | 375.2118048 | 0.527225329 | 2.84E-06 | 5.24E-05 |
| ENSMUST00000148960.7,Myl12a-001,protein_coding,ENSMUSG00000024048.14,protein_coding | 29839.70466 | 0.196267463 | 2.86E-06 | 5.29E-05 |
| ENSMUST00000204097.2,Klhl5-001,protein_coding,ENSMUSG00000054920.12,protein_coding | 5270.981342 | 0.191789952 | 2.87E-06 | 5.29E-05 |
| ENSMUST00000126136.7,Hes6-005,retained_intron,ENSMUSG00000067071.8,protein_coding | 170.0074654 | -0.731399121 | 2.88E-06 | 5.32E-05 |
| ENSMUST00000044825.4,Ndufaf5-001,protein_coding,ENSMUSG00000027384.6,protein_coding | 1795.35256 | 0.186098594 | 2.89E-06 | 5.33E-05 |
| ENSMUST00000040280.13,Slc25a23-001,protein_coding,ENSMUSG00000046329.14,protein_coding | 558.3552401 | -0.366819985 | 2.90E-06 | 5.35E-05 |
| ENSMUST00000121661.7,Smim14-004,protein_coding,ENSMUSG00000037822.12,protein_coding | 3281.079737 | 0.281075865 | 2.91E-06 | 5.36E-05 |
| ENSMUST00000078665.12,Dhps-001,protein_coding,ENSMUSG00000060038.14,protein_coding | 5183.034642 | 0.228345638 | 2.92E-06 | 5.38E-05 |
| ENSMUST00000163313.1,Baalc-201,protein_coding,ENSMUSG00000022296.8,protein_coding | 411.5431291 | -0.537609788 | 2.92E-06 | 5.38E-05 |
| ENSMUST00000044113.10,Ago2-001,protein_coding,ENSMUSG00000036698.10,protein_coding | 38479.41666 | 0.129277613 | 2.93E-06 | 5.39E-05 |
| ENSMUST00000052355.13,Nup37-201,protein_coding,ENSMUSG00000035351.15,protein_coding | 5773.8178 | 0.224656745 | 2.93E-06 | 5.39E-05 |
| ENSMUST00000056950.13,Rin3-001,protein_coding,ENSMUSG00000044456.16,protein_coding | 2460.133263 | 0.255311464 | 2.95E-06 | 5.42E-05 |
| ENSMUST00000031291.8,Asphd2-001,protein_coding,ENSMUSG00000029348.11,protein_coding | 622.1913874 | -0.357244962 | 2.96E-06 | 5.43E-05 |
| ENSMUST00000157028.1,Arfip2-009,nonsense_mediated_decay,ENSMUSG00000030881.15,protein_coding | 205.0419229 | 0.567344491 | 2.97E-06 | 5.45E-05 |
| ENSMUST00000063117.8,Gstz1-201,protein_coding,ENSMUSG00000021033.10,protein_coding | 3428.935796 | 0.274734671 | 3.00E-06 | 5.51E-05 |
| ENSMUST00000153397.7,Hira-003,nonsense_mediated_decay,ENSMUSG00000022702.15,protein_coding | 255.3270035 | 0.819937966 | 3.02E-06 | 5.55E-05 |
| ENSMUST00000006774.10,Gtf2h1-001,protein_coding,ENSMUSG00000006599.13,protein_coding | 4978.729639 | -0.26242516 | 3.06E-06 | 5.62E-05 |
| ENSMUST00000143096.1,Etfb-002,processed_transcript,ENSMUSG00000004610.4,protein_coding | 37.11579602 | 0.849796337 | 3.07E-06 | 5.63E-05 |
| ENSMUST00000203290.2,Zyx-003,retained_intron,ENSMUSG00000029860.16,protein_coding | 687.377964 | -0.369620117 | 3.07E-06 | 5.63E-05 |
| ENSMUST00000057692.10,Luc7l2-001,protein_coding,ENSMUSG00000029823.16,protein_coding | 9179.870649 | 0.269671816 | 3.09E-06 | 5.66E-05 |
| ENSMUST00000117506.8,Ube4a-003,protein_coding,ENSMUSG00000059890.16,protein_coding | 2586.655647 | -0.287558488 | 3.09E-06 | 5.67E-05 |
| ENSMUST00000020153.8,Hbs1l-201,protein_coding,ENSMUSG00000019977.15,protein_coding | 15394.58194 | 0.134910669 | 3.10E-06 | 5.67E-05 |
| ENSMUST00000048490.11,Larp1b-002,protein_coding,ENSMUSG00000025762.14,protein_coding | 1115.694987 | 0.279225411 | 3.11E-06 | 5.69E-05 |
| ENSMUST00000023497.2,Lmln-001,protein_coding,ENSMUSG00000022802.2,protein_coding | 565.9596045 | -0.368353627 | 3.12E-06 | 5.71E-05 |
| ENSMUST00000020248.15,Ccdc53-001,protein_coding,ENSMUSG00000020056.16,protein_coding | 1482.217888 | 0.285977731 | 3.18E-06 | 5.80E-05 |
| ENSMUST00000024811.7,Pim1-001,protein_coding,ENSMUSG00000024014.7,protein_coding | 5700.774284 | 0.293842228 | 3.18E-06 | 5.80E-05 |
| ENSMUST00000141630.7,Tyms-002,nonsense_mediated_decay,ENSMUSG00000025747.12,protein_coding | 455.5300967 | -0.613649427 | 3.18E-06 | 5.81E-05 |
| ENSMUST00000037918.11,Tmem33-001,protein_coding,ENSMUSG00000037720.16,protein_coding | 2842.955547 | -0.612151833 | 3.22E-06 | 5.88E-05 |
| ENSMUST00000027629.9,Tfcp2l1-001,protein_coding,ENSMUSG00000026380.10,protein_coding | 403.9672196 | 0.582784153 | 3.23E-06 | 5.90E-05 |
| ENSMUST00000160281.7,Pole4-005,protein_coding,ENSMUSG00000030042.15,protein_coding | 1861.633601 | 0.326201994 | 3.25E-06 | 5.92E-05 |
| ENSMUST00000064495.7,Hook2-001,protein_coding,ENSMUSG00000052566.8,protein_coding | 2131.616085 | 0.334181539 | 3.24E-06 | 5.92E-05 |
| ENSMUST00000095062.9,Lmbrd1-002,protein_coding,ENSMUSG00000073725.8,protein_coding | 669.2926624 | -0.50196917 | 3.25E-06 | 5.92E-05 |
| ENSMUST00000112536.7,Zfp932-003,protein_coding,ENSMUSG00000066613.14,protein_coding | 99.71276209 | -0.705182459 | 3.26E-06 | 5.95E-05 |
| ENSMUST00000018246.4,Hist1h2bc-001,protein_coding,ENSMUSG00000018102.4,protein_coding | 1580.954896 | -0.402432913 | 3.28E-06 | 5.97E-05 |
| ENSMUST00000155805.1,Rps16-003,retained_intron,ENSMUSG00000037563.15,protein_coding | 41830.58051 | 0.247100928 | 3.28E-06 | 5.97E-05 |
| ENSMUST00000046908.9,Oser1-001,protein_coding,ENSMUSG00000035399.12,protein_coding | 4380.678337 | 0.217830969 | 3.28E-06 | 5.97E-05 |
| ENSMUST00000145839.1,Nf1-005,processed_transcript,ENSMUSG00000020716.15,protein_coding | 382.741878 | 0.40884512 | 3.29E-06 | 5.98E-05 |
| ENSMUST00000116621.1,Ndufb11-001,protein_coding,ENSMUSG00000031059.9,protein_coding | 10183.02813 | 0.212157937 | 3.30E-06 | 6.00E-05 |
| ENSMUST00000134458.7,Macf1-008,protein_coding,ENSMUSG00000028649.17,protein_coding | 3349.775982 | -0.472855757 | 3.31E-06 | 6.01E-05 |
| ENSMUST00000154790.1,Tk1-006,retained_intron,ENSMUSG00000025574.13,protein_coding | 256.7636301 | 0.487779408 | 3.31E-06 | 6.01E-05 |
| ENSMUST00000004156.8,Map3k11-201,protein_coding,ENSMUSG00000004054.8,protein_coding | 7261.288286 | -0.14741992 | 3.33E-06 | 6.05E-05 |
| ENSMUST00000040259.7,Atp6v0a4-001,protein_coding,ENSMUSG00000038600.12,protein_coding | 458.6969133 | -0.765925726 | 3.34E-06 | 6.06E-05 |
| ENSMUST00000015124.14,Tsen15-001,protein_coding,ENSMUSG00000014980.14,protein_coding | 2854.794671 | 0.296390911 | 3.34E-06 | 6.06E-05 |
| ENSMUST00000034276.12,Cox4i1-001,protein_coding,ENSMUSG00000031818.12,protein_coding | 35265.25085 | 0.214110295 | 3.36E-06 | 6.10E-05 |
| ENSMUST00000064639.14,Ubap2l-004,protein_coding,ENSMUSG00000042520.16,protein_coding | 11262.92934 | 0.216314544 | 3.36E-06 | 6.10E-05 |
| ENSMUST00000022808.13,Pabpn1-001,protein_coding,ENSMUSG00000022194.15,protein_coding | 13103.44267 | 0.163310567 | 3.39E-06 | 6.13E-05 |
| ENSMUST00000133449.7,Snhg17-001,lincRNA,ENSMUSG00000085385.7,processed_transcript | 555.4011287 | 0.503283732 | 3.40E-06 | 6.15E-05 |
| ENSMUST00000142127.7,Mettl5-002,nonsense_mediated_decay,ENSMUSG00000051730.15,protein_coding | 234.4940669 | -0.712134158 | 3.41E-06 | 6.16E-05 |
| ENSMUST00000036877.9,Dennd2a-001,protein_coding,ENSMUSG00000038456.9,protein_coding | 2825.203195 | -0.18662148 | 3.41E-06 | 6.16E-05 |
| ENSMUST00000034049.4,Slc25a4-001,protein_coding,ENSMUSG00000031633.4,protein_coding | 43632.26323 | -0.204378068 | 3.40E-06 | 6.16E-05 |
| ENSMUST00000055704.11,Gnai2-001,protein_coding,ENSMUSG00000032562.13,protein_coding | 44782.31086 | 0.184821069 | 3.41E-06 | 6.16E-05 |
| ENSMUST00000177669.1,Pfn1-003,retained_intron,ENSMUSG00000018293.4,protein_coding | 1386.85127 | -0.306913081 | 3.41E-06 | 6.17E-05 |
| ENSMUST00000156921.1,Rpl27a-003,nonsense_mediated_decay,ENSMUSG00000046364.14,protein_coding | 4141.840543 | 0.301057773 | 3.42E-06 | 6.17E-05 |
| ENSMUST00000201688.3,Ubl3-005,processed_transcript,ENSMUSG00000001687.15,protein_coding | 921.4255063 | 0.348851902 | 3.42E-06 | 6.18E-05 |
| ENSMUST00000096194.8,Tmem2-202,protein_coding,ENSMUSG00000024754.12,protein_coding | 1207.321267 | -0.420990712 | 3.42E-06 | 6.18E-05 |
| ENSMUST00000049241.8,Hoxb4-001,protein_coding,ENSMUSG00000038692.8,protein_coding | 1101.303696 | -0.243090762 | 3.44E-06 | 6.20E-05 |
| ENSMUST00000171384.7,Tm4sf1-201,protein_coding,ENSMUSG00000027800.14,protein_coding | 10904.18797 | -0.378123738 | 3.44E-06 | 6.20E-05 |
| ENSMUST00000023487.4,Arhgap31-001,protein_coding,ENSMUSG00000022799.4,protein_coding | 2599.79914 | -0.277294289 | 3.44E-06 | 6.20E-05 |
| ENSMUST00000058845.7,Basp1-201,protein_coding,ENSMUSG00000045763.7,protein_coding | 5719.029813 | -0.260150191 | 3.44E-06 | 6.20E-05 |
| ENSMUST00000025739.13,Uhrf2-001,protein_coding,ENSMUSG00000024817.13,protein_coding | 15757.83136 | 0.239374014 | 3.45E-06 | 6.21E-05 |
| ENSMUST00000080371.6,Mtss1-201,protein_coding,ENSMUSG00000022353.9,protein_coding | 1151.95641 | -0.232252419 | 3.47E-06 | 6.25E-05 |

| | | | | |
|---|---|---|---|---|
| ENSMUST00000013220.7,Amotl1-001,protein_coding,ENSMUSG00000013076.16,protein_coding | 21060.80698 | 0.252773339 | 3.49E-06 | 6.28E-05 |
| ENSMUST00000055433.4,Spsb4-001,protein_coding,ENSMUSG00000046997.5,protein_coding | 94.79056865 | -0.672168092 | 3.49E-06 | 6.28E-05 |
| ENSMUST00000161989.1,Ppp3cb-002,protein_coding,ENSMUSG00000021816.10,protein_coding | 1987.307586 | -0.199933674 | 3.50E-06 | 6.29E-05 |
| ENSMUST00000118729.7,Zfp740-002,protein_coding,ENSMUSG00000046897.16,protein_coding | 11321.85341 | 0.268052905 | 3.50E-06 | 6.29E-05 |
| ENSMUST00000024846.12,Myl12a-006,protein_coding,ENSMUSG00000024048.14,protein_coding | 60741.496 | 0.201654666 | 3.50E-06 | 6.29E-05 |
| ENSMUST00000064407.9,Ikbkg-006,protein_coding,ENSMUSG00000004221.16,protein_coding | 5615.227075 | -0.142195865 | 3.50E-06 | 6.29E-05 |
| ENSMUST00000105584.9,Acap3-201,protein_coding,ENSMUSG00000029033.16,protein_coding | 3408.283824 | -0.27521262 | 3.51E-06 | 6.30E-05 |
| ENSMUST00000097950.8,Pde4b-002,protein_coding,ENSMUSG00000028525.16,protein_coding | 530.5358341 | -0.461464766 | 3.54E-06 | 6.36E-05 |
| ENSMUST00000065512.10,Rtkn-001,protein_coding,ENSMUSG00000034930.16,protein_coding | 2668.649277 | 0.288476606 | 3.55E-06 | 6.37E-05 |
| ENSMUST00000168195.7,Runx1-004,protein_coding,ENSMUSG00000022952.16,protein_coding | 891.0528626 | 0.428304146 | 3.57E-06 | 6.40E-05 |
| ENSMUST00000031542.12,Oasl2-001,protein_coding,ENSMUSG00000029561.17,protein_coding | 709.4425532 | 0.481317216 | 3.57E-06 | 6.41E-05 |
| ENSMUST00000154340.7,Rpl12-003,retained_intron,ENSMUSG00000038900.17,protein_coding | 2597.970352 | 0.259761114 | 3.60E-06 | 6.44E-05 |
| ENSMUST00000175674.1,Gm20667-001,sense_overlapping,ENSMUSG00000093726.1,sense_overlapping | 348.1957396 | -0.439052112 | 3.62E-06 | 6.47E-05 |
| ENSMUST00000074285.7,Meis2-014,protein_coding,ENSMUSG00000027210.20,protein_coding | 1266.591991 | 0.407280866 | 3.64E-06 | 6.51E-05 |
| ENSMUST00000073781.11,E2f7-001,protein_coding,ENSMUSG00000020185.16,protein_coding | 4020.840185 | -0.256094151 | 3.64E-06 | 6.52E-05 |
| ENSMUST00000116280.8,Knop1-201,protein_coding,ENSMUSG00000030980.17,protein_coding | 6860.859132 | -0.264558788 | 3.66E-06 | 6.54E-05 |
| ENSMUST00000035166.11,Uba5-001,protein_coding,ENSMUSG00000032557.12,protein_coding | 2536.888898 | -0.215135517 | 3.66E-06 | 6.54E-05 |
| ENSMUST00000063517.4,Spats2-201,protein_coding,ENSMUSG00000051934.4,protein_coding | 14381.63078 | 0.237730761 | 3.66E-06 | 6.54E-05 |
| ENSMUST00000205539.1,Snhg1-002,retained_intron,ENSMUSG00000108414.1,processed_transcript | 571.3731825 | -0.511397302 | 3.66E-06 | 6.54E-05 |
| ENSMUST00000057216.14,Mtch2-005,processed_transcript,ENSMUSG00000027282.17,protein_coding | 2937.90612 | 0.261504983 | 3.68E-06 | 6.56E-05 |
| ENSMUST00000031224.14,Tgfbr3-001,protein_coding,ENSMUSG00000029287.14,protein_coding | 1343.076411 | 0.267675999 | 3.68E-06 | 6.56E-05 |
| ENSMUST00000136758.1,Sugp2-003,retained_intron,ENSMUSG00000036054.15,protein_coding | 329.6189434 | 0.510846346 | 3.68E-06 | 6.56E-05 |
| ENSMUST00000086294.10,Ppp1cc-002,protein_coding,ENSMUSG00000004455.16,protein_coding | 9760.99515 | 0.237755809 | 3.69E-06 | 6.58E-05 |
| ENSMUST00000130216.1,Srsf3-001,protein_coding,ENSMUSG00000071172.12,protein_coding | 16051.64212 | -0.415805332 | 3.71E-06 | 6.61E-05 |
| ENSMUST00000032422.5,Creld1-001,protein_coding,ENSMUSG00000030284.11,protein_coding | 5473.846545 | 0.201775955 | 3.71E-06 | 6.61E-05 |
| ENSMUST00000111773.9,Rbms3-001,protein_coding,ENSMUSG00000039607.15,protein_coding | 553.4564625 | -0.50239956 | 3.73E-06 | 6.65E-05 |
| ENSMUST00000102588.9,Tmem39b-001,protein_coding,ENSMUSG00000053730.15,protein_coding | 1967.59385 | 0.238146495 | 3.75E-06 | 6.67E-05 |
| ENSMUST00000089217.10,Uba1-001,protein_coding,ENSMUSG00000001924.15,protein_coding | 13325.27488 | 0.371243293 | 3.76E-06 | 6.69E-05 |
| ENSMUST00000062740.13,Acp1-201,protein_coding,ENSMUSG00000044573.14,protein_coding | 9072.078484 | 0.203886149 | 3.76E-06 | 6.69E-05 |
| ENSMUST00000176528.7,Arhgef2-027,processed_transcript,ENSMUSG00000028059.15,protein_coding | 186.7224557 | 0.772616494 | 3.77E-06 | 6.70E-05 |
| ENSMUST00000022176.14,Hmgcr-001,protein_coding,ENSMUSG00000021670.14,protein_coding | 20564.50162 | 0.268071369 | 3.78E-06 | 6.72E-05 |
| ENSMUST00000117299.8,Selenoh-001,protein_coding,ENSMUSG00000076437.10,protein_coding | 9631.552971 | 0.216563896 | 3.78E-06 | 6.72E-05 |
| ENSMUST00000036350.2,Cdnf-001,protein_coding,ENSMUSG00000039496.8,protein_coding | 146.8024003 | 0.611003556 | 3.81E-06 | 6.76E-05 |
| ENSMUST00000000756.5,Rpl13-001,protein_coding,ENSMUSG00000000740.12,protein_coding | 166796.9763 | 0.20715167 | 3.84E-06 | 6.81E-05 |
| ENSMUST00000023747.13,Nckap5l-001,protein_coding,ENSMUSG00000023009.14,protein_coding | 3331.457393 | -0.179594409 | 3.86E-06 | 6.85E-05 |
| ENSMUST00000115453.1,Cav1-002,protein_coding,ENSMUSG00000007655.16,protein_coding | 3204.095136 | 0.292560337 | 3.88E-06 | 6.89E-05 |
| ENSMUST00000032335.12,Atf7ip-001,protein_coding,ENSMUSG00000030213.12,protein_coding | 10355.14666 | -0.122505929 | 3.89E-06 | 6.89E-05 |
| ENSMUST00000060531.15,Lrrc8d-001,protein_coding,ENSMUSG00000046079.16,protein_coding | 2577.111704 | 0.280130602 | 3.90E-06 | 6.90E-05 |
| ENSMUST00000125280.7,Zdhhc13-002,protein_coding,ENSMUSG00000030471.17,protein_coding | 89.58268369 | -0.860808267 | 3.90E-06 | 6.90E-05 |
| ENSMUST00000113997.8,Ppp6r3-202,protein_coding,ENSMUSG00000024908.14,protein_coding | 12172.97688 | -0.221104581 | 3.90E-06 | 6.90E-05 |
| ENSMUST00000182540.1,Pdlim1-005,retained_intron,ENSMUSG00000055044.12,protein_coding | 3064.184339 | 0.312484625 | 3.92E-06 | 6.94E-05 |
| ENSMUST00000008032.13,Crlf1-001,protein_coding,ENSMUSG00000007888.15,protein_coding | 7121.096983 | -0.230444998 | 3.95E-06 | 7.00E-05 |
| ENSMUST00000087315.13,Vars-002,protein_coding,ENSMUSG00000007029.16,protein_coding | 72948.11853 | 0.219371258 | 3.96E-06 | 7.01E-05 |
| ENSMUST00000111558.7,Ablim1-020,protein_coding,ENSMUSG00000025085.16,protein_coding | 35.79728271 | -0.85907648 | 3.97E-06 | 7.01E-05 |
| ENSMUST00000199014.4,Hax1-002,retained_intron,ENSMUSG00000027944.13,protein_coding | 763.1904089 | 0.43490247 | 3.97E-06 | 7.01E-05 |
| ENSMUST00000128518.1,Gm12854-001,processed_pseudogene,ENSMUSG00000078238.4,processed_pseudogene | 322.7884989 | -0.586762024 | 3.97E-06 | 7.01E-05 |
| ENSMUST00000001460.13,Snd1-001,protein_coding,ENSMUSG00000001424.14,protein_coding | 63334.02482 | 0.214964653 | 3.97E-06 | 7.01E-05 |
| ENSMUST00000040561.5,Rundc1-001,protein_coding,ENSMUSG00000035007.5,protein_coding | 1867.203854 | -0.224557988 | 3.97E-06 | 7.01E-05 |
| ENSMUST00000028735.7,Jag1-001,protein_coding,ENSMUSG00000027276.7,protein_coding | 1681.904481 | 0.350529897 | 3.98E-06 | 7.03E-05 |
| ENSMUST00000112720.7,Apex2-001,protein_coding,ENSMUSG00000025269.16,protein_coding | 349.4857421 | -0.408783047 | 4.01E-06 | 7.08E-05 |
| ENSMUST00000190630.6,Tardbp-021,protein_coding,ENSMUSG00000041459.15,protein_coding | 218.3734145 | 0.768403612 | 4.03E-06 | 7.10E-05 |
| ENSMUST00000064571.10,Afap1-001,protein_coding,ENSMUSG00000029094.12,protein_coding | 5363.501523 | 0.277815389 | 4.03E-06 | 7.11E-05 |
| ENSMUST00000170939.1,Ptk2-202,protein_coding,ENSMUSG00000022607.14,protein_coding | 2170.257197 | 0.4153234 | 4.04E-06 | 7.11E-05 |
| ENSMUST00000027601.10,Mcm6-001,protein_coding,ENSMUSG00000026355.11,protein_coding | 39237.87234 | 0.200392968 | 4.05E-06 | 7.13E-05 |
| ENSMUST00000173357.7,Nelfe-001,protein_coding,ENSMUSG00000024369.16,protein_coding | 4322.918009 | 0.25848866 | 4.06E-06 | 7.14E-05 |
| ENSMUST00000142664.2,P2rx4-004,protein_coding,ENSMUSG00000029470.15,protein_coding | 409.7063487 | 0.55669956 | 4.09E-06 | 7.20E-05 |
| ENSMUST00000217434.1,Dpp8-003,nonsense_mediated_decay,ENSMUSG00000032393.15,protein_coding | 609.9622661 | -0.494246248 | 4.10E-06 | 7.21E-05 |
| ENSMUST00000026986.6,Higd2a-001,protein_coding,ENSMUSG00000025868.6,protein_coding | 5291.4727 | 0.178173075 | 4.11E-06 | 7.22E-05 |
| ENSMUST00000201702.3,Syngap1-004,protein_coding,ENSMUSG00000067629.11,protein_coding | 317.3099618 | -0.432847293 | 4.12E-06 | 7.24E-05 |
| ENSMUST00000105692.1,Ctnnbip1-002,protein_coding,ENSMUSG00000028988.13,protein_coding | 627.2247742 | 0.3900442 | 4.13E-06 | 7.25E-05 |
| ENSMUST00000037099.8,Clic4-001,protein_coding,ENSMUSG00000037242.8,protein_coding | 90975.847 | 0.189134958 | 4.13E-06 | 7.26E-05 |
| ENSMUST00000018522.12,Cuedc1-001,protein_coding,ENSMUSG00000018378.13,protein_coding | 4600.974565 | 0.203488894 | 4.13E-06 | 7.26E-05 |
| ENSMUST00000107409.4,Mfge8-002,protein_coding,ENSMUSG00000030605.15,protein_coding | 13408.68798 | 0.20115983 | 4.16E-06 | 7.29E-05 |
| ENSMUST00000065162.10,Fam53a-006,protein_coding,ENSMUSG00000037339.17,protein_coding | 580.1688091 | 0.395603201 | 4.19E-06 | 7.34E-05 |
| ENSMUST00000170895.2,Guk1-006,protein_coding,ENSMUSG00000020444.19,protein_coding | 830.0248796 | 0.519718014 | 4.21E-06 | 7.38E-05 |
| ENSMUST00000045855.8,Hebp1-001,protein_coding,ENSMUSG00000042770.8,protein_coding | 727.6472667 | 0.326700285 | 4.22E-06 | 7.40E-05 |
| ENSMUST00000210489.1,Srek1-003,protein_coding,ENSMUSG00000032621.8,protein_coding | 738.2690876 | -0.646013437 | 4.23E-06 | 7.41E-05 |
| ENSMUST00000130325.7,Lsm6-002,protein_coding,ENSMUSG00000031683.16,protein_coding | 1412.770627 | -0.306845149 | 4.28E-06 | 7.49E-05 |
| ENSMUST00000113211.7,Rpl36a-001,protein_coding,ENSMUSG00000079435.9,protein_coding | 26025.59549 | 0.180407185 | 4.29E-06 | 7.51E-05 |
| ENSMUST00000043422.7,Tapbpl-001,protein_coding,ENSMUSG00000038213.7,protein_coding | 705.7445094 | 0.298079298 | 4.34E-06 | 7.60E-05 |
| ENSMUST00000022528.4,Pinx1-201,protein_coding,ENSMUSG00000021958.4,protein_coding | 1346.354609 | -0.327182467 | 4.38E-06 | 7.66E-05 |
| ENSMUST00000091257.10,Mfn1-001,protein_coding,ENSMUSG00000027668.13,protein_coding | 3247.327438 | -0.234534658 | 4.42E-06 | 7.73E-05 |
| ENSMUST00000034313.12,Ntpcr-001,protein_coding,ENSMUSG00000031851.14,protein_coding | 1541.0476 | 0.269778744 | 4.43E-06 | 7.75E-05 |
| ENSMUST00000103133.3,Smarce1-001,protein_coding,ENSMUSG00000037935.16,protein_coding | 17463.19177 | 0.161170678 | 4.44E-06 | 7.76E-05 |
| ENSMUST00000108485.8,Atp2a3-202,protein_coding,ENSMUSG00000020788.15,protein_coding | 326.5472076 | 0.773420432 | 4.47E-06 | 7.80E-05 |
| ENSMUST00000108799.9,Tpd52l2-009,protein_coding,ENSMUSG00000000827.18,protein_coding | 2345.748279 | 0.279323277 | 4.55E-06 | 7.93E-05 |
| ENSMUST00000164209.8,Rbm4-006,protein_coding,ENSMUSG00000094936.7,protein_coding | 501.0796191 | 0.373299101 | 4.55E-06 | 7.93E-05 |
| ENSMUST00000031038.10,Ociad1-004,protein_coding,ENSMUSG00000029152.13,protein_coding | 13244.99686 | 0.205886119 | 4.56E-06 | 7.94E-05 |
| ENSMUST00000110952.9,Rcbtb2-201,protein_coding,ENSMUSG00000022106.14,protein_coding | 3434.416465 | 0.214928527 | 4.56E-06 | 7.94E-05 |
| ENSMUST00000161074.7,Hint3-001,protein_coding,ENSMUSG00000019791.12,protein_coding | 517.3712649 | 0.416203519 | 4.64E-06 | 8.07E-05 |
| ENSMUST00000041892.12,Wdr19-005,protein_coding,ENSMUSG00000037890.13,protein_coding | 1094.69736 | 0.410682226 | 4.65E-06 | 8.10E-05 |
| ENSMUST00000107682.1,Tmem154-001,protein_coding,ENSMUSG00000056498.13,protein_coding | 391.7527915 | 0.458394243 | 4.67E-06 | 8.13E-05 |
| ENSMUST00000131713.7,Srp19-005,processed_transcript,ENSMUSG00000014504.16,protein_coding | 171.1018049 | -0.748154517 | 4.67E-06 | 8.13E-05 |
| ENSMUST00000109943.9,Mtmr3-201,protein_coding,ENSMUSG00000034354.17,protein_coding | 6719.456424 | 0.234988646 | 4.68E-06 | 8.14E-05 |
| ENSMUST00000191237.6,Tmem183a-003,nonsense_mediated_decay,ENSMUSG00000042305.12,protein_coding | 2142.058623 | 0.333422768 | 4.70E-06 | 8.16E-05 |
| ENSMUST00000194677.5,Lamb3-005,protein_coding,ENSMUSG00000026639.18,protein_coding | 55.93782218 | 0.853815019 | 4.71E-06 | 8.18E-05 |
| ENSMUST00000191462.1,Smim10l1-001,protein_coding,ENSMUSG00000072704.6,protein_coding | 151.52335 | 0.690156519 | 4.73E-06 | 8.21E-05 |
| ENSMUST00000040069.8,Colec12-201,protein_coding,ENSMUSG00000036103.8,protein_coding | 1208.655367 | -0.350657719 | 4.75E-06 | 8.24E-05 |
| ENSMUST00000020856.4,Bzw2-201,protein_coding,ENSMUSG00000020547.4,protein_coding | 15362.62613 | 0.20519961 | 4.75E-06 | 8.25E-05 |
| ENSMUST00000162530.1,Luc7l2-013,retained_intron,ENSMUSG00000029823.16,protein_coding | 276.0018243 | -0.723471149 | 4.78E-06 | 8.29E-05 |
| ENSMUST00000053020.7,Neurl1b-001,protein_coding,ENSMUSG00000034413.14,protein_coding | 1145.801242 | 0.343915917 | 4.80E-06 | 8.32E-05 |
| ENSMUST00000168064.1,Vgll3-201,protein_coding,ENSMUSG00000091243.1,protein_coding | 2313.920655 | -0.420222428 | 4.80E-06 | 8.32E-05 |

| | | | | |
|---|---|---|---|---|
| ENSMUST00000029013.9,Rae1-001,protein_coding,ENSMUSG00000027509.11,protein_coding | 11369.38322 | 0.212119957 | 4.81E-06 | 8.34E-05 |
| ENSMUST00000002152.12,Bbc3-001,protein_coding,ENSMUSG00000002083.13,protein_coding | 522.6050408 | 0.533542427 | 4.82E-06 | 8.34E-05 |
| ENSMUST00000174512.2,Zfp101-002,protein_coding,ENSMUSG00000055240.16,protein_coding | 373.2847176 | -0.743021658 | 4.83E-06 | 8.36E-05 |
| ENSMUST00000040824.1,Dpm3-201,protein_coding,ENSMUSG00000042737.9,protein_coding | 1880.16705 | 0.323359063 | 4.85E-06 | 8.40E-05 |
| ENSMUST00000030841.9,Klhl7-002,protein_coding,ENSMUSG00000028986.12,protein_coding | 1767.00314 | -0.425523881 | 4.86E-06 | 8.40E-05 |
| ENSMUST00000003245.9,Sri-006,retained_intron,ENSMUSG00000003161.15,protein_coding | 6673.338707 | 0.248241862 | 4.88E-06 | 8.44E-05 |
| ENSMUST00000127300.1,Calm1-004,retained_intron,ENSMUSG00000001175.13,protein_coding | 2751.955419 | 0.276996716 | 4.89E-06 | 8.45E-05 |
| ENSMUST00000179114.7,Rabep1-006,protein_coding,ENSMUSG00000020817.16,protein_coding | 440.3516086 | 0.462494937 | 4.92E-06 | 8.50E-05 |
| ENSMUST00000099353.4,Sfr1-201,protein_coding,ENSMUSG00000025066.9,protein_coding | 25561.72013 | 0.178507756 | 4.98E-06 | 8.61E-05 |
| ENSMUST00000030145.8,Dcaf12-001,protein_coding,ENSMUSG00000028436.8,protein_coding | 8147.599661 | 0.167317742 | 5.00E-06 | 8.63E-05 |
| ENSMUST00000188524.1,Eef1b2-009,protein_coding,ENSMUSG00000025967.16,protein_coding | 22831.12327 | 0.259844339 | 5.00E-06 | 8.64E-05 |
| ENSMUST00000129204.7,Dnaja1-002,processed_transcript,ENSMUSG00000028410.13,protein_coding | 538.9816852 | -0.534770614 | 5.01E-06 | 8.64E-05 |
| ENSMUST00000004868.5,Mtfp1-001,protein_coding,ENSMUSG00000004748.5,protein_coding | 397.9035319 | -0.354757479 | 5.01E-06 | 8.65E-05 |
| ENSMUST00000173084.7,Ube2i-002,protein_coding,ENSMUSG00000015120.15,protein_coding | 1368.026118 | -0.415236227 | 5.03E-06 | 8.67E-05 |
| ENSMUST00000112871.7,Cxcl12-002,protein_coding,ENSMUSG00000061353.11,protein_coding | 5587.515259 | -0.362478884 | 5.03E-06 | 8.67E-05 |
| ENSMUST00000120938.1,Gm3244-001,processed_pseudogene,ENSMUSG00000083380.1,processed_pseudogene | 3745.015543 | 0.240351277 | 5.04E-06 | 8.68E-05 |
| ENSMUST00000165007.7,Klhdc3-202,protein_coding,ENSMUSG00000063576.11,protein_coding | 726.2480354 | 0.483533535 | 5.06E-06 | 8.70E-05 |
| ENSMUST00000216351.1,Parp6-003,nonsense_mediated_decay,ENSMUSG00000025237.16,protein_coding | 586.73994 | 0.414106781 | 5.09E-06 | 8.76E-05 |
| ENSMUST00000025506.6,Rbm22-001,protein_coding,ENSMUSG00000024604.7,protein_coding | 9347.546144 | 0.196959159 | 5.09E-06 | 8.76E-05 |
| ENSMUST00000109566.8,Dlgap4-005,protein_coding,ENSMUSG00000061689.15,protein_coding | 2554.737823 | 0.267933131 | 5.10E-06 | 8.76E-05 |
| ENSMUST00000106152.2,Eva1b-001,protein_coding,ENSMUSG00000050212.4,protein_coding | 1341.057864 | -0.276512386 | 5.11E-06 | 8.78E-05 |
| ENSMUST00000032272.12,Adipor2-001,protein_coding,ENSMUSG00000030168.13,protein_coding | 12725.75838 | 0.184992658 | 5.12E-06 | 8.80E-05 |
| ENSMUST00000211113.1,Nkd1-002,protein_coding,ENSMUSG00000031661.12,protein_coding | 134.925024 | 0.675910731 | 5.16E-06 | 8.86E-05 |
| ENSMUST00000028880.9,Slc20a1-001,protein_coding,ENSMUSG00000027397.14,protein_coding | 7126.715021 | -0.308914139 | 5.19E-06 | 8.90E-05 |
| ENSMUST00000149495.1,Ebp-003,retained_intron,ENSMUSG00000031168.14,protein_coding | 53.12208496 | 0.849738111 | 5.19E-06 | 8.90E-05 |
| ENSMUST00000139623.1,H3f3b-002,processed_transcript,ENSMUSG00000016559.14,protein_coding | 4864.722526 | 0.395907992 | 5.23E-06 | 8.97E-05 |
| ENSMUST00000028648.2,Syt13-001,protein_coding,ENSMUSG00000027220.2,protein_coding | 467.4748913 | -0.565446628 | 5.23E-06 | 8.97E-05 |
| ENSMUST00000216649.1,Zbtb44-001,protein_coding,ENSMUSG00000047412.16,protein_coding | 868.0429023 | 0.292251162 | 5.23E-06 | 8.97E-05 |
| ENSMUST00000113620.9,Sh3glb2-003,protein_coding,ENSMUSG00000026860.16,protein_coding | 362.1965217 | 0.786555198 | 5.25E-06 | 8.99E-05 |
| ENSMUST00000019660.10,Zkscan1-001,protein_coding,ENSMUSG00000029729.12,protein_coding | 2093.113461 | -0.288069355 | 5.25E-06 | 8.99E-05 |
| ENSMUST00000102699.7,Mutyh-001,protein_coding,ENSMUSG00000028687.17,protein_coding | 746.0376671 | 0.387257819 | 5.26E-06 | 9.01E-05 |
| ENSMUST00000143698.7,Frmd4b-007,processed_transcript,ENSMUSG00000030064.16,protein_coding | 1055.385308 | -0.650745118 | 5.27E-06 | 9.02E-05 |
| ENSMUST00000026698.7,Podxl-001,protein_coding,ENSMUSG00000025608.9,protein_coding | 5736.013225 | 0.132226564 | 5.28E-06 | 9.02E-05 |
| ENSMUST00000007449.8,Akr1b7-001,protein_coding,ENSMUSG00000052131.7,protein_coding | 55.06712762 | 0.818630704 | 5.27E-06 | 9.02E-05 |
| ENSMUST00000188674.6,Fn1-005,protein_coding,ENSMUSG00000026193.15,protein_coding | 3160.400002 | 0.848309113 | 5.28E-06 | 9.02E-05 |
| ENSMUST00000129113.7,Arhgef11-002,protein_coding,ENSMUSG00000041977.18,protein_coding | 3132.719529 | 0.394807534 | 5.28E-06 | 9.03E-05 |
| ENSMUST00000048145.11,Rbfox2-201,protein_coding,ENSMUSG00000033565.16,protein_coding | 10961.079 | 0.302011477 | 5.29E-06 | 9.04E-05 |
| ENSMUST00000030636.10,Stmn1-001,protein_coding,ENSMUSG00000028832.11,protein_coding | 21741.28577 | 0.144022072 | 5.30E-06 | 9.06E-05 |
| ENSMUST00000090614.10,Arhgap1-003,protein_coding,ENSMUSG00000027247.16,protein_coding | 12042.17174 | 0.188381708 | 5.31E-06 | 9.06E-05 |
| ENSMUST00000201185.3,Gsn-005,protein_coding,ENSMUSG00000026879.14,protein_coding | 61128.36709 | 0.249431876 | 5.31E-06 | 9.07E-05 |
| ENSMUST00000026072.3,Trub1-201,protein_coding,ENSMUSG00000025086.12,protein_coding | 1083.12238 | 0.277631408 | 5.38E-06 | 9.18E-05 |
| ENSMUST00000102476.4,B2m-001,protein_coding,ENSMUSG00000060802.8,protein_coding | 15435.07894 | 0.166969376 | 5.38E-06 | 9.18E-05 |
| ENSMUST00000007559.14,Gatad1-001,protein_coding,ENSMUSG00000007415.15,protein_coding | 10530.15622 | 0.207935583 | 5.40E-06 | 9.20E-05 |
| ENSMUST00000135912.7,Fam149a-002,protein_coding,ENSMUSG00000070044.15,protein_coding | 1122.504011 | -0.3264153 | 5.45E-06 | 9.29E-05 |
| ENSMUST00000077066.7,Tmem151a-001,protein_coding,ENSMUSG00000061451.13,protein_coding | 5884.736851 | 0.232528536 | 5.53E-06 | 9.42E-05 |
| ENSMUST00000102838.9,Sptbn1-002,protein_coding,ENSMUSG00000020315.18,protein_coding | 14739.67371 | -0.204343322 | 5.54E-06 | 9.43E-05 |
| ENSMUST00000025161.8,Tapbp-001,protein_coding,ENSMUSG00000024308.14,protein_coding | 6804.89241 | 0.201950005 | 5.55E-06 | 9.44E-05 |
| ENSMUST00000161012.7,Tcf19-001,protein_coding,ENSMUSG00000050410.15,protein_coding | 2630.026057 | -0.315380783 | 5.55E-06 | 9.44E-05 |
| ENSMUST00000069221.11,Syncrip-001,protein_coding,ENSMUSG00000032423.12,protein_coding | 1079.96616 | -0.669847729 | 5.58E-06 | 9.49E-05 |
| ENSMUST00000099556.1,Fam122a-201,protein_coding,ENSMUSG00000074922.1,protein_coding | 1328.131383 | 0.20490809 | 5.63E-06 | 9.56E-05 |
| ENSMUST00000031822.12,Abcg2-001,protein_coding,ENSMUSG00000029802.13,protein_coding | 513.0792911 | -0.339179896 | 5.64E-06 | 9.58E-05 |
| ENSMUST00000026148.8,Cbr2-001,protein_coding,ENSMUSG00000025150.10,protein_coding | 106.0466847 | 0.596581509 | 5.64E-06 | 9.58E-05 |
| ENSMUST00000115655.7,Pqbp1-004,protein_coding,ENSMUSG00000031157.10,protein_coding | 2066.901037 | 0.222484854 | 5.65E-06 | 9.59E-05 |
| ENSMUST00000061681.7,Gm7334-201,protein_coding,ENSMUSG00000044645.7,protein_coding | 670.0292838 | -0.502372583 | 5.66E-06 | 9.60E-05 |
| ENSMUST00000200766.3,Pds5a-005,retained_intron,ENSMUSG00000029202.12,protein_coding | 392.8163165 | -0.54030006 | 5.67E-06 | 9.61E-05 |
| ENSMUST00000049343.14,Pold1-001,protein_coding,ENSMUSG00000038644.14,protein_coding | 16981.24449 | 0.200665844 | 5.67E-06 | 9.61E-05 |
| ENSMUST00000034349.9,Nae1-001,protein_coding,ENSMUSG00000031878.18,protein_coding | 3442.771281 | 0.382970536 | 5.69E-06 | 9.64E-05 |
| ENSMUST00000033763.14,Naa10-001,protein_coding,ENSMUSG00000031388.14,protein_coding | 1673.522729 | -0.282879447 | 5.72E-06 | 9.70E-05 |
| ENSMUST00000032736.10,Mtmr10-001,protein_coding,ENSMUSG00000030522.14,protein_coding | 6439.295302 | -0.345200881 | 5.74E-06 | 9.73E-05 |
| ENSMUST00000124580.1,Irak1-016,protein_coding,ENSMUSG00000031392.18,protein_coding | 78.59769582 | -0.800007547 | 5.75E-06 | 9.73E-05 |
| ENSMUST00000162031.7,Txndc9-001,protein_coding,ENSMUSG00000058407.12,protein_coding | 1387.97815 | -0.316424155 | 5.79E-06 | 9.79E-05 |
| ENSMUST00000121111.8,P3h1-001,protein_coding,ENSMUSG00000028641.16,protein_coding | 4585.607433 | 0.232116586 | 5.79E-06 | 9.80E-05 |
| ENSMUST00000009036.10,Vdac3-001,protein_coding,ENSMUSG00000008892.12,protein_coding | 23660.32356 | 0.200376295 | 5.80E-06 | 9.81E-05 |
| ENSMUST00000036333.13,Prkx-001,protein_coding,ENSMUSG00000035725.13,protein_coding | 2268.274804 | -0.195062 | 5.80E-06 | 9.81E-05 |
| ENSMUST00000015511.14,Plxnd1-001,protein_coding,ENSMUSG00000030123.15,protein_coding | 3130.592851 | 0.216133067 | 5.84E-06 | 9.87E-05 |
| ENSMUST00000151715.1,Tcp1-006,retained_intron,ENSMUSG00000068039.12,protein_coding | 3976.483414 | 0.286754105 | 5.85E-06 | 9.88E-05 |
| ENSMUST00000172298.7,Rab11a-001,protein_coding,ENSMUSG00000004771.12,protein_coding | 1564.47041 | -0.423157951 | 5.88E-06 | 9.92E-05 |
| ENSMUST00000117462.1,Rhod-002,protein_coding,ENSMUSG00000041845.10,protein_coding | 1776.036058 | -0.378086715 | 5.88E-06 | 9.92E-05 |
| ENSMUST00000073575.11,Slc39a13-001,protein_coding,ENSMUSG00000002105.15,protein_coding | 1958.308149 | -0.30103451 | 5.88E-06 | 9.92E-05 |
| ENSMUST00000036852.7,Recql4-201,protein_coding,ENSMUSG00000033762.7,protein_coding | 11187.4001 | 0.420455995 | 5.89E-06 | 9.93E-05 |
| ENSMUST00000167546.1,Usf1-201,protein_coding,ENSMUSG00000026641.13,protein_coding | 2916.656657 | 0.206195893 | 5.91E-06 | 9.96E-05 |
| ENSMUST00000134351.1,Uck1-002,retained_intron,ENSMUSG00000002550.16,protein_coding | 72.33907655 | 0.767003059 | 5.91E-06 | 9.97E-05 |
| ENSMUST00000091924.9,Polr3gl-001,protein_coding,ENSMUSG00000028104.14,protein_coding | 1395.311381 | 0.237149714 | 5.92E-06 | 9.97E-05 |
| ENSMUST00000150459.1,1500009L16Rik-201,protein_coding,ENSMUSG00000087651.2,protein_coding | 69.31900798 | -0.736385444 | 5.95E-06 | 0.000100144 |
| ENSMUST00000045713.3,Nacad-001,protein_coding,ENSMUSG00000041073.10,protein_coding | 291.193186 | -0.551864302 | 5.95E-06 | 0.000100145 |
| ENSMUST00000134965.7,Immp2l-002,protein_coding,ENSMUSG00000056899.10,protein_coding | 211.1238191 | 0.668883601 | 5.96E-06 | 0.000100191 |
| ENSMUST00000178353.1,Gm21992-004,nonsense_mediated_decay,ENSMUSG00000096370.8,protein_coding | 117.62324 | -0.726827033 | 5.96E-06 | 0.000100256 |
| ENSMUST00000165408.7,Baiap2l2-001,protein_coding,ENSMUSG00000018126.15,protein_coding | 273.9612065 | -0.556651033 | 5.97E-06 | 0.000100309 |
| ENSMUST00000137103.7,Serpinf1-006,protein_coding,ENSMUSG00000000753.15,protein_coding | 632.9522238 | 0.400005576 | 5.98E-06 | 0.000100394 |
| ENSMUST00000052281.5,E130012A19Rik-001,protein_coding,ENSMUSG00000043439.5,protein_coding | 2230.43858 | -0.210090329 | 6.01E-06 | 0.000100971 |
| ENSMUST00000179022.7,Lmnb2-201,protein_coding,ENSMUSG00000062075.13,protein_coding | 10030.99483 | 0.172191377 | 6.03E-06 | 0.000101318 |
| ENSMUST00000059250.6,Brms1l-201,protein_coding,ENSMUSG00000012076.7,protein_coding | 2962.495045 | -0.174635222 | 6.04E-06 | 0.000101331 |
| ENSMUST00000004143.2,Stat5b-001,protein_coding,ENSMUSG00000020919.11,protein_coding | 5521.499915 | 0.195467984 | 6.04E-06 | 0.000101389 |
| ENSMUST00000022690.9,Fam160b2-201,protein_coding,ENSMUSG00000022095.9,protein_coding | 1851.53724 | -0.265761067 | 6.06E-06 | 0.000101698 |
| ENSMUST00000045477.4,Stk32a-201,protein_coding,ENSMUSG00000039954.8,protein_coding | 266.6400649 | 0.493748438 | 6.08E-06 | 0.000101982 |
| ENSMUST00000204095.2,Zfp775-001,protein_coding,ENSMUSG00000007216.6,protein_coding | 1163.417448 | 0.30531284 | 6.09E-06 | 0.00010212 |
| ENSMUST00000022416.14,Anxa11-001,protein_coding,ENSMUSG00000021866.15,protein_coding | 10162.49682 | 0.20379017 | 6.10E-06 | 0.000102204 |
| ENSMUST00000037913.8,Rmi2-002,protein_coding,ENSMUSG00000037991.9,protein_coding | 913.3001041 | 0.310749496 | 6.10E-06 | 0.000102204 |
| ENSMUST00000134921.1,C230014O12Rik-001,lincRNA,ENSMUSG00000087461.2,lincRNA | 34.72805576 | -0.834360307 | 6.12E-06 | 0.000102355 |
| ENSMUST00000102976.3,Gata3-001,protein_coding,ENSMUSG00000015619.10,protein_coding | 1567.046219 | 0.381414885 | 6.19E-06 | 0.000103601 |
| ENSMUST00000040260.10,Frrs1-005,protein_coding,ENSMUSG00000033386.10,protein_coding | 2133.932045 | 0.214868085 | 6.20E-06 | 0.000103604 |
| ENSMUST00000092834.11,Synrg-001,protein_coding,ENSMUSG00000034940.15,protein_coding | 1203.863726 | -0.25138679 | 6.20E-06 | 0.000103682 |

| | | | | |
|---|---|---|---|---|
| ENSMUST00000058383.8,Paip2b-001,protein_coding,ENSMUSG00000045896.14,protein_coding | 14806.13436 | 0.176609737 | 6.21E-06 | 0.000103732 |
| ENSMUST00000209238.1,Flt3l-004,nonsense_mediated_decay,ENSMUSG00000110206.1,protein_coding | 23.53416191 | 0.843194316 | 6.24E-06 | 0.000104181 |
| ENSMUST00000171121.8,Ppt2-006,protein_coding,ENSMUSG00000015474.16,protein_coding | 3583.999133 | 0.236658939 | 6.27E-06 | 0.000104574 |
| ENSMUST00000151938.7,Brcc3-003,retained_intron,ENSMUSG00000031201.17,protein_coding | 169.2674754 | -0.752192034 | 6.26E-06 | 0.000104574 |
| ENSMUST00000043050.8,Chst12-001,protein_coding,ENSMUSG00000036599.10,protein_coding | 2540.166062 | -0.269773794 | 6.28E-06 | 0.000104723 |
| ENSMUST00000130645.7,Mtfr1-001,protein_coding,ENSMUSG00000027601.13,protein_coding | 9767.446695 | 0.274681891 | 6.29E-06 | 0.000104929 |
| ENSMUST00000162602.7,Pard3-015,protein_coding,ENSMUSG00000025812.17,protein_coding | 288.8670749 | 0.693216197 | 6.29E-06 | 0.000104929 |
| ENSMUST00000030643.2,Extl1-001,protein_coding,ENSMUSG00000028838.11,protein_coding | 675.1371642 | 0.723975834 | 6.36E-06 | 0.000105984 |
| ENSMUST00000148330.7,Clk1-004,nonsense_mediated_decay,ENSMUSG00000026034.17,protein_coding | 176.2852246 | -0.832260592 | 6.38E-06 | 0.000106229 |
| ENSMUST00000151880.7,Slu7-002,protein_coding,ENSMUSG00000020409.10,protein_coding | 2037.998401 | 0.179800048 | 6.38E-06 | 0.000106229 |
| ENSMUST00000034713.8,Ldlr-001,protein_coding,ENSMUSG00000032193.9,protein_coding | 25744.37676 | 0.225151305 | 6.40E-06 | 0.000106521 |
| ENSMUST00000042288.6,Asb13-001,protein_coding,ENSMUSG00000033781.6,protein_coding | 548.7331173 | -0.414570327 | 6.40E-06 | 0.000106569 |
| ENSMUST00000203730.2,Ppfibp1-012,protein_coding,ENSMUSG00000016487.15,protein_coding | 1068.282879 | 0.269888741 | 6.49E-06 | 0.000107956 |
| ENSMUST00000199060.1,Gm17494-002,retained_intron,ENSMUSG00000057359.7,processed_transcript | 280.5944202 | 0.449346587 | 6.50E-06 | 0.000108002 |
| ENSMUST00000066408.5,1700016K19Rik-001,protein_coding,ENSMUSG00000053783.5,protein_coding | 31.52250893 | 0.825547423 | 6.51E-06 | 0.000108215 |
| ENSMUST00000023736.8,Lmbr1l-001,protein_coding,ENSMUSG00000022999.14,protein_coding | 1235.683636 | 0.363964694 | 6.52E-06 | 0.000108346 |
| ENSMUST00000092566.7,Slc16a10-001,protein_coding,ENSMUSG00000019838.11,protein_coding | 1578.675088 | 0.41518411 | 6.55E-06 | 0.000108754 |
| ENSMUST00000026976.11,Get4-001,protein_coding,ENSMUSG00000025858.12,protein_coding | 5774.277729 | -0.199056673 | 6.56E-06 | 0.000108878 |
| ENSMUST00000134102.1,Ube2o-002,processed_transcript,ENSMUSG00000020802.8,protein_coding | 322.1812026 | 0.632969594 | 6.59E-06 | 0.000109339 |
| ENSMUST00000130870.1,Gm14226-001,protein_coding,ENSMUSG00000084897.1,protein_coding | 373.6850643 | -0.493778335 | 6.62E-06 | 0.000109861 |
| ENSMUST00000058022.5,Tpcn2-001,protein_coding,ENSMUSG00000048677.9,protein_coding | 701.3128159 | -0.459336194 | 6.63E-06 | 0.000109988 |
| ENSMUST00000105545.10,Phactr2-201,protein_coding,ENSMUSG00000062866.15,protein_coding | 661.4111247 | -0.459468527 | 6.64E-06 | 0.000110034 |
| ENSMUST00000161994.1,Ciart-005,protein_coding,ENSMUSG00000038550.10,protein_coding | 239.6514004 | 0.666988244 | 6.65E-06 | 0.000110224 |
| ENSMUST00000084030.5,Tacc1-002,protein_coding,ENSMUSG00000065954.11,protein_coding | 4803.437039 | -0.274965933 | 6.66E-06 | 0.000110279 |
| ENSMUST00000092735.11,Med1-004,protein_coding,ENSMUSG00000018160.15,protein_coding | 770.9952275 | 0.551469606 | 6.70E-06 | 0.000110899 |
| ENSMUST00000150416.7,Wsb1-005,retained_intron,ENSMUSG00000017677.11,protein_coding | 1174.713931 | -0.455474963 | 6.72E-06 | 0.000111147 |
| ENSMUST00000144000.7,Srsf9-002,processed_transcript,ENSMUSG00000029538.14,protein_coding | 49.75527473 | 0.833904453 | 6.72E-06 | 0.000111222 |
| ENSMUST00000113952.9,Wdr5-001,protein_coding,ENSMUSG00000026917.15,protein_coding | 15519.74222 | 0.194287647 | 6.77E-06 | 0.000111969 |
| ENSMUST00000204885.1,Hnrnpa2b1-008,retained_intron,ENSMUSG00000004980.16,protein_coding | 3135.302136 | -0.449704806 | 6.78E-06 | 0.0001121 |
| ENSMUST00000190080.6,Trerf1-001,processed_transcript,ENSMUSG00000064043.13,processed_transcript | 1332.956782 | -0.326638635 | 6.82E-06 | 0.000112746 |
| ENSMUST00000122865.1,Sik3-002,protein_coding,ENSMUSG00000034135.15,protein_coding | 2726.320286 | -0.217906671 | 6.84E-06 | 0.000112984 |
| ENSMUST00000205651.1,Fbrs-002,retained_intron,ENSMUSG00000042423.9,protein_coding | 217.946683 | -0.493081084 | 6.90E-06 | 0.00011389 |
| ENSMUST00000026427.6,Esyt1-201,protein_coding,ENSMUSG00000025366.7,protein_coding | 15705.84886 | 0.22418032 | 6.91E-06 | 0.000113973 |
| ENSMUST00000100635.4,Fam84b-001,protein_coding,ENSMUSG00000072568.4,protein_coding | 1194.713546 | 0.553991071 | 6.91E-06 | 0.000113973 |
| ENSMUST00000015893.12,Anp32e-002,protein_coding,ENSMUSG00000015749.12,protein_coding | 3687.458602 | 0.295128697 | 6.92E-06 | 0.000114193 |
| ENSMUST00000080797.7,Cdh3-001,protein_coding,ENSMUSG00000061048.8,protein_coding | 5635.391779 | -0.178276802 | 6.93E-06 | 0.00011424 |
| ENSMUST00000212616.1,Csnk2a2-010,protein_coding,ENSMUSG00000046707.9,protein_coding | 2618.241934 | 0.226475277 | 6.94E-06 | 0.000114424 |
| ENSMUST00000102926.4,Anp32b-001,protein_coding,ENSMUSG00000028333.10,protein_coding | 69260.05887 | 0.19254469 | 6.97E-06 | 0.000114847 |
| ENSMUST00000135960.7,Wars2-002,nonsense_mediated_decay,ENSMUSG00000004233.14,protein_coding | 1088.994786 | 0.35339548 | 7.00E-06 | 0.000115243 |
| ENSMUST00000199397.1,Miga1-004,protein_coding,ENSMUSG00000054942.13,protein_coding | 243.8629842 | -0.471409273 | 7.01E-06 | 0.00011541 |
| ENSMUST00000193053.5,Wdr20-001,protein_coding,ENSMUSG00000037957.14,protein_coding | 298.1900255 | 0.493505936 | 7.01E-06 | 0.00011541 |
| ENSMUST00000037012.2,Kcna4-001,protein_coding,ENSMUSG00000042604.5,protein_coding | 69.15745136 | -0.729179388 | 7.05E-06 | 0.000115947 |
| ENSMUST00000018002.12,Ift52-001,protein_coding,ENSMUSG00000017858.12,protein_coding | 5073.023261 | 0.222393534 | 7.05E-06 | 0.000115947 |
| ENSMUST00000118012.8,Gm14295-001,protein_coding,ENSMUSG00000078877.11,protein_coding | 405.4627908 | -0.538874512 | 7.06E-06 | 0.000115952 |
| ENSMUST00000020964.5,Fkbp1b-201,protein_coding,ENSMUSG00000020635.7,protein_coding | 336.9730277 | 0.380619026 | 7.06E-06 | 0.000115952 |
| ENSMUST00000059589.5,Rtn4r-001,protein_coding,ENSMUSG00000043811.5,protein_coding | 149.3861297 | 0.615395894 | 7.08E-06 | 0.000116315 |
| ENSMUST00000094077.4,Kdm6b-001,protein_coding,ENSMUSG00000018476.7,protein_coding | 10112.93231 | 0.288871689 | 7.10E-06 | 0.000116601 |
| ENSMUST00000131867.7,Dkc1-004,processed_transcript,ENSMUSG00000031403.14,protein_coding | 304.3846071 | 0.403999034 | 7.13E-06 | 0.00011707 |
| ENSMUST00000029908.7,Faxc-001,protein_coding,ENSMUSG00000028246.13,protein_coding | 372.1620139 | -0.518755452 | 7.16E-06 | 0.000117493 |
| ENSMUST00000212634.1,Pard6a-004,processed_transcript,ENSMUSG00000005699.16,protein_coding | 255.5780468 | 0.560999288 | 7.21E-06 | 0.000118219 |
| ENSMUST00000034457.8,Urb2-001,protein_coding,ENSMUSG00000031976.14,protein_coding | 3763.187789 | -0.26656067 | 7.24E-06 | 0.000118599 |
| ENSMUST00000039630.5,Ror1-001,protein_coding,ENSMUSG00000035305.5,protein_coding | 331.4362038 | -0.572331696 | 7.26E-06 | 0.000118938 |
| ENSMUST00000168401.1,Lpl-002,nonsense_mediated_decay,ENSMUSG00000015568.16,protein_coding | 1341.569797 | 0.389725325 | 7.26E-06 | 0.000118954 |
| ENSMUST00000185398.6,Mdm4-004,nonsense_mediated_decay,ENSMUSG00000054387.13,protein_coding | 95.60129443 | 0.829385935 | 7.27E-06 | 0.000118954 |
| ENSMUST00000105579.7,Sdf4-002,protein_coding,ENSMUSG00000029076.14,protein_coding | 16137.0779 | 0.241570337 | 7.35E-06 | 0.000120253 |
| ENSMUST00000109896.7,Ninl-001,protein_coding,ENSMUSG00000068115.13,protein_coding | 3891.802201 | -0.186706371 | 7.37E-06 | 0.00012055 |
| ENSMUST00000114849.1,Uhrf1bp1-001,protein_coding,ENSMUSG00000039512.11,protein_coding | 12905.48448 | 0.246582679 | 7.39E-06 | 0.000120879 |
| ENSMUST00000033920.5,Aga-001,protein_coding,ENSMUSG00000031521.5,protein_coding | 2139.552762 | 0.267270627 | 7.42E-06 | 0.000121298 |
| ENSMUST00000165684.7,Med16-003,protein_coding,ENSMUSG00000013833.15,protein_coding | 2521.211175 | -0.307256237 | 7.44E-06 | 0.000121515 |
| ENSMUST00000093078.12,Banp-002,protein_coding,ENSMUSG00000025316.16,protein_coding | 2474.32884 | 0.244702621 | 7.51E-06 | 0.000122665 |
| ENSMUST00000138979.1,Vma21-004,processed_transcript,ENSMUSG00000073131.12,protein_coding | 190.1476463 | 0.786822004 | 7.59E-06 | 0.000123958 |
| ENSMUST00000125435.1,Gpc6-002,protein_coding,ENSMUSG00000058571.10,protein_coding | 382.0207596 | 0.511662222 | 7.65E-06 | 0.000124825 |
| ENSMUST00000050990.9,Asap2-006,protein_coding,ENSMUSG00000052632.15,protein_coding | 1757.674466 | 0.263759819 | 7.65E-06 | 0.000124849 |
| ENSMUST00000193901.5,Pcdh9-004,protein_coding,ENSMUSG00000055421.8,protein_coding | 1151.482449 | -0.460026331 | 7.70E-06 | 0.000125517 |
| ENSMUST00000039476.14,Arhgef11-001,protein_coding,ENSMUSG00000041977.18,protein_coding | 7622.155811 | 0.400795191 | 7.71E-06 | 0.000125734 |
| ENSMUST00000079644.12,Ybx1-001,protein_coding,ENSMUSG00000028639.14,protein_coding | 184056.0006 | 0.232110752 | 7.72E-06 | 0.000125734 |
| ENSMUST00000021158.3,Txndc17-001,protein_coding,ENSMUSG00000020803.3,protein_coding | 5737.674785 | -0.157703556 | 7.72E-06 | 0.000125734 |
| ENSMUST00000176670.7,Cpsf6-005,protein_coding,ENSMUSG00000055531.12,protein_coding | 4706.002324 | 0.156922613 | 7.74E-06 | 0.00012608 |
| ENSMUST00000058884.8,Parn-201,protein_coding,ENSMUSG00000022685.8,protein_coding | 8028.700907 | 0.270055183 | 7.81E-06 | 0.000127207 |
| ENSMUST00000039956.5,Slc41a2-001,protein_coding,ENSMUSG00000034591.5,protein_coding | 973.2766324 | -0.295800993 | 7.82E-06 | 0.000127261 |
| ENSMUST00000027687.7,Ube2t-001,protein_coding,ENSMUSG00000026429.8,protein_coding | 2302.192513 | 0.235463496 | 7.83E-06 | 0.000127283 |
| ENSMUST00000135242.7,Fyn-004,protein_coding,ENSMUSG00000019843.14,protein_coding | 546.8899891 | 0.665083827 | 7.82E-06 | 0.000127283 |
| ENSMUST00000119251.7,Mef2d-002,protein_coding,ENSMUSG00000001419.17,protein_coding | 4358.497325 | 0.202250892 | 7.83E-06 | 0.000127289 |
| ENSMUST00000166711.7,Adsl-002,nonsense_mediated_decay,ENSMUSG00000022407.9,protein_coding | 8665.689382 | 0.185730921 | 7.85E-06 | 0.00012755 |
| ENSMUST00000172132.8,Tpi1-001,protein_coding,ENSMUSG00000023456.14,protein_coding | 210909.2083 | 0.207620178 | 7.88E-06 | 0.000128017 |
| ENSMUST00000118009.1,Naf1-001,protein_coding,ENSMUSG00000014907.10,protein_coding | 14086.40855 | 0.183683111 | 7.88E-06 | 0.000128017 |
| ENSMUST00000164296.7,Brd3-202,protein_coding,ENSMUSG00000026918.16,protein_coding | 2411.671104 | 0.452947165 | 7.89E-06 | 0.000128032 |
| ENSMUST00000170134.7,Rangap1-202,protein_coding,ENSMUSG00000022391.15,protein_coding | 1586.568845 | 0.333228849 | 7.90E-06 | 0.000128111 |
| ENSMUST00000135846.1,Enpp1-002,protein_coding,ENSMUSG00000037370.13,protein_coding | 490.1714117 | 0.469615002 | 7.95E-06 | 0.000128957 |
| ENSMUST00000040314.11,Rbm17-001,protein_coding,ENSMUSG00000037197.11,protein_coding | 17801.82293 | 0.132192568 | 7.96E-06 | 0.000129053 |
| ENSMUST00000024857.12,Lbh-001,protein_coding,ENSMUSG00000024063.12,protein_coding | 3610.743788 | -0.26287449 | 8.05E-06 | 0.000130509 |
| ENSMUST00000199320.4,Cenpa-009,protein_coding,ENSMUSG00000029177.9,protein_coding | 1142.087032 | -0.349396785 | 8.07E-06 | 0.000130704 |
| ENSMUST00000170578.1,Rgmb-201,protein_coding,ENSMUSG00000048027.7,protein_coding | 2392.888503 | 0.181783491 | 8.08E-06 | 0.000130915 |
| ENSMUST00000048308.5,C130074G19Rik-001,protein_coding,ENSMUSG00000039349.5,protein_coding | 240.3535452 | 0.607502563 | 8.10E-06 | 0.000131117 |
| ENSMUST00000032143.7,Rpn1-001,protein_coding,ENSMUSG00000030062.7,protein_coding | 47307.7897 | 0.147225595 | 8.10E-06 | 0.000131117 |
| ENSMUST00000147574.7,Add1-007,protein_coding,ENSMUSG00000029106.14,protein_coding | 227.9890758 | 0.568148322 | 8.20E-06 | 0.000132684 |
| ENSMUST00000086876.5,Pot1b-201,protein_coding,ENSMUSG00000024174.8,protein_coding | 825.1109766 | -0.295630456 | 8.22E-06 | 0.000132884 |
| ENSMUST00000004774.3,Aqp1-001,protein_coding,ENSMUSG00000004655.5,protein_coding | 2027.715805 | 0.289730938 | 8.28E-06 | 0.000133844 |
| ENSMUST00000151966.7,Prpsap2-002,processed_transcript,ENSMUSG00000020528.14,protein_coding | 1317.455873 | 0.444142214 | 8.29E-06 | 0.00013389 |
| ENSMUST00000065906.8,Ggcx-001,protein_coding,ENSMUSG00000053460.8,protein_coding | 3290.271675 | -0.206679881 | 8.32E-06 | 0.000134365 |
| ENSMUST00000110586.9,Casc4-201,protein_coding,ENSMUSG00000060227.15,protein_coding | 1180.048731 | -0.601504367 | 8.34E-06 | 0.000134585 |
| ENSMUST00000067268.14,Pxn-001,protein_coding,ENSMUSG00000029528.18,protein_coding | 15520.08716 | 0.171011906 | 8.39E-06 | 0.000135428 |

| | | | | |
|---|---|---|---|---|
| ENSMUST00000022196.3,Ap3b1-201,protein_coding,ENSMUSG00000021686.4,protein_coding | 13593.47255 | 0.188080893 | 8.39E-06 | 0.000135428 |
| ENSMUST00000028816.8,Tmem230-001,protein_coding,ENSMUSG00000027341.10,protein_coding | 2565.780145 | 0.25694228 | 8.40E-06 | 0.000135506 |
| ENSMUST00000028696.4,Ddb2-001,protein_coding,ENSMUSG00000002109.14,protein_coding | 772.0016675 | 0.497251277 | 8.42E-06 | 0.00013576 |
| ENSMUST00000043676.11,Sspo-201,protein_coding,ENSMUSG00000029797.13,protein_coding | 42.66475747 | 0.822882617 | 8.43E-06 | 0.000135845 |
| ENSMUST00000120243.7,Csf1-002,protein_coding,ENSMUSG00000014599.10,protein_coding | 1028.457938 | -0.562105517 | 8.44E-06 | 0.000135992 |
| ENSMUST00000153641.7,Surf4-003,nonsense_mediated_decay,ENSMUSG00000014867.9,protein_coding | 1085.839534 | 0.314270495 | 8.46E-06 | 0.000136304 |
| ENSMUST00000201530.1,Dusp11-008,nonsense_mediated_decay,ENSMUSG00000030002.15,protein_coding | 1010.803964 | 0.373336499 | 8.53E-06 | 0.000137328 |
| ENSMUST00000030254.14,Leprot-001,protein_coding,ENSMUSG00000035212.14,protein_coding | 4169.020462 | 0.179499941 | 8.56E-06 | 0.00013779 |
| ENSMUST00000186583.6,Ash1l-001,protein_coding,ENSMUSG00000028053.13,protein_coding | 3838.170817 | -0.432174988 | 8.57E-06 | 0.000137806 |
| ENSMUST00000208749.1,Serpinh1-005,protein_coding,ENSMUSG00000070436.12,protein_coding | 533.9016241 | -0.374405428 | 8.59E-06 | 0.000138174 |
| ENSMUST00000020273.15,Supv3l1-001,protein_coding,ENSMUSG00000020079.15,protein_coding | 7305.30874 | 0.239703202 | 8.62E-06 | 0.000138497 |
| ENSMUST00000045243.14,Pde4dip-005,protein_coding,ENSMUSG00000038170.15,protein_coding | 11584.7824 | 0.272920014 | 8.65E-06 | 0.000138896 |
| ENSMUST00000119902.5,Tspan2-002,protein_coding,ENSMUSG00000027858.13,protein_coding | 30.55311873 | -0.830176573 | 8.65E-06 | 0.000138896 |
| ENSMUST00000048002.6,B4galnt4-001,protein_coding,ENSMUSG00000055629.5,protein_coding | 393.4977058 | -0.386957562 | 8.65E-06 | 0.000138896 |
| ENSMUST00000202370.1,Gm9794-001,processed_pseudogene,ENSMUSG00000107176.1,processed_pseudogene | 821.4285357 | -0.478788088 | 8.67E-06 | 0.000139121 |
| ENSMUST00000137029.1,Fbln2-003,retained_intron,ENSMUSG00000064080.12,protein_coding | 12736.5298 | 0.22778707 | 8.69E-06 | 0.000139363 |
| ENSMUST00000109267.8,Map3k1-001,protein_coding,ENSMUSG00000021754.17,protein_coding | 2389.134405 | -0.25572171 | 8.76E-06 | 0.00014054 |
| ENSMUST00000195855.5,Pex2-003,protein_coding,ENSMUSG00000040374.13,protein_coding | 176.2559417 | -0.808263821 | 8.78E-06 | 0.000140783 |
| ENSMUST00000026436.9,Baiap2-001,protein_coding,ENSMUSG00000025372.16,protein_coding | 4449.788094 | 0.216072083 | 8.79E-06 | 0.000140877 |
| ENSMUST00000161032.7,Ipo9-003,nonsense_mediated_decay,ENSMUSG00000041879.13,protein_coding | 4259.950835 | 0.446475279 | 8.82E-06 | 0.000141353 |
| ENSMUST00000025542.9,Psat1-001,protein_coding,ENSMUSG00000024640.9,protein_coding | 84524.35354 | 0.240310589 | 8.83E-06 | 0.000141373 |
| ENSMUST00000022224.15,Trappc13-003,protein_coding,ENSMUSG00000021711.16,protein_coding | 1095.924782 | -0.399882962 | 8.87E-06 | 0.000141915 |
| ENSMUST00000168100.1,BC100451-201,protein_coding,ENSMUSG00000076433.4,protein_coding | 213.2841429 | -0.648732951 | 8.91E-06 | 0.000142599 |
| ENSMUST00000115870.8,Epc1-001,protein_coding,ENSMUSG00000024240.13,protein_coding | 567.6050132 | 0.504763491 | 8.91E-06 | 0.000142599 |
| ENSMUST00000019927.6,Trmt11-001,protein_coding,ENSMUSG00000019792.8,protein_coding | 793.4229955 | -0.582142377 | 8.93E-06 | 0.000142792 |
| ENSMUST00000053252.8,Ctxn1-001,protein_coding,ENSMUSG00000048644.8,protein_coding | 2022.704648 | -0.231178229 | 8.95E-06 | 0.000143129 |
| ENSMUST00000210792.1,Rpl13a-008,retained_intron,ENSMUSG00000074129.13,protein_coding | 4182.584736 | 0.335426494 | 8.98E-06 | 0.000143449 |
| ENSMUST00000156902.1,Rai1-004,processed_transcript,ENSMUSG00000062115.15,protein_coding | 193.3791737 | -0.5032215 | 9.06E-06 | 0.000144685 |
| ENSMUST00000067538.5,Ptprs-001,protein_coding,ENSMUSG00000013236.16,protein_coding | 29681.61074 | 0.195626578 | 9.11E-06 | 0.000145407 |
| ENSMUST00000031221.11,Cdc7-001,protein_coding,ENSMUSG00000029283.17,protein_coding | 1510.752576 | -0.307850043 | 9.15E-06 | 0.0001461 |
| ENSMUST00000143496.7,Chpf-004,retained_intron,ENSMUSG00000032997.16,protein_coding | 220.3105799 | -0.564754767 | 9.19E-06 | 0.00014669 |
| ENSMUST00000147570.2,Snrpb-002,nonsense_mediated_decay,ENSMUSG00000027404.15,protein_coding | 256.8239363 | 0.721438241 | 9.20E-06 | 0.000146749 |
| ENSMUST00000019199.13,Plod1-001,protein_coding,ENSMUSG00000019055.15,protein_coding | 5730.71782 | 0.148706104 | 9.21E-06 | 0.000146774 |
| ENSMUST00000068487.11,Rad18-002,protein_coding,ENSMUSG00000030254.16,protein_coding | 1244.34495 | -0.388413045 | 9.22E-06 | 0.000146892 |
| ENSMUST00000174651.1,Gm20604-001,protein_coding,ENSMUSG00000041716.7,protein_coding | 901.9331248 | -0.353957846 | 9.24E-06 | 0.000147186 |
| ENSMUST00000151513.1,Eif1-002,protein_coding,ENSMUSG00000035530.13,protein_coding | 192.6862868 | 0.584577097 | 9.26E-06 | 0.000147504 |
| ENSMUST00000107100.2,Map6-202,protein_coding,ENSMUSG00000055407.14,protein_coding | 2315.604408 | 0.22269882 | 9.30E-06 | 0.000147997 |
| ENSMUST00000044581.13,1700025G04Rik-001,protein_coding,ENSMUSG00000032666.16,protein_coding | 3874.494025 | -0.205502532 | 9.31E-06 | 0.000148126 |
| ENSMUST00000212301.1,Hip1-010,protein_coding,ENSMUSG00000039959.13,protein_coding | 103.3132671 | 0.809757136 | 9.32E-06 | 0.000148163 |
| ENSMUST00000120943.7,Spsb3-001,protein_coding,ENSMUSG00000024160.16,protein_coding | 669.0041283 | -0.795464307 | 9.32E-06 | 0.000148163 |
| ENSMUST00000030581.9,Azin2-001,protein_coding,ENSMUSG00000028789.16,protein_coding | 282.405102 | 0.480761383 | 9.33E-06 | 0.000148277 |
| ENSMUST00000064447.11,Nup188-001,protein_coding,ENSMUSG00000052533.14,protein_coding | 49464.42355 | 0.196376295 | 9.38E-06 | 0.000149108 |
| ENSMUST00000025279.5,Npc1-001,protein_coding,ENSMUSG00000024413.13,protein_coding | 5751.480069 | 0.320747455 | 9.41E-06 | 0.000149542 |
| ENSMUST00000026448.9,Faap100-001,protein_coding,ENSMUSG00000025384.15,protein_coding | 2041.151506 | -0.215861884 | 9.43E-06 | 0.000149688 |
| ENSMUST00000001711.5,Hoxc6-001,protein_coding,ENSMUSG00000001661.5,protein_coding | 1740.819295 | -0.213908814 | 9.43E-06 | 0.000149694 |
| ENSMUST00000129021.8,Aptx-002,nonsense_mediated_decay,ENSMUSG00000028411.15,protein_coding | 1533.76795 | 0.22773731 | 9.47E-06 | 0.000150262 |
| ENSMUST00000159759.1,AI837181-001,protein_coding,ENSMUSG00000047423.9,protein_coding | 4581.829337 | -0.277053332 | 9.55E-06 | 0.000151428 |
| ENSMUST00000193675.1,Vim-006,protein_coding,ENSMUSG00000026728.9,protein_coding | 80158.55184 | 0.2485158 | 9.55E-06 | 0.000151441 |
| ENSMUST00000064200.7,Tmem106c-201,protein_coding,ENSMUSG00000052369.7,protein_coding | 4002.033454 | 0.17307131 | 9.56E-06 | 0.000151508 |
| ENSMUST00000029989.10,Rnf20-001,protein_coding,ENSMUSG00000028309.14,protein_coding | 7943.08063 | 0.271264129 | 9.59E-06 | 0.000151878 |
| ENSMUST00000105886.7,Dhdds-007,protein_coding,ENSMUSG00000012117.13,protein_coding | 206.6407721 | 0.727235625 | 9.60E-06 | 0.000151946 |
| ENSMUST00000177592.1,Ddi2-201,protein_coding,ENSMUSG00000078515.4,protein_coding | 6150.554149 | 0.194782686 | 9.59E-06 | 0.000151946 |
| ENSMUST00000027231.12,Slc9a2-001,protein_coding,ENSMUSG00000026062.12,protein_coding | 9515.884788 | 0.176960584 | 9.63E-06 | 0.000152422 |
| ENSMUST00000027846.7,Tada1-001,protein_coding,ENSMUSG00000026563.13,protein_coding | 1561.958473 | 0.242099336 | 9.66E-06 | 0.00015281 |
| ENSMUST00000182597.7,Birc6-004,protein_coding,ENSMUSG00000024073.14,protein_coding | 7163.946517 | -0.59171822 | 9.66E-06 | 0.00015281 |
| ENSMUST00000010298.6,Spire2-001,protein_coding,ENSMUSG00000010154.7,protein_coding | 2796.848689 | 0.221354922 | 9.67E-06 | 0.000152943 |
| ENSMUST00000088373.10,Srgap3-001,protein_coding,ENSMUSG00000030257.16,protein_coding | 1953.762735 | 0.226419649 | 9.68E-06 | 0.000153038 |
| ENSMUST00000142718.7,Myef2-005,protein_coding,ENSMUSG00000027201.16,protein_coding | 2955.246868 | -0.657496705 | 9.71E-06 | 0.000153389 |
| ENSMUST00000031344.12,Mphosph9-001,protein_coding,ENSMUSG00000038126.17,protein_coding | 2820.899131 | -0.178744123 | 9.80E-06 | 0.000154759 |
| ENSMUST00000107247.7,Nfib-003,protein_coding,ENSMUSG00000008575.17,protein_coding | 477.557752 | -0.825309187 | 9.83E-06 | 0.000155122 |
| ENSMUST00000003115.8,Cdkn1b-002,protein_coding,ENSMUSG00000003031.14,protein_coding | 4561.509356 | 0.349539314 | 9.85E-06 | 0.000155461 |
| ENSMUST00000036865.12,Fanci-001,protein_coding,ENSMUSG00000039187.16,protein_coding | 4394.756158 | -0.162055757 | 9.94E-06 | 0.000156753 |
| ENSMUST00000053202.11,Rhbdl2-201,protein_coding,ENSMUSG00000043333.12,protein_coding | 65.10929926 | -0.741577353 | 9.97E-06 | 0.000157211 |
| ENSMUST00000031287.10,Tpst2-001,protein_coding,ENSMUSG00000029344.14,protein_coding | 9837.619434 | -0.189430652 | 1.00E-05 | 0.000157903 |
| ENSMUST00000116641.8,Lmf1-001,protein_coding,ENSMUSG00000002279.18,protein_coding | 1756.609795 | 0.317910631 | 1.00E-05 | 0.00015822 |
| ENSMUST00000202196.3,Sept11-007,protein_coding,ENSMUSG00000058013.11,protein_coding | 321.2407776 | 0.710976658 | 1.00E-05 | 0.000158228 |
| ENSMUST00000035484.10,Cdv3-001,protein_coding,ENSMUSG00000032803.15,protein_coding | 12547.12341 | -0.224355126 | 1.01E-05 | 0.000158391 |
| ENSMUST00000115520.7,Ica1-004,protein_coding,ENSMUSG00000062995.12,protein_coding | 1347.006758 | 0.356350076 | 1.01E-05 | 0.000158883 |
| ENSMUST00000001709.2,Hoxc5-001,protein_coding,ENSMUSG00000022485.3,protein_coding | 477.7318536 | -0.366204425 | 1.01E-05 | 0.000159244 |
| ENSMUST00000094782.9,Inpp5b-001,protein_coding,ENSMUSG00000028894.18,protein_coding | 9281.960935 | 0.22341496 | 1.02E-05 | 0.000159531 |
| ENSMUST00000092648.12,Ezh2-010,protein_coding,ENSMUSG00000029687.16,protein_coding | 5079.109826 | 0.278135804 | 1.01E-05 | 0.000159531 |
| ENSMUST00000038572.14,Hnrnpl-201,protein_coding,ENSMUSG00000015165.16,protein_coding | 22250.78707 | 0.270142003 | 1.02E-05 | 0.000159531 |
| ENSMUST00000035458.14,Ate1-002,protein_coding,ENSMUSG00000030850.17,protein_coding | 1219.832589 | -0.491088562 | 1.02E-05 | 0.00015978 |
| ENSMUST00000029175.13,Src-001,protein_coding,ENSMUSG00000027646.15,protein_coding | 12927.20176 | 0.200756117 | 1.02E-05 | 0.000159824 |
| ENSMUST00000074259.14,Nrm-001,protein_coding,ENSMUSG00000059791.14,protein_coding | 2803.121933 | 0.246213058 | 1.02E-05 | 0.000160002 |
| ENSMUST00000021903.2,Gadd45g-001,protein_coding,ENSMUSG00000021453.2,protein_coding | 2370.874519 | -0.413197214 | 1.02E-05 | 0.000160055 |
| ENSMUST00000108196.7,Capns1-002,protein_coding,ENSMUSG00000001794.13,protein_coding | 8721.816896 | 0.399653939 | 1.02E-05 | 0.00016031 |
| ENSMUST00000003152.13,Stk11-001,protein_coding,ENSMUSG00000003068.16,protein_coding | 8773.870762 | -0.130956331 | 1.02E-05 | 0.00016031 |
| ENSMUST00000077856.12,Rpl27-001,protein_coding,ENSMUSG00000063316.13,protein_coding | 52954.01169 | 0.202269098 | 1.02E-05 | 0.00016031 |
| ENSMUST00000051301.4,Pura-001,protein_coding,ENSMUSG00000043991.4,protein_coding | 3560.051703 | -0.40038012 | 1.02E-05 | 0.000160374 |
| ENSMUST00000048073.8,Pigs-001,protein_coding,ENSMUSG00000041958.10,protein_coding | 11198.47366 | 0.1658195 | 1.03E-05 | 0.000160841 |
| ENSMUST00000109521.3,Polr2c-001,protein_coding,ENSMUSG00000031783.10,protein_coding | 16276.67516 | 0.200622797 | 1.03E-05 | 0.000161704 |
| ENSMUST00000075606.4,Ehbp1l1-202,protein_coding,ENSMUSG00000024937.14,protein_coding | 4890.004645 | 0.226442916 | 1.03E-05 | 0.000161871 |
| ENSMUST00000120566.1,Gm12183-001,processed_pseudogene,ENSMUSG00000082454.3,processed_pseudogene | 255.2717613 | -0.547184993 | 1.04E-05 | 0.00016221 |
| ENSMUST00000140312.7,Son-005,nonsense_mediated_decay,ENSMUSG00000022961.17,protein_coding | 2944.140102 | -0.707413473 | 1.04E-05 | 0.000162769 |
| ENSMUST00000030118.9,Dnaja1-001,protein_coding,ENSMUSG00000028410.13,protein_coding | 15334.60607 | -0.324044049 | 1.04E-05 | 0.000162936 |
| ENSMUST00000048044.11,Trub2-001,protein_coding,ENSMUSG00000039826.13,protein_coding | 4446.966619 | -0.283058237 | 1.04E-05 | 0.000163089 |
| ENSMUST00000112621.1,Ccdc93-002,protein_coding,ENSMUSG00000026339.18,protein_coding | 1028.146128 | 0.354677015 | 1.05E-05 | 0.000163824 |
| ENSMUST00000110189.8,Smox-005,protein_coding,ENSMUSG00000027333.18,protein_coding | 31.97539071 | -0.820975906 | 1.05E-05 | 0.000163888 |
| ENSMUST00000167866.1,Idh3a-201,protein_coding,ENSMUSG00000032279.11,protein_coding | 1501.100534 | -0.417080303 | 1.05E-05 | 0.000163888 |
| ENSMUST00000029644.15,Ppa2-001,protein_coding,ENSMUSG00000028013.16,protein_coding | 4407.201116 | 0.238881803 | 1.06E-05 | 0.000165142 |
| ENSMUST00000023151.5,Bcl6-001,protein_coding,ENSMUSG00000022508.5,protein_coding | 2218.959981 | 0.20316338 | 1.06E-05 | 0.000165142 |

| | | | | |
|---|---|---|---|---|
| ENSMUST00000170157.7,Slc3a2-001,protein_coding,ENSMUSG00000010095.13,protein_coding | 2591.197696 | 0.374600192 | 1.06E-05 | 0.000165241 |
| ENSMUST00000031692.11,Bcl7b-001,protein_coding,ENSMUSG00000029681.12,protein_coding | 3233.817882 | 0.236943142 | 1.06E-05 | 0.000165512 |
| ENSMUST00000212441.1,Csnk2a2-006,nonsense_mediated_decay,ENSMUSG00000046707.9,protein_coding | 1300.982075 | 0.310952462 | 1.06E-05 | 0.000165512 |
| ENSMUST00000163075.7,Pip5k1c-003,protein_coding,ENSMUSG00000034902.17,protein_coding | 7606.567565 | -0.203409143 | 1.06E-05 | 0.000165512 |
| ENSMUST00000047099.11,Pif1-001,protein_coding,ENSMUSG00000041064.13,protein_coding | 828.859999 | -0.739107888 | 1.06E-05 | 0.000165516 |
| ENSMUST00000185145.7,Lig1-014,protein_coding,ENSMUSG00000056394.17,protein_coding | 1534.965903 | 0.377647039 | 1.06E-05 | 0.0001657 |
| ENSMUST00000212771.1,Tomm20-002,protein_coding,ENSMUSG00000093904.2,protein_coding | 9971.395648 | 0.258941078 | 1.07E-05 | 0.000166744 |
| ENSMUST00000046038.7,Slc25a28-201,protein_coding,ENSMUSG00000040414.7,protein_coding | 4614.474382 | 0.248632485 | 1.07E-05 | 0.000166997 |
| ENSMUST00000118470.7,Skil-001,protein_coding,ENSMUSG00000027660.16,protein_coding | 5876.88391 | -0.34579313 | 1.08E-05 | 0.000168632 |
| ENSMUST00000217570.1,Adpgk-001,protein_coding,ENSMUSG00000025236.11,protein_coding | 3628.971476 | 0.356203504 | 1.08E-05 | 0.000168632 |
| ENSMUST00000138561.7,Cdk5rap2-006,protein_coding,ENSMUSG00000039298.16,protein_coding | 181.0475804 | 0.598032133 | 1.09E-05 | 0.000168935 |
| ENSMUST00000045884.16,Klhdc4-001,protein_coding,ENSMUSG00000040263.16,protein_coding | 5258.633286 | -0.266319002 | 1.09E-05 | 0.000169363 |
| ENSMUST00000138323.1,Pfdn2-003,processed_transcript,ENSMUSG00000006412.10,protein_coding | 2747.835835 | 0.267521836 | 1.09E-05 | 0.000169669 |
| ENSMUST00000030192.4,Hint2-001,protein_coding,ENSMUSG00000028470.10,protein_coding | 1322.85616 | 0.265316333 | 1.10E-05 | 0.000170068 |
| ENSMUST00000123139.1,Mrps35-005,processed_transcript,ENSMUSG00000040112.9,protein_coding | 1510.997284 | 0.222702881 | 1.10E-05 | 0.000170068 |
| ENSMUST00000125919.7,Dlat-004,processed_transcript,ENSMUSG00000000168.9,protein_coding | 1346.161571 | 0.263618064 | 1.10E-05 | 0.000170068 |
| ENSMUST00000085741.1,Ttc32-201,protein_coding,ENSMUSG00000066637.1,protein_coding | 467.1513317 | 0.336083561 | 1.10E-05 | 0.000170068 |
| ENSMUST00000100339.8,Commd6-001,protein_coding,ENSMUSG00000075486.9,protein_coding | 1212.24456 | 0.283466462 | 1.10E-05 | 0.000170109 |
| ENSMUST00000073441.12,Smg7-002,protein_coding,ENSMUSG00000042772.15,protein_coding | 8100.013697 | 0.185543503 | 1.10E-05 | 0.000170968 |
| ENSMUST00000194252.5,Ncam1-006,protein_coding,ENSMUSG00000039542.16,protein_coding | 945.1062352 | 0.425672555 | 1.10E-05 | 0.000170998 |
| ENSMUST00000103148.7,Pcgf2-002,protein_coding,ENSMUSG00000018537.16,protein_coding | 1747.009505 | 0.315124219 | 1.11E-05 | 0.000172053 |
| ENSMUST00000155901.7,Cnot6l-001,protein_coding,ENSMUSG00000034724.17,protein_coding | 4784.643619 | -0.441431386 | 1.12E-05 | 0.000172914 |
| ENSMUST00000108127.3,3110043O21Rik-001,protein_coding,ENSMUSG00000028300.14,protein_coding | 2509.390545 | 0.362703156 | 1.12E-05 | 0.000173116 |
| ENSMUST00000152733.1,Ybx1-002,processed_transcript,ENSMUSG00000028639.14,protein_coding | 1012.63089 | -0.776220226 | 1.12E-05 | 0.000173116 |
| ENSMUST00000109136.2,Cass4-003,protein_coding,ENSMUSG00000074570.13,protein_coding | 110.373823 | 0.808953864 | 1.12E-05 | 0.000173235 |
| ENSMUST00000031621.10,Ccz1-001,protein_coding,ENSMUSG00000029617.10,protein_coding | 9193.684724 | -0.137410739 | 1.12E-05 | 0.000173394 |
| ENSMUST00000105256.8,Ctdsp2-202,protein_coding,ENSMUSG00000078429.8,protein_coding | 14603.1622 | 0.146943044 | 1.14E-05 | 0.000175504 |
| ENSMUST00000133422.7,Afdn-016,retained_intron,ENSMUSG00000068036.14,protein_coding | 51.37351251 | -0.799919394 | 1.14E-05 | 0.000175504 |
| ENSMUST00000173524.7,Dtnb-010,processed_transcript,ENSMUSG00000071454.13,protein_coding | 40.33625627 | 0.801401194 | 1.15E-05 | 0.000177393 |
| ENSMUST00000021592.15,Cdca7l-001,protein_coding,ENSMUSG00000021175.15,protein_coding | 6796.181391 | 0.162305945 | 1.15E-05 | 0.000177679 |
| ENSMUST00000181045.7,4933404O12Rik-001,lincRNA,ENSMUSG00000097908.7,lincRNA | 838.2200906 | 0.270668068 | 1.16E-05 | 0.000178236 |
| ENSMUST00000017922.8,Cox7c-003,processed_transcript,ENSMUSG00000017778.14,protein_coding | 740.7794523 | 0.299026579 | 1.16E-05 | 0.000179369 |
| ENSMUST00000075253.12,Cyp4f13-001,protein_coding,ENSMUSG00000024055.14,protein_coding | 647.3692064 | 0.343578769 | 1.17E-05 | 0.000179854 |
| ENSMUST00000132983.2,Srsf1-004,retained_intron,ENSMUSG00000018379.17,protein_coding | 2409.777944 | -0.267725249 | 1.17E-05 | 0.000179949 |
| ENSMUST00000001672.11,Ifrd1-001,protein_coding,ENSMUSG00000001627.11,protein_coding | 8805.514833 | 0.154829691 | 1.17E-05 | 0.000180673 |
| ENSMUST00000119559.7,Lancl1-003,protein_coding,ENSMUSG00000026000.16,protein_coding | 3900.356155 | 0.252612718 | 1.17E-05 | 0.000180784 |
| ENSMUST00000174113.7,Sumo3-008,nonsense_mediated_decay,ENSMUSG00000020265.15,protein_coding | 368.9460285 | 0.560734871 | 1.18E-05 | 0.00018111 |
| ENSMUST00000094065.4,Tmem256-001,protein_coding,ENSMUSG00000070394.10,protein_coding | 2171.236196 | 0.29868188 | 1.18E-05 | 0.00018122 |
| ENSMUST00000025243.4,Iws1-001,protein_coding,ENSMUSG00000024384.4,protein_coding | 2740.155748 | -0.467695635 | 1.20E-05 | 0.000184312 |
| ENSMUST00000020365.14,Mum1-001,protein_coding,ENSMUSG00000020156.16,protein_coding | 5574.491783 | 0.198447756 | 1.20E-05 | 0.000184519 |
| ENSMUST00000001059.8,Ern1-001,protein_coding,ENSMUSG00000020715.9,protein_coding | 10291.38127 | 0.273131524 | 1.20E-05 | 0.000184519 |
| ENSMUST00000130278.1,Slc43a3-003,protein_coding,ENSMUSG00000027074.14,protein_coding | 392.2798207 | 0.406585988 | 1.20E-05 | 0.000184548 |
| ENSMUST00000151120.8,Ctsd-001,protein_coding,ENSMUSG00000007891.16,protein_coding | 24269.5116 | 0.194655742 | 1.20E-05 | 0.000184624 |
| ENSMUST00000081650.13,Rpl3-001,protein_coding,ENSMUSG00000060036.13,protein_coding | 173672.4711 | 0.209436264 | 1.20E-05 | 0.000184761 |
| ENSMUST00000074525.9,Gulp1-001,protein_coding,ENSMUSG00000056870.9,protein_coding | 690.5782934 | -0.463232217 | 1.20E-05 | 0.000184763 |
| ENSMUST00000134929.7,RP23-220F20.2-003,protein_coding,ENSMUSG00000110344.1,protein_coding | 995.8823514 | -0.276337755 | 1.21E-05 | 0.000185029 |
| ENSMUST00000059802.6,Rpl29-002,protein_coding,ENSMUSG00000048758.14,protein_coding | 17453.94191 | 0.186264968 | 1.21E-05 | 0.000185439 |
| ENSMUST00000106291.9,Ube2d3-002,protein_coding,ENSMUSG00000078578.9,protein_coding | 19674.43582 | 0.151065963 | 1.21E-05 | 0.000185922 |
| ENSMUST00000200058.4,Anapc5-010,retained_intron,ENSMUSG00000029472.13,protein_coding | 96.57855927 | 0.689452323 | 1.22E-05 | 0.000186734 |
| ENSMUST00000033121.6,Nomo1-001,protein_coding,ENSMUSG00000030835.6,protein_coding | 32141.39468 | 0.197280383 | 1.22E-05 | 0.000186735 |
| ENSMUST00000032194.10,Bhlhe40-001,protein_coding,ENSMUSG00000030103.11,protein_coding | 9628.409637 | 0.175818103 | 1.22E-05 | 0.000187106 |
| ENSMUST00000004634.6,Slc1a4-001,protein_coding,ENSMUSG00000020142.12,protein_coding | 3831.8859 | 0.661455845 | 1.23E-05 | 0.000188262 |
| ENSMUST00000037853.4,Kdelc2-001,protein_coding,ENSMUSG00000034487.4,protein_coding | 3800.497788 | 0.262472789 | 1.25E-05 | 0.000191265 |
| ENSMUST00000005255.8,Wisp1-001,protein_coding,ENSMUSG00000005124.9,protein_coding | 6010.486433 | -0.212822302 | 1.25E-05 | 0.000191265 |
| ENSMUST00000111962.7,Hnrnpa3-001,protein_coding,ENSMUSG00000059005.13,protein_coding | 20208.44533 | -0.493791037 | 1.25E-05 | 0.000191318 |
| ENSMUST00000207849.1,Serpinh1-004,protein_coding,ENSMUSG00000070436.12,protein_coding | 58808.64604 | 0.206070457 | 1.25E-05 | 0.000191318 |
| ENSMUST00000056482.13,Itsn1-003,protein_coding,ENSMUSG00000022957.19,protein_coding | 7190.336832 | -0.181720197 | 1.25E-05 | 0.000191318 |
| ENSMUST00000179062.7,Mbtps2-201,protein_coding,ENSMUSG00000046873.18,protein_coding | 229.3441632 | -0.687172137 | 1.25E-05 | 0.000191539 |
| ENSMUST00000208581.1,Qars-032,protein_coding,ENSMUSG00000032604.16,protein_coding | 249.2920995 | -0.714894542 | 1.26E-05 | 0.00019247 |
| ENSMUST00000150183.8,Ccdc34-004,protein_coding,ENSMUSG00000027160.16,protein_coding | 58.0253605 | 0.742046366 | 1.26E-05 | 0.000192676 |
| ENSMUST00000051867.6,Lsm6-001,protein_coding,ENSMUSG00000031683.16,protein_coding | 1941.52767 | -0.304546899 | 1.26E-05 | 0.000192676 |
| ENSMUST00000209509.1,Nucb1-006,retained_intron,ENSMUSG00000030824.17,protein_coding | 954.9552338 | 0.32496031 | 1.26E-05 | 0.000192969 |
| ENSMUST00000056032.7,E4f1-201,protein_coding,ENSMUSG00000024137.8,protein_coding | 2458.117727 | 0.320325454 | 1.27E-05 | 0.000193115 |
| ENSMUST00000058578.7,Pgrmc2-001,protein_coding,ENSMUSG00000049940.7,protein_coding | 10137.03428 | 0.169494992 | 1.28E-05 | 0.000194779 |
| ENSMUST00000165935.1,Pphln1-204,protein_coding,ENSMUSG00000036167.15,protein_coding | 670.5697977 | -0.541450784 | 1.28E-05 | 0.000195263 |
| ENSMUST00000115331.9,Prkcsh-003,protein_coding,ENSMUSG00000003402.14,protein_coding | 6914.341508 | 0.233540069 | 1.30E-05 | 0.000198155 |
| ENSMUST00000212645.1,Wwp2-009,retained_intron,ENSMUSG00000031930.11,protein_coding | 178.7968487 | 0.753188381 | 1.30E-05 | 0.000198465 |
| ENSMUST00000085585.11,Lsm14a-001,protein_coding,ENSMUSG00000066568.12,protein_coding | 12887.16634 | 0.227786238 | 1.30E-05 | 0.000198638 |
| ENSMUST00000060711.7,Epm2aip1-001,protein_coding,ENSMUSG00000046785.8,protein_coding | 2767.20211 | -0.370135485 | 1.31E-05 | 0.000198756 |
| ENSMUST00000023353.3,Mcm4-201,protein_coding,ENSMUSG00000022673.4,protein_coding | 36054.33931 | 0.132926911 | 1.31E-05 | 0.000198794 |
| ENSMUST00000036407.4,Fam160b1-201,protein_coding,ENSMUSG00000033478.4,protein_coding | 7745.622867 | 0.158079177 | 1.31E-05 | 0.000198817 |
| ENSMUST00000081170.8,Sgcb-001,protein_coding,ENSMUSG00000029156.11,protein_coding | 1425.554088 | 0.247621188 | 1.31E-05 | 0.000199035 |
| ENSMUST00000001834.3,Rtcb-001,protein_coding,ENSMUSG00000001783.3,protein_coding | 26777.3411 | 0.187941041 | 1.31E-05 | 0.000199035 |
| ENSMUST00000087052.10,Fam92a-002,protein_coding,ENSMUSG00000028218.19,protein_coding | 1306.263434 | -0.473894212 | 1.32E-05 | 0.000199842 |
| ENSMUST00000004097.15,Cux1-001,protein_coding,ENSMUSG00000029705.17,protein_coding | 8468.620132 | -0.196359623 | 1.32E-05 | 0.00020011 |
| ENSMUST00000113690.7,Tpm1-013,protein_coding,ENSMUSG00000032366.15,protein_coding | 828.8929546 | -0.322315553 | 1.32E-05 | 0.000200179 |
| ENSMUST00000034947.6,Ppib-001,protein_coding,ENSMUSG00000032383.7,protein_coding | 22502.69965 | 0.186552433 | 1.33E-05 | 0.000202447 |
| ENSMUST00000051186.8,Prkaa1-201,protein_coding,ENSMUSG00000050697.8,protein_coding | 10041.72501 | -0.232908414 | 1.33E-05 | 0.000202447 |
| ENSMUST00000045296.5,Siah1a-001,protein_coding,ENSMUSG00000036840.5,protein_coding | 2148.602322 | 0.316795878 | 1.34E-05 | 0.000203782 |
| ENSMUST00000025846.14,Ppp6r3-201,protein_coding,ENSMUSG00000024908.14,protein_coding | 1995.119885 | -0.372698392 | 1.35E-05 | 0.000204019 |
| ENSMUST00000050552.14,Bzw1-001,protein_coding,ENSMUSG00000051223.14,protein_coding | 29193.31987 | -0.300305912 | 1.35E-05 | 0.000204593 |
| ENSMUST00000107425.7,Aen-001,protein_coding,ENSMUSG00000030609.18,protein_coding | 352.9047514 | -0.654773534 | 1.35E-05 | 0.000205083 |
| ENSMUST00000077915.9,Rpl38-002,protein_coding,ENSMUSG00000057322.12,protein_coding | 9297.875619 | 0.204756972 | 1.36E-05 | 0.000206064 |
| ENSMUST00000025773.3,Pold4-201,protein_coding,ENSMUSG00000024854.3,protein_coding | 3215.87144 | 0.232197293 | 1.37E-05 | 0.000206865 |
| ENSMUST00000180459.2,Eif3c-003,protein_coding,ENSMUSG00000030738.11,protein_coding | 4046.213543 | 0.282238456 | 1.37E-05 | 0.000207497 |
| ENSMUST00000213610.1,Fignl2-001,protein_coding,ENSMUSG00000095440.2,protein_coding | 604.3225468 | -0.357109908 | 1.37E-05 | 0.000207497 |
| ENSMUST00000025903.5,Rps6ka4-201,protein_coding,ENSMUSG00000024952.12,protein_coding | 3980.824862 | -0.324208607 | 1.37E-05 | 0.000207828 |
| ENSMUST00000110329.7,Azin1-002,protein_coding,ENSMUSG00000037458.14,protein_coding | 13454.76414 | -0.209777417 | 1.38E-05 | 0.000208533 |
| ENSMUST00000040783.10,Usp40-003,protein_coding,ENSMUSG00000005501.14,protein_coding | 3848.11428 | -0.143045202 | 1.39E-05 | 0.000210343 |
| ENSMUST00000051934.5,Gskip-201,protein_coding,ENSMUSG00000044715.6,protein_coding | 1623.690037 | -0.225316669 | 1.40E-05 | 0.000211204 |
| ENSMUST00000170068.8,Sirt2-009,protein_coding,ENSMUSG00000015149.14,protein_coding | 290.3002736 | 0.649302398 | 1.40E-05 | 0.000211374 |
| ENSMUST00000102917.10,Col15a1-001,protein_coding,ENSMUSG00000028339.17,protein_coding | 552.0838436 | 0.413020417 | 1.41E-05 | 0.000212317 |

| | | | | |
|---|---|---|---|---|
| ENSMUST00000199531.4,Sh3glb1-002,protein_coding,ENSMUSG00000037062.13,protein_coding | 1449.141984 | 0.311634306 | 1.43E-05 | 0.000215133 |
| ENSMUST00000030455.14,Akr1a1-001,protein_coding,ENSMUSG00000028692.14,protein_coding | 51688.37324 | 0.206176882 | 1.43E-05 | 0.000216355 |
| ENSMUST00000024755.5,Clic5-201,protein_coding,ENSMUSG00000023959.8,protein_coding | 359.5823723 | 0.437626713 | 1.44E-05 | 0.00021698 |
| ENSMUST00000153038.7,Ywhae-003,retained_intron,ENSMUSG00000020849.13,protein_coding | 6230.161473 | 0.176130375 | 1.44E-05 | 0.000217244 |
| ENSMUST00000119896.1,Ssr3-002,protein_coding,ENSMUSG00000027828.12,protein_coding | 169.9495443 | -0.685169577 | 1.45E-05 | 0.000217762 |
| ENSMUST00000153230.7,Pdlim7-006,retained_intron,ENSMUSG00000021493.15,protein_coding | 679.6912081 | -0.357572458 | 1.45E-05 | 0.000217762 |
| ENSMUST00000201069.1,Haus3-003,retained_intron,ENSMUSG00000079555.2,protein_coding | 117.9547971 | -0.722933511 | 1.45E-05 | 0.000218736 |
| ENSMUST00000173075.1,H2-K1-010,protein_coding,ENSMUSG00000061232.15,protein_coding | 197.5633703 | 0.608378335 | 1.45E-05 | 0.000218916 |
| ENSMUST00000043775.7,Kdm3b-201,protein_coding,ENSMUSG00000038773.10,protein_coding | 11160.05159 | 0.186738692 | 1.46E-05 | 0.000219851 |
| ENSMUST00000015434.14,Ubl4a-005,nonsense_mediated_decay,ENSMUSG00000015290.15,protein_coding | 716.9108542 | 0.411657626 | 1.46E-05 | 0.000219855 |
| ENSMUST00000109654.7,Grb10-009,protein_coding,ENSMUSG00000020176.17,protein_coding | 305.8914326 | 0.751562298 | 1.47E-05 | 0.000221122 |
| ENSMUST00000143203.7,Stxbp4-001,protein_coding,ENSMUSG00000020546.14,protein_coding | 2152.209398 | -0.230078285 | 1.47E-05 | 0.000221516 |
| ENSMUST00000049385.13,Eif1-001,protein_coding,ENSMUSG00000035530.13,protein_coding | 93152.83136 | 0.196699248 | 1.48E-05 | 0.00022248 |
| ENSMUST00000043990.13,Edc3-001,protein_coding,ENSMUSG00000038957.13,protein_coding | 3025.257187 | 0.212660622 | 1.48E-05 | 0.00022264 |
| ENSMUST00000129427.1,Pdk1-008,processed_transcript,ENSMUSG00000006494.11,protein_coding | 93.41899213 | 0.729002239 | 1.49E-05 | 0.000223362 |
| ENSMUST00000049312.13,Tmem55b-002,protein_coding,ENSMUSG00000035953.13,protein_coding | 3480.722778 | 0.270648883 | 1.49E-05 | 0.000223367 |
| ENSMUST00000043893.12,Tbc1d1-001,protein_coding,ENSMUSG00000029174.18,protein_coding | 805.1862601 | -0.460434378 | 1.49E-05 | 0.000223671 |
| ENSMUST00000142964.7,Mpg-002,nonsense_mediated_decay,ENSMUSG00000020287.15,protein_coding | 289.6837351 | 0.468125467 | 1.49E-05 | 0.000223728 |
| ENSMUST00000060067.11,Dnaja3-001,protein_coding,ENSMUSG00000004069.16,protein_coding | 3279.78129 | 0.439327536 | 1.50E-05 | 0.000224404 |
| ENSMUST00000030862.4,Draxin-001,protein_coding,ENSMUSG00000029005.4,protein_coding | 117.5386641 | -0.705012912 | 1.50E-05 | 0.000224418 |
| ENSMUST00000170720.1,Sptlc2-003,processed_transcript,ENSMUSG00000021036.10,protein_coding | 1252.713198 | 0.276144256 | 1.50E-05 | 0.000224418 |
| ENSMUST00000106633.9,Slc39a11-004,protein_coding,ENSMUSG00000041654.15,protein_coding | 612.7041562 | 0.534297444 | 1.50E-05 | 0.000224763 |
| ENSMUST00000053913.12,Dgkq-001,protein_coding,ENSMUSG00000004815.12,protein_coding | 975.05255 | -0.354347841 | 1.50E-05 | 0.000225219 |
| ENSMUST00000154266.2,Tes-004,protein_coding,ENSMUSG00000029552.19,protein_coding | 255.6413231 | 0.667823545 | 1.50E-05 | 0.000225267 |
| ENSMUST00000106406.8,Kdm4a-001,protein_coding,ENSMUSG00000033326.15,protein_coding | 15393.55388 | 0.308398968 | 1.51E-05 | 0.000225466 |
| ENSMUST00000020500.13,Appl2-001,protein_coding,ENSMUSG00000020263.14,protein_coding | 1483.715733 | 0.267736282 | 1.51E-05 | 0.00022552 |
| ENSMUST00000156130.1,Fbxw2-009,processed_transcript,ENSMUSG00000035949.15,protein_coding | 180.5700856 | -0.478073925 | 1.51E-05 | 0.000226237 |
| ENSMUST00000106354.8,Sept9-006,protein_coding,ENSMUSG00000059248.13,protein_coding | 6270.268676 | 0.242158766 | 1.51E-05 | 0.000226237 |
| ENSMUST00000063414.8,Tbc1d22a-201,protein_coding,ENSMUSG00000051864.9,protein_coding | 5018.818983 | 0.186052426 | 1.52E-05 | 0.000226701 |
| ENSMUST00000148968.7,Tmem29-002,processed_transcript,ENSMUSG00000041353.12,protein_coding | 260.8985991 | -0.606252477 | 1.52E-05 | 0.000226759 |
| ENSMUST00000045110.13,Dstyk-001,protein_coding,ENSMUSG00000042046.15,protein_coding | 4882.388371 | 0.211661804 | 1.54E-05 | 0.000230362 |
| ENSMUST00000156700.1,Kctd9-002,nonsense_mediated_decay,ENSMUSG00000034327.17,protein_coding | 1791.339686 | 0.284855034 | 1.54E-05 | 0.00023061 |
| ENSMUST00000106769.7,St7l-002,protein_coding,ENSMUSG00000045576.16,protein_coding | 734.3065891 | 0.411744478 | 1.55E-05 | 0.000231123 |
| ENSMUST00000108980.8,Gm14305-001,protein_coding,ENSMUSG00000078878.9,protein_coding | 492.2327523 | -0.641786554 | 1.55E-05 | 0.000231417 |
| ENSMUST00000026416.13,Cdk2-202,protein_coding,ENSMUSG00000025358.15,protein_coding | 4694.214841 | 0.19197279 | 1.56E-05 | 0.000232146 |
| ENSMUST00000170516.8,Rps6ka4-202,protein_coding,ENSMUSG00000024952.12,protein_coding | 11244.96912 | -0.202151817 | 1.56E-05 | 0.000232439 |
| ENSMUST00000139846.1,Eif1-003,processed_transcript,ENSMUSG00000035530.13,protein_coding | 1065.691377 | 0.2888703 | 1.56E-05 | 0.000233186 |
| ENSMUST00000069536.11,Tcf7l1-001,protein_coding,ENSMUSG00000055799.13,protein_coding | 3939.095615 | 0.156538961 | 1.57E-05 | 0.000233466 |
| ENSMUST00000047910.14,Metap2-001,protein_coding,ENSMUSG00000036112.15,protein_coding | 2054.737257 | -0.459235449 | 1.57E-05 | 0.000233799 |
| ENSMUST00000162004.7,Hps1-001,protein_coding,ENSMUSG00000025188.15,protein_coding | 3085.68426 | 0.257100897 | 1.57E-05 | 0.000234174 |
| ENSMUST00000030263.8,St3gal3-001,protein_coding,ENSMUSG00000028538.12,protein_coding | 1712.236608 | -0.289914733 | 1.57E-05 | 0.000234289 |
| ENSMUST00000097625.9,Pam-201,protein_coding,ENSMUSG00000026335.16,protein_coding | 1202.68018 | -0.58403321 | 1.58E-05 | 0.000234571 |
| ENSMUST00000177371.7,Asap1-004,protein_coding,ENSMUSG00000022377.16,protein_coding | 11217.29503 | -0.198738665 | 1.58E-05 | 0.000234578 |
| ENSMUST00000054599.7,Sprr1a-001,protein_coding,ENSMUSG00000050359.7,protein_coding | 378.2239003 | -0.429503015 | 1.58E-05 | 0.000235034 |
| ENSMUST00000185370.6,Spp0-005,nonsense_mediated_decay,ENSMUSG00000029797.13,protein_coding | 99.10154696 | 0.75929151 | 1.59E-05 | 0.000236371 |
| ENSMUST00000067098.6,Ffar4-201,protein_coding,ENSMUSG00000054200.6,protein_coding | 167.2224062 | 0.523202384 | 1.60E-05 | 0.000237029 |
| ENSMUST00000001706.6,Hoxc9-001,protein_coding,ENSMUSG00000036139.6,protein_coding | 1639.249207 | -0.24027446 | 1.60E-05 | 0.00023727 |
| ENSMUST00000030003.9,Exosc3-001,protein_coding,ENSMUSG00000028322.11,protein_coding | 3962.048766 | 0.178504177 | 1.61E-05 | 0.000238459 |
| ENSMUST00000041683.8,Usp22-001,protein_coding,ENSMUSG00000042506.15,protein_coding | 28132.26161 | 0.16527177 | 1.61E-05 | 0.000238952 |
| ENSMUST00000065371.13,Fam53b-006,protein_coding,ENSMUSG00000030956.15,protein_coding | 467.2008918 | 0.422612254 | 1.62E-05 | 0.000239768 |
| ENSMUST00000025300.12,Nr3c1-002,protein_coding,ENSMUSG00000024431.14,protein_coding | 246.8070815 | 0.64605053 | 1.62E-05 | 0.000239984 |
| ENSMUST00000108433.7,Slc43a2-002,protein_coding,ENSMUSG00000038178.16,protein_coding | 828.3392975 | -0.309601646 | 1.62E-05 | 0.000240405 |
| ENSMUST00000021170.8,Mxra7-001,protein_coding,ENSMUSG00000020814.13,protein_coding | 2793.035217 | -0.308230884 | 1.63E-05 | 0.000241461 |
| ENSMUST00000167140.7,Mpst-202,protein_coding,ENSMUSG00000071711.11,protein_coding | 2725.691898 | 0.249113172 | 1.64E-05 | 0.000243627 |
| ENSMUST00000124678.1,Mlec-002,processed_transcript,ENSMUSG00000048578.11,protein_coding | 731.766446 | -0.283413226 | 1.65E-05 | 0.000244289 |
| ENSMUST00000211690.1,Ankrd10-006,retained_intron,ENSMUSG00000031508.14,protein_coding | 621.3607989 | -0.451306827 | 1.66E-05 | 0.000245718 |
| ENSMUST00000136139.7,Gabpb2-001,protein_coding,ENSMUSG00000038766.16,protein_coding | 3159.122845 | -0.32790024 | 1.66E-05 | 0.000245732 |
| ENSMUST00000046259.13,Sun2-002,protein_coding,ENSMUSG00000042524.13,protein_coding | 7880.01338 | 0.280509775 | 1.66E-05 | 0.000246035 |
| ENSMUST00000058716.13,Sun1-001,protein_coding,ENSMUSG00000036817.14,protein_coding | 8023.438674 | 0.197465184 | 1.66E-05 | 0.000246165 |
| ENSMUST00000051925.4,Prdx6-002,protein_coding,ENSMUSG00000026701.15,protein_coding | 1298.429063 | 0.448996396 | 1.67E-05 | 0.000247018 |
| ENSMUST00000145480.1,Strn-003,protein_coding,ENSMUSG00000024077.15,protein_coding | 81.91537275 | 0.733272436 | 1.67E-05 | 0.000247046 |
| ENSMUST00000156862.1,Tead3-001,protein_coding,ENSMUSG00000002249.18,protein_coding | 2576.043127 | -0.191570802 | 1.67E-05 | 0.00024732 |
| ENSMUST00000177527.1,Comtd1-004,protein_coding,ENSMUSG00000021773.10,protein_coding | 334.8543892 | -0.435366029 | 1.68E-05 | 0.0002477 |
| ENSMUST00000087517.9,Dido1-002,protein_coding,ENSMUSG00000038914.15,protein_coding | 3188.619645 | -0.483967921 | 1.69E-05 | 0.000249015 |
| ENSMUST00000058119.8,Arxes2-001,protein_coding,ENSMUSG00000048040.8,protein_coding | 383.1745387 | -0.371742153 | 1.69E-05 | 0.000249015 |
| ENSMUST00000162819.7,Wdr26-001,protein_coding,ENSMUSG00000038733.13,protein_coding | 8640.025164 | -0.326549632 | 1.70E-05 | 0.000250523 |
| ENSMUST00000028771.7,Nusap1-002,protein_coding,ENSMUSG00000027306.15,protein_coding | 1388.694906 | -0.499666533 | 1.70E-05 | 0.000250523 |
| ENSMUST00000121455.2,Gm12346-001,transcribed_processed_pseudogene,ENSMUSG00000083899.2,transcribed_processed_pseudogene | 605.266781 | -0.593887346 | 1.72E-05 | 0.000254029 |
| ENSMUST00000182232.7,Snhg5-002,lincRNA,ENSMUSG00000097195.9,processed_transcript | 715.1503743 | 0.538874044 | 1.73E-05 | 0.000255099 |
| ENSMUST00000106609.7,Clcc1-002,protein_coding,ENSMUSG00000027884.16,protein_coding | 3302.012634 | 0.163457181 | 1.73E-05 | 0.000255263 |
| ENSMUST00000005406.10,App-201,protein_coding,ENSMUSG00000022892.10,protein_coding | 15511.65026 | 0.206715622 | 1.74E-05 | 0.000255861 |
| ENSMUST00000058089.12,Raly-001,protein_coding,ENSMUSG00000027593.15,protein_coding | 25365.18963 | 0.206066793 | 1.74E-05 | 0.00025631 |
| ENSMUST00000060989.8,Sorl1-001,protein_coding,ENSMUSG00000049313.8,protein_coding | 415.4683084 | -0.336980824 | 1.74E-05 | 0.000256601 |
| ENSMUST00000109594.7,Slc1a4-201,protein_coding,ENSMUSG00000020142.12,protein_coding | 7779.938628 | 0.434314172 | 1.75E-05 | 0.000256933 |
| ENSMUST00000036187.8,Qsox2-001,protein_coding,ENSMUSG00000036327.18,protein_coding | 1440.919934 | 0.26579741 | 1.75E-05 | 0.000258073 |
| ENSMUST00000105364.7,Ndufs7-001,protein_coding,ENSMUSG00000020153.14,protein_coding | 450.7513851 | -0.438573871 | 1.76E-05 | 0.000258586 |
| ENSMUST00000022587.9,Tsc22d1-001,protein_coding,ENSMUSG00000022010.19,protein_coding | 31898.49217 | -0.252761978 | 1.76E-05 | 0.000258655 |
| ENSMUST00000129900.1,Senp5-003,protein_coding,ENSMUSG00000022772.11,protein_coding | 945.5798334 | 0.397879211 | 1.76E-05 | 0.000259209 |
| ENSMUST00000193460.5,Diexf-007,protein_coding,ENSMUSG00000016181.9,protein_coding | 345.4196703 | 0.365244525 | 1.78E-05 | 0.000261172 |
| ENSMUST00000010191.12,Hyal2-201,protein_coding,ENSMUSG00000010047.12,protein_coding | 2735.950587 | -0.23617209 | 1.78E-05 | 0.000261461 |
| ENSMUST00000093517.6,Casp3-003,protein_coding,ENSMUSG00000031628.9,protein_coding | 1089.179672 | -0.42441617 | 1.80E-05 | 0.000263867 |
| ENSMUST00000027381.12,Pecr-001,protein_coding,ENSMUSG00000026189.13,protein_coding | 1316.71579 | -0.249756109 | 1.80E-05 | 0.000264255 |
| ENSMUST00000055006.11,Trim16-001,protein_coding,ENSMUSG00000047821.16,protein_coding | 2041.920475 | 0.291615877 | 1.81E-05 | 0.000265002 |
| ENSMUST00000178948.1,Gm21967-201,protein_coding,ENSMUSG00000094114.1,protein_coding | 701.7207041 | -0.314001171 | 1.81E-05 | 0.000265277 |
| ENSMUST00000167921.1,Zbtb7c-202,protein_coding,ENSMUSG00000044646.14,protein_coding | 464.8760377 | 0.498752453 | 1.81E-05 | 0.000265549 |
| ENSMUST00000021941.7,Mxd3-001,protein_coding,ENSMUSG00000021485.13,protein_coding | 1285.586223 | -0.290004306 | 1.82E-05 | 0.000266678 |
| ENSMUST00000021796.7,Edn1-201,protein_coding,ENSMUSG00000021367.7,protein_coding | 97.8426349 | -0.614847653 | 1.82E-05 | 0.000266915 |
| ENSMUST00000026818.11,Sec11a-001,protein_coding,ENSMUSG00000025724.12,protein_coding | 4627.519328 | 0.16539201 | 1.82E-05 | 0.000266982 |
| ENSMUST00000163588.7,Ston1-201,protein_coding,ENSMUSG00000033855.15,protein_coding | 7949.057951 | -0.241934528 | 1.82E-05 | 0.000266982 |
| ENSMUST00000160225.1,Zfp579-003,processed_transcript,ENSMUSG00000051550.10,protein_coding | 267.8851589 | -0.611341393 | 1.83E-05 | 0.000268574 |
| ENSMUST00000034204.10,Nudt21-001,protein_coding,ENSMUSG00000031754.10,protein_coding | 4987.765575 | -0.276992103 | 1.84E-05 | 0.000268609 |
| ENSMUST00000093191.2,Spdl1-001,protein_coding,ENSMUSG00000069910.2,protein_coding | 3219.759499 | -0.325037751 | 1.84E-05 | 0.000269375 |

| | | | | |
|---|---|---|---|---|
| ENSMUST00000167977.7,Nfatc1-203,protein_coding,ENSMUSG00000033016.15,protein_coding | 1086.739242 | 0.758384901 | 1.85E-05 | 0.000270246 |
| ENSMUST00000066264.12,Ech1-001,protein_coding,ENSMUSG00000053898.12,protein_coding | 4513.304992 | 0.246043317 | 1.85E-05 | 0.00027083 |
| ENSMUST00000122850.1,Araf-007,protein_coding,ENSMUSG00000001127.12,protein_coding | 111.1255228 | 0.796814712 | 1.85E-05 | 0.000271024 |
| ENSMUST00000065302.7,Cenpj-001,protein_coding,ENSMUSG00000064128.6,protein_coding | 862.9085537 | -0.517308812 | 1.86E-05 | 0.000271923 |
| ENSMUST00000036854.3,Efhd2-001,protein_coding,ENSMUSG00000040659.3,protein_coding | 8961.886268 | 0.191664311 | 1.87E-05 | 0.000272709 |
| ENSMUST00000005711.5,Chmp2a-001,protein_coding,ENSMUSG00000033916.5,protein_coding | 3941.869923 | 0.202845721 | 1.87E-05 | 0.000272882 |
| ENSMUST00000168163.7,Parpbp-002,protein_coding,ENSMUSG00000035365.15,protein_coding | 152.789213 | -0.582542524 | 1.87E-05 | 0.000273047 |
| ENSMUST00000074840.11,Preb-001,protein_coding,ENSMUSG00000045302.13,protein_coding | 1399.249926 | -0.301050762 | 1.89E-05 | 0.000274965 |
| ENSMUST00000126014.1,Cep85-008,retained_intron,ENSMUSG00000037443.13,protein_coding | 42.76896238 | 0.783494166 | 1.89E-05 | 0.000275372 |
| ENSMUST00000144308.7,Zfp384-002,processed_transcript,ENSMUSG00000038346.18,protein_coding | 83.19957381 | -0.701619725 | 1.91E-05 | 0.00027773 |
| ENSMUST00000127401.7,Slc1a5-002,retained_intron,ENSMUSG00000001918.17,protein_coding | 5049.229784 | 0.29691873 | 1.91E-05 | 0.00027773 |
| ENSMUST00000146306.1,Shc1-009,processed_transcript,ENSMUSG00000042626.13,protein_coding | 360.0084379 | 0.655065475 | 1.92E-05 | 0.000279145 |
| ENSMUST00000100538.2,Zc3h3-201,protein_coding,ENSMUSG00000075600.2,protein_coding | 3888.701426 | 0.156043041 | 1.92E-05 | 0.000279176 |
| ENSMUST00000097873.9,C77080-001,protein_coding,ENSMUSG00000050390.12,protein_coding | 1109.262154 | -0.653957275 | 1.93E-05 | 0.000280516 |
| ENSMUST00000024636.14,Fgfr1op-002,protein_coding,ENSMUSG00000069135.11,protein_coding | 2091.50685 | -0.309663316 | 1.93E-05 | 0.000280516 |
| ENSMUST00000010804.3,Plcd1-002,protein_coding,ENSMUSG00000010660.3,protein_coding | 1213.388593 | -0.31388635 | 1.93E-05 | 0.000280892 |
| ENSMUST00000027980.7,Mcm10-006,protein_coding,ENSMUSG00000026669.14,protein_coding | 10962.01653 | 0.15644756 | 1.93E-05 | 0.00028091 |
| ENSMUST00000084097.11,Aurkaip1-001,protein_coding,ENSMUSG00000065990.12,protein_coding | 3792.435512 | -0.252715043 | 1.94E-05 | 0.000281414 |
| ENSMUST00000021324.2,Map3k14-001,protein_coding,ENSMUSG00000020941.7,protein_coding | 2182.245865 | -0.220788179 | 1.94E-05 | 0.000281414 |
| ENSMUST00000057551.13,Slbp-001,protein_coding,ENSMUSG00000004642.13,protein_coding | 7473.758755 | -0.214331971 | 1.95E-05 | 0.000282473 |
| ENSMUST00000171588.1,Lifr-203,protein_coding,ENSMUSG00000054263.11,protein_coding | 4250.434609 | -0.373104974 | 1.94E-05 | 0.000282473 |
| ENSMUST00000153251.7,Fam98c-009,protein_coding,ENSMUSG00000030590.15,protein_coding | 58.27939194 | 0.766074931 | 1.95E-05 | 0.000282552 |
| ENSMUST00000193611.5,Srsf6-004,retained_intron,ENSMUSG00000016921.14,protein_coding | 3291.175851 | 0.233379636 | 1.95E-05 | 0.00028343 |
| ENSMUST00000067646.11,Ilf3-001,protein_coding,ENSMUSG00000032178.14,protein_coding | 7636.036799 | 0.203384566 | 1.95E-05 | 0.000283601 |
| ENSMUST00000167488.8,Bend3-001,protein_coding,ENSMUSG00000038214.15,protein_coding | 4290.957014 | -0.197388015 | 1.96E-05 | 0.000283645 |
| ENSMUST00000154306.1,Vapb-002,processed_transcript,ENSMUSG00000054455.4,protein_coding | 2875.015296 | 0.25675948 | 1.96E-05 | 0.000284452 |
| ENSMUST00000112905.3,Thap4-002,nonsense_mediated_decay,ENSMUSG00000026279.14,protein_coding | 4789.685213 | 0.191100695 | 1.96E-05 | 0.000284621 |
| ENSMUST00000058109.8,Mrps7-001,protein_coding,ENSMUSG00000046756.10,protein_coding | 10569.06912 | -0.232834061 | 1.97E-05 | 0.000284621 |
| ENSMUST00000115050.7,Spire1-203,protein_coding,ENSMUSG00000024533.15,protein_coding | 3037.463501 | 0.315955333 | 1.96E-05 | 0.000284621 |
| ENSMUST00000103188.9,1700037H04Rik-001,protein_coding,ENSMUSG00000027327.16,protein_coding | 6864.890081 | 0.208646748 | 1.97E-05 | 0.000284815 |
| ENSMUST00000082175.6,Trim68-001,protein_coding,ENSMUSG00000073968.4,protein_coding | 233.0555958 | 0.718156123 | 1.97E-05 | 0.00028496 |
| ENSMUST00000141055.1,Otub2-003,retained_intron,ENSMUSG00000021203.15,protein_coding | 230.9684246 | 0.555157534 | 1.98E-05 | 0.00028598 |
| ENSMUST00000170209.7,Mms19-007,retained_intron,ENSMUSG00000025159.8,protein_coding | 390.6654346 | -0.436367464 | 1.98E-05 | 0.000286712 |
| ENSMUST00000109166.7,Aida-001,protein_coding,ENSMUSG00000042901.10,protein_coding | 1714.489629 | -0.311810633 | 2.00E-05 | 0.000288731 |
| ENSMUST00000111248.7,Ext2-007,protein_coding,ENSMUSG00000027198.16,protein_coding | 74.22754645 | 0.713067417 | 2.02E-05 | 0.000291729 |
| ENSMUST00000154078.2,Akap5-201,protein_coding,ENSMUSG00000021057.15,protein_coding | 102.425786 | -0.735294869 | 2.02E-05 | 0.000292043 |
| ENSMUST00000028623.12,Ext2-001,protein_coding,ENSMUSG00000027198.16,protein_coding | 15877.563 | 0.18334899 | 2.03E-05 | 0.000292558 |
| ENSMUST00000059955.14,Yars2-001,protein_coding,ENSMUSG00000022792.16,protein_coding | 2663.615617 | -0.332602286 | 2.04E-05 | 0.000294159 |
| ENSMUST00000036206.13,Ccdc92-001,protein_coding,ENSMUSG00000037979.13,protein_coding | 1104.577568 | -0.351941617 | 2.04E-05 | 0.000294181 |
| ENSMUST00000147397.1,Ears2-003,retained_intron,ENSMUSG00000030871.11,protein_coding | 115.799716 | 0.764844101 | 2.04E-05 | 0.00029463 |
| ENSMUST00000126315.7,Thbs3-002,retained_intron,ENSMUSG00000028047.11,protein_coding | 98.01145003 | -0.639048896 | 2.04E-05 | 0.000294662 |
| ENSMUST00000131070.1,Ide-001,protein_coding,ENSMUSG00000056999.14,protein_coding | 14227.24186 | -0.144822482 | 2.07E-05 | 0.000298 |
| ENSMUST00000175692.7,Rwdd4a-003,nonsense_mediated_decay,ENSMUSG00000031568.16,protein_coding | 1922.488833 | 0.257323667 | 2.07E-05 | 0.00029829 |
| ENSMUST00000186531.1,2810013P06Rik-001,bidirectional_promoter_lncRNA,ENSMUSG00000099881.1,bidirectional_promoter_lncRNA | 1081.23019 | 0.24587019 | 2.07E-05 | 0.000298466 |
| ENSMUST00000129909.7,Uba52-003,protein_coding,ENSMUSG00000090137.8,protein_coding | 7691.46575 | 0.22562265 | 2.08E-05 | 0.000299103 |
| ENSMUST00000022806.9,Bcl2l2-001,protein_coding,ENSMUSG00000089682.9,protein_coding | 3806.833521 | 0.18277136 | 2.08E-05 | 0.000299103 |
| ENSMUST00000042345.6,Ctnna1-201,protein_coding,ENSMUSG00000037815.6,protein_coding | 50795.95502 | 0.167699065 | 2.10E-05 | 0.000301563 |
| ENSMUST00000033577.10,Pbdc1-001,protein_coding,ENSMUSG00000031226.13,protein_coding | 2752.087136 | -0.293919902 | 2.09E-05 | 0.000301563 |
| ENSMUST00000019997.10,Tnfaip3-001,protein_coding,ENSMUSG00000019850.11,protein_coding | 2937.438511 | -0.30027097 | 2.10E-05 | 0.000301819 |
| ENSMUST00000154679.7,Polr3c-005,protein_coding,ENSMUSG00000028099.8,protein_coding | 646.0384704 | -0.488634089 | 2.10E-05 | 0.00030219 |
| ENSMUST00000126184.1,Ywhaz-006,protein_coding,ENSMUSG00000022285.16,protein_coding | 195.2796606 | 0.682683378 | 2.10E-05 | 0.00030219 |
| ENSMUST00000071874.6,Ano6-201,protein_coding,ENSMUSG00000064210.6,protein_coding | 22697.19784 | -0.137301371 | 2.11E-05 | 0.000303918 |
| ENSMUST00000102948.9,E2f3-001,protein_coding,ENSMUSG00000016477.17,protein_coding | 4329.185977 | -0.167997287 | 2.12E-05 | 0.000305018 |
| ENSMUST00000098613.8,Usp3-002,protein_coding,ENSMUSG00000032376.12,protein_coding | 808.6827547 | 0.483526651 | 2.13E-05 | 0.000305146 |
| ENSMUST00000111113.2,Wdr35-002,protein_coding,ENSMUSG00000066643.12,protein_coding | 3276.040514 | -0.152469228 | 2.13E-05 | 0.000305146 |
| ENSMUST00000151894.7,Pfkp-004,retained_intron,ENSMUSG00000021196.13,protein_coding | 464.9053255 | 0.444668971 | 2.13E-05 | 0.000305146 |
| ENSMUST00000048854.7,Zfp647-201,protein_coding,ENSMUSG00000054967.6,protein_coding | 964.6825933 | 0.272376643 | 2.13E-05 | 0.000305146 |
| ENSMUST00000201775.1,Gm20186-003,lincRNA,ENSMUSG00000106874.1,lincRNA | 1071.045629 | -0.596612467 | 2.13E-05 | 0.00030525 |
| ENSMUST00000176686.7,Cpsf6-002,protein_coding,ENSMUSG00000055531.12,protein_coding | 3673.250987 | 0.23602138 | 2.14E-05 | 0.000306475 |
| ENSMUST00000171770.8,Fut8-202,protein_coding,ENSMUSG00000021065.15,protein_coding | 1916.659213 | 0.32198959 | 2.14E-05 | 0.000306777 |
| ENSMUST00000072735.7,Fam173a-201,protein_coding,ENSMUSG00000057411.8,protein_coding | 1802.387977 | 0.250522206 | 2.15E-05 | 0.000308445 |
| ENSMUST00000099967.9,Fyn-002,protein_coding,ENSMUSG00000019843.14,protein_coding | 3402.870486 | 0.370816779 | 2.16E-05 | 0.000308878 |
| ENSMUST00000113369.2,Rcor2-201,protein_coding,ENSMUSG00000024968.13,protein_coding | 1726.396158 | 0.234706215 | 2.18E-05 | 0.000312871 |
| ENSMUST00000021900.13,Sema4d-002,protein_coding,ENSMUSG00000021451.15,protein_coding | 622.064055 | 0.458274607 | 2.19E-05 | 0.000313173 |
| ENSMUST00000061143.14,Map7d1-002,protein_coding,ENSMUSG00000028849.17,protein_coding | 9574.560686 | 0.166752035 | 2.20E-05 | 0.000314127 |
| ENSMUST00000179954.7,Rpl31-005,protein_coding,ENSMUSG00000073702.11,protein_coding | 1005.889509 | -0.263035907 | 2.21E-05 | 0.000315833 |
| ENSMUST00000122216.7,Bola3-002,protein_coding,ENSMUSG00000045160.14,protein_coding | 2580.661295 | 0.216764657 | 2.21E-05 | 0.000315853 |
| ENSMUST00000093321.11,Grb10-005,protein_coding,ENSMUSG00000020176.17,protein_coding | 21887.94253 | 0.203295837 | 2.21E-05 | 0.000315853 |
| ENSMUST00000086423.4,Gm10184-201,protein_coding,ENSMUSG00000066878.4,protein_coding | 973.1943632 | -0.492336392 | 2.21E-05 | 0.000316353 |
| ENSMUST00000180726.1,E030030I06Rik-001,processed_transcript,ENSMUSG00000097327.7,protein_coding | 35.30119095 | 0.758706564 | 2.22E-05 | 0.000317367 |
| ENSMUST00000117890.7,Spsb3-003,protein_coding,ENSMUSG00000024160.16,protein_coding | 1305.42125 | 0.261361436 | 2.22E-05 | 0.000317592 |
| ENSMUST00000127260.7,Mbd2-003,nonsense_mediated_decay,ENSMUSG00000024513.16,protein_coding | 50.45466746 | 0.758123961 | 2.22E-05 | 0.000317592 |
| ENSMUST00000048309.11,Camsap2-201,protein_coding,ENSMUSG00000041570.14,protein_coding | 3783.748935 | -0.351702436 | 2.23E-05 | 0.000318502 |
| ENSMUST00000137446.7,Tm9sf4-002,processed_transcript,ENSMUSG00000068040.10,protein_coding | 261.2634451 | 0.607238129 | 2.23E-05 | 0.000318654 |
| ENSMUST00000059018.13,Fbxo31-001,protein_coding,ENSMUSG00000052934.14,protein_coding | 4216.952692 | 0.224962609 | 2.25E-05 | 0.000320498 |
| ENSMUST00000204592.2,Cct7-007,protein_coding,ENSMUSG00000030007.8,protein_coding | 21188.37382 | 0.276387484 | 2.25E-05 | 0.000321095 |
| ENSMUST00000106489.8,Eef2k-002,protein_coding,ENSMUSG00000035064.17,protein_coding | 1552.687917 | 0.334036733 | 2.25E-05 | 0.000321186 |
| ENSMUST00000169504.7,Tdrd3-002,protein_coding,ENSMUSG00000022019.15,protein_coding | 603.2884328 | -0.495676742 | 2.27E-05 | 0.000322862 |
| ENSMUST00000153936.1,Wipi2-002,retained_intron,ENSMUSG00000029578.15,protein_coding | 374.6862146 | 0.430406023 | 2.28E-05 | 0.000324079 |
| ENSMUST00000068508.12,Spsb3-002,protein_coding,ENSMUSG00000024160.16,protein_coding | 514.7772323 | 0.62184164 | 2.29E-05 | 0.000325713 |
| ENSMUST00000161017.7,Kif26b-002,protein_coding,ENSMUSG00000026494.12,protein_coding | 77.95069278 | -0.649291416 | 2.29E-05 | 0.000326146 |
| ENSMUST00000140447.3,1810010H24Rik-001,protein_coding,ENSMUSG00000078607.5,protein_coding | 157.9643245 | 0.550460592 | 2.29E-05 | 0.000326233 |
| ENSMUST00000023351.10,Mzt2-002,protein_coding,ENSMUSG00000022671.12,protein_coding | 1198.018349 | 0.28172188 | 2.31E-05 | 0.000329069 |
| ENSMUST00000128161.7,Rgs12-004,retained_intron,ENSMUSG00000029101.14,protein_coding | 35.49181621 | 0.770503818 | 2.31E-05 | 0.000329084 |
| ENSMUST00000203302.1,Edem1-004,retained_intron,ENSMUSG00000030104.9,protein_coding | 229.6571788 | 0.584576488 | 2.32E-05 | 0.000329904 |
| ENSMUST00000107420.7,Rgs3-010,protein_coding,ENSMUSG00000059810.18,protein_coding | 189.9311189 | 0.673077672 | 2.33E-05 | 0.000330491 |
| ENSMUST00000151368.1,Qsox1-004,retained_intron,ENSMUSG00000033684.14,protein_coding | 28.60667385 | -0.779560109 | 2.34E-05 | 0.0003327 |
| ENSMUST00000059080.6,Rps21-001,protein_coding,ENSMUSG00000039001.12,protein_coding | 41001.67053 | 0.171414872 | 2.34E-05 | 0.0003327 |
| ENSMUST00000078690.12,Sun1-004,protein_coding,ENSMUSG00000036817.14,protein_coding | 361.0745616 | 0.523364438 | 2.34E-05 | 0.000332722 |
| ENSMUST00000207858.1,Arhgap33-003,protein_coding,ENSMUSG00000036882.7,protein_coding | 313.1261925 | 0.599591987 | 2.35E-05 | 0.000333482 |
| ENSMUST00000027783.13,Desi2-001,protein_coding,ENSMUSG00000026502.13,protein_coding | 4404.872715 | 0.203592008 | 2.36E-05 | 0.000334237 |
| ENSMUST00000025505.5,Dctn4-201,protein_coding,ENSMUSG00000024603.7,protein_coding | 8959.970851 | 0.136545763 | 2.36E-05 | 0.000334927 |

| | | | | |
|---|---|---|---|---|
| ENSMUST00000187103.1,Tuba4a-005,retained_intron,ENSMUSG00000026202.13,protein_coding | 788.852659 | -0.358466269 | 2.37E-05 | 0.000335869 |
| ENSMUST00000166820.7,R3hdm2-001,protein_coding,ENSMUSG00000025404.15,protein_coding | 6786.311968 | 0.261969764 | 2.37E-05 | 0.000335991 |
| ENSMUST00000200585.4,Fgf2-007,protein_coding,ENSMUSG00000037225.13,protein_coding | 711.7854866 | -0.532816065 | 2.39E-05 | 0.000339075 |
| ENSMUST00000208190.1,Nap1l4-002,protein_coding,ENSMUSG00000059119.9,protein_coding | 13904.52488 | -0.201489996 | 2.40E-05 | 0.000340162 |
| ENSMUST00000025239.7,Bin1-201,protein_coding,ENSMUSG00000024381.15,protein_coding | 410.7596991 | 0.541414329 | 2.41E-05 | 0.000340899 |
| ENSMUST00000027684.10,Arl8a-001,protein_coding,ENSMUSG00000026426.10,protein_coding | 8707.434164 | 0.197806306 | 2.41E-05 | 0.000341423 |
| ENSMUST00000177437.7,Foxp1-021,protein_coding,ENSMUSG00000030067.17,protein_coding | 4863.309466 | 0.170926646 | 2.42E-05 | 0.000342672 |
| ENSMUST00000047799.12,Acad11-001,protein_coding,ENSMUSG00000090150.9,protein_coding | 1320.787384 | 0.247866265 | 2.42E-05 | 0.000342672 |
| ENSMUST00000015858.11,Cers2-001,protein_coding,ENSMUSG00000015714.11,protein_coding | 28243.54328 | 0.171902287 | 2.42E-05 | 0.000342806 |
| ENSMUST00000106757.7,Nol11-001,protein_coding,ENSMUSG00000018433.14,protein_coding | 1944.265758 | -0.294223547 | 2.43E-05 | 0.000343403 |
| ENSMUST00000110888.7,Tsc22d1-006,protein_coding,ENSMUSG00000022010.19,protein_coding | 597.4499086 | -0.496506905 | 2.43E-05 | 0.000343878 |
| ENSMUST00000043314.9,Cyp2s1-001,protein_coding,ENSMUSG00000040703.11,protein_coding | 2207.6898 | 0.266209932 | 2.44E-05 | 0.000344827 |
| ENSMUST00000118788.7,Hagh-001,protein_coding,ENSMUSG00000024158.17,protein_coding | 4600.111921 | 0.237566993 | 2.45E-05 | 0.000345351 |
| ENSMUST00000044148.2,Alg2-001,protein_coding,ENSMUSG00000039740.6,protein_coding | 4471.874318 | 0.281203753 | 2.46E-05 | 0.000346927 |
| ENSMUST00000050702.8,Ptprq-001,protein_coding,ENSMUSG00000035916.7,protein_coding | 104.4123426 | -0.59480447 | 2.47E-05 | 0.000348822 |
| ENSMUST00000021450.4,Sgpp1-201,protein_coding,ENSMUSG00000021054.5,protein_coding | 3777.310951 | -0.199094381 | 2.47E-05 | 0.000348822 |
| ENSMUST00000042608.7,Acd-002,protein_coding,ENSMUSG00000038000.10,protein_coding | 1266.400331 | 0.298556874 | 2.48E-05 | 0.000349092 |
| ENSMUST00000090586.11,Phf21a-203,protein_coding,ENSMUSG00000058318.15,protein_coding | 474.7768307 | -0.787108587 | 2.48E-05 | 0.000349193 |
| ENSMUST00000041859.7,Gmds-001,protein_coding,ENSMUSG00000038372.13,protein_coding | 2345.502367 | 0.214466281 | 2.48E-05 | 0.000349193 |
| ENSMUST00000111943.7,Mapk8-004,protein_coding,ENSMUSG00000021936.13,protein_coding | 1012.321212 | 0.303774772 | 2.48E-05 | 0.000349193 |
| ENSMUST00000000356.8,Dazap2-201,protein_coding,ENSMUSG00000000346.8,protein_coding | 23574.4745 | 0.182308171 | 2.48E-05 | 0.000349193 |
| ENSMUST00000053917.4,Ccny-201,protein_coding,ENSMUSG00000024286.7,protein_coding | 8415.244761 | 0.127030169 | 2.48E-05 | 0.000349361 |
| ENSMUST00000149105.1,Tmem161a-003,protein_coding,ENSMUSG00000002342.17,protein_coding | 136.2370088 | 0.646186978 | 2.49E-05 | 0.00035005 |
| ENSMUST00000212377.1,Slc7a6-003,nonsense_mediated_decay,ENSMUSG00000031904.5,protein_coding | 3948.328984 | 0.278384986 | 2.49E-05 | 0.000350178 |
| ENSMUST00000179165.7,Fzd6-202,protein_coding,ENSMUSG00000022297.14,protein_coding | 4479.770885 | -0.27438025 | 2.51E-05 | 0.000352476 |
| ENSMUST00000027125.6,Coq10b-001,protein_coding,ENSMUSG00000025981.13,protein_coding | 2615.298164 | -0.321403364 | 2.53E-05 | 0.00035548 |
| ENSMUST00000128803.7,Spg7-002,retained_intron,ENSMUSG00000000738.18,protein_coding | 271.2765011 | -0.457871141 | 2.53E-05 | 0.000356003 |
| ENSMUST00000097713.1,Plekhm3-001,protein_coding,ENSMUSG00000051344.13,protein_coding | 1683.598969 | -0.173949549 | 2.53E-05 | 0.000356082 |
| ENSMUST00000024762.2,Rsph9-001,protein_coding,ENSMUSG00000023966.8,protein_coding | 250.2961704 | -0.448690142 | 2.53E-05 | 0.00035611 |
| ENSMUST00000202308.3,Sept11-004,protein_coding,ENSMUSG00000058013.11,protein_coding | 1649.993329 | 0.317076083 | 2.54E-05 | 0.000356168 |
| ENSMUST00000025385.6,Hsd17b4-001,protein_coding,ENSMUSG00000024507.6,protein_coding | 5431.378245 | 0.183016016 | 2.54E-05 | 0.000356432 |
| ENSMUST00000110527.8,Fut10-004,protein_coding,ENSMUSG00000046152.16,protein_coding | 413.4046661 | -0.497848111 | 2.54E-05 | 0.000356567 |
| ENSMUST00000134101.7,Mrps5-003,retained_intron,ENSMUSG00000027374.12,protein_coding | 2133.836412 | 0.260313311 | 2.54E-05 | 0.000356686 |
| ENSMUST00000024701.7,Pkmyt1-201,protein_coding,ENSMUSG00000023908.7,protein_coding | 6277.342544 | 0.23581857 | 2.55E-05 | 0.000357263 |
| ENSMUST00000059672.7,Pik3ap1-201,protein_coding,ENSMUSG00000025017.9,protein_coding | 109.1219145 | -0.616364909 | 2.55E-05 | 0.000357263 |
| ENSMUST00000160866.1,Herpud1-005,retained_intron,ENSMUSG00000031770.16,protein_coding | 175.6723632 | 0.608808867 | 2.56E-05 | 0.00035814 |
| ENSMUST00000001802.9,Naglu-001,protein_coding,ENSMUSG00000001751.9,protein_coding | 2057.831286 | 0.348435892 | 2.56E-05 | 0.000358812 |
| ENSMUST00000009102.8,Vps72-001,protein_coding,ENSMUSG00000008958.14,protein_coding | 9268.137885 | 0.159909496 | 2.56E-05 | 0.000358969 |
| ENSMUST00000115579.7,Sgce-001,protein_coding,ENSMUSG00000004631.15,protein_coding | 3284.090353 | 0.220734151 | 2.58E-05 | 0.000361077 |
| ENSMUST00000029380.13,Wwtr1-001,protein_coding,ENSMUSG00000027803.14,protein_coding | 18124.72524 | 0.208415991 | 2.58E-05 | 0.000361143 |
| ENSMUST00000034240.14,Kifc3-001,protein_coding,ENSMUSG00000031788.14,protein_coding | 4396.103775 | -0.161668573 | 2.59E-05 | 0.000362033 |
| ENSMUST00000073109.11,Ctdspl-001,protein_coding,ENSMUSG00000047409.13,protein_coding | 2795.755269 | 0.168468959 | 2.59E-05 | 0.000362335 |
| ENSMUST00000031034.11,Nrbp1-001,protein_coding,ENSMUSG00000029148.15,protein_coding | 10766.65975 | 0.204362557 | 2.59E-05 | 0.000362584 |
| ENSMUST00000060834.11,Alkbh6-001,protein_coding,ENSMUSG00000042831.13,protein_coding | 727.2840255 | -0.314864152 | 2.60E-05 | 0.000362969 |
| ENSMUST00000100329.9,Ndor1-011,protein_coding,ENSMUSG00000006471.18,protein_coding | 1521.430331 | -0.271602738 | 2.60E-05 | 0.000363632 |
| ENSMUST00000105470.7,Man1a-002,protein_coding,ENSMUSG00000003746.15,protein_coding | 1340.977643 | 0.304727553 | 2.61E-05 | 0.000364456 |
| ENSMUST00000058488.8,Tmed3-001,protein_coding,ENSMUSG00000032353.9,protein_coding | 3257.731366 | 0.278831818 | 2.61E-05 | 0.000365076 |
| ENSMUST00000146692.7,Tnfrsf22-001,protein_coding,ENSMUSG00000010751.15,protein_coding | 367.4184886 | 0.405372035 | 2.62E-05 | 0.000365166 |
| ENSMUST00000113757.7,Odf2-004,protein_coding,ENSMUSG00000026790.19,protein_coding | 2650.719024 | 0.260104586 | 2.62E-05 | 0.000366102 |
| ENSMUST00000031795.7,Fkbp9-001,protein_coding,ENSMUSG00000029781.7,protein_coding | 26911.5711 | 0.169492527 | 2.63E-05 | 0.000366997 |
| ENSMUST00000148128.1,Lmo4-005,retained_intron,ENSMUSG00000028266.17,protein_coding | 103.1773623 | -0.65788921 | 2.65E-05 | 0.000370327 |
| ENSMUST00000155047.5,Fkbp14-004,nonsense_mediated_decay,ENSMUSG00000038074.16,protein_coding | 42.43569459 | 0.781988323 | 2.66E-05 | 0.00037034 |
| ENSMUST00000130201.7,Mrpl15-002,protein_coding,ENSMUSG00000033845.13,protein_coding | 1627.425432 | 0.30654638 | 2.66E-05 | 0.000371484 |
| ENSMUST00000125383.7,Ccdc47-005,protein_coding,ENSMUSG00000078622.8,protein_coding | 283.8944918 | -0.493044861 | 2.67E-05 | 0.000372086 |
| ENSMUST00000045617.14,Hpse-001,protein_coding,ENSMUSG00000035273.14,protein_coding | 2264.508632 | 0.323454412 | 2.68E-05 | 0.000372669 |
| ENSMUST00000033086.7,Phkg2-001,protein_coding,ENSMUSG00000030815.11,protein_coding | 1080.365313 | 0.250509305 | 2.68E-05 | 0.000372866 |
| ENSMUST00000165876.7,Map4-002,protein_coding,ENSMUSG00000032479.15,protein_coding | 22371.03515 | 0.228685924 | 2.68E-05 | 0.000373001 |
| ENSMUST00000216863.1,Actn4-012,protein_coding,ENSMUSG00000054808.15,protein_coding | 264.0095966 | 0.727608383 | 2.70E-05 | 0.000375215 |
| ENSMUST00000112501.8,Tank-202,protein_coding,ENSMUSG00000064289.15,protein_coding | 506.659555 | -0.403267578 | 2.71E-05 | 0.000376404 |
| ENSMUST00000070878.8,Fkbp2-201,protein_coding,ENSMUSG00000056629.15,protein_coding | 1094.877991 | -0.346981837 | 2.71E-05 | 0.0003771 |
| ENSMUST00000207736.1,Bub3-006,nonsense_mediated_decay,ENSMUSG00000066979.6,protein_coding | 453.8349556 | 0.414760482 | 2.71E-05 | 0.000377471 |
| ENSMUST00000040599.14,Nav1-201,protein_coding,ENSMUSG00000009418.15,protein_coding | 2359.615474 | -0.371340822 | 2.73E-05 | 0.000379691 |
| ENSMUST00000092996.4,Sco1-001,protein_coding,ENSMUSG00000069844.12,protein_coding | 2954.416635 | 0.205552652 | 2.73E-05 | 0.00037996 |
| ENSMUST00000037633.14,Zc3h7a-001,protein_coding,ENSMUSG00000037965.14,protein_coding | 2904.329338 | -0.189170935 | 2.74E-05 | 0.000380355 |
| ENSMUST00000030795.9,Abcf2-001,protein_coding,ENSMUSG00000028953.10,protein_coding | 37994.73306 | 0.169696325 | 2.74E-05 | 0.000380505 |
| ENSMUST00000035481.4,Chchd5-001,protein_coding,ENSMUSG00000037938.4,protein_coding | 2014.856691 | 0.27897323 | 2.75E-05 | 0.000381204 |
| ENSMUST00000042724.7,Usp38-001,protein_coding,ENSMUSG00000038250.9,protein_coding | 3384.936106 | -0.191789304 | 2.75E-05 | 0.000381411 |
| ENSMUST00000031838.8,Igf2bp3-001,protein_coding,ENSMUSG00000029814.10,protein_coding | 414.510306 | -0.330102855 | 2.76E-05 | 0.000382997 |
| ENSMUST00000081381.5,Mbtps1-002,protein_coding,ENSMUSG00000031835.16,protein_coding | 23498.34961 | 0.139625317 | 2.76E-05 | 0.000383445 |
| ENSMUST00000146384.7,Eri3-006,processed_transcript,ENSMUSG00000033423.16,protein_coding | 184.4070977 | 0.608256464 | 2.78E-05 | 0.000384808 |
| ENSMUST00000114726.7,Rbmx-002,protein_coding,ENSMUSG00000031134.16,protein_coding | 4517.122366 | 0.347722264 | 2.78E-05 | 0.000385234 |
| ENSMUST00000140824.1,Thsd4-004,protein_coding,ENSMUSG00000032289.15,protein_coding | 29.22139671 | 0.774256117 | 2.79E-05 | 0.000387101 |
| ENSMUST00000100794.9,Myo18a-004,protein_coding,ENSMUSG00000000631.20,protein_coding | 2632.951656 | -0.35246871 | 2.80E-05 | 0.000387743 |
| ENSMUST00000015816.3,Mrpl32-201,protein_coding,ENSMUSG00000015672.3,protein_coding | 3536.639159 | 0.207506765 | 2.81E-05 | 0.000388797 |
| ENSMUST00000109011.7,Gm6710-001,protein_coding,ENSMUSG00000078887.10,protein_coding | 270.9198446 | -0.636627932 | 2.82E-05 | 0.000389952 |
| ENSMUST00000047419.7,Tspo-001,protein_coding,ENSMUSG00000041736.7,protein_coding | 15879.06487 | 0.230013551 | 2.82E-05 | 0.000389952 |
| ENSMUST00000035612.5,Ccar2-201,protein_coding,ENSMUSG00000033712.5,protein_coding | 10054.46468 | 0.18006624 | 2.83E-05 | 0.000391641 |
| ENSMUST00000166682.8,Atxn2l-003,protein_coding,ENSMUSG00000032637.15,protein_coding | 622.088179 | 0.473276941 | 2.84E-05 | 0.000392832 |
| ENSMUST00000134575.7,2410004B18Rik-002,nonsense_mediated_decay,ENSMUSG00000036873.13,protein_coding | 492.5166666 | 0.452244999 | 2.84E-05 | 0.000393129 |
| ENSMUST00000133338.7,Atp11a-004,nonsense_mediated_decay,ENSMUSG00000031441.15,protein_coding | 1370.152178 | 0.373008524 | 2.85E-05 | 0.000393556 |
| ENSMUST00000102696.4,Rps8-001,protein_coding,ENSMUSG00000047675.15,protein_coding | 173637.9052 | 0.186384755 | 2.85E-05 | 0.000393571 |
| ENSMUST00000132419.1,Kdm2b-017,retained_intron,ENSMUSG00000029475.17,protein_coding | 71.44870588 | -0.752933959 | 2.85E-05 | 0.000393571 |
| ENSMUST00000076921.6,Arl16-001,protein_coding,ENSMUSG00000057594.12,protein_coding | 441.2010378 | 0.418392103 | 2.85E-05 | 0.000393571 |
| ENSMUST00000036195.12,Slc35d1-002,protein_coding,ENSMUSG00000028521.17,protein_coding | 1879.869325 | 0.30714863 | 2.87E-05 | 0.000395746 |
| ENSMUST00000004054.12,Kpna1-001,protein_coding,ENSMUSG00000022905.12,protein_coding | 7048.694418 | -0.226197091 | 2.87E-05 | 0.000395806 |
| ENSMUST00000169905.1,Mtdh-006,protein_coding,ENSMUSG00000022255.15,protein_coding | 192.2413929 | -0.613939107 | 2.88E-05 | 0.000396668 |
| ENSMUST00000031879.4,Tcaf2-001,protein_coding,ENSMUSG00000029851.5,protein_coding | 1238.633314 | -0.243030742 | 2.88E-05 | 0.000397668 |
| ENSMUST00000131556.1,Arfgef1-001,protein_coding,ENSMUSG00000067851.11,protein_coding | 1185.523301 | 0.659257212 | 2.89E-05 | 0.000397835 |
| ENSMUST00000215907.1,Smco4-001,protein_coding,ENSMUSG00000058173.12,protein_coding | 544.9044587 | 0.450429485 | 2.89E-05 | 0.000398901 |
| ENSMUST00000004968.10,Plod3-001,protein_coding,ENSMUSG00000004846.10,protein_coding | 11018.41016 | 0.163583902 | 2.92E-05 | 0.000401606 |
| ENSMUST00000133992.7,Yars-003,nonsense_mediated_decay,ENSMUSG00000028811.11,protein_coding | 40.44878841 | 0.770456473 | 2.94E-05 | 0.000405436 |
| ENSMUST00000106344.7,Tnrc6c-001,protein_coding,ENSMUSG00000025571.13,protein_coding | 3955.178867 | 0.227432001 | 3.01E-05 | 0.000414623 |

| | | | | |
|---|---|---|---|---|
| ENSMUST00000051139.12,Rsbn1-002,protein_coding,ENSMUSG00000044098.14,protein_coding | 1573.191918 | 0.379429399 | 3.02E-05 | 0.000416109 |
| ENSMUST00000095267.5,Jund-001,protein_coding,ENSMUSG00000071076.6,protein_coding | 13814.51079 | -0.145028177 | 3.03E-05 | 0.000416221 |
| ENSMUST00000153611.1,Ddx3x-003,retained_intron,ENSMUSG00000000787.12,protein_coding | 137.1953726 | -0.772463664 | 3.03E-05 | 0.000416992 |
| ENSMUST00000209426.1,Ruvbl2-004,protein_coding,ENSMUSG00000003868.14,protein_coding | 300.0400871 | 0.511355588 | 3.03E-05 | 0.000417222 |
| ENSMUST00000048162.9,Bsdc1-001,protein_coding,ENSMUSG00000040859.14,protein_coding | 3471.661702 | 0.225930606 | 3.05E-05 | 0.000418967 |
| ENSMUST00000040298.4,Creg1-003,protein_coding,ENSMUSG00000040713.12,protein_coding | 514.5669252 | 0.389321779 | 3.05E-05 | 0.000419224 |
| ENSMUST00000023486.14,Tfrc-001,protein_coding,ENSMUSG00000022797.15,protein_coding | 11951.59196 | -0.56397397 | 3.06E-05 | 0.000420037 |
| ENSMUST00000114801.7,Fance-004,protein_coding,ENSMUSG00000007570.15,protein_coding | 210.1394914 | 0.574484775 | 3.06E-05 | 0.000420265 |
| ENSMUST00000115233.1,Ppp1r2-002,protein_coding,ENSMUSG00000047714.14,protein_coding | 2353.961702 | 0.198593164 | 3.06E-05 | 0.000420361 |
| ENSMUST00000147105.7,Myef2-003,protein_coding,ENSMUSG00000027201.16,protein_coding | 3406.289915 | 0.272530002 | 3.07E-05 | 0.000421865 |
| ENSMUST00000170060.2,Zfp839-201,protein_coding,ENSMUSG00000021271.4,protein_coding | 962.0897852 | 0.288634343 | 3.08E-05 | 0.00042315 |
| ENSMUST00000018851.13,Dync1h1-001,protein_coding,ENSMUSG00000018707.13,protein_coding | 101425.1242 | 0.178818586 | 3.12E-05 | 0.000427716 |
| ENSMUST00000127793.1,Uba52-007,retained_intron,ENSMUSG00000090137.8,protein_coding | 257.9219897 | 0.437144437 | 3.12E-05 | 0.000428381 |
| ENSMUST00000027339.13,Smap1-001,protein_coding,ENSMUSG00000026155.13,protein_coding | 7069.740934 | 0.177176583 | 3.14E-05 | 0.000430409 |
| ENSMUST00000185184.1,Tcf24-001,protein_coding,ENSMUSG00000099032.2,protein_coding | 644.2347615 | -0.28408143 | 3.17E-05 | 0.00043339 |
| ENSMUST00000078459.6,Memo1-201,protein_coding,ENSMUSG00000058704.7,protein_coding | 8383.371523 | 0.184740696 | 3.16E-05 | 0.00043339 |
| ENSMUST00000150267.1,Napg-007,protein_coding,ENSMUSG00000024581.12,protein_coding | 135.4163415 | 0.568722605 | 3.16E-05 | 0.00043339 |
| ENSMUST00000002073.12,Ltbp2-001,protein_coding,ENSMUSG00000002020.15,protein_coding | 238.9633982 | -0.705902081 | 3.17E-05 | 0.000433654 |
| ENSMUST00000121791.7,Tm9sf1-008,protein_coding,ENSMUSG00000002320.15,protein_coding | 687.1578809 | 0.773244333 | 3.17E-05 | 0.000433696 |
| ENSMUST00000029694.13,Arhgef2-001,protein_coding,ENSMUSG00000028059.15,protein_coding | 11631.83662 | -0.34896326 | 3.18E-05 | 0.000434606 |
| ENSMUST00000162246.8,Hmgn3-001,protein_coding,ENSMUSG00000066456.14,protein_coding | 579.9463988 | -0.402910063 | 3.18E-05 | 0.000434801 |
| ENSMUST00000149610.2,Tpi1-004,protein_coding,ENSMUSG00000023456.14,protein_coding | 4495.119603 | 0.408028134 | 3.19E-05 | 0.000436066 |
| ENSMUST00000130500.7,Bnip3-004,protein_coding,ENSMUSG00000078566.8,protein_coding | 1000.433403 | 0.477013692 | 3.19E-05 | 0.000436151 |
| ENSMUST00000023024.7,Tef-006,protein_coding,ENSMUSG00000022389.14,protein_coding | 4302.782292 | 0.193728223 | 3.20E-05 | 0.000436599 |
| ENSMUST00000109384.9,Tomm34-001,protein_coding,ENSMUSG00000018322.10,protein_coding | 7017.688179 | -0.198474371 | 3.20E-05 | 0.00043665 |
| ENSMUST00000029610.8,Hadh-001,protein_coding,ENSMUSG00000027984.8,protein_coding | 10548.51074 | 0.20490031 | 3.20E-05 | 0.000436675 |
| ENSMUST00000163754.7,Xpnpep3-001,protein_coding,ENSMUSG00000022401.11,protein_coding | 6314.314005 | 0.167012416 | 3.20E-05 | 0.000436675 |
| ENSMUST00000025759.7,Eif1ad-201,protein_coding,ENSMUSG00000024841.7,protein_coding | 12581.75596 | 0.202476659 | 3.20E-05 | 0.000436675 |
| ENSMUST00000159327.1,Myc-004,protein_coding,ENSMUSG00000022346.15,protein_coding | 1390.438027 | 0.743228884 | 3.22E-05 | 0.000439098 |
| ENSMUST00000029085.8,Mrgbp-001,protein_coding,ENSMUSG00000027569.15,protein_coding | 1593.492118 | -0.223125768 | 3.22E-05 | 0.000439582 |
| ENSMUST00000115251.7,Tnpo3-201,protein_coding,ENSMUSG00000012535.14,protein_coding | 1373.607411 | 0.603759655 | 3.23E-05 | 0.000440242 |
| ENSMUST00000009039.5,Rpl30-002,protein_coding,ENSMUSG00000058600.13,protein_coding | 35941.29642 | 0.128987943 | 3.24E-05 | 0.000440829 |
| ENSMUST00000026661.3,Tk1-001,protein_coding,ENSMUSG00000025574.13,protein_coding | 12659.60504 | -0.240361715 | 3.25E-05 | 0.000442309 |
| ENSMUST00000077693.9,Eif2b4-001,protein_coding,ENSMUSG00000029145.12,protein_coding | 1699.022717 | -0.273855548 | 3.25E-05 | 0.000442736 |
| ENSMUST00000197993.1,Anapc7-006,retained_intron,ENSMUSG00000029466.11,protein_coding | 66.73652679 | 0.714161867 | 3.25E-05 | 0.000442736 |
| ENSMUST00000038422.6,Irf2bpl-201,protein_coding,ENSMUSG00000034168.6,protein_coding | 7201.839743 | 0.117677391 | 3.25E-05 | 0.000442908 |
| ENSMUST00000082424.10,Selenop-201,protein_coding,ENSMUSG00000064373.11,protein_coding | 1701.117969 | -0.316831819 | 3.27E-05 | 0.000444249 |
| ENSMUST00000150372.1,Psmb2-003,processed_transcript,ENSMUSG00000028837.8,protein_coding | 73.01630575 | 0.710610906 | 3.29E-05 | 0.000446921 |
| ENSMUST00000024905.9,Ralbp1-201,protein_coding,ENSMUSG00000024096.16,protein_coding | 9961.517927 | 0.156284859 | 3.36E-05 | 0.000456439 |
| ENSMUST00000091299.6,Cdk7-201,protein_coding,ENSMUSG00000069089.7,protein_coding | 5037.809168 | -0.16427737 | 3.36E-05 | 0.000456607 |
| ENSMUST00000069792.13,Nab1-001,protein_coding,ENSMUSG00000002881.14,protein_coding | 6289.532832 | -0.393184308 | 3.37E-05 | 0.000457547 |
| ENSMUST00000176797.7,Hnrnpk-021,protein_coding,ENSMUSG00000021546.17,protein_coding | 2078.021321 | 0.526430573 | 3.37E-05 | 0.000458068 |
| ENSMUST00000119746.7,Usf3-001,protein_coding,ENSMUSG00000068284.14,protein_coding | 5929.141946 | -0.301824073 | 3.38E-05 | 0.000459005 |
| ENSMUST00000211981.1,Wdr59-003,protein_coding,ENSMUSG00000031959.15,protein_coding | 1205.973016 | 0.484847098 | 3.39E-05 | 0.000460713 |
| ENSMUST00000152037.2,Uba2-007,protein_coding,ENSMUSG00000052997.15,protein_coding | 130.412339 | 0.577741238 | 3.40E-05 | 0.000461402 |
| ENSMUST00000047425.4,Sntb2-002,protein_coding,ENSMUSG00000041308.6,protein_coding | 2416.196681 | 0.261142995 | 3.40E-05 | 0.00046166 |
| ENSMUST00000123141.1,Mex3d-002,protein_coding,ENSMUSG00000048696.10,protein_coding | 677.1486743 | 0.488367406 | 3.41E-05 | 0.000461986 |
| ENSMUST00000035295.5,Degs1-001,protein_coding,ENSMUSG00000038633.5,protein_coding | 15655.04729 | 0.176184076 | 3.42E-05 | 0.000463669 |
| ENSMUST00000181986.7,Rbm6-003,nonsense_mediated_decay,ENSMUSG00000032582.14,protein_coding | 468.6605572 | 0.562074313 | 3.43E-05 | 0.000464395 |
| ENSMUST00000022272.12,Kctd6-201,protein_coding,ENSMUSG00000021752.12,protein_coding | 986.5895994 | 0.386445913 | 3.45E-05 | 0.000467608 |
| ENSMUST00000177051.7,Hnrnpk-018,retained_intron,ENSMUSG00000021546.17,protein_coding | 2991.427117 | 0.372481788 | 3.45E-05 | 0.000467654 |
| ENSMUST00000029643.14,Gar1-001,protein_coding,ENSMUSG00000028010.16,protein_coding | 4710.330116 | 0.247665695 | 3.46E-05 | 0.000467867 |
| ENSMUST00000133885.1,Hmgn1-004,protein_coding,ENSMUSG00000040681.14,protein_coding | 510.5744618 | 0.693725354 | 3.46E-05 | 0.000468265 |
| ENSMUST00000033570.5,Igbp1-001,protein_coding,ENSMUSG00000031221.7,protein_coding | 5973.534447 | 0.148523082 | 3.46E-05 | 0.00046853 |
| ENSMUST00000042196.3,Vwa1-001,protein_coding,ENSMUSG00000042116.3,protein_coding | 246.0376198 | -0.50885936 | 3.48E-05 | 0.000470559 |
| ENSMUST00000145001.1,Polr3gl-002,protein_coding,ENSMUSG00000028104.14,protein_coding | 127.8309051 | 0.612722125 | 3.48E-05 | 0.000471156 |
| ENSMUST00000150063.8,Gnb2-004,protein_coding,ENSMUSG00000029713.15,protein_coding | 665.0555729 | -0.411215102 | 3.49E-05 | 0.000472257 |
| ENSMUST00000210853.1,Cyth2-003,protein_coding,ENSMUSG00000003269.16,protein_coding | 2760.689454 | 0.196370936 | 3.51E-05 | 0.000474191 |
| ENSMUST00000039990.5,P3h2-001,protein_coding,ENSMUSG00000038168.5,protein_coding | 1288.116689 | 0.301575624 | 3.51E-05 | 0.000474234 |
| ENSMUST00000079625.10,Tor3a-001,protein_coding,ENSMUSG00000060519.11,protein_coding | 938.6370855 | 0.41509617 | 3.52E-05 | 0.000475041 |
| ENSMUST00000182877.7,Xpnpep1-002,processed_transcript,ENSMUSG00000025027.17,protein_coding | 15708.74328 | 0.192903252 | 3.52E-05 | 0.000475041 |
| ENSMUST00000150285.7,Slc35d1-001,protein_coding,ENSMUSG00000028521.17,protein_coding | 1671.739205 | 0.265993539 | 3.56E-05 | 0.000480441 |
| ENSMUST00000188593.1,Tuba4a-003,protein_coding,ENSMUSG00000026202.13,protein_coding | 2714.056752 | -0.302119108 | 3.56E-05 | 0.000480814 |
| ENSMUST00000081202.5,Manbal-001,protein_coding,ENSMUSG00000063019.5,protein_coding | 4458.503289 | 0.196641445 | 3.57E-05 | 0.000481696 |
| ENSMUST00000014640.7,Ankrd28-201,protein_coding,ENSMUSG00000014496.7,protein_coding | 9735.058214 | -0.196491762 | 3.57E-05 | 0.000481696 |
| ENSMUST00000113904.8,Zcchc4-001,protein_coding,ENSMUSG00000029179.14,protein_coding | 2098.363592 | 0.162809657 | 3.59E-05 | 0.000484153 |
| ENSMUST00000068505.9,Capn2-001,protein_coding,ENSMUSG00000026509.15,protein_coding | 55794.68326 | 0.176652998 | 3.59E-05 | 0.00048436 |
| ENSMUST00000045078.12,Grhpr-001,protein_coding,ENSMUSG00000035637.14,protein_coding | 4721.032259 | 0.232429642 | 3.60E-05 | 0.000485134 |
| ENSMUST00000058245.4,Fgfr1op2-004,protein_coding,ENSMUSG00000040242.14,protein_coding | 2564.630272 | 0.170510981 | 3.62E-05 | 0.000487772 |
| ENSMUST00000089414.10,Kctd17-001,protein_coding,ENSMUSG00000033287.15,protein_coding | 3542.871325 | -0.181036486 | 3.63E-05 | 0.000488705 |
| ENSMUST00000115668.9,Otud5-002,protein_coding,ENSMUSG00000031154.15,protein_coding | 3309.120322 | 0.195796881 | 3.64E-05 | 0.000489604 |
| ENSMUST00000047951.8,Fbxo2-001,protein_coding,ENSMUSG00000041556.8,protein_coding | 637.6159237 | -0.431885739 | 3.64E-05 | 0.00049 |
| ENSMUST00000198415.1,Fbrsl1-007,processed_transcript,ENSMUSG00000043323.16,protein_coding | 1236.950726 | -0.207795874 | 3.66E-05 | 0.000491774 |
| ENSMUST00000171702.7,Racgap1-001,protein_coding,ENSMUSG00000023015.13,protein_coding | 9609.635436 | 0.239413771 | 3.66E-05 | 0.000492238 |
| ENSMUST00000093883.3,Arl14ep-002,protein_coding,ENSMUSG00000027122.15,protein_coding | 1324.625498 | 0.400126945 | 3.66E-05 | 0.000492244 |
| ENSMUST00000152009.1,Itga6-003,processed_transcript,ENSMUSG00000027111.15,protein_coding | 273.3503625 | 0.493714945 | 3.69E-05 | 0.000495554 |
| ENSMUST00000002881.3,Nudt14-201,protein_coding,ENSMUSG00000002804.3,protein_coding | 3196.335613 | 0.225069781 | 3.69E-05 | 0.00049564 |
| ENSMUST00000063562.7,Mipep-201,protein_coding,ENSMUSG00000021993.9,protein_coding | 3611.415802 | 0.198895408 | 3.69E-05 | 0.000496002 |
| ENSMUST00000120281.7,Mmp11-002,protein_coding,ENSMUSG00000000901.15,protein_coding | 788.8200049 | 0.511597662 | 3.70E-05 | 0.00049694 |
| ENSMUST00000160430.2,Luc7l2-012,retained_intron,ENSMUSG00000029823.16,protein_coding | 1273.039628 | -0.476647854 | 3.71E-05 | 0.000497619 |
| ENSMUST00000110339.4,Gm20498-003,processed_transcript,ENSMUSG00000021139.17,protein_coding | 331.9415143 | 0.411003724 | 3.71E-05 | 0.000498175 |
| ENSMUST00000129995.7,Etv4-008,processed_transcript,ENSMUSG00000017724.14,protein_coding | 1222.411407 | 0.248163037 | 3.72E-05 | 0.000498401 |
| ENSMUST00000183271.1,Dennd2d-001,protein_coding,ENSMUSG00000027901.12,protein_coding | 174.7994368 | 0.66218578 | 3.73E-05 | 0.000499514 |
| ENSMUST00000043552.16,Serpinb6a-001,protein_coding,ENSMUSG00000060147.15,protein_coding | 17172.1062 | 0.224081617 | 3.73E-05 | 0.000499514 |
| ENSMUST00000168621.2,Ptprj-201,protein_coding,ENSMUSG00000025314.16,protein_coding | 2776.734199 | 0.162395345 | 3.75E-05 | 0.000503013 |
| ENSMUST00000045281.12,Smg6-001,protein_coding,ENSMUSG00000038290.15,protein_coding | 5825.61038 | 0.171316626 | 3.77E-05 | 0.000505561 |
| ENSMUST00000184864.7,Erc1-009,protein_coding,ENSMUSG00000030172.15,protein_coding | 1606.663366 | -0.382692112 | 3.78E-05 | 0.000506183 |
| ENSMUST00000102752.9,Homer1-002,protein_coding,ENSMUSG00000007617.17,protein_coding | 290.4396784 | -0.632822006 | 3.78E-05 | 0.00050624 |
| ENSMUST00000020524.14,Rhbdf1-006,protein_coding,ENSMUSG00000020282.18,protein_coding | 4009.77165 | 0.259543707 | 3.79E-05 | 0.00050762 |
| ENSMUST00000176014.7,Asap1-008,protein_coding,ENSMUSG00000022377.16,protein_coding | 2286.409838 | 0.368723414 | 3.80E-05 | 0.000509084 |
| ENSMUST00000168521.1,E130317F20Rik-001,processed_transcript,ENSMUSG00000091994.1,processed_transcript | 406.2232084 | -0.363514105 | 3.81E-05 | 0.000509622 |
| ENSMUST00000150174.7,Stard5-005,retained_intron,ENSMUSG00000046027.17,protein_coding | 103.9805166 | 0.753772309 | 3.82E-05 | 0.000510975 |

| | | | | |
|---|---|---|---|---|
| ENSMUST00000048923.6,Spred3-001,protein_coding,ENSMUSG00000037239.8,protein_coding | 1518.337569 | -0.223108577 | 3.82E-05 | 0.000511076 |
| ENSMUST00000105870.7,Pafah2-001,protein_coding,ENSMUSG00000037366.14,protein_coding | 594.616473 | -0.32181768 | 3.86E-05 | 0.000515463 |
| ENSMUST00000069486.12,Gemin6-001,protein_coding,ENSMUSG00000055760.12,protein_coding | 2121.275159 | -0.270880359 | 3.86E-05 | 0.000515869 |
| ENSMUST00000153661.1,Tatdn2-003,protein_coding,ENSMUSG00000056952.13,protein_coding | 71.93296184 | 0.761917946 | 3.87E-05 | 0.0005169 |
| ENSMUST00000087031.5,Xpo5-201,protein_coding,ENSMUSG00000067150.5,protein_coding | 26550.97921 | 0.170271059 | 3.87E-05 | 0.000517353 |
| ENSMUST00000028408.2,Hat1-201,protein_coding,ENSMUSG00000027018.11,protein_coding | 11617.10638 | -0.185665861 | 3.89E-05 | 0.000519079 |
| ENSMUST00000114372.2,Hcfc1-005,protein_coding,ENSMUSG00000031386.14,protein_coding | 17527.56679 | -0.208431079 | 3.91E-05 | 0.000521246 |
| ENSMUST00000110408.1,Ropn1l-201,protein_coding,ENSMUSG00000022236.9,protein_coding | 241.8461027 | 0.480058593 | 3.93E-05 | 0.000523935 |
| ENSMUST00000060253.3,Fem1a-201,protein_coding,ENSMUSG00000043683.3,protein_coding | 6299.70859 | -0.166759201 | 3.93E-05 | 0.000524365 |
| ENSMUST00000162945.1,Spata13-002,protein_coding,ENSMUSG00000021990.15,protein_coding | 769.9883523 | -0.251618886 | 3.93E-05 | 0.000524468 |
| ENSMUST00000032174.11,Klf15-001,protein_coding,ENSMUSG00000030087.11,protein_coding | 149.1993044 | 0.751224344 | 3.95E-05 | 0.000526604 |
| ENSMUST00000096430.10,Zhx2-001,protein_coding,ENSMUSG00000071757.10,protein_coding | 1757.288397 | 0.328163164 | 3.98E-05 | 0.000529852 |
| ENSMUST00000040787.12,Ankrd13c-001,protein_coding,ENSMUSG00000039988.14,protein_coding | 3594.18219 | -0.284411891 | 3.98E-05 | 0.000530102 |
| ENSMUST00000032264.8,Gabarapl1-001,protein_coding,ENSMUSG00000030161.8,protein_coding | 13147.74836 | 0.175396038 | 3.98E-05 | 0.000530277 |
| ENSMUST00000039522.7,Apobr-001,protein_coding,ENSMUSG00000042759.12,protein_coding | 915.4570215 | -0.267470018 | 3.99E-05 | 0.000531189 |
| ENSMUST00000047711.12,Vezt-002,protein_coding,ENSMUSG00000036099.16,protein_coding | 1402.345942 | 0.238793531 | 4.00E-05 | 0.000532111 |
| ENSMUST00000173198.7,Magohb-003,nonsense_mediated_decay,ENSMUSG00000030188.12,protein_coding | 100.9923744 | 0.632668961 | 4.01E-05 | 0.000533363 |
| ENSMUST00000135927.1,Ddb2-005,processed_transcript,ENSMUSG00000002109.14,protein_coding | 263.9599817 | 0.652966781 | 4.01E-05 | 0.000533788 |
| ENSMUST00000153513.7,Setd5-011,processed_transcript,ENSMUSG00000034269.12,protein_coding | 2263.822009 | 0.265436051 | 4.02E-05 | 0.000534749 |
| ENSMUST00000025417.8,Snx24-201,protein_coding,ENSMUSG00000024535.15,protein_coding | 92.03147145 | 0.69852366 | 4.04E-05 | 0.000536806 |
| ENSMUST00000106726.9,0610037L13Rik-013,protein_coding,ENSMUSG00000028608.15,protein_coding | 271.583902 | -0.52133307 | 4.05E-05 | 0.00053835 |
| ENSMUST00000149808.1,Ucp2-010,processed_transcript,ENSMUSG00000033685.13,protein_coding | 47.25274283 | 0.712494231 | 4.06E-05 | 0.00053867 |
| ENSMUST00000018361.9,Pmp22-001,protein_coding,ENSMUSG00000018217.12,protein_coding | 1531.238655 | 0.297828577 | 4.06E-05 | 0.000538697 |
| ENSMUST00000072008.10,Sh3rf2-001,protein_coding,ENSMUSG00000057719.10,protein_coding | 786.2211694 | -0.451060147 | 4.06E-05 | 0.000538698 |
| ENSMUST00000135877.7,Eef1b2-004,nonsense_mediated_decay,ENSMUSG00000025967.16,protein_coding | 30599.32029 | 0.217610408 | 4.07E-05 | 0.000540213 |
| ENSMUST00000177978.1,1700047I17Rik2-201,protein_coding,ENSMUSG00000094103.1,protein_coding | 3702.578312 | 0.236791449 | 4.08E-05 | 0.000541582 |
| ENSMUST00000041314.14,Paip2-001,protein_coding,ENSMUSG00000037058.14,protein_coding | 8231.649753 | 0.168579739 | 4.10E-05 | 0.000543358 |
| ENSMUST00000040990.5,Vwa8-201,protein_coding,ENSMUSG00000058997.7,protein_coding | 5423.79387 | 0.158893558 | 4.10E-05 | 0.000543403 |
| ENSMUST00000119606.7,Zbtb7a-002,protein_coding,ENSMUSG00000035011.15,protein_coding | 537.516574 | -0.483713225 | 4.10E-05 | 0.000543761 |
| ENSMUST00000151796.2,Fkbp4-004,protein_coding,ENSMUSG00000030357.10,protein_coding | 4165.77323 | 0.292874296 | 4.10E-05 | 0.000543828 |
| ENSMUST00000088896.9,Tmcc1-001,protein_coding,ENSMUSG00000030126.17,protein_coding | 1398.57717 | -0.324093644 | 4.11E-05 | 0.000544939 |
| ENSMUST00000108210.8,Ttc14-008,protein_coding,ENSMUSG00000027677.17,protein_coding | 628.4582755 | -0.585870083 | 4.12E-05 | 0.00054578 |
| ENSMUST00000139210.7,Socs2-002,nonsense_mediated_decay,ENSMUSG00000020027.18,protein_coding | 172.1730737 | 0.600035711 | 4.12E-05 | 0.00054578 |
| ENSMUST00000136252.7,Parl-002,nonsense_mediated_decay,ENSMUSG00000033918.15,protein_coding | 223.3853182 | 0.704103028 | 4.12E-05 | 0.00054578 |
| ENSMUST00000041565.10,Ift172-001,protein_coding,ENSMUSG00000038564.10,protein_coding | 7095.354176 | 0.237950617 | 4.13E-05 | 0.000545836 |
| ENSMUST00000117901.7,Zmym3-002,protein_coding,ENSMUSG00000031310.16,protein_coding | 540.0554573 | -0.36752903 | 4.13E-05 | 0.000545836 |
| ENSMUST00000150387.7,Fkbp4-003,protein_coding,ENSMUSG00000030357.10,protein_coding | 91.30969865 | 0.765313803 | 4.13E-05 | 0.000546223 |
| ENSMUST00000123013.1,Nol12-003,protein_coding,ENSMUSG00000033099.9,protein_coding | 415.0945602 | 0.42325169 | 4.16E-05 | 0.000550251 |
| ENSMUST00000190476.1,Ing5-003,protein_coding,ENSMUSG00000026283.13,protein_coding | 128.4289961 | 0.687615548 | 4.16E-05 | 0.000550286 |
| ENSMUST00000021078.2,Fdxr-001,protein_coding,ENSMUSG00000018861.8,protein_coding | 2123.655264 | 0.259860064 | 4.17E-05 | 0.000551258 |
| ENSMUST00000069453.8,Paqr3-001,protein_coding,ENSMUSG00000055725.11,protein_coding | 541.3182611 | 0.417439268 | 4.19E-05 | 0.000552673 |
| ENSMUST00000171165.7,Ube2f-002,protein_coding,ENSMUSG00000034343.14,protein_coding | 859.4412476 | -0.287989789 | 4.19E-05 | 0.000553082 |
| ENSMUST00000134907.7,Anxa2-004,protein_coding,ENSMUSG00000032231.14,protein_coding | 457.7672386 | 0.591311671 | 4.19E-05 | 0.000553202 |
| ENSMUST00000031617.12,Rpl6-001,protein_coding,ENSMUSG00000029614.13,protein_coding | 186419.1003 | 0.173034779 | 4.20E-05 | 0.000553776 |
| ENSMUST00000199134.1,Cxxc4-003,retained_intron,ENSMUSG00000044365.15,protein_coding | 572.3828988 | -0.292071799 | 4.21E-05 | 0.000555738 |
| ENSMUST00000015239.9,Fbxw5-001,protein_coding,ENSMUSG00000015095.12,protein_coding | 5957.561246 | -0.150862935 | 4.23E-05 | 0.000557172 |
| ENSMUST00000040907.6,Decr2-001,protein_coding,ENSMUSG00000036775.12,protein_coding | 2605.627244 | 0.202628401 | 4.23E-05 | 0.000557582 |
| ENSMUST00000171569.7,Ndrg1-002,retained_intron,ENSMUSG00000005125.12,protein_coding | 102.53721 | 0.751223617 | 4.25E-05 | 0.00056025 |
| ENSMUST00000177035.7,Asap1-007,protein_coding,ENSMUSG00000022377.16,protein_coding | 1233.546916 | -0.564114752 | 4.30E-05 | 0.00056578 |
| ENSMUST00000034015.10,Msmo1-001,protein_coding,ENSMUSG00000031604.10,protein_coding | 18033.12136 | 0.257420245 | 4.30E-05 | 0.000566177 |
| ENSMUST00000055131.12,Usp48-001,protein_coding,ENSMUSG00000043411.15,protein_coding | 3737.981299 | -0.262834461 | 4.32E-05 | 0.000568747 |
| ENSMUST00000187938.6,Fn1-007,protein_coding,ENSMUSG00000026193.15,protein_coding | 11699.1539 | 0.722072327 | 4.32E-05 | 0.000568951 |
| ENSMUST00000147223.7,Entpd6-003,retained_intron,ENSMUSG00000033068.16,protein_coding | 166.5314652 | 0.5311261 | 4.33E-05 | 0.000568968 |
| ENSMUST00000177513.1,Rpl41-006,retained_intron,ENSMUSG00000093674.7,protein_coding | 2336.275926 | -0.262054578 | 4.33E-05 | 0.000568968 |
| ENSMUST00000120531.7,Stxbp6-002,protein_coding,ENSMUSG00000046314.15,protein_coding | 431.6254673 | -0.495203844 | 4.33E-05 | 0.000569259 |
| ENSMUST00000043429.11,Nfyc-001,protein_coding,ENSMUSG00000032897.17,protein_coding | 5945.306083 | -0.219577613 | 4.34E-05 | 0.000570472 |
| ENSMUST00000109280.1,Pwwp2a-001,protein_coding,ENSMUSG00000044950.7,protein_coding | 69.83128473 | 0.71061905 | 4.35E-05 | 0.000571614 |
| ENSMUST00000020204.4,Ntn4-001,protein_coding,ENSMUSG00000020019.4,protein_coding | 1867.221143 | -0.221916045 | 4.36E-05 | 0.000572231 |
| ENSMUST00000090171.6,Adh7-001,protein_coding,ENSMUSG00000055301.8,protein_coding | 891.43874 | 0.452737518 | 4.39E-05 | 0.000575853 |
| ENSMUST00000211932.1,Tcf25-008,protein_coding,ENSMUSG00000001472.17,protein_coding | 4524.209324 | -0.215807992 | 4.39E-05 | 0.000576228 |
| ENSMUST00000022371.9,Ap3m1-002,protein_coding,ENSMUSG00000021824.12,protein_coding | 2225.287468 | -0.344204178 | 4.39E-05 | 0.000576302 |
| ENSMUST00000182812.7,Sec31a-005,protein_coding,ENSMUSG00000035325.16,protein_coding | 3807.606858 | -0.171851114 | 4.40E-05 | 0.000577116 |
| ENSMUST00000115492.1,Gpr85-002,protein_coding,ENSMUSG00000048216.13,protein_coding | 489.754099 | 0.568113181 | 4.42E-05 | 0.000580114 |
| ENSMUST00000017567.13,Zfp207-002,protein_coding,ENSMUSG00000017421.18,protein_coding | 13207.54217 | -0.218687143 | 4.44E-05 | 0.000582206 |
| ENSMUST00000046470.15,Mettl22-001,protein_coding,ENSMUSG00000039345.15,protein_coding | 1142.021189 | 0.322089276 | 4.44E-05 | 0.000582579 |
| ENSMUST00000079659.11,U2surp-002,protein_coding,ENSMUSG00000032407.14,protein_coding | 1618.195095 | -0.486801555 | 4.49E-05 | 0.000587999 |
| ENSMUST00000003681.7,Sec14l2-006,protein_coding,ENSMUSG00000003585.13,protein_coding | 166.8057259 | -0.563866247 | 4.50E-05 | 0.000589244 |
| ENSMUST00000155949.1,6530402F18Rik-001,processed_transcript,ENSMUSG00000079499.9,processed_transcript | 168.6179519 | -0.536064523 | 4.51E-05 | 0.000589915 |
| ENSMUST00000182715.7,Tmem161a-012,nonsense_mediated_decay,ENSMUSG00000002342.17,protein_coding | 46.36389246 | 0.757464136 | 4.51E-05 | 0.000589915 |
| ENSMUST00000130643.7,Srpk1-001,protein_coding,ENSMUSG00000004865.15,protein_coding | 11712.53071 | 0.142420811 | 4.51E-05 | 0.000590182 |
| ENSMUST00000164884.8,Fam96b-001,protein_coding,ENSMUSG00000031879.14,protein_coding | 4179.928619 | 0.198936823 | 4.52E-05 | 0.000590757 |
| ENSMUST00000212024.1,Wdfy3-008,protein_coding,ENSMUSG00000043940.15,protein_coding | 3711.959997 | -0.242648712 | 4.52E-05 | 0.000591263 |
| ENSMUST00000034927.12,Zic1-001,protein_coding,ENSMUSG00000032368.14,protein_coding | 611.2229034 | -0.344758483 | 4.53E-05 | 0.00059197 |
| ENSMUST00000008094.9,Abhd8-001,protein_coding,ENSMUSG00000007950.9,protein_coding | 6797.578765 | -0.210915936 | 4.54E-05 | 0.000593021 |
| ENSMUST00000057198.8,5730508B09Rik-002,protein_coding,ENSMUSG00000050549.8,protein_coding | 294.1472639 | 0.557688173 | 4.54E-05 | 0.000593847 |
| ENSMUST00000092213.10,Nr2c1-002,protein_coding,ENSMUSG00000005897.14,protein_coding | 691.914565 | 0.450876289 | 4.55E-05 | 0.000594007 |
| ENSMUST00000209212.1,Rpl13a-014,nonsense_mediated_decay,ENSMUSG00000074129.13,protein_coding | 3559.752593 | 0.3918577 | 4.55E-05 | 0.000594851 |
| ENSMUST00000051011.13,Tmem198b-001,protein_coding,ENSMUSG00000047090.13,protein_coding | 1548.10786 | -0.169903142 | 4.56E-05 | 0.000594862 |
| ENSMUST00000111704.7,Rassf8-001,protein_coding,ENSMUSG00000030259.12,protein_coding | 5358.815305 | 0.115762381 | 4.60E-05 | 0.00060038 |
| ENSMUST00000198239.1,Fdps-003,retained_intron,ENSMUSG00000059743.12,protein_coding | 720.6668891 | 0.407487256 | 4.61E-05 | 0.00060207 |
| ENSMUST00000051737.7,Ap1ar-001,protein_coding,ENSMUSG00000074238.6,protein_coding | 7637.877187 | 0.165670133 | 4.62E-05 | 0.000602874 |
| ENSMUST00000034581.3,Cadm1-006,protein_coding,ENSMUSG00000032076.19,protein_coding | 783.8486836 | 0.571703928 | 4.63E-05 | 0.000603834 |
| ENSMUST00000040110.7,Chst11-001,protein_coding,ENSMUSG00000034612.7,protein_coding | 4694.073236 | -0.199721986 | 4.63E-05 | 0.000603834 |
| ENSMUST00000112484.9,Ptpn6-002,protein_coding,ENSMUSG00000004266.15,protein_coding | 1295.236472 | 0.335852434 | 4.64E-05 | 0.000604205 |
| ENSMUST00000127800.1,Ago1-002,nonsense_mediated_decay,ENSMUSG00000041530.16,protein_coding | 153.8406223 | 0.69368903 | 4.65E-05 | 0.000606246 |
| ENSMUST00000114159.8,Nacc2-002,protein_coding,ENSMUSG00000026932.14,protein_coding | 2644.806315 | -0.47812825 | 4.67E-05 | 0.00060771 |
| ENSMUST00000203404.1,Cbx3-008,retained_intron,ENSMUSG00000029836.15,protein_coding | 211.5473432 | -0.502893722 | 4.67E-05 | 0.000607867 |
| ENSMUST00000026254.13,Gbf1-001,protein_coding,ENSMUSG00000025224.14,protein_coding | 12888.25178 | 0.160498371 | 4.67E-05 | 0.000607867 |
| ENSMUST00000074465.8,Aqp9-001,protein_coding,ENSMUSG00000032204.13,protein_coding | 91.12152621 | 0.712302411 | 4.69E-05 | 0.000610006 |
| ENSMUST00000105097.2,Zswim6-201,protein_coding,ENSMUSG00000032846.9,protein_coding | 770.6247902 | 0.263154972 | 4.70E-05 | 0.000611618 |
| ENSMUST00000033335.5,Akip1-001,protein_coding,ENSMUSG00000031023.5,protein_coding | 1382.982505 | 0.227772372 | 4.72E-05 | 0.000613482 |
| ENSMUST00000112177.7,Sfmbt1-004,protein_coding,ENSMUSG00000006527.14,protein_coding | 1055.846377 | 0.41170028 | 4.72E-05 | 0.000613552 |

| | | | | |
|---|---|---|---|---|
| ENSMUST00000028593.10,Prrg4-001,protein_coding,ENSMUSG00000027171.10,protein_coding | 269.6507041 | -0.416669453 | 4.77E-05 | 0.000619309 |
| ENSMUST00000025856.16,Lrp5-001,protein_coding,ENSMUSG00000024913.16,protein_coding | 14186.61923 | 0.141608683 | 4.77E-05 | 0.000619897 |
| ENSMUST00000026292.14,Huwe1-002,protein_coding,ENSMUSG00000025261.17,protein_coding | 60602.8473 | -0.207741649 | 4.80E-05 | 0.000622869 |
| ENSMUST00000205440.1,Sh2b1-008,protein_coding,ENSMUSG00000030733.10,protein_coding | 1717.797262 | 0.278943034 | 4.80E-05 | 0.000623096 |
| ENSMUST00000142256.1,Pdcd6ip-006,retained_intron,ENSMUSG00000032504.15,protein_coding | 33.37833264 | -0.738587181 | 4.81E-05 | 0.000624852 |
| ENSMUST00000044250.3,Alkbh5-001,protein_coding,ENSMUSG00000042650.3,protein_coding | 27435.22462 | 0.138009727 | 4.82E-05 | 0.000625026 |
| ENSMUST00000019614.12,Xab2-001,protein_coding,ENSMUSG00000019470.12,protein_coding | 9293.075027 | 0.192961246 | 4.82E-05 | 0.000625359 |
| ENSMUST00000174391.7,Syncrip-004,protein_coding,ENSMUSG00000032423.12,protein_coding | 4477.126568 | -0.523303354 | 4.84E-05 | 0.000627164 |
| ENSMUST00000071106.4,Gpatch2l-201,protein_coding,ENSMUSG00000021254.8,protein_coding | 3329.90615 | 0.175921532 | 4.88E-05 | 0.000632599 |
| ENSMUST00000165853.1,Ptp4a2-201,protein_coding,ENSMUSG00000028788.14,protein_coding | 2207.297146 | -0.69021616 | 4.89E-05 | 0.000633225 |
| ENSMUST00000204177.2,Mlf2-003,retained_intron,ENSMUSG00000030120.14,protein_coding | 2263.644602 | -0.22292742 | 4.89E-05 | 0.000633326 |
| ENSMUST00000034022.3,Sap30-001,protein_coding,ENSMUSG00000031609.3,protein_coding | 2723.161737 | 0.150146121 | 4.89E-05 | 0.000633326 |
| ENSMUST00000041362.11,Mavs-001,protein_coding,ENSMUSG00000037523.13,protein_coding | 5114.968906 | 0.178147889 | 4.91E-05 | 0.000635232 |
| ENSMUST00000139028.1,Pask-002,retained_intron,ENSMUSG00000026274.11,protein_coding | 692.9742575 | 0.471889438 | 5.00E-05 | 0.00064694 |
| ENSMUST00000148682.1,Tnks1bp1-005,processed_transcript,ENSMUSG00000033955.12,protein_coding | 1202.306067 | 0.616226635 | 5.00E-05 | 0.000647605 |
| ENSMUST00000038608.13,Mboat7-001,protein_coding,ENSMUSG00000035596.14,protein_coding | 4124.587148 | 0.163515643 | 5.01E-05 | 0.000648088 |
| ENSMUST00000000326.11,Bcl6b-001,protein_coding,ENSMUSG00000000317.11,protein_coding | 95.01045245 | -0.563649668 | 5.02E-05 | 0.000649061 |
| ENSMUST00000203638.2,Itpr1-005,protein_coding,ENSMUSG00000030102.11,protein_coding | 415.6732637 | -0.337802898 | 5.02E-05 | 0.000649076 |
| ENSMUST00000174609.8,Rps18-006,protein_coding,ENSMUSG00000008668.14,protein_coding | 1653.974588 | 0.267826622 | 5.02E-05 | 0.000649076 |
| ENSMUST00000164806.5,Adsl-003,protein_coding,ENSMUSG00000022407.9,protein_coding | 10009.49576 | 0.199954178 | 5.04E-05 | 0.000650669 |
| ENSMUST00000193338.5,Pmf1-002,protein_coding,ENSMUSG00000028066.15,protein_coding | 261.5392574 | 0.555097941 | 5.05E-05 | 0.000652211 |
| ENSMUST00000111343.1,Ran-002,protein_coding,ENSMUSG00000029430.13,protein_coding | 12563.08945 | 0.234095394 | 5.06E-05 | 0.000653269 |
| ENSMUST00000109587.8,Rbm39-001,protein_coding,ENSMUSG00000027620.16,protein_coding | 2435.810268 | -0.572621245 | 5.06E-05 | 0.000653571 |
| ENSMUST00000114224.7,Gng12-007,protein_coding,ENSMUSG00000036402.13,protein_coding | 225.5993361 | 0.614760594 | 5.07E-05 | 0.00065457 |
| ENSMUST00000112272.1,Wnt5a-002,protein_coding,ENSMUSG00000021994.14,protein_coding | 843.8851838 | -0.367093941 | 5.07E-05 | 0.00065457 |
| ENSMUST00000182892.7,Pcmtd1-004,retained_intron,ENSMUSG00000051285.17,protein_coding | 231.8405529 | -0.664375531 | 5.10E-05 | 0.000657669 |
| ENSMUST00000185621.2,Kansl3-007,nonsense_mediated_decay,ENSMUSG00000010453.12,protein_coding | 70.01105089 | 0.660746964 | 5.11E-05 | 0.000658894 |
| ENSMUST00000004428.13,Clcn5-001,protein_coding,ENSMUSG00000004317.14,protein_coding | 6405.307617 | 0.171965396 | 5.11E-05 | 0.00065912 |
| ENSMUST00000007257.9,Clic1-001,protein_coding,ENSMUSG00000007041.9,protein_coding | 32444.19002 | 0.204509619 | 5.12E-05 | 0.000659613 |
| ENSMUST00000020323.5,Avpr1a-001,protein_coding,ENSMUSG00000020123.5,protein_coding | 41.44153295 | -0.74752131 | 5.12E-05 | 0.000660014 |
| ENSMUST00000065308.11,Azin1-001,protein_coding,ENSMUSG00000037458.14,protein_coding | 1101.945256 | -0.576438393 | 5.13E-05 | 0.000660226 |
| ENSMUST00000066469.13,Cope-001,protein_coding,ENSMUSG00000055681.14,protein_coding | 9170.534946 | -0.192344406 | 5.15E-05 | 0.00066313 |
| ENSMUST00000022727.8,Tgds-201,protein_coding,ENSMUSG00000022130.9,protein_coding | 3037.963088 | 0.241417213 | 5.15E-05 | 0.00066313 |
| ENSMUST00000165971.1,Sept10-201,protein_coding,ENSMUSG00000019917.9,protein_coding | 4571.983462 | 0.276314239 | 5.17E-05 | 0.000664743 |
| ENSMUST00000105847.7,Kdm1a-002,protein_coding,ENSMUSG00000036940.15,protein_coding | 9545.982711 | 0.13858708 | 5.17E-05 | 0.000665052 |
| ENSMUST00000048155.15,Rbm25-002,protein_coding,ENSMUSG00000010608.15,protein_coding | 7931.172756 | -0.385985205 | 5.17E-05 | 0.000665052 |
| ENSMUST00000150843.7,Pank2-001,protein_coding,ENSMUSG00000037514.17,protein_coding | 2735.521392 | -0.357752874 | 5.18E-05 | 0.000665651 |
| ENSMUST00000160681.1,Hcfc2-002,protein_coding,ENSMUSG00000020246.13,protein_coding | 888.9902717 | -0.224770468 | 5.18E-05 | 0.000665651 |
| ENSMUST00000144378.1,N4bp2l2-002,retained_intron,ENSMUSG00000029655.17,protein_coding | 2242.157076 | 0.229172397 | 5.19E-05 | 0.000667266 |
| ENSMUST00000038816.12,Cxcl10-001,protein_coding,ENSMUSG00000034855.13,protein_coding | 1019.416415 | 0.315068881 | 5.21E-05 | 0.000669233 |
| ENSMUST00000151702.7,Rpl10-003,protein_coding,ENSMUSG00000008682.13,protein_coding | 27356.43491 | 0.215481466 | 5.22E-05 | 0.000669758 |
| ENSMUST00000145600.1,Atg2a-002,protein_coding,ENSMUSG00000024773.14,protein_coding | 1404.505872 | -0.462192033 | 5.22E-05 | 0.00067065 |
| ENSMUST00000022212.7,Plk2-201,protein_coding,ENSMUSG00000021701.7,protein_coding | 14530.61002 | 0.154939986 | 5.24E-05 | 0.000672008 |
| ENSMUST00000144349.1,Psip1-005,retained_intron,ENSMUSG00000028484.16,protein_coding | 289.7584299 | -0.48393273 | 5.25E-05 | 0.00067349 |
| ENSMUST00000001703.7,Hoxc8-001,protein_coding,ENSMUSG00000001657.7,protein_coding | 277.3079932 | -0.374439486 | 5.25E-05 | 0.000673851 |
| ENSMUST00000127305.1,Epn3-001,protein_coding,ENSMUSG00000010080.15,protein_coding | 1143.524127 | -0.308702614 | 5.26E-05 | 0.000674519 |
| ENSMUST00000167244.1,Immt-007,retained_intron,ENSMUSG00000052337.15,protein_coding | 166.781418 | -0.567472551 | 5.27E-05 | 0.000675388 |
| ENSMUST00000150227.7,Adarb1-009,retained_intron,ENSMUSG00000020262.15,protein_coding | 133.4081531 | -0.519743746 | 5.29E-05 | 0.000677242 |
| ENSMUST00000021187.11,Dhrs13-001,protein_coding,ENSMUSG00000020834.17,protein_coding | 2520.328309 | -0.232989907 | 5.29E-05 | 0.000677242 |
| ENSMUST00000137581.1,Snrpc-002,retained_intron,ENSMUSG00000024217.9,protein_coding | 323.5504063 | 0.384269312 | 5.29E-05 | 0.000677242 |
| ENSMUST00000202567.1,Preb-002,protein_coding,ENSMUSG00000045302.13,protein_coding | 8005.346439 | 0.136140351 | 5.30E-05 | 0.000678657 |
| ENSMUST00000067924.12,Lrrc8c-001,protein_coding,ENSMUSG00000054720.12,protein_coding | 5895.629572 | -0.153519007 | 5.30E-05 | 0.000678657 |
| ENSMUST00000199617.1,Cenpa-011,nonsense_mediated_decay,ENSMUSG00000029177.9,protein_coding | 423.9536055 | -0.357745106 | 5.30E-05 | 0.000678788 |
| ENSMUST00000067496.6,Atpif1-001,protein_coding,ENSMUSG00000054428.12,protein_coding | 9574.011209 | 0.14595686 | 5.32E-05 | 0.000680634 |
| ENSMUST00000090134.11,Pdlim5-010,protein_coding,ENSMUSG00000028273.15,protein_coding | 202.8766426 | 0.554978725 | 5.33E-05 | 0.000681838 |
| ENSMUST00000194111.5,Cr1l-003,protein_coding,ENSMUSG00000016481.15,protein_coding | 347.829215 | 0.488570129 | 5.34E-05 | 0.000683326 |
| ENSMUST00000062753.2,D030056L22Rik-202,protein_coding,ENSMUSG00000047044.7,protein_coding | 5621.883187 | 0.195409485 | 5.36E-05 | 0.000685229 |
| ENSMUST00000087086.6,Pax3-001,protein_coding,ENSMUSG00000004872.15,protein_coding | 2115.240636 | -0.290842346 | 5.37E-05 | 0.00068613 |
| ENSMUST00000024897.8,Vapa-201,protein_coding,ENSMUSG00000024091.8,protein_coding | 16735.5978 | 0.132511194 | 5.37E-05 | 0.000686262 |
| ENSMUST00000030606.13,Rcan3-001,protein_coding,ENSMUSG00000059713.12,protein_coding | 4335.176512 | 0.180390842 | 5.40E-05 | 0.000689413 |
| ENSMUST00000100196.8,Alg1-002,protein_coding,ENSMUSG00000039427.14,protein_coding | 3224.471408 | 0.17568092 | 5.42E-05 | 0.000692235 |
| ENSMUST00000029049.6,Chmp4c-001,protein_coding,ENSMUSG00000027536.6,protein_coding | 946.5663924 | -0.306649941 | 5.45E-05 | 0.000694941 |
| ENSMUST00000016696.12,Foxred2-001,protein_coding,ENSMUSG00000016552.13,protein_coding | 7947.446416 | 0.249119478 | 5.45E-05 | 0.000695139 |
| ENSMUST00000075164.10,Kif21b-002,protein_coding,ENSMUSG00000041642.18,protein_coding | 4028.548277 | 0.550917231 | 5.45E-05 | 0.000695314 |
| ENSMUST00000072836.6,Sphk2-002,protein_coding,ENSMUSG00000057342.15,protein_coding | 811.8635428 | 0.385252884 | 5.45E-05 | 0.000695411 |
| ENSMUST00000150683.1,Hdac5-008,retained_intron,ENSMUSG00000008855.17,protein_coding | 340.7701154 | -0.369509201 | 5.46E-05 | 0.00069547 |
| ENSMUST00000032728.8,Tarsl2-001,protein_coding,ENSMUSG00000030515.9,protein_coding | 2885.616162 | 0.171114532 | 5.50E-05 | 0.000700579 |
| ENSMUST00000034326.6,Atp13a1-201,protein_coding,ENSMUSG00000031862.10,protein_coding | 9413.006408 | -0.136051935 | 5.50E-05 | 0.000700579 |
| ENSMUST00000148553.1,Eef1b2-005,processed_transcript,ENSMUSG00000025967.16,protein_coding | 2214.410663 | 0.218641237 | 5.52E-05 | 0.000703243 |
| ENSMUST00000110731.3,Kdelr2-001,protein_coding,ENSMUSG00000079111.3,protein_coding | 19756.01332 | 0.143925697 | 5.53E-05 | 0.000703923 |
| ENSMUST00000145709.7,Rac1-003,processed_transcript,ENSMUSG00000001847.14,protein_coding | 7670.95895 | 0.155488643 | 5.55E-05 | 0.000706898 |
| ENSMUST00000109607.9,Cpne1-002,protein_coding,ENSMUSG00000074643.12,protein_coding | 14694.27273 | 0.202846794 | 5.59E-05 | 0.000711502 |
| ENSMUST00000052631.7,Snai1-001,protein_coding,ENSMUSG00000042821.7,protein_coding | 2974.547531 | -0.287314379 | 5.59E-05 | 0.000711502 |
| ENSMUST00000136960.7,Sh2b3-003,nonsense_mediated_decay,ENSMUSG00000042594.16,protein_coding | 3458.057463 | 0.163853964 | 5.60E-05 | 0.000711742 |
| ENSMUST00000114297.4,Vopp1-002,protein_coding,ENSMUSG00000037788.14,protein_coding | 4177.503742 | 0.221055145 | 5.60E-05 | 0.000711742 |
| ENSMUST00000093292.10,Sertad2-001,protein_coding,ENSMUSG00000049800.13,protein_coding | 1943.577715 | 0.239679085 | 5.62E-05 | 0.000714866 |
| ENSMUST00000034197.4,St3gal2-001,protein_coding,ENSMUSG00000031749.12,protein_coding | 6762.160252 | -0.162378471 | 5.64E-05 | 0.000716242 |
| ENSMUST00000201448.3,Slc37a3-002,nonsense_mediated_decay,ENSMUSG00000029924.12,protein_coding | 2762.77959 | 0.189209978 | 5.68E-05 | 0.000721917 |
| ENSMUST00000138822.7,Clk2-004,retained_intron,ENSMUSG00000068917.12,protein_coding | 269.7985575 | -0.618172439 | 5.72E-05 | 0.000725788 |
| ENSMUST00000172122.7,Rbm26-004,protein_coding,ENSMUSG00000022119.15,protein_coding | 832.3387079 | -0.53683753 | 5.72E-05 | 0.000725958 |
| ENSMUST00000026032.5,Pcgf6-201,protein_coding,ENSMUSG00000025050.7,protein_coding | 3071.394445 | 0.187360071 | 5.74E-05 | 0.000728202 |
| ENSMUST00000031017.10,Fosl2-001,protein_coding,ENSMUSG00000029135.10,protein_coding | 4490.05664 | -0.250569996 | 5.74E-05 | 0.000728629 |
| ENSMUST00000054197.6,S1pr2-001,protein_coding,ENSMUSG00000043895.6,protein_coding | 3119.326104 | -0.248430928 | 5.75E-05 | 0.000728629 |
| ENSMUST00000047203.8,Rnf126-201,protein_coding,ENSMUSG00000035890.8,protein_coding | 16445.21825 | 0.153545718 | 5.75E-05 | 0.000728629 |
| ENSMUST00000108817.4,Hist3h2a-001,protein_coding,ENSMUSG00000078851.4,protein_coding | 490.9401643 | -0.338006232 | 5.75E-05 | 0.000728903 |
| ENSMUST00000055226.12,Fn1-001,protein_coding,ENSMUSG00000026193.15,protein_coding | 102248.9875 | 0.211807581 | 5.77E-05 | 0.000730728 |
| ENSMUST00000060223.2,Zadh2-201,protein_coding,ENSMUSG00000049090.2,protein_coding | 4803.137193 | 0.17535852 | 5.78E-05 | 0.000731951 |
| ENSMUST00000201987.1,Pds5a-002,retained_intron,ENSMUSG00000029202.12,protein_coding | 338.1170147 | -0.602568191 | 5.81E-05 | 0.00073632 |
| ENSMUST00000197232.1,Aph1a-003,retained_intron,ENSMUSG00000015750.15,protein_coding | 410.9886128 | -0.479617685 | 5.83E-05 | 0.000737623 |
| ENSMUST00000047923.11,Sec24d-001,protein_coding,ENSMUSG00000039234.11,protein_coding | 13042.38386 | 0.184688006 | 5.84E-05 | 0.000739265 |
| ENSMUST00000040608.4,3930402G23Rik-001,lincRNA,ENSMUSG00000038917.4,lincRNA | 134.7564665 | -0.536974341 | 5.85E-05 | 0.000740412 |
| ENSMUST00000147797.7,Nol9-003,retained_intron,ENSMUSG00000028948.16,protein_coding | 199.9036909 | -0.437414346 | 5.88E-05 | 0.000744268 |

| | | | | |
|---|---|---|---|---|
| ENSMUST00000002303.11,Rhoc-001,protein_coding,ENSMUSG00000002233.13,protein_coding | 14754.22781 | 0.208745973 | 5.90E-05 | 0.000745737 |
| ENSMUST00000139782.7,Mthfsd-008,protein_coding,ENSMUSG00000031816.15,protein_coding | 694.0690363 | -0.367460305 | 5.90E-05 | 0.000745922 |
| ENSMUST00000104999.3,Nrarp-001,protein_coding,ENSMUSG00000078202.3,protein_coding | 1149.022938 | -0.319321244 | 5.91E-05 | 0.000746904 |
| ENSMUST00000180489.7,Mirt1-201,lincRNA,ENSMUSG00000097636.7,lincRNA | 279.0763223 | -0.450877977 | 5.96E-05 | 0.000752445 |
| ENSMUST00000107283.7,Snx27-002,protein_coding,ENSMUSG00000028136.15,protein_coding | 4214.228622 | 0.283848892 | 5.97E-05 | 0.000754292 |
| ENSMUST00000046719.13,Nfya-001,protein_coding,ENSMUSG00000023994.13,protein_coding | 4790.59918 | -0.250146065 | 5.97E-05 | 0.000754292 |
| ENSMUST00000079033.4,Bloc1s2-201,protein_coding,ENSMUSG00000057506.4,protein_coding | 4098.015886 | 0.161418659 | 5.98E-05 | 0.000754292 |
| ENSMUST00000030547.14,Hspg2-001,protein_coding,ENSMUSG00000028763.17,protein_coding | 89197.23221 | 0.145271965 | 5.99E-05 | 0.00075599 |
| ENSMUST00000034763.9,Rp9-001,protein_coding,ENSMUSG00000032239.10,protein_coding | 6119.81635 | 0.168951113 | 6.00E-05 | 0.000757252 |
| ENSMUST00000002683.2,Ccdc97-001,protein_coding,ENSMUSG00000002608.8,protein_coding | 5426.628514 | -0.186395427 | 6.01E-05 | 0.000757583 |
| ENSMUST00000003061.13,Bcam-001,protein_coding,ENSMUSG00000002980.14,protein_coding | 1107.970901 | -0.255498681 | 6.01E-05 | 0.000757815 |
| ENSMUST00000128318.7,Clk2-005,nonsense_mediated_decay,ENSMUSG00000068917.12,protein_coding | 173.7280213 | -0.747730004 | 6.02E-05 | 0.000758089 |
| ENSMUST00000212810.1,Dok4-002,protein_coding,ENSMUSG00000040631.8,protein_coding | 173.710944 | -0.69970211 | 6.02E-05 | 0.000758089 |
| ENSMUST00000132561.7,BC029214-010,processed_transcript,ENSMUSG00000047617.11,protein_coding | 73.62449577 | 0.668088479 | 6.02E-05 | 0.000758489 |
| ENSMUST00000211366.1,Snrnp70-004,retained_intron,ENSMUSG00000063511.11,protein_coding | 1651.95534 | -0.194504299 | 6.02E-05 | 0.000758489 |
| ENSMUST00000107818.8,Ankrd40-001,protein_coding,ENSMUSG00000020864.13,protein_coding | 5600.231736 | -0.212743876 | 6.03E-05 | 0.000759171 |
| ENSMUST00000065784.5,Ddx19b-002,protein_coding,ENSMUSG00000033658.16,protein_coding | 1126.772521 | 0.39431113 | 6.04E-05 | 0.00076083 |
| ENSMUST00000114911.7,Enox2-004,protein_coding,ENSMUSG00000031109.16,protein_coding | 430.4441282 | 0.511920396 | 6.05E-05 | 0.00076143 |
| ENSMUST00000147545.7,Ccdc6-001,protein_coding,ENSMUSG00000048701.13,protein_coding | 19174.58868 | 0.128481661 | 6.08E-05 | 0.000765264 |
| ENSMUST00000212320.1,Rfxank-002,protein_coding,ENSMUSG00000036120.10,protein_coding | 649.0126041 | 0.281457426 | 6.10E-05 | 0.000766574 |
| ENSMUST00000100143.9,Rc3h2-201,protein_coding,ENSMUSG00000075376.10,protein_coding | 3099.570416 | -0.548449292 | 6.11E-05 | 0.000767556 |
| ENSMUST00000029677.8,Zbtb7b-001,protein_coding,ENSMUSG00000028042.15,protein_coding | 7636.132977 | 0.271429205 | 6.11E-05 | 0.000767556 |
| ENSMUST00000001451.10,Smg5-001,protein_coding,ENSMUSG00000001415.10,protein_coding | 30542.17621 | -0.1809522 | 6.11E-05 | 0.000767766 |
| ENSMUST00000167020.7,Wac-001,protein_coding,ENSMUSG00000024283.14,protein_coding | 551.7506127 | -0.392531523 | 6.15E-05 | 0.000772197 |
| ENSMUST00000162903.6,Aebp2-008,protein_coding,ENSMUSG00000030232.17,protein_coding | 503.9602241 | -0.550591995 | 6.16E-05 | 0.000772612 |
| ENSMUST00000068056.11,Ddx39b-001,protein_coding,ENSMUSG00000019432.15,protein_coding | 35800.75532 | 0.159418823 | 6.16E-05 | 0.000772612 |
| ENSMUST00000131820.1,Myl12b-002,retained_intron,ENSMUSG00000034868.8,protein_coding | 526.8827234 | 0.310352876 | 6.15E-05 | 0.000772612 |
| ENSMUST00000038770.3,Vasn-001,protein_coding,ENSMUSG00000039646.5,protein_coding | 3835.204354 | -0.140432662 | 6.16E-05 | 0.000773179 |
| ENSMUST00000140115.1,Sdccag3-008,processed_transcript,ENSMUSG00000026927.17,protein_coding | 137.0283788 | 0.544120206 | 6.18E-05 | 0.000775 |
| ENSMUST00000188142.6,Peak1-002,protein_coding,ENSMUSG00000074305.9,protein_coding | 4844.927727 | -0.214827536 | 6.19E-05 | 0.000775725 |
| ENSMUST00000142536.1,Pced1a-006,retained_intron,ENSMUSG00000037773.14,protein_coding | 291.5001857 | 0.421452596 | 6.22E-05 | 0.000780153 |
| ENSMUST00000106251.9,Fus-001,protein_coding,ENSMUSG00000030795.18,protein_coding | 384.5549218 | -0.393997341 | 6.23E-05 | 0.000780631 |
| ENSMUST00000023044.5,Fam83f-201,protein_coding,ENSMUSG00000022408.6,protein_coding | 130.2091954 | 0.583473309 | 6.23E-05 | 0.000780791 |
| ENSMUST00000067530.4,Vapb-001,protein_coding,ENSMUSG00000054455.4,protein_coding | 17777.2842 | 0.173255534 | 6.24E-05 | 0.000781461 |
| ENSMUST00000152471.1,Snx21-002,protein_coding,ENSMUSG00000050373.13,protein_coding | 194.4229003 | 0.521878305 | 6.25E-05 | 0.000782789 |
| ENSMUST00000035158.15,Trf-001,protein_coding,ENSMUSG00000032554.15,protein_coding | 5000.493587 | -0.206321251 | 6.27E-05 | 0.000784823 |
| ENSMUST00000174583.7,Gigyf2-012,processed_transcript,ENSMUSG00000048000.15,protein_coding | 182.86237 | -0.607864693 | 6.28E-05 | 0.000785307 |
| ENSMUST00000200587.4,Klf3-002,retained_intron,ENSMUSG00000029178.14,protein_coding | 658.26554 | 0.591083681 | 6.28E-05 | 0.000786159 |
| ENSMUST00000088512.11,Rnps1-201,protein_coding,ENSMUSG00000034681.16,protein_coding | 6827.776812 | 0.256598737 | 6.30E-05 | 0.000787265 |
| ENSMUST00000119945.7,Mrps10-005,protein_coding,ENSMUSG00000034729.16,protein_coding | 2958.991852 | 0.187135389 | 6.32E-05 | 0.000789947 |
| ENSMUST00000074807.6,Lrig3-201,protein_coding,ENSMUSG00000020105.8,protein_coding | 1135.347165 | 0.349125747 | 6.34E-05 | 0.000792595 |
| ENSMUST00000169826.1,Rps24-202,protein_coding,ENSMUSG00000025290.16,protein_coding | 36766.2249 | -0.133944484 | 6.35E-05 | 0.00079403 |
| ENSMUST00000215998.1,Stt3a-009,processed_transcript,ENSMUSG00000032116.18,protein_coding | 4986.047805 | 0.165084197 | 6.38E-05 | 0.000796568 |
| ENSMUST00000108340.8,Asph-002,protein_coding,ENSMUSG00000028207.18,protein_coding | 2990.794448 | 0.539450734 | 6.38E-05 | 0.000797116 |
| ENSMUST00000143545.1,Gar1-005,retained_intron,ENSMUSG00000028010.16,protein_coding | 1654.983836 | 0.246126843 | 6.41E-05 | 0.00079987 |
| ENSMUST00000100705.10,Dusp14-002,protein_coding,ENSMUSG00000018648.15,protein_coding | 1175.799155 | -0.234118571 | 6.42E-05 | 0.000800734 |
| ENSMUST00000086167.11,Nfrkb-001,protein_coding,ENSMUSG00000042185.14,protein_coding | 3556.705716 | 0.237938777 | 6.44E-05 | 0.000803294 |
| ENSMUST00000107399.8,Nt5c3b-202,protein_coding,ENSMUSG00000017176.17,protein_coding | 92.36242279 | 0.742353972 | 6.44E-05 | 0.000803294 |
| ENSMUST00000040359.5,Arsi-001,protein_coding,ENSMUSG00000036412.5,protein_coding | 175.7865302 | -0.485436774 | 6.45E-05 | 0.000804284 |
| ENSMUST00000019224.7,Mfsd3-201,protein_coding,ENSMUSG00000019080.7,protein_coding | 457.0671102 | -0.358747241 | 6.47E-05 | 0.0008065 |
| ENSMUST00000102555.10,Dnajc24-001,protein_coding,ENSMUSG00000027166.14,protein_coding | 1377.308099 | 0.187650796 | 6.48E-05 | 0.000807044 |
| ENSMUST00000214054.1,Taf1d-006,nonsense_mediated_decay,ENSMUSG00000031939.16,protein_coding | 737.5700217 | -0.452074232 | 6.48E-05 | 0.000807044 |
| ENSMUST00000023737.4,Dhh-201,protein_coding,ENSMUSG00000023000.4,protein_coding | 53.60237243 | -0.644910877 | 6.48E-05 | 0.000807044 |
| ENSMUST00000149277.7,Atat1-003,protein_coding,ENSMUSG00000024426.17,protein_coding | 373.0168213 | -0.510077196 | 6.48E-05 | 0.000807044 |
| ENSMUST00000063249.8,Xrcc1-201,protein_coding,ENSMUSG00000051768.9,protein_coding | 8295.100364 | 0.141991954 | 6.50E-05 | 0.000808616 |
| ENSMUST00000028599.7,Cstf3-001,protein_coding,ENSMUSG00000027176.8,protein_coding | 4533.376086 | -0.184535675 | 6.50E-05 | 0.000808639 |
| ENSMUST00000058764.5,Lsm14b-002,protein_coding,ENSMUSG00000039108.14,protein_coding | 1641.24429 | 0.246424665 | 6.50E-05 | 0.000808639 |
| ENSMUST00000097837.10,Rap1gap-001,protein_coding,ENSMUSG00000041351.16,protein_coding | 1134.637836 | -0.288923319 | 6.50E-05 | 0.000808639 |
| ENSMUST00000072239.13,Tmpo-001,protein_coding,ENSMUSG00000019961.17,protein_coding | 19949.49114 | -0.251520909 | 6.50E-05 | 0.000808639 |
| ENSMUST00000006692.5,Mvd-001,protein_coding,ENSMUSG00000006517.5,protein_coding | 8966.564367 | 0.272117001 | 6.53E-05 | 0.000812299 |
| ENSMUST00000021207.6,Rflnb-001,protein_coding,ENSMUSG00000020846.6,protein_coding | 2938.934675 | 0.193846013 | 6.55E-05 | 0.000813477 |
| ENSMUST00000133602.7,1700037H04Rik-005,protein_coding,ENSMUSG00000027327.16,protein_coding | 53.5789818 | 0.685642673 | 6.56E-05 | 0.000815014 |
| ENSMUST00000156041.1,Nde1-004,retained_intron,ENSMUSG00000022678.16,protein_coding | 33.93751683 | -0.740206073 | 6.59E-05 | 0.000818204 |
| ENSMUST00000130836.7,Cyb5r1-004,processed_transcript,ENSMUSG00000026456.18,protein_coding | 62.31373907 | 0.743020182 | 6.60E-05 | 0.000819136 |
| ENSMUST00000147441.7,Cdk12-006,processed_transcript,ENSMUSG00000003119.14,protein_coding | 396.1286686 | -0.327864687 | 6.60E-05 | 0.000819826 |
| ENSMUST00000016383.9,Lonrf3-001,protein_coding,ENSMUSG00000016239.11,protein_coding | 1359.78515 | 0.315368507 | 6.62E-05 | 0.000821133 |
| ENSMUST00000096420.2,Gm14698-001,protein_coding,ENSMUSG00000071748.2,protein_coding | 1455.555844 | -0.577236461 | 6.62E-05 | 0.000821555 |
| ENSMUST00000152447.8,Nt5c3-002,nonsense_mediated_decay,ENSMUSG00000029780.14,protein_coding | 316.1802767 | 0.633705968 | 6.63E-05 | 0.000821956 |
| ENSMUST00000115443.7,Met-001,protein_coding,ENSMUSG00000009376.15,protein_coding | 22770.08586 | -0.246463081 | 6.63E-05 | 0.000822511 |
| ENSMUST00000037418.5,Tmed8-201,protein_coding,ENSMUSG00000034111.5,protein_coding | 944.8417896 | -0.260980657 | 6.64E-05 | 0.000823278 |
| ENSMUST00000203749.1,Aup1-015,retained_intron,ENSMUSG00000068328.9,protein_coding | 387.8408477 | 0.458797526 | 6.65E-05 | 0.00082355 |
| ENSMUST00000029128.3,Map1lc3a-001,protein_coding,ENSMUSG00000027602.9,protein_coding | 3352.588257 | 0.215918143 | 6.66E-05 | 0.00082502 |
| ENSMUST00000097646.3,2700062C07Rik-201,protein_coding,ENSMUSG00000024273.7,protein_coding | 1533.022515 | 0.224472873 | 6.68E-05 | 0.000826738 |
| ENSMUST00000164416.7,Pcid2-001,protein_coding,ENSMUSG00000038542.14,protein_coding | 8517.833607 | 0.166372956 | 6.68E-05 | 0.000827151 |
| ENSMUST00000044766.14,4930402H24Rik-001,protein_coding,ENSMUSG00000027309.18,protein_coding | 3462.331268 | 0.175925777 | 6.68E-05 | 0.000827195 |
| ENSMUST00000112205.1,Gorasp2-003,protein_coding,ENSMUSG00000014959.12,protein_coding | 2054.98218 | 0.316932261 | 6.71E-05 | 0.000830484 |
| ENSMUST00000023212.14,Maf1-001,protein_coding,ENSMUSG00000022553.15,protein_coding | 7155.177552 | 0.203158394 | 6.73E-05 | 0.000832316 |
| ENSMUST00000027487.14,Rnpepl1-001,protein_coding,ENSMUSG00000026269.14,protein_coding | 3627.555299 | -0.143463861 | 6.74E-05 | 0.000832934 |
| ENSMUST00000000724.14,Kat2b-001,protein_coding,ENSMUSG00000000708.14,protein_coding | 2071.198288 | -0.25518162 | 6.74E-05 | 0.000832934 |
| ENSMUST00000105372.7,Gpx4-001,protein_coding,ENSMUSG00000075706.10,protein_coding | 31615.56256 | 0.18257192 | 6.75E-05 | 0.000833752 |
| ENSMUST00000155712.8,Ncor1-003,protein_coding,ENSMUSG00000018501.17,protein_coding | 3627.460421 | -0.321148341 | 6.75E-05 | 0.000833752 |
| ENSMUST00000021689.12,Evl-201,protein_coding,ENSMUSG00000021262.14,protein_coding | 5878.275057 | 0.202270742 | 6.75E-05 | 0.000833752 |
| ENSMUST00000122384.7,Mras-002,protein_coding,ENSMUSG00000032470.16,protein_coding | 1147.720459 | 0.318129978 | 6.76E-05 | 0.00083518 |
| ENSMUST00000140405.8,Tmem134-006,processed_transcript,ENSMUSG00000024845.17,processed_transcript | 1961.197584 | 0.256629531 | 6.78E-05 | 0.000836876 |
| ENSMUST00000164765.1,Actb-005,retained_intron,ENSMUSG00000029580.14,protein_coding | 1375.974735 | -0.408667176 | 6.86E-05 | 0.000847102 |
| ENSMUST00000139649.7,Plekhm3-002,nonsense_mediated_decay,ENSMUSG00000051344.13,protein_coding | 167.1510309 | 0.742961758 | 6.87E-05 | 0.000847512 |
| ENSMUST00000147736.1,Brd3-006,protein_coding,ENSMUSG00000026918.16,protein_coding | 77.40679055 | 0.741892763 | 6.88E-05 | 0.000848213 |
| ENSMUST00000102598.3,Rbbp4-001,protein_coding,ENSMUSG00000057236.10,protein_coding | 53650.50098 | 0.170064623 | 6.91E-05 | 0.000852207 |
| ENSMUST00000124789.7,Adamtsl5-004,retained_intron,ENSMUSG00000043822.18,protein_coding | 534.8496159 | -0.398070382 | 6.91E-05 | 0.000852207 |
| ENSMUST00000154617.7,Rps6kb1-001,protein_coding,ENSMUSG00000020516.15,protein_coding | 3250.110928 | -0.467697747 | 6.97E-05 | 0.000858627 |
| ENSMUST00000159315.1,Ncor1-025,protein_coding,ENSMUSG00000018501.17,protein_coding | 344.4085224 | 0.352102374 | 6.98E-05 | 0.000859479 |
| ENSMUST00000003720.4,Crot-001,protein_coding,ENSMUSG00000003623.4,protein_coding | 2386.940056 | -0.227208868 | 6.98E-05 | 0.000859606 |

| | | | | |
|---|---|---|---|---|
| ENSMUST00000048274.10,Ttc39b-002,protein_coding,ENSMUSG00000038172.14,protein_coding | 1190.871459 | 0.304746425 | 7.01E-05 | 0.000863149 |
| ENSMUST00000039655.2,Tulp4-001,protein_coding,ENSMUSG00000034377.10,protein_coding | 8893.094877 | -0.342802608 | 7.02E-05 | 0.000864327 |
| ENSMUST00000182380.7,Pvt1-008,lincRNA,ENSMUSG00000097039.8,lincRNA | 40.77224314 | 0.554943674 | 7.03E-05 | 0.000864552 |
| ENSMUST00000112094.7,Pbrm1-003,protein_coding,ENSMUSG00000042323.16,protein_coding | 1815.295023 | -0.577238577 | 7.05E-05 | 0.00086678 |
| ENSMUST00000145391.7,Pdzd7-001,protein_coding,ENSMUSG00000074818.12,protein_coding | 131.7498202 | 0.57676158 | 7.06E-05 | 0.000867803 |
| ENSMUST00000066163.2,A930004D18Rik-001,protein_coding,ENSMUSG00000054057.2,protein_coding | 378.220149 | -0.429659622 | 7.06E-05 | 0.000867854 |
| ENSMUST00000107562.1,Cacnb1-004,protein_coding,ENSMUSG00000020882.17,protein_coding | 120.7882182 | -0.725779723 | 7.06E-05 | 0.000868116 |
| ENSMUST00000075220.12,Cdyl-201,protein_coding,ENSMUSG00000059288.13,protein_coding | 102.0688167 | 0.709700887 | 7.07E-05 | 0.000869161 |
| ENSMUST00000164430.6,Nfkb1-201,protein_coding,ENSMUSG00000028163.17,protein_coding | 4385.375624 | 0.211837964 | 7.09E-05 | 0.000870785 |
| ENSMUST00000192773.5,Dclk2-005,nonsense_mediated_decay,ENSMUSG00000028078.14,protein_coding | 93.6923533 | -0.545474791 | 7.10E-05 | 0.000871543 |
| ENSMUST00000124306.1,Adra1b-003,processed_transcript,ENSMUSG00000050541.14,protein_coding | 480.4558081 | 0.367476008 | 7.10E-05 | 0.000871543 |
| ENSMUST00000167574.1,Adra1b-201,protein_coding,ENSMUSG00000050541.14,protein_coding | 480.4558081 | 0.367476008 | 7.10E-05 | 0.000871543 |
| ENSMUST00000176957.7,Esd-002,protein_coding,ENSMUSG00000021996.16,protein_coding | 36653.83036 | 0.126793677 | 7.10E-05 | 0.000871543 |
| ENSMUST00000115168.8,Cep19-001,protein_coding,ENSMUSG00000035790.15,protein_coding | 1189.871332 | -0.299912608 | 7.11E-05 | 0.00087273 |
| ENSMUST00000160180.7,Cyb5a-001,protein_coding,ENSMUSG00000024646.13,protein_coding | 725.2272833 | -0.351121152 | 7.12E-05 | 0.000873793 |
| ENSMUST00000126000.7,Atg13-005,processed_transcript,ENSMUSG00000027244.14,protein_coding | 111.5565527 | 0.684535817 | 7.14E-05 | 0.000875908 |
| ENSMUST00000078912.6,Ampd2-003,protein_coding,ENSMUSG00000027889.17,protein_coding | 1705.922235 | 0.309523792 | 7.16E-05 | 0.000878084 |
| ENSMUST00000107208.7,Nbr1-005,protein_coding,ENSMUSG00000017119.19,protein_coding | 2243.824726 | 0.450267181 | 7.17E-05 | 0.000878084 |
| ENSMUST00000077867.9,Abcc5-004,protein_coding,ENSMUSG00000022822.15,protein_coding | 155.968265 | 0.630411482 | 7.17E-05 | 0.000878084 |
| ENSMUST00000084935.10,Pgm2l1-001,protein_coding,ENSMUSG00000030729.17,protein_coding | 831.6346932 | -0.487196631 | 7.19E-05 | 0.000880246 |
| ENSMUST00000149168.1,Slc6a9-002,nonsense_mediated_decay,ENSMUSG00000028542.17,protein_coding | 427.1632874 | 0.550189783 | 7.20E-05 | 0.000881464 |
| ENSMUST00000028608.12,Nat10-001,protein_coding,ENSMUSG00000027185.14,protein_coding | 12651.51915 | 0.182388716 | 7.23E-05 | 0.000884578 |
| ENSMUST00000078404.13,Pcbp2-202,protein_coding,ENSMUSG00000056851.13,protein_coding | 10294.43332 | -0.230739548 | 7.27E-05 | 0.000889209 |
| ENSMUST00000203907.2,Osbpl3-011,protein_coding,ENSMUSG00000029822.15,protein_coding | 100.9269564 | 0.735670058 | 7.28E-05 | 0.000890177 |
| ENSMUST00000166117.1,Gen1-201,protein_coding,ENSMUSG00000051235.8,protein_coding | 1577.085117 | -0.418327202 | 7.28E-05 | 0.000890494 |
| ENSMUST00000098346.4,Man2a2-001,protein_coding,ENSMUSG00000038886.10,protein_coding | 10862.87026 | 0.197130566 | 7.30E-05 | 0.000892634 |
| ENSMUST00000050312.2,Mapk8ip1-001,protein_coding,ENSMUSG00000027223.15,protein_coding | 2128.842108 | -0.300223923 | 7.33E-05 | 0.000895376 |
| ENSMUST00000072566.4,Nme2-002,protein_coding,ENSMUSG00000020857.11,protein_coding | 19263.63285 | 0.248878054 | 7.33E-05 | 0.000895376 |
| ENSMUST00000110483.8,Stard13-001,protein_coding,ENSMUSG00000016128.14,protein_coding | 1339.202796 | -0.415808977 | 7.34E-05 | 0.000897005 |
| ENSMUST00000038614.11,Ypel3-001,protein_coding,ENSMUSG00000042675.15,protein_coding | 884.372357 | 0.372100914 | 7.37E-05 | 0.000899854 |
| ENSMUST00000058868.8,9530077C05Rik-001,protein_coding,ENSMUSG00000036411.10,protein_coding | 192.8376833 | -0.477014918 | 7.37E-05 | 0.000899854 |
| ENSMUST00000156974.7,5530601H04Rik-004,processed_transcript,ENSMUSG00000087174.7,processed_transcript | 118.5542098 | -0.636612633 | 7.37E-05 | 0.000899854 |
| ENSMUST00000034822.11,Acsbg1-001,protein_coding,ENSMUSG00000032281.11,protein_coding | 4066.701905 | 0.217213062 | 7.39E-05 | 0.000901508 |
| ENSMUST00000110698.8,Ghr-002,protein_coding,ENSMUSG00000055737.12,protein_coding | 712.0140985 | 0.264349562 | 7.42E-05 | 0.000904475 |
| ENSMUST00000108807.8,Zgpat-002,protein_coding,ENSMUSG00000027582.16,protein_coding | 1421.332936 | 0.225964384 | 7.42E-05 | 0.000904914 |
| ENSMUST00000135909.7,Chd3-003,retained_intron,ENSMUSG00000018474.17,protein_coding | 509.6844536 | -0.485857787 | 7.42E-05 | 0.000904914 |
| ENSMUST00000038211.12,Slfn9-001,protein_coding,ENSMUSG00000069793.12,protein_coding | 2192.605567 | -0.555637651 | 7.43E-05 | 0.000904914 |
| ENSMUST00000023099.6,Slc38a2-201,protein_coding,ENSMUSG00000022462.6,protein_coding | 58851.10959 | 0.125442885 | 7.43E-05 | 0.000904914 |
| ENSMUST00000020203.6,Snrpf-001,protein_coding,ENSMUSG00000020018.6,protein_coding | 11563.12394 | 0.098826396 | 7.43E-05 | 0.000905319 |
| ENSMUST00000181062.7,Agrn-008,retained_intron,ENSMUSG00000041936.18,protein_coding | 321.6206297 | 0.422630488 | 7.44E-05 | 0.000905431 |
| ENSMUST00000020909.3,Laptm4a-001,protein_coding,ENSMUSG00000020585.4,protein_coding | 23400.57418 | 0.178672539 | 7.44E-05 | 0.000905908 |
| ENSMUST00000136970.7,Crtc2-002,processed_transcript,ENSMUSG00000027936.18,protein_coding | 1714.720553 | 0.339137019 | 7.47E-05 | 0.000908965 |
| ENSMUST00000020717.11,Arf5-001,protein_coding,ENSMUSG00000020440.13,protein_coding | 16250.15546 | 0.203109999 | 7.47E-05 | 0.000908965 |
| ENSMUST00000163401.8,Rfx7-001,protein_coding,ENSMUSG00000037674.15,protein_coding | 4742.666558 | -0.264922419 | 7.51E-05 | 0.000912938 |
| ENSMUST00000170811.7,Rnf14-202,protein_coding,ENSMUSG00000060450.13,protein_coding | 4425.411526 | 0.187308435 | 7.51E-05 | 0.000913371 |
| ENSMUST00000167571.7,Gspt1-001,protein_coding,ENSMUSG00000062203.14,protein_coding | 7042.807069 | -0.205139451 | 7.53E-05 | 0.000915074 |
| ENSMUST00000015903.10,Cnih1-001,protein_coding,ENSMUSG00000015759.10,protein_coding | 19006.78837 | 0.147737453 | 7.56E-05 | 0.000918245 |
| ENSMUST00000054979.9,Aff1-001,protein_coding,ENSMUSG00000029313.18,protein_coding | 17402.31554 | 0.154657785 | 7.56E-05 | 0.000918635 |
| ENSMUST00000034472.15,Jam3-002,protein_coding,ENSMUSG00000031990.15,protein_coding | 925.6406104 | -0.253774273 | 7.57E-05 | 0.000919249 |
| ENSMUST00000127316.7,Stam2-002,protein_coding,ENSMUSG00000055371.17,protein_coding | 777.465221 | 0.326622849 | 7.57E-05 | 0.000919491 |
| ENSMUST00000105964.1,Gmeb1-003,protein_coding,ENSMUSG00000028901.13,protein_coding | 2132.97234 | 0.225560248 | 7.65E-05 | 0.000928518 |
| ENSMUST00000154609.2,Hmg20b-009,protein_coding,ENSMUSG00000020232.17,protein_coding | 113.4251618 | 0.710739078 | 7.65E-05 | 0.000928518 |
| ENSMUST00000033184.5,Tpp1-001,protein_coding,ENSMUSG00000030894.5,protein_coding | 5818.416867 | 0.156847237 | 7.66E-05 | 0.000929094 |
| ENSMUST00000145285.1,P2rx3-004,processed_transcript,ENSMUSG00000027071.14,protein_coding | 24.51889674 | 0.738258339 | 7.66E-05 | 0.000929227 |
| ENSMUST00000018909.3,Fxr2-006,protein_coding,ENSMUSG00000018765.5,protein_coding | 29000.72085 | 0.202005758 | 7.67E-05 | 0.000930149 |
| ENSMUST00000017430.11,Glod4-001,protein_coding,ENSMUSG00000017286.15,protein_coding | 6245.189185 | 0.198188465 | 7.68E-05 | 0.00093058 |
| ENSMUST00000125482.1,Ciz1-015,protein_coding,ENSMUSG00000039205.16,protein_coding | 409.1540679 | 0.408799251 | 7.69E-05 | 0.000931368 |
| ENSMUST00000129200.1,Cnpy3-002,nonsense_mediated_decay,ENSMUSG00000023973.13,protein_coding | 177.1228167 | 0.730609304 | 7.69E-05 | 0.000931862 |
| ENSMUST00000211235.1,Nup98-003,protein_coding,ENSMUSG00000063550.8,protein_coding | 3448.979692 | 0.176398466 | 7.71E-05 | 0.000933596 |
| ENSMUST00000105861.7,Srrm1-003,protein_coding,ENSMUSG00000028809.16,protein_coding | 16322.38052 | -0.203371438 | 7.75E-05 | 0.000938061 |
| ENSMUST00000144859.3,Selenof-003,processed_transcript,ENSMUSG00000037072.15,protein_coding | 8787.173986 | 0.236542001 | 7.77E-05 | 0.000940041 |
| ENSMUST00000113643.7,Phyhd1-004,protein_coding,ENSMUSG00000079484.12,protein_coding | 57.01542887 | 0.731464843 | 7.80E-05 | 0.000943853 |
| ENSMUST00000134254.1,0610037L13Rik-010,protein_coding,ENSMUSG00000028608.15,protein_coding | 21.26220631 | 0.733099239 | 7.80E-05 | 0.000943853 |
| ENSMUST00000204588.1,RP24-470M22.10-001,TEC,ENSMUSG00000107838.1,TEC | 178.7625842 | 0.47903207 | 7.81E-05 | 0.000944741 |
| ENSMUST00000097644.8,Hdac4-001,protein_coding,ENSMUSG00000026313.16,protein_coding | 3679.613288 | 0.204361615 | 7.82E-05 | 0.000945471 |
| ENSMUST00000061537.11,2810025M15Rik-002,processed_transcript,ENSMUSG00000049881.13,transcribed_processed_pseudogene | 3048.40289 | 0.214528667 | 7.83E-05 | 0.000946532 |
| ENSMUST00000148295.7,Fkbp1a-010,retained_intron,ENSMUSG00000032966.14,protein_coding | 545.4607819 | 0.384362306 | 7.84E-05 | 0.000947001 |
| ENSMUST00000034465.8,2810004N23Rik-001,protein_coding,ENSMUSG00000031984.8,protein_coding | 7440.353692 | 0.220063525 | 7.84E-05 | 0.000947306 |
| ENSMUST00000063980.7,Zkscan7-001,protein_coding,ENSMUSG00000063488.6,protein_coding | 287.6830421 | 0.367879452 | 7.84E-05 | 0.000947306 |
| ENSMUST00000134632.1,Ran-003,processed_transcript,ENSMUSG00000029430.13,protein_coding | 3776.576064 | 0.200768936 | 7.85E-05 | 0.000948045 |
| ENSMUST00000032508.10,Fkbp4-001,protein_coding,ENSMUSG00000030357.10,protein_coding | 81394.16632 | 0.149659337 | 7.86E-05 | 0.000948045 |
| ENSMUST00000130801.7,Trim26-008,protein_coding,ENSMUSG00000024457.16,protein_coding | 148.7977556 | 0.736090527 | 7.86E-05 | 0.000948045 |
| ENSMUST00000160561.2,2700081O15Rik-001,retained_intron,ENSMUSG00000053080.10,protein_coding | 301.6105681 | -0.668467305 | 7.85E-05 | 0.000948045 |
| ENSMUST00000051221.12,Ankrd40-003,protein_coding,ENSMUSG00000020864.13,protein_coding | 2269.901197 | 0.209762224 | 7.86E-05 | 0.000948267 |
| ENSMUST00000164034.7,Rhobtb1-011,protein_coding,ENSMUSG00000019944.14,protein_coding | 169.7909243 | 0.670696113 | 7.88E-05 | 0.000949839 |
| ENSMUST00000076265.12,Upf3b-001,protein_coding,ENSMUSG00000036572.16,protein_coding | 2038.483903 | -0.445366799 | 7.91E-05 | 0.000953339 |
| ENSMUST00000075650.6,Gm9761-001,processed_pseudogene,ENSMUSG00000034437.8,processed_pseudogene | 202.2761562 | -0.599979644 | 7.94E-05 | 0.000956509 |
| ENSMUST00000186914.1,Srrm2-004,processed_transcript,ENSMUSG00000039218.16,protein_coding | 1402.954256 | -0.344134413 | 7.96E-05 | 0.000958665 |
| ENSMUST00000126116.2,Capns1-201,protein_coding,ENSMUSG00000001794.13,protein_coding | 10950.35608 | 0.247733236 | 8.00E-05 | 0.000963279 |
| ENSMUST00000198826.4,Zcchc8-006,nonsense_mediated_decay,ENSMUSG00000029427.11,protein_coding | 686.5101508 | 0.296207803 | 8.01E-05 | 0.000964065 |
| ENSMUST00000184951.7,Mphosph9-011,protein_coding,ENSMUSG00000038126.17,protein_coding | 685.4185665 | 0.519699402 | 8.01E-05 | 0.000964269 |
| ENSMUST00000162848.1,Lss-004,retained_intron,ENSMUSG00000033105.12,protein_coding | 900.2784474 | 0.419958737 | 8.01E-05 | 0.00096445 |
| ENSMUST00000015333.11,Casd1-001,protein_coding,ENSMUSG00000015189.12,protein_coding | 3380.442411 | -0.287254584 | 8.04E-05 | 0.000967589 |
| ENSMUST00000019991.7,Tube1-001,protein_coding,ENSMUSG00000019845.8,protein_coding | 1171.19236 | 0.344526301 | 8.05E-05 | 0.000968214 |
| ENSMUST00000022148.6,Mccc2-201,protein_coding,ENSMUSG00000021646.8,protein_coding | 2715.94313 | 0.202817762 | 8.06E-05 | 0.000968631 |
| ENSMUST00000027493.3,Pask-001,protein_coding,ENSMUSG00000026274.11,protein_coding | 4303.684923 | 0.256728317 | 8.07E-05 | 0.000969486 |
| ENSMUST00000080356.9,Snrpa-001,protein_coding,ENSMUSG00000061479.15,protein_coding | 7503.626158 | 0.151853379 | 8.08E-05 | 0.000971131 |
| ENSMUST00000108337.7,Asph-006,protein_coding,ENSMUSG00000028207.18,protein_coding | 1086.294061 | -0.659974739 | 8.09E-05 | 0.00097125 |
| ENSMUST00000106648.7,Mical2-005,retained_intron,ENSMUSG00000038244.13,protein_coding | 282.6915497 | -0.511230196 | 8.09E-05 | 0.00097125 |
| ENSMUST00000023101.9,Slc38a4-201,protein_coding,ENSMUSG00000022464.13,protein_coding | 3060.220145 | -0.274181237 | 8.09E-05 | 0.00097125 |
| ENSMUST00000115160.9,Tmem209-001,protein_coding,ENSMUSG00000029782.18,protein_coding | 5975.293469 | -0.217834925 | 8.13E-05 | 0.000975644 |
| ENSMUST00000130411.6,Srsf6-001,protein_coding,ENSMUSG00000016921.14,protein_coding | 15576.17547 | -0.245811631 | 8.15E-05 | 0.000977404 |

| | | | | |
|---|---|---|---|---|
| ENSMUST00000077783.4,0610030E20Rik-001,protein_coding,ENSMUSG00000058706.5,protein_coding | 4022.986582 | -0.1708823 | 8.15E-05 | 0.000977404 |
| ENSMUST00000019362.14,Dvl2-001,protein_coding,ENSMUSG00000020888.16,protein_coding | 8883.026658 | 0.163222221 | 8.16E-05 | 0.000978548 |
| ENSMUST00000191651.5,Rabgap1l-007,protein_coding,ENSMUSG00000026721.15,protein_coding | 34.52065633 | -0.735300321 | 8.18E-05 | 0.000980439 |
| ENSMUST00000018755.9,Pdlim4-001,protein_coding,ENSMUSG00000020388.12,protein_coding | 422.1663658 | -0.36741033 | 8.21E-05 | 0.000984269 |
| ENSMUST00000100655.4,9130401M01Rik-001,protein_coding,ENSMUSG00000101892.1,protein_coding | 7376.809254 | 0.206707426 | 8.22E-05 | 0.000985248 |
| ENSMUST00000081688.12,Srgap1-201,protein_coding,ENSMUSG00000020121.15,protein_coding | 1092.641114 | -0.370766916 | 8.27E-05 | 0.000990903 |
| ENSMUST00000029722.6,Rps3a1-001,protein_coding,ENSMUSG00000028081.6,protein_coding | 113120.0101 | 0.116834754 | 8.29E-05 | 0.000992954 |
| ENSMUST00000077817.7,Hnrnph1-006,protein_coding,ENSMUSG00000007850.16,protein_coding | 2096.456029 | -0.431101559 | 8.31E-05 | 0.000994905 |
| ENSMUST00000028761.4,Ubox5-001,protein_coding,ENSMUSG00000027300.9,protein_coding | 1879.876138 | 0.257737178 | 8.32E-05 | 0.000996001 |
| ENSMUST00000022696.6,Xpo7-201,protein_coding,ENSMUSG00000022100.13,protein_coding | 22213.6685 | 0.139463887 | 8.32E-05 | 0.000996134 |
| ENSMUST00000113696.7,Tpm1-015,protein_coding,ENSMUSG00000032366.15,protein_coding | 1284.32396 | 0.403990253 | 8.35E-05 | 0.000999454 |
| ENSMUST00000176272.1,Arhgef2-029,processed_transcript,ENSMUSG00000028059.15,protein_coding | 658.1694933 | 0.589240103 | 8.36E-05 | 0.000999873 |
| ENSMUST00000125153.8,Epb41l1-003,protein_coding,ENSMUSG00000027624.17,protein_coding | 402.6310326 | -0.371236875 | 8.37E-05 | 0.001001221 |
| ENSMUST00000038475.8,Fa2h-001,protein_coding,ENSMUSG00000033579.16,protein_coding | 54.27080599 | 0.698936159 | 8.40E-05 | 0.001004296 |
| ENSMUST00000130729.2,Ssrp1-002,protein_coding,ENSMUSG00000027067.16,protein_coding | 2932.639262 | 0.33301316 | 8.41E-05 | 0.001005075 |
| ENSMUST00000206414.1,Tsku-002,protein_coding,ENSMUSG00000049580.12,protein_coding | 2491.812571 | 0.261013159 | 8.41E-05 | 0.001005075 |
| ENSMUST00000059078.3,Fam43a-001,protein_coding,ENSMUSG00000046546.3,protein_coding | 753.6203387 | -0.353816628 | 8.41E-05 | 0.001005075 |
| ENSMUST00000174260.7,Vars-004,protein_coding,ENSMUSG00000007029.16,protein_coding | 2341.338839 | 0.213648863 | 8.43E-05 | 0.001006058 |
| ENSMUST00000035194.7,Mapkapk3-001,protein_coding,ENSMUSG00000032577.16,protein_coding | 1395.110102 | -0.318931954 | 8.45E-05 | 0.001009007 |
| ENSMUST00000074945.6,Agap1-003,protein_coding,ENSMUSG00000055013.15,protein_coding | 2465.184573 | -0.295321958 | 8.47E-05 | 0.001011368 |
| ENSMUST00000052865.15,Tmem123-001,protein_coding,ENSMUSG00000050912.15,protein_coding | 6383.09913 | 0.203402253 | 8.49E-05 | 0.0010131 |
| ENSMUST00000161262.7,AW554918-001,processed_transcript,ENSMUSG00000033632.15,protein_coding | 41.52990171 | 0.695850502 | 8.49E-05 | 0.001013166 |
| ENSMUST00000021670.14,Ylpm1-001,protein_coding,ENSMUSG00000021244.15,protein_coding | 8234.48721 | -0.183818608 | 8.53E-05 | 0.001016615 |
| ENSMUST00000112585.7,Apobec1-002,protein_coding,ENSMUSG00000040613.14,protein_coding | 155.2457441 | 0.657128674 | 8.56E-05 | 0.00102044 |
| ENSMUST00000075578.6,Abcb10-001,protein_coding,ENSMUSG00000031974.8,protein_coding | 3278.698245 | -0.185728754 | 8.56E-05 | 0.00102044 |
| ENSMUST00000002923.8,Adprh-201,protein_coding,ENSMUSG00000002844.8,protein_coding | 6324.465996 | 0.225360763 | 8.56E-05 | 0.00102044 |
| ENSMUST00000025183.8,Ring1-001,protein_coding,ENSMUSG00000024325.8,protein_coding | 2171.790504 | 0.229040733 | 8.60E-05 | 0.001024681 |
| ENSMUST00000137705.7,Fam53a-005,nonsense_mediated_decay,ENSMUSG00000037339.17,protein_coding | 49.66243981 | 0.668758298 | 8.61E-05 | 0.001025058 |
| ENSMUST00000042818.9,Pim3-201,protein_coding,ENSMUSG00000035828.10,protein_coding | 20045.08929 | 0.223854555 | 8.62E-05 | 0.001026116 |
| ENSMUST00000110529.5,Flt1-002,protein_coding,ENSMUSG00000029648.13,protein_coding | 642.4174091 | -0.36404878 | 8.62E-05 | 0.001026144 |
| ENSMUST00000020922.7,Trib2-001,protein_coding,ENSMUSG00000020601.7,protein_coding | 2884.158725 | -0.188905683 | 8.63E-05 | 0.001027176 |
| ENSMUST00000022469.12,Nisch-004,protein_coding,ENSMUSG00000021910.15,protein_coding | 30823.15267 | 0.214610554 | 8.68E-05 | 0.001032778 |
| ENSMUST00000146375.5,Kmt2e-003,nonsense_mediated_decay,ENSMUSG00000029004.15,protein_coding | 52.21160309 | 0.700524368 | 8.69E-05 | 0.001033773 |
| ENSMUST00000113805.7,Pnkd-003,protein_coding,ENSMUSG00000026179.14,protein_coding | 1874.731413 | -0.206648181 | 8.74E-05 | 0.001038744 |
| ENSMUST00000144742.5,Cenpa-001,protein_coding,ENSMUSG00000029177.9,protein_coding | 4682.561016 | -0.243993704 | 8.74E-05 | 0.001038744 |
| ENSMUST00000197711.1,Dnajb14-004,protein_coding,ENSMUSG00000074212.7,protein_coding | 88.43518565 | 0.731591348 | 8.75E-05 | 0.001039076 |
| ENSMUST00000105821.2,Kif17-004,protein_coding,ENSMUSG00000028758.12,protein_coding | 41.00701869 | 0.731854442 | 8.75E-05 | 0.001039076 |
| ENSMUST00000170953.2,Rps13-001,protein_coding,ENSMUSG00000090862.3,protein_coding | 48933.4239 | 0.169690708 | 8.77E-05 | 0.00104124 |
| ENSMUST00000118459.1,Gm13835-001,processed_pseudogene,ENSMUSG00000086922.2,processed_pseudogene | 249.9198487 | -0.39355604 | 8.79E-05 | 0.001043627 |
| ENSMUST00000132705.7,Cdca2-002,protein_coding,ENSMUSG00000048922.18,protein_coding | 288.4967973 | 0.731918113 | 8.83E-05 | 0.001047901 |
| ENSMUST00000122222.7,Casz1-004,protein_coding,ENSMUSG00000028977.16,protein_coding | 101.3822388 | -0.583257001 | 8.85E-05 | 0.001050123 |
| ENSMUST00000030642.2,Psmb2-001,protein_coding,ENSMUSG00000028837.8,protein_coding | 16682.41097 | 0.168492738 | 8.86E-05 | 0.001050372 |
| ENSMUST00000117654.2,Tbrg1-001,protein_coding,ENSMUSG00000011114.18,protein_coding | 11722.18013 | 0.172223937 | 8.97E-05 | 0.001063672 |
| ENSMUST00000105082.8,Plekha6-201,protein_coding,ENSMUSG00000041757.16,protein_coding | 567.4874248 | -0.512575118 | 8.98E-05 | 0.001063804 |
| ENSMUST00000107112.1,Capn5-002,protein_coding,ENSMUSG00000035547.14,protein_coding | 424.2536312 | 0.518000647 | 8.98E-05 | 0.001063804 |
| ENSMUST00000049032.12,Gmeb2-001,protein_coding,ENSMUSG00000038705.13,protein_coding | 6008.561438 | 0.199586509 | 9.01E-05 | 0.001066609 |
| ENSMUST00000030401.13,Ndufs5-001,protein_coding,ENSMUSG00000028648.13,protein_coding | 5666.783033 | 0.158526193 | 9.01E-05 | 0.001066609 |
| ENSMUST00000199632.1,4833413G10Rik-001,antisense,ENSMUSG00000104950.1,antisense | 82.6125917 | -0.606998841 | 9.01E-05 | 0.001066609 |
| ENSMUST00000207932.1,Prss23-001,protein_coding,ENSMUSG00000039405.7,protein_coding | 6893.204648 | -0.24235031 | 9.01E-05 | 0.001066609 |
| ENSMUST00000091201.6,Arl15-001,protein_coding,ENSMUSG00000042348.9,protein_coding | 1461.094701 | 0.226895272 | 9.01E-05 | 0.001066609 |
| ENSMUST00000103015.3,Narf-001,protein_coding,ENSMUSG00000000056.7,protein_coding | 2059.646348 | 0.211107863 | 9.02E-05 | 0.001067043 |
| ENSMUST00000038172.14,Mtmr12-001,protein_coding,ENSMUSG00000039458.14,protein_coding | 3274.886105 | 0.18325669 | 9.02E-05 | 0.001067043 |
| ENSMUST00000159801.7,Tbc1d31-002,retained_intron,ENSMUSG00000022364.14,protein_coding | 1839.543124 | 0.44382845 | 9.02E-05 | 0.001067043 |
| ENSMUST00000114200.9,Fam219b-001,protein_coding,ENSMUSG00000032305.15,protein_coding | 1930.936656 | 0.364784237 | 9.08E-05 | 0.001073172 |
| ENSMUST00000142589.5,Frmd4b-002,retained_intron,ENSMUSG00000030064.16,protein_coding | 71.10406775 | -0.6879693 | 9.10E-05 | 0.001076031 |
| ENSMUST00000127674.7,Ppp1r16a-002,retained_intron,ENSMUSG00000033819.15,protein_coding | 137.8155628 | -0.695875112 | 9.12E-05 | 0.001077917 |
| ENSMUST00000002318.7,Zfp523-005,protein_coding,ENSMUSG00000024220.13,protein_coding | 37.7320211 | 0.689921519 | 9.12E-05 | 0.001077917 |
| ENSMUST00000120080.7,Adcy9-002,protein_coding,ENSMUSG00000005580.11,protein_coding | 832.7025161 | 0.377321933 | 9.13E-05 | 0.001078141 |
| ENSMUST00000217624.1,6030419C18Rik-003,retained_intron,ENSMUSG00000066607.5,protein_coding | 194.4326631 | 0.651317829 | 9.14E-05 | 0.001079113 |
| ENSMUST00000168129.8,Trappc12-202,protein_coding,ENSMUSG00000020628.15,protein_coding | 2522.192186 | 0.226611347 | 9.27E-05 | 0.001094577 |
| ENSMUST00000188519.1,Gm7160-002,lincRNA,ENSMUSG00000099843.1,lincRNA | 327.764551 | 0.366292666 | 9.28E-05 | 0.001094807 |
| ENSMUST00000115519.7,Ica1-006,protein_coding,ENSMUSG00000062995.12,protein_coding | 1675.719055 | 0.357819037 | 9.30E-05 | 0.001096528 |
| ENSMUST00000129285.7,Pdgfc-002,nonsense_mediated_decay,ENSMUSG00000028019.9,protein_coding | 908.1015959 | 0.302312159 | 9.31E-05 | 0.001097899 |
| ENSMUST00000140305.7,Gtf2a2-007,protein_coding,ENSMUSG00000033543.17,protein_coding | 25.67779088 | -0.727935633 | 9.34E-05 | 0.001100914 |
| ENSMUST00000021347.11,Sel1l-001,protein_coding,ENSMUSG00000020964.14,protein_coding | 8603.407486 | 0.247799413 | 9.35E-05 | 0.001101549 |
| ENSMUST00000015501.10,Clpx-001,protein_coding,ENSMUSG00000015357.10,protein_coding | 3540.018131 | 0.159133313 | 9.35E-05 | 0.001101808 |
| ENSMUST00000034742.7,Ccnb2-001,protein_coding,ENSMUSG00000032218.7,protein_coding | 15238.99889 | 0.10986288 | 9.38E-05 | 0.001104669 |
| ENSMUST00000160560.1,Sharpin-010,nonsense_mediated_decay,ENSMUSG00000022552.15,protein_coding | 111.2879807 | -0.530163559 | 9.38E-05 | 0.001105297 |
| ENSMUST00000023072.6,Parvb-001,protein_coding,ENSMUSG00000022438.6,protein_coding | 29649.19247 | 0.147515714 | 9.41E-05 | 0.001108267 |
| ENSMUST00000037708.9,Asic4-001,protein_coding,ENSMUSG00000033007.15,protein_coding | 313.2264277 | 0.508891717 | 9.47E-05 | 0.001114287 |
| ENSMUST00000193384.1,Tada3-007,protein_coding,ENSMUSG00000048930.12,protein_coding | 54.559066 | 0.651989271 | 9.47E-05 | 0.001114457 |
| ENSMUST00000136872.7,Mtch2-001,protein_coding,ENSMUSG00000027282.17,protein_coding | 5381.964748 | 0.163334279 | 9.48E-05 | 0.001115297 |
| ENSMUST00000131374.7,Rps15a-001,protein_coding,ENSMUSG00000008683.16,protein_coding | 12455.37243 | 0.182719882 | 9.52E-05 | 0.001119165 |
| ENSMUST00000164022.7,Glod4-007,protein_coding,ENSMUSG00000017286.15,protein_coding | 766.4468086 | 0.421492548 | 9.57E-05 | 0.001124658 |
| ENSMUST00000167459.1,Mbnl2-202,protein_coding,ENSMUSG00000022139.16,protein_coding | 11250.89059 | -0.42307866 | 9.59E-05 | 0.001127202 |
| ENSMUST00000099547.3,Fam8a1-001,protein_coding,ENSMUSG00000069237.8,protein_coding | 4490.051958 | -0.280279779 | 9.60E-05 | 0.001128367 |
| ENSMUST00000154160.1,Crls1-004,protein_coding,ENSMUSG00000027357.16,protein_coding | 180.8767393 | 0.721374327 | 9.63E-05 | 0.001131191 |
| ENSMUST00000186364.1,Plscr1-002,protein_coding,ENSMUSG00000032369.13,protein_coding | 5015.116535 | 0.184366066 | 9.63E-05 | 0.001131265 |
| ENSMUST00000109572.1,Epb41l1-008,protein_coding,ENSMUSG00000027624.17,protein_coding | 35.34153461 | -0.72777506 | 9.65E-05 | 0.001132722 |
| ENSMUST00000111786.8,Mapkapk5-005,protein_coding,ENSMUSG00000029454.15,protein_coding | 1527.041622 | 0.262956441 | 9.65E-05 | 0.001133369 |
| ENSMUST00000046533.7,Prr7-201,protein_coding,ENSMUSG00000034686.7,protein_coding | 903.2966822 | -0.27258194 | 9.67E-05 | 0.001134358 |
| ENSMUST00000113359.7,Frmd4b-005,protein_coding,ENSMUSG00000030064.16,protein_coding | 2552.283313 | -0.349598059 | 9.67E-05 | 0.001134373 |
| ENSMUST00000181570.1,Gpr137b-ps-201,lincRNA,ENSMUSG00000097715.1,lincRNA | 706.354968 | 0.432691061 | 9.67E-05 | 0.001134373 |
| ENSMUST00000115351.9,Rab3d-001,protein_coding,ENSMUSG00000019066.13,protein_coding | 2452.623158 | 0.241922542 | 9.70E-05 | 0.001136547 |
| ENSMUST00000177738.7,Rbm10-203,protein_coding,ENSMUSG00000031060.16,protein_coding | 3027.341323 | 0.178376002 | 9.70E-05 | 0.001136547 |
| ENSMUST00000055931.4,Dusp18-201,protein_coding,ENSMUSG00000047205.12,protein_coding | 4245.59515 | 0.244781908 | 9.72E-05 | 0.001139002 |
| ENSMUST00000136631.1,Nrip2-003,retained_intron,ENSMUSG00000001520.12,protein_coding | 100.9940892 | 0.614886462 | 9.75E-05 | 0.001141784 |
| ENSMUST00000161075.7,Desi2-002,protein_coding,ENSMUSG00000026502.13,protein_coding | 2714.868018 | 0.244429017 | 9.76E-05 | 0.001143372 |
| ENSMUST00000123640.1,Lck-004,retained_intron,ENSMUSG00000000409.14,protein_coding | 145.6681829 | 0.557951837 | 9.79E-05 | 0.001146 |
| ENSMUST00000183887.1,Acaca-008,protein_coding,ENSMUSG00000020532.18,protein_coding | 317.0619422 | 0.451396374 | 9.79E-05 | 0.001146 |
| ENSMUST00000204087.1,Rab11fip5-001,protein_coding,ENSMUSG00000051343.11,protein_coding | 3825.031887 | 0.202480621 | 9.79E-05 | 0.001146108 |
| ENSMUST00000001327.9,Itgb7-001,protein_coding,ENSMUSG00000001281.9,protein_coding | 1297.521784 | 0.278533157 | 9.84E-05 | 0.001150891 |

| | | | | |
|---|---|---|---|---|
| ENSMUST00000032956.9,Sgf29-001,protein_coding,ENSMUSG00000030714.14,protein_coding | 1706.481283 | 0.204862481 | 9.85E-05 | 0.001152583 |
| ENSMUST00000058714.9,Cd24a-001,protein_coding,ENSMUSG00000047139.9,protein_coding | 24036.90474 | -0.310071086 | 9.89E-05 | 0.001156816 |
| ENSMUST00000111775.7,Xpr1-003,protein_coding,ENSMUSG00000026469.14,protein_coding | 1630.663504 | 0.362345302 | 9.90E-05 | 0.001157392 |
| ENSMUST00000036210.6,Poglut1-001,protein_coding,ENSMUSG00000034064.14,protein_coding | 5026.446069 | 0.184527334 | 9.90E-05 | 0.001157392 |
| ENSMUST00000034244.8,Tmem38a-001,protein_coding,ENSMUSG00000031791.8,protein_coding | 627.8905541 | -0.392579011 | 9.91E-05 | 0.00115782 |
| ENSMUST00000032566.2,Qpctl-001,protein_coding,ENSMUSG00000030407.2,protein_coding | 1106.124985 | -0.223022686 | 9.92E-05 | 0.001159215 |
| ENSMUST00000036049.5,Hsdl1-001,protein_coding,ENSMUSG00000034189.5,protein_coding | 2426.074544 | 0.178407689 | 9.94E-05 | 0.001160296 |
| ENSMUST00000140045.1,Sephs1-004,processed_transcript,ENSMUSG00000026662.13,protein_coding | 512.8014917 | 0.427212024 | 9.95E-05 | 0.001162056 |
| ENSMUST00000049382.5,Gatad2b-201,protein_coding,ENSMUSG00000042390.9,protein_coding | 1576.04526 | 0.272053448 | 9.97E-05 | 0.001163216 |
| ENSMUST00000171561.7,Mms19-001,protein_coding,ENSMUSG00000025159.8,protein_coding | 29.54731639 | -0.633125535 | 9.98E-05 | 0.001165105 |
| ENSMUST00000024594.8,Agpat4-001,protein_coding,ENSMUSG00000023827.8,protein_coding | 4507.658301 | 0.148419793 | 0.000100381 | 0.001171001 |
| ENSMUST00000023156.12,Il1rap-004,protein_coding,ENSMUSG00000022514.14,protein_coding | 780.7905202 | -0.339171698 | 0.000100493 | 0.001171993 |
| ENSMUST00000135645.1,Cops7a-007,retained_intron,ENSMUSG00000030127.15,protein_coding | 1044.253489 | 0.374541304 | 0.000100537 | 0.001172187 |
| ENSMUST00000126135.1,Plk3-002,processed_transcript,ENSMUSG00000028680.14,protein_coding | 153.0775513 | -0.593342915 | 0.000100692 | 0.001173675 |
| ENSMUST00000051259.9,Adgrg3-001,protein_coding,ENSMUSG00000060470.7,protein_coding | 266.3330278 | -0.403908127 | 0.000100936 | 0.001176199 |
| ENSMUST00000136269.7,Rpl7a-002,processed_transcript,ENSMUSG00000062647.16,protein_coding | 3780.271782 | 0.251435133 | 0.000101294 | 0.00117509 |
| ENSMUST00000126794.1,Rpl28-003,protein_coding,ENSMUSG00000030432.12,protein_coding | 6255.204794 | 0.23035599 | 0.000101278 | 0.001179509 |
| ENSMUST00000108829.7,Zfp672-003,protein_coding,ENSMUSG00000049755.14,protein_coding | 155.5572477 | -0.627742495 | 0.000101302 | 0.001179509 |
| ENSMUST00000156461.1,Anp32a-005,protein_coding,ENSMUSG00000032249.14,protein_coding | 556.7554862 | 0.327656321 | 0.000101648 | 0.001183214 |
| ENSMUST00000017622.11,Zc3h18-001,protein_coding,ENSMUSG00000017478.15,protein_coding | 6463.217805 | 0.284705553 | 0.000101916 | 0.001186018 |
| ENSMUST00000044277.9,Chmp4b-001,protein_coding,ENSMUSG00000038467.15,protein_coding | 25878.09109 | 0.170735564 | 0.000101984 | 0.001186491 |
| ENSMUST00000029025.8,Zdbf2-201,protein_coding,ENSMUSG00000027520.15,protein_coding | 27.89115684 | -0.716314882 | 0.000102013 | 0.001186501 |
| ENSMUST00000124720.7,Tmem260-005,protein_coding,ENSMUSG00000036339.17,protein_coding | 142.3777078 | 0.706150062 | 0.000102231 | 0.001188721 |
| ENSMUST00000142259.1,Lztr1-003,protein_coding,ENSMUSG00000022761.10,protein_coding | 271.9281244 | -0.498995036 | 0.000102347 | 0.001189747 |
| ENSMUST00000056181.6,Snx21-001,protein_coding,ENSMUSG00000050373.13,protein_coding | 577.1905668 | -0.360261764 | 0.000102526 | 0.0011915 |
| ENSMUST00000188882.1,Tsn-005,retained_intron,ENSMUSG00000026374.14,protein_coding | 1741.9129 | 0.203248638 | 0.000102815 | 0.001194543 |
| ENSMUST00000041382.6,Fuk-001,protein_coding,ENSMUSG00000033703.9,protein_coding | 3188.488746 | -0.21929707 | 0.000103001 | 0.001196377 |
| ENSMUST00000001790.5,Cldn15-201,protein_coding,ENSMUSG00000001739.14,protein_coding | 139.3726525 | -0.708215741 | 0.000103079 | 0.001196962 |
| ENSMUST00000079286.3,S100a7a-001,protein_coding,ENSMUSG00000063767.4,protein_coding | 115.2912072 | -0.530917948 | 0.000103138 | 0.001197321 |
| ENSMUST00000197513.1,Agap3-003,protein_coding,ENSMUSG00000023353.14,protein_coding | 3889.695959 | 0.201169887 | 0.00010328 | 0.001198655 |
| ENSMUST00000109085.7,Gnas-014,protein_coding,ENSMUSG00000027523.19,protein_coding | 27178.09411 | 0.195220214 | 0.000103332 | 0.001198927 |
| ENSMUST00000164826.7,Card10-001,protein_coding,ENSMUSG00000033170.14,protein_coding | 1850.186109 | -0.262196947 | 0.000103972 | 0.001206028 |
| ENSMUST00000024931.4,Tbc1d24-201,protein_coding,ENSMUSG00000036473.15,protein_coding | 124.8371016 | -0.581362531 | 0.000104038 | 0.001206472 |
| ENSMUST00000106149.7,Ssx2ip-004,protein_coding,ENSMUSG00000036825.12,protein_coding | 366.4092311 | 0.56432447 | 0.000105445 | 0.001222455 |
| ENSMUST00000141568.1,Zfand5-002,retained_intron,ENSMUSG00000024750.10,protein_coding | 225.2661402 | 0.538013426 | 0.000105715 | 0.001225256 |
| ENSMUST00000179474.8,Fam160a2-001,protein_coding,ENSMUSG00000044465.19,protein_coding | 958.9188904 | -0.352368105 | 0.000105757 | 0.001225415 |
| ENSMUST00000206403.1,Gm44950-001,TEC,ENSMUSG00000108350.1,TEC | 750.4710472 | -0.24226962 | 0.000105982 | 0.001227689 |
| ENSMUST00000027897.7,Smyd2-001,protein_coding,ENSMUSG00000026603.13,protein_coding | 11188.45907 | 0.173812874 | 0.000106186 | 0.001229728 |
| ENSMUST00000081926.12,Zfp341-001,protein_coding,ENSMUSG00000059842.12,protein_coding | 910.893406 | 0.255277203 | 0.000107279 | 0.001242045 |
| ENSMUST00000212976.1,Adgrg1-010,protein_coding,ENSMUSG00000031785.16,protein_coding | 151.3030672 | 0.639223313 | 0.000107409 | 0.001243222 |
| ENSMUST00000108721.7,Atpaf2-001,protein_coding,ENSMUSG00000042709.14,protein_coding | 3499.594306 | -0.212761516 | 0.000107444 | 0.001243292 |
| ENSMUST00000105907.8,Tmem222-001,protein_coding,ENSMUSG00000028857.16,protein_coding | 5726.485021 | 0.21610912 | 0.000108052 | 0.001249991 |
| ENSMUST00000050970.3,Polr1d-001,protein_coding,ENSMUSG00000029642.11,protein_coding | 7940.784757 | -0.197967315 | 0.000108522 | 0.001255086 |
| ENSMUST00000005532.7,Nid1-201,protein_coding,ENSMUSG00000005397.7,protein_coding | 40699.09294 | 0.155580916 | 0.000108743 | 0.001257306 |
| ENSMUST00000109149.8,Rnd1-003,protein_coding,ENSMUSG00000054855.13,protein_coding | 319.6455031 | 0.504587381 | 0.000109035 | 0.001260347 |
| ENSMUST00000003313.8,Grk5-201,protein_coding,ENSMUSG00000003228.8,protein_coding | 384.3443061 | -0.363202653 | 0.000109086 | 0.001260589 |
| ENSMUST00000085272.6,Htatip2-001,protein_coding,ENSMUSG00000039745.8,protein_coding | 586.1077539 | 0.540192407 | 0.000109661 | 0.001266903 |
| ENSMUST00000201226.1,Brca2-007,retained_intron,ENSMUSG00000041147.10,protein_coding | 23.15545217 | 0.721299583 | 0.000109805 | 0.001268223 |
| ENSMUST00000107353.2,Zfp592-001,protein_coding,ENSMUSG00000005621.11,protein_coding | 4213.467849 | 0.129476319 | 0.000109834 | 0.001268223 |
| ENSMUST00000037270.3,Fam91a1-201,protein_coding,ENSMUSG00000037119.3,protein_coding | 24694.50762 | -0.175839991 | 0.00010987 | 0.001268292 |
| ENSMUST00000103232.1,2510039O18Rik-001,protein_coding,ENSMUSG00000044496.6,protein_coding | 7563.283026 | 0.12999873 | 0.000110057 | 0.00127011 |
| ENSMUST00000103032.10,Llgl2-001,protein_coding,ENSMUSG00000020782.18,protein_coding | 877.2377588 | -0.321069322 | 0.000110233 | 0.001271797 |
| ENSMUST00000040234.8,Tsen2-001,protein_coding,ENSMUSG00000042389.13,protein_coding | 2553.035593 | 0.229179869 | 0.000110716 | 0.001277036 |
| ENSMUST00000195714.1,Gm37305-001,antisense,ENSMUSG00000103041.1,antisense | 428.1596682 | 0.306444388 | 0.000110783 | 0.001277459 |
| ENSMUST00000111070.3,Pnpla6-201,protein_coding,ENSMUSG00000004565.15,protein_coding | 615.8624807 | 0.719889085 | 0.000111197 | 0.001281887 |
| ENSMUST00000139322.7,Hspa4-003,retained_intron,ENSMUSG00000020361.13,protein_coding | 3234.058209 | 0.218171819 | 0.000111502 | 0.001285068 |
| ENSMUST00000125786.7,9430015G10Rik-002,processed_transcript,ENSMUSG00000059939.13,protein_coding | 1546.572611 | 0.191130325 | 0.000111626 | 0.00128615 |
| ENSMUST00000102645.3,Med19-001,protein_coding,ENSMUSG00000027080.6,protein_coding | 7718.253872 | 0.15258962 | 0.000111723 | 0.00128692 |
| ENSMUST00000030896.14,Tprgl-001,protein_coding,ENSMUSG00000029030.14,protein_coding | 3787.145283 | 0.215352116 | 0.00011218 | 0.001291835 |
| ENSMUST00000025101.8,Dync2li1-201,protein_coding,ENSMUSG00000024253.8,protein_coding | 2547.068076 | 0.203936132 | 0.000112249 | 0.001292294 |
| ENSMUST00000093043.6,Gas8-001,protein_coding,ENSMUSG00000040220.10,protein_coding | 2857.827302 | -0.206337123 | 0.000112323 | 0.001292792 |
| ENSMUST00000159110.7,Apaf1-002,protein_coding,ENSMUSG00000019979.12,protein_coding | 1617.878322 | -0.498077546 | 0.000112417 | 0.001293529 |
| ENSMUST00000023958.9,P3h3-001,protein_coding,ENSMUSG00000023191.9,protein_coding | 2646.47679 | 0.214745481 | 0.000112466 | 0.001293745 |
| ENSMUST00000168445.1,Cyb561a3-201,protein_coding,ENSMUSG00000034445.7,protein_coding | 1782.507139 | 0.193644714 | 0.000112676 | 0.001295819 |
| ENSMUST00000150157.1,Srsf3-005,retained_intron,ENSMUSG00000071172.12,protein_coding | 4225.054856 | -0.306859523 | 0.000114175 | 0.001312706 |
| ENSMUST00000185408.6,Fn1-012,retained_intron,ENSMUSG00000026193.15,protein_coding | 1006.306977 | -0.357613173 | 0.000114299 | 0.001313429 |
| ENSMUST00000074038.5,Acp1-202,protein_coding,ENSMUSG00000044573.14,protein_coding | 5825.523271 | -0.17590098 | 0.000114286 | 0.001313429 |
| ENSMUST00000061508.7,Zfp358-001,protein_coding,ENSMUSG00000047264.8,protein_coding | 3567.786448 | 0.186988298 | 0.00011473 | 0.001318027 |
| ENSMUST00000028072.12,Dnajc1-008,nonsense_mediated_decay,ENSMUSG00000026740.12,protein_coding | 329.4574679 | 0.613978535 | 0.000115173 | 0.001322774 |
| ENSMUST00000156876.7,Mfsd6-001,protein_coding,ENSMUSG00000041439.15,protein_coding | 879.532983 | 0.304312858 | 0.000115301 | 0.00132388 |
| ENSMUST00000024123.8,Agap3-001,protein_coding,ENSMUSG00000023353.14,protein_coding | 3588.959314 | -0.213345465 | 0.000115419 | 0.001324889 |
| ENSMUST00000004140.10,Gstm1-001,protein_coding,ENSMUSG00000058135.12,protein_coding | 8835.8608 | 0.153377962 | 0.000115595 | 0.001326555 |
| ENSMUST00000025762.14,Banf1-201,protein_coding,ENSMUSG00000024844.14,protein_coding | 26538.93054 | 0.159360443 | 0.000115646 | 0.001326786 |
| ENSMUST00000057921.9,Arf2-001,protein_coding,ENSMUSG00000062421.13,protein_coding | 2026.528698 | -0.249736928 | 0.00011573 | 0.001327397 |
| ENSMUST00000029421.5,Ptx3-001,protein_coding,ENSMUSG00000027832.5,protein_coding | 800.6597258 | 0.545075687 | 0.000115856 | 0.001328487 |
| ENSMUST00000043305.13,Wdtc1-001,protein_coding,ENSMUSG00000037622.14,protein_coding | 7286.168781 | 0.143027123 | 0.000115935 | 0.001328688 |
| ENSMUST00000127423.1,Mtch1-007,processed_transcript,ENSMUSG00000024012.17,protein_coding | 80.22717584 | 0.673400462 | 0.000115905 | 0.001328688 |
| ENSMUST00000103029.9,Rhbdf2-003,protein_coding,ENSMUSG00000020806.15,protein_coding | 1745.339329 | 0.253568627 | 0.00011598 | 0.001328849 |
| ENSMUST00000122817.7,Becn1-003,retained_intron,ENSMUSG00000035086.13,protein_coding | 1149.606896 | 0.310187408 | 0.000116201 | 0.001331025 |
| ENSMUST00000144698.7,Gpx4-006,processed_transcript,ENSMUSG00000075706.10,protein_coding | 149.7883413 | 0.589212917 | 0.00011641 | 0.001333064 |
| ENSMUST00000041968.10,Tmem135-001,protein_coding,ENSMUSG00000039428.11,protein_coding | 2388.323508 | 0.177103132 | 0.000116734 | 0.001336418 |
| ENSMUST00000202217.2,Sept11-006,protein_coding,ENSMUSG00000058013.11,protein_coding | 2504.231257 | 0.288841625 | 0.000116849 | 0.001337022 |
| ENSMUST00000187609.1,Nupr1-006,nonsense_mediated_decay,ENSMUSG00000030717.9,protein_coding | 202.4618556 | 0.651210942 | 0.000116843 | 0.001337022 |
| ENSMUST00000179802.1,Cmtm4-001,protein_coding,ENSMUSG00000096188.1,protein_coding | 18586.95279 | 0.159939756 | 0.000117008 | 0.001338482 |
| ENSMUST00000175776.7,Eps15-005,protein_coding,ENSMUSG00000028552.13,protein_coding | 1668.837727 | 0.260476192 | 0.000117102 | 0.001339205 |
| ENSMUST00000076316.5,Arhgap10-001,protein_coding,ENSMUSG00000037148.8,protein_coding | 8339.070807 | 0.144004276 | 0.000117602 | 0.001344572 |
| ENSMUST00000111557.7,Dennd5b-001,protein_coding,ENSMUSG00000030313.15,protein_coding | 2900.357205 | -0.309296984 | 0.000117645 | 0.001344704 |
| ENSMUST00000106211.1,Selenof-002,protein_coding,ENSMUSG00000037072.15,protein_coding | 11276.17359 | 0.310683545 | 0.000118282 | 0.001351267 |
| ENSMUST00000107824.8,Zcchc7-003,protein_coding,ENSMUSG00000035649.17,protein_coding | 1896.304255 | -0.303118394 | 0.000118269 | 0.001351267 |
| ENSMUST00000211172.1,Cars2-012,retained_intron,ENSMUSG00000056228.10,protein_coding | 238.817411 | -0.424984987 | 0.000118582 | 0.001353974 |
| ENSMUST00000049908.10,Ssbp4-001,protein_coding,ENSMUSG00000070003.7,protein_coding | 4256.805853 | -0.183387035 | 0.000118554 | 0.001353974 |
| ENSMUST00000033734.13,Ap1s2-002,protein_coding,ENSMUSG00000031367.15,protein_coding | 490.8832773 | 0.355562565 | 0.00011881 | 0.001356219 |

| | | | | |
|---|---|---|---|---|
| ENSMUST00000028293.11,Sdccag3-001,protein_coding,ENSMUSG00000026927.17,protein_coding | 2703.624682 | -0.363380426 | 0.000119012 | 0.00135816 |
| ENSMUST00000133777.1,Ptgs1-005,retained_intron,ENSMUSG00000047250.13,protein_coding | 140.9753172 | 0.486912382 | 0.000119212 | 0.001359359 |
| ENSMUST00000100652.9,Gtf2ird1-002,protein_coding,ENSMUSG00000023079.14,protein_coding | 504.519166 | -0.717311 | 0.00011918 | 0.001359359 |
| ENSMUST00000150327.7,Vegfa-007,retained_intron,ENSMUSG00000023951.17,protein_coding | 836.9097337 | 0.470812611 | 0.000119177 | 0.001359359 |
| ENSMUST00000198140.4,Pfkfb4-002,protein_coding,ENSMUSG00000025648.17,protein_coding | 3241.603391 | -0.412924579 | 0.000119501 | 0.001362304 |
| ENSMUST00000033489.7,Praf2-001,protein_coding,ENSMUSG00000031149.7,protein_coding | 6865.747473 | 0.148908661 | 0.000119616 | 0.001363246 |
| ENSMUST00000194666.1,Ndufaf3-002,retained_intron,ENSMUSG00000070283.4,protein_coding | 81.85175889 | 0.615353833 | 0.00011997 | 0.001366916 |
| ENSMUST00000151346.7,Macf1-009,protein_coding,ENSMUSG00000028649.17,protein_coding | 23088.16499 | -0.240654704 | 0.000120105 | 0.001368095 |
| ENSMUST00000072875.14,Srsf11-002,protein_coding,ENSMUSG00000055436.18,protein_coding | 9406.191604 | -0.154902397 | 0.000120168 | 0.001368451 |
| ENSMUST00000182250.1,Gm26982-001,antisense,ENSMUSG00000097994.1,antisense | 190.2492826 | -0.471604617 | 0.000120474 | 0.00137121 |
| ENSMUST00000003521.9,Rps11-001,protein_coding,ENSMUSG00000003429.11,protein_coding | 63177.94048 | 0.176368846 | 0.000120449 | 0.00137121 |
| ENSMUST00000046425.15,Txlna-201,protein_coding,ENSMUSG00000053841.15,protein_coding | 10667.01911 | -0.176312145 | 0.000120607 | 0.001372358 |
| ENSMUST00000031390.9,Mmp17-001,protein_coding,ENSMUSG00000029436.9,protein_coding | 560.6290129 | 0.373358375 | 0.000120886 | 0.001375056 |
| ENSMUST00000043520.4,Fbxo25-001,protein_coding,ENSMUSG00000038365.8,protein_coding | 2427.528751 | 0.191158592 | 0.000120908 | 0.001375056 |
| ENSMUST00000071169.8,Rcc2-001,protein_coding,ENSMUSG00000040945.13,protein_coding | 5546.665733 | 0.4373163 | 0.000121061 | 0.001376435 |
| ENSMUST00000002735.7,Clpp-201,protein_coding,ENSMUSG00000002660.7,protein_coding | 7675.750212 | 0.148788631 | 0.000121132 | 0.001376883 |
| ENSMUST00000002572.4,Slc9a3r2-201,protein_coding,ENSMUSG00000002504.14,protein_coding | 3947.896418 | -0.19962154 | 0.000121229 | 0.001377616 |
| ENSMUST00000048544.12,Bicd2-201,protein_coding,ENSMUSG00000037933.15,protein_coding | 804.1012447 | -0.271516807 | 0.000121288 | 0.001377931 |
| ENSMUST00000130454.7,Tead4-002,protein_coding,ENSMUSG00000030353.15,protein_coding | 556.1830836 | 0.403293337 | 0.000121405 | 0.001378891 |
| ENSMUST00000096990.9,Cdc45-003,protein_coding,ENSMUSG00000000028.14,protein_coding | 3032.810204 | 0.32868783 | 0.000121702 | 0.001381903 |
| ENSMUST00000164538.1,Otog-001,protein_coding,ENSMUSG00000009487.10,protein_coding | 35.26074946 | 0.711913598 | 0.000122556 | 0.001391233 |
| ENSMUST00000023775.7,Cela1-201,protein_coding,ENSMUSG00000023031.7,protein_coding | 835.4006156 | -0.329766393 | 0.000122687 | 0.001392356 |
| ENSMUST00000022245.8,Mrps30-001,protein_coding,ENSMUSG00000021731.9,protein_coding | 9650.391649 | 0.160697342 | 0.000123132 | 0.001397035 |
| ENSMUST00000037337.9,Tbc1d19-002,protein_coding,ENSMUSG00000039178.9,protein_coding | 1821.618403 | -0.261721308 | 0.000123526 | 0.001401134 |
| ENSMUST00000006435.7,Atp6v1b2-001,protein_coding,ENSMUSG00000006273.14,protein_coding | 13519.55496 | 0.193559096 | 0.000123627 | 0.001401915 |
| ENSMUST00000030078.11,Hsdl2-001,protein_coding,ENSMUSG00000028383.17,protein_coding | 1981.911375 | -0.19423816 | 0.000123666 | 0.001401979 |
| ENSMUST00000017821.11,Wsb1-001,protein_coding,ENSMUSG00000017677.11,protein_coding | 4497.542686 | -0.279554578 | 0.000123861 | 0.001403819 |
| ENSMUST00000132347.7,Trrap-003,retained_intron,ENSMUSG00000045482.16,protein_coding | 997.3053666 | 0.327412911 | 0.000124062 | 0.00140573 |
| ENSMUST00000087342.12,Rps28-002,protein_coding,ENSMUSG00000067288.12,protein_coding | 12379.94925 | 0.159448299 | 0.000124263 | 0.001407638 |
| ENSMUST00000095048.5,Rab7-003,protein_coding,ENSMUSG00000079477.9,protein_coding | 7627.035754 | 0.214324069 | 0.000124619 | 0.001409897 |
| ENSMUST00000026092.8,3110040N11Rik-001,protein_coding,ENSMUSG00000025102.14,protein_coding | 702.5751532 | 0.331352253 | 0.000124562 | 0.001409897 |
| ENSMUST00000199868.4,Shisa5-021,protein_coding,ENSMUSG00000025647.16,protein_coding | 635.9504792 | 0.445357478 | 0.000124504 | 0.001409897 |
| ENSMUST00000059551.4,Slc39a3-002,protein_coding,ENSMUSG00000046822.12,protein_coding | 742.85745 | -0.343424324 | 0.00012466 | 0.001409897 |
| ENSMUST00000115805.1,Ntan1-004,protein_coding,ENSMUSG00000022681.14,protein_coding | 115.6162679 | 0.651068954 | 0.000124563 | 0.001409897 |
| ENSMUST00000163107.7,Wiz-003,nonsense_mediated_decay,ENSMUSG00000024050.15,protein_coding | 6572.603708 | 0.211057639 | 0.000124679 | 0.001409897 |
| ENSMUST00000005975.6,Gpr108-201,protein_coding,ENSMUSG00000005823.8,protein_coding | 4035.957821 | 0.177877222 | 0.000124692 | 0.001409897 |
| ENSMUST00000093120.12,Kars-002,protein_coding,ENSMUSG00000031948.14,protein_coding | 7752.692586 | -0.212008077 | 0.000124754 | 0.001410233 |
| ENSMUST00000170310.1,Arpp19-004,protein_coding,ENSMUSG00000007656.13,protein_coding | 581.6474929 | -0.304532801 | 0.00012499 | 0.001412531 |
| ENSMUST00000046675.11,Iqcc-001,protein_coding,ENSMUSG00000040795.12,protein_coding | 2013.631802 | 0.214499543 | 0.000126359 | 0.001427625 |
| ENSMUST00000033882.9,Chrna6-001,protein_coding,ENSMUSG00000031491.10,protein_coding | 186.6718755 | -0.582118765 | 0.000126451 | 0.001428294 |
| ENSMUST00000203519.2,Magi1-007,protein_coding,ENSMUSG00000045095.16,protein_coding | 1862.396328 | 0.162967797 | 0.000126666 | 0.001430344 |
| ENSMUST00000121618.1,Lhfpl2-003,protein_coding,ENSMUSG00000045312.11,protein_coding | 1500.522916 | 0.640038519 | 0.000126774 | 0.001431185 |
| ENSMUST00000045372.5,Bpgm-001,protein_coding,ENSMUSG00000038871.5,protein_coding | 2108.902261 | -0.225831246 | 0.000127067 | 0.001433902 |
| ENSMUST00000129229.1,Pwwp2a-002,processed_transcript,ENSMUSG00000044950.7,protein_coding | 565.876359 | -0.273622409 | 0.000127081 | 0.001433902 |
| ENSMUST00000162372.7,Slc30a9-001,protein_coding,ENSMUSG00000029221.15,protein_coding | 2711.846448 | 0.162929498 | 0.000127369 | 0.001436769 |
| ENSMUST00000186474.1,2410022M11Rik-001,antisense,ENSMUSG00000100017.1,antisense | 210.9287253 | 0.426209188 | 0.000127428 | 0.001437064 |
| ENSMUST00000146177.7,Fmr1-004,retained_intron,ENSMUSG00000000838.17,protein_coding | 113.8074363 | -0.592532004 | 0.000127487 | 0.001437354 |
| ENSMUST00000128750.1,Usp46-003,processed_transcript,ENSMUSG00000054814.14,protein_coding | 396.0257707 | 0.352479162 | 0.000127643 | 0.001438737 |
| ENSMUST00000155824.1,Ppia-003,retained_intron,ENSMUSG00000071866.12,protein_coding | 1188.080533 | -0.210624553 | 0.000127898 | 0.001441231 |
| ENSMUST00000098816.9,Slc7a2-003,protein_coding,ENSMUSG00000031596.15,protein_coding | 512.2922699 | 0.471866865 | 0.000127994 | 0.001441939 |
| ENSMUST00000145653.7,2410002F23Rik-014,protein_coding,ENSMUSG00000045411.16,protein_coding | 192.5992724 | 0.456160341 | 0.000128056 | 0.001442257 |
| ENSMUST00000090924.12,Trim46-003,protein_coding,ENSMUSG00000042766.17,protein_coding | 831.6459258 | 0.310915275 | 0.00012818 | 0.00144327 |
| ENSMUST00000119527.7,Iffo1-001,protein_coding,ENSMUSG00000038271.17,protein_coding | 263.8422631 | -0.710758223 | 0.000128223 | 0.001443379 |
| ENSMUST00000211564.1,Gm10575-004,retained_intron,ENSMUSG00000073787.4,processed_transcript | 144.9220755 | -0.598999581 | 0.0001286 | 0.001447249 |
| ENSMUST00000121585.1,Gm15542-001,processed_pseudogene,ENSMUSG00000083396.1,processed_pseudogene | 497.3072184 | -0.486922861 | 0.000128717 | 0.001448189 |
| ENSMUST00000133119.1,Csnk2a1-005,protein_coding,ENSMUSG00000074698.10,protein_coding | 147.1941359 | 0.601965642 | 0.000128798 | 0.001448716 |
| ENSMUST00000198128.1,S100a6-002,protein_coding,ENSMUSG00000001025.8,protein_coding | 73699.21824 | 0.206944246 | 0.000129384 | 0.001454924 |
| ENSMUST00000002029.12,Emd-001,protein_coding,ENSMUSG00000001964.14,protein_coding | 8260.95241 | 0.186848929 | 0.000129417 | 0.001454924 |
| ENSMUST00000111294.7,Phf21a-004,protein_coding,ENSMUSG00000058318.15,protein_coding | 880.5312146 | 0.463432995 | 0.000129689 | 0.001457342 |
| ENSMUST00000040873.11,Srrt-201,protein_coding,ENSMUSG00000037364.11,protein_coding | 11065.27558 | -0.17902663 | 0.0001297 | 0.001457342 |
| ENSMUST00000182415.1,Gm9892-001,processed_pseudogene,ENSMUSG00000052825.6,processed_pseudogene | 293.4227364 | -0.464831726 | 0.000129764 | 0.00145768 |
| ENSMUST00000200541.4,Aldh2-006,protein_coding,ENSMUSG00000029455.14,protein_coding | 2703.157504 | 0.177772889 | 0.000130072 | 0.001460759 |
| ENSMUST00000210495.1,Snrnp70-015,retained_intron,ENSMUSG00000063511.11,protein_coding | 175.8034715 | 0.555578892 | 0.000130143 | 0.00146117 |
| ENSMUST00000025379.13,Dpysl3-002,protein_coding,ENSMUSG00000024501.19,protein_coding | 517.1890892 | 0.304798182 | 0.000130184 | 0.001461257 |
| ENSMUST00000204056.1,Uba3-005,nonsense_mediated_decay,ENSMUSG00000030061.16,protein_coding | 233.2122962 | -0.5343479 | 0.000130448 | 0.001463837 |
| ENSMUST00000049648.8,Kbtbd3-002,protein_coding,ENSMUSG00000025893.8,protein_coding | 42.61750086 | -0.713992861 | 0.00013064 | 0.001465608 |
| ENSMUST00000200305.1,Rps27-008,retained_intron,ENSMUSG00000090733.6,protein_coding | 483.9490378 | 0.375650621 | 0.000130912 | 0.001468273 |
| ENSMUST00000030769.5,Psmc2-001,protein_coding,ENSMUSG00000028932.7,protein_coding | 23425.25112 | 0.148415436 | 0.000131141 | 0.001470463 |
| ENSMUST00000040330.14,Cep44-001,protein_coding,ENSMUSG00000038215.15,protein_coding | 1295.38038 | 0.323459711 | 0.000131195 | 0.001470687 |
| ENSMUST00000036380.13,Atp6v0b-001,protein_coding,ENSMUSG00000033379.13,protein_coding | 7857.787167 | 0.176981409 | 0.000131298 | 0.001471451 |
| ENSMUST00000192057.5,Smg5-002,retained_intron,ENSMUSG00000001415.10,protein_coding | 830.0734438 | -0.338350891 | 0.000131637 | 0.001474819 |
| ENSMUST00000064848.5,Nup107-201,protein_coding,ENSMUSG00000052798.12,protein_coding | 20592.54455 | 0.167149852 | 0.000131667 | 0.001474819 |
| ENSMUST00000203813.1,Rpn1-003,retained_intron,ENSMUSG00000030062.7,protein_coding | 5983.031452 | 0.187478034 | 0.00013199 | 0.001478057 |
| ENSMUST00000102507.9,Capzb-002,protein_coding,ENSMUSG00000028745.18,protein_coding | 701.5032386 | -0.32922991 | 0.000132742 | 0.001486091 |
| ENSMUST00000101347.9,Mat2b-001,protein_coding,ENSMUSG00000042032.13,protein_coding | 3335.80207 | -0.203836956 | 0.000133087 | 0.001489569 |
| ENSMUST00000185466.1,Ndufa10-006,retained_intron,ENSMUSG00000026260.12,protein_coding | 2020.006683 | 0.227083196 | 0.000133246 | 0.001490572 |
| ENSMUST00000214758.1,Ilf3-010,protein_coding,ENSMUSG00000032178.14,protein_coding | 3050.254256 | 0.371148134 | 0.000133228 | 0.001490572 |
| ENSMUST00000176777.7,Unc13a-003,retained_intron,ENSMUSG00000034799.16,protein_coding | 71.12634602 | 0.628046411 | 0.000133523 | 0.001493284 |
| ENSMUST00000073471.12,Rpl26-001,protein_coding,ENSMUSG00000060938.14,protein_coding | 75551.39016 | 0.125553754 | 0.000133561 | 0.001493321 |
| ENSMUST00000033495.14,Pim2-001,protein_coding,ENSMUSG00000031155.15,protein_coding | 1079.928989 | -0.216111839 | 0.000133963 | 0.001497419 |
| ENSMUST00000102656.3,Foxo6-001,protein_coding,ENSMUSG00000052135.8,protein_coding | 1086.637264 | -0.292017498 | 0.000134044 | 0.001497935 |
| ENSMUST00000020959.7,Rnaseh1-201,protein_coding,ENSMUSG00000020630.8,protein_coding | 1710.682268 | 0.186350969 | 0.000134623 | 0.00150402 |
| ENSMUST00000062829.8,Hoxa6-001,protein_coding,ENSMUSG00000043219.9,protein_coding | 240.4487842 | -0.429185495 | 0.000134693 | 0.001504408 |
| ENSMUST00000124638.7,Setdb1-002,retained_intron,ENSMUSG00000015697.14,protein_coding | 1136.577262 | -0.220095467 | 0.000134809 | 0.001505318 |
| ENSMUST00000162681.1,Pam-011,protein_coding,ENSMUSG00000026335.16,protein_coding | 68.44240517 | -0.6751442 | 0.000134894 | 0.001505878 |
| ENSMUST00000115467.10,Tes-001,protein_coding,ENSMUSG00000029552.19,protein_coding | 29600.35632 | 0.170241499 | 0.000134986 | 0.001506511 |
| ENSMUST00000151863.7,B4galt3-008,protein_coding,ENSMUSG00000052423.14,protein_coding | 142.9122902 | 0.569709204 | 0.000135282 | 0.001509033 |
| ENSMUST00000146285.7,Sfpq-006,processed_transcript,ENSMUSG00000028820.13,protein_coding | 2204.166615 | 0.294190144 | 0.00013527 | 0.001509033 |
| ENSMUST00000169385.1,Rai14-202,protein_coding,ENSMUSG00000022246.13,protein_coding | 33828.89095 | 0.14197652 | 0.000135603 | 0.001512224 |
| ENSMUST00000058860.13,Usf2-001,protein_coding,ENSMUSG00000058239.13,protein_coding | 4855.064903 | 0.234759797 | 0.000135802 | 0.001514044 |
| ENSMUST00000102632.6,Sort1-001,protein_coding,ENSMUSG00000068747.14,protein_coding | 2588.363309 | -0.332724053 | 0.000135844 | 0.001514122 |
| ENSMUST00000108751.7,Mprip-003,protein_coding,ENSMUSG00000005417.17,protein_coding | 79008.33448 | 0.130036286 | 0.000136033 | 0.001515835 |

| | | | | |
|---|---|---|---|---|
| ENSMUST00000207548.1,Gm45133-001,TEC,ENSMUSG00000109274.1,TEC | 234.7180954 | -0.479738639 | 0.000136318 | 0.001518622 |
| ENSMUST00000131214.1,Sqstm1-007,retained_intron,ENSMUSG00000015837.15,protein_coding | 374.059916 | 0.456439122 | 0.000136551 | 0.001520822 |
| ENSMUST00000117394.1,Taok2-002,protein_coding,ENSMUSG00000059981.12,protein_coding | 3229.499361 | -0.190891363 | 0.000136606 | 0.001521044 |
| ENSMUST00000032879.14,Rab30-001,protein_coding,ENSMUSG00000030643.14,protein_coding | 250.3962476 | -0.543001622 | 0.000136646 | 0.001521096 |
| ENSMUST00000069456.10,Ncor1-010,protein_coding,ENSMUSG00000018501.17,protein_coding | 802.2221125 | -0.356707085 | 0.000136689 | 0.001521183 |
| ENSMUST00000088646.11,Wnk1-021,protein_coding,ENSMUSG00000045962.16,protein_coding | 7604.221641 | 0.381924549 | 0.000136746 | 0.00152142 |
| ENSMUST00000047179.5,Zfp598-201,protein_coding,ENSMUSG00000041130.9,protein_coding | 12558.5254 | 0.257818069 | 0.000136999 | 0.001523845 |
| ENSMUST00000176216.8,Cux1-004,protein_coding,ENSMUSG00000029705.17,protein_coding | 3013.471987 | 0.421062437 | 0.000137775 | 0.00153208 |
| ENSMUST00000079496.8,Ube2s-001,protein_coding,ENSMUSG00000060860.8,protein_coding | 53048.13548 | 0.145262577 | 0.000138043 | 0.001534661 |
| ENSMUST00000032749.11,Vps33b-001,protein_coding,ENSMUSG00000030534.13,protein_coding | 936.0499451 | -0.269595416 | 0.000139332 | 0.001547385 |
| ENSMUST00000035034.9,Mrps22-001,protein_coding,ENSMUSG00000032459.9,protein_coding | 4113.776335 | 0.186819902 | 0.000139286 | 0.001547385 |
| ENSMUST00000108657.3,Trp53-002,protein_coding,ENSMUSG00000059552.13,protein_coding | 4712.863723 | 0.19978467 | 0.000139326 | 0.001547385 |
| ENSMUST00000025800.14,Pip5k1b-001,protein_coding,ENSMUSG00000024867.14,protein_coding | 547.9580737 | 0.318641757 | 0.000139247 | 0.001547385 |
| ENSMUST00000115600.3,Gnb1l-008,nonsense_mediated_decay,ENSMUSG00000000884.16,protein_coding | 853.5899724 | 0.360367677 | 0.000139386 | 0.001547589 |
| ENSMUST00000032485.6,Mrpl51-001,protein_coding,ENSMUSG00000030335.6,protein_coding | 2501.634399 | -0.237147974 | 0.000139624 | 0.001549834 |
| ENSMUST00000054956.14,Stambpl1-001,protein_coding,ENSMUSG00000024776.17,protein_coding | 2753.447829 | 0.288753638 | 0.000140177 | 0.001555575 |
| ENSMUST00000117956.1,Zbtb7a-004,protein_coding,ENSMUSG00000035011.15,protein_coding | 1567.132646 | 0.247997233 | 0.000140344 | 0.001557023 |
| ENSMUST00000135232.7,Faf2-004,protein_coding,ENSMUSG00000025873.15,protein_coding | 246.137615 | 0.622331652 | 0.00014074 | 0.001561017 |
| ENSMUST00000192615.5,Gnai2-003,protein_coding,ENSMUSG00000032562.13,protein_coding | 9194.533079 | -0.180300826 | 0.000140933 | 0.001562754 |
| ENSMUST00000025181.16,H2-K1-001,protein_coding,ENSMUSG00000061232.15,protein_coding | 5198.83779 | 0.223271071 | 0.000141403 | 0.001567559 |
| ENSMUST00000210074.1,Arntl-003,protein_coding,ENSMUSG00000055116.8,protein_coding | 40.33372643 | 0.704424373 | 0.000141581 | 0.001569127 |
| ENSMUST00000044443.14,Ank2-001,protein_coding,ENSMUSG00000032826.17,protein_coding | 5463.25273 | 0.304327525 | 0.000141692 | 0.001569955 |
| ENSMUST00000126716.8,Srsf11-008,nonsense_mediated_decay,ENSMUSG00000055436.18,protein_coding | 165.0554945 | 0.570715731 | 0.000142031 | 0.001572905 |
| ENSMUST00000074957.4,Bud13-001,protein_coding,ENSMUSG00000032077.5,protein_coding | 2925.589983 | 0.160095424 | 0.000142029 | 0.001572905 |
| ENSMUST00000020446.10,Tmbim4-001,protein_coding,ENSMUSG00000020225.10,protein_coding | 2353.600289 | -0.189886731 | 0.000142113 | 0.001573399 |
| ENSMUST00000181108.1,Cbx3-ps6-001,processed_pseudogene,ENSMUSG00000057886.5,processed_pseudogene | 474.4394679 | -0.37447713 | 0.000142176 | 0.001573702 |
| ENSMUST00000134846.7,Cenpa-004,nonsense_mediated_decay,ENSMUSG00000029177.9,protein_coding | 263.9526816 | 0.599242106 | 0.000142234 | 0.001573935 |
| ENSMUST00000057610.6,Daam2-201,protein_coding,ENSMUSG00000040260.7,protein_coding | 17905.4259 | -0.174968406 | 0.000142296 | 0.001574211 |
| ENSMUST00000034277.13,Emc8-001,protein_coding,ENSMUSG00000031819.13,protein_coding | 2805.12708 | -0.267612287 | 0.00014302 | 0.001581199 |
| ENSMUST00000211187.1,Trak1-011,protein_coding,ENSMUSG00000032536.11,protein_coding | 1780.170511 | 0.216669993 | 0.000143046 | 0.001581199 |
| ENSMUST00000152665.1,Rpl10a-003,retained_intron,ENSMUSG00000037805.14,protein_coding | 489.7827407 | 0.382367303 | 0.000143035 | 0.001581199 |
| ENSMUST00000152463.7,Sat1-002,nonsense_mediated_decay,ENSMUSG00000025283.15,protein_coding | 86.21824198 | 0.690407704 | 0.000143074 | 0.001581199 |
| ENSMUST00000063882.11,Dcun1d4-001,protein_coding,ENSMUSG00000051674.14,protein_coding | 1471.55283 | -0.209556 | 0.000143227 | 0.001582485 |
| ENSMUST00000113824.7,Clpx-201,protein_coding,ENSMUSG00000015357.10,protein_coding | 684.090405 | -0.512098428 | 0.000143563 | 0.001585788 |
| ENSMUST00000210467.1,Ldha-012,protein_coding,ENSMUSG00000063229.15,protein_coding | 145.658196 | 0.705596173 | 0.000143808 | 0.001588085 |
| ENSMUST00000003029.13,Timm44-001,protein_coding,ENSMUSG00000002949.15,protein_coding | 8337.568121 | 0.132895998 | 0.000143949 | 0.001588875 |
| ENSMUST00000076505.3,Pyroxd2-201,protein_coding,ENSMUSG00000060224.3,protein_coding | 351.867939 | 0.354591768 | 0.000143953 | 0.001588875 |
| ENSMUST00000178893.7,Crlf3-007,nonsense_mediated_decay,ENSMUSG00000017561.16,protein_coding | 181.2655968 | 0.701087204 | 0.000144062 | 0.001589665 |
| ENSMUST00000041686.9,Nudt22-001,protein_coding,ENSMUSG00000037349.9,protein_coding | 1287.036408 | -0.262432196 | 0.000144437 | 0.001593402 |
| ENSMUST00000112934.7,Rc3h2-001,protein_coding,ENSMUSG00000075376.10,protein_coding | 3022.424387 | -0.546060962 | 0.000144542 | 0.001594147 |
| ENSMUST00000167195.7,Slc29a1-016,protein_coding,ENSMUSG00000023942.15,protein_coding | 115.1100375 | 0.611775598 | 0.000144707 | 0.001595561 |
| ENSMUST00000072889.11,Carmil1-001,protein_coding,ENSMUSG00000021338.17,protein_coding | 1346.128006 | 0.195799196 | 0.000145744 | 0.001606578 |
| ENSMUST00000094894.3,Trabd2b-001,protein_coding,ENSMUSG00000070867.4,protein_coding | 966.2705976 | -0.252580528 | 0.000145971 | 0.001608672 |
| ENSMUST00000051145.14,Wdr47-001,protein_coding,ENSMUSG00000040389.15,protein_coding | 1036.566216 | -0.313737349 | 0.000146189 | 0.001610381 |
| ENSMUST00000146838.2,Dlx4-002,processed_transcript,ENSMUSG00000020871.8,protein_coding | 259.3913543 | -0.388955004 | 0.000146201 | 0.001610381 |
| ENSMUST00000200119.2,Usp30-004,protein_coding,ENSMUSG00000029592.11,protein_coding | 1235.678289 | 0.178271519 | 0.000147094 | 0.001619393 |
| ENSMUST00000142424.1,Mrpl3-004,nonsense_mediated_decay,ENSMUSG00000032563.16,protein_coding | 5552.339443 | 0.194614783 | 0.000147087 | 0.001619393 |
| ENSMUST00000142512.7,Cald1-007,protein_coding,ENSMUSG00000029761.16,protein_coding | 2165.03959 | -0.363841908 | 0.000147635 | 0.001624925 |
| ENSMUST00000025439.3,Me2-201,protein_coding,ENSMUSG00000024556.3,protein_coding | 12426.94234 | 0.178579804 | 0.000147817 | 0.001626515 |
| ENSMUST00000155986.1,Slc25a4-002,retained_intron,ENSMUSG00000031633.4,protein_coding | 1929.88084 | -0.40968932 | 0.000147863 | 0.001626602 |
| ENSMUST00000176438.8,Zeb2-001,protein_coding,ENSMUSG00000026872.17,protein_coding | 1257.421579 | -0.407806545 | 0.000148505 | 0.001632417 |
| ENSMUST00000148871.1,Ptpn11-003,processed_transcript,ENSMUSG00000043733.14,protein_coding | 494.6489139 | 0.580364973 | 0.000148468 | 0.001632417 |
| ENSMUST00000049353.8,Zfp692-001,protein_coding,ENSMUSG00000037243.17,protein_coding | 959.2495762 | 0.316591954 | 0.000148433 | 0.001632417 |
| ENSMUST00000107920.9,Srsf1-006,protein_coding,ENSMUSG00000018379.17,protein_coding | 6938.956545 | -0.221322055 | 0.000149457 | 0.001642463 |
| ENSMUST00000041614.14,Ttc13-001,protein_coding,ENSMUSG00000037300.18,protein_coding | 1242.818967 | 0.327548112 | 0.000149686 | 0.001644559 |
| ENSMUST00000081940.10,Uba52-005,protein_coding,ENSMUSG00000090137.8,protein_coding | 21604.20927 | 0.18581959 | 0.000149817 | 0.001645583 |
| ENSMUST00000078803.3,Tonsl-201,protein_coding,ENSMUSG00000059323.13,protein_coding | 6596.732589 | 0.16288388 | 0.000150261 | 0.001650031 |
| ENSMUST00000133077.7,Ctsa-005,retained_intron,ENSMUSG00000017760.15,protein_coding | 131.472278 | -0.667969543 | 0.000150434 | 0.001650825 |
| ENSMUST00000095754.9,Tia1-010,protein_coding,ENSMUSG00000071337.11,protein_coding | 248.161308 | 0.696130856 | 0.000150452 | 0.001650825 |
| ENSMUST00000110316.2,Galnt7-002,protein_coding,ENSMUSG00000031608.13,protein_coding | 625.5104116 | -0.434690564 | 0.000150375 | 0.001650825 |
| ENSMUST00000025170.9,Wdr46-001,protein_coding,ENSMUSG00000024312.9,protein_coding | 12469.41628 | -0.171262417 | 0.000150486 | 0.001650825 |
| ENSMUST00000027052.12,Stau2-003,protein_coding,ENSMUSG00000025920.19,protein_coding | 176.4745291 | 0.638294332 | 0.00015057 | 0.001651316 |
| ENSMUST00000144493.1,Eif4a2-020,retained_intron,ENSMUSG00000022884.14,protein_coding | 381.5304798 | -0.510952594 | 0.000151466 | 0.001660725 |
| ENSMUST00000102781.9,Jak1-001,protein_coding,ENSMUSG00000028530.14,protein_coding | 18854.93991 | 0.128171538 | 0.000151542 | 0.00166113 |
| ENSMUST00000109485.8,Ralgapb-009,protein_coding,ENSMUSG00000027652.15,protein_coding | 2144.336288 | 0.3898728 | 0.000151806 | 0.001663608 |
| ENSMUST00000058437.13,Rpl7-001,protein_coding,ENSMUSG00000043716.13,protein_coding | 72150.28786 | 0.121184576 | 0.000151865 | 0.00166383 |
| ENSMUST00000030460.14,Gpbp1l1-001,protein_coding,ENSMUSG00000034042.16,protein_coding | 3776.338278 | 0.162716795 | 0.000151985 | 0.001664722 |
| ENSMUST00000106908.8,Pde4b-001,protein_coding,ENSMUSG00000028525.16,protein_coding | 1612.811927 | -0.373467903 | 0.000152189 | 0.001666526 |
| ENSMUST00000216914.1,Higd1a-003,protein_coding,ENSMUSG00000038412.8,protein_coding | 177.6931333 | 0.527925996 | 0.000152554 | 0.001670104 |
| ENSMUST00000188485.1,Gm29650-001,lincRNA,ENSMUSG00000099876.1,lincRNA | 23.1156408 | -0.706480142 | 0.00015286 | 0.001673026 |
| ENSMUST00000042057.11,Midn-001,protein_coding,ENSMUSG00000035621.13,protein_coding | 8278.07518 | -0.259828842 | 0.000153616 | 0.001680868 |
| ENSMUST00000165389.7,Bnip2-201,protein_coding,ENSMUSG00000011958.17,protein_coding | 1690.670177 | 0.293678987 | 0.00015432 | 0.001687889 |
| ENSMUST00000124870.7,Fance-003,retained_intron,ENSMUSG00000007570.15,protein_coding | 139.2898326 | 0.629747647 | 0.000154336 | 0.001687889 |
| ENSMUST00000027748.7,Rgs16-001,protein_coding,ENSMUSG00000026475.7,protein_coding | 192.4031276 | -0.419119479 | 0.000154515 | 0.00168935 |
| ENSMUST00000106110.9,Wdr45b-004,protein_coding,ENSMUSG00000025173.13,protein_coding | 125.6643251 | 0.70124452 | 0.000154548 | 0.00168935 |
| ENSMUST00000054536.10,Pqlc3-001,protein_coding,ENSMUSG00000045679.13,protein_coding | 1112.768001 | 0.197130981 | 0.000154588 | 0.001689357 |
| ENSMUST00000034761.14,Ice2-001,protein_coding,ENSMUSG00000032235.15,protein_coding | 1068.745258 | -0.315072395 | 0.00015482 | 0.001691463 |
| ENSMUST00000141158.7,Adgrl1-001,protein_coding,ENSMUSG00000013033.16,protein_coding | 2740.371196 | -0.160059154 | 0.000155456 | 0.001697982 |
| ENSMUST00000032491.14,Tnfrsf1a-001,protein_coding,ENSMUSG00000030341.17,protein_coding | 14512.64599 | 0.128350331 | 0.000155962 | 0.001703074 |
| ENSMUST00000027693.7,Rab29-001,protein_coding,ENSMUSG00000026433.13,protein_coding | 268.0526033 | -0.448335613 | 0.000156214 | 0.001705145 |
| ENSMUST00000122292.1,Gm9816-001,processed_pseudogene,ENSMUSG00000085783.3,processed_pseudogene | 215.8035716 | -0.572716473 | 0.00015623 | 0.001705145 |
| ENSMUST00000020970.8,Rsad2-001,protein_coding,ENSMUSG00000020641.15,protein_coding | 823.9551855 | 0.441218988 | 0.000156353 | 0.00170605 |
| ENSMUST00000116514.3,Wbp11-001,protein_coding,ENSMUSG00000030216.12,protein_coding | 32628.57103 | 0.144445119 | 0.000156681 | 0.001709196 |
| ENSMUST00000103099.7,Nbr1-001,protein_coding,ENSMUSG00000017119.19,protein_coding | 624.3843444 | -0.438905403 | 0.000156841 | 0.001710506 |
| ENSMUST00000171229.7,Eci2-001,protein_coding,ENSMUSG00000021417.14,protein_coding | 1922.459902 | 0.226722118 | 0.000156997 | 0.001711773 |
| ENSMUST00000020794.5,Ska2-001,protein_coding,ENSMUSG00000020492.11,protein_coding | 6121.281663 | -0.299479359 | 0.000157043 | 0.001711841 |
| ENSMUST00000161240.3,Galnt4-001,protein_coding,ENSMUSG00000090035.3,protein_coding | 1893.425985 | -0.23825257 | 0.000157127 | 0.001712328 |
| ENSMUST00000048937.5,Leo1-001,protein_coding,ENSMUSG00000042487.6,protein_coding | 4328.361983 | 0.175258248 | 0.00015733 | 0.001714107 |
| ENSMUST00000144843.7,Slain2-001,protein_coding,ENSMUSG00000036087.18,protein_coding | 5218.898764 | 0.279556815 | 0.000157817 | 0.001718973 |
| ENSMUST00000143977.7,Camsap1-003,nonsense_mediated_decay,ENSMUSG00000026933.17,protein_coding | 4283.808991 | 0.149982725 | 0.000159044 | 0.001731907 |
| ENSMUST00000064391.11,Cpne8-001,protein_coding,ENSMUSG00000052560.15,protein_coding | 1608.105572 | -0.257767699 | 0.00015954 | 0.001736868 |
| ENSMUST00000143180.7,Fbxl2-002,nonsense_mediated_decay,ENSMUSG00000032507.15,protein_coding | 744.4855823 | 0.323488401 | 0.000159664 | 0.001737771 |

| | | | | |
|---|---|---|---|---|
| ENSMUST00000136646.1,Rere-007,protein_coding,ENSMUSG00000039852.16,protein_coding | 271.8714915 | 0.52193565 | 0.000160197 | 0.00174313 |
| ENSMUST00000073067.10,Slc10a3-002,protein_coding,ENSMUSG00000032806.13,protein_coding | 929.8003624 | -0.334654262 | 0.000160456 | 0.00174551 |
| ENSMUST00000092155.11,Col11a1-001,protein_coding,ENSMUSG00000027966.20,protein_coding | 2078.706702 | -0.311221054 | 0.000160569 | 0.001746303 |
| ENSMUST00000141230.7,Iffo1-004,retained_intron,ENSMUSG00000038271.17,protein_coding | 97.86252008 | -0.622316669 | 0.000161682 | 0.001757511 |
| ENSMUST00000047379.14,Ptger4-001,protein_coding,ENSMUSG00000039942.15,protein_coding | 301.511419 | -0.525232991 | 0.000161673 | 0.001757511 |
| ENSMUST00000014321.4,Tvp23b-001,protein_coding,ENSMUSG00000014177.10,protein_coding | 2562.850288 | 0.211496156 | 0.000161887 | 0.001759296 |
| ENSMUST00000206388.1,Lsm14a-006,nonsense_mediated_decay,ENSMUSG00000066568.12,protein_coding | 1885.427658 | 0.253380811 | 0.000162557 | 0.00176614 |
| ENSMUST00000094762.9,Relb-001,protein_coding,ENSMUSG00000002983.17,protein_coding | 1258.361264 | 0.389598795 | 0.000164191 | 0.001783435 |
| ENSMUST00000211933.1,Alkbh8-001,protein_coding,ENSMUSG00000025899.14,protein_coding | 645.0474384 | -0.566795124 | 0.000164572 | 0.001787123 |
| ENSMUST00000040844.15,Ankrd27-001,protein_coding,ENSMUSG00000034867.16,protein_coding | 5714.961236 | 0.179263634 | 0.000164803 | 0.001789181 |
| ENSMUST00000027688.14,Rassf5-001,protein_coding,ENSMUSG00000026430.16,protein_coding | 448.396135 | 0.436116695 | 0.000165117 | 0.001792145 |
| ENSMUST00000041761.6,Prss23-002,protein_coding,ENSMUSG00000039405.7,protein_coding | 1481.214885 | -0.316431183 | 0.000165246 | 0.001793086 |
| ENSMUST00000090824.11,Zak-001,protein_coding,ENSMUSG00000004085.14,protein_coding | 1858.39476 | 0.283226789 | 0.000165553 | 0.001795963 |
| ENSMUST00000108572.1,Zfp579-201,protein_coding,ENSMUSG00000051550.10,protein_coding | 220.2077471 | 0.665762372 | 0.000166335 | 0.001803993 |
| ENSMUST00000085555.6,Diexf-002,protein_coding,ENSMUSG00000016181.9,protein_coding | 3226.649523 | 0.383001948 | 0.000166521 | 0.001805562 |
| ENSMUST00000212426.1,Mcm5-002,protein_coding,ENSMUSG00000005410.6,protein_coding | 33188.16744 | 0.169232432 | 0.000166795 | 0.001808074 |
| ENSMUST00000012627.4,Rpa3-001,protein_coding,ENSMUSG00000012483.4,protein_coding | 4294.575707 | 0.194064166 | 0.000166848 | 0.001808195 |
| ENSMUST00000140267.8,Tmem134-001,processed_transcript,ENSMUSG00000024845.17,processed_transcript | 3639.126416 | 0.250850242 | 0.000167474 | 0.001814517 |
| ENSMUST00000019803.8,Ccdc12-001,protein_coding,ENSMUSG00000019659.8,protein_coding | 3169.374826 | -0.169012156 | 0.000167764 | 0.001817207 |
| ENSMUST00000041768.6,Akr1c14-201,protein_coding,ENSMUSG00000033715.14,protein_coding | 77.72676693 | -0.656697954 | 0.000168112 | 0.00182052 |
| ENSMUST00000117536.7,Paics-002,protein_coding,ENSMUSG00000029247.17,protein_coding | 24795.13329 | 0.150188732 | 0.00016829 | 0.001821985 |
| ENSMUST00000070653.12,Slc16a3-001,protein_coding,ENSMUSG00000025161.16,protein_coding | 24176.62412 | 0.150623352 | 0.000168462 | 0.001823392 |
| ENSMUST00000100036.9,Sgk1-201,protein_coding,ENSMUSG00000019970.15,protein_coding | 269.9774036 | -0.661955045 | 0.00016863 | 0.001824752 |
| ENSMUST00000044833.8,Oas3-001,protein_coding,ENSMUSG00000032661.9,protein_coding | 311.9539437 | 0.357827317 | 0.000168768 | 0.001825512 |
| ENSMUST00000044423.3,Brip1-001,protein_coding,ENSMUSG00000034329.16,protein_coding | 2800.127125 | -0.330924442 | 0.000168805 | 0.001825512 |
| ENSMUST00000102745.9,Tnfaip2-001,protein_coding,ENSMUSG00000021281.15,protein_coding | 29875.767 | 0.187883987 | 0.000168828 | 0.001825512 |
| ENSMUST00000111785.8,Ssfa2-001,protein_coding,ENSMUSG00000027007.16,protein_coding | 8597.769005 | 0.146520645 | 0.000169019 | 0.001827121 |
| ENSMUST00000116343.2,Erf-002,protein_coding,ENSMUSG00000040857.15,protein_coding | 610.434266 | -0.284341712 | 0.000170195 | 0.001838482 |
| ENSMUST00000102602.7,Trappc1-002,protein_coding,ENSMUSG00000049299.14,protein_coding | 1778.137362 | -0.269083709 | 0.000170198 | 0.001838482 |
| ENSMUST00000047131.15,Ipo4-001,protein_coding,ENSMUSG00000002319.15,protein_coding | 31438.11324 | -0.156780333 | 0.000170134 | 0.001838482 |
| ENSMUST00000097666.3,Cab39-002,protein_coding,ENSMUSG00000036707.16,protein_coding | 286.0834995 | 0.5334129 | 0.000170311 | 0.001839235 |
| ENSMUST00000137880.1,Cdca7-003,processed_transcript,ENSMUSG00000055612.15,protein_coding | 126.820436 | 0.640593256 | 0.000170373 | 0.001839449 |
| ENSMUST00000191127.6,Ly6e-008,protein_coding,ENSMUSG00000022587.14,protein_coding | 1985.327027 | 0.190521829 | 0.000170437 | 0.001839674 |
| ENSMUST00000188027.1,Soat1-005,protein_coding,ENSMUSG00000026600.12,protein_coding | 888.9669757 | 0.261174338 | 0.000171107 | 0.001846449 |
| ENSMUST00000163441.7,Tex264-001,protein_coding,ENSMUSG00000040813.16,protein_coding | 1044.710286 | 0.263861267 | 0.000171396 | 0.001848633 |
| ENSMUST00000131862.7,Etv4-010,processed_transcript,ENSMUSG00000017724.14,protein_coding | 251.3925721 | 0.540781918 | 0.000171354 | 0.001848633 |
| ENSMUST00000186745.1,Ivns1abp-004,nonsense_mediated_decay,ENSMUSG00000023150.14,protein_coding | 1380.818199 | -0.402855847 | 0.000171873 | 0.00185332 |
| ENSMUST00000053456.10,Sept6-002,protein_coding,ENSMUSG00000050379.15,protein_coding | 2691.72768 | 0.180801797 | 0.000172184 | 0.001856209 |
| ENSMUST00000102884.9,Egfr-002,protein_coding,ENSMUSG00000020122.16,protein_coding | 1219.94248 | 0.291419266 | 0.000172238 | 0.001856328 |
| ENSMUST00000146574.1,Bckdk-002,retained_intron,ENSMUSG00000030802.14,protein_coding | 572.1017318 | 0.300674543 | 0.000172989 | 0.001863953 |
| ENSMUST00000137717.1,Copg1-002,retained_intron,ENSMUSG00000030058.17,protein_coding | 520.6507541 | -0.406664798 | 0.000173666 | 0.001870775 |
| ENSMUST00000028928.7,Gzf1-001,protein_coding,ENSMUSG00000027439.9,protein_coding | 5249.748841 | 0.112837541 | 0.000173739 | 0.001871098 |
| ENSMUST00000212378.1,Rpl18a-002,protein_coding,ENSMUSG00000045128.9,protein_coding | 2913.685611 | 0.542004535 | 0.000174407 | 0.001877821 |
| ENSMUST00000202218.3,Gtf3c2-005,retained_intron,ENSMUSG00000106864.3,protein_coding | 623.1284726 | 0.309364362 | 0.000174761 | 0.001881033 |
| ENSMUST00000129270.7,Slc25a1-006,retained_intron,ENSMUSG00000003528.14,protein_coding | 1001.607013 | 0.455149494 | 0.000174793 | 0.001881033 |
| ENSMUST00000033999.7,Frg1-001,protein_coding,ENSMUSG00000031590.8,protein_coding | 4018.554182 | 0.177804672 | 0.000175091 | 0.001883775 |
| ENSMUST00000084124.6,Pgd-001,protein_coding,ENSMUSG00000028961.15,protein_coding | 37679.87054 | -0.18598048 | 0.000175469 | 0.001887364 |
| ENSMUST00000196304.1,Gatad1-004,protein_coding,ENSMUSG00000007415.15,protein_coding | 4451.602884 | 0.196554151 | 0.000175772 | 0.001890155 |
| ENSMUST00000190051.1,Trf-011,retained_intron,ENSMUSG00000032554.15,protein_coding | 406.0679718 | -0.332545803 | 0.000175942 | 0.001891508 |
| ENSMUST00000069805.13,Atp8b2-001,protein_coding,ENSMUSG00000060671.12,protein_coding | 2911.036203 | -0.38046353 | 0.000177115 | 0.001903645 |
| ENSMUST00000028600.13,Hipk3-001,protein_coding,ENSMUSG00000027177.14,protein_coding | 6250.89652 | -0.378006254 | 0.000177324 | 0.001905418 |
| ENSMUST00000102694.3,Ptgfrn-001,protein_coding,ENSMUSG00000027864.9,protein_coding | 21314.87379 | 0.096433902 | 0.000178919 | 0.001922073 |
| ENSMUST00000169333.1,Gpr107-005,retained_intron,ENSMUSG00000000194.13,protein_coding | 97.33828103 | 0.621003852 | 0.000179467 | 0.001927476 |
| ENSMUST00000148159.2,Tnfaip8-001,protein_coding,ENSMUSG00000062210.13,protein_coding | 1094.101814 | -0.336799475 | 0.000179591 | 0.001928325 |
| ENSMUST00000203437.1,Gm43890-001,TEC,ENSMUSG00000108053.1,TEC | 329.632566 | 0.481900425 | 0.000180033 | 0.001932593 |
| ENSMUST00000029270.9,Ccna2-001,protein_coding,ENSMUSG00000027715.9,protein_coding | 33451.11713 | -0.141307968 | 0.000180405 | 0.001936105 |
| ENSMUST00000143514.2,Arfip1-003,protein_coding,ENSMUSG00000074513.9,protein_coding | 1592.135755 | 0.240607087 | 0.000180829 | 0.001940171 |
| ENSMUST00000033418.7,Il13ra1-001,protein_coding,ENSMUSG00000017057.9,protein_coding | 1407.994235 | -0.356683621 | 0.000180957 | 0.001941065 |
| ENSMUST00000031250.13,Nudt9-001,protein_coding,ENSMUSG00000029310.13,protein_coding | 5721.408705 | 0.176321941 | 0.00018153 | 0.001946716 |
| ENSMUST00000175783.1,Tshz1-002,processed_transcript,ENSMUSG00000046982.10,protein_coding | 1669.962169 | -0.265078736 | 0.000181602 | 0.001947012 |
| ENSMUST00000015003.9,E2f4-001,protein_coding,ENSMUSG00000014859.9,protein_coding | 14669.61582 | 0.125636268 | 0.00018184 | 0.001949077 |
| ENSMUST00000201207.1,Ppp1cb-003,processed_transcript,ENSMUSG00000014956.15,protein_coding | 466.7405225 | -0.408986116 | 0.000181922 | 0.001949466 |
| ENSMUST00000044475.4,Ogt-001,protein_coding,ENSMUSG00000034160.13,protein_coding | 8056.35445 | -0.203834046 | 0.000182064 | 0.001950509 |
| ENSMUST00000099513.7,Hcn2-001,protein_coding,ENSMUSG00000020331.9,protein_coding | 579.7476329 | -0.261438264 | 0.000182257 | 0.001952089 |
| ENSMUST00000120194.1,Fam104a-001,protein_coding,ENSMUSG00000041629.7,protein_coding | 9367.635774 | 0.175711663 | 0.000182333 | 0.001952414 |
| ENSMUST00000135603.1,Rprd1b-007,protein_coding,ENSMUSG00000027651.16,protein_coding | 21.91450283 | 0.581930382 | 0.00018266 | 0.001955431 |
| ENSMUST00000049836.13,Specc1-001,protein_coding,ENSMUSG00000042331.13,protein_coding | 897.6320767 | -0.277733203 | 0.000182893 | 0.001957438 |
| ENSMUST00000141682.1,Glis2-003,protein_coding,ENSMUSG00000014303.13,protein_coding | 33.56925387 | 0.690546638 | 0.000182998 | 0.001958069 |
| ENSMUST00000079773.12,Csnk1g2-201,protein_coding,ENSMUSG00000003345.15,protein_coding | 9884.197833 | 0.182646707 | 0.00018337 | 0.001961565 |
| ENSMUST00000146441.7,Asph-004,protein_coding,ENSMUSG00000028207.18,protein_coding | 154.0595205 | -0.503970592 | 0.000183524 | 0.001962728 |
| ENSMUST00000102711.8,Gemin5-002,protein_coding,ENSMUSG00000037275.14,protein_coding | 7083.719786 | 0.216629275 | 0.000184581 | 0.001973538 |
| ENSMUST00000088480.3,Rpl15-ps3-001,processed_pseudogene,ENSMUSG00000061167.9,processed_pseudogene | 1659.816222 | -0.379988057 | 0.000185007 | 0.001977114 |
| ENSMUST00000179301.7,Tnpo1-003,protein_coding,ENSMUSG00000009470.16,protein_coding | 8278.762581 | 0.143136265 | 0.000184994 | 0.001977114 |
| ENSMUST00000114863.8,D17Wsu92e-001,protein_coding,ENSMUSG00000056692.12,protein_coding | 19305.70479 | 0.16289989 | 0.00018526 | 0.001979328 |
| ENSMUST00000123533.7,Mrps2-003,processed_transcript,ENSMUSG00000035772.11,protein_coding | 285.5221697 | 0.422580743 | 0.000185454 | 0.001980909 |
| ENSMUST00000141355.2,Aldoa-004,protein_coding,ENSMUSG00000030695.14,protein_coding | 808.1137779 | 0.391399907 | 0.000185586 | 0.001981544 |
| ENSMUST00000174281.8,Bag6-005,protein_coding,ENSMUSG00000024392.17,protein_coding | 5345.031263 | -0.272101928 | 0.000185606 | 0.001981544 |
| ENSMUST00000183076.1,Gm5345-001,processed_pseudogene,ENSMUSG00000098050.1,processed_pseudogene | 117.6535021 | 0.495806008 | 0.0001857 | 0.001982019 |
| ENSMUST00000130572.1,Cobl-002,retained_intron,ENSMUSG00000020173.17,protein_coding | 20.87689377 | -0.697750915 | 0.000185742 | 0.001982019 |
| ENSMUST00000042857.13,Satb2-002,protein_coding,ENSMUSG00000038331.15,protein_coding | 555.7109804 | -0.383520943 | 0.000185898 | 0.001983194 |
| ENSMUST00000171237.7,Cox16-007,processed_transcript,ENSMUSG00000091803.7,protein_coding | 405.4519233 | -0.365891261 | 0.000186272 | 0.001986684 |
| ENSMUST00000166735.7,Papola-007,protein_coding,ENSMUSG00000021111.15,protein_coding | 956.2401329 | 0.448812797 | 0.000186639 | 0.001990108 |
| ENSMUST00000102593.10,Eif3i-001,protein_coding,ENSMUSG00000028798.16,protein_coding | 26616.68923 | 0.188250609 | 0.000186743 | 0.001990724 |
| ENSMUST00000186811.1,Atg4b-011,nonsense_mediated_decay,ENSMUSG00000026280.17,protein_coding | 57.52237432 | 0.672589598 | 0.00018776 | 0.002001074 |
| ENSMUST00000056710.9,Aph1a-002,protein_coding,ENSMUSG00000015750.15,protein_coding | 4668.584321 | 0.247057838 | 0.00018813 | 0.002004515 |
| ENSMUST00000103079.3,Cttn-001,protein_coding,ENSMUSG00000031078.15,protein_coding | 25445.21993 | 0.108777275 | 0.000188232 | 0.002004642 |
| ENSMUST00000062213.11,Sfxn3-201,protein_coding,ENSMUSG00000025212.16,protein_coding | 6182.271019 | -0.148117879 | 0.000188235 | 0.002004642 |
| ENSMUST00000211646.1,Rab11fip1-002,processed_transcript,ENSMUSG00000031488.14,protein_coding | 29.77849266 | 0.696310126 | 0.000188902 | 0.002010927 |
| ENSMUST00000067384.4,Rhob-201,protein_coding,ENSMUSG00000054364.4,protein_coding | 5132.90514 | -0.275573278 | 0.000188919 | 0.002010927 |
| ENSMUST00000193987.1,Gpx1-003,protein_coding,ENSMUSG00000063856.7,protein_coding | 7061.735568 | 0.194511807 | 0.000189055 | 0.002011382 |
| ENSMUST00000020208.4,Fgd6-001,protein_coding,ENSMUSG00000020021.4,protein_coding | 3358.807289 | 0.160780881 | 0.000189038 | 0.002011382 |
| ENSMUST00000034567.3,Dlat-001,protein_coding,ENSMUSG00000000168.9,protein_coding | 9457.358896 | 0.143487861 | 0.000189226 | 0.002012711 |

| | | | | |
|---|---|---|---|---|
| ENSMUST00000150802.7,Zzz3-006,processed_transcript,ENSMUSG00000039068.16,protein_coding | 119.2452209 | 0.636280522 | 0.000190934 | 0.00203037 |
| ENSMUST00000092033.2,Rbms2-201,protein_coding,ENSMUSG00000040043.15,protein_coding | 14269.22207 | 0.160117829 | 0.000191444 | 0.00203529 |
| ENSMUST00000123292.1,Dynll1-005,processed_transcript,ENSMUSG00000009013.5,protein_coding | 5053.250554 | -0.287188401 | 0.000192131 | 0.002042091 |
| ENSMUST00000152288.1,Emd-006,nonsense_mediated_decay,ENSMUSG00000001964.14,protein_coding | 1907.002509 | 0.323305686 | 0.000192638 | 0.002046977 |
| ENSMUST00000057416.6,Tmem251-201,protein_coding,ENSMUSG00000046675.6,protein_coding | 909.5356122 | -0.250403691 | 0.000192727 | 0.002047418 |
| ENSMUST00000021514.8,Ppm1a-201,protein_coding,ENSMUSG00000021096.10,protein_coding | 12449.58887 | 0.200664923 | 0.000192818 | 0.002047877 |
| ENSMUST00000015950.11,Qdpr-001,protein_coding,ENSMUSG00000015806.12,protein_coding | 5674.044661 | 0.193841605 | 0.000193066 | 0.002048492 |
| ENSMUST00000087324.6,Pgm1-001,protein_coding,ENSMUSG00000029171.12,protein_coding | 6787.491149 | 0.197183705 | 0.000193025 | 0.002048492 |
| ENSMUST00000162397.2,Chic2-002,retained_intron,ENSMUSG00000029229.8,protein_coding | 976.8251343 | 0.239435739 | 0.000193022 | 0.002048492 |
| ENSMUST00000166518.7,Ssr4-201,protein_coding,ENSMUSG00000002014.12,protein_coding | 3721.77034 | 0.230592105 | 0.000193014 | 0.002048492 |
| ENSMUST00000106307.8,Hivep3-001,protein_coding,ENSMUSG00000028634.16,protein_coding | 1790.036008 | -0.223314125 | 0.000193144 | 0.002048816 |
| ENSMUST00000051803.6,Aldh3b1-201,protein_coding,ENSMUSG00000024885.8,protein_coding | 2374.859344 | -0.248557377 | 0.000193333 | 0.002050311 |
| ENSMUST00000029825.13,Depdc1a-003,protein_coding,ENSMUSG00000028175.15,protein_coding | 32.50600002 | 0.687781595 | 0.000193716 | 0.002053359 |
| ENSMUST00000096232.5,Cebpd-001,protein_coding,ENSMUSG00000071637.5,protein_coding | 481.4961709 | 0.325133337 | 0.000193711 | 0.002053359 |
| ENSMUST00000022865.15,Mtdh-001,protein_coding,ENSMUSG00000022255.15,protein_coding | 17356.68607 | -0.251484466 | 0.000194132 | 0.002057265 |
| ENSMUST00000029651.10,Gstcd-001,protein_coding,ENSMUSG00000028018.15,protein_coding | 4885.213458 | 0.249977085 | 0.000194226 | 0.00205775 |
| ENSMUST00000021847.14,Pxdc1-001,nonsense_mediated_decay,ENSMUSG00000021411.15,protein_coding | 127.6328286 | -0.607239175 | 0.000195186 | 0.002067417 |
| ENSMUST00000152579.7,Fstl1-006,processed_transcript,ENSMUSG00000022816.11,protein_coding | 111.1409125 | -0.619455322 | 0.000195263 | 0.002067726 |
| ENSMUST00000107152.8,Hdac5-001,protein_coding,ENSMUSG00000008855.17,protein_coding | 4663.510015 | -0.166767493 | 0.000195796 | 0.002072859 |
| ENSMUST00000051454.3,Fam171b-001,protein_coding,ENSMUSG00000048388.3,protein_coding | 3044.52051 | 0.348004908 | 0.000196004 | 0.00207455 |
| ENSMUST00000146303.1,Rps9-007,retained_intron,ENSMUSG00000006333.14,protein_coding | 169.7674277 | -0.519294598 | 0.000196771 | 0.002081128 |
| ENSMUST00000078988.8,Lpp-202,protein_coding,ENSMUSG00000033306.14,protein_coding | 7570.373069 | -0.473838338 | 0.000196739 | 0.002081128 |
| ENSMUST00000047206.5,Plekhh2-201,protein_coding,ENSMUSG00000040852.5,protein_coding | 5121.235378 | -0.119015972 | 0.000196759 | 0.002081128 |
| ENSMUST00000089581.10,Pced1a-001,protein_coding,ENSMUSG00000037773.14,protein_coding | 180.7681429 | -0.620585653 | 0.000196857 | 0.002081534 |
| ENSMUST00000128220.7,Hnrnph1-015,processed_transcript,ENSMUSG00000007850.16,protein_coding | 39.41772289 | -0.684809827 | 0.000197172 | 0.002084348 |
| ENSMUST00000060419.1,Gjb4-201,protein_coding,ENSMUSG00000046623.8,protein_coding | 1189.099058 | -0.428165096 | 0.000197632 | 0.002088595 |
| ENSMUST00000215006.1,Anln-006,protein_coding,ENSMUSG00000036777.8,protein_coding | 466.1712141 | -0.347861377 | 0.000197671 | 0.002088595 |
| ENSMUST00000049257.7,Ddx52-001,protein_coding,ENSMUSG00000020677.7,protein_coding | 7227.296951 | -0.238923678 | 0.000197791 | 0.002089357 |
| ENSMUST00000058163.10,Pus10-002,protein_coding,ENSMUSG00000020280.12,protein_coding | 1859.333817 | -0.420774249 | 0.000197932 | 0.002090329 |
| ENSMUST00000204150.1,Phf14-010,retained_intron,ENSMUSG00000029629.17,protein_coding | 114.265566 | -0.575832092 | 0.000198995 | 0.002101042 |
| ENSMUST00000160653.1,Txndc5-003,protein_coding,ENSMUSG00000038991.16,protein_coding | 1197.40354 | 0.326120335 | 0.000199132 | 0.00210197 |
| ENSMUST00000106108.8,Zmym4-001,protein_coding,ENSMUSG00000042446.16,protein_coding | 5700.333305 | -0.224126614 | 0.000199269 | 0.002102896 |
| ENSMUST00000063683.6,Tagap1-201,protein_coding,ENSMUSG00000052031.7,protein_coding | 5081.394177 | 0.245725959 | 0.000199669 | 0.002106608 |
| ENSMUST00000029374.7,Nbea-001,protein_coding,ENSMUSG00000027799.12,protein_coding | 5223.261012 | -0.351416074 | 0.000199908 | 0.002108444 |
| ENSMUST00000115157.7,Tmem209-005,protein_coding,ENSMUSG00000029782.18,protein_coding | 4743.723188 | 0.264261309 | 0.000199941 | 0.002108444 |
| ENSMUST00000086635.8,Pitpnb-001,protein_coding,ENSMUSG00000050017.11,protein_coding | 20294.47305 | 0.144716046 | 0.000200629 | 0.002115178 |
| ENSMUST00000205152.2,Aup1-007,retained_intron,ENSMUSG00000068328.9,protein_coding | 126.3395198 | 0.655343996 | 0.000201027 | 0.002118851 |
| ENSMUST00000043321.5,Mapk9-006,protein_coding,ENSMUSG00000020366.18,protein_coding | 364.1856155 | 0.509090154 | 0.000201865 | 0.002127164 |
| ENSMUST00000026917.9,Nrp1-001,protein_coding,ENSMUSG00000025810.9,protein_coding | 3260.677494 | -0.285048084 | 0.000202135 | 0.002129491 |
| ENSMUST00000130926.7,Mta1-004,processed_transcript,ENSMUSG00000021144.14,protein_coding | 727.9420096 | 0.601322407 | 0.000202761 | 0.00213556 |
| ENSMUST00000078975.7,G6pc3-002,protein_coding,ENSMUSG00000034793.15,protein_coding | 1613.666891 | -0.210999079 | 0.000203204 | 0.002139709 |
| ENSMUST00000114459.7,Hnrnpa2b1-006,protein_coding,ENSMUSG00000004980.16,protein_coding | 7742.247536 | 0.325243922 | 0.0002033 | 0.00214019 |
| ENSMUST00000099575.3,Grem1-001,protein_coding,ENSMUSG00000074934.3,protein_coding | 5771.426936 | -0.354963273 | 0.000204072 | 0.0021478 |
| ENSMUST00000130386.7,Als2cl-001,protein_coding,ENSMUSG00000044037.15,protein_coding | 331.9148522 | 0.634455296 | 0.000204135 | 0.002147928 |
| ENSMUST00000087268.6,Abca12-001,protein_coding,ENSMUSG00000050296.14,protein_coding | 78.33660629 | 0.689332141 | 0.000204276 | 0.002148431 |
| ENSMUST00000037127.14,Eri3-001,protein_coding,ENSMUSG00000033423.16,protein_coding | 6908.411716 | 0.171214085 | 0.000204332 | 0.002148431 |
| ENSMUST00000025743.6,Mrpl21-001,protein_coding,ENSMUSG00000024829.12,protein_coding | 5104.153223 | 0.137326389 | 0.000204288 | 0.002148431 |
| ENSMUST00000116375.1,Cstf1-001,protein_coding,ENSMUSG00000027498.14,protein_coding | 8518.864889 | 0.164654966 | 0.000204716 | 0.002151942 |
| ENSMUST00000049519.3,Irgm1-001,protein_coding,ENSMUSG00000046879.7,protein_coding | 1323.559606 | -0.268956903 | 0.000204996 | 0.002154361 |
| ENSMUST00000165563.7,Micu1-001,protein_coding,ENSMUSG00000020111.15,protein_coding | 14275.98432 | 0.205452752 | 0.000205359 | 0.002157647 |
| ENSMUST00000121869.7,Dzip3-001,protein_coding,ENSMUSG00000064061.13,protein_coding | 1904.686516 | -0.527261073 | 0.000205651 | 0.002159659 |
| ENSMUST00000172302.7,Vldlr-002,protein_coding,ENSMUSG00000024924.14,protein_coding | 2879.640643 | 0.450175254 | 0.000205604 | 0.002159659 |
| ENSMUST00000161821.3,Marc2-003,protein_coding,ENSMUSG00000073481.9,protein_coding | 1087.143225 | 0.291735032 | 0.000205733 | 0.002159996 |
| ENSMUST00000145271.1,Mrps5-005,retained_intron,ENSMUSG00000027374.12,protein_coding | 2306.434313 | 0.173169477 | 0.000206013 | 0.002162409 |
| ENSMUST00000131786.7,Huwe1-006,retained_intron,ENSMUSG00000025261.17,protein_coding | 1340.634944 | -0.335859497 | 0.000207074 | 0.002173013 |
| ENSMUST00000067458.6,Sema5a-201,protein_coding,ENSMUSG00000022231.9,protein_coding | 1103.418637 | 0.262948671 | 0.000207876 | 0.002180898 |
| ENSMUST00000019726.6,Plin3-201,protein_coding,ENSMUSG00000024197.9,protein_coding | 7662.015619 | 0.169724118 | 0.000208623 | 0.002188205 |
| ENSMUST00000034896.12,Mto1-001,protein_coding,ENSMUSG00000032342.13,protein_coding | 1803.580594 | -0.162945578 | 0.000209826 | 0.002200017 |
| ENSMUST00000115677.7,Tfe3-002,protein_coding,ENSMUSG00000000134.17,protein_coding | 2841.487504 | 0.284056464 | 0.000209852 | 0.002200017 |
| ENSMUST00000216558.1,Ubp1-002,protein_coding,ENSMUSG00000009741.15,protein_coding | 1060.688924 | 0.437798532 | 0.000209993 | 0.002200962 |
| ENSMUST00000171366.6,Disp1-003,protein_coding,ENSMUSG00000030768.12,protein_coding | 2758.455406 | -0.276756219 | 0.000210141 | 0.002201979 |
| ENSMUST00000049248.5,Malt1-001,protein_coding,ENSMUSG00000032688.7,protein_coding | 1102.635652 | -0.309117122 | 0.000210236 | 0.00220244 |
| ENSMUST00000026027.5,Taf5-201,protein_coding,ENSMUSG00000025049.5,protein_coding | 3042.072879 | 0.148128951 | 0.000210817 | 0.002207988 |
| ENSMUST00000021164.3,Fam64a-001,protein_coding,ENSMUSG00000020808.3,protein_coding | 9962.906945 | 0.155583379 | 0.00021093 | 0.002208637 |
| ENSMUST00000020263.13,Hnrnph3-001,protein_coding,ENSMUSG00000020069.16,protein_coding | 1167.080017 | -0.442990376 | 0.000211337 | 0.002212359 |
| ENSMUST00000165164.8,Pcgf1-001,protein_coding,ENSMUSG00000069678.10,protein_coding | 1034.385792 | 0.374117237 | 0.000212131 | 0.002220126 |
| ENSMUST00000141504.7,Recql-003,protein_coding,ENSMUSG00000030243.16,protein_coding | 126.5930418 | -0.523903682 | 0.0002122 | 0.002220313 |
| ENSMUST00000009693.14,Srp14-001,protein_coding,ENSMUSG00000009549.14,protein_coding | 8698.90539 | 0.172495132 | 0.000212629 | 0.002224262 |
| ENSMUST00000190998.1,Fam132b-002,protein_coding,ENSMUSG00000047443.14,protein_coding | 61.41195461 | 0.610561444 | 0.000213102 | 0.002228668 |
| ENSMUST00000131147.1,Gm13387-002,lincRNA,ENSMUSG00000087028.7,lincRNA | 133.660539 | -0.556825544 | 0.000213177 | 0.002228909 |
| ENSMUST00000052012.13,Phka1-001,protein_coding,ENSMUSG00000034055.16,protein_coding | 5134.381815 | -0.168836384 | 0.000213675 | 0.002233578 |
| ENSMUST00000167466.1,Sel1l-006,protein_coding,ENSMUSG00000020964.14,protein_coding | 3289.88077 | 0.241404591 | 0.000214128 | 0.002237766 |
| ENSMUST00000055485.11,Lsm14b-001,protein_coding,ENSMUSG00000039108.14,protein_coding | 8497.105004 | 0.14359261 | 0.000214304 | 0.002239062 |
| ENSMUST00000176015.4,Uchl1-002,retained_intron,ENSMUSG00000029223.12,protein_coding | 375.7270889 | -0.512196267 | 0.000214364 | 0.002239141 |
| ENSMUST00000195728.1,Rpl22l1-008,retained_intron,ENSMUSG00000039221.10,protein_coding | 160.2813005 | -0.480184377 | 0.000214516 | 0.002240135 |
| ENSMUST00000136335.7,Primpol-003,nonsense_mediated_decay,ENSMUSG00000038225.15,protein_coding | 457.128045 | 0.567510183 | 0.000214563 | 0.002240135 |
| ENSMUST00000162413.7,Tra2b-003,nonsense_mediated_decay,ENSMUSG00000022858.15,protein_coding | 2143.261037 | 0.42497051 | 0.000214831 | 0.002242391 |
| ENSMUST00000198437.4,Dclk1-005,protein_coding,ENSMUSG00000027797.15,protein_coding | 133.4481458 | -0.632650798 | 0.000215737 | 0.002250755 |
| ENSMUST00000109930.2,Ascc2-002,protein_coding,ENSMUSG00000020412.16,protein_coding | 4945.131153 | 0.206304164 | 0.000215698 | 0.002250755 |
| ENSMUST00000038893.5,Rcc2-201,protein_coding,ENSMUSG00000040945.13,protein_coding | 62679.12774 | 0.12646229 | 0.000215989 | 0.002252833 |
| ENSMUST00000154832.1,Gga3-011,processed_transcript,ENSMUSG00000020740.13,protein_coding | 297.5664398 | -0.374568548 | 0.000216353 | 0.002256085 |
| ENSMUST00000021181.12,0610010K14Rik-008,protein_coding,ENSMUSG00000020831.18,protein_coding | 2263.386171 | 0.233686896 | 0.000216497 | 0.002257045 |
| ENSMUST00000085358.11,Tex9-003,protein_coding,ENSMUSG00000090626.9,protein_coding | 135.6031509 | -0.559030843 | 0.000216645 | 0.002258037 |
| ENSMUST00000120044.8,Cngb1-002,protein_coding,ENSMUSG00000031789.17,protein_coding | 49.01259041 | 0.672086968 | 0.000217728 | 0.002268772 |
| ENSMUST00000138301.7,Frmd4b-011,retained_intron,ENSMUSG00000030064.16,protein_coding | 182.5867287 | -0.568573507 | 0.000217798 | 0.002268953 |
| ENSMUST00000107869.8,Tom1l1-008,protein_coding,ENSMUSG00000020541.12,protein_coding | 511.6771173 | 0.471130537 | 0.000218024 | 0.002270761 |
| ENSMUST00000064187.11,Thra-001,protein_coding,ENSMUSG00000058756.13,protein_coding | 5098.586605 | 0.15648342 | 0.000218219 | 0.002272248 |
| ENSMUST00000131487.7,Pogk-002,protein_coding,ENSMUSG00000040596.15,protein_coding | 179.2275852 | -0.660031819 | 0.000218551 | 0.002275149 |
| ENSMUST00000100784.8,Flot2-201,protein_coding,ENSMUSG00000061981.14,protein_coding | 377.3011264 | 0.612019928 | 0.000218814 | 0.002276791 |
| ENSMUST00000033751.7,Vegfd-001,protein_coding,ENSMUSG00000031380.10,protein_coding | 337.5752019 | -0.379152967 | 0.000218795 | 0.002276791 |
| ENSMUST00000111269.1,Sap18-006,protein_coding,ENSMUSG00000021963.16,protein_coding | 3336.984441 | 0.184112985 | 0.000219517 | 0.002283553 |
| ENSMUST00000215842.1,Bag1-202,protein_coding,ENSMUSG00000028416.12,protein_coding | 4804.105946 | 0.308143321 | 0.000219981 | 0.002287823 |

| | | | | |
|---|---|---|---|---|
| ENSMUST00000024954.9,Epas1-201,protein_coding,ENSMUSG00000024140.9,protein_coding | 134.9492045 | -0.462211421 | 0.000220457 | 0.002292225 |
| ENSMUST00000030436.11,Pnrc2-001,protein_coding,ENSMUSG00000028675.12,protein_coding | 7732.248619 | 0.241490587 | 0.000220763 | 0.002294846 |
| ENSMUST00000045140.4,Tcaf1-002,protein_coding,ENSMUSG00000036667.14,protein_coding | 604.3711017 | -0.334131993 | 0.000221078 | 0.00229757 |
| ENSMUST00000129711.1,Aurkaip1-005,processed_transcript,ENSMUSG00000065990.12,protein_coding | 65.23273078 | 0.612468391 | 0.000222231 | 0.002308988 |
| ENSMUST00000139132.1,Exoc4-006,processed_transcript,ENSMUSG00000029763.17,protein_coding | 33.07981219 | 0.689167951 | 0.000222599 | 0.002312257 |
| ENSMUST00000004686.12,Pex11g-001,protein_coding,ENSMUSG00000069633.12,protein_coding | 602.4891433 | 0.265087121 | 0.000222657 | 0.002312306 |
| ENSMUST00000196449.4,Vcam1-002,protein_coding,ENSMUSG00000027962.14,protein_coding | 289.1622881 | -0.512855788 | 0.000222905 | 0.002314321 |
| ENSMUST00000164828.7,Pex2-201,protein_coding,ENSMUSG00000040374.13,protein_coding | 223.583746 | -0.58745273 | 0.000223229 | 0.002317127 |
| ENSMUST00000062041.5,Ispd-201,protein_coding,ENSMUSG00000043153.5,protein_coding | 416.2443553 | 0.336317495 | 0.000224039 | 0.00232497 |
| ENSMUST00000075326.10,Fnbp1-007,protein_coding,ENSMUSG00000075415.13,protein_coding | 44.38445559 | 0.675591728 | 0.000224335 | 0.002327485 |
| ENSMUST00000028644.10,Cd82-002,protein_coding,ENSMUSG00000027215.13,protein_coding | 1624.089711 | 0.327310755 | 0.000224574 | 0.002329405 |
| ENSMUST00000002064.14,Blvra-001,protein_coding,ENSMUSG00000001999.15,protein_coding | 4345.461181 | 0.182707395 | 0.00022464 | 0.002329527 |
| ENSMUST00000125536.1,Ing2-002,processed_transcript,ENSMUSG00000063049.4,protein_coding | 690.411803 | 0.337883093 | 0.000224704 | 0.002329631 |
| ENSMUST00000035216.10,Uba7-001,protein_coding,ENSMUSG00000032596.14,protein_coding | 167.101879 | 0.632872203 | 0.000225192 | 0.002334127 |
| ENSMUST00000071642.10,Mdn1-001,protein_coding,ENSMUSG00000058006.12,protein_coding | 25475.76589 | -0.184066816 | 0.000225337 | 0.002335068 |
| ENSMUST00000140466.7,Dync2h1-001,protein_coding,ENSMUSG00000047193.16,protein_coding | 651.9260345 | -0.536750713 | 0.000226463 | 0.002346163 |
| ENSMUST00000040519.11,Apopt1-001,protein_coding,ENSMUSG00000037787.13,protein_coding | 2913.192574 | 0.218762679 | 0.00022656 | 0.002346612 |
| ENSMUST00000117330.7,Ick-006,protein_coding,ENSMUSG00000009828.15,protein_coding | 383.0996658 | 0.541907833 | 0.000226731 | 0.002347222 |
| ENSMUST00000001700.6,Hoxc13-001,protein_coding,ENSMUSG00000001655.6,protein_coding | 1003.707807 | -0.215734031 | 0.000226783 | 0.002347222 |
| ENSMUST00000066368.12,Mdfi-001,protein_coding,ENSMUSG00000032717.14,protein_coding | 173.3927101 | -0.445700015 | 0.000226702 | 0.002347222 |
| ENSMUST00000048946.6,Card19-201,protein_coding,ENSMUSG00000037960.10,protein_coding | 6582.389676 | 0.180377716 | 0.00022698 | 0.002348696 |
| ENSMUST00000002091.5,Bcap31-001,protein_coding,ENSMUSG00000002015.5,protein_coding | 13000.50461 | 0.147425742 | 0.000227193 | 0.002350336 |
| ENSMUST00000086519.11,Rplp0-001,protein_coding,ENSMUSG00000067274.10,protein_coding | 456651.443 | 0.181935865 | 0.000227934 | 0.002357436 |
| ENSMUST00000030805.13,Park7-001,protein_coding,ENSMUSG00000028964.14,protein_coding | 17611.91131 | 0.177962509 | 0.000228095 | 0.002358532 |
| ENSMUST00000069004.12,Dcbld1-201,protein_coding,ENSMUSG00000019891.14,protein_coding | 1602.759472 | 0.253800876 | 0.000228275 | 0.002359825 |
| ENSMUST00000184535.7,1700037H04Rik-003,nonsense_mediated_decay,ENSMUSG00000027327.16,protein_coding | 288.3917787 | 0.474309152 | 0.000229322 | 0.002368942 |
| ENSMUST00000112148.3,Foxm1-003,protein_coding,ENSMUSG00000001517.14,protein_coding | 2259.588833 | 0.398553911 | 0.00022929 | 0.002368942 |
| ENSMUST00000061468.8,Bcl7c-001,protein_coding,ENSMUSG00000030814.17,protein_coding | 6512.40933 | 0.159087848 | 0.000229277 | 0.002368942 |
| ENSMUST00000035987.7,Map3k9-201,protein_coding,ENSMUSG00000042724.7,protein_coding | 406.0024958 | 0.363191277 | 0.000229533 | 0.002370554 |
| ENSMUST00000039197.8,Slc44a3-001,protein_coding,ENSMUSG00000039865.8,protein_coding | 136.0019411 | 0.526464694 | 0.000229767 | 0.002371833 |
| ENSMUST00000112294.8,Ap1s2-003,protein_coding,ENSMUSG00000031367.15,protein_coding | 568.8727642 | -0.408540658 | 0.000229748 | 0.002371833 |
| ENSMUST00000106941.8,Tlk2-006,protein_coding,ENSMUSG00000020694.16,protein_coding | 8583.206818 | 0.266816993 | 0.000229853 | 0.002372158 |
| ENSMUST00000038677.3,Rrp12-201,protein_coding,ENSMUSG00000035049.3,protein_coding | 11771.58691 | -0.196224088 | 0.000230177 | 0.00237493 |
| ENSMUST00000053087.3,Klhl23-001,protein_coding,ENSMUSG00000042155.3,protein_coding | 1806.566846 | -0.212889025 | 0.000231593 | 0.00238866 |
| ENSMUST00000129269.7,Stag1-001,protein_coding,ENSMUSG00000037286.15,protein_coding | 3698.908317 | -0.396943675 | 0.000231619 | 0.00238866 |
| ENSMUST00000132357.7,Fam117a-006,processed_transcript,ENSMUSG00000038893.12,protein_coding | 66.01767791 | 0.670330105 | 0.000231928 | 0.002391271 |
| ENSMUST00000113272.7,Eng-005,protein_coding,ENSMUSG00000026814.16,protein_coding | 139.7627383 | -0.521919239 | 0.000232408 | 0.00239565 |
| ENSMUST00000102482.4,Gm12481-001,processed_pseudogene,ENSMUSG00000043770.10,processed_pseudogene | 372.8559914 | -0.662402617 | 0.000232525 | 0.002396285 |
| ENSMUST00000141874.7,Zfp106-002,retained_intron,ENSMUSG00000027288.16,protein_coding | 770.6753037 | -0.258351901 | 0.000233189 | 0.002401396 |
| ENSMUST00000172356.7,Thsd7a-201,protein_coding,ENSMUSG00000032625.14,protein_coding | 925.8328552 | -0.478683653 | 0.000233144 | 0.002401396 |
| ENSMUST00000112933.1,Cntf-001,protein_coding,ENSMUSG00000079415.2,protein_coding | 95.16768136 | -0.644930636 | 0.000233111 | 0.002401396 |
| ENSMUST00000045029.14,Pigk-002,protein_coding,ENSMUSG00000039047.17,protein_coding | 1026.954748 | 0.374121151 | 0.000233925 | 0.002408404 |
| ENSMUST00000029780.11,Ptbp2-201,protein_coding,ENSMUSG00000028134.11,protein_coding | 1782.205585 | -0.462455923 | 0.000234019 | 0.002408799 |
| ENSMUST00000145132.7,Ttll4-010,processed_transcript,ENSMUSG00000033257.14,protein_coding | 2634.78454 | -0.191255226 | 0.00023498 | 0.002418104 |
| ENSMUST00000171158.7,Sdc1-201,protein_coding,ENSMUSG00000020592.14,protein_coding | 8080.523804 | 0.147902296 | 0.00023554 | 0.002423294 |
| ENSMUST00000007046.7,Tmco6-201,protein_coding,ENSMUSG00000006850.7,protein_coding | 1133.400758 | 0.265508552 | 0.000236552 | 0.002433123 |
| ENSMUST00000067218.13,Parp3-001,protein_coding,ENSMUSG00000023249.15,protein_coding | 196.2686511 | -0.561403799 | 0.000236758 | 0.002434663 |
| ENSMUST00000084846.11,Srrm1-001,protein_coding,ENSMUSG00000028809.16,protein_coding | 5352.530962 | 0.345393616 | 0.000236921 | 0.002435754 |
| ENSMUST00000015981.11,Sqstm1-002,protein_coding,ENSMUSG00000015837.15,protein_coding | 2708.881913 | 0.30115759 | 0.00023751 | 0.002441231 |
| ENSMUST00000135337.7,Cep44-003,protein_coding,ENSMUSG00000038215.15,protein_coding | 202.9962798 | 0.49283505 | 0.000237654 | 0.002442122 |
| ENSMUST00000028291.14,Inpp5e-003,processed_transcript,ENSMUSG00000026925.16,protein_coding | 1138.040992 | 0.42354691 | 0.000238913 | 0.002454472 |
| ENSMUST00000067689.7,Tnrc6b-201,protein_coding,ENSMUSG00000047888.9,protein_coding | 12353.33963 | -0.248753765 | 0.000239163 | 0.002456456 |
| ENSMUST00000113539.7,Zxdc-003,protein_coding,ENSMUSG00000034430.16,protein_coding | 1194.038529 | 0.26472042 | 0.000240904 | 0.002473748 |
| ENSMUST00000075181.10,Plekha1-001,protein_coding,ENSMUSG00000040268.17,protein_coding | 764.4079165 | 0.346608842 | 0.000241216 | 0.00247636 |
| ENSMUST00000201421.3,Sept11-005,protein_coding,ENSMUSG00000058013.11,protein_coding | 820.9384644 | 0.321109083 | 0.000243757 | 0.002501855 |
| ENSMUST00000027149.11,Lancl1-001,protein_coding,ENSMUSG00000026000.16,protein_coding | 329.5058399 | 0.365307244 | 0.000244167 | 0.002505461 |
| ENSMUST00000211212.1,Ppp1r15a-003,protein_coding,ENSMUSG00000040435.12,protein_coding | 105.0282559 | 0.684630242 | 0.000244318 | 0.00250582 |
| ENSMUST00000066743.10,Adnp2-001,protein_coding,ENSMUSG00000053950.10,protein_coding | 5911.060397 | 0.177427723 | 0.000244309 | 0.00250582 |
| ENSMUST00000185913.6,Ttk-004,protein_coding,ENSMUSG00000038379.15,protein_coding | 141.3000763 | -0.684593226 | 0.000244891 | 0.002511094 |
| ENSMUST00000198748.4,Ube2d3-005,nonsense_mediated_decay,ENSMUSG00000078578.9,protein_coding | 3874.981828 | 0.173085319 | 0.000245525 | 0.002517 |
| ENSMUST00000100819.11,Tmem254c-001,protein_coding,ENSMUSG00000072680.11,protein_coding | 1423.233952 | 0.251388075 | 0.000245742 | 0.002518618 |
| ENSMUST00000137400.7,Rps14-004,protein_coding,ENSMUSG00000024608.10,protein_coding | 10754.43736 | 0.276275271 | 0.000246049 | 0.002521163 |
| ENSMUST00000054294.6,Fzd1-001,protein_coding,ENSMUSG00000044674.6,protein_coding | 1443.462831 | -0.224864086 | 0.000247758 | 0.002538075 |
| ENSMUST00000051572.7,Sdpr-001,protein_coding,ENSMUSG00000045954.7,protein_coding | 3747.838916 | -0.316653687 | 0.000247972 | 0.002539663 |
| ENSMUST00000109187.1,Hsp25-ps1-201,pseudogene,ENSMUSG00000078915.1,pseudogene | 73.10885309 | -0.642130326 | 0.000248656 | 0.002546065 |
| ENSMUST00000182959.7,Ank2-015,protein_coding,ENSMUSG00000032826.17,protein_coding | 2093.052665 | 0.361407777 | 0.000249036 | 0.002549347 |
| ENSMUST00000153162.7,Ppp4r1l-ps-003,retained_intron,ENSMUSG00000055897.13,transcribed_unprocessed_pseudogene | 543.4324644 | 0.26668624 | 0.000249217 | 0.002550593 |
| ENSMUST00000103129.8,Dsn1-001,protein_coding,ENSMUSG00000027635.15,protein_coding | 3711.397964 | 0.189177745 | 0.000249566 | 0.002553556 |
| ENSMUST00000162768.7,Zfand2b-002,protein_coding,ENSMUSG00000026197.12,protein_coding | 222.7030808 | 0.441572537 | 0.000250649 | 0.002564027 |
| ENSMUST00000030541.12,Hp1bp3-201,protein_coding,ENSMUSG00000028759.13,protein_coding | 22002.05623 | 0.165910143 | 0.000251329 | 0.00257037 |
| ENSMUST00000032069.7,Add2-201,protein_coding,ENSMUSG00000030000.10,protein_coding | 1507.575425 | 0.411471325 | 0.000251712 | 0.002573306 |
| ENSMUST00000022153.6,Ptcd2-201,protein_coding,ENSMUSG00000021650.6,protein_coding | 3868.115321 | 0.190450034 | 0.000251735 | 0.002573306 |
| ENSMUST00000006123.10,Tuft1-001,protein_coding,ENSMUSG00000005968.14,protein_coding | 6223.232612 | -0.224907411 | 0.000252279 | 0.002578257 |
| ENSMUST00000199335.1,St7l-009,nonsense_mediated_decay,ENSMUSG00000045576.16,protein_coding | 112.1962955 | 0.577537211 | 0.000252378 | 0.002578655 |
| ENSMUST00000011058.8,Pdhx-001,protein_coding,ENSMUSG00000010914.10,protein_coding | 3867.833582 | 0.163624263 | 0.000252523 | 0.00257952 |
| ENSMUST00000023687.7,Ifngr2-001,protein_coding,ENSMUSG00000022965.7,protein_coding | 6126.200468 | 0.177767556 | 0.00025309 | 0.002584696 |
| ENSMUST00000105249.8,R3hdm2-006,protein_coding,ENSMUSG00000025404.15,protein_coding | 10068.24375 | 0.178056997 | 0.000253171 | 0.002584916 |
| ENSMUST00000171533.7,Nrep-203,protein_coding,ENSMUSG00000042834.14,protein_coding | 124.3915044 | -0.681169574 | 0.000253614 | 0.002588825 |
| ENSMUST00000004200.8,Cwc15-001,protein_coding,ENSMUSG00000004096.9,protein_coding | 5300.620945 | 0.13329723 | 0.000254264 | 0.002594838 |
| ENSMUST00000023259.14,Lynx1-001,protein_coding,ENSMUSG00000022594.14,protein_coding | 621.2728342 | -0.37847793 | 0.000254436 | 0.002595983 |
| ENSMUST00000121902.1,1700030K09Rik-002,protein_coding,ENSMUSG00000052794.13,protein_coding | 579.6544245 | 0.282859616 | 0.000254932 | 0.002600428 |
| ENSMUST00000099820.9,Lgals8-004,protein_coding,ENSMUSG00000057554.13,protein_coding | 1893.999243 | 0.284608614 | 0.000255276 | 0.002603317 |
| ENSMUST00000174759.7,Pear1-004,protein_coding,ENSMUSG00000028073.15,protein_coding | 209.1541803 | -0.442421735 | 0.000255502 | 0.002605008 |
| ENSMUST00000194546.5,Igsf10-001,protein_coding,ENSMUSG00000036334.12,protein_coding | 58.32158854 | -0.679066905 | 0.000256261 | 0.00261213 |
| ENSMUST00000114050.7,Tmsb10-001,protein_coding,ENSMUSG00000079523.8,protein_coding | 100913.6787 | 0.19333987 | 0.000257517 | 0.00262431 |
| ENSMUST00000049089.5,Twist1-201,protein_coding,ENSMUSG00000035799.5,protein_coding | 3865.147384 | 0.136004155 | 0.000259686 | 0.00264579 |
| ENSMUST00000029885.4,Cpne3-001,protein_coding,ENSMUSG00000028228.5,protein_coding | 5836.909143 | -0.266224868 | 0.000259853 | 0.002646857 |
| ENSMUST00000053183.10,Arf3-001,protein_coding,ENSMUSG00000051853.9,protein_coding | 36606.74919 | -0.100079711 | 0.000260193 | 0.002649696 |
| ENSMUST00000191097.1,Snrpd3-002,retained_intron,ENSMUSG00000020180.9,protein_coding | 1639.953057 | -0.261291226 | 0.000260562 | 0.00265283 |
| ENSMUST00000169169.7,Tnnc1-001,protein_coding,ENSMUSG00000091898.8,protein_coding | 120.6291137 | 0.554907183 | 0.000260626 | 0.002652856 |
| ENSMUST00000026016.12,Fundc1-001,protein_coding,ENSMUSG00000025040.13,protein_coding | 1778.965314 | -0.452894293 | 0.00026126 | 0.002658674 |
| ENSMUST00000131814.7,Taf13-002,processed_transcript,ENSMUSG00000048100.13,protein_coding | 138.0498446 | -0.543067894 | 0.000261476 | 0.00266025 |

| | | | | |
|---|---|---|---|---|
| ENSMUST00000140527.1,Rpl30-004,processed_transcript,ENSMUSG00000058600.13,protein_coding | 10350.1645 | 0.196275527 | 0.000261618 | 0.002661061 |
| ENSMUST00000161825.7,Gfm2-008,protein_coding,ENSMUSG00000021666.16,protein_coding | 227.2947614 | -0.42095664 | 0.000261767 | 0.002661947 |
| ENSMUST00000094216.4,Mesdc1-001,protein_coding,ENSMUSG00000070462.4,protein_coding | 4071.905684 | -0.126129814 | 0.000262097 | 0.002664677 |
| ENSMUST00000108195.9,Dnajc19-002,protein_coding,ENSMUSG00000027679.13,protein_coding | 2709.064358 | 0.206738174 | 0.000262704 | 0.002670211 |
| ENSMUST00000148333.7,Steap2-002,protein_coding,ENSMUSG00000015653.13,protein_coding | 992.8078045 | 0.530504858 | 0.000263348 | 0.002676124 |
| ENSMUST00000032726.13,Tm2d3-002,protein_coding,ENSMUSG00000078681.10,protein_coding | 1158.873668 | 0.251865585 | 0.000263895 | 0.00268105 |
| ENSMUST00000034328.12,Ints10-001,protein_coding,ENSMUSG00000031864.16,protein_coding | 2386.368772 | 0.188145728 | 0.000264004 | 0.002681533 |
| ENSMUST00000034955.7,Spg21-001,protein_coding,ENSMUSG00000032388.7,protein_coding | 6038.727301 | 0.194941076 | 0.000264102 | 0.002681893 |
| ENSMUST00000099996.4,Ttc30b-001,protein_coding,ENSMUSG00000075273.4,protein_coding | 490.9268625 | -0.250262027 | 0.000264204 | 0.002682292 |
| ENSMUST00000183037.1,Gpx4-007,protein_coding,ENSMUSG00000075706.10,protein_coding | 12505.45303 | 0.230947166 | 0.000264344 | 0.002683078 |
| ENSMUST00000147076.7,Rrp1b-002,retained_intron,ENSMUSG00000058392.12,protein_coding | 1106.786496 | -0.243824882 | 0.000264882 | 0.002687913 |
| ENSMUST00000150582.7,Stam2-003,retained_intron,ENSMUSG00000055371.17,protein_coding | 707.5336453 | 0.314498835 | 0.000265079 | 0.002689274 |
| ENSMUST00000205336.1,Aldoa-011,protein_coding,ENSMUSG00000030695.14,protein_coding | 34.62647752 | 0.670602077 | 0.000265702 | 0.002694958 |
| ENSMUST00000216812.1,Kif23-012,processed_transcript,ENSMUSG00000032254.10,protein_coding | 456.7077805 | -0.309664919 | 0.000266125 | 0.002698612 |
| ENSMUST00000194171.5,Usp19-017,protein_coding,ENSMUSG00000006676.16,protein_coding | 96.09123628 | 0.474159909 | 0.000266393 | 0.002700693 |
| ENSMUST00000078676.5,Uqcrh-001,protein_coding,ENSMUSG00000063882.12,protein_coding | 8365.89872 | -0.270282904 | 0.00026652 | 0.002701344 |
| ENSMUST00000043551.10,Ankib1-001,protein_coding,ENSMUSG00000040351.11,protein_coding | 7174.920697 | -0.210694367 | 0.000267051 | 0.002706089 |
| ENSMUST00000004396.12,Atp6v1f-001,protein_coding,ENSMUSG00000004285.12,protein_coding | 11448.95864 | 0.174286493 | 0.000268489 | 0.002720021 |
| ENSMUST00000212062.1,Nup93-006,retained_intron,ENSMUSG00000032939.14,protein_coding | 79.86552725 | 0.642458668 | 0.000270117 | 0.002735869 |
| ENSMUST00000122965.7,Elp4-001,protein_coding,ENSMUSG00000027167.11,protein_coding | 2485.070256 | 0.145521236 | 0.000272097 | 0.002755275 |
| ENSMUST00000044970.6,Mgat3-201,protein_coding,ENSMUSG00000042428.6,protein_coding | 3478.504037 | -0.210943863 | 0.000272965 | 0.002763413 |
| ENSMUST00000043493.5,Ice1-201,protein_coding,ENSMUSG00000034525.6,protein_coding | 10382.2468 | 0.199745444 | 0.000273637 | 0.002769565 |
| ENSMUST00000086851.1,Hes6-001,protein_coding,ENSMUSG00000067071.8,protein_coding | 1665.414272 | 0.253619755 | 0.000274008 | 0.002772677 |
| ENSMUST00000027071.6,Lactb2-001,protein_coding,ENSMUSG00000025937.6,protein_coding | 4913.708961 | 0.204759445 | 0.000274178 | 0.002773741 |
| ENSMUST00000075724.8,Rfxank-001,protein_coding,ENSMUSG00000036120.10,protein_coding | 925.2512537 | -0.231571999 | 0.000275003 | 0.00278125 |
| ENSMUST00000034543.4,Rpusd4-001,protein_coding,ENSMUSG00000032044.4,protein_coding | 1850.253831 | 0.164091634 | 0.000275049 | 0.002781249 |
| ENSMUST00000026227.2,Twnk-201,protein_coding,ENSMUSG00000025209.4,protein_coding | 5745.950799 | 0.140292241 | 0.000275358 | 0.002783711 |
| ENSMUST00000134053.7,Alas1-004,processed_transcript,ENSMUSG00000032786.15,protein_coding | 3025.456072 | 0.208384494 | 0.000276241 | 0.00279199 |
| ENSMUST00000066070.6,Eif3k-001,protein_coding,ENSMUSG00000053565.10,protein_coding | 17202.18289 | 0.157824816 | 0.000276384 | 0.002792124 |
| ENSMUST00000123575.1,Prpsap2-003,processed_transcript,ENSMUSG00000020528.14,protein_coding | 65.84285315 | 0.586234528 | 0.000276374 | 0.002792124 |
| ENSMUST00000048216.5,Abhd13-001,protein_coding,ENSMUSG00000040396.12,protein_coding | 2413.557071 | -0.217919085 | 0.000277188 | 0.002799582 |
| ENSMUST00000034949.9,Csnk1g1-001,protein_coding,ENSMUSG00000032384.16,protein_coding | 3604.336975 | -0.213326205 | 0.000277534 | 0.00280242 |
| ENSMUST00000102536.10,Rpl11-001,protein_coding,ENSMUSG00000059291.15,protein_coding | 60328.8848 | 0.165592873 | 0.0002776 | 0.002802434 |
| ENSMUST00000198502.1,Tpst2-007,protein_coding,ENSMUSG00000029344.14,protein_coding | 111.5580691 | 0.638566629 | 0.000277972 | 0.002805534 |
| ENSMUST00000011400.7,Adam19-001,protein_coding,ENSMUSG00000011256.16,protein_coding | 1774.638008 | -0.161113622 | 0.000278128 | 0.002806449 |
| ENSMUST00000025052.12,Rps10-002,protein_coding,ENSMUSG00000052146.15,protein_coding | 474.6733465 | 0.541510489 | 0.000278989 | 0.002814471 |
| ENSMUST00000042410.4,Cdk6-201,protein_coding,ENSMUSG00000040274.11,protein_coding | 1385.894043 | 0.317521864 | 0.000279057 | 0.002814505 |
| ENSMUST00000053266.10,Mcm3-001,protein_coding,ENSMUSG00000041859.10,protein_coding | 47544.4607 | 0.168596442 | 0.000280004 | 0.002821406 |
| ENSMUST00000186543.6,Srgap2-004,protein_coding,ENSMUSG00000026425.15,protein_coding | 3158.185528 | 0.210116413 | 0.000279955 | 0.002821406 |
| ENSMUST00000142554.7,Abl1-007,nonsense_mediated_decay,ENSMUSG00000026842.16,protein_coding | 32.91150973 | 0.674817563 | 0.000279819 | 0.002821406 |
| ENSMUST00000165965.7,Mark2-005,protein_coding,ENSMUSG00000024969.15,protein_coding | 1250.257732 | 0.365396399 | 0.000279983 | 0.002821406 |
| ENSMUST00000059458.4,Maml1-001,protein_coding,ENSMUSG00000050567.16,protein_coding | 7992.643641 | 0.128752429 | 0.000280351 | 0.002824248 |
| ENSMUST00000099945.5,Amd1-001,protein_coding,ENSMUSG00000075232.5,protein_coding | 15493.95713 | -0.195766563 | 0.000280449 | 0.002824573 |
| ENSMUST00000051589.8,Aasdhppt-001,protein_coding,ENSMUSG00000025894.11,protein_coding | 2447.784747 | -0.248311029 | 0.000281044 | 0.0028299 |
| ENSMUST00000118952.7,Papd5-002,protein_coding,ENSMUSG00000036779.12,protein_coding | 2997.213162 | 0.36631129 | 0.000283122 | 0.002850157 |
| ENSMUST00000114337.7,Nif3l1-002,protein_coding,ENSMUSG00000026036.17,protein_coding | 728.7723203 | 0.224653893 | 0.000284214 | 0.002860481 |
| ENSMUST00000029453.12,Vangl1-001,protein_coding,ENSMUSG00000027860.15,protein_coding | 1257.292878 | -0.228901669 | 0.000284828 | 0.002865991 |
| ENSMUST00000133986.1,Zfp740-007,protein_coding,ENSMUSG00000046897.16,protein_coding | 125.760222 | 0.673727321 | 0.000285494 | 0.002872023 |
| ENSMUST00000060427.4,Ier2-201,protein_coding,ENSMUSG00000053560.4,protein_coding | 5676.622806 | 0.195121179 | 0.00028596 | 0.00287604 |
| ENSMUST00000026900.13,Hgs-201,protein_coding,ENSMUSG00000025793.15,protein_coding | 9817.968037 | 0.146007151 | 0.000286156 | 0.002877338 |
| ENSMUST00000046514.12,Eprs-001,protein_coding,ENSMUSG00000026615.14,protein_coding | 72811.4371 | 0.232844523 | 0.000286887 | 0.002884015 |
| ENSMUST00000085852.10,Zfp68-002,protein_coding,ENSMUSG00000058291.14,protein_coding | 390.8918735 | -0.55679705 | 0.00028797 | 0.002893334 |
| ENSMUST00000033973.13,Rwdd4a-001,protein_coding,ENSMUSG00000031568.16,protein_coding | 3037.330052 | -0.223309401 | 0.000287997 | 0.002893334 |
| ENSMUST00000044551.7,Ick-001,protein_coding,ENSMUSG00000009828.15,protein_coding | 611.2276199 | -0.454233455 | 0.000288016 | 0.002893334 |
| ENSMUST00000114354.9,Irak1-001,protein_coding,ENSMUSG00000031392.18,protein_coding | 7529.701185 | -0.1833367 | 0.000288162 | 0.002894122 |
| ENSMUST00000133444.7,Tial1-013,nonsense_mediated_decay,ENSMUSG00000030846.15,protein_coding | 62.47128429 | 0.672399719 | 0.000288576 | 0.002897607 |
| ENSMUST00000108229.1,Pnisr-002,protein_coding,ENSMUSG00000028248.15,protein_coding | 476.9037915 | -0.469349151 | 0.000289212 | 0.002903319 |
| ENSMUST00000187040.6,Srcap-001,protein_coding,ENSMUSG00000053877.12,protein_coding | 10339.51482 | -0.189679386 | 0.000289511 | 0.002905643 |
| ENSMUST00000141557.7,Trappc13-002,protein_coding,ENSMUSG00000021711.16,protein_coding | 911.9500867 | -0.45291802 | 0.000289804 | 0.002907899 |
| ENSMUST00000212461.1,Golga4-011,protein_coding,ENSMUSG00000038708.10,protein_coding | 29.23596996 | -0.672923691 | 0.000290215 | 0.002911351 |
| ENSMUST00000169948.1,Rchy1-004,protein_coding,ENSMUSG00000029397.15,protein_coding | 1130.674842 | 0.358028167 | 0.000291904 | 0.002927606 |
| ENSMUST00000052457.14,Mtss1l-001,protein_coding,ENSMUSG00000033763.14,protein_coding | 2743.496158 | 0.312121882 | 0.000292134 | 0.002929237 |
| ENSMUST00000054030.7,Acy3-201,protein_coding,ENSMUSG00000024866.9,protein_coding | 1413.103641 | 0.215621695 | 0.000292661 | 0.00293383 |
| ENSMUST00000044567.3,Alg5-001,protein_coding,ENSMUSG00000036632.9,protein_coding | 4138.066477 | 0.178352342 | 0.000293042 | 0.002936972 |
| ENSMUST00000178399.1,Ripk3-203,protein_coding,ENSMUSG00000022221.13,protein_coding | 517.2649667 | 0.654752725 | 0.000293891 | 0.002944789 |
| ENSMUST00000034728.8,Dock6-007,protein_coding,ENSMUSG00000032198.9,protein_coding | 2461.311953 | 0.207568476 | 0.000295049 | 0.002955706 |
| ENSMUST00000007800.7,Arpp19-007,protein_coding,ENSMUSG00000007656.13,protein_coding | 1239.526589 | -0.328676015 | 0.000295178 | 0.00295631 |
| ENSMUST00000090355.10,Rbm33-002,non_stop_decay,ENSMUSG00000048271.14,protein_coding | 599.3186542 | 0.291678539 | 0.000295254 | 0.002956383 |
| ENSMUST00000198322.4,Ubap2l-016,protein_coding,ENSMUSG00000042520.16,protein_coding | 6147.711323 | 0.174927264 | 0.00029547 | 0.002957687 |
| ENSMUST00000125704.7,2010107G23Rik-002,protein_coding,ENSMUSG00000020083.13,protein_coding | 57.84005094 | 0.588435049 | 0.000295521 | 0.002957687 |
| ENSMUST00000112961.9,Lhx6-001,protein_coding,ENSMUSG00000026890.19,protein_coding | 52.13760908 | -0.584116553 | 0.00029633 | 0.00296493 |
| ENSMUST00000126022.7,Mdm2-007,processed_transcript,ENSMUSG00000020184.14,protein_coding | 73.32097961 | -0.649862349 | 0.000296383 | 0.00296493 |
| ENSMUST00000090759.4,Acp6-001,protein_coding,ENSMUSG00000028093.15,protein_coding | 3402.39848 | 0.190907439 | 0.000296586 | 0.002966269 |
| ENSMUST00000171088.7,Prr5l-202,protein_coding,ENSMUSG00000032841.15,protein_coding | 113.9105033 | -0.545083823 | 0.000296678 | 0.002966501 |
| ENSMUST00000132449.7,Pdhx-003,protein_coding,ENSMUSG00000010914.10,protein_coding | 457.5429859 | 0.395638188 | 0.000297321 | 0.00297155 |
| ENSMUST00000145794.2,Gm15663-001,lincRNA,ENSMUSG00000085282.8,processed_transcript | 48.15786309 | 0.639636478 | 0.000297309 | 0.00297155 |
| ENSMUST00000134860.7,Uxs1-005,processed_transcript,ENSMUSG00000057363.12,protein_coding | 29.23892284 | 0.671941134 | 0.000297573 | 0.002973379 |
| ENSMUST00000192666.1,Mfsd7b-002,nonsense_mediated_decay,ENSMUSG00000066595.9,protein_coding | 143.1005899 | 0.67416893 | 0.000298332 | 0.002978892 |
| ENSMUST00000156821.7,Dusp11-004,processed_transcript,ENSMUSG00000030002.15,protein_coding | 349.7852671 | 0.509288061 | 0.000298326 | 0.002978892 |
| ENSMUST00000208222.1,Sesn3-001,protein_coding,ENSMUSG00000032009.8,protein_coding | 2449.255862 | -0.249776668 | 0.000298227 | 0.002978892 |
| ENSMUST00000112307.3,Ppm1b-204,protein_coding,ENSMUSG00000061130.12,protein_coding | 801.6391219 | 0.334035909 | 0.000298733 | 0.002981514 |
| ENSMUST00000052761.8,Idh3g-001,protein_coding,ENSMUSG00000002010.17,protein_coding | 9569.536732 | 0.170726407 | 0.000298705 | 0.002981514 |
| ENSMUST00000140157.7,Nubp1-005,processed_transcript,ENSMUSG00000022503.10,protein_coding | 454.4018436 | 0.373765929 | 0.0002995 | 0.002988478 |
| ENSMUST00000001699.7,Hoxc10-001,protein_coding,ENSMUSG00000022484.7,protein_coding | 1012.339894 | -0.239016786 | 0.000301568 | 0.003008409 |
| ENSMUST00000045153.10,Pik3ip1-001,protein_coding,ENSMUSG00000034614.14,protein_coding | 402.2416677 | 0.341133495 | 0.000302503 | 0.003017041 |
| ENSMUST00000061586.10,Zfp507-001,protein_coding,ENSMUSG00000044452.10,protein_coding | 4985.197618 | 0.137063919 | 0.000303894 | 0.003030213 |
| ENSMUST00000070181.6,Itpkb-001,protein_coding,ENSMUSG00000038855.10,protein_coding | 4330.798912 | -0.120515875 | 0.000304532 | 0.003035874 |
| ENSMUST00000067124.5,Lpin1-201,protein_coding,ENSMUSG00000020593.14,protein_coding | 774.2471063 | 0.507731293 | 0.000304747 | 0.003037313 |
| ENSMUST00000094421.10,Cep250-202,protein_coding,ENSMUSG00000038241.16,protein_coding | 2271.996908 | -0.34313724 | 0.000305389 | 0.003042299 |
| ENSMUST00000020673.2,Vdac1-005,protein_coding,ENSMUSG00000020402.11,protein_coding | 11033.94959 | 0.228212789 | 0.000305337 | 0.003042299 |
| ENSMUST00000041282.12,Trim37-001,protein_coding,ENSMUSG00000018548.15,protein_coding | 9310.856654 | -0.183474225 | 0.000305631 | 0.003044005 |
| ENSMUST00000031391.8,Bcl7a-001,protein_coding,ENSMUSG00000029438.9,protein_coding | 794.0585162 | 0.403924704 | 0.000308471 | 0.003071578 |

| | | | | |
|---|---|---|---|---|
| ENSMUST00000151254.1,1110004F10Rik-003,nonsense_mediated_decay,ENSMUSG00000030663.12,protein_coding | 287.9326522 | 0.568528267 | 0.000309344 | 0.003079563 |
| ENSMUST00000140809.1,Qsox1-003,retained_intron,ENSMUSG00000033684.14,protein_coding | 87.14418355 | -0.512919941 | 0.000309836 | 0.003083751 |
| ENSMUST00000195394.5,Dcaf8-002,retained_intron,ENSMUSG00000026554.15,protein_coding | 604.3472894 | 0.397047326 | 0.000310293 | 0.003087577 |
| ENSMUST00000142871.1,4930458D05Rik-001,antisense,ENSMUSG00000087611.1,antisense | 180.5267257 | 0.404102545 | 0.000310726 | 0.003091175 |
| ENSMUST00000144211.1,Srrm3-001,protein_coding,ENSMUSG00000039860.19,protein_coding | 59.08963204 | -0.65003645 | 0.000311359 | 0.003096761 |
| ENSMUST00000099356.8,Arrdc3-001,protein_coding,ENSMUSG00000074794.9,protein_coding | 2383.044342 | -0.230680934 | 0.000311812 | 0.00310054 |
| ENSMUST00000167149.7,Nfe2l1-203,protein_coding,ENSMUSG00000038615.17,protein_coding | 6527.456414 | 0.6588219 | 0.00031234 | 0.003105079 |
| ENSMUST00000123762.1,Tor1a-004,retained_intron,ENSMUSG00000026849.18,protein_coding | 193.4144191 | 0.564499578 | 0.00031325 | 0.003113405 |
| ENSMUST00000033934.4,Mrps31-001,protein_coding,ENSMUSG00000031533.4,protein_coding | 3660.719214 | 0.225957597 | 0.000313623 | 0.003116389 |
| ENSMUST00000156729.1,Crcp-002,processed_transcript,ENSMUSG00000025532.13,protein_coding | 219.6677083 | 0.458381176 | 0.000314149 | 0.003120902 |
| ENSMUST00000182793.7,Atg7-004,protein_coding,ENSMUSG00000030314.16,protein_coding | 1747.68436 | 0.260116483 | 0.000314922 | 0.003127857 |
| ENSMUST00000107423.1,Aen-003,protein_coding,ENSMUSG00000030609.18,protein_coding | 516.3677917 | -0.288066449 | 0.000315688 | 0.003134741 |
| ENSMUST00000025999.6,Add3-201,protein_coding,ENSMUSG00000025026.14,protein_coding | 440.3319009 | 0.638139242 | 0.000316082 | 0.003137934 |
| ENSMUST00000156263.1,Rad23b-003,processed_transcript,ENSMUSG00000028426.10,protein_coding | 5226.232013 | 0.17701665 | 0.000316402 | 0.00314038 |
| ENSMUST00000027425.15,Itm2c-001,protein_coding,ENSMUSG00000026223.15,protein_coding | 10596.74331 | 0.200016194 | 0.000316533 | 0.003140957 |
| ENSMUST00000130290.1,Nt5dc1-004,processed_transcript,ENSMUSG00000039480.14,protein_coding | 190.0804875 | 0.597797333 | 0.000317799 | 0.003152799 |
| ENSMUST00000003622.14,Slc25a1-001,protein_coding,ENSMUSG00000003528.14,protein_coding | 14481.88783 | 0.19992241 | 0.000318667 | 0.003160676 |
| ENSMUST00000014922.4,Fhod1-001,protein_coding,ENSMUSG00000014778.10,protein_coding | 4368.611821 | 0.123227002 | 0.000319593 | 0.003169128 |
| ENSMUST00000200988.1,Tigar-002,nonsense_mediated_decay,ENSMUSG00000038028.9,protein_coding | 192.9920787 | 0.527448108 | 0.000320813 | 0.0031805 |
| ENSMUST00000182659.7,Rbm5-001,protein_coding,ENSMUSG00000032580.12,protein_coding | 4017.877576 | 0.289387304 | 0.000323376 | 0.003205169 |
| ENSMUST00000217034.1,Ddx6-001,protein_coding,ENSMUSG00000032097.10,protein_coding | 20141.18209 | -0.188594432 | 0.000323537 | 0.003206021 |
| ENSMUST00000166615.2,Wwp2-001,protein_coding,ENSMUSG00000031930.11,protein_coding | 4910.901827 | -0.187090362 | 0.000324056 | 0.003210432 |
| ENSMUST00000150485.1,Rpl22-006,processed_transcript,ENSMUSG00000028936.15,protein_coding | 676.037424 | 0.315021723 | 0.000324551 | 0.003214592 |
| ENSMUST00000030154.6,Nudt2-001,protein_coding,ENSMUSG00000028443.6,protein_coding | 1775.671647 | 0.167667308 | 0.000324782 | 0.003216138 |
| ENSMUST00000139306.1,Pmepa1-002,protein_coding,ENSMUSG00000038400.15,protein_coding | 1305.779078 | -0.301215708 | 0.000325463 | 0.003221914 |
| ENSMUST00000192584.1,Ncam1-003,protein_coding,ENSMUSG00000039542.16,protein_coding | 289.9327953 | 0.508727232 | 0.000325515 | 0.003221914 |
| ENSMUST00000022147.14,Smn1-001,protein_coding,ENSMUSG00000021645.16,protein_coding | 6384.115957 | 0.162924931 | 0.000325696 | 0.003222969 |
| ENSMUST00000032557.14,Mark2-001,protein_coding,ENSMUSG00000024969.15,protein_coding | 5785.796883 | 0.170760637 | 0.000326261 | 0.003227813 |
| ENSMUST00000019649.3,Ubb-001,protein_coding,ENSMUSG00000019505.7,protein_coding | 158809.3217 | 0.196592997 | 0.000326725 | 0.003231663 |
| ENSMUST00000209812.1,Rpl13a-017,nonsense_mediated_decay,ENSMUSG00000074129.13,protein_coding | 15781.22036 | 0.184146581 | 0.000327977 | 0.0032433 |
| ENSMUST00000058407.5,Uqcr10-001,protein_coding,ENSMUSG00000059534.8,protein_coding | 7755.993895 | 0.165295129 | 0.000328309 | 0.003245838 |
| ENSMUST00000028063.6,Pter-002,protein_coding,ENSMUSG00000026730.12,protein_coding | 648.788392 | 0.3523561 | 0.000328604 | 0.003248008 |
| ENSMUST00000039517.10,Syde2-201,protein_coding,ENSMUSG00000036863.12,protein_coding | 905.5300645 | -0.369197423 | 0.000329261 | 0.003253753 |
| ENSMUST00000005120.11,Ccdc130-002,protein_coding,ENSMUSG00000004994.12,protein_coding | 753.6578954 | 0.332033485 | 0.000329853 | 0.003257844 |
| ENSMUST00000154584.7,Eef1d-001,protein_coding,ENSMUSG00000055762.15,protein_coding | 2149.036487 | -0.423533946 | 0.000329813 | 0.003257844 |
| ENSMUST00000100375.10,Poldip3-003,protein_coding,ENSMUSG00000041815.13,protein_coding | 4813.170484 | 0.331578689 | 0.000329902 | 0.003257844 |
| ENSMUST00000124659.1,Jup-003,processed_transcript,ENSMUSG00000001552.14,protein_coding | 73.16277048 | -0.579469427 | 0.000330075 | 0.003258809 |
| ENSMUST00000201902.3,Ccnd2-002,protein_coding,ENSMUSG00000000184.12,protein_coding | 2790.147748 | 0.263295473 | 0.000331014 | 0.003267331 |
| ENSMUST00000022586.1,Nufip1-201,protein_coding,ENSMUSG00000022009.1,protein_coding | 5561.316255 | 0.155747941 | 0.000332015 | 0.003276461 |
| ENSMUST00000199300.4,Camk2d-001,protein_coding,ENSMUSG00000053819.16,protein_coding | 1242.863296 | 0.247531733 | 0.00033275 | 0.003282956 |
| ENSMUST00000004050.6,Mmd-001,protein_coding,ENSMUSG00000003948.17,protein_coding | 12916.36894 | 0.200995378 | 0.000332865 | 0.003283342 |
| ENSMUST00000137615.7,Relb-004,retained_intron,ENSMUSG00000002983.17,protein_coding | 47.3686589 | -0.624470201 | 0.000334359 | 0.003297126 |
| ENSMUST00000113993.7,Itsn1-005,protein_coding,ENSMUSG00000022957.19,protein_coding | 1642.780101 | 0.366950366 | 0.000334416 | 0.003297126 |
| ENSMUST00000003035.11,Disp1-002,protein_coding,ENSMUSG00000030768.12,protein_coding | 3446.371341 | 0.186130723 | 0.00033458 | 0.003297996 |
| ENSMUST00000061882.8,Mcat-201,protein_coding,ENSMUSG00000048755.8,protein_coding | 3807.826651 | 0.187619414 | 0.000335089 | 0.003302256 |
| ENSMUST00000018965.3,Avpi1-001,protein_coding,ENSMUSG00000018821.3,protein_coding | 8380.024611 | 0.177191388 | 0.000335324 | 0.003303811 |
| ENSMUST00000027173.14,Wdr12-001,protein_coding,ENSMUSG00000026019.15,protein_coding | 495.1251889 | -0.269174395 | 0.000335554 | 0.003305319 |
| ENSMUST00000077853.3,Prpf4b-201,protein_coding,ENSMUSG00000021413.8,protein_coding | 4523.421503 | -0.40185922 | 0.000336165 | 0.003310584 |
| ENSMUST00000176535.7,Ect2-008,protein_coding,ENSMUSG00000027699.19,protein_coding | 150.8474455 | -0.648324408 | 0.000337258 | 0.003320583 |
| ENSMUST00000060805.6,Fam120a-201,protein_coding,ENSMUSG00000038014.7,protein_coding | 62755.19889 | 0.115161538 | 0.000337484 | 0.003322057 |
| ENSMUST00000214311.1,Kif15-008,nonsense_mediated_decay,ENSMUSG00000036768.6,protein_coding | 27.99609494 | -0.667743827 | 0.000338131 | 0.003327658 |
| ENSMUST00000136909.7,Acin1-016,processed_transcript,ENSMUSG00000022185.18,protein_coding | 96.89574155 | 0.56843487 | 0.000338388 | 0.003329424 |
| ENSMUST00000064795.5,Egr1-001,protein_coding,ENSMUSG00000038418.7,protein_coding | 1069.825773 | 0.336416543 | 0.00033862 | 0.003330948 |
| ENSMUST00000098858.10,Kcnu1-001,protein_coding,ENSMUSG00000031576.16,protein_coding | 249.2539461 | -0.433728561 | 0.00033924 | 0.003336289 |
| ENSMUST00000075954.7,Taf1b-201,protein_coding,ENSMUSG00000059669.7,protein_coding | 1698.677506 | 0.184816268 | 0.000339726 | 0.003340301 |
| ENSMUST00000138226.7,Anp32a-003,processed_transcript,ENSMUSG00000032249.14,protein_coding | 105.3790943 | 0.668228107 | 0.000340645 | 0.00334857 |
| ENSMUST00000107417.8,Shc1-003,protein_coding,ENSMUSG00000042626.13,protein_coding | 6891.403733 | 0.284494519 | 0.000341022 | 0.003351511 |
| ENSMUST00000212650.1,Acd-004,nonsense_mediated_decay,ENSMUSG00000038000.10,protein_coding | 1621.241997 | 0.245561538 | 0.000342828 | 0.003368496 |
| ENSMUST00000054459.10,Tmem201-006,protein_coding,ENSMUSG00000044700.15,protein_coding | 961.8166062 | 0.271175929 | 0.000343295 | 0.00337231 |
| ENSMUST00000188641.6,Tceb1-002,protein_coding,ENSMUSG00000079658.9,protein_coding | 1996.810428 | -0.3872398 | 0.000343506 | 0.003373615 |
| ENSMUST00000090227.4,March6-201,protein_coding,ENSMUSG00000039100.9,protein_coding | 18622.92744 | 0.198912547 | 0.000343627 | 0.003374034 |
| ENSMUST00000108593.7,Ctdnep1-001,protein_coding,ENSMUSG00000018559.16,protein_coding | 15799.43718 | 0.117105313 | 0.000343809 | 0.003375054 |
| ENSMUST00000039156.6,Phpt1-001,protein_coding,ENSMUSG00000036504.6,protein_coding | 5101.285007 | -0.184015246 | 0.000344278 | 0.003378111 |
| ENSMUST00000031707.13,Aass-001,protein_coding,ENSMUSG00000029695.13,protein_coding | 33.43288864 | -0.655096012 | 0.00034427 | 0.003378111 |
| ENSMUST00000135003.1,Zmym4-002,protein_coding,ENSMUSG00000042446.16,protein_coding | 925.5556569 | -0.503885571 | 0.000344661 | 0.003381105 |
| ENSMUST00000060251.7,Higd1a-001,protein_coding,ENSMUSG00000038412.8,protein_coding | 2817.463448 | 0.168346748 | 0.000345672 | 0.003390244 |
| ENSMUST00000036181.14,Car14-001,protein_coding,ENSMUSG00000038526.14,protein_coding | 40.75600999 | -0.652243304 | 0.000346041 | 0.003391545 |
| ENSMUST00000206349.1,Ino80e-014,protein_coding,ENSMUSG00000030689.15,protein_coding | 1344.330937 | 0.223477836 | 0.000345968 | 0.003391545 |
| ENSMUST00000023225.6,Grina-201,protein_coding,ENSMUSG00000022564.6,protein_coding | 22634.2441 | 0.168658018 | 0.000345927 | 0.003391545 |
| ENSMUST00000141274.1,Ndufa5-002,processed_transcript,ENSMUSG00000023089.12,protein_coding | 2510.568874 | -0.21094349 | 0.000346289 | 0.00339269 |
| ENSMUST00000117513.7,Slc29a3-001,protein_coding,ENSMUSG00000020100.15,protein_coding | 2455.260931 | 0.219828747 | 0.000346315 | 0.00339269 |
| ENSMUST00000179939.7,Atp2a2-003,nonsense_mediated_decay,ENSMUSG00000029467.15,protein_coding | 4216.712336 | -0.219034967 | 0.000346743 | 0.003395546 |
| ENSMUST00000081635.12,St7-010,protein_coding,ENSMUSG00000029534.17,protein_coding | 688.1838298 | 0.427358038 | 0.000346764 | 0.003395546 |
| ENSMUST00000016086.9,Gtf2ird2-001,protein_coding,ENSMUSG00000015942.9,protein_coding | 942.798745 | -0.213314772 | 0.000347003 | 0.003396677 |
| ENSMUST00000045054.10,Tcaf1-001,protein_coding,ENSMUSG00000036667.14,protein_coding | 1311.993971 | -0.370394647 | 0.000347038 | 0.003396677 |
| ENSMUST00000126897.1,Exosc10-004,retained_intron,ENSMUSG00000017264.16,protein_coding | 311.3843639 | 0.352686754 | 0.000347859 | 0.003403937 |
| ENSMUST00000125859.7,Chka-007,retained_intron,ENSMUSG00000024843.15,protein_coding | 248.055908 | 0.422750385 | 0.00034829 | 0.003407387 |
| ENSMUST00000102660.7,Sestd1-001,protein_coding,ENSMUSG00000042272.17,protein_coding | 3013.173717 | -0.310696787 | 0.000348853 | 0.003412119 |
| ENSMUST00000046389.4,Rbm45-001,protein_coding,ENSMUSG00000042369.8,protein_coding | 3105.276232 | -0.169077991 | 0.000348942 | 0.003412208 |
| ENSMUST00000043848.10,Ncapd2-001,protein_coding,ENSMUSG00000038252.13,protein_coding | 63780.66829 | 0.118039968 | 0.000349491 | 0.003416031 |
| ENSMUST00000139049.1,Prps1l3-001,protein_coding,ENSMUSG00000079104.4,protein_coding | 3187.078151 | -0.15998971 | 0.000349429 | 0.003416031 |
| ENSMUST00000029910.11,Nsmaf-001,protein_coding,ENSMUSG00000028245.15,protein_coding | 4774.625098 | 0.142910652 | 0.000349752 | 0.003417808 |
| ENSMUST00000150020.1,Huwe1-005,retained_intron,ENSMUSG00000025261.17,protein_coding | 1666.887194 | -0.222652108 | 0.000350659 | 0.00342589 |
| ENSMUST00000214361.1,Adpgk-003,retained_intron,ENSMUSG00000025236.11,protein_coding | 1488.664289 | 0.243718442 | 0.000352349 | 0.003441617 |
| ENSMUST00000026256.8,Fbxl15-201,protein_coding,ENSMUSG00000025226.11,protein_coding | 597.0353182 | -0.433149038 | 0.000353124 | 0.003448409 |
| ENSMUST00000128724.7,Iqsec2-005,processed_transcript,ENSMUSG00000041115.16,protein_coding | 162.9225133 | 0.467018704 | 0.000353564 | 0.003451919 |
| ENSMUST00000080856.13,Ipo11-001,protein_coding,ENSMUSG00000042590.16,protein_coding | 9013.954919 | -0.138300268 | 0.000354015 | 0.003455542 |
| ENSMUST00000141668.7,Hnrnph1-014,processed_transcript,ENSMUSG00000007850.16,protein_coding | 93.31776093 | -0.554603348 | 0.000354888 | 0.003463178 |
| ENSMUST00000144283.1,Lgals8-005,protein_coding,ENSMUSG00000057554.13,protein_coding | 430.791298 | 0.400887787 | 0.000354958 | 0.003463178 |
| ENSMUST00000110384.8,Fam65b-003,protein_coding,ENSMUSG00000036006.19,protein_coding | 2976.799111 | 0.194182887 | 0.000355216 | 0.003464904 |
| ENSMUST00000196698.4,Sfswap-002,nonsense_mediated_decay,ENSMUSG00000029439.14,protein_coding | 579.6994936 | 0.568210268 | 0.000355832 | 0.003470124 |
| ENSMUST00000209927.1,Rpl13a-025,protein_coding,ENSMUSG00000074129.13,protein_coding | 32348.4465 | 0.212738143 | 0.000356058 | 0.003471544 |

| | | | | |
|---|---|---|---|---|
| ENSMUST00000132734.7,Ppargc1a-001,protein_coding,ENSMUSG00000029167.13,protein_coding | 106.1143781 | -0.571590453 | 0.00035738 | 0.003483647 |
| ENSMUST00000026039.14,Gk-001,protein_coding,ENSMUSG00000025059.16,protein_coding | 432.1193392 | -0.47959952 | 0.000357969 | 0.003488594 |
| ENSMUST00000128889.1,Tpt1-004,retained_intron,ENSMUSG00000060126.14,protein_coding | 49.71786855 | -0.641281523 | 0.000359593 | 0.003503635 |
| ENSMUST00000040555.14,Rnf10-003,protein_coding,ENSMUSG00000041740.16,protein_coding | 6808.513158 | 0.156017842 | 0.00035985 | 0.003505219 |
| ENSMUST00000207889.1,Mical3-018,protein_coding,ENSMUSG00000051586.16,protein_coding | 6685.876557 | 0.140212742 | 0.000359934 | 0.003505219 |
| ENSMUST00000141268.7,Eef1d-010,protein_coding,ENSMUSG00000055762.15,protein_coding | 7160.429056 | 0.161955068 | 0.00036 | 0.003505219 |
| ENSMUST00000030915.10,Morn1-001,protein_coding,ENSMUSG00000029049.14,protein_coding | 74.81687691 | 0.577085069 | 0.000360694 | 0.003511177 |
| ENSMUST00000110708.2,Srp54b-201,protein_coding,ENSMUSG00000079108.5,protein_coding | 6390.061217 | -0.328397937 | 0.000360927 | 0.003512651 |
| ENSMUST00000036328.8,Zfhx2-001,protein_coding,ENSMUSG00000040721.9,protein_coding | 645.5369509 | -0.259063105 | 0.000361015 | 0.003512717 |
| ENSMUST00000027153.5,Acadl-001,protein_coding,ENSMUSG00000026003.5,protein_coding | 10632.46698 | 0.111823112 | 0.000361268 | 0.00351438 |
| ENSMUST00000049412.11,Stau1-005,protein_coding,ENSMUSG00000039536.16,protein_coding | 1462.748575 | 0.315781292 | 0.00036183 | 0.003519059 |
| ENSMUST00000109110.3,Rab22a-003,protein_coding,ENSMUSG00000027519.10,protein_coding | 1354.762676 | 0.175429883 | 0.000362045 | 0.003520354 |
| ENSMUST00000207892.1,Slc9a6-005,protein_coding,ENSMUSG00000060681.15,protein_coding | 87.65981965 | 0.648920802 | 0.000362147 | 0.003520548 |
| ENSMUST00000078528.6,C1qbp-001,protein_coding,ENSMUSG00000018446.8,protein_coding | 40167.34175 | 0.155847752 | 0.000362245 | 0.003520706 |
| ENSMUST00000173665.7,Trim39-004,nonsense_mediated_decay,ENSMUSG00000045409.16,protein_coding | 94.12164229 | -0.66219592 | 0.00036401 | 0.003537063 |
| ENSMUST00000030622.2,Syf2-001,protein_coding,ENSMUSG00000028821.8,protein_coding | 4091.963898 | 0.220828205 | 0.000364315 | 0.003539225 |
| ENSMUST00000019276.10,BC005537-001,protein_coding,ENSMUSG00000019132.10,protein_coding | 15890.63302 | 0.12723871 | 0.0003647 | 0.003542165 |
| ENSMUST00000030361.10,Tmem59-001,protein_coding,ENSMUSG00000028618.11,protein_coding | 8588.557781 | 0.212006395 | 0.000365174 | 0.00354597 |
| ENSMUST00000041361.13,Zfp335-001,protein_coding,ENSMUSG00000039834.17,protein_coding | 6101.142972 | 0.135029977 | 0.000365408 | 0.003547438 |
| ENSMUST00000093801.9,Plscr1-001,protein_coding,ENSMUSG00000032369.13,protein_coding | 1859.544336 | 0.237138707 | 0.00036626 | 0.003552843 |
| ENSMUST00000087399.4,Tnxb-001,protein_coding,ENSMUSG00000033327.16,protein_coding | 684.9637463 | 0.375075115 | 0.00036618 | 0.003552843 |
| ENSMUST00000033506.12,Wdr13-001,protein_coding,ENSMUSG00000031166.13,protein_coding | 1830.202147 | 0.20022363 | 0.00036625 | 0.003552843 |
| ENSMUST00000082177.12,Kdm5c-004,protein_coding,ENSMUSG00000025332.14,protein_coding | 5224.122921 | 0.203474564 | 0.000366295 | 0.003552843 |
| ENSMUST00000080353.2,Ing2-001,protein_coding,ENSMUSG00000063049.4,protein_coding | 1377.953832 | 0.262972054 | 0.000366445 | 0.003553501 |
| ENSMUST00000133117.7,Plpp5-004,nonsense_mediated_decay,ENSMUSG00000031570.17,protein_coding | 671.0442759 | 0.429711737 | 0.000366767 | 0.003555824 |
| ENSMUST00000118047.1,Gm14681-001,processed_pseudogene,ENSMUSG00000081603.1,processed_pseudogene | 678.5842923 | -0.434682347 | 0.000367776 | 0.003564806 |
| ENSMUST00000118307.1,Gm12191-001,processed_pseudogene,ENSMUSG00000083061.1,processed_pseudogene | 371.8417578 | -0.396568296 | 0.000368254 | 0.003568633 |
| ENSMUST00000059231.3,Frat2-001,protein_coding,ENSMUSG00000047604.3,protein_coding | 463.5610012 | 0.324485121 | 0.000368977 | 0.003574832 |
| ENSMUST00000114265.8,Clic3-001,protein_coding,ENSMUSG00000015093.15,protein_coding | 69.10171469 | 0.588194906 | 0.000369213 | 0.003575492 |
| ENSMUST00000144811.1,RP23-33N12.2-003,processed_transcript,ENSMUSG00000069755.5,transcribed_unprocessed_pseudogene | 116.5301683 | -0.472267892 | 0.000369226 | 0.003575492 |
| ENSMUST00000184744.1,Kctd12-001,protein_coding,ENSMUSG00000098557.1,protein_coding | 71.26060741 | -0.578148608 | 0.000369294 | 0.003575492 |
| ENSMUST00000029201.13,Pik3ca-201,protein_coding,ENSMUSG00000027665.13,protein_coding | 131.9292099 | -0.550485195 | 0.000369711 | 0.003577921 |
| ENSMUST00000138401.1,Dek-003,processed_transcript,ENSMUSG00000021377.14,protein_coding | 115.6990559 | -0.642233842 | 0.000369703 | 0.003577921 |
| ENSMUST00000168463.7,Cox16-009,protein_coding,ENSMUSG00000091803.7,protein_coding | 263.6750435 | 0.531622597 | 0.000369948 | 0.003579406 |
| ENSMUST00000033006.13,Nsmce1-001,protein_coding,ENSMUSG00000030750.13,protein_coding | 4980.539885 | 0.192973581 | 0.000370494 | 0.003583881 |
| ENSMUST00000034989.14,Me1-001,protein_coding,ENSMUSG00000032418.15,protein_coding | 21627.95753 | 0.185410969 | 0.000370837 | 0.003586394 |
| ENSMUST00000150541.1,Timm17b-003,retained_intron,ENSMUSG00000031158.11,protein_coding | 1611.23277 | 0.210061089 | 0.000371466 | 0.003591671 |
| ENSMUST00000106447.8,Eif2b3-201,protein_coding,ENSMUSG00000028683.14,protein_coding | 694.9782365 | 0.241191192 | 0.000372193 | 0.003597895 |
| ENSMUST00000028921.5,Xrn2-001,protein_coding,ENSMUSG00000027433.5,protein_coding | 18685.56248 | 0.143560334 | 0.000372478 | 0.003599839 |
| ENSMUST00000006956.8,Saa3-001,protein_coding,ENSMUSG00000040026.8,protein_coding | 35.28804183 | 0.6635983 | 0.000373612 | 0.003609993 |
| ENSMUST00000172088.7,Zfp239-201,protein_coding,ENSMUSG00000042097.17,protein_coding | 923.0569538 | -0.203532131 | 0.000374307 | 0.003615894 |
| ENSMUST00000025921.14,Mark2-006,protein_coding,ENSMUSG00000024969.15,protein_coding | 997.6044193 | -0.257339757 | 0.00037451 | 0.00361704 |
| ENSMUST00000109266.10,Zmynd8-201,protein_coding,ENSMUSG00000039671.18,protein_coding | 2280.47548 | -0.438064171 | 0.000374628 | 0.003617371 |
| ENSMUST00000170078.7,Myg1-002,retained_intron,ENSMUSG00000001285.12,protein_coding | 518.8541462 | 0.526084948 | 0.000375371 | 0.003623728 |
| ENSMUST00000185644.6,Tfdp2-008,protein_coding,ENSMUSG00000032411.15,protein_coding | 112.934659 | 0.659181999 | 0.000375685 | 0.003625948 |
| ENSMUST00000195059.5,Enah-004,protein_coding,ENSMUSG00000022995.16,protein_coding | 517.4367142 | 0.588958334 | 0.000375932 | 0.003626992 |
| ENSMUST00000147563.8,Zdhhc3-003,protein_coding,ENSMUSG00000025786.16,protein_coding | 2096.663157 | -0.194156663 | 0.000375961 | 0.003626992 |
| ENSMUST00000079223.10,Fggy-001,protein_coding,ENSMUSG00000028573.18,protein_coding | 473.8157503 | 0.282881334 | 0.000377062 | 0.003636419 |
| ENSMUST00000167731.7,Myc-006,protein_coding,ENSMUSG00000022346.15,protein_coding | 5629.242804 | 0.166661865 | 0.000377108 | 0.003636419 |
| ENSMUST00000162163.7,Gas5-006,retained_intron,ENSMUSG00000053332.13,processed_transcript | 784.2624688 | -0.497179868 | 0.000377477 | 0.003639168 |
| ENSMUST00000130048.7,Vegfb-002,protein_coding,ENSMUSG00000024962.13,protein_coding | 2055.574325 | 0.267001451 | 0.000377621 | 0.003639743 |
| ENSMUST00000184770.7,Tcf12-009,retained_intron,ENSMUSG00000032228.16,protein_coding | 80.45885194 | -0.5766807 | 0.000379949 | 0.003661362 |
| ENSMUST00000025290.5,Impact-201,protein_coding,ENSMUSG00000024423.5,protein_coding | 1922.827547 | -0.300133625 | 0.000380322 | 0.003664134 |
| ENSMUST00000045689.13,Mrpl15-003,nonsense_mediated_decay,ENSMUSG00000033845.13,protein_coding | 795.8260568 | 0.225065184 | 0.000381041 | 0.003670186 |
| ENSMUST00000119081.1,Mcmbp-002,protein_coding,ENSMUSG00000048170.14,protein_coding | 5276.471788 | 0.187946023 | 0.000381121 | 0.003670186 |
| ENSMUST00000168499.8,Nlrx1-004,protein_coding,ENSMUSG00000032109.15,protein_coding | 593.9966087 | 0.534074138 | 0.00038205 | 0.003678312 |
| ENSMUST00000101004.8,Per1-003,protein_coding,ENSMUSG00000020893.17,protein_coding | 843.8387259 | 0.208261443 | 0.000384188 | 0.003697357 |
| ENSMUST00000166977.8,Pla2g6-002,protein_coding,ENSMUSG00000042632.16,protein_coding | 2476.088029 | 0.205128263 | 0.0003842 | 0.003697357 |
| ENSMUST00000108246.8,Wwp1-001,protein_coding,ENSMUSG00000041058.15,protein_coding | 2831.822612 | -0.429835557 | 0.000384801 | 0.003702312 |
| ENSMUST00000105417.9,Dip2a-201,protein_coding,ENSMUSG00000020231.15,protein_coding | 658.3751095 | 0.528399419 | 0.000385253 | 0.003705003 |
| ENSMUST00000035444.9,Chrm1-001,protein_coding,ENSMUSG00000032773.9,protein_coding | 31.66566069 | -0.660169412 | 0.000385188 | 0.003705003 |
| ENSMUST00000122402.2,Farp2-002,protein_coding,ENSMUSG00000034066.13,protein_coding | 1423.766546 | -0.314211574 | 0.000385927 | 0.003710654 |
| ENSMUST00000105387.7,Agpat3-004,protein_coding,ENSMUSG00000001211.14,protein_coding | 5614.703541 | 0.197799321 | 0.000386161 | 0.003712078 |
| ENSMUST00000214659.1,Cdk19-003,nonsense_mediated_decay,ENSMUSG00000038481.13,protein_coding | 969.6704187 | 0.392871213 | 0.000386495 | 0.003714461 |
| ENSMUST00000069098.6,Soga1-001,protein_coding,ENSMUSG00000055485.6,protein_coding | 7199.868401 | -0.124409639 | 0.000387095 | 0.003718721 |
| ENSMUST00000207106.1,Picalm-010,retained_intron,ENSMUSG00000039361.11,protein_coding | 232.3945758 | -0.555672462 | 0.000387174 | 0.003718721 |
| ENSMUST00000039610.9,Xylb-001,protein_coding,ENSMUSG00000035769.9,protein_coding | 429.5913581 | -0.457842176 | 0.000387198 | 0.003718721 |
| ENSMUST00000033444.10,Stk26-001,protein_coding,ENSMUSG00000031112.10,protein_coding | 1072.86472 | -0.469742629 | 0.000387715 | 0.003722864 |
| ENSMUST00000027056.11,Prex2-001,protein_coding,ENSMUSG00000048960.13,protein_coding | 62.95986015 | -0.630108769 | 0.000387933 | 0.003724124 |
| ENSMUST00000069965.8,Cdk17-002,protein_coding,ENSMUSG00000020015.10,protein_coding | 4083.754842 | 0.124325075 | 0.000388062 | 0.003724527 |
| ENSMUST00000124949.1,Tasp1-003,retained_intron,ENSMUSG00000039033.11,protein_coding | 238.7945775 | -0.36042111 | 0.000388455 | 0.003727472 |
| ENSMUST00000182481.7,Plce1-002,protein_coding,ENSMUSG00000024998.17,protein_coding | 2911.717593 | -0.213774288 | 0.000388794 | 0.003729892 |
| ENSMUST00000182017.1,Rps13-ps2-001,processed_pseudogene,ENSMUSG00000069972.6,processed_pseudogene | 275.2294916 | -0.42866738 | 0.000389428 | 0.003735145 |
| ENSMUST00000204696.2,Smarcad1-003,protein_coding,ENSMUSG00000029920.9,protein_coding | 207.5982364 | 0.540493589 | 0.000390566 | 0.003745222 |
| ENSMUST00000069097.12,Ago3-001,protein_coding,ENSMUSG00000028842.15,protein_coding | 3548.827634 | -0.448875314 | 0.000391016 | 0.003748702 |
| ENSMUST00000116307.7,Echdc2-003,protein_coding,ENSMUSG00000028601.18,protein_coding | 51.79513068 | -0.647722836 | 0.000392416 | 0.003761289 |
| ENSMUST00000162240.1,Opa1-007,protein_coding,ENSMUSG00000038084.16,protein_coding | 167.8939483 | 0.59756188 | 0.000392754 | 0.003763689 |
| ENSMUST00000186889.6,Cacna1c-001,protein_coding,ENSMUSG00000051331.14,protein_coding | 100.5442893 | -0.513514792 | 0.00039339 | 0.003768944 |
| ENSMUST00000026225.14,Sema4g-001,protein_coding,ENSMUSG00000025207.15,protein_coding | 222.9979799 | -0.557739484 | 0.000394365 | 0.003777449 |
| ENSMUST00000159879.1,Trove2-001,protein_coding,ENSMUSG00000018199.8,protein_coding | 5570.272059 | -0.465837699 | 0.000394863 | 0.003781376 |
| ENSMUST00000115259.2,Tmem60-001,protein_coding,ENSMUSG00000045435.9,protein_coding | 1123.481868 | -0.200576352 | 0.000395715 | 0.003788412 |
| ENSMUST00000178136.7,Abcc3-201,protein_coding,ENSMUSG00000020865.16,protein_coding | 3689.766666 | 0.292598723 | 0.000395774 | 0.003788412 |
| ENSMUST00000189728.6,Thap4-006,protein_coding,ENSMUSG00000026279.14,protein_coding | 591.1037762 | 0.403804498 | 0.000396804 | 0.003796979 |
| ENSMUST00000168276.7,Atp8b2-006,protein_coding,ENSMUSG00000060671.12,protein_coding | 1556.312333 | -0.415745636 | 0.000396845 | 0.003796979 |
| ENSMUST00000185483.6,Nemp2-001,nonsense_mediated_decay,ENSMUSG00000043015.15,protein_coding | 153.2824174 | 0.556924422 | 0.000397934 | 0.003806553 |
| ENSMUST00000084264.11,Txlna-001,protein_coding,ENSMUSG00000053841.15,protein_coding | 2791.169381 | -0.35017508 | 0.000398128 | 0.003807558 |
| ENSMUST00000109366.7,Fbxw11-001,protein_coding,ENSMUSG00000020271.17,protein_coding | 9558.420262 | 0.153051178 | 0.000398324 | 0.003808584 |
| ENSMUST00000118733.7,Ank1-001,protein_coding,ENSMUSG00000031543.18,protein_coding | 465.2788648 | -0.341702572 | 0.000399412 | 0.00381814 |
| ENSMUST00000115511.8,Phf14-006,protein_coding,ENSMUSG00000029629.17,protein_coding | 4698.086726 | -0.266384278 | 0.00039975 | 0.003819456 |
| ENSMUST00000171057.1,Immt-008,protein_coding,ENSMUSG00000052337.15,protein_coding | 318.0091392 | -0.387915223 | 0.0003998 | 0.003819456 |
| ENSMUST00000187814.6,Cipc-001,protein_coding,ENSMUSG00000034157.12,protein_coding | 1081.102891 | -0.242395168 | 0.000399904 | 0.003819456 |
| ENSMUST00000038369.10,Cipc-201,protein_coding,ENSMUSG00000034157.12,protein_coding | 1081.102891 | -0.242395168 | 0.000399904 | 0.003819456 |

| | | | | |
|---|---|---|---|---|
| ENSMUST00000130942.7,Echdc2-004,nonsense_mediated_decay,ENSMUSG00000028601.18,protein_coding | 202.437761 | 0.485346371 | 0.000400699 | 0.003826194 |
| ENSMUST00000006035.12,Ergic3-001,protein_coding,ENSMUSG00000005881.13,protein_coding | 13897.68037 | 0.1451983 | 0.000401101 | 0.003828334 |
| ENSMUST00000004206.9,Eif3g-001,protein_coding,ENSMUSG00000070319.6,protein_coding | 21513.73045 | 0.157614954 | 0.000401094 | 0.003828334 |
| ENSMUST00000027532.12,Scly-001,protein_coding,ENSMUSG00000026307.12,protein_coding | 1567.95519 | 0.207918866 | 0.000402049 | 0.003834837 |
| ENSMUST00000141283.7,Mthfr-002,protein_coding,ENSMUSG00000029009.17,protein_coding | 391.1039374 | 0.395384457 | 0.000401886 | 0.003834837 |
| ENSMUST00000035813.8,Ppp1r9a-001,protein_coding,ENSMUSG00000032827.15,protein_coding | 93.94558861 | -0.567004419 | 0.000402028 | 0.003834837 |
| ENSMUST00000019944.8,Adat2-001,protein_coding,ENSMUSG00000019808.8,protein_coding | 960.0836038 | 0.235976463 | 0.000402288 | 0.003836262 |
| ENSMUST00000189552.1,Prim2-002,retained_intron,ENSMUSG00000026134.11,protein_coding | 29.14247369 | -0.660354881 | 0.00040384 | 0.003850212 |
| ENSMUST00000006496.14,Rps9-001,protein_coding,ENSMUSG00000006333.14,protein_coding | 99585.78629 | 0.164568768 | 0.000404927 | 0.003859722 |
| ENSMUST00000170908.7,Dtwd1-201,protein_coding,ENSMUSG00000023330.12,protein_coding | 2986.39849 | 0.34643196 | 0.000405061 | 0.003860144 |
| ENSMUST00000020403.5,Csrp2-201,protein_coding,ENSMUSG00000020186.6,protein_coding | 1084.695946 | 0.204713259 | 0.000406459 | 0.003872606 |
| ENSMUST00000037609.7,Hgsnat-001,protein_coding,ENSMUSG00000037260.8,protein_coding | 5640.902928 | 0.14931229 | 0.000406837 | 0.003875351 |
| ENSMUST00000166312.7,Luc7l3-004,protein_coding,ENSMUSG00000020863.15,protein_coding | 674.1236289 | -0.506933856 | 0.000407301 | 0.003878911 |
| ENSMUST00000155910.2,Peg3-003,retained_intron,ENSMUSG00000002265.16,protein_coding | 75.21877712 | -0.604399962 | 0.000407473 | 0.003879695 |
| ENSMUST00000029316.15,Exosc8-001,protein_coding,ENSMUSG00000027752.15,protein_coding | 7977.692828 | 0.121505204 | 0.000407617 | 0.003879748 |
| ENSMUST00000108496.8,Slc1a5-001,protein_coding,ENSMUSG00000001918.17,protein_coding | 8741.838876 | 0.316511175 | 0.000407659 | 0.003879748 |
| ENSMUST00000204881.2,Isy1-005,nonsense_mediated_decay,ENSMUSG00000030056.8,protein_coding | 2696.06934 | 0.203208659 | 0.000408674 | 0.003888548 |
| ENSMUST00000162512.7,Gtpbp8-001,protein_coding,ENSMUSG00000022668.10,protein_coding | 793.6562475 | -0.251050071 | 0.000408799 | 0.003888875 |
| ENSMUST00000023171.7,Hes1-001,protein_coding,ENSMUSG00000022528.7,protein_coding | 2054.482903 | -0.294491987 | 0.000409526 | 0.003894932 |
| ENSMUST00000068333.13,Farsb-001,protein_coding,ENSMUSG00000026245.16,protein_coding | 17525.66318 | 0.134196621 | 0.000411172 | 0.003909719 |
| ENSMUST00000122242.7,Tor3a-006,nonsense_mediated_decay,ENSMUSG00000060519.11,protein_coding | 297.5472003 | 0.451798675 | 0.000411692 | 0.003912934 |
| ENSMUST00000188964.6,Tor3a-201,protein_coding,ENSMUSG00000060519.11,protein_coding | 297.5472003 | 0.451798675 | 0.000411692 | 0.003912934 |
| ENSMUST00000192147.1,Gm37691-001,TEC,ENSMUSG00000104348.1,TEC | 43.4652319 | 0.647585368 | 0.000412216 | 0.003917054 |
| ENSMUST00000082094.4,Ptcd3-001,protein_coding,ENSMUSG00000063884.6,protein_coding | 13427.27282 | -0.158430829 | 0.000412476 | 0.003917789 |
| ENSMUST00000004756.13,Wwox-001,protein_coding,ENSMUSG00000004637.15,protein_coding | 801.7180964 | 0.318269261 | 0.000412386 | 0.003917789 |
| ENSMUST00000018549.7,Mrm1-001,protein_coding,ENSMUSG00000018405.7,protein_coding | 1216.371113 | 0.305078992 | 0.00041387 | 0.00393017 |
| ENSMUST00000028351.8,Dph7-001,protein_coding,ENSMUSG00000026975.10,protein_coding | 2157.910869 | 0.233561084 | 0.000414388 | 0.003934216 |
| ENSMUST00000099626.4,Fibin-001,protein_coding,ENSMUSG00000074971.4,protein_coding | 99.82035626 | 0.621090475 | 0.000416063 | 0.003949253 |
| ENSMUST00000153107.1,Cep41-008,retained_intron,ENSMUSG00000029790.16,protein_coding | 43.53972341 | 0.658875505 | 0.000416261 | 0.003950129 |
| ENSMUST00000204545.2,Mpp6-009,protein_coding,ENSMUSG00000038388.14,protein_coding | 1339.877343 | -0.242142284 | 0.000416339 | 0.003950129 |
| ENSMUST00000031841.8,Tra2a-004,protein_coding,ENSMUSG00000029817.11,protein_coding | 7528.145904 | -0.25717017 | 0.000416453 | 0.003950341 |
| ENSMUST00000036951.12,Pebp1-001,protein_coding,ENSMUSG00000032959.12,protein_coding | 60175.83216 | 0.173760609 | 0.000416605 | 0.003950912 |
| ENSMUST00000151689.7,Magt1-001,protein_coding,ENSMUSG00000031232.16,protein_coding | 4850.619987 | -0.284034132 | 0.000416776 | 0.003951663 |
| ENSMUST00000131779.7,Pank2-002,processed_transcript,ENSMUSG00000037514.17,protein_coding | 178.7325146 | 0.656463296 | 0.000417075 | 0.003953627 |
| ENSMUST00000111697.8,Ctnnd1-202,protein_coding,ENSMUSG00000034101.14,protein_coding | 1675.745499 | 0.481442816 | 0.000417751 | 0.00395916 |
| ENSMUST00000106116.1,Ncdn-002,protein_coding,ENSMUSG00000028833.13,protein_coding | 129.0521111 | 0.658422241 | 0.000418818 | 0.003966804 |
| ENSMUST00000006397.6,Epor-001,protein_coding,ENSMUSG00000006235.6,protein_coding | 157.7259945 | -0.464524348 | 0.000418814 | 0.003966804 |
| ENSMUST00000095347.11,Brd2-005,protein_coding,ENSMUSG00000024335.19,protein_coding | 13852.72497 | 0.203575292 | 0.000418834 | 0.003966804 |
| ENSMUST00000126211.1,Snrpa-003,protein_coding,ENSMUSG00000061479.15,protein_coding | 2067.782854 | 0.291507321 | 0.000419326 | 0.003970343 |
| ENSMUST00000034012.9,Cnot7-001,protein_coding,ENSMUSG00000031601.16,protein_coding | 6905.224007 | -0.255337146 | 0.000419392 | 0.003970343 |
| ENSMUST00000172607.1,Calu-010,protein_coding,ENSMUSG00000029767.16,protein_coding | 44.01418061 | 0.658240748 | 0.000419958 | 0.00397395 |
| ENSMUST00000108479.1,Dmwd-003,protein_coding,ENSMUSG00000030410.16,protein_coding | 4392.0736 | 0.15846656 | 0.000419909 | 0.00397395 |
| ENSMUST00000081291.12,Cyhr1-001,protein_coding,ENSMUSG00000053929.16,protein_coding | 6696.771025 | -0.150743958 | 0.000420586 | 0.00397902 |
| ENSMUST00000112925.7,Ift122-001,protein_coding,ENSMUSG00000030323.13,protein_coding | 2944.092949 | 0.256227374 | 0.000420716 | 0.003979373 |
| ENSMUST00000110110.8,Zfp169-201,protein_coding,ENSMUSG00000050954.15,protein_coding | 54.37948911 | -0.645298216 | 0.000421221 | 0.003983278 |
| ENSMUST00000025453.14,Smad2-002,protein_coding,ENSMUSG00000024563.15,protein_coding | 1031.642358 | 0.50021564 | 0.000421734 | 0.003987256 |
| ENSMUST00000129233.7,Rnf181-006,nonsense_mediated_decay,ENSMUSG00000055850.12,protein_coding | 128.3141248 | 0.540985766 | 0.000422365 | 0.003992339 |
| ENSMUST00000099546.5,Chst14-001,protein_coding,ENSMUSG00000074916.5,protein_coding | 3214.86519 | 0.139398826 | 0.000423353 | 0.004000806 |
| ENSMUST00000156505.7,Fance-008,protein_coding,ENSMUSG00000007570.15,protein_coding | 68.51873205 | 0.597771061 | 0.000424341 | 0.004009258 |
| ENSMUST00000090993.7,Arpc5l-004,nonsense_mediated_decay,ENSMUSG00000026755.14,protein_coding | 453.5742063 | 0.363103228 | 0.000424717 | 0.004011935 |
| ENSMUST00000102659.1,Sestd1-002,protein_coding,ENSMUSG00000042272.17,protein_coding | 3136.684087 | 0.148484998 | 0.000425219 | 0.004015793 |
| ENSMUST00000196173.1,Gm43096-001,processed_pseudogene,ENSMUSG00000104882.1,processed_pseudogene | 23.61281497 | -0.655218855 | 0.000425407 | 0.004016684 |
| ENSMUST00000030568.13,Sema3c-001,protein_coding,ENSMUSG00000028780.13,protein_coding | 8134.476527 | -0.351511318 | 0.000425625 | 0.004017866 |
| ENSMUST00000093490.8,Clcn3-001,protein_coding,ENSMUSG00000004319.15,protein_coding | 3701.283012 | -0.343156204 | 0.000425724 | 0.004017917 |
| ENSMUST00000202907.1,Phf19-005,protein_coding,ENSMUSG00000026873.9,protein_coding | 19.98874025 | 0.65732406 | 0.000425884 | 0.004018547 |
| ENSMUST00000102726.7,Cobll1-002,protein_coding,ENSMUSG00000034903.18,protein_coding | 3538.509917 | -0.332631745 | 0.000426613 | 0.00402454 |
| ENSMUST00000123765.7,Macf1-003,protein_coding,ENSMUSG00000028649.17,protein_coding | 688.163358 | -0.465229892 | 0.000426967 | 0.004027001 |
| ENSMUST00000046288.14,Ndufv3-001,protein_coding,ENSMUSG00000024038.16,protein_coding | 2014.290213 | 0.183898997 | 0.000428005 | 0.004035901 |
| ENSMUST00000202991.3,Ube2a-002,protein_coding,ENSMUSG00000016308.12,protein_coding | 1826.533991 | 0.218177105 | 0.000428516 | 0.004039839 |
| ENSMUST00000133316.7,Cenpa-002,nonsense_mediated_decay,ENSMUSG00000029177.9,protein_coding | 10643.54219 | 0.181598307 | 0.000428987 | 0.004043388 |
| ENSMUST00000150330.7,Necap2-002,processed_transcript,ENSMUSG00000028923.14,protein_coding | 106.7783524 | 0.625161439 | 0.000429563 | 0.004047939 |
| ENSMUST00000052068.10,Rbp1-001,protein_coding,ENSMUSG00000046402.10,protein_coding | 2640.7917 | 0.257040662 | 0.00043092 | 0.004059837 |
| ENSMUST00000125918.1,Tmem94-007,protein_coding,ENSMUSG00000020747.17,protein_coding | 302.6598392 | 0.423955286 | 0.000431145 | 0.004061066 |
| ENSMUST00000187834.1,Rbbp5-006,retained_intron,ENSMUSG00000026439.14,protein_coding | 564.7864077 | 0.408741191 | 0.000431586 | 0.004064334 |
| ENSMUST00000022980.3,Ndufb9-201,protein_coding,ENSMUSG00000022354.3,protein_coding | 16512.87785 | 0.18178622 | 0.000432388 | 0.004070994 |
| ENSMUST00000201992.1,Gm43597-001,lincRNA,ENSMUSG00000107096.3,lincRNA | 88.43037015 | -0.564936377 | 0.0004333 | 0.004078115 |
| ENSMUST00000046950.12,Phf8-001,protein_coding,ENSMUSG00000041229.15,protein_coding | 2229.773042 | 0.171022419 | 0.000433334 | 0.004078115 |
| ENSMUST00000152479.1,Anks6-002,processed_transcript,ENSMUSG00000066191.11,protein_coding | 93.64659945 | -0.525599289 | 0.00043438 | 0.004087065 |
| ENSMUST00000022504.11,Mapk8-002,protein_coding,ENSMUSG00000021936.13,protein_coding | 1892.08942 | 0.189417999 | 0.000434738 | 0.004089538 |
| ENSMUST00000030074.7,Ugcg-001,protein_coding,ENSMUSG00000028381.8,protein_coding | 4679.851786 | -0.221350301 | 0.000435133 | 0.004092357 |
| ENSMUST00000131696.2,Nol9-002,nonsense_mediated_decay,ENSMUSG00000028948.16,protein_coding | 163.9134589 | -0.633271079 | 0.00043867 | 0.004123821 |
| ENSMUST00000202926.1,Gm43844-001,TEC,ENSMUSG00000106858.1,TEC | 49.274916 | -0.618632746 | 0.000438596 | 0.004123821 |
| ENSMUST00000179297.2,Prpf19-002,protein_coding,ENSMUSG00000024735.13,protein_coding | 1058.302357 | 0.306942515 | 0.000439615 | 0.004131803 |
| ENSMUST00000129993.2,Fam188a-007,protein_coding,ENSMUSG00000026767.12,protein_coding | 94.18332924 | 0.544483182 | 0.000440304 | 0.004136584 |
| ENSMUST00000166917.1,Oxr1-205,protein_coding,ENSMUSG00000022307.15,protein_coding | 203.5866838 | -0.538908849 | 0.000440316 | 0.004136584 |
| ENSMUST00000070143.12,Zhx1-001,protein_coding,ENSMUSG00000022361.14,protein_coding | 2633.153996 | -0.38374209 | 0.000440554 | 0.004137923 |
| ENSMUST00000199875.1,Zcchc8-002,retained_intron,ENSMUSG00000029427.11,protein_coding | 2347.222489 | 0.214216702 | 0.000441944 | 0.004150066 |
| ENSMUST00000098025.10,Srcap-202,protein_coding,ENSMUSG00000053877.12,protein_coding | 643.2283434 | -0.334374325 | 0.000442618 | 0.004155493 |
| ENSMUST00000046186.7,Spon2-001,protein_coding,ENSMUSG00000037379.7,protein_coding | 246.0415265 | -0.443726139 | 0.000443612 | 0.00416392 |
| ENSMUST00000135144.1,Gm15998-001,antisense,ENSMUSG00000086389.1,antisense | 41.57447875 | -0.609961882 | 0.000443995 | 0.004166604 |
| ENSMUST00000040999.13,Aox3-001,protein_coding,ENSMUSG00000064294.12,protein_coding | 357.1470585 | -0.3336105 | 0.000444807 | 0.004173312 |
| ENSMUST00000201266.3,Ube2k-003,protein_coding,ENSMUSG00000029203.16,protein_coding | 400.1650929 | 0.641726896 | 0.000444952 | 0.004173766 |
| ENSMUST00000143082.3,Saal1-001,protein_coding,ENSMUSG00000006763.14,protein_coding | 489.3405785 | -0.382241289 | 0.000446117 | 0.004183778 |
| ENSMUST00000044660.5,Armc5-001,protein_coding,ENSMUSG00000042178.6,protein_coding | 1910.133463 | -0.160439241 | 0.000447791 | 0.004197654 |
| ENSMUST00000112437.7,Atl2-202,protein_coding,ENSMUSG00000059811.12,protein_coding | 309.2583418 | -0.418911914 | 0.000447714 | 0.004197654 |
| ENSMUST00000094695.11,Rmdn3-001,protein_coding,ENSMUSG00000070730.11,protein_coding | 4659.159869 | 0.133383862 | 0.000448193 | 0.004200062 |
| ENSMUST00000051065.5,Gprin3-001,protein_coding,ENSMUSG00000045441.5,protein_coding | 98.63097964 | -0.574589615 | 0.000448243 | 0.004200062 |
| ENSMUST00000031486.13,Prkab1-001,protein_coding,ENSMUSG00000029513.14,protein_coding | 4037.902439 | 0.281178081 | 0.000448407 | 0.004200485 |
| ENSMUST00000111835.2,Styx-201,protein_coding,ENSMUSG00000053205.8,protein_coding | 292.8605389 | -0.528671163 | 0.000448484 | 0.004200485 |
| ENSMUST00000079634.7,Exosc5-001,protein_coding,ENSMUSG00000061286.7,protein_coding | 2846.659305 | 0.209258958 | 0.000448738 | 0.004201078 |
| ENSMUST00000020034.5,Echdc1-001,protein_coding,ENSMUSG00000019883.11,protein_coding | 978.9299283 | -0.345573315 | 0.000448742 | 0.004201078 |
| ENSMUST00000144064.4,Oasl2-007,nonsense_mediated_decay,ENSMUSG00000029561.17,protein_coding | 814.0031936 | 0.396609018 | 0.000450433 | 0.004215993 |

| | | | | |
|---|---|---|---|---|
| ENSMUST00000069772.15,Tmem143-001,protein_coding,ENSMUSG00000002781.19,protein_coding | 1384.391517 | 0.401607143 | 0.000450546 | 0.004216128 |
| ENSMUST00000014743.9,Csf1-001,protein_coding,ENSMUSG00000014599.10,protein_coding | 15403.29215 | -0.157555087 | 0.000451005 | 0.004219512 |
| ENSMUST00000111118.7,Tcp11l1-001,protein_coding,ENSMUSG00000027175.10,protein_coding | 1615.554096 | -0.218954509 | 0.000451643 | 0.004224557 |
| ENSMUST00000111670.8,Ctnnd1-201,protein_coding,ENSMUSG00000034101.14,protein_coding | 3161.352399 | 0.386076334 | 0.000452003 | 0.004227007 |
| ENSMUST00000121044.5,Btc-001,protein_coding,ENSMUSG00000082361.6,protein_coding | 2117.02528 | -0.32142234 | 0.000452467 | 0.00422951 |
| ENSMUST00000111473.7,Cdk2ap1-004,protein_coding,ENSMUSG00000029394.11,protein_coding | 207.4245933 | -0.529090323 | 0.000452401 | 0.00422951 |
| ENSMUST00000088552.6,Myl9-001,protein_coding,ENSMUSG00000067818.6,protein_coding | 172.7666908 | -0.429759915 | 0.000452894 | 0.004232579 |
| ENSMUST00000182884.7,Ank3-001,protein_coding,ENSMUSG00000069601.13,protein_coding | 1208.459856 | -0.28166259 | 0.000454041 | 0.004242384 |
| ENSMUST00000139299.7,Eif4a1-006,processed_transcript,ENSMUSG00000059796.16,protein_coding | 164.1322119 | 0.65451624 | 0.000454581 | 0.004246507 |
| ENSMUST00000124834.7,Crls1-002,nonsense_mediated_decay,ENSMUSG00000027357.16,protein_coding | 294.2020889 | 0.57970192 | 0.000455465 | 0.004252917 |
| ENSMUST00000108928.1,Gm14326-003,protein_coding,ENSMUSG00000078862.10,protein_coding | 177.7083841 | -0.580799155 | 0.000455403 | 0.004252917 |
| ENSMUST00000107251.8,Pi4kb-002,protein_coding,ENSMUSG00000038861.14,protein_coding | 1430.033638 | 0.339375952 | 0.000456104 | 0.004257964 |
| ENSMUST00000111711.7,Tfpi-004,protein_coding,ENSMUSG00000027082.15,protein_coding | 33.73523824 | -0.651557384 | 0.000456469 | 0.004260443 |
| ENSMUST00000050491.10,Glrx2-011,nonsense_mediated_decay,ENSMUSG00000018196.18,protein_coding | 25.04588876 | -0.653665675 | 0.000457202 | 0.004266363 |
| ENSMUST00000046502.6,Rad54l2-001,protein_coding,ENSMUSG00000040661.6,protein_coding | 10036.73545 | 0.121936775 | 0.000457859 | 0.004271569 |
| ENSMUST00000079703.9,Nsmce2-201,protein_coding,ENSMUSG00000059586.9,protein_coding | 4840.517424 | 0.125369507 | 0.000458937 | 0.004280698 |
| ENSMUST00000045521.7,Dtx4-201,protein_coding,ENSMUSG00000039982.7,protein_coding | 1505.889592 | 0.251326695 | 0.000459721 | 0.004287073 |
| ENSMUST00000045628.13,R3hdm4-201,protein_coding,ENSMUSG00000035781.13,protein_coding | 9686.248709 | 0.147840913 | 0.000460588 | 0.004294236 |
| ENSMUST00000141650.7,Slc43a3-005,protein_coding,ENSMUSG00000027074.14,protein_coding | 899.6163713 | 0.407193355 | 0.000462875 | 0.004314621 |
| ENSMUST00000113144.7,Gprasp1-001,protein_coding,ENSMUSG00000043384.15,protein_coding | 559.6392736 | -0.500095069 | 0.00046338 | 0.004318389 |
| ENSMUST00000022946.5,Rida-001,protein_coding,ENSMUSG00000022323.11,protein_coding | 701.649964 | -0.388631077 | 0.000463793 | 0.004321308 |
| ENSMUST00000034203.16,Cog4-001,protein_coding,ENSMUSG00000031753.16,protein_coding | 8337.619128 | -0.146594232 | 0.000464153 | 0.004323721 |
| ENSMUST00000032114.7,Mogs-001,protein_coding,ENSMUSG00000030036.8,protein_coding | 11466.86433 | 0.145433013 | 0.000464764 | 0.004328481 |
| ENSMUST00000032477.5,Necap1-001,protein_coding,ENSMUSG00000030327.8,protein_coding | 12468.89446 | 0.175805485 | 0.000464948 | 0.004328988 |
| ENSMUST00000113112.1,Bex3-002,protein_coding,ENSMUSG00000046432.12,protein_coding | 2946.89285 | 0.19863244 | 0.00046502 | 0.004328988 |
| ENSMUST00000184272.2,Gm9780-201,protein_coding,ENSMUSG00000094800.8,protein_coding | 82.32621554 | 0.54588791 | 0.000465493 | 0.004332455 |
| ENSMUST00000030169.14,Stoml2-001,protein_coding,ENSMUSG00000028455.15,protein_coding | 8989.700555 | 0.180872231 | 0.000466237 | 0.004338442 |
| ENSMUST00000109108.8,Wwox-004,protein_coding,ENSMUSG00000004637.15,protein_coding | 99.57303272 | 0.642388337 | 0.000468182 | 0.004355601 |
| ENSMUST00000210297.1,Emp3-004,protein_coding,ENSMUSG00000040212.12,protein_coding | 644.4031827 | 0.544254693 | 0.000468713 | 0.004359596 |
| ENSMUST00000125660.2,Guf1-007,retained_intron,ENSMUSG00000029208.16,protein_coding | 221.8896295 | -0.51644416 | 0.000469238 | 0.004363534 |
| ENSMUST00000006973.11,Kat2a-001,protein_coding,ENSMUSG00000020918.14,protein_coding | 14548.08902 | 0.152222077 | 0.000469366 | 0.004363784 |
| ENSMUST00000097526.2,Tor1aip2-007,protein_coding,ENSMUSG00000050565.16,protein_coding | 2121.770617 | -0.28016482 | 0.000469527 | 0.004364343 |
| ENSMUST00000021567.5,Pcnx-201,protein_coding,ENSMUSG00000021140.8,protein_coding | 6229.80244 | 0.17996222 | 0.000469779 | 0.004365741 |
| ENSMUST00000003318.11,Dvl3-001,protein_coding,ENSMUSG00000003233.11,protein_coding | 2961.0293 | 0.156343776 | 0.000470171 | 0.004368442 |
| ENSMUST00000059204.9,Fbxl7-201,protein_coding,ENSMUSG00000043556.9,protein_coding | 207.6080804 | -0.394767683 | 0.000470312 | 0.004368807 |
| ENSMUST00000196704.1,Tm4sf1-005,retained_intron,ENSMUSG00000027800.14,protein_coding | 110.2581746 | -0.610025345 | 0.000471245 | 0.004376528 |
| ENSMUST00000036111.9,Mrps35-001,protein_coding,ENSMUSG00000040112.9,protein_coding | 3852.143823 | 0.190085399 | 0.000471379 | 0.004376828 |
| ENSMUST00000038368.8,Id1-001,protein_coding,ENSMUSG00000042745.9,protein_coding | 15360.92571 | -0.262498786 | 0.00047223 | 0.004383785 |
| ENSMUST00000160027.7,Psma3-001,protein_coding,ENSMUSG00000060073.9,protein_coding | 924.8379446 | -0.353524085 | 0.000474031 | 0.004399552 |
| ENSMUST00000165489.7,Npepps-013,retained_intron,ENSMUSG00000001441.13,protein_coding | 261.0078736 | 0.568338467 | 0.000474602 | 0.004403908 |
| ENSMUST00000040523.8,Adamts2-001,protein_coding,ENSMUSG00000036545.8,protein_coding | 3767.66414 | 0.127173367 | 0.000474832 | 0.004405086 |
| ENSMUST00000008594.8,Nutf2-001,protein_coding,ENSMUSG00000008450.8,protein_coding | 22098.58814 | 0.126532479 | 0.000474993 | 0.004405634 |
| ENSMUST00000186993.1,Snhg17-009,lincRNA,ENSMUSG00000085385.7,processed_transcript | 946.5749275 | -0.240402538 | 0.000476921 | 0.004422564 |
| ENSMUST00000084241.11,Wasf2-001,protein_coding,ENSMUSG00000028868.13,protein_coding | 20975.73956 | 0.135425158 | 0.000477316 | 0.004425269 |
| ENSMUST00000094756.10,Rbm47-006,protein_coding,ENSMUSG00000070780.11,protein_coding | 62.75804792 | -0.609529844 | 0.000477573 | 0.004426701 |
| ENSMUST00000066991.5,Adm2-201,protein_coding,ENSMUSG00000054136.5,protein_coding | 933.5953063 | 0.25326536 | 0.000477708 | 0.004427002 |
| ENSMUST00000076228.2,Wbscr16-001,protein_coding,ENSMUSG00000061979.8,protein_coding | 4623.62132 | -0.135519822 | 0.000477986 | 0.004428625 |
| ENSMUST00000069808.12,Nrf1-202,protein_coding,ENSMUSG00000058440.14,protein_coding | 1311.127335 | 0.436675793 | 0.000478096 | 0.004428692 |
| ENSMUST00000183451.1,Nedd4-008,retained_intron,ENSMUSG00000032216.14,protein_coding | 67.51267174 | -0.537973278 | 0.000478402 | 0.004430354 |
| ENSMUST00000099384.2,Brd9-201,protein_coding,ENSMUSG00000057649.5,protein_coding | 3952.214833 | -0.195432366 | 0.000478482 | 0.004430354 |
| ENSMUST00000142942.7,Ube2d-ps-001,processed_transcript,ENSMUSG00000083844.8,transcribed_unprocessed_pseudogene | 434.9275911 | 0.406849737 | 0.000478878 | 0.004433074 |
| ENSMUST00000037333.16,Ccnd3-001,protein_coding,ENSMUSG00000034165.16,protein_coding | 73.64972007 | 0.618205687 | 0.000479819 | 0.004440829 |
| ENSMUST00000108407.9,Abr-201,protein_coding,ENSMUSG00000017631.18,protein_coding | 12155.3009 | -0.147401812 | 0.000480851 | 0.004449423 |
| ENSMUST00000157045.7,Jtb-002,retained_intron,ENSMUSG00000027937.15,protein_coding | 73.1523385 | -0.610061983 | 0.000481061 | 0.004450409 |
| ENSMUST00000044038.9,Kyat1-001,protein_coding,ENSMUSG00000039648.14,protein_coding | 609.1361206 | 0.463752108 | 0.000481576 | 0.004453933 |
| ENSMUST00000105851.8,Pithd1-001,protein_coding,ENSMUSG00000028669.15,protein_coding | 4688.458718 | 0.245055788 | 0.000481649 | 0.004453933 |
| ENSMUST00000172732.7,Rfc1-004,protein_coding,ENSMUSG00000029191.16,protein_coding | 4649.858952 | -0.231730291 | 0.000482651 | 0.004461282 |
| ENSMUST00000023854.9,Fhl1-001,protein_coding,ENSMUSG00000023092.16,protein_coding | 3627.634348 | -0.185226004 | 0.000482618 | 0.004461282 |
| ENSMUST00000128341.1,Myo9a-003,protein_coding,ENSMUSG00000039585.15,protein_coding | 991.1570703 | -0.648739411 | 0.000483472 | 0.004467914 |
| ENSMUST00000160722.7,B230118H07Rik-008,protein_coding,ENSMUSG00000027165.16,protein_coding | 513.5663932 | -0.487407562 | 0.000483889 | 0.004470803 |
| ENSMUST00000178079.7,Rpl31-003,protein_coding,ENSMUSG00000073702.11,protein_coding | 35513.3886 | 0.125638354 | 0.000484898 | 0.004479167 |
| ENSMUST00000066091.13,Smad1-001,protein_coding,ENSMUSG00000031681.16,protein_coding | 5861.031493 | 0.230342671 | 0.000485075 | 0.004479844 |
| ENSMUST00000161170.1,Zfp9-001,protein_coding,ENSMUSG00000072623.6,protein_coding | 3916.181549 | -0.226820479 | 0.0004859 | 0.0044865 |
| ENSMUST00000085671.9,Zkscan5-001,protein_coding,ENSMUSG00000055991.15,protein_coding | 1090.653319 | -0.24947292 | 0.000486323 | 0.004489438 |
| ENSMUST00000100015.5,Ola1-002,protein_coding,ENSMUSG00000027108.15,protein_coding | 2054.742937 | 0.239286727 | 0.000486528 | 0.004489534 |
| ENSMUST00000208172.1,Nup62-005,protein_coding,ENSMUSG00000109511.1,protein_coding | 233.4201677 | 0.604510279 | 0.000486646 | 0.004489534 |
| ENSMUST00000107159.2,Nars2-002,protein_coding,ENSMUSG00000018995.12,protein_coding | 370.735157 | 0.376940121 | 0.000486587 | 0.004489534 |
| ENSMUST00000117423.8,Kif2a-005,protein_coding,ENSMUSG00000021693.19,protein_coding | 3906.289664 | 0.169674165 | 0.000487252 | 0.004494162 |
| ENSMUST00000138603.8,Hap1-005,protein_coding,ENSMUSG00000006930.15,protein_coding | 433.6177246 | -0.313968756 | 0.000488358 | 0.004503398 |
| ENSMUST00000146892.1,Mvb12b-003,processed_transcript,ENSMUSG00000038740.9,protein_coding | 191.9420102 | 0.469764822 | 0.000490315 | 0.004520478 |
| ENSMUST00000106259.8,Gaa-001,protein_coding,ENSMUSG00000025579.14,protein_coding | 8866.825105 | 0.221340801 | 0.000491272 | 0.00452833 |
| ENSMUST00000020979.7,Bcap29-201,protein_coding,ENSMUSG00000020650.14,protein_coding | 196.7505555 | -0.560670571 | 0.000491478 | 0.004529257 |
| ENSMUST00000188859.6,Sf3b1-006,retained_intron,ENSMUSG00000025982.13,protein_coding | 43.43765219 | 0.647746985 | 0.000493642 | 0.004548227 |
| ENSMUST00000211765.1,Nucb1-001,protein_coding,ENSMUSG00000030824.17,protein_coding | 3816.347608 | 0.176185212 | 0.000493953 | 0.004550113 |
| ENSMUST00000073945.5,Vkorc1l1-003,protein_coding,ENSMUSG00000066735.8,protein_coding | 386.3207155 | 0.392167912 | 0.000494524 | 0.004554407 |
| ENSMUST00000127370.2,Zrsr2-005,protein_coding,ENSMUSG00000031370.12,protein_coding | 169.0508891 | -0.461272955 | 0.000495304 | 0.004560607 |
| ENSMUST00000114533.8,Slc4a1ap-201,protein_coding,ENSMUSG00000029141.16,protein_coding | 2530.235178 | 0.173651958 | 0.000497967 | 0.004584146 |
| ENSMUST00000070642.3,Cebpb-001,protein_coding,ENSMUSG00000056501.3,protein_coding | 24759.77514 | 0.192636106 | 0.000498404 | 0.004587191 |
| ENSMUST00000068156.6,Vps13a-201,protein_coding,ENSMUSG00000046230.9,protein_coding | 5952.468548 | -0.475334008 | 0.000498576 | 0.004587793 |
| ENSMUST00000028914.8,Polr3f-001,protein_coding,ENSMUSG00000027427.13,protein_coding | 4022.625426 | -0.227511256 | 0.000499024 | 0.004590932 |
| ENSMUST00000003669.7,Homer3-002,protein_coding,ENSMUSG00000003573.15,protein_coding | 1055.87136 | -0.646679045 | 0.000500996 | 0.004608093 |
| ENSMUST00000027286.6,Coa5-001,protein_coding,ENSMUSG00000026112.7,protein_coding | 13170.3711 | 0.163719501 | 0.000502312 | 0.004619212 |
| ENSMUST00000034808.11,Nnmt-001,protein_coding,ENSMUSG00000032271.13,protein_coding | 664.2876673 | 0.366220568 | 0.000506037 | 0.004652467 |
| ENSMUST00000000090.7,Cox5a-001,protein_coding,ENSMUSG00000000088.7,protein_coding | 11762.98309 | 0.173279831 | 0.000506583 | 0.004656497 |
| ENSMUST00000035155.7,Rab6b-001,protein_coding,ENSMUSG00000032549.7,protein_coding | 382.8448145 | -0.392023077 | 0.000506833 | 0.0046578 |
| ENSMUST00000115388.8,Adam22-202,protein_coding,ENSMUSG00000040537.17,protein_coding | 335.350862 | -0.528622616 | 0.000508142 | 0.004668836 |
| ENSMUST00000116380.8,Rpn2-001,protein_coding,ENSMUSG00000027642.15,protein_coding | 18621.98981 | 0.135410798 | 0.000508749 | 0.004672411 |
| ENSMUST00000125982.1,Serpinf1-005,protein_coding,ENSMUSG00000000753.15,protein_coding | 431.9682975 | 0.385776356 | 0.00050872 | 0.004672411 |
| ENSMUST00000077458.6,Stxbp1-003,protein_coding,ENSMUSG00000026797.16,protein_coding | 5083.333277 | 0.198783822 | 0.000510447 | 0.004686354 |
| ENSMUST00000025811.4,Yif1a-201,protein_coding,ENSMUSG00000024875.4,protein_coding | 5636.863993 | 0.167874416 | 0.000510593 | 0.004686354 |
| ENSMUST00000011896.6,Pgam1-201,protein_coding,ENSMUSG00000011752.6,protein_coding | 165137.8198 | 0.172034474 | 0.000510531 | 0.004686354 |
| ENSMUST00000159388.7,Vangl1-201,protein_coding,ENSMUSG00000027860.15,protein_coding | 2555.700705 | 0.143857999 | 0.000514209 | 0.004718532 |

| | | | | |
|---|---|---|---|---|
| ENSMUST00000148488.1,Phgdh-002,protein_coding,ENSMUSG00000053398.11,protein_coding | 625.1919067 | 0.626605226 | 0.000514913 | 0.004723985 |
| ENSMUST00000124246.3,Gt(ROSA)26Sor-002,antisense,ENSMUSG00000086429.9,antisense | 983.956617 | -0.347047226 | 0.000516585 | 0.004738232 |
| ENSMUST00000180252.1,Tmem151b-201,protein_coding,ENSMUSG00000096847.1,protein_coding | 41.26600806 | -0.644801237 | 0.000516686 | 0.004738232 |
| ENSMUST00000107800.2,Trmt10b-002,protein_coding,ENSMUSG00000035601.9,protein_coding | 323.8197833 | 0.3373938 | 0.000517688 | 0.004746413 |
| ENSMUST00000162505.7,Tbp-001,protein_coding,ENSMUSG00000014767.16,protein_coding | 311.4434668 | -0.429247446 | 0.000518947 | 0.004756944 |
| ENSMUST00000085427.5,Zfp865-002,protein_coding,ENSMUSG00000074405.7,protein_coding | 1493.025331 | 0.189635891 | 0.000520142 | 0.004766879 |
| ENSMUST00000143813.1,0610009L18Rik-001,protein_coding,ENSMUSG00000043644.4,protein_coding | 77.21399728 | 0.519794068 | 0.000520513 | 0.004769261 |
| ENSMUST00000138800.7,Ppdpf-005,retained_intron,ENSMUSG00000016344.14,protein_coding | 388.6387514 | 0.375566695 | 0.00052112 | 0.004773809 |
| ENSMUST00000114993.8,Cnot4-005,protein_coding,ENSMUSG00000038784.13,protein_coding | 3757.979836 | 0.170295582 | 0.000521599 | 0.004777178 |
| ENSMUST00000099498.8,Ccdc80-001,protein_coding,ENSMUSG00000022665.14,protein_coding | 1138.857633 | -0.277842623 | 0.000524661 | 0.004804204 |
| ENSMUST00000050201.10,Mapk3-003,protein_coding,ENSMUSG00000063065.13,protein_coding | 69.51020701 | -0.610974963 | 0.000525794 | 0.004813554 |
| ENSMUST00000040154.8,Cox6a1-001,protein_coding,ENSMUSG00000041697.8,protein_coding | 21995.43663 | 0.141801506 | 0.000527087 | 0.004824057 |
| ENSMUST00000211882.1,Stox2-010,protein_coding,ENSMUSG00000038143.16,protein_coding | 442.5261389 | -0.387981823 | 0.000527166 | 0.004824057 |
| ENSMUST00000185482.6,Atg4b-010,nonsense_mediated_decay,ENSMUSG00000026280.17,protein_coding | 1097.653104 | 0.251035341 | 0.000527894 | 0.004829701 |
| ENSMUST00000091420.5,Skida1-001,protein_coding,ENSMUSG00000054074.9,protein_coding | 143.3364888 | -0.558173759 | 0.000528346 | 0.004832803 |
| ENSMUST00000064941.6,Nr1d1-001,protein_coding,ENSMUSG00000020889.11,protein_coding | 2487.456494 | 0.176062351 | 0.000528895 | 0.004836797 |
| ENSMUST00000149366.1,Ndc1-002,nonsense_mediated_decay,ENSMUSG00000028614.14,protein_coding | 417.4653013 | 0.459000418 | 0.000529149 | 0.004838094 |
| ENSMUST00000126303.7,Asns-003,protein_coding,ENSMUSG00000029752.12,protein_coding | 105.4249468 | 0.630366485 | 0.00053161 | 0.00485853 |
| ENSMUST00000181162.2,Gm7389-001,processed_pseudogene,ENSMUSG00000097657.2,processed_pseudogene | 1142.569401 | 0.395601029 | 0.000531569 | 0.00485853 |
| ENSMUST00000117148.1,Gm7488-001,processed_pseudogene,ENSMUSG00000039617.8,processed_pseudogene | 1940.652516 | 0.238133442 | 0.000532553 | 0.004865087 |
| ENSMUST00000114387.7,Naa10-004,protein_coding,ENSMUSG00000031388.14,protein_coding | 1388.553029 | 0.271177402 | 0.00053245 | 0.004865087 |
| ENSMUST00000026990.5,Thoc3-001,protein_coding,ENSMUSG00000025872.5,protein_coding | 8406.699417 | -0.162937119 | 0.00053349 | 0.004872612 |
| ENSMUST00000062092.5,Cdca4-201,protein_coding,ENSMUSG00000047832.8,protein_coding | 9731.331538 | 0.127746477 | 0.000533959 | 0.004875857 |
| ENSMUST00000108985.8,Gm14306-004,protein_coding,ENSMUSG00000078879.10,protein_coding | 252.2508254 | -0.488602858 | 0.000535044 | 0.004884731 |
| ENSMUST00000115850.1,Cdh2-003,protein_coding,ENSMUSG00000024304.14,protein_coding | 13324.70623 | -0.115751845 | 0.000535673 | 0.004889436 |
| ENSMUST00000124909.1,Trrap-009,retained_intron,ENSMUSG00000045482.16,protein_coding | 174.0877597 | 0.514889816 | 0.000536102 | 0.004892317 |
| ENSMUST00000057725.9,Samhd1-001,protein_coding,ENSMUSG00000027639.16,protein_coding | 10906.19964 | -0.172558727 | 0.000537297 | 0.004902183 |
| ENSMUST00000008999.11,Hdac5-002,protein_coding,ENSMUSG00000008855.17,protein_coding | 107.7912631 | -0.632492083 | 0.000539262 | 0.004918817 |
| ENSMUST00000079121.3,Mrpl18-001,protein_coding,ENSMUSG00000057388.9,protein_coding | 7615.923668 | -0.246334228 | 0.000539349 | 0.004918817 |
| ENSMUST00000105992.8,Pum1-202,protein_coding,ENSMUSG00000028580.15,protein_coding | 1572.75472 | -0.501736042 | 0.000539665 | 0.00492053 |
| ENSMUST00000164666.5,Mcc-002,protein_coding,ENSMUSG00000071856.10,protein_coding | 2097.164579 | -0.210057439 | 0.000539765 | 0.00492053 |
| ENSMUST00000072607.8,Ints1-004,protein_coding,ENSMUSG00000029547.10,protein_coding | 19876.01329 | -0.161928595 | 0.000540718 | 0.004928169 |
| ENSMUST00000131688.7,Mboat7-003,retained_intron,ENSMUSG00000035596.14,protein_coding | 702.9386687 | -0.307746245 | 0.000540981 | 0.004929521 |
| ENSMUST00000087622.5,Plekha5-001,protein_coding,ENSMUSG00000030231.11,protein_coding | 543.9089296 | 0.36918051 | 0.000541682 | 0.004934862 |
| ENSMUST00000122385.2,Gm15975-001,processed_pseudogene,ENSMUSG00000083902.2,processed_pseudogene | 67.68623106 | -0.568529829 | 0.000543197 | 0.004947617 |
| ENSMUST00000211593.1,Dock1-002,protein_coding,ENSMUSG00000058325.6,protein_coding | 198.8501174 | -0.436204187 | 0.000543426 | 0.004948658 |
| ENSMUST00000130521.1,Pcsk4-006,retained_intron,ENSMUSG00000020131.14,protein_coding | 154.6284292 | 0.568073479 | 0.000544375 | 0.004956251 |
| ENSMUST00000074541.5,Jazf1-001,protein_coding,ENSMUSG00000063568.11,protein_coding | 185.8045925 | 0.396474252 | 0.00054465 | 0.004957706 |
| ENSMUST00000181711.1,Gm19426-001,protein_coding,ENSMUSG00000097187.1,protein_coding | 289.6431207 | -0.341930325 | 0.000545745 | 0.004966626 |
| ENSMUST00000160359.1,Plod2-002,protein_coding,ENSMUSG00000032374.14,protein_coding | 4533.889621 | -0.180167833 | 0.000545969 | 0.004967616 |
| ENSMUST00000166021.7,Gpatch4-201,protein_coding,ENSMUSG00000028069.16,protein_coding | 234.5761995 | -0.51399008 | 0.000546633 | 0.004972601 |
| ENSMUST00000076991.6,4930453N24Rik-001,protein_coding,ENSMUSG00000059920.9,protein_coding | 3179.876231 | -0.168443482 | 0.000546938 | 0.004974331 |
| ENSMUST00000023501.14,Maats1-002,protein_coding,ENSMUSG00000022805.16,protein_coding | 98.61709432 | 0.514074237 | 0.000547543 | 0.004978777 |
| ENSMUST00000212338.1,Csnk2a2-007,retained_intron,ENSMUSG00000046707.9,protein_coding | 157.004253 | -0.521892734 | 0.000548474 | 0.004986187 |
| ENSMUST00000032359.14,Aebp2-001,protein_coding,ENSMUSG00000030232.17,protein_coding | 3343.899365 | -0.328837625 | 0.000548851 | 0.004988565 |
| ENSMUST00000023806.12,Soat2-001,protein_coding,ENSMUSG00000023045.12,protein_coding | 312.4623763 | 0.479846848 | 0.000550651 | 0.005003865 |
| ENSMUST00000148969.7,Nup188-007,nonsense_mediated_decay,ENSMUSG00000052533.14,protein_coding | 61.95575646 | 0.630590686 | 0.000551478 | 0.005010322 |
| ENSMUST00000176600.7,Pan3-002,protein_coding,ENSMUSG00000029647.15,protein_coding | 1165.933185 | -0.326757465 | 0.000552491 | 0.005018468 |
| ENSMUST00000049031.5,Fam63b-001,protein_coding,ENSMUSG00000042444.10,protein_coding | 2026.680325 | -0.432133706 | 0.000553488 | 0.005026463 |
| ENSMUST00000141755.7,Mettl16-001,protein_coding,ENSMUSG00000010554.14,protein_coding | 6149.430859 | -0.115020169 | 0.000553932 | 0.00502944 |
| ENSMUST00000156110.7,Gak-005,protein_coding,ENSMUSG00000062234.14,protein_coding | 950.3442576 | 0.436993373 | 0.000554383 | 0.00503247 |
| ENSMUST00000033321.10,Cask-002,protein_coding,ENSMUSG00000031012.17,protein_coding | 6000.317401 | 0.21774233 | 0.000555192 | 0.005038748 |
| ENSMUST00000077755.10,Arpc5-001,protein_coding,ENSMUSG00000008475.13,protein_coding | 9632.274 | -0.158247913 | 0.000555868 | 0.005043818 |
| ENSMUST00000081980.6,Med9-001,protein_coding,ENSMUSG00000061650.6,protein_coding | 4398.893663 | 0.162934628 | 0.000556056 | 0.005044461 |
| ENSMUST00000047815.12,Slc9a8-001,protein_coding,ENSMUSG00000039463.14,protein_coding | 2723.987472 | -0.180411987 | 0.000556209 | 0.005044791 |
| ENSMUST00000099678.4,Fjx1-001,protein_coding,ENSMUSG00000075012.4,protein_coding | 147.001951 | -0.45887963 | 0.000559112 | 0.005069602 |
| ENSMUST00000036719.11,Prex1-001,protein_coding,ENSMUSG00000039621.13,protein_coding | 534.080135 | -0.236176801 | 0.00055918 | 0.005069602 |
| ENSMUST00000213012.1,RP23-114B10.6-001,processed_transcript,ENSMUSG00000110444.1,transcribed_unprocessed_pseudogene | 414.5251497 | -0.582593092 | 0.000560414 | 0.005078952 |
| ENSMUST00000123791.7,Sbf1-001,protein_coding,ENSMUSG00000036529.17,protein_coding | 8861.326528 | 0.123053516 | 0.000560447 | 0.005078952 |
| ENSMUST00000198126.1,Adh5-007,nonsense_mediated_decay,ENSMUSG00000028138.8,protein_coding | 1879.7998 | 0.335361378 | 0.000562181 | 0.005092981 |
| ENSMUST00000177652.1,Ap3m1-ps-201,pseudogene,ENSMUSG00000094683.1,pseudogene | 29.02915921 | -0.640134879 | 0.000562232 | 0.005092981 |
| ENSMUST00000100812.3,Git1-004,protein_coding,ENSMUSG00000011877.13,protein_coding | 10338.81834 | 0.119913147 | 0.000562714 | 0.005096275 |
| ENSMUST00000112289.8,Zrsr2-001,protein_coding,ENSMUSG00000031370.12,protein_coding | 1870.39743 | 0.241884195 | 0.000565074 | 0.005116566 |
| ENSMUST00000015920.11,Med22-002,protein_coding,ENSMUSG00000015776.12,protein_coding | 435.9993937 | 0.493666231 | 0.000565625 | 0.005120484 |
| ENSMUST00000048653.9,Cpne2-001,protein_coding,ENSMUSG00000034361.10,protein_coding | 6210.113436 | 0.102168438 | 0.000567478 | 0.005136177 |
| ENSMUST00000091186.6,Intu-001,protein_coding,ENSMUSG00000060798.7,protein_coding | 1088.002224 | -0.218270367 | 0.000569103 | 0.005149805 |
| ENSMUST00000149210.1,Zfp106-004,retained_intron,ENSMUSG00000027288.16,protein_coding | 570.2202731 | 0.30328196 | 0.000570941 | 0.005165345 |
| ENSMUST00000152724.1,Usp49-002,protein_coding,ENSMUSG00000090115.7,protein_coding | 276.040907 | 0.372070126 | 0.000571438 | 0.005168762 |
| ENSMUST00000094835.8,1110051M20Rik-003,protein_coding,ENSMUSG00000040591.18,protein_coding | 1685.091787 | -0.233160527 | 0.000571861 | 0.005171501 |
| ENSMUST00000057627.15,Spata2-001,protein_coding,ENSMUSG00000047030.15,protein_coding | 4498.371521 | 0.239773033 | 0.000573132 | 0.005181902 |
| ENSMUST00000039818.9,Aldh4a1-001,protein_coding,ENSMUSG00000028737.15,protein_coding | 1739.156623 | -0.20974658 | 0.000573517 | 0.005184294 |
| ENSMUST00000204991.2,Firre-007,processed_transcript,ENSMUSG00000085396.7,processed_transcript | 109.9266804 | 0.583652023 | 0.000573856 | 0.005186274 |
| ENSMUST00000043333.8,Tada3-005,protein_coding,ENSMUSG00000048930.12,protein_coding | 4009.796281 | 0.203271017 | 0.000574027 | 0.00518673 |
| ENSMUST00000113338.8,Ciz1-201,protein_coding,ENSMUSG00000039205.16,protein_coding | 3280.774168 | 0.337377961 | 0.00057425 | 0.005187648 |
| ENSMUST00000079125.6,Scand1-201,protein_coding,ENSMUSG00000046229.9,protein_coding | 5678.229894 | 0.154263481 | 0.000574772 | 0.005191278 |
| ENSMUST00000066379.10,Chchd3-001,protein_coding,ENSMUSG00000053768.13,protein_coding | 14928.75648 | 0.163445751 | 0.000577032 | 0.005209507 |
| ENSMUST00000171999.7,Gcat-203,protein_coding,ENSMUSG00000006378.13,protein_coding | 2479.401532 | 0.241621674 | 0.000576933 | 0.005209507 |
| ENSMUST00000198535.1,Gm43437-001,TEC,ENSMUSG00000104965.1,TEC | 42.89790602 | -0.627784177 | 0.000577558 | 0.005213157 |
| ENSMUST00000043338.9,Sft2d2-001,protein_coding,ENSMUSG00000040848.11,protein_coding | 7408.517108 | 0.231020025 | 0.0005792 | 0.005226882 |
| ENSMUST00000138696.7,Atpaf2-003,processed_transcript,ENSMUSG00000042709.14,protein_coding | 442.9656313 | 0.292092002 | 0.000579324 | 0.005226908 |
| ENSMUST00000025430.10,Setbp1-001,protein_coding,ENSMUSG00000024548.11,protein_coding | 1489.385075 | -0.300351445 | 0.000580783 | 0.005238974 |
| ENSMUST00000196109.4,Srrt-020,protein_coding,ENSMUSG00000037364.11,protein_coding | 57.02851382 | -0.639695923 | 0.000581799 | 0.005247044 |
| ENSMUST00000213957.1,Spg21-002,protein_coding,ENSMUSG00000032388.7,protein_coding | 2293.559225 | 0.200415042 | 0.000582311 | 0.00525056 |
| ENSMUST00000017859.9,Pgs1-004,nonsense_mediated_decay,ENSMUSG00000017715.11,protein_coding | 4640.609946 | 0.125828607 | 0.000582842 | 0.005254247 |
| ENSMUST00000159986.7,Wtap-004,protein_coding,ENSMUSG00000060475.12,protein_coding | 261.7440005 | -0.415006145 | 0.00058335 | 0.00525772 |
| ENSMUST00000192762.5,Rfwd2-009,protein_coding,ENSMUSG00000040782.12,protein_coding | 769.9327912 | 0.418606156 | 0.000583517 | 0.005258129 |
| ENSMUST00000165318.3,Rbak-001,protein_coding,ENSMUSG00000061898.10,protein_coding | 267.5567557 | 0.462100024 | 0.000584927 | 0.005269732 |
| ENSMUST00000009120.7,Gabpa-001,protein_coding,ENSMUSG00000008976.16,protein_coding | 3930.092393 | -0.35622734 | 0.000585331 | 0.005272269 |
| ENSMUST00000216284.1,Cep164-005,protein_coding,ENSMUSG00000043987.18,protein_coding | 91.59607064 | 0.636851695 | 0.000587032 | 0.00528648 |
| ENSMUST00000011315.9,Vipr2-001,protein_coding,ENSMUSG00000011171.11,protein_coding | 33.13764815 | 0.636255661 | 0.00058791 | 0.00529328 |
| ENSMUST00000089162.4,Edem1-001,protein_coding,ENSMUSG00000030104.9,protein_coding | 13406.34374 | 0.164369298 | 0.000589668 | 0.005307995 |
| ENSMUST00000029140.11,Procr-001,protein_coding,ENSMUSG00000027611.12,protein_coding | 2506.027859 | 0.196508008 | 0.00059158 | 0.005324098 |

| | | | | |
|---|---|---|---|---|
| ENSMUST00000112889.7,Tmem164-003,protein_coding,ENSMUSG00000047045.17,protein_coding | 211.326162 | 0.390016069 | 0.000591807 | 0.005325027 |
| ENSMUST00000215875.1,Gnb5-002,protein_coding,ENSMUSG00000032192.9,protein_coding | 702.0274979 | -0.334535871 | 0.000592336 | 0.005328669 |
| ENSMUST00000187951.6,Edem3-002,protein_coding,ENSMUSG00000043019.12,protein_coding | 113.9494873 | -0.62505613 | 0.000593951 | 0.005338735 |
| ENSMUST00000030623.7,Sfpq-001,protein_coding,ENSMUSG00000028820.13,protein_coding | 45938.15943 | 0.179832833 | 0.000593943 | 0.005338735 |
| ENSMUST00000152196.1,Asxl2-007,processed_transcript,ENSMUSG00000037486.17,protein_coding | 653.5470151 | -0.448074312 | 0.000593858 | 0.005338735 |
| ENSMUST00000164298.7,Rcbtb2-011,nonsense_mediated_decay,ENSMUSG00000022106.14,protein_coding | 219.2937047 | -0.639946668 | 0.000593821 | 0.005338735 |
| ENSMUST00000071329.7,Bckdha-001,protein_coding,ENSMUSG00000060376.7,protein_coding | 2767.773739 | 0.221841144 | 0.000595557 | 0.005352053 |
| ENSMUST00000070552.13,Osbp2-001,protein_coding,ENSMUSG00000020435.17,protein_coding | 294.9284492 | 0.318228976 | 0.000596207 | 0.005356782 |
| ENSMUST00000035157.9,Srprb-001,protein_coding,ENSMUSG00000032553.14,protein_coding | 8073.431348 | 0.15664239 | 0.000596807 | 0.005361051 |
| ENSMUST00000105813.7,Mul1-002,protein_coding,ENSMUSG00000041241.12,protein_coding | 355.0089877 | -0.34176623 | 0.000596936 | 0.00536109 |
| ENSMUST00000144900.1,Tfe3-013,protein_coding,ENSMUSG00000000134.17,protein_coding | 216.1295245 | 0.444818099 | 0.000597875 | 0.005368402 |
| ENSMUST00000106998.7,Clpb-003,protein_coding,ENSMUSG00000001829.17,protein_coding | 5957.122077 | 0.217533015 | 0.000598922 | 0.005376679 |
| ENSMUST00000042944.7,Arap3-201,protein_coding,ENSMUSG00000024451.8,protein_coding | 671.211063 | -0.218568924 | 0.000599621 | 0.00538184 |
| ENSMUST00000212964.1,Zfp821-001,protein_coding,ENSMUSG00000031728.10,protein_coding | 725.8740272 | 0.295301826 | 0.000600455 | 0.005388076 |
| ENSMUST00000182813.1,Pdlim1-004,retained_intron,ENSMUSG00000055044.12,protein_coding | 1879.531862 | 0.292207576 | 0.000600567 | 0.005388076 |
| ENSMUST00000111315.7,Adamts4-001,protein_coding,ENSMUSG00000006403.12,protein_coding | 2615.706356 | -0.153780344 | 0.000603178 | 0.005410378 |
| ENSMUST00000132618.1,Eif1-005,processed_transcript,ENSMUSG00000035530.13,protein_coding | 2716.9598 | 0.269903174 | 0.000603633 | 0.005413327 |
| ENSMUST00000069511.6,Abcd2-201,protein_coding,ENSMUSG00000055782.8,protein_coding | 130.8623927 | -0.473519193 | 0.000603808 | 0.005413775 |
| ENSMUST00000047186.9,Wipi1-002,protein_coding,ENSMUSG00000041895.15,protein_coding | 1374.557178 | -0.210808022 | 0.000604445 | 0.005418359 |
| ENSMUST00000035371.8,Sv2a-001,protein_coding,ENSMUSG00000038486.9,protein_coding | 1079.87547 | -0.192175528 | 0.000605118 | 0.005423099 |
| ENSMUST00000022595.7,Rgcc-001,protein_coding,ENSMUSG00000022018.7,protein_coding | 56.29006015 | 0.574521283 | 0.000605226 | 0.005423099 |
| ENSMUST00000047734.14,Zfp281-001,protein_coding,ENSMUSG00000041483.14,protein_coding | 2120.406266 | -0.35546332 | 0.000606408 | 0.005432554 |
| ENSMUST00000171606.7,Agtpbp1-003,protein_coding,ENSMUSG00000021557.14,protein_coding | 137.9962119 | -0.551933909 | 0.000607521 | 0.0054414 |
| ENSMUST00000072948.10,Acox1-002,protein_coding,ENSMUSG00000020777.16,protein_coding | 1304.38523 | -0.306986246 | 0.000607911 | 0.005443752 |
| ENSMUST00000134142.7,Tia1-008,retained_intron,ENSMUSG00000071337.11,protein_coding | 581.6376549 | -0.413252381 | 0.000610251 | 0.005463576 |
| ENSMUST00000027409.9,Des-001,protein_coding,ENSMUSG00000026208.9,protein_coding | 434.4840535 | 0.293134778 | 0.000610514 | 0.00546479 |
| ENSMUST00000028475.8,Clp1-001,protein_coding,ENSMUSG00000027079.15,protein_coding | 1759.506637 | -0.191389292 | 0.000610833 | 0.005466513 |
| ENSMUST00000114475.8,Brd4-202,protein_coding,ENSMUSG00000024002.16,protein_coding | 686.9180115 | 0.361910392 | 0.000613078 | 0.005485465 |
| ENSMUST00000042732.5,Fbxo45-001,protein_coding,ENSMUSG00000035764.5,protein_coding | 4086.554196 | -0.168554489 | 0.000613236 | 0.005485739 |
| ENSMUST00000192506.1,Gapdh-009,retained_intron,ENSMUSG00000057666.18,protein_coding | 4239.490852 | 0.152145717 | 0.000613692 | 0.005488678 |
| ENSMUST00000103198.10,Nop56-001,protein_coding,ENSMUSG00000027405.16,protein_coding | 53378.36921 | 0.152757035 | 0.000613932 | 0.005489679 |
| ENSMUST00000120299.3,Syngr4-003,protein_coding,ENSMUSG00000040231.16,protein_coding | 64.72794755 | 0.584780865 | 0.000615313 | 0.005498976 |
| ENSMUST00000036554.12,Abcc4-201,protein_coding,ENSMUSG00000032849.13,protein_coding | 10038.5227 | 0.15880293 | 0.000615355 | 0.005498976 |
| ENSMUST00000025356.2,Mal2-201,protein_coding,ENSMUSG00000024479.2,protein_coding | 2678.567994 | -0.233327977 | 0.000615149 | 0.005498976 |
| ENSMUST00000038701.7,Zfp324-001,protein_coding,ENSMUSG00000004500.9,protein_coding | 608.6023176 | -0.326011064 | 0.000616914 | 0.005511765 |
| ENSMUST00000199377.1,Eif3b-003,protein_coding,ENSMUSG00000056076.13,protein_coding | 972.4398662 | 0.49886745 | 0.000619858 | 0.005536573 |
| ENSMUST00000044604.14,Crybg3-001,protein_coding,ENSMUSG00000022723.15,protein_coding | 1826.193739 | -0.286591831 | 0.000619948 | 0.005536573 |
| ENSMUST00000017692.14,Suz12-001,protein_coding,ENSMUSG00000017548.15,protein_coding | 10154.88296 | -0.431972605 | 0.000621282 | 0.005547338 |
| ENSMUST00000178636.1,Larp1-201,protein_coding,ENSMUSG00000037331.15,protein_coding | 63205.55737 | 0.08600827 | 0.000621667 | 0.005549624 |
| ENSMUST00000166407.7,Ubxn1-202,protein_coding,ENSMUSG00000071655.11,protein_coding | 13355.1501 | 0.158358535 | 0.000621815 | 0.005549788 |
| ENSMUST00000111155.1,Hspb1-002,protein_coding,ENSMUSG00000004951.10,protein_coding | 79.02824539 | -0.518386638 | 0.00062246 | 0.005554388 |
| ENSMUST00000033915.8,Gpm6a-001,protein_coding,ENSMUSG00000031517.8,protein_coding | 149.4880888 | -0.482335795 | 0.000622588 | 0.005554388 |
| ENSMUST00000114049.1,Tmsb10-002,protein_coding,ENSMUSG00000079523.8,protein_coding | 3227.819887 | 0.243435321 | 0.00062323 | 0.005558965 |
| ENSMUST00000117786.7,Csnk1e-001,protein_coding,ENSMUSG00000022433.18,protein_coding | 14431.65223 | 0.154386585 | 0.000623411 | 0.00555943 |
| ENSMUST00000006625.7,Rbm14-001,protein_coding,ENSMUSG00000006456.10,protein_coding | 12026.20163 | 0.126579082 | 0.000624323 | 0.005566404 |
| ENSMUST00000151280.7,Hacd4-001,protein_coding,ENSMUSG00000028497.12,protein_coding | 427.287865 | -0.439974787 | 0.00062582 | 0.005578602 |
| ENSMUST00000132193.7,Alkbh6-002,retained_intron,ENSMUSG00000042831.13,protein_coding | 433.5466345 | 0.318195368 | 0.0006271 | 0.005588852 |
| ENSMUST00000033673.6,Nono-001,protein_coding,ENSMUSG00000031311.17,protein_coding | 63892.83263 | 0.166030325 | 0.000627706 | 0.005593097 |
| ENSMUST00000145514.7,Gls-005,retained_intron,ENSMUSG00000026103.14,protein_coding | 292.3313475 | -0.573844385 | 0.00062784 | 0.00559313 |
| ENSMUST00000132813.7,Mrpl27-002,processed_transcript,ENSMUSG00000024414.13,protein_coding | 67.38593578 | 0.624367331 | 0.000628781 | 0.005600355 |
| ENSMUST00000135795.7,Tctn3-002,protein_coding,ENSMUSG00000025008.15,protein_coding | 69.04799487 | 0.637811673 | 0.000629085 | 0.005601902 |
| ENSMUST00000088102.11,Zfx-001,protein_coding,ENSMUSG00000079509.10,protein_coding | 1426.987344 | -0.597615205 | 0.00062957 | 0.005605063 |
| ENSMUST00000176751.1,Gm20683-001,nonsense_mediated_decay,ENSMUSG00000093593.1,protein_coding | 175.1620905 | -0.482339607 | 0.00063009 | 0.005608533 |
| ENSMUST00000177779.7,Psap-203,protein_coding,ENSMUSG00000004207.14,protein_coding | 293.5430564 | 0.490279694 | 0.000630848 | 0.005614119 |
| ENSMUST00000047963.6,Rabepk-007,nonsense_mediated_decay,ENSMUSG00000070953.13,protein_coding | 495.8538472 | 0.4119309 | 0.000631239 | 0.005616436 |
| ENSMUST00000141118.8,Alas1-001,protein_coding,ENSMUSG00000032786.15,protein_coding | 549.2743949 | 0.382259666 | 0.000632366 | 0.0056253 |
| ENSMUST00000136532.1,Cggbp1-002,retained_intron,ENSMUSG00000054604.7,protein_coding | 1493.807378 | 0.240940488 | 0.000635292 | 0.005650164 |
| ENSMUST00000111231.9,B230118H07Rik-001,protein_coding,ENSMUSG00000027165.16,protein_coding | 81.53141297 | -0.63176756 | 0.00063553 | 0.005651115 |
| ENSMUST00000074734.12,Vill-002,protein_coding,ENSMUSG00000038775.14,protein_coding | 367.6992356 | 0.571696073 | 0.000635916 | 0.005653377 |
| ENSMUST00000199754.4,Gatad2b-004,protein_coding,ENSMUSG00000042390.9,protein_coding | 13768.47286 | 0.136666945 | 0.000638789 | 0.005677747 |
| ENSMUST00000161568.7,Tmem167-001,protein_coding,ENSMUSG00000012422.14,protein_coding | 4005.874006 | -0.362097706 | 0.000639321 | 0.005681303 |
| ENSMUST00000017783.12,Rab11fip4-001,protein_coding,ENSMUSG00000017639.13,protein_coding | 167.7001871 | -0.424525246 | 0.000639593 | 0.005682539 |
| ENSMUST00000126034.3,Slc9b2-003,retained_intron,ENSMUSG00000037994.14,protein_coding | 205.7912845 | 0.35725461 | 0.000639994 | 0.005684934 |
| ENSMUST00000193210.5,Cox5b-001,protein_coding,ENSMUSG00000061518.10,protein_coding | 13090.80673 | 0.129096227 | 0.000641387 | 0.00569496 |
| ENSMUST00000098651.5,Pias1-001,protein_coding,ENSMUSG00000032405.10,protein_coding | 2973.232932 | -0.163598242 | 0.000641266 | 0.00569496 |
| ENSMUST00000002805.13,Naa20-002,protein_coding,ENSMUSG00000002728.14,protein_coding | 407.6366025 | -0.411589007 | 0.000641671 | 0.005696306 |
| ENSMUST00000121465.2,Fuca2-001,protein_coding,ENSMUSG00000019810.15,protein_coding | 221.9986838 | -0.400275917 | 0.000642136 | 0.005699255 |
| ENSMUST00000125830.1,Pygo2-003,processed_transcript,ENSMUSG00000047824.13,protein_coding | 166.9453206 | 0.511234717 | 0.000644094 | 0.005715458 |
| ENSMUST00000113565.2,Obsl1-010,protein_coding,ENSMUSG00000026211.17,protein_coding | 2775.593379 | -0.198501294 | 0.000645026 | 0.005721366 |
| ENSMUST00000155341.1,AU040320-010,processed_transcript,ENSMUSG00000028830.14,protein_coding | 62.18586105 | 0.61110681 | 0.000644999 | 0.005721366 |
| ENSMUST00000001836.9,Pwp1-201,protein_coding,ENSMUSG00000001785.12,protein_coding | 11826.01478 | 0.154917748 | 0.000645245 | 0.005722131 |
| ENSMUST00000036225.14,Mpp6-003,protein_coding,ENSMUSG00000038388.14,protein_coding | 577.5917911 | 0.570637538 | 0.000646997 | 0.005736487 |
| ENSMUST00000213417.1,Clk3-009,retained_intron,ENSMUSG00000032316.8,protein_coding | 125.9047499 | 0.514336898 | 0.000647282 | 0.005736651 |
| ENSMUST00000154232.8,Brd2-003,nonsense_mediated_decay,ENSMUSG00000024335.19,protein_coding | 8567.00145 | 0.164125839 | 0.000647266 | 0.005736651 |
| ENSMUST00000091831.12,Smad2-003,protein_coding,ENSMUSG00000024563.15,protein_coding | 539.796023 | 0.43996954 | 0.000649138 | 0.005751915 |
| ENSMUST00000046073.15,Kdm2b-001,protein_coding,ENSMUSG00000029475.17,protein_coding | 552.9046726 | 0.486318914 | 0.000649356 | 0.00575222 |
| ENSMUST00000098350.9,Scap-001,protein_coding,ENSMUSG00000032485.14,protein_coding | 12209.42327 | 0.132766308 | 0.00064944 | 0.00575222 |
| ENSMUST00000049527.6,Prkar1a-001,protein_coding,ENSMUSG00000020612.16,protein_coding | 52373.53226 | 0.145339867 | 0.000650737 | 0.005762527 |
| ENSMUST00000164404.7,Vars2-004,nonsense_mediated_decay,ENSMUSG00000038838.15,protein_coding | 115.752365 | 0.635309055 | 0.00065093 | 0.005763049 |
| ENSMUST00000132828.1,Napa-004,retained_intron,ENSMUSG00000006024.13,protein_coding | 83.59377376 | 0.631330297 | 0.000651511 | 0.005767001 |
| ENSMUST00000039173.12,2310035C23Rik-002,protein_coding,ENSMUSG00000026319.13,protein_coding | 759.9630629 | -0.336444663 | 0.000652744 | 0.005775993 |
| ENSMUST00000202361.3,Hsph1-001,protein_coding,ENSMUSG00000029657.15,protein_coding | 11821.89506 | -0.329234509 | 0.000652795 | 0.005775993 |
| ENSMUST00000146409.7,Slc26a2-001,protein_coding,ENSMUSG00000034320.14,protein_coding | 1394.617049 | -0.294862123 | 0.000652947 | 0.005776152 |
| ENSMUST00000166627.7,Zfp414-002,protein_coding,ENSMUSG00000073423.9,protein_coding | 462.8938309 | 0.416955482 | 0.000653511 | 0.005779953 |
| ENSMUST00000090952.10,Fmnl2-001,protein_coding,ENSMUSG00000036053.16,protein_coding | 4118.20423 | 0.18406723 | 0.000654454 | 0.005787104 |
| ENSMUST00000143411.1,Dctn6-003,protein_coding,ENSMUSG00000031516.11,protein_coding | 72.12551872 | 0.542285797 | 0.000654621 | 0.005787392 |
| ENSMUST00000119417.1,Gm5844-001,processed_pseudogene,ENSMUSG00000082896.1,processed_pseudogene | 135.9253462 | -0.54009672 | 0.000655198 | 0.005790119 |
| ENSMUST00000138127.7,Zfp318-002,protein_coding,ENSMUSG00000015597.16,protein_coding | 35.076885 | 0.63319851 | 0.000655124 | 0.005790119 |
| ENSMUST00000125645.1,Nrd1-009,protein_coding,ENSMUSG00000053510.12,protein_coding | 105.2129577 | -0.571186398 | 0.000655666 | 0.005793062 |
| ENSMUST00000032168.6,Sec61a1-001,protein_coding,ENSMUSG00000030082.14,protein_coding | 63918.64933 | 0.143111382 | 0.000655877 | 0.005793229 |
| ENSMUST00000111820.3,Tmppe-001,protein_coding,ENSMUSG00000079260.3,protein_coding | 612.0914945 | -0.301031686 | 0.000655954 | 0.005793229 |
| ENSMUST00000055018.10,Lpar1-001,protein_coding,ENSMUSG00000038668.14,protein_coding | 710.0430352 | -0.369669667 | 0.000656971 | 0.005801019 |

| | | | | |
|---|---|---|---|---|
| ENSMUST00000135768.1,Pus3-002,retained_intron,ENSMUSG00000032103.10,protein_coding | 172.511285 | -0.525251308 | 0.000657911 | 0.005808127 |
| ENSMUST00000109236.8,Stau1-002,protein_coding,ENSMUSG00000039536.16,protein_coding | 3664.096531 | 0.344251003 | 0.000658277 | 0.005810165 |
| ENSMUST00000036156.5,Ipo13-001,protein_coding,ENSMUSG00000033365.14,protein_coding | 5375.318333 | -0.135169455 | 0.000659078 | 0.005816042 |
| ENSMUST00000151187.7,Tmem209-002,nonsense_mediated_decay,ENSMUSG00000029782.18,protein_coding | 148.2532052 | 0.503246448 | 0.000659431 | 0.005817967 |
| ENSMUST00000031143.12,Srd5a3-001,protein_coding,ENSMUSG00000029233.15,protein_coding | 2439.277435 | 0.251108684 | 0.00065983 | 0.005819331 |
| ENSMUST00000163966.7,Hsp90ab1-007,nonsense_mediated_decay,ENSMUSG00000023944.14,protein_coding | 11651.06322 | 0.180827105 | 0.000659856 | 0.005819331 |
| ENSMUST00000132102.1,Gtf3a-003,protein_coding,ENSMUSG00000016503.16,protein_coding | 3349.945163 | 0.231949592 | 0.000662848 | 0.005844518 |
| ENSMUST00000135780.1,Camta1-004,processed_transcript,ENSMUSG00000014592.20,protein_coding | 300.6087118 | 0.303268634 | 0.000663917 | 0.00585275 |
| ENSMUST00000029671.8,Ifi44-001,protein_coding,ENSMUSG00000028037.13,protein_coding | 155.6005211 | 0.60481298 | 0.000664305 | 0.005854974 |
| ENSMUST00000153990.7,Stradb-003,nonsense_mediated_decay,ENSMUSG00000026027.13,protein_coding | 460.9461705 | 0.424622009 | 0.000665009 | 0.005859976 |
| ENSMUST00000039061.14,Trappc11-001,protein_coding,ENSMUSG00000038102.15,protein_coding | 7725.314675 | 0.114831142 | 0.000667332 | 0.005879246 |
| ENSMUST00000151351.1,Ptgs1-004,retained_intron,ENSMUSG00000047250.13,protein_coding | 126.6191724 | 0.431515896 | 0.000667552 | 0.005879296 |
| ENSMUST00000199317.1,Lamtor5-001,protein_coding,ENSMUSG00000087260.5,protein_coding | 4534.615132 | 0.174877547 | 0.000667611 | 0.005879296 |
| ENSMUST00000102638.7,Ampd2-001,protein_coding,ENSMUSG00000027889.17,protein_coding | 2402.361541 | -0.354009844 | 0.000669034 | 0.005890616 |
| ENSMUST00000058265.7,C1galt1c1-001,protein_coding,ENSMUSG00000048970.9,protein_coding | 814.7882975 | -0.46823532 | 0.000669285 | 0.005891622 |
| ENSMUST00000133696.7,Map4k2-003,retained_intron,ENSMUSG00000024948.14,protein_coding | 161.9683314 | 0.443412173 | 0.000669871 | 0.00589558 |
| ENSMUST00000106731.2,Lrp8-003,protein_coding,ENSMUSG00000028613.14,protein_coding | 3074.304168 | -0.251217537 | 0.000671403 | 0.005907854 |
| ENSMUST00000107396.7,Atp8b2-201,protein_coding,ENSMUSG00000060671.12,protein_coding | 2997.744871 | -0.327710034 | 0.000673896 | 0.00592858 |
| ENSMUST00000200323.1,Stard3nl-003,protein_coding,ENSMUSG00000003062.12,protein_coding | 97.18287767 | 0.491260289 | 0.000674085 | 0.005929031 |
| ENSMUST00000112932.1,Zbtb6-001,protein_coding,ENSMUSG00000066798.3,protein_coding | 562.3213126 | -0.466381294 | 0.000674709 | 0.005933308 |
| ENSMUST00000135091.1,1500011K16Rik-001,protein_coding,ENSMUSG00000051319.6,protein_coding | 1472.220303 | 0.231414078 | 0.000675193 | 0.005936343 |
| ENSMUST00000159806.1,Lmo7-006,protein_coding,ENSMUSG00000033060.15,protein_coding | 4337.026029 | -0.228820971 | 0.00067615 | 0.005943547 |
| ENSMUST00000034093.14,Eif2ak3-001,protein_coding,ENSMUSG00000031668.14,protein_coding | 3551.579081 | 0.14711033 | 0.000677048 | 0.00595023 |
| ENSMUST00000212737.1,Wwp2-010,processed_transcript,ENSMUSG00000031930.11,protein_coding | 171.0078279 | -0.50235257 | 0.000677279 | 0.005951038 |
| ENSMUST00000205335.1,Mat2a-002,protein_coding,ENSMUSG00000053907.10,protein_coding | 7936.143478 | 0.189271871 | 0.000677969 | 0.005955886 |
| ENSMUST00000105421.8,Specc1l-202,protein_coding,ENSMUSG00000033444.13,protein_coding | 12964.35515 | 0.122562025 | 0.000678533 | 0.005959624 |
| ENSMUST00000135660.7,Stoml2-005,protein_coding,ENSMUSG00000028455.15,protein_coding | 4481.71943 | 0.190649595 | 0.000682202 | 0.00599063 |
| ENSMUST00000138104.8,Fgfr1-005,nonsense_mediated_decay,ENSMUSG00000031565.18,protein_coding | 467.8525736 | 0.273039541 | 0.000682541 | 0.005992388 |
| ENSMUST00000137935.1,Htra3-003,protein_coding,ENSMUSG00000029096.15,protein_coding | 307.2289381 | 0.394826654 | 0.00068323 | 0.00599721 |
| ENSMUST00000190457.1,2210011C24Rik-001,protein_coding,ENSMUSG00000074217.8,protein_coding | 251.3234545 | -0.400747755 | 0.000684056 | 0.006003232 |
| ENSMUST00000042122.14,Ssbp2-004,protein_coding,ENSMUSG00000003992.14,protein_coding | 2411.533058 | 0.181243454 | 0.000684197 | 0.006003247 |
| ENSMUST00000144310.1,Ubald2-002,processed_transcript,ENSMUSG00000050628.6,protein_coding | 152.0394761 | 0.594941458 | 0.00068607 | 0.006018455 |
| ENSMUST00000105650.7,Gpr153-002,protein_coding,ENSMUSG00000042804.13,protein_coding | 188.9697473 | -0.501058981 | 0.000687073 | 0.0060248 |
| ENSMUST00000124329.1,2810403D21Rik-001,lincRNA,ENSMUSG00000086629.8,processed_transcript | 63.4018707 | 0.542524235 | 0.000686969 | 0.0060248 |
| ENSMUST00000165199.7,Arsa-001,protein_coding,ENSMUSG00000022620.14,protein_coding | 1583.628751 | 0.216037229 | 0.000687508 | 0.006027387 |
| ENSMUST00000162175.7,Exoc5-001,protein_coding,ENSMUSG00000061244.13,protein_coding | 3137.247587 | -0.44501097 | 0.000688134 | 0.006031645 |
| ENSMUST00000080560.8,Sema3f-001,protein_coding,ENSMUSG00000034684.12,protein_coding | 609.6031495 | -0.429708306 | 0.000692668 | 0.006070149 |
| ENSMUST00000046399.6,Apmap-001,protein_coding,ENSMUSG00000033096.7,protein_coding | 3514.598126 | 0.177825259 | 0.000695809 | 0.00609644 |
| ENSMUST00000050516.12,Ascc1-201,protein_coding,ENSMUSG00000044475.13,protein_coding | 2162.449421 | 0.242559227 | 0.000696016 | 0.006097011 |
| ENSMUST00000033923.13,Dlc1-002,protein_coding,ENSMUSG00000031523.16,protein_coding | 11439.58128 | -0.164161532 | 0.000697282 | 0.006106855 |
| ENSMUST00000113682.8,Myg1-001,protein_coding,ENSMUSG00000001285.12,protein_coding | 5393.933723 | -0.17050728 | 0.000698832 | 0.006119189 |
| ENSMUST00000150717.7,Htra1-003,retained_intron,ENSMUSG00000006205.13,protein_coding | 312.5688841 | 0.509232339 | 0.000699518 | 0.006123952 |
| ENSMUST00000038090.11,Ttll7-001,retained_intron,ENSMUSG00000036745.15,protein_coding | 247.4934557 | 0.623212565 | 0.000699933 | 0.006126333 |
| ENSMUST00000084979.10,Myo7a-002,protein_coding,ENSMUSG00000030761.15,protein_coding | 411.5874611 | 0.600851248 | 0.000701352 | 0.006137505 |
| ENSMUST00000058856.8,Scd4-001,protein_coding,ENSMUSG00000050195.8,protein_coding | 60.44537185 | 0.585058521 | 0.000703196 | 0.006152397 |
| ENSMUST00000138621.1,Rpl6-006,retained_intron,ENSMUSG00000029614.13,protein_coding | 170.474893 | -0.459609178 | 0.000703341 | 0.006152407 |
| ENSMUST00000040321.12,Trmt10a-001,protein_coding,ENSMUSG00000004127.12,protein_coding | 1169.672766 | -0.194358457 | 0.000705054 | 0.006166147 |
| ENSMUST00000022998.13,Mtbp-001,protein_coding,ENSMUSG00000022369.13,protein_coding | 11870.53349 | 0.151228682 | 0.000705406 | 0.006167967 |
| ENSMUST00000093217.8,B3gnt9-201,protein_coding,ENSMUSG00000069920.10,protein_coding | 573.8834524 | -0.308945183 | 0.000707657 | 0.00618639 |
| ENSMUST00000020312.12,Mcu-001,protein_coding,ENSMUSG00000009647.13,protein_coding | 16644.87171 | 0.158989341 | 0.000709141 | 0.006198108 |
| ENSMUST00000028910.8,Mgme1-002,protein_coding,ENSMUSG00000027424.11,protein_coding | 777.7637636 | 0.279196039 | 0.000709404 | 0.006199145 |
| ENSMUST00000026681.6,Tma16-001,protein_coding,ENSMUSG00000025591.6,protein_coding | 1750.928954 | 0.208092861 | 0.000710312 | 0.006205822 |
| ENSMUST00000151130.7,Shf-002,protein_coding,ENSMUSG00000033256.14,protein_coding | 784.7454873 | 0.287157267 | 0.00071062 | 0.006206812 |
| ENSMUST00000117066.7,Pigt-003,protein_coding,ENSMUSG00000017721.15,protein_coding | 48.56474784 | 0.631949917 | 0.000710714 | 0.006206812 |
| ENSMUST00000159785.1,Wnk1-009,protein_coding,ENSMUSG00000045962.16,protein_coding | 27.13583148 | 0.63181761 | 0.000711283 | 0.00620955 |
| ENSMUST00000186193.6,Pgm3-004,retained_intron,ENSMUSG00000056131.13,protein_coding | 230.2636326 | 0.505546082 | 0.000711316 | 0.00620955 |
| ENSMUST00000148219.8,Drap1-009,protein_coding,ENSMUSG00000024914.16,protein_coding | 1551.544915 | 0.209755766 | 0.000711962 | 0.006213934 |
| ENSMUST00000215547.1,Dse-003,protein_coding,ENSMUSG00000039497.8,protein_coding | 32.84473901 | -0.631438046 | 0.000712139 | 0.006214215 |
| ENSMUST00000120016.1,Zbtb9-001,protein_coding,ENSMUSG00000079605.4,protein_coding | 1743.159426 | 0.166274925 | 0.000712893 | 0.006219538 |
| ENSMUST00000189220.6,Agfg1-001,protein_coding,ENSMUSG00000026159.13,protein_coding | 5059.867695 | -0.319581922 | 0.000713231 | 0.006221221 |
| ENSMUST00000077977.13,Miga2-001,protein_coding,ENSMUSG00000026858.18,protein_coding | 4410.031471 | -0.180362523 | 0.000715031 | 0.006234397 |
| ENSMUST00000163048.7,Krr1-001,protein_coding,ENSMUSG00000063334.16,protein_coding | 721.1703706 | -0.352796881 | 0.000714889 | 0.006234397 |
| ENSMUST00000132590.7,Rad18-003,processed_transcript,ENSMUSG00000030254.16,protein_coding | 3655.635866 | -0.180780764 | 0.000715373 | 0.006234855 |
| ENSMUST00000180072.7,Atp5h-201,protein_coding,ENSMUSG00000034566.10,protein_coding | 1803.894261 | 0.367373706 | 0.000715256 | 0.006234855 |
| ENSMUST00000156513.1,Csnk1e-007,retained_intron,ENSMUSG00000022433.18,protein_coding | 181.1097522 | -0.478258163 | 0.0007156 | 0.006235573 |
| ENSMUST00000173341.7,Gtf2i-002,protein_coding,ENSMUSG00000060261.15,protein_coding | 13133.2698 | 0.226968249 | 0.000716487 | 0.006242034 |
| ENSMUST00000147512.1,Trp53-003,processed_transcript,ENSMUSG00000059552.13,protein_coding | 330.9867898 | -0.347736572 | 0.000718096 | 0.006254785 |
| ENSMUST00000006181.6,Napa-001,protein_coding,ENSMUSG00000006024.13,protein_coding | 14268.52202 | 0.15813711 | 0.000718389 | 0.006254812 |
| ENSMUST00000172906.7,Ankrd11-002,nonsense_mediated_decay,ENSMUSG00000035569.17,protein_coding | 7988.580827 | 0.3552129 | 0.000718334 | 0.006254812 |
| ENSMUST00000185176.7,Lzts1-001,protein_coding,ENSMUSG00000036306.13,protein_coding | 95.887318 | 0.592806012 | 0.000719347 | 0.006261886 |
| ENSMUST00000126491.1,Mrps5-004,retained_intron,ENSMUSG00000027374.12,protein_coding | 1054.689656 | 0.225025789 | 0.000720597 | 0.006271498 |
| ENSMUST00000038341.7,Bub1b-001,protein_coding,ENSMUSG00000040084.9,protein_coding | 22399.41968 | 0.105048327 | 0.000721336 | 0.006276662 |
| ENSMUST00000038675.6,Mrpl23-001,protein_coding,ENSMUSG00000037772.13,protein_coding | 1981.553719 | 0.253974437 | 0.000721508 | 0.006276884 |
| ENSMUST00000030849.12,Fam126a-001,protein_coding,ENSMUSG00000028995.14,protein_coding | 3897.345748 | 0.157939311 | 0.000721843 | 0.006278529 |
| ENSMUST00000139699.7,Exoc7-004,processed_transcript,ENSMUSG00000020792.15,protein_coding | 104.7124705 | -0.631137828 | 0.000722722 | 0.006284911 |
| ENSMUST00000019470.13,Psme3-001,protein_coding,ENSMUSG00000078652.9,protein_coding | 24351.90316 | 0.134705442 | 0.000723818 | 0.006293166 |
| ENSMUST00000010904.4,Phlda2-001,protein_coding,ENSMUSG00000010760.5,protein_coding | 152.560058 | 0.415293592 | 0.000724954 | 0.006301775 |
| ENSMUST00000112522.8,Mbtps2-003,protein_coding,ENSMUSG00000046873.18,protein_coding | 67.43975776 | -0.615613177 | 0.000726648 | 0.006315217 |
| ENSMUST00000197349.1,Zfp316-002,retained_intron,ENSMUSG00000046658.16,protein_coding | 77.38225729 | 0.630003928 | 0.000727042 | 0.006316099 |
| ENSMUST00000095273.6,Nr2c2ap-001,protein_coding,ENSMUSG00000071078.6,protein_coding | 2308.208142 | 0.228320355 | 0.000726917 | 0.006316099 |
| ENSMUST00000099441.4,Slc35d2-201,protein_coding,ENSMUSG00000033114.10,protein_coding | 699.610267 | 0.22613278 | 0.000727926 | 0.006322503 |
| ENSMUST00000191046.6,Epb41l5-001,protein_coding,ENSMUSG00000026383.14,protein_coding | 65.74739253 | -0.556087574 | 0.000728847 | 0.006329223 |
| ENSMUST00000211389.1,Ankrd10-005,retained_intron,ENSMUSG00000031508.14,protein_coding | 345.2071363 | -0.443607545 | 0.00072927 | 0.006331618 |
| ENSMUST00000028650.8,Pex16-001,protein_coding,ENSMUSG00000027222.14,protein_coding | 1434.756018 | 0.245345242 | 0.000729784 | 0.006334804 |
| ENSMUST00000028341.10,Anapc2-001,protein_coding,ENSMUSG00000026965.12,protein_coding | 12310.87053 | -0.143239916 | 0.000729966 | 0.0063351 |
| ENSMUST00000070166.5,Gramd3-201,protein_coding,ENSMUSG00000001700.9,protein_coding | 496.4846995 | 0.273064096 | 0.000730415 | 0.00633772 |
| ENSMUST00000040248.7,Spidr-201,protein_coding,ENSMUSG00000041974.9,protein_coding | 4220.272495 | 0.144669212 | 0.000731752 | 0.006348044 |
| SMUST00000139264.1,2610005L07Rik-002,transcribed_unprocessed_pseudogene,ENSMUSG00000071793.12,transcribed_unprocessed_pseudoge | 2390.796217 | 0.351080229 | 0.000732451 | 0.006352831 |
| ENSMUST00000198389.4,Dcun1d1-002,protein_coding,ENSMUSG00000027708.14,protein_coding | 1235.05139 | -0.451320823 | 0.000732814 | 0.006354698 |
| ENSMUST00000142493.1,Wbp1-007,retained_intron,ENSMUSG00000030035.14,protein_coding | 241.7123056 | -0.527834213 | 0.000734115 | 0.0063647 |
| ENSMUST00000040853.10,Oxsr1-001,protein_coding,ENSMUSG00000036737.12,protein_coding | 9119.165883 | 0.164210042 | 0.000734607 | 0.006367676 |
| ENSMUST00000121655.1,Got2-ps1-001,processed_pseudogene,ENSMUSG00000080935.3,processed_pseudogene | 340.3002247 | -0.38468865 | 0.000735153 | 0.006371128 |

| | | | | |
|---|---|---|---|---|
| ENSMUST00000114136.8,Pml-002,protein_coding,ENSMUSG00000036986.16,protein_coding | 1249.062624 | 0.288551083 | 0.000735784 | 0.006375315 |
| ENSMUST00000113145.7,Gprasp1-003,protein_coding,ENSMUSG00000043384.15,protein_coding | 514.1592972 | -0.475194368 | 0.000736412 | 0.006379473 |
| ENSMUST00000084296.9,Cdca8-001,protein_coding,ENSMUSG00000028873.16,protein_coding | 13222.23866 | 0.136451013 | 0.000737642 | 0.006388844 |
| ENSMUST00000046897.12,Trim14-001,protein_coding,ENSMUSG00000039853.18,protein_coding | 125.3553539 | 0.483428156 | 0.000738231 | 0.006392654 |
| ENSMUST00000045887.8,Cisd1-001,protein_coding,ENSMUSG00000037710.8,protein_coding | 7242.530092 | 0.155374993 | 0.000739809 | 0.006405036 |
| ENSMUST00000047935.7,Tspyl4-001,protein_coding,ENSMUSG00000039485.8,protein_coding | 2730.353574 | 0.26061023 | 0.000740138 | 0.006406589 |
| ENSMUST00000099443.10,Zfyve27-003,protein_coding,ENSMUSG00000018820.13,protein_coding | 3157.246724 | 0.193269741 | 0.000740985 | 0.006412634 |
| ENSMUST00000132378.1,2010001A14Rik-002,processed_transcript,ENSMUSG00000087165.7,processed_transcript | 28.69943202 | -0.629162433 | 0.000743725 | 0.006435051 |
| ENSMUST00000019749.3,Hoxd8-001,protein_coding,ENSMUSG00000027102.4,protein_coding | 1307.195186 | -0.248951427 | 0.000745741 | 0.006451202 |
| ENSMUST00000201908.3,Ociad2-006,protein_coding,ENSMUSG00000029153.11,protein_coding | 25.44587919 | -0.611447793 | 0.000745939 | 0.006451618 |
| ENSMUST00000190874.6,Nadk2-003,processed_transcript,ENSMUSG00000022253.14,protein_coding | 329.2781224 | 0.458023002 | 0.000748389 | 0.006471504 |
| ENSMUST00000038794.4,Dpp9-201,protein_coding,ENSMUSG00000001229.7,protein_coding | 14915.18612 | 0.104612298 | 0.000751716 | 0.00649897 |
| ENSMUST00000025893.5,Arl2-001,protein_coding,ENSMUSG00000024944.5,protein_coding | 6009.385467 | 0.175981941 | 0.000752101 | 0.006500994 |
| ENSMUST00000065977.10,Nrd1-001,protein_coding,ENSMUSG00000053510.12,protein_coding | 9215.734487 | 0.230633537 | 0.000752768 | 0.006504312 |
| ENSMUST00000119946.7,Pus7-001,protein_coding,ENSMUSG00000057541.14,protein_coding | 9845.40363 | 0.167392006 | 0.000752787 | 0.006504312 |
| ENSMUST00000038411.4,Adat3-201,protein_coding,ENSMUSG00000035370.7,protein_coding | 569.6372421 | 0.295388803 | 0.000754067 | 0.00651406 |
| ENSMUST00000013807.7,Pten-001,protein_coding,ENSMUSG00000013663.7,protein_coding | 10372.76093 | -0.351737914 | 0.000755041 | 0.006521168 |
| ENSMUST00000147660.7,Rpl9-004,retained_intron,ENSMUSG00000047215.14,protein_coding | 99.56205322 | -0.480719321 | 0.000756922 | 0.006536103 |
| ENSMUST00000118853.7,Fam198b-001,protein_coding,ENSMUSG00000027955.16,protein_coding | 1665.93383 | -0.448476876 | 0.000757461 | 0.00653944 |
| ENSMUST00000171644.7,Pparg-001,protein_coding,ENSMUSG00000000440.12,protein_coding | 2658.077279 | 0.178231831 | 0.000757913 | 0.006542036 |
| ENSMUST00000033913.10,Dctn6-001,protein_coding,ENSMUSG00000031516.11,protein_coding | 2042.37911 | -0.278682938 | 0.000759064 | 0.006550659 |
| ENSMUST00000103175.7,Ube2j2-003,protein_coding,ENSMUSG00000023286.16,protein_coding | 4978.762746 | 0.180007878 | 0.00076119 | 0.006567686 |
| ENSMUST00000149359.1,Mkl2-001,protein_coding,ENSMUSG00000009569.14,protein_coding | 3853.068381 | -0.178134337 | 0.000761682 | 0.006570615 |
| ENSMUST00000122845.7,Copa-008,retained_intron,ENSMUSG00000026553.17,protein_coding | 167.6842199 | -0.577511669 | 0.000765247 | 0.006600043 |
| ENSMUST00000185148.7,Mrpl20-005,protein_coding,ENSMUSG00000029066.12,protein_coding | 69.96443143 | 0.628069097 | 0.000766096 | 0.006606048 |
| ENSMUST00000176047.7,Atpaf1-001,protein_coding,ENSMUSG00000028710.16,protein_coding | 1940.613467 | -0.170743539 | 0.00076649 | 0.006608119 |
| ENSMUST00000118869.7,Ick-004,protein_coding,ENSMUSG00000009828.15,protein_coding | 1860.300677 | 0.247532378 | 0.000767262 | 0.00661345 |
| ENSMUST00000136079.7,Ciz1-008,protein_coding,ENSMUSG00000039205.16,protein_coding | 132.5201431 | 0.553834259 | 0.000767964 | 0.006618176 |
| ENSMUST00000108431.2,Myo1c-004,protein_coding,ENSMUSG00000017774.19,protein_coding | 11205.43135 | 0.145270817 | 0.000768211 | 0.006618976 |
| ENSMUST00000022446.5,Eaf1-201,protein_coding,ENSMUSG00000021890.5,protein_coding | 15163.94947 | 0.186574599 | 0.000768474 | 0.006619916 |
| ENSMUST00000133930.7,BC030867-001,nonsense_mediated_decay,ENSMUSG00000034773.16,protein_coding | 94.29625197 | 0.619986962 | 0.000768761 | 0.006621063 |
| ENSMUST00000160780.1,Hrh1-005,protein_coding,ENSMUSG00000053004.9,protein_coding | 565.7658691 | 0.402075899 | 0.000769304 | 0.006624416 |
| ENSMUST00000043815.15,Nup205-002,protein_coding,ENSMUSG00000038759.15,protein_coding | 72939.94368 | 0.130979453 | 0.000771031 | 0.006637957 |
| ENSMUST00000034827.9,Imp3-001,protein_coding,ENSMUSG00000032288.9,protein_coding | 4185.428568 | 0.141408072 | 0.00077287 | 0.006652465 |
| ENSMUST00000149486.1,Bptf-003,protein_coding,ENSMUSG00000040481.17,protein_coding | 31.331064 | 0.61942981 | 0.00077321 | 0.006654054 |
| ENSMUST00000191450.6,Tardbp-009,nonsense_mediated_decay,ENSMUSG00000041459.15,protein_coding | 426.8890259 | 0.403005005 | 0.000773669 | 0.006656677 |
| ENSMUST00000049507.5,Pcsk9-001,protein_coding,ENSMUSG00000044254.6,protein_coding | 1396.561084 | 0.31002992 | 0.00077394 | 0.00665768 |
| ENSMUST00000052236.12,Fbxo10-001,protein_coding,ENSMUSG00000048232.12,protein_coding | 2217.591831 | 0.165386349 | 0.000774269 | 0.006658341 |
| ENSMUST00000032487.13,Vamp1-001,protein_coding,ENSMUSG00000030337.16,protein_coding | 36.34615168 | 0.586544918 | 0.000774326 | 0.006658341 |
| ENSMUST00000129369.1,Rfc5-005,retained_intron,ENSMUSG00000029363.14,protein_coding | 349.8198244 | -0.32821363 | 0.000776029 | 0.006671647 |
| ENSMUST00000025642.13,Prpf19-001,protein_coding,ENSMUSG00000024735.13,protein_coding | 28655.4344 | 0.143028922 | 0.000777505 | 0.006683006 |
| ENSMUST00000216281.1,Cep63-011,protein_coding,ENSMUSG00000032534.17,protein_coding | 80.82215876 | 0.445563047 | 0.000778513 | 0.006690328 |
| ENSMUST00000182179.7,Nktr-003,nonsense_mediated_decay,ENSMUSG00000032525.15,protein_coding | 76.48934973 | 0.561105597 | 0.000779251 | 0.006695338 |
| ENSMUST00000109418.1,Fitm2-001,protein_coding,ENSMUSG00000048486.6,protein_coding | 4131.066651 | -0.157072123 | 0.000780748 | 0.00670686 |
| ENSMUST00000070801.10,Alg13-001,protein_coding,ENSMUSG00000041718.15,protein_coding | 168.5229225 | -0.541401702 | 0.000780954 | 0.006707294 |
| ENSMUST00000039442.11,Alg14-001,protein_coding,ENSMUSG00000039887.11,protein_coding | 1175.121162 | -0.23559784 | 0.000782095 | 0.006715747 |
| ENSMUST00000091614.6,Nfx1-007,protein_coding,ENSMUSG00000028423.15,protein_coding | 4921.09628 | 0.170088703 | 0.000782485 | 0.006717758 |
| ENSMUST00000108324.3,Ccne2-002,protein_coding,ENSMUSG00000028212.16,protein_coding | 1664.233271 | -0.495428159 | 0.000782657 | 0.0067179 |
| ENSMUST00000144920.3,Cmas-003,nonsense_mediated_decay,ENSMUSG00000030282.14,protein_coding | 457.8000531 | -0.261742137 | 0.000782951 | 0.00671908 |
| ENSMUST00000102650.9,Akap10-001,protein_coding,ENSMUSG00000047804.15,protein_coding | 3912.463837 | -0.110146332 | 0.000783585 | 0.006723178 |
| ENSMUST00000204169.1,Gm45140-001,protein_coding,ENSMUSG00000107928.1,protein_coding | 296.5036953 | 0.417543103 | 0.000784578 | 0.006730359 |
| ENSMUST00000022292.9,Samd8-001,protein_coding,ENSMUSG00000021770.10,protein_coding | 312.096416 | 0.527961429 | 0.000785506 | 0.00673698 |
| ENSMUST00000108940.2,Gm14403-201,protein_coding,ENSMUSG00000094786.1,protein_coding | 28.61255852 | -0.621211768 | 0.00078629 | 0.006741018 |
| ENSMUST00000182008.7,Ank2-003,protein_coding,ENSMUSG00000032826.17,protein_coding | 804.9986725 | 0.253115102 | 0.000786139 | 0.006741018 |
| ENSMUST00000173867.7,Cct4-001,protein_coding,ENSMUSG00000007739.10,protein_coding | 28261.33716 | 0.085242173 | 0.000786904 | 0.006744933 |
| ENSMUST00000142701.2,Brap-007,protein_coding,ENSMUSG00000029458.14,protein_coding | 640.2844818 | 0.326760667 | 0.000787246 | 0.006746528 |
| ENSMUST00000107491.10,Dap3-002,protein_coding,ENSMUSG00000068921.14,protein_coding | 5797.232378 | 0.186628367 | 0.000787528 | 0.006747595 |
| ENSMUST00000201972.1,Bend4-001,protein_coding,ENSMUSG00000092060.4,protein_coding | 1038.425975 | -0.570438812 | 0.000789934 | 0.006766862 |
| ENSMUST00000192388.1,Gm36958-001,TEC,ENSMUSG00000102204.1,TEC | 42.6070879 | -0.616514698 | 0.000792047 | 0.006783619 |
| ENSMUST00000114547.7,Cadm1-003,protein_coding,ENSMUSG00000032076.19,protein_coding | 10102.36077 | -0.147498161 | 0.000793017 | 0.006790572 |
| ENSMUST00000107556.9,Tsacc-001,protein_coding,ENSMUSG00000010538.14,protein_coding | 127.7494918 | 0.45347327 | 0.000797358 | 0.006826383 |
| ENSMUST00000121430.1,Gm12715-001,processed_pseudogene,ENSMUSG00000059195.10,processed_pseudogene | 5522.554365 | -0.382111594 | 0.000798096 | 0.006831346 |
| ENSMUST00000174552.7,Dip2c-001,protein_coding,ENSMUSG00000048264.15,protein_coding | 3767.762743 | 0.229653064 | 0.000798346 | 0.006832131 |
| ENSMUST00000198527.1,4632415L05Rik-001,lincRNA,ENSMUSG00000048106.5,lincRNA | 1296.888686 | 0.185618085 | 0.000800066 | 0.006845484 |
| ENSMUST00000048716.10,Fgd3-002,protein_coding,ENSMUSG00000037946.11,protein_coding | 343.2410044 | -0.490862092 | 0.000800662 | 0.006849223 |
| ENSMUST00000111194.7,Cd44-201,protein_coding,ENSMUSG00000005087.17,protein_coding | 33099.2105 | 0.150427637 | 0.000801272 | 0.006852921 |
| ENSMUST00000079012.12,March8-001,protein_coding,ENSMUSG00000025702.15,protein_coding | 3926.906052 | 0.358725029 | 0.000801413 | 0.006852921 |
| ENSMUST00000010899.13,Cars-001,protein_coding,ENSMUSG00000010755.17,protein_coding | 3309.694764 | 0.177288309 | 0.000801749 | 0.00685444 |
| ENSMUST00000127252.7,Rpn2-003,processed_transcript,ENSMUSG00000027642.15,protein_coding | 26636.07444 | 0.159588341 | 0.00080221 | 0.006857018 |
| ENSMUST00000028580.11,Ccdc34-001,protein_coding,ENSMUSG00000027160.16,protein_coding | 5907.627278 | 0.20649479 | 0.000803213 | 0.006863763 |
| ENSMUST00000123084.7,Senp3-008,retained_intron,ENSMUSG00000005204.12,protein_coding | 79.17524444 | 0.507546651 | 0.000803318 | 0.006863763 |
| ENSMUST00000099482.3,Grk6-202,protein_coding,ENSMUSG00000074886.10,protein_coding | 338.0856132 | -0.539696661 | 0.000803731 | 0.006865929 |
| ENSMUST00000105799.7,Arhgef10l-003,protein_coding,ENSMUSG00000040964.16,protein_coding | 255.3007024 | 0.622866653 | 0.000805078 | 0.006874705 |
| ENSMUST00000031678.9,Tspan12-001,protein_coding,ENSMUSG00000029669.14,protein_coding | 885.3836523 | -0.37324893 | 0.000804984 | 0.006874705 |
| ENSMUST00000035090.13,Fbxl2-001,protein_coding,ENSMUSG00000032507.15,protein_coding | 211.2604381 | 0.392360694 | 0.000805413 | 0.006876204 |
| ENSMUST00000168527.7,Dennd1b-201,protein_coding,ENSMUSG00000056268.15,protein_coding | 2299.759238 | -0.330786697 | 0.000807168 | 0.006889819 |
| ENSMUST00000025827.8,Minpp1-201,protein_coding,ENSMUSG00000024896.8,protein_coding | 7795.828674 | 0.11668411 | 0.000807498 | 0.006891266 |
| ENSMUST00000013633.11,Fgfrl1-001,protein_coding,ENSMUSG00000008090.14,protein_coding | 3893.883344 | 0.172168462 | 0.000808892 | 0.006900634 |
| ENSMUST00000025751.9,Ighmbp2-001,protein_coding,ENSMUSG00000024831.13,protein_coding | 4107.18566 | 0.156058354 | 0.000808916 | 0.006900634 |
| ENSMUST00000101426.10,Tmem176a-001,protein_coding,ENSMUSG00000023367.14,protein_coding | 1007.24412 | -0.291931609 | 0.000809518 | 0.006904404 |
| ENSMUST00000074425.12,Efcab14-001,protein_coding,ENSMUSG00000034210.13,protein_coding | 2378.908156 | -0.152058306 | 0.000810983 | 0.006915525 |
| ENSMUST00000110639.7,Map1a-002,protein_coding,ENSMUSG00000027254.13,protein_coding | 2223.182565 | -0.204773465 | 0.000814796 | 0.00694639 |
| ENSMUST00000196921.4,Fdps-005,protein_coding,ENSMUSG00000059743.12,protein_coding | 226.2295597 | 0.516303245 | 0.000814984 | 0.00694639 |
| ENSMUST00000212421.1,Slc7a6-004,protein_coding,ENSMUSG00000031904.5,protein_coding | 23.33017797 | 0.601155567 | 0.000815087 | 0.00694639 |
| ENSMUST00000026439.13,Nabp2-001,protein_coding,ENSMUSG00000025374.13,protein_coding | 7273.148133 | 0.14924109 | 0.000816656 | 0.006958386 |
| ENSMUST00000165561.2,Smim13-001,protein_coding,ENSMUSG00000091264.2,protein_coding | 2202.179073 | 0.215037777 | 0.000817384 | 0.00696321 |
| ENSMUST00000103209.9,Mmp28-003,protein_coding,ENSMUSG00000020682.17,protein_coding | 578.456179 | 0.2897046 | 0.000817748 | 0.006964935 |
| ENSMUST00000014445.6,Pam16-001,protein_coding,ENSMUSG00000014301.12,protein_coding | 2931.219497 | 0.137075166 | 0.000818578 | 0.006970624 |
| ENSMUST00000182709.7,Rnf4-002,protein_coding,ENSMUSG00000029110.13,protein_coding | 11684.64161 | 0.124278871 | 0.000820489 | 0.006985517 |
| ENSMUST00000092723.10,Arid1b-002,protein_coding,ENSMUSG00000069729.13,protein_coding | 7317.224734 | 0.120597417 | 0.000820781 | 0.00698662 |
| ENSMUST00000068975.5,Zfp180-002,protein_coding,ENSMUSG00000057101.8,protein_coding | 3006.622391 | 0.20887975 | 0.000820983 | 0.006986958 |
| ENSMUST00000134775.2,Ehmt1-004,nonsense_mediated_decay,ENSMUSG00000036893.17,protein_coding | 42.50987499 | -0.62342861 | 0.000822304 | 0.006996816 |

| | | | | |
|---|---|---|---|---|
| ENSMUST00000110167.4,Ndufa13-001,protein_coding,ENSMUSG00000036199.9,protein_coding | 10959.52638 | 0.160228968 | 0.000824703 | 0.007015839 |
| ENSMUST00000029566.8,Efna1-001,protein_coding,ENSMUSG00000027954.9,protein_coding | 1278.76404 | 0.200849062 | 0.000827499 | 0.007038238 |
| ENSMUST00000162500.7,Rps6kc1-002,retained_intron,ENSMUSG00000089872.10,protein_coding | 96.30323084 | 0.58867055 | 0.000828409 | 0.007044583 |
| ENSMUST00000001185.13,Gmcl1-001,protein_coding,ENSMUSG00000001157.13,protein_coding | 4262.289862 | -0.150000971 | 0.000830256 | 0.0070589 |
| ENSMUST00000131869.7,Camk2d-003,processed_transcript,ENSMUSG00000053819.16,protein_coding | 893.6526754 | 0.461076632 | 0.00083162 | 0.007069101 |
| ENSMUST00000130103.1,Mesdc2-002,protein_coding,ENSMUSG00000038503.15,protein_coding | 738.5347788 | 0.259728418 | 0.000831825 | 0.007069443 |
| ENSMUST00000035214.10,Ip6k1-001,protein_coding,ENSMUSG00000032594.11,protein_coding | 3918.691281 | -0.14055189 | 0.000832733 | 0.007075765 |
| ENSMUST00000030675.7,Mrps15-001,protein_coding,ENSMUSG00000028861.13,protein_coding | 6828.333145 | 0.087661606 | 0.00083357 | 0.007078686 |
| ENSMUST00000024004.8,Ccl25-002,protein_coding,ENSMUSG00000023235.14,protein_coding | 148.7385649 | 0.591426119 | 0.000833423 | 0.007078686 |
| ENSMUST00000153524.1,Calm1-003,retained_intron,ENSMUSG00000001175.13,protein_coding | 4074.962887 | 0.167070401 | 0.000833518 | 0.007078686 |
| ENSMUST00000143965.7,Calm2-003,retained_intron,ENSMUSG00000036438.12,protein_coding | 212.2033903 | -0.503341403 | 0.000834334 | 0.007083775 |
| ENSMUST00000023179.5,Zfp7-201,protein_coding,ENSMUSG00000033669.7,protein_coding | 3644.990634 | 0.125762883 | 0.000835395 | 0.00708999 |
| ENSMUST00000023598.14,Rfc4-001,protein_coding,ENSMUSG00000022881.14,protein_coding | 5227.524407 | 0.145650722 | 0.000835232 | 0.00708999 |
| ENSMUST00000210564.1,Chmp1b-001,protein_coding,ENSMUSG00000109901.1,protein_coding | 1799.022707 | -0.285171368 | 0.000837265 | 0.007104455 |
| ENSMUST00000038913.15,Cnot3-002,protein_coding,ENSMUSG00000035632.16,protein_coding | 9226.838789 | 0.156192073 | 0.000839571 | 0.007122622 |
| ENSMUST00000016115.4,Actr8-201,protein_coding,ENSMUSG00000015971.4,protein_coding | 4595.188851 | 0.168964716 | 0.000839752 | 0.007122755 |
| ENSMUST00000092143.11,Msrb3-001,protein_coding,ENSMUSG00000051236.13,protein_coding | 4135.568957 | -0.151061907 | 0.00084269 | 0.007146264 |
| ENSMUST00000130802.7,Ikbkg-009,protein_coding,ENSMUSG00000004221.16,protein_coding | 100.3868492 | 0.615491288 | 0.000843648 | 0.00715298 |
| ENSMUST00000057669.15,Mapk3-001,protein_coding,ENSMUSG00000063065.13,protein_coding | 22756.48789 | 0.125340482 | 0.000844667 | 0.007160207 |
| ENSMUST00000032108.10,Wdr54-006,processed_transcript,ENSMUSG00000030032.12,protein_coding | 562.856637 | 0.284502207 | 0.000845171 | 0.007163073 |
| ENSMUST00000030243.7,Prkaa2-001,protein_coding,ENSMUSG00000028518.8,protein_coding | 188.9571982 | -0.418286646 | 0.000845377 | 0.007163404 |
| ENSMUST00000135886.1,Zfp688-002,retained_intron,ENSMUSG00000045251.13,protein_coding | 178.5436212 | 0.399983717 | 0.000847517 | 0.007180129 |
| ENSMUST00000112414.7,Acrbp-002,protein_coding,ENSMUSG00000072770.9,protein_coding | 100.5743691 | 0.507059127 | 0.000850222 | 0.00720021 |
| ENSMUST00000133170.1,Taf15-005,protein_coding,ENSMUSG00000020680.15,protein_coding | 1434.321478 | 0.382944081 | 0.000850079 | 0.00720021 |
| ENSMUST00000187507.6,Soat1-004,protein_coding,ENSMUSG00000026600.12,protein_coding | 610.1720705 | 0.325742985 | 0.000850744 | 0.007201792 |
| ENSMUST00000058585.13,Tceanc2-010,protein_coding,ENSMUSG00000028619.15,protein_coding | 496.4272887 | 0.309757555 | 0.000850739 | 0.007201792 |
| ENSMUST00000186764.6,Nab1-003,protein_coding,ENSMUSG00000002881.14,protein_coding | 730.7746334 | -0.451876351 | 0.00085278 | 0.007217609 |
| ENSMUST00000147779.7,Ddx31-002,processed_transcript,ENSMUSG00000026806.15,protein_coding | 54.76620312 | 0.578185946 | 0.000854035 | 0.007226815 |
| ENSMUST00000014642.9,Ankrd52-001,protein_coding,ENSMUSG00000014498.9,protein_coding | 20639.83423 | -0.096698698 | 0.000854869 | 0.007232444 |
| ENSMUST00000167304.7,Efemp2-004,protein_coding,ENSMUSG00000024909.14,protein_coding | 225.6221141 | -0.461587104 | 0.000857107 | 0.007249953 |
| ENSMUST00000200455.1,Gltscr2-004,processed_transcript,ENSMUSG00000041560.12,protein_coding | 1666.582006 | 0.151098581 | 0.000857477 | 0.007251659 |
| ENSMUST00000142429.1,Gm13056-001,lincRNA,ENSMUSG00000085395.1,lincRNA | 642.0228359 | 0.279948576 | 0.000861174 | 0.007281494 |
| ENSMUST00000060945.11,Aff4-001,protein_coding,ENSMUSG00000049470.13,protein_coding | 20638.03763 | -0.32495833 | 0.000861596 | 0.007283634 |
| ENSMUST00000107476.7,Mef2a-004,protein_coding,ENSMUSG00000030557.17,protein_coding | 1063.060274 | -0.306895044 | 0.000862819 | 0.00729254 |
| ENSMUST00000202868.3,Ctbp1-001,protein_coding,ENSMUSG00000037373.12,protein_coding | 20547.12332 | 0.144033683 | 0.000863354 | 0.007295628 |
| ENSMUST00000067232.9,Ctnnd1-002,protein_coding,ENSMUSG00000034101.14,protein_coding | 311.7789024 | 0.606319636 | 0.000865965 | 0.007316253 |
| ENSMUST00000156045.7,Hdac7-009,nonsense_mediated_decay,ENSMUSG00000022475.18,protein_coding | 1408.158397 | 0.351856571 | 0.000866846 | 0.007322267 |
| ENSMUST00000060820.7,Nkx3-2-001,protein_coding,ENSMUSG00000049691.8,protein_coding | 95.77622751 | -0.501248215 | 0.000870403 | 0.007350867 |
| ENSMUST00000043204.7,Fbxo33-001,protein_coding,ENSMUSG00000035329.7,protein_coding | 1724.222256 | -0.248775282 | 0.000870581 | 0.007350924 |
| ENSMUST00000034547.5,Acat1-001,protein_coding,ENSMUSG00000032047.5,protein_coding | 6237.490393 | 0.127465896 | 0.000871863 | 0.007360311 |
| ENSMUST00000152830.8,Satb1-002,protein_coding,ENSMUSG00000023927.15,protein_coding | 238.0868502 | -0.361072401 | 0.000872828 | 0.007367013 |
| ENSMUST00000144309.1,AU022252-003,processed_transcript,ENSMUSG00000078584.9,protein_coding | 148.4844047 | 0.395076722 | 0.000873861 | 0.007374284 |
| ENSMUST00000053766.13,Amfr-001,protein_coding,ENSMUSG00000031751.14,protein_coding | 26512.48644 | 0.202409842 | 0.000874185 | 0.007375569 |
| ENSMUST00000069009.6,Maf-001,protein_coding,ENSMUSG00000055435.6,protein_coding | 506.4387589 | -0.261028703 | 0.00087478 | 0.007376259 |
| ENSMUST00000212923.1,Gas8-002,protein_coding,ENSMUSG00000040220.10,protein_coding | 273.8126061 | 0.348818901 | 0.000874575 | 0.007376259 |
| ENSMUST00000118077.7,Vezt-007,protein_coding,ENSMUSG00000036099.16,protein_coding | 699.4245509 | 0.303565815 | 0.000874639 | 0.007376259 |
| ENSMUST00000188747.1,Gm28229-001,sense_intronic,ENSMUSG00000101335.1,sense_intronic | 218.9095074 | -0.583780891 | 0.000877419 | 0.007397056 |
| ENSMUST00000132010.7,Trp53inp1-002,processed_transcript,ENSMUSG00000028211.11,protein_coding | 290.7205279 | 0.620763305 | 0.000880403 | 0.007420764 |
| ENSMUST00000072750.12,Hnrnpd-005,protein_coding,ENSMUSG00000000568.15,protein_coding | 4713.720392 | 0.269039571 | 0.000885669 | 0.007460763 |
| ENSMUST00000166468.1,Ptprk-201,protein_coding,ENSMUSG00000019889.9,protein_coding | 9595.343581 | 0.136716298 | 0.000885663 | 0.007460763 |
| ENSMUST00000065113.13,Glis3-003,nonsense_mediated_decay,ENSMUSG00000052942.13,protein_coding | 321.6406069 | -0.475349098 | 0.000885601 | 0.007460763 |
| ENSMUST00000099294.8,Ablim1-001,protein_coding,ENSMUSG00000025085.16,protein_coding | 797.897368 | -0.386203581 | 0.000885942 | 0.007461607 |
| ENSMUST00000033176.6,Uqcrc2-001,protein_coding,ENSMUSG00000030884.12,protein_coding | 13234.34688 | 0.139292471 | 0.000887649 | 0.007474522 |
| ENSMUST00000028135.14,Fam129b-001,protein_coding,ENSMUSG00000026796.16,protein_coding | 52232.02112 | 0.12668977 | 0.000888077 | 0.007475874 |
| ENSMUST00000111198.8,Cd44-004,protein_coding,ENSMUSG00000005087.17,protein_coding | 56.60345613 | 0.597072167 | 0.000888331 | 0.007475874 |
| ENSMUST00000029624.14,Ccdc109b-001,protein_coding,ENSMUSG00000027994.14,protein_coding | 3303.796416 | 0.194186193 | 0.000888214 | 0.007475874 |
| ENSMUST00000029925.9,Ndufaf4-001,protein_coding,ENSMUSG00000028261.13,protein_coding | 535.050766 | -0.289810718 | 0.000888801 | 0.00747691 |
| ENSMUST00000180863.1,Gm26809-201,lincRNA,ENSMUSG00000097815.1,lincRNA | 64.44817802 | -0.537853328 | 0.000888765 | 0.00747691 |
| ENSMUST00000118800.7,Afap1l2-005,protein_coding,ENSMUSG00000025083.17,protein_coding | 293.7017814 | 0.30920858 | 0.000889088 | 0.007477861 |
| ENSMUST00000069620.9,Per2-001,protein_coding,ENSMUSG00000055866.9,protein_coding | 1158.148712 | 0.171164864 | 0.000893497 | 0.007513472 |
| ENSMUST00000192173.1,Gm37233-001,lincRNA,ENSMUSG00000102336.1,lincRNA | 209.5053343 | -0.427740202 | 0.0008939 | 0.007515392 |
| ENSMUST00000151131.7,Uap1l1-002,processed_transcript,ENSMUSG00000026956.15,protein_coding | 170.0091401 | 0.473001381 | 0.000894306 | 0.007517336 |
| ENSMUST00000081241.1,Gm10039-001,processed_pseudogene,ENSMUSG00000091478.2,processed_pseudogene | 142.0891721 | -0.500065529 | 0.000895382 | 0.007523446 |
| ENSMUST00000110294.1,Elmsan1-001,protein_coding,ENSMUSG00000042507.14,protein_coding | 4297.199539 | 0.291282544 | 0.000895287 | 0.007523446 |
| ENSMUST00000147822.7,Commd7-003,retained_intron,ENSMUSG00000056941.17,protein_coding | 259.8734367 | 0.435494475 | 0.000896347 | 0.007528614 |
| ENSMUST00000176886.1,Tsc22d1-010,nonsense_mediated_decay,ENSMUSG00000022010.19,protein_coding | 192.6827116 | -0.404447401 | 0.000896303 | 0.007528614 |
| ENSMUST00000031434.7,Rnf34-001,protein_coding,ENSMUSG00000029474.7,protein_coding | 2801.010818 | 0.172097883 | 0.000896875 | 0.007531394 |
| ENSMUST00000073456.7,Nsa2-201,protein_coding,ENSMUSG00000060739.7,protein_coding | 16744.28193 | 0.108192486 | 0.000897028 | 0.007531394 |
| ENSMUST00000154208.7,Dhrs3-001,protein_coding,ENSMUSG00000066026.14,protein_coding | 252.2773113 | -0.537443245 | 0.000899606 | 0.007550676 |
| ENSMUST00000035988.14,Txndc5-001,protein_coding,ENSMUSG00000038991.16,protein_coding | 39091.63854 | 0.129895194 | 0.000899851 | 0.007550676 |
| ENSMUST00000162708.6,Unc93b1-001,protein_coding,ENSMUSG00000036908.16,protein_coding | 2057.956888 | 0.232001546 | 0.000899812 | 0.007550676 |
| ENSMUST00000006137.8,Trap1-001,protein_coding,ENSMUSG00000005981.11,protein_coding | 26213.28367 | 0.141838031 | 0.000901089 | 0.007559593 |
| ENSMUST00000134626.7,Hacl1-004,nonsense_mediated_decay,ENSMUSG00000021884.17,protein_coding | 59.62364519 | 0.615830663 | 0.0009014 | 0.00756073 |
| ENSMUST00000097443.9,Gpatch2-002,processed_transcript,ENSMUSG00000039210.16,protein_coding | 151.1629541 | -0.436963176 | 0.000902656 | 0.007569791 |
| ENSMUST00000171266.7,Ndrg1-005,nonsense_mediated_decay,ENSMUSG00000005125.12,protein_coding | 30.02332278 | 0.59967056 | 0.000904101 | 0.007580427 |
| ENSMUST00000094398.12,Uevld-001,protein_coding,ENSMUSG00000043262.16,protein_coding | 4098.386275 | 0.161135592 | 0.000904957 | 0.007586128 |
| ENSMUST00000147096.1,Phpt1-002,processed_transcript,ENSMUSG00000036504.6,protein_coding | 859.6443214 | 0.232192662 | 0.000905732 | 0.007591151 |
| ENSMUST00000027464.8,Mrpl44-001,protein_coding,ENSMUSG00000026248.9,protein_coding | 3249.393359 | 0.17192231 | 0.000906679 | 0.00759761 |
| ENSMUST00000034190.10,Vac14-002,protein_coding,ENSMUSG00000010936.11,protein_coding | 6248.001008 | 0.136191848 | 0.000907885 | 0.007606228 |
| ENSMUST00000129157.1,Ppil1-002,processed_transcript,ENSMUSG00000024007.14,protein_coding | 56.17076824 | -0.555794461 | 0.000908979 | 0.007613914 |
| ENSMUST00000029570.8,Mfsd14a-001,protein_coding,ENSMUSG00000089911.4,protein_coding | 5880.756806 | -0.14943195 | 0.000909829 | 0.007619556 |
| ENSMUST00000026446.2,Cnpy2-201,protein_coding,ENSMUSG00000025381.2,protein_coding | 8181.843016 | 0.177041166 | 0.000911584 | 0.007632767 |
| ENSMUST00000031351.10,Arl6ip4-001,protein_coding,ENSMUSG00000029404.15,protein_coding | 7164.034747 | 0.151373087 | 0.000917847 | 0.007683708 |
| ENSMUST00000212920.1,Cyb5b-002,processed_transcript,ENSMUSG00000031924.4,protein_coding | 3342.397455 | 0.294642733 | 0.000918299 | 0.007686002 |
| ENSMUST00000060175.7,BC030336-001,protein_coding,ENSMUSG00000046096.7,protein_coding | 1369.927457 | -0.306609335 | 0.000919086 | 0.007691095 |
| ENSMUST00000034552.7,Fdx1-001,protein_coding,ENSMUSG00000032051.9,protein_coding | 2286.637429 | 0.166360307 | 0.000920951 | 0.007705203 |
| ENSMUST00000184932.7,Pank2-004,nonsense_mediated_decay,ENSMUSG00000037514.17,protein_coding | 988.7954608 | 0.398951389 | 0.000922516 | 0.007716796 |
| ENSMUST00000109100.1,Fam186b-201,protein_coding,ENSMUSG00000078907.1,protein_coding | 56.85986853 | 0.595366448 | 0.000923123 | 0.00772037 |
| ENSMUST00000174874.7,Cobl-008,protein_coding,ENSMUSG00000020173.17,protein_coding | 654.3015237 | -0.592136788 | 0.000924082 | 0.007726893 |
| ENSMUST00000080178.12,Pabpc4-001,protein_coding,ENSMUSG00000011257.19,protein_coding | 25465.28318 | 0.146765785 | 0.000924493 | 0.007728824 |
| ENSMUST00000123201.7,Vezt-005,protein_coding,ENSMUSG00000036099.16,protein_coding | 143.4169361 | 0.458357708 | 0.000929348 | 0.007766394 |
| ENSMUST00000045540.3,Socs7-001,protein_coding,ENSMUSG00000038485.5,protein_coding | 7551.749738 | 0.14283833 | 0.000929308 | 0.007766394 |

| | | | | |
|---|---|---|---|---|
| ENSMUST00000012104.6,Ccnt1-201,protein_coding,ENSMUSG00000011960.12,protein_coding | 1530.567588 | 0.329684108 | 0.000931785 | 0.007785255 |
| ENSMUST00000161064.7,Atxn2-002,protein_coding,ENSMUSG00000042605.15,protein_coding | 5414.233174 | 0.137761481 | 0.000932179 | 0.00778703 |
| ENSMUST00000069695.8,Tmem150a-001,protein_coding,ENSMUSG00000055912.8,protein_coding | 2263.443017 | -0.156695213 | 0.000932754 | 0.007789176 |
| ENSMUST00000211991.1,Nfatc3-003,protein_coding,ENSMUSG00000031902.10,protein_coding | 1021.63354 | -0.336097723 | 0.000932798 | 0.007789176 |
| ENSMUST00000036355.12,Phldb2-001,protein_coding,ENSMUSG00000033149.16,protein_coding | 30617.30172 | -0.249354432 | 0.000933947 | 0.007797261 |
| ENSMUST00000113099.9,Gapvd1-202,protein_coding,ENSMUSG00000026867.18,protein_coding | 3382.46895 | 0.340549681 | 0.000935099 | 0.007802342 |
| ENSMUST00000060447.12,Mettl5-001,protein_coding,ENSMUSG00000051730.15,protein_coding | 993.5065431 | 0.222810957 | 0.000935006 | 0.007802342 |
| ENSMUST00000025036.10,Kdm4b-001,protein_coding,ENSMUSG00000024201.12,protein_coding | 4966.351069 | 0.196025423 | 0.000935008 | 0.007802342 |
| ENSMUST00000132197.7,Chmp6-002,protein_coding,ENSMUSG00000025371.12,protein_coding | 83.43482797 | 0.521781863 | 0.000939625 | 0.007838582 |
| ENSMUST00000099931.10,Sesn1-001,protein_coding,ENSMUSG00000038332.13,protein_coding | 554.1446722 | -0.292556147 | 0.000941209 | 0.007850277 |
| ENSMUST00000140871.1,Gatad1-003,protein_coding,ENSMUSG00000007415.15,protein_coding | 130.7677209 | -0.475716327 | 0.000941669 | 0.007852595 |
| ENSMUST00000134585.7,Cbx1-003,nonsense_mediated_decay,ENSMUSG00000018666.13,protein_coding | 7395.915569 | -0.156074915 | 0.000942075 | 0.007854456 |
| ENSMUST00000020461.14,Nfic-002,protein_coding,ENSMUSG00000055053.17,protein_coding | 9839.377928 | -0.118387403 | 0.000942746 | 0.007858533 |
| ENSMUST00000109218.6,Slc9a8-004,protein_coding,ENSMUSG00000039463.14,protein_coding | 358.7393791 | 0.374368701 | 0.000946337 | 0.007886937 |
| ENSMUST00000120428.7,Trim24-002,protein_coding,ENSMUSG00000029833.17,protein_coding | 4633.830859 | 0.157535428 | 0.000947566 | 0.007895649 |
| ENSMUST00000134059.1,Gm13073-001,lincRNA,ENSMUSG00000084807.1,lincRNA | 55.63109827 | 0.607530646 | 0.000953082 | 0.007940077 |
| ENSMUST00000119274.1,Stk38-002,protein_coding,ENSMUSG00000024006.16,protein_coding | 615.21299 | 0.380641677 | 0.000954741 | 0.007952356 |
| ENSMUST00000035121.13,Amotl2-001,protein_coding,ENSMUSG00000032531.15,protein_coding | 33473.56928 | -0.131652722 | 0.000956583 | 0.00796616 |
| ENSMUST00000081387.10,Birc5-001,protein_coding,ENSMUSG00000017716.15,protein_coding | 12974.68116 | -0.167607123 | 0.000956952 | 0.007967687 |
| ENSMUST00000005817.8,Tomm40l-001,protein_coding,ENSMUSG00000005674.9,protein_coding | 66.18659321 | 0.561551952 | 0.000958473 | 0.007978814 |
| ENSMUST00000137767.1,Gosr1-002,processed_transcript,ENSMUSG00000010392.8,protein_coding | 308.0267341 | 0.293792898 | 0.00096134 | 0.008001132 |
| ENSMUST00000042953.8,Nbas-201,protein_coding,ENSMUSG00000020576.9,protein_coding | 11508.45629 | 0.161000755 | 0.000962466 | 0.008008953 |
| ENSMUST00000119816.8,Pgap2-024,nonsense_mediated_decay,ENSMUSG00000030990.18,protein_coding | 114.7809515 | 0.587874755 | 0.000963277 | 0.008013942 |
| ENSMUST00000002889.4,Flii-001,protein_coding,ENSMUSG00000002812.4,protein_coding | 21260.64826 | 0.12321642 | 0.000963438 | 0.008013942 |
| ENSMUST00000202241.3,Mpv17-002,protein_coding,ENSMUSG00000107283.3,protein_coding | 282.7483658 | 0.424113924 | 0.000963967 | 0.008016794 |
| ENSMUST00000166033.5,Ube2d3-003,protein_coding,ENSMUSG00000078578.9,protein_coding | 4601.576588 | -0.289031179 | 0.000964478 | 0.00801949 |
| ENSMUST00000168247.1,Zfp110-201,protein_coding,ENSMUSG00000058638.14,protein_coding | 1440.854448 | 0.314306959 | 0.000965743 | 0.008028462 |
| ENSMUST00000184635.7,Hnrnpll-001,protein_coding,ENSMUSG00000024095.15,protein_coding | 9812.887102 | -0.156620669 | 0.000966061 | 0.008029558 |
| ENSMUST00000072899.8,Cgnl1-201,protein_coding,ENSMUSG00000032232.14,protein_coding | 6282.719802 | 0.124749494 | 0.000966301 | 0.008029998 |
| ENSMUST00000118544.7,Armt1-002,protein_coding,ENSMUSG00000061759.15,protein_coding | 468.2942108 | -0.298000406 | 0.000967079 | 0.00803491 |
| ENSMUST00000105501.1,Foxo3-003,protein_coding,ENSMUSG00000048756.11,protein_coding | 70.8330477 | 0.56778512 | 0.000969559 | 0.008053965 |
| ENSMUST00000087951.5,Vps37c-201,protein_coding,ENSMUSG00000048832.8,protein_coding | 3436.304662 | 0.157752012 | 0.000973076 | 0.00808162 |
| ENSMUST00000142675.7,Pcolce-006,protein_coding,ENSMUSG00000029718.14,protein_coding | 1030.59133 | -0.269130124 | 0.000974674 | 0.008093329 |
| ENSMUST00000075896.6,Tsnax-001,protein_coding,ENSMUSG00000056820.7,protein_coding | 12688.03243 | 0.14892756 | 0.000975101 | 0.008095316 |
| ENSMUST00000169082.1,Aqp5-202,protein_coding,ENSMUSG00000044217.16,protein_coding | 778.7165293 | -0.328331275 | 0.000975758 | 0.008099208 |
| ENSMUST00000134980.1,Rnd2-003,processed_transcript,ENSMUSG00000001313.12,protein_coding | 45.16970683 | 0.608012071 | 0.000978243 | 0.008118266 |
| ENSMUST00000045286.8,Tmem59l-001,protein_coding,ENSMUSG00000035964.8,protein_coding | 239.072233 | -0.375752186 | 0.000981649 | 0.008143392 |
| ENSMUST00000038926.12,Baz1a-201,protein_coding,ENSMUSG00000035021.13,protein_coding | 318.2927713 | -0.561822307 | 0.000981525 | 0.008143392 |
| ENSMUST00000149441.7,Mpnd-001,protein_coding,ENSMUSG00000003199.16,protein_coding | 6227.126371 | 0.173184479 | 0.000983772 | 0.008159431 |
| ENSMUST00000138147.1,Ccdc43-004,processed_transcript,ENSMUSG00000020925.16,protein_coding | 75.83757792 | -0.499887704 | 0.000983994 | 0.008159704 |
| ENSMUST00000025204.5,Pfdn1-201,protein_coding,ENSMUSG00000024346.5,protein_coding | 8809.566538 | 0.121789572 | 0.000985419 | 0.008169943 |
| ENSMUST00000002790.13,Cse1l-001,protein_coding,ENSMUSG00000002718.14,protein_coding | 27079.08729 | -0.162572442 | 0.000985934 | 0.008172638 |
| ENSMUST00000020896.14,Tspan13-001,protein_coding,ENSMUSG00000020577.15,protein_coding | 806.0578244 | 0.258749831 | 0.000987253 | 0.008181996 |
| ENSMUST00000093450.5,Ano8-001,protein_coding,ENSMUSG00000034863.9,protein_coding | 3157.971652 | -0.161080234 | 0.000987898 | 0.008185767 |
| ENSMUST00000036019.4,Fras1-001,protein_coding,ENSMUSG00000034687.8,protein_coding | 608.838823 | -0.284007726 | 0.000990703 | 0.008205854 |
| ENSMUST00000004375.15,Phb2-001,protein_coding,ENSMUSG00000004264.17,protein_coding | 50023.10131 | 0.142841978 | 0.000990568 | 0.008205854 |
| ENSMUST00000005064.13,Pias4-001,protein_coding,ENSMUSG00000004934.13,protein_coding | 5874.220619 | 0.160899304 | 0.000993077 | 0.008223934 |
| ENSMUST00000142376.1,Osbpl2-002,retained_intron,ENSMUSG00000039050.8,protein_coding | 300.2650331 | 0.474124572 | 0.000994107 | 0.008230873 |
| ENSMUST00000044007.2,Oxld1-001,protein_coding,ENSMUSG00000039670.2,protein_coding | 617.8440056 | 0.276018756 | 0.000994598 | 0.008233362 |
| ENSMUST00000197188.4,Ankle2-002,protein_coding,ENSMUSG00000029501.14,protein_coding | 5810.357989 | 0.172243596 | 0.000995352 | 0.008238016 |
| ENSMUST00000100203.9,Dip2b-001,protein_coding,ENSMUSG00000023026.15,protein_coding | 6080.711964 | 0.129106213 | 0.000996168 | 0.008243188 |
| ENSMUST00000129466.7,Tpm1-019,retained_intron,ENSMUSG00000032366.15,protein_coding | 135.2066192 | -0.46433786 | 0.000996484 | 0.008244213 |
| ENSMUST00000030356.9,Lrp8-006,protein_coding,ENSMUSG00000028613.14,protein_coding | 2784.831467 | 0.246271141 | 0.000997002 | 0.008246914 |
| ENSMUST00000123952.8,Cfap74-003,protein_coding,ENSMUSG00000078490.9,protein_coding | 104.6256917 | -0.494075864 | 0.000998529 | 0.008257647 |
| ENSMUST00000044189.15,Senp1-001,protein_coding,ENSMUSG00000033075.17,protein_coding | 7425.510249 | -0.10681566 | 0.000998683 | 0.008257647 |
| ENSMUST00000133168.7,Spice1-003,retained_intron,ENSMUSG00000043065.12,protein_coding | 677.7729025 | 0.250845229 | 0.000999158 | 0.008258403 |
| ENSMUST00000164253.1,Csnk1a1-010,protein_coding,ENSMUSG00000024576.14,protein_coding | 54.53498088 | 0.583957966 | 0.000999014 | 0.008258403 |
| ENSMUST00000102546.3,Il22ra1-001,protein_coding,ENSMUSG00000037157.8,protein_coding | 964.0699271 | -0.285981209 | 0.000999621 | 0.008260645 |
| ENSMUST00000024963.9,Mcfd2-001,protein_coding,ENSMUSG00000024150.11,protein_coding | 7882.887551 | 0.202250636 | 0.001000865 | 0.008269339 |
| ENSMUST00000049040.13,Bri3bp-001,protein_coding,ENSMUSG00000037905.13,protein_coding | 12463.57833 | 0.122337062 | 0.001001442 | 0.008272514 |
| ENSMUST00000175673.1,Senp6-003,retained_intron,ENSMUSG00000034252.14,protein_coding | 60.43818612 | -0.604270778 | 0.001005058 | 0.0082992 |
| ENSMUST00000176207.7,Hnrnpk-008,protein_coding,ENSMUSG00000021546.17,protein_coding | 4491.02873 | 0.240090728 | 0.001004922 | 0.0082992 |
| ENSMUST00000081560.4,Sdhc-002,protein_coding,ENSMUSG00000058076.12,protein_coding | 6977.042081 | 0.161745891 | 0.001005265 | 0.008299317 |
| ENSMUST00000048209.15,Ldha-001,protein_coding,ENSMUSG00000063229.15,protein_coding | 258505.7923 | 0.167155191 | 0.001006589 | 0.008308653 |
| ENSMUST00000115352.9,Tceb1-001,protein_coding,ENSMUSG00000079658.9,protein_coding | 1573.503944 | -0.443071802 | 0.001007792 | 0.008316989 |
| ENSMUST00000176923.7,Wdr90-006,protein_coding,ENSMUSG00000073434.11,protein_coding | 784.469803 | -0.394500461 | 0.001008824 | 0.008323911 |
| ENSMUST00000101198.8,Tmem164-005,protein_coding,ENSMUSG00000047045.17,protein_coding | 1846.307961 | 0.522556592 | 0.001009164 | 0.008325123 |
| ENSMUST00000185117.7,Tcf12-018,protein_coding,ENSMUSG00000032228.16,protein_coding | 2916.880431 | 0.285690215 | 0.001011491 | 0.008342718 |
| ENSMUST00000109682.8,Dynlrb1-001,protein_coding,ENSMUSG00000047459.14,protein_coding | 8258.619159 | 0.121228142 | 0.001012535 | 0.008346527 |
| ENSMUST00000047208.11,Dzip1-202,protein_coding,ENSMUSG00000042156.15,protein_coding | 293.8205178 | -0.530780594 | 0.0010123 | 0.008346527 |
| ENSMUST00000025649.8,Ddb1-201,protein_coding,ENSMUSG00000024740.9,protein_coding | 230393.2201 | 0.14908127 | 0.001012369 | 0.008346527 |
| ENSMUST00000129459.1,Htra3-004,protein_coding,ENSMUSG00000029096.15,protein_coding | 387.13597 | 0.286003191 | 0.001014414 | 0.00836042 |
| ENSMUST00000045907.14,2700049A03Rik-201,protein_coding,ENSMUSG00000034601.16,protein_coding | 1016.924585 | -0.492209653 | 0.001015919 | 0.00837122 |
| ENSMUST00000111774.1,Xpr1-002,protein_coding,ENSMUSG00000026469.14,protein_coding | 198.4874838 | 0.579779165 | 0.001016947 | 0.008378083 |
| ENSMUST00000168892.7,Zfp595-001,protein_coding,ENSMUSG00000057842.13,protein_coding | 131.5991944 | -0.437581256 | 0.001018957 | 0.008393038 |
| ENSMUST00000034030.14,Tmem184c-001,protein_coding,ENSMUSG00000031617.16,protein_coding | 2635.76958 | -0.357679907 | 0.001019508 | 0.008395973 |
| ENSMUST00000194127.5,Ptpn14-004,retained_intron,ENSMUSG00000026604.17,protein_coding | 123.1712805 | -0.606229852 | 0.001020135 | 0.008399525 |
| ENSMUST00000027438.7,Ncl-001,protein_coding,ENSMUSG00000026234.12,protein_coding | 33384.07014 | -0.290755406 | 0.001020482 | 0.008400775 |
| ENSMUST00000111535.7,Amn1-002,protein_coding,ENSMUSG00000068250.13,protein_coding | 314.8011272 | 0.415730997 | 0.001020689 | 0.008400874 |
| ENSMUST00000028856.2,Mall-001,protein_coding,ENSMUSG00000027377.2,protein_coding | 33.53938391 | 0.611895049 | 0.001021773 | 0.008406581 |
| ENSMUST00000203609.2,Zfp746-006,protein_coding,ENSMUSG00000057691.8,protein_coding | 3482.316189 | 0.226567894 | 0.001021768 | 0.008406581 |
| ENSMUST00000151492.1,Foxk2-003,retained_intron,ENSMUSG00000039275.9,protein_coding | 163.8421515 | 0.596347188 | 0.001022446 | 0.008410514 |
| ENSMUST00000053744.7,2410016O06Rik-201,protein_coding,ENSMUSG00000046791.7,protein_coding | 3496.812542 | 0.124223684 | 0.001024053 | 0.008422124 |
| ENSMUST00000024620.6,Chd1-201,protein_coding,ENSMUSG00000023852.13,protein_coding | 5133.480531 | -0.262866076 | 0.001026248 | 0.008438559 |
| ENSMUST00000117223.3,Dnajc19-003,protein_coding,ENSMUSG00000027679.13,protein_coding | 57.54747667 | -0.559514213 | 0.001027172 | 0.008444548 |
| ENSMUST00000186613.1,Fn1-014,nonsense_mediated_decay,ENSMUSG00000026193.15,protein_coding | 252.2782765 | 0.350546113 | 0.001028548 | 0.008453657 |
| ENSMUST00000142735.7,Rpl22-003,processed_transcript,ENSMUSG00000028936.15,protein_coding | 736.8753101 | 0.292685119 | 0.001028763 | 0.008453657 |
| ENSMUST00000032198.10,Usp18-001,protein_coding,ENSMUSG00000030107.10,protein_coding | 150.6552805 | -0.459877107 | 0.001029066 | 0.008453657 |
| ENSMUST00000163761.7,Fam184a-006,protein_coding,ENSMUSG00000019856.13,protein_coding | 88.35460491 | -0.603807174 | 0.001029005 | 0.008453657 |
| ENSMUST00000169582.2,Usf3-201,protein_coding,ENSMUSG00000068284.14,protein_coding | 620.4641661 | 0.611493855 | 0.001029323 | 0.008454159 |
| ENSMUST00000142191.1,Tiprl-003,processed_transcript,ENSMUSG00000040843.10,protein_coding | 1018.393871 | 0.213843228 | 0.001029929 | 0.008457522 |
| ENSMUST00000124810.7,Ppig-004,processed_transcript,ENSMUSG00000042133.16,protein_coding | 96.96244438 | -0.5679598 | 0.001030785 | 0.008461321 |

| | | | | |
|---|---|---|---|---|
| ENSMUST00000144153.7,Usp36-003,protein_coding,ENSMUSG00000033909.17,protein_coding | 1984.029826 | 0.280199054 | 0.001030722 | 0.008461321 |
| ENSMUST00000022460.10,Galnt15-001,protein_coding,ENSMUSG00000021903.11,protein_coding | 85.95793083 | -0.498227729 | 0.001031047 | 0.00846186 |
| ENSMUST00000057258.10,Zfp518b-001,protein_coding,ENSMUSG00000046572.10,protein_coding | 3357.293277 | 0.292834352 | 0.001033658 | 0.008476823 |
| ENSMUST00000204136.1,9430076C15Rik-003,processed_transcript,ENSMUSG00000053007.9,protein_coding | 37.39430368 | 0.605104894 | 0.001033577 | 0.008476823 |
| ENSMUST00000092892.9,Ankrd13b-201,protein_coding,ENSMUSG00000037907.16,protein_coding | 825.5280868 | 0.235880648 | 0.001033562 | 0.008476823 |
| ENSMUST00000037391.11,Cd99l2-001,protein_coding,ENSMUSG00000035776.14,protein_coding | 4371.341924 | 0.186498072 | 0.001033219 | 0.008476823 |
| ENSMUST00000209556.1,Rpl18-006,protein_coding,ENSMUSG00000059070.16,protein_coding | 630.979456 | 0.302323802 | 0.001035579 | 0.008490957 |
| ENSMUST00000169383.7,Gtpbp2-002,nonsense_mediated_decay,ENSMUSG00000023952.13,protein_coding | 373.4129952 | 0.493113738 | 0.001039667 | 0.008522853 |
| ENSMUST00000196712.1,Stard3nl-002,retained_intron,ENSMUSG00000003062.12,protein_coding | 352.3008053 | 0.351168318 | 0.001041297 | 0.008534589 |
| ENSMUST00000161952.1,Rnf8-002,processed_transcript,ENSMUSG00000090083.10,protein_coding | 33.73575599 | -0.600861433 | 0.001041653 | 0.00853588 |
| ENSMUST00000022908.8,Slc25a32-201,protein_coding,ENSMUSG00000022299.8,protein_coding | 3247.45876 | -0.199685663 | 0.001043117 | 0.008544665 |
| ENSMUST00000033754.14,Piga-001,protein_coding,ENSMUSG00000031381.16,protein_coding | 832.678827 | -0.287299259 | 0.001043122 | 0.008544665 |
| ENSMUST00000167759.7,Atxn2l-002,protein_coding,ENSMUSG00000032637.15,protein_coding | 17541.54789 | 0.221153765 | 0.001044102 | 0.008551069 |
| ENSMUST00000052691.8,1600012H06Rik-003,protein_coding,ENSMUSG00000050088.9,protein_coding | 515.4418848 | -0.39806552 | 0.001044828 | 0.008555385 |
| ENSMUST00000150892.1,Clip3-003,protein_coding,ENSMUSG00000013921.15,protein_coding | 430.8012392 | -0.252544566 | 0.00104636 | 0.008566301 |
| ENSMUST00000037620.13,Mospd3-001,protein_coding,ENSMUSG00000037221.13,protein_coding | 154.7305529 | -0.50472923 | 0.001051605 | 0.008607607 |
| ENSMUST00000123693.1,Ptgr2-003,nonsense_mediated_decay,ENSMUSG00000072946.12,protein_coding | 323.2830053 | 0.518993129 | 0.001058199 | 0.008659932 |
| ENSMUST00000113756.7,Odf2-002,protein_coding,ENSMUSG00000026790.19,protein_coding | 8135.105644 | 0.255934704 | 0.001058607 | 0.008661621 |
| ENSMUST00000093166.10,Cyfip2-001,protein_coding,ENSMUSG00000020340.16,protein_coding | 70.79964566 | -0.58008906 | 0.001058968 | 0.00866293 |
| ENSMUST00000077890.11,Iqce-002,protein_coding,ENSMUSG00000036555.14,protein_coding | 1213.570382 | -0.200145452 | 0.001060513 | 0.008673921 |
| ENSMUST00000109113.7,Clk4-003,protein_coding,ENSMUSG00000020385.16,protein_coding | 175.2473957 | -0.517309228 | 0.001061567 | 0.008680891 |
| ENSMUST00000212352.1,Pard6a-006,protein_coding,ENSMUSG00000005699.16,protein_coding | 616.8828475 | 0.48919397 | 0.001061952 | 0.008682391 |
| ENSMUST00000155392.2,Prr14l-004,nonsense_mediated_decay,ENSMUSG00000054280.14,protein_coding | 81.7003626 | -0.570301915 | 0.001064355 | 0.008700382 |
| ENSMUST00000107866.7,Tmem8b-001,protein_coding,ENSMUSG00000078716.9,protein_coding | 837.2890401 | 0.392961801 | 0.001069822 | 0.008743416 |
| ENSMUST00000132459.7,Prrc2b-002,protein_coding,ENSMUSG00000039262.16,protein_coding | 5334.368839 | 0.175385795 | 0.001071232 | 0.008753276 |
| ENSMUST00000078090.11,Mllt3-001,protein_coding,ENSMUSG00000028496.17,protein_coding | 592.7283755 | -0.453949894 | 0.001071909 | 0.008757097 |
| ENSMUST00000165418.7,Zfp148-202,protein_coding,ENSMUSG00000022811.16,protein_coding | 4633.38144 | -0.406950547 | 0.001072107 | 0.008757097 |
| ENSMUST00000034524.4,Rexo2-001,protein_coding,ENSMUSG00000032026.7,protein_coding | 5706.311026 | 0.191868538 | 0.001075595 | 0.00878392 |
| ENSMUST00000050067.9,Hectd3-001,protein_coding,ENSMUSG00000046861.9,protein_coding | 2479.6965 | 0.189351132 | 0.00107888 | 0.008809079 |
| ENSMUST00000056578.6,Bri3-001,protein_coding,ENSMUSG00000047843.16,protein_coding | 15645.24152 | 0.157234915 | 0.001081047 | 0.008825096 |
| ENSMUST00000110457.7,Myo10-001,protein_coding,ENSMUSG00000022272.17,protein_coding | 43375.14651 | -0.139693519 | 0.001082475 | 0.008835078 |
| ENSMUST00000080138.12,Acbd6-003,protein_coding,ENSMUSG00000033701.13,protein_coding | 5572.420281 | 0.15723297 | 0.001083331 | 0.00883537 |
| ENSMUST00000086831.3,Pkd2-001,protein_coding,ENSMUSG00000034462.9,protein_coding | 6228.803205 | -0.233447996 | 0.001083241 | 0.00883537 |
| ENSMUST00000107128.7,Myo7a-001,protein_coding,ENSMUSG00000030761.15,protein_coding | 1481.274065 | 0.189599286 | 0.001083112 | 0.00883537 |
| ENSMUST00000014339.14,Dnajc7-001,protein_coding,ENSMUSG00000014195.15,protein_coding | 14039.67531 | 0.142782458 | 0.001083164 | 0.00883537 |
| ENSMUST00000108583.8,Kmt5c-002,protein_coding,ENSMUSG00000059851.15,protein_coding | 796.6168145 | 0.316999266 | 0.00108356 | 0.008835558 |
| ENSMUST00000024747.13,Vegfa-008,protein_coding,ENSMUSG00000023951.17,protein_coding | 4955.846767 | 0.250089453 | 0.001084138 | 0.008838595 |
| ENSMUST00000060895.5,Ugp2-002,protein_coding,ENSMUSG00000001891.16,protein_coding | 2957.780338 | -0.255737801 | 0.001086754 | 0.008858248 |
| ENSMUST00000050073.12,Dnajb4-002,protein_coding,ENSMUSG00000028035.13,protein_coding | 2987.865123 | -0.285365668 | 0.001088628 | 0.008871846 |
| ENSMUST00000125107.7,0610037L13Rik-003,protein_coding,ENSMUSG00000028608.15,protein_coding | 458.5595348 | -0.236543359 | 0.001089876 | 0.008878653 |
| ENSMUST00000170141.2,Orc6-002,protein_coding,ENSMUSG00000031697.12,protein_coding | 567.6367662 | -0.431172158 | 0.001089713 | 0.008878653 |
| ENSMUST00000067205.14,Kif21a-201,protein_coding,ENSMUSG00000022629.16,protein_coding | 70.57670027 | 0.602412422 | 0.001090777 | 0.008882632 |
| ENSMUST00000172334.7,Smagp-202,protein_coding,ENSMUSG00000053559.12,protein_coding | 288.4062024 | 0.438799669 | 0.001090732 | 0.008882632 |
| ENSMUST00000205568.1,Maz-003,protein_coding,ENSMUSG00000030678.7,protein_coding | 6565.221865 | 0.203056322 | 0.001093775 | 0.008905363 |
| ENSMUST00000060490.10,Homer1-006,protein_coding,ENSMUSG00000007617.17,protein_coding | 388.251309 | -0.391265066 | 0.001094625 | 0.008910598 |
| ENSMUST00000091853.10,Net1-201,protein_coding,ENSMUSG00000021215.14,protein_coding | 4578.724839 | -0.103126747 | 0.001096255 | 0.008922174 |
| ENSMUST00000026666.12,Gaa-002,protein_coding,ENSMUSG00000025579.14,protein_coding | 9500.901327 | 0.178984855 | 0.001096932 | 0.008925856 |
| ENSMUST00000181149.1,B230354K17Rik-201,lincRNA,ENSMUSG00000097119.1,lincRNA | 1648.088505 | -0.363768498 | 0.001097122 | 0.008925856 |
| ENSMUST00000030140.2,Ikbkap-001,protein_coding,ENSMUSG00000028431.11,protein_coding | 12941.84444 | -0.133985319 | 0.001097587 | 0.008927956 |
| ENSMUST00000148371.7,Otub2-002,protein_coding,ENSMUSG00000021203.15,protein_coding | 42.61197498 | 0.579997552 | 0.001098391 | 0.008932808 |
| ENSMUST00000199887.1,Sfswap-003,retained_intron,ENSMUSG00000029439.14,protein_coding | 690.5734164 | -0.265258937 | 0.00109936 | 0.008938998 |
| ENSMUST00000164957.7,Prr13-004,protein_coding,ENSMUSG00000023048.13,protein_coding | 8283.132635 | 0.140456103 | 0.001100249 | 0.008944533 |
| ENSMUST00000135625.7,Zfp106-003,protein_coding,ENSMUSG00000027288.16,protein_coding | 1240.857853 | 0.230923022 | 0.001100854 | 0.008947761 |
| ENSMUST00000176423.1,Cux1-017,protein_coding,ENSMUSG00000029705.17,protein_coding | 712.1976911 | 0.194308982 | 0.001101913 | 0.008954684 |
| ENSMUST00000034946.14,Snx1-001,protein_coding,ENSMUSG00000032382.14,protein_coding | 7841.910339 | 0.167281964 | 0.001103871 | 0.008968899 |
| ENSMUST00000067103.2,Taf8-201,protein_coding,ENSMUSG00000023980.6,protein_coding | 2988.42092 | 0.132206795 | 0.001105172 | 0.008977772 |
| ENSMUST00000185332.1,Atf7ip-004,retained_intron,ENSMUSG00000030213.12,protein_coding | 285.9409716 | 0.39628583 | 0.001106918 | 0.008990263 |
| ENSMUST00000028403.2,Cybrd1-001,protein_coding,ENSMUSG00000027015.4,protein_coding | 99.04404716 | -0.501960424 | 0.001107537 | 0.008991896 |
| ENSMUST00000164504.1,Cfap97-201,protein_coding,ENSMUSG00000031631.15,protein_coding | 609.6294632 | -0.535094532 | 0.001107444 | 0.008991896 |
| ENSMUST00000028083.5,Psmb7-001,protein_coding,ENSMUSG00000026750.6,protein_coding | 29553.06967 | 0.14246261 | 0.001108998 | 0.009002064 |
| ENSMUST00000086399.5,Icam1-001,protein_coding,ENSMUSG00000037405.8,protein_coding | 328.7031506 | -0.370795626 | 0.001110834 | 0.009015266 |
| ENSMUST00000085461.3,Klk8-001,protein_coding,ENSMUSG00000064023.4,protein_coding | 191.5382851 | 0.384335593 | 0.001112336 | 0.009025753 |
| ENSMUST00000030142.3,Epb41l4b-001,protein_coding,ENSMUSG00000028434.12,protein_coding | 1779.927551 | 0.268852052 | 0.001115821 | 0.009050967 |
| ENSMUST00000179936.7,Epn2-202,protein_coding,ENSMUSG00000001036.17,protein_coding | 84.42797851 | 0.488830654 | 0.001115864 | 0.009050967 |
| ENSMUST00000216076.1,Rps25-004,protein_coding,ENSMUSG00000009927.9,protein_coding | 9545.251029 | 0.174195436 | 0.001116755 | 0.00905649 |
| ENSMUST00000167741.7,Cox7a2l-202,protein_coding,ENSMUSG00000024248.13,protein_coding | 57.69455029 | 0.592455358 | 0.001117385 | 0.009059888 |
| ENSMUST00000198931.1,Cdkl5-003,protein_coding,ENSMUSG00000031292.14,protein_coding | 609.1057482 | -0.435066354 | 0.001118202 | 0.009064805 |
| ENSMUST00000143362.1,Ttc39b-004,retained_intron,ENSMUSG00000038172.14,protein_coding | 364.4745067 | 0.438397397 | 0.001119095 | 0.009070027 |
| ENSMUST00000023143.13,Litaf-001,protein_coding,ENSMUSG00000022500.14,protein_coding | 7508.324632 | 0.141182031 | 0.001119267 | 0.009070027 |
| ENSMUST00000194508.1,Gm36936-001,TEC,ENSMUSG00000102411.1,TEC | 214.5860342 | -0.381052413 | 0.001119886 | 0.009073334 |
| ENSMUST00000186182.1,Psmd8-201,protein_coding,ENSMUSG00000030591.17,protein_coding | 67.05150301 | 0.606297887 | 0.001122942 | 0.009096382 |
| ENSMUST00000056391.14,Men1-003,protein_coding,ENSMUSG00000024947.16,protein_coding | 4152.286165 | 0.16798568 | 0.001123207 | 0.009096814 |
| ENSMUST00000119603.1,Stambpl1-003,protein_coding,ENSMUSG00000024776.17,protein_coding | 532.3685545 | 0.522766207 | 0.001124437 | 0.009105065 |
| ENSMUST00000203687.1,Itpr1-006,protein_coding,ENSMUSG00000030102.11,protein_coding | 27.89695352 | 0.607820315 | 0.001125805 | 0.009112714 |
| ENSMUST00000111878.7,Eps8-002,protein_coding,ENSMUSG00000015766.14,protein_coding | 8193.302677 | 0.192569815 | 0.001125788 | 0.009112714 |
| ENSMUST00000100219.4,Dolk-001,protein_coding,ENSMUSG00000075419.4,protein_coding | 2203.634158 | -0.186805019 | 0.001126988 | 0.009120362 |
| ENSMUST00000038890.5,Dennd4a-001,protein_coding,ENSMUSG00000053641.9,protein_coding | 5445.790972 | 0.376169488 | 0.001127174 | 0.009120362 |
| ENSMUST00000034534.12,Ets1-001,protein_coding,ENSMUSG00000032035.15,protein_coding | 8120.587534 | -0.147477401 | 0.001128501 | 0.009129382 |
| ENSMUST00000027108.12,Eef1b2-006,nonsense_mediated_decay,ENSMUSG00000025967.16,protein_coding | 385.1603177 | 0.446597587 | 0.001129816 | 0.009138303 |
| ENSMUST00000034226.7,Fam192a-001,protein_coding,ENSMUSG00000031774.8,protein_coding | 3771.405575 | 0.134895926 | 0.001131577 | 0.009150826 |
| ENSMUST00000160786.7,Mff-002,protein_coding,ENSMUSG00000026150.14,protein_coding | 927.0586703 | 0.353613503 | 0.00113239 | 0.009155685 |
| ENSMUST00000031565.14,Fscn1-001,protein_coding,ENSMUSG00000029581.14,protein_coding | 59959.8165 | -0.136109245 | 0.001133445 | 0.009162489 |
| ENSMUST00000040766.7,Tmed10-201,protein_coding,ENSMUSG00000021248.8,protein_coding | 19517.28406 | 0.114057038 | 0.001133852 | 0.009164062 |
| ENSMUST00000116415.8,Mthfsd-001,protein_coding,ENSMUSG00000031816.15,protein_coding | 329.5022552 | -0.410433122 | 0.001134955 | 0.009171256 |
| ENSMUST00000197052.1,Orai2-005,protein_coding,ENSMUSG00000039747.11,protein_coding | 239.8585444 | -0.489719783 | 0.001135828 | 0.009176588 |
| ENSMUST00000210901.1,Rtp4-007,protein_coding,ENSMUSG00000033355.6,protein_coding | 177.0382015 | 0.410509989 | 0.001136788 | 0.009182615 |
| ENSMUST00000097816.8,Crocc-001,protein_coding,ENSMUSG00000040860.16,protein_coding | 1450.212528 | 0.316480454 | 0.001137393 | 0.009185783 |
| ENSMUST00000030399.6,Rragc-001,protein_coding,ENSMUSG00000028646.16,protein_coding | 10206.9922 | 0.148791542 | 0.00113829 | 0.009191306 |
| ENSMUST00000043813.1,Nudt15-201,protein_coding,ENSMUSG00000033405.3,protein_coding | 183.5518769 | -0.413353479 | 0.001138828 | 0.009193922 |
| ENSMUST00000182580.7,Snhg6-001,lincRNA,ENSMUSG00000098234.7,lincRNA | 2588.999658 | 0.181899672 | 0.001139351 | 0.009196418 |
| ENSMUST00000059580.10,Kmt5a-001,protein_coding,ENSMUSG00000049327.17,protein_coding | 17287.08711 | 0.146887763 | 0.001140588 | 0.009204681 |
| ENSMUST00000034000.14,Asah1-001,protein_coding,ENSMUSG00000031591.14,protein_coding | 8536.876712 | 0.176808099 | 0.001140839 | 0.009204978 |

| | | | | |
|---|---|---|---|---|
| ENSMUST00000078323.12,Tmem175-003,protein_coding,ENSMUSG00000013495.15,protein_coding | 953.7659428 | 0.178877163 | 0.001144063 | 0.009229261 |
| ENSMUST00000040336.11,Slc22a23-001,protein_coding,ENSMUSG00000038267.14,protein_coding | 3446.67169 | -0.126899467 | 0.001145388 | 0.009238217 |
| ENSMUST00000115632.9,Mre11a-002,protein_coding,ENSMUSG00000031928.15,protein_coding | 1769.958828 | -0.20986244 | 0.001146504 | 0.009245488 |
| ENSMUST00000029674.7,Efna4-001,protein_coding,ENSMUSG00000028040.7,protein_coding | 615.5087356 | -0.308757834 | 0.001148412 | 0.009259135 |
| ENSMUST00000140679.7,Uba52-004,protein_coding,ENSMUSG00000090137.8,protein_coding | 1560.804297 | 0.229543304 | 0.001149093 | 0.009262891 |
| ENSMUST00000151559.1,Uqcr10-002,processed_transcript,ENSMUSG00000059534.8,protein_coding | 1611.522343 | 0.187751724 | 0.001149849 | 0.009267257 |
| ENSMUST00000154938.1,Kdm5c-003,retained_intron,ENSMUSG00000025332.14,protein_coding | 53.53839157 | -0.595247558 | 0.001151182 | 0.009276263 |
| ENSMUST00000188797.6,Epha4-001,protein_coding,ENSMUSG00000026235.14,protein_coding | 528.5203083 | 0.470724838 | 0.001152757 | 0.009287214 |
| ENSMUST00000031254.8,Klhl8-001,protein_coding,ENSMUSG00000029312.12,protein_coding | 619.4743067 | 0.264959004 | 0.001153127 | 0.009288457 |
| ENSMUST00000043237.13,Trp53inp2-001,protein_coding,ENSMUSG00000038375.15,protein_coding | 2402.962903 | -0.20007408 | 0.001158223 | 0.009327758 |
| ENSMUST00000199878.4,Papss1-002,protein_coding,ENSMUSG00000028032.13,protein_coding | 3707.563119 | 0.173122524 | 0.001162491 | 0.009360382 |
| ENSMUST00000109353.8,Tubgcp6-001,protein_coding,ENSMUSG00000051786.14,protein_coding | 5211.675427 | -0.178035732 | 0.001163663 | 0.00936806 |
| ENSMUST00000128653.7,Pcsk4-002,nonsense_mediated_decay,ENSMUSG00000020131.14,protein_coding | 49.35323675 | 0.604438437 | 0.001165004 | 0.009377105 |
| ENSMUST00000058183.8,Ndufaf6-001,protein_coding,ENSMUSG00000050323.14,protein_coding | 1115.884938 | 0.212930102 | 0.001166322 | 0.009385961 |
| ENSMUST00000068725.9,Marc2-001,protein_coding,ENSMUSG00000073481.9,protein_coding | 2813.551893 | 0.177178864 | 0.001169122 | 0.009403943 |
| ENSMUST00000098552.9,Tmod2-001,protein_coding,ENSMUSG00000032186.15,protein_coding | 457.6947658 | -0.382128295 | 0.001169431 | 0.009403943 |
| ENSMUST00000092673.10,Sgk1-003,protein_coding,ENSMUSG00000019970.15,protein_coding | 902.8212518 | -0.286286253 | 0.001169192 | 0.009403943 |
| ENSMUST00000174164.1,Ddx39b-005,retained_intron,ENSMUSG00000019432.15,protein_coding | 5321.386067 | 0.162757035 | 0.001169321 | 0.009403943 |
| ENSMUST00000133748.1,Cers2-007,retained_intron,ENSMUSG00000015714.11,protein_coding | 168.5634545 | 0.351431638 | 0.001172318 | 0.009425398 |
| ENSMUST00000215420.1,Slc37a4-003,protein_coding,ENSMUSG00000032114.9,protein_coding | 514.8043592 | 0.325970955 | 0.001172791 | 0.009426267 |
| ENSMUST00000177024.7,Traf7-009,retained_intron,ENSMUSG00000052752.15,protein_coding | 197.796633 | -0.38213927 | 0.001172864 | 0.009426267 |
| ENSMUST00000058994.5,Tram1l1-001,protein_coding,ENSMUSG00000044528.5,protein_coding | 945.1408418 | -0.202447118 | 0.001175258 | 0.00944375 |
| ENSMUST00000134522.7,Zkscan17-004,processed_transcript,ENSMUSG00000020472.14,protein_coding | 345.1158619 | 0.447723759 | 0.001182634 | 0.009501245 |
| ENSMUST00000013886.8,Ppp1r12c-001,protein_coding,ENSMUSG00000019254.16,protein_coding | 6427.422643 | 0.146690407 | 0.001183631 | 0.009507477 |
| ENSMUST00000110320.8,Zc3h6-001,protein_coding,ENSMUSG00000042851.17,protein_coding | 706.4452288 | 0.431434203 | 0.001185689 | 0.009520458 |
| ENSMUST00000126604.7,Nipa2-009,protein_coding,ENSMUSG00000030452.16,protein_coding | 100.418202 | 0.48723237 | 0.001185611 | 0.009520458 |
| ENSMUST00000060634.13,Rpusd3-002,protein_coding,ENSMUSG00000051169.14,protein_coding | 158.8720052 | 0.565630069 | 0.00118917 | 0.009546627 |
| ENSMUST00000154322.7,Siglecg-004,retained_intron,ENSMUSG00000030468.12,protein_coding | 112.1681934 | -0.560984091 | 0.001191691 | 0.009565085 |
| ENSMUST00000159615.1,Cdk8-004,retained_intron,ENSMUSG00000029635.15,protein_coding | 103.2415293 | -0.59235147 | 0.001192359 | 0.009566875 |
| ENSMUST00000033875.9,Prosc-001,protein_coding,ENSMUSG00000031485.15,protein_coding | 1377.925951 | -0.220394216 | 0.001192291 | 0.009566875 |
| ENSMUST00000103212.9,Ube4b-001,protein_coding,ENSMUSG00000028960.18,protein_coding | 22948.07383 | 0.115021321 | 0.00119296 | 0.009568507 |
| ENSMUST00000179937.1,Tnfaip8-010,protein_coding,ENSMUSG00000062210.13,protein_coding | 114.5124186 | -0.461149874 | 0.001193007 | 0.009568507 |
| ENSMUST00000109510.8,Tcf20-202,protein_coding,ENSMUSG00000041852.14,protein_coding | 717.0405433 | 0.474675861 | 0.001193789 | 0.009572996 |
| ENSMUST00000109794.1,Pofut1-006,protein_coding,ENSMUSG00000046020.13,protein_coding | 330.5680042 | 0.528119074 | 0.001194323 | 0.009575494 |
| ENSMUST00000144258.7,Pkn1-004,protein_coding,ENSMUSG00000057672.16,protein_coding | 4189.52992 | 0.257650934 | 0.001195024 | 0.009579333 |
| ENSMUST00000124152.7,Myo19-007,processed_transcript,ENSMUSG00000020527.14,protein_coding | 276.9734531 | -0.563987634 | 0.001201199 | 0.009627038 |
| ENSMUST00000162479.1,Gtpbp8-004,nonsense_mediated_decay,ENSMUSG00000022668.10,protein_coding | 78.84702304 | 0.563855256 | 0.001205295 | 0.009658063 |
| ENSMUST00000121725.7,Gda-003,protein_coding,ENSMUSG00000058624.12,protein_coding | 278.3473149 | 0.396127188 | 0.001205607 | 0.009658767 |
| ENSMUST00000176149.7,Evi5l-001,protein_coding,ENSMUSG00000011832.15,protein_coding | 811.6201225 | 0.413346938 | 0.001206632 | 0.009665181 |
| ENSMUST00000094331.4,Ndnl2-001,protein_coding,ENSMUSG00000070520.4,protein_coding | 2204.489477 | 0.207542507 | 0.001210052 | 0.009690775 |
| ENSMUST00000132458.1,Ankrd24-012,protein_coding,ENSMUSG00000054708.17,protein_coding | 484.6482609 | 0.281925893 | 0.001211277 | 0.009698779 |
| ENSMUST00000161543.1,Nop56-003,retained_intron,ENSMUSG00000027405.16,protein_coding | 743.0484398 | -0.288778671 | 0.001211978 | 0.009700322 |
| ENSMUST00000080812.13,Zc3hc1-002,protein_coding,ENSMUSG00000039130.18,protein_coding | 10716.66989 | 0.160165193 | 0.001212684 | 0.009700322 |
| ENSMUST00000031810.14,Cep41-001,protein_coding,ENSMUSG00000029790.16,protein_coding | 1970.436225 | 0.134029858 | 0.001212657 | 0.009700322 |
| ENSMUST00000204302.2,Mical3-003,protein_coding,ENSMUSG00000051586.16,protein_coding | 204.6764733 | -0.389882889 | 0.001212788 | 0.009700322 |
| ENSMUST00000128512.1,Samd14-002,processed_transcript,ENSMUSG00000047181.12,protein_coding | 288.1145944 | 0.513973138 | 0.001212822 | 0.009700322 |
| ENSMUST00000093907.10,Sept9-001,protein_coding,ENSMUSG00000059248.13,protein_coding | 1719.700485 | 0.379024198 | 0.001212498 | 0.009700322 |
| ENSMUST00000140687.1,Gm15501-001,transcribed_processed_pseudogene,ENSMUSG00000087412.4,transcribed_processed_pseudogene | 967.7316796 | -0.385569541 | 0.001214118 | 0.009708888 |
| ENSMUST00000046404.6,Ubr7-201,protein_coding,ENSMUSG00000041712.6,protein_coding | 7222.35474 | 0.168916314 | 0.001216782 | 0.00972838 |
| ENSMUST00000181524.7,Gm19705-003,lincRNA,ENSMUSG00000097113.7,lincRNA | 55.6769381 | -0.598792006 | 0.001220484 | 0.009752544 |
| ENSMUST00000208429.1,Adgrg6-004,protein_coding,ENSMUSG00000039116.5,protein_coding | 489.308536 | 0.351938493 | 0.001220386 | 0.009752544 |
| ENSMUST00000140886.2,Vps53-003,processed_transcript,ENSMUSG00000017288.15,protein_coding | 21.88242151 | 0.599608927 | 0.001220229 | 0.009752544 |
| ENSMUST00000030184.11,Tpm2-002,protein_coding,ENSMUSG00000028464.16,protein_coding | 142.2960621 | -0.575260033 | 0.001224818 | 0.00978173 |
| ENSMUST00000021595.8,Psmc1-201,protein_coding,ENSMUSG00000021178.8,protein_coding | 24018.59208 | 0.134674502 | 0.001224452 | 0.00978173 |
| ENSMUST00000169061.7,Drosha-201,protein_coding,ENSMUSG00000022191.15,protein_coding | 1218.41346 | -0.428228526 | 0.0012246 | 0.00978173 |
| ENSMUST00000163623.1,Nedd9-202,protein_coding,ENSMUSG00000021365.14,protein_coding | 1690.787216 | -0.158660711 | 0.001227443 | 0.009800878 |
| ENSMUST00000154088.1,Usp14-006,retained_intron,ENSMUSG00000047879.16,protein_coding | 30.33471166 | -0.599433059 | 0.001228147 | 0.009804006 |
| ENSMUST00000183939.1,Nxf1-003,protein_coding,ENSMUSG00000010097.13,protein_coding | 113.627687 | 0.587100616 | 0.001228291 | 0.009804006 |
| ENSMUST00000167549.1,Endod1-001,protein_coding,ENSMUSG00000037419.9,protein_coding | 357.8978511 | 0.336616426 | 0.001228744 | 0.009805807 |
| ENSMUST00000155257.7,Mrto4-004,protein_coding,ENSMUSG00000028741.13,protein_coding | 81.65218276 | 0.480528688 | 0.001229769 | 0.009812164 |
| ENSMUST00000180835.1,6030442K20Rik-001,antisense,ENSMUSG00000097760.2,processed_transcript | 54.27021983 | -0.568206222 | 0.001231191 | 0.009819869 |
| ENSMUST00000036340.11,Fancd2-001,protein_coding,ENSMUSG00000034023.16,protein_coding | 6647.795676 | 0.14650787 | 0.001231185 | 0.009819869 |
| ENSMUST00000045761.6,Lzts3-201,protein_coding,ENSMUSG00000037703.14,protein_coding | 1964.952144 | -0.225466461 | 0.001232777 | 0.009830698 |
| ENSMUST00000122456.7,Rbm18-004,protein_coding,ENSMUSG00000026889.12,protein_coding | 113.9427948 | -0.536747009 | 0.00123301 | 0.009830739 |
| ENSMUST00000113155.1,Setd5-002,protein_coding,ENSMUSG00000034269.12,protein_coding | 9515.110951 | 0.278438301 | 0.001242666 | 0.009904056 |
| ENSMUST00000169591.7,Cyp4f16-001,protein_coding,ENSMUSG00000048440.15,protein_coding | 501.8204741 | -0.538121067 | 0.001242625 | 0.009904056 |
| ENSMUST00000035283.10,Nup88-003,protein_coding,ENSMUSG00000040667.11,protein_coding | 353.6139947 | 0.596168601 | 0.001246541 | 0.009933097 |
| ENSMUST00000110079.8,Tasp1-001,protein_coding,ENSMUSG00000039033.11,protein_coding | 435.8633378 | -0.434862145 | 0.001251435 | 0.009968405 |
| ENSMUST00000022597.13,Naa16-001,protein_coding,ENSMUSG00000022020.14,protein_coding | 2277.235102 | -0.210605411 | 0.00125138 | 0.009968405 |
| ENSMUST00000174313.7,Slit2-002,protein_coding,ENSMUSG00000031558.15,protein_coding | 344.7594183 | 0.404986183 | 0.001253027 | 0.009977868 |
| ENSMUST00000085658.4,6030419C18Rik-001,protein_coding,ENSMUSG00000066607.5,protein_coding | 2312.296525 | 0.215223857 | 0.001253086 | 0.009977868 |
| ENSMUST00000029692.14,Rit1-001,protein_coding,ENSMUSG00000028057.14,protein_coding | 1440.054779 | 0.153799996 | 0.001253432 | 0.009978776 |
| ENSMUST00000129160.7,Etv4-011,processed_transcript,ENSMUSG00000017724.14,protein_coding | 90.02288892 | 0.581782661 | 0.001255298 | 0.009991786 |
