## Supplementary Table 2 for "Loss of the RNA helicase *Dhx15* impairs endothelial energy metabolism, lymphatic drainage and tumor metastasis in mice"

| Accession number | Description | Log2 mean wild-type | Log2 mean siL-DHX15 | Log Fold-change ("WT" minus "siL-DHX15-LEC") | p-value |
|---|---|---|---|---|---|
| P07724 | Serum albumin OS=Mus musculus GN=Alb PE=1 SV=3 | 23.31 | 22.83 | 0.47 | 0.00674738 |
| O88888 | Amyloid beta A4 precursor protein-binding family A member 3 OS=Mus musculus GN=Apba3 PE=1 SV=1 | 24.34 | 23.22 | 1.12 | 0.00713274 |
| P35396 | Peroxisome proliferator-activated receptor delta OS=Mus musculus GN=Ppard PE=2 SV=1 | 23.05 | 22.03 | 1.02 | 0.00544691 |
| P31938 | Dual specificity mitogen-activated protein kinase kinase 1 OS=Mus musculus GN=Map2k1 PE=1 SV=2 | 22.29 | 21.37 | 0.92 | 0.00909615 |
| P62737 | Actin, aortic smooth muscle OS=Mus musculus GN=Acta2 PE=1 SV=1 | 24.80 | 23.96 | 0.84 | 8.19041E-05 |
| Q9CPT0 | Apoptosis facilitator Bcl-2-like protein 14 OS=Mus musculus GN=Bcl2l14 PE=1 SV=1 | 22.12 | 21.53 | 0.60 | 0.00824637 |
| Q9JLC8 | Sacsin OS=Mus musculus GN=Sacs PE=1 SV=2 | 22.43 | 21.96 | 0.47 | 0.00346342 |
| P10853 | Histone H2B type 1-F/J/L OS=Mus musculus GN=Hist1h2bf PE=1 SV=2 | 24.46 | 24.01 | 0.44 | 0.00646284 |
| P23818 | Glutamate receptor 1 OS=Mus musculus GN=Gria1 PE=1 SV=1 | 23.32 | 22.93 | 0.39 | 0.006028 |
| P10126 | Elongation factor 1-alpha 1 OS=Mus musculus GN=Eef1a1 PE=1 SV=3 | 23.28 | 23.00 | 0.28 | 0.0041607 |
| A2ASS6 | Titin OS=Mus musculus GN=Ttn PE=1 SV=1 | 26.32 | 26.07 | 0.25 | 0.000880608 |
| P14069 | Protein S100-A6 OS=Mus musculus GN=S100a6 PE=1 SV=3 | 26.15 | 25.97 | 0.18 | 0.00855212 |
| P62983 | Ubiquitin-40S ribosomal protein S27a OS=Mus musculus GN=Rps27a PE=1 SV=2 | 24.85 | 25.10 | -0.24 | 0.00870254 |
| P32261 | Antithrombin-III OS=Mus musculus GN=Serpinc1 PE=1 SV=1 | 26.42 | 26.68 | -0.27 | 0.00805759 |
| P63017 | Heat shock cognate 71 kDa protein OS=Mus musculus GN=Hspa8 PE=1 SV=1 | 24.90 | 25.18 | -0.28 | 0.00439213 |
| Q8CEZ1 | Single-pass membrane and coiled-coil domain-containing protein 1 OS=Mus musculus GN=Smco1 PE=2 SV=3 | 28.71 | 29.04 | -0.33 | 0.00395603 |
| Q6ZWY8 | Thymosin beta-10 OS=Mus musculus GN=Tmsb10 PE=1 SV=3 | 26.96 | 27.34 | -0.37 | 0.00516854 |
| sp | SWISS-PROT:P81187\|Fragment | 25.56 | 25.95 | -0.39 | 0.00723766 |
| P16045 | Galectin-1 OS=Mus musculus GN=Lgals1 PE=1 SV=3 | 25.40 | 25.79 | -0.39 | 0.0066363 |
| Q9CQN1 | Heat shock protein 75 kDa, mitochondrial OS=Mus musculus GN=Trap1 PE=1 SV=1 | 22.76 | 23.17 | -0.40 | 0.000996274 |
| Q9R1C7 | Pre-mRNA-processing factor 40 homolog A OS=Mus musculus GN=Prpf40a PE=1 SV=1 | 22.64 | 23.04 | -0.41 | 0.00709296 |
| A2AIV8 | Caspase recruitment domain-containing protein 9 OS=Mus musculus GN=Card9 PE=1 SV=1 | 28.10 | 28.51 | -0.41 | 0.000959653 |
| P17742 | Peptidyl-prolyl cis-trans isomerase A OS=Mus musculus GN=Ppia PE=1 SV=2 | 27.32 | 27.74 | -0.41 | 0.00099352 |
| P20152 | Vimentin OS=Mus musculus GN=Vim PE=1 SV=3 | 25.55 | 25.99 | -0.44 | 0.00221991 |
| P11499 | Heat shock protein HSP 90-beta OS=Mus musculus GN=Hsp90ab1 PE=1 SV=3 | 22.68 | 23.21 | -0.52 | 0.00210939 |
| Q61124 | Battenin OS=Mus musculus GN=Cln3 PE=1 SV=2 | 24.38 | 24.93 | -0.55 | 0.0075191 |
| Q9EQW7 | Kinesin-like protein KIF13A OS=Mus musculus GN=Kif13a PE=1 SV=1 | 24.61 | 25.20 | -0.59 | 2.63659E-05 |
| Q6ZPE2 | Myotubularin-related protein 5 OS=Mus musculus GN=Sbf1 PE=1 SV=2 | 27.15 | 27.74 | -0.59 | 0.00144266 |
| P26350 | Prothymosin alpha OS=Mus musculus GN=Ptma PE=1 SV=2 | 24.92 | 25.56 | -0.63 | 0.00162608 |
| Q61937 | Nucleophosmin OS=Mus musculus GN=Npm1 PE=1 SV=1 | 24.67 | 25.47 | -0.80 | 0.00348053 |
| P10639 | Thioredoxin OS=Mus musculus GN=Txn PE=1 SV=3 | 22.68 | 23.52 | -0.84 | 1.48244E-06 |
| P16858 | Glyceraldehyde-3-phosphate dehydrogenase OS=Mus musculus GN=Gapdh PE=1 SV=2 | 23.49 | 24.35 | -0.86 | 9.35024E-06 |
| P29699 | Alpha-2-HS-glycoprotein OS=Mus musculus GN=Ahsg PE=1 SV=1 | 22.51 | 23.52 | -1.01 | 0.00814973 |
| P63101 | 14-3-3 protein zeta/delta OS=Mus musculus GN=Ywhaz PE=1 SV=1 | 26.68 | 27.46 | -0.78 | 0.0104003 |
| P35700 | Peroxiredoxin-1 OS=Mus musculus GN=Prdx1 PE=1 SV=1 | 23.93 | 24.21 | -0.28 | 0.0131758 |
| P17182 | Alpha-enolase OS=Mus musculus GN=Eno1 PE=1 SV=3 | 24.22 | 24.66 | -0.44 | 0.0135079 |
| P99027 | 60S acidic ribosomal protein P2 OS=Mus musculus GN=Rplp2 PE=1 SV=3 | 26.28 | 26.94 | -0.65 | 0.0140773 |
| P09405 | Nucleolin OS=Mus musculus GN=Ncl PE=1 SV=2 | 22.52 | 22.99 | -0.47 | 0.0146771 |
| Q9CY62 | E3 ubiquitin-protein ligase RNF181 OS=Mus musculus GN=Rnf181 PE=1 SV=1 | 24.72 | 25.33 | -0.61 | 0.0167309 |
| P62827 | GTP-binding nuclear protein Ran OS=Mus musculus GN=Ran PE=1 SV=3 | 25.33 | 25.67 | -0.34 | 0.0167966 |
| P58252 | Elongation factor 2 OS=Mus musculus GN=Eef2 PE=1 SV=2 | 22.79 | 22.39 | 0.40 | 0.0179994 |
| P56394 | Cytochrome c oxidase copper chaperone OS=Mus musculus GN=Cox17 PE=1 SV=2 | 24.85 | 25.12 | -0.27 | 0.0183319 |
| P05064 | Fructose-bisphosphate aldolase A OS=Mus musculus GN=Aldoa PE=1 SV=2 | 24.37 | 23.92 | 0.45 | 0.0185559 |
| P62204 | Calmodulin OS=Mus musculus GN=Calm1 PE=1 SV=2 | 22.96 | 22.10 | 0.86 | 0.0188586 |
| O88569 | Heterogeneous nuclear ribonucleoproteins A2/B1 OS=Mus musculus GN=Hnrnpa2b1 PE=1 SV=2 | 25.62 | 25.84 | -0.22 | 0.0189241 |
| Q9D8N0 | Elongation factor 1-gamma OS=Mus musculus GN=Eef1g PE=1 SV=3 | 25.19 | 25.56 | -0.38 | 0.0198974 |
| P62806 | Histone H4 OS=Mus musculus GN=Hist1h4a PE=1 SV=2 | 23.65 | 23.34 | 0.32 | 0.020848 |
| P84099 | 60S ribosomal protein L19 OS=Mus musculus GN=Rpl19 PE=1 SV=1 | 26.16 | 26.37 | -0.21 | 0.0209767 |
| P21550 | Beta-enolase OS=Mus musculus GN=Eno3 PE=1 SV=3 | 21.59 | 22.24 | -0.65 | 0.0220349 |
| P62962 | Profilin-1 OS=Mus musculus GN=Pfn1 PE=1 SV=2 | 24.43 | 24.78 | -0.34 | 0.0234102 |
| P63028 | Translationally-controlled tumor protein OS=Mus musculus GN=Tpt1 PE=1 SV=1 | 22.69 | 23.44 | -0.75 | 0.0239483 |
| Q6URW6 | Myosin-14 OS=Mus musculus GN=Myh14 PE=1 SV=1 | 25.61 | 25.80 | -0.19 | 0.0244003 |
| P43274 | Histone H1.4 OS=Mus musculus GN=Hist1h1e PE=1 SV=2 | 21.17 | 21.74 | -0.56 | 0.0251253 |
| Q9ERE8 | Mesoderm development candidate 1 OS=Mus musculus GN=Mesdc1 PE=1 SV=1 | 22.42 | 22.05 | 0.37 | 0.0257104 |
| P63158 | High mobility group protein B1 OS=Mus musculus GN=Hmgb1 PE=1 SV=2 | 23.07 | 23.46 | -0.39 | 0.0260307 |
| P60843 | Eukaryotic initiation factor 4A-I OS=Mus musculus GN=Eif4a1 PE=1 SV=1 | 25.83 | 26.08 | -0.25 | 0.027568 |
| Q9EQU5 | Protein SET OS=Mus musculus GN=Set PE=1 SV=1 | 23.47 | 22.92 | 0.54 | 0.0293399 |
| P43276 | Histone H1.5 OS=Mus musculus GN=Hist1h1b PE=1 SV=2 | 22.58 | 22.89 | -0.32 | 0.0312889 |
| P14206 | 40S ribosomal protein SA OS=Mus musculus GN=Rpsa PE=1 SV=4 | 24.42 | 24.90 | -0.49 | 0.0315155 |
| P49312 | Heterogeneous nuclear ribonucleoprotein A1 OS=Mus musculus GN=Hnrnpa1 PE=1 SV=2 | 24.25 | 24.75 | -0.50 | 0.0315586 |
| C8YR32 | Lipoxygenase homology domain-containing protein 1 OS=Mus musculus GN=Loxhd1 PE=2 SV=1 | 27.99 | 28.25 | -0.27 | 0.0327487 |
| P52480 | Pyruvate kinase PKM OS=Mus musculus GN=Pkm PE=1 SV=4 | 23.05 | 23.81 | -0.76 | 0.0336031 |
| Q01768 | Nucleoside diphosphate kinase B OS=Mus musculus GN=Nme2 PE=1 SV=1 | 26.08 | 25.47 | 0.61 | 0.0359094 |
| P34884 | Macrophage migration inhibitory factor OS=Mus musculus GN=Mif PE=1 SV=2 | 25.47 | 25.26 | 0.21 | 0.0359861 |
| O70251 | Elongation factor 1-beta OS=Mus musculus GN=Eef1b PE=1 SV=5 | 20.96 | 21.29 | -0.34 | 0.0360416 |
| P47955 | 60S acidic ribosomal protein P1 OS=Mus musculus GN=Rplp1 PE=1 SV=1 | 24.69 | 24.03 | 0.66 | 0.0365638 |
| P47962 | 60S ribosomal protein L5 OS=Mus musculus GN=Rpl5 PE=1 SV=3 | 22.37 | 21.85 | 0.52 | 0.0369921 |
| P68368 | Tubulin alpha-4A chain OS=Mus musculus GN=Tuba4a PE=1 SV=1 | 22.59 | 23.15 | -0.56 | 0.0401247 |
| Q60692 | Proteasome subunit beta type-6 OS=Mus musculus GN=Psmb6 PE=1 SV=3 | 25.68 | 26.17 | -0.48 | 0.0428003 |
| O08807 | Peroxiredoxin-4 OS=Mus musculus GN=Prdx4 PE=1 SV=1 | 22.17 | 22.41 | -0.24 | 0.0429028 |
| P17751 | Triosephosphate isomerase OS=Mus musculus GN=Tpi1 PE=1 SV=4 | 24.73 | 24.94 | -0.21 | 0.0429457 |
| P09411 | Phosphoglycerate kinase 1 OS=Mus musculus GN=Pgk1 PE=1 SV=4 | 24.02 | 25.28 | -1.26 | 0.0432845 |
| P63242 | Eukaryotic translation initiation factor 5A-1 OS=Mus musculus GN=Eif5a PE=1 SV=2 | 22.32 | 22.72 | -0.40 | 0.043626 |
| P99024 | Tubulin beta-5 chain OS=Mus musculus GN=Tubb5 PE=1 SV=1 | 24.30 | 24.78 | -0.49 | 0.0447078 |
| Q9R0P3 | S-formylglutathione hydrolase OS=Mus musculus GN=Esd PE=1 SV=1 | 27.95 | 27.76 | 0.19 | 0.0449522 |
| Q60864 | Stress-induced-phosphoprotein 1 OS=Mus musculus GN=Stip1 PE=1 SV=1 | 22.90 | 22.64 | 0.26 | 0.0451283 |
| P57776 | Elongation factor 1-delta OS=Mus musculus GN=Eef1d PE=1 SV=3 | 22.97 | 22.61 | 0.35 | 0.0464474 |
| Q8BJ42 | Disks large-associated protein 2 OS=Mus musculus GN=Dlgap2 PE=1 SV=2 | 24.63 | 25.18 | -0.55 | 0.0466368 |
| P50543 | Protein S100-A11 OS=Mus musculus GN=S100a11 PE=1 SV=1 | 26.41 | 26.57 | -0.16 | 0.0471694 |
| P11679 | Keratin, type II cytoskeletal 8 OS=Mus musculus GN=Krt8 PE=1 SV=4 | 27.20 | 26.73 | 0.47 | 0.0482842 |
| P67984 | 60S ribosomal protein L22 OS=Mus musculus GN=Rpl22 PE=1 SV=2 | 21.96 | 21.47 | 0.49 | 0.0489795 |
| Q61703 | Inter-alpha-trypsin inhibitor heavy chain H2 OS=Mus musculus GN=Itih2 PE=1 SV=1 | ND | 27.61 | NA | NA |
| Q99KH8 | Serine/threonine-protein kinase 24 OS=Mus musculus GN=Stk24 PE=1 SV=1 | ND | 21.67 | NA | NA |
| Q8CGC6 | RNA-binding protein 28 OS=Mus musculus GN=Rbm28 PE=1 SV=4 | ND | 21.01 | NA | NA |
| Q8BJW5 | Nucleolar protein 11 OS=Mus musculus GN=Nol11 PE=2 SV=1 | ND | 20.84 | NA | NA |
| Q60715 | Prolyl 4-hydroxylase subunit alpha-1 OS=Mus musculus GN=P4ha1 PE=1 SV=2 | ND | 21.88 | NA | NA |
| P59759 | MKL/myocardin-like protein 2 OS=Mus musculus GN=Mkl2 PE=1 SV=1 | ND | 22.46 | NA | NA |
| Q99MI1 | ELKS/Rab6-interacting/CAST family member 1 OS=Mus musculus GN=Erc1 PE=1 SV=1 | ND | 20.89 | NA | NA |
| P14901 | Heme oxygenase 1 OS=Mus musculus GN=Hmox1 PE=1 SV=1 | ND | 21.95 | NA | NA |
| O35129 | Prohibitin-2 OS=Mus musculus GN=Phb2 PE=1 SV=1 | ND | 22.26 | NA | NA |
| P29391 | Ferritin light chain 1 OS=Mus musculus GN=Ftl1 PE=1 SV=2 | ND | 22.43 | NA | NA |
| Q9JLI8 | Squamous cell carcinoma antigen recognized by T-cells 3 OS=Mus musculus GN=Sart3 PE=1 SV=1 | ND | 20.89 | NA | NA |
| Q6P5E4 | UDP-glucose:glycoprotein glucosyltransferase 1 OS=Mus musculus GN=Uggt1 PE=1 SV=4 | ND | 20.84 | NA | NA |
| Q8CGK3 | Lon protease homolog, mitochondrial OS=Mus musculus GN=Lonp1 PE=1 SV=2 | ND | 21.53 | NA | NA |
| Q64378 | Peptidyl-prolyl cis-trans isomerase FKBP5 OS=Mus musculus GN=Fkbp5 PE=1 SV=1 | ND | 23.04 | NA | NA |
| Q9DBG6 | Dolichyl-diphosphooligosaccharide--protein glycosyltransferase subunit 2 OS=Mus musculus GN=Rpn2 PE=1 SV=1 | ND | 21.76 | NA | NA |
| P14576 | Signal recognition particle 54 kDa protein OS=Mus musculus GN=Srp54 PE=1 SV=2 | ND | 21.33 | NA | NA |
| P97429 | Annexin A4 OS=Mus musculus GN=Anxa4 PE=1 SV=4 | ND | 22.42 | NA | NA |
| Q8K009 | Mitochondrial 10-formyltetrahydrofolate dehydrogenase OS=Mus musculus GN=Aldh1l2 PE=1 SV=2 | ND | 21.97 | NA | NA |
| P54726 | UV excision repair protein RAD23 homolog A OS=Mus musculus GN=Rad23a PE=1 SV=2 | ND | 22.02 | NA | NA |
| Q62351 | Transferrin receptor protein 1 OS=Mus musculus GN=Tfrc PE=1 SV=1 | ND | 21.40 | NA | NA |
| Q9CSU0 | Regulation of nuclear pre-mRNA domain-containing protein 1B OS=Mus musculus GN=Rprd1b PE=1 SV=2 | ND | 20.92 | NA | NA |
| Q60841 | Reelin OS=Mus musculus GN=Reln PE=1 SV=3 | ND | 26.35 | NA | NA |
| Q8K5B2 | Multiple coagulation factor deficiency protein 2 homolog OS=Mus musculus GN=Mcfd2 PE=1 SV=1 | ND | 23.19 | NA | NA |
| A2AMZ4 | ADH dehydrogenase [ubiquinone] 1 alpha subcomplex assembly factor 8 OS=Mus musculus GN=Ndufaf8 PE=1 SV= | 25.85 | ND | NA | NA |
| Q80Y83 | Dixin OS=Mus musculus GN=Dixdc1 PE=1 SV=1 | 23.24 | ND | NA | NA |
| P0CL69 | Zinc finger protein 703 OS=Mus musculus GN=Znf703 PE=1 SV=1 | 23.02 | ND | NA | NA |
| Q9JKP5 | Muscleblind-like protein 1 OS=Mus musculus GN=Mbnl1 PE=1 SV=1 | 21.20 | ND | NA | NA |
| Q8CF98 | Collectin-10 OS=Mus musculus GN=Colec10 PE=2 SV=1 | 18.04 | ND | NA | NA |
| Q70IV5 | Synemin OS=Mus musculus GN=Synm PE=1 SV=2 | 18.54 | ND | NA | NA |
| Q8C5S7 | Meiotic recombination protein REC8 homolog OS=Mus musculus GN=Rec8 PE=1 SV=1 | 25.72 | ND | NA | NA |
| Q64310 | Surfeit locus protein 4 OS=Mus musculus GN=Surf4 PE=1 SV=1 | 21.52 | ND | NA | NA |

| | | | | | |
|---|---|---|---|---|---|
| Q571C7 | Transcription factor TFIIIB component B'' homolog OS=Mus musculus GN=Bdp1 PE=2 SV=2 | 25.41 | ND | NA | NA |
| P62874 | Guanine nucleotide-binding protein G(I)/G(S)/G(T) subunit beta-1 OS=Mus musculus GN=Gnb1 PE=1 SV=3 | 22.38 | ND | NA | NA |
| Q9ET30 | Transmembrane 9 superfamily member 3 OS=Mus musculus GN=Tm9sf3 PE=1 SV=1 | 22.64 | ND | NA | NA |
| Q91WG2 | Rab GTPase-binding effector protein 2 OS=Mus musculus GN=Rabep2 PE=1 SV=3 | 20.40 | ND | NA | NA |
| Q5SW19 | Clustered mitochondria protein homolog OS=Mus musculus GN=Cluh PE=1 SV=2 | 21.87 | ND | NA | NA |
| Q9WVK4 | EH domain-containing protein 1 OS=Mus musculus GN=Ehd1 PE=1 SV=1 | 21.24 | ND | NA | NA |
| O35286 | Pre-mRNA-splicing factor ATP-dependent RNA helicase DHX15 OS=Mus musculus GN=Dhx15 PE=1 SV=2 | 23.15 | ND | NA | NA |

ND= not detectable
NA= not applicable
