## Supplementary Table 3 for "Loss of the RNA helicase *Dhx15* impairs endothelial energy metabolism, lymphatic drainage and tumor metastasis in mice"

| Gene ID | gene Symbol | chr | strand | long Exon Start base | long Exon End base | Short Exon Start base | Short Exon End base | FDR | Inclusion Level Difference |
|---|---|---|---|---|---|---|---|---|---|
| ENSMUSG00000022822.15 | Abcc5 | chr16 | - | 20404488 | 20404708 | 20404565 | 20404708 | 0.00567811924814 | 0.051 |
| ENSMUSG00000053411.16 | Cbx7 | chr15 | - | 79918276 | 79918476 | 79918322 | 79918476 | 0.0139411233857 | -0.051 |
| ENSMUSG00000046897.16 | Zfp740 | chr15 | + | 102204570 | 102205086 | 102204570 | 102204883 | 0.0274336736997 | -0.054 |
| ENSMUSG00000058799.14 | Nap1l1 | chr10 | + | 111490027 | 111490135 | 111490027 | 111490037 | 0.000552179743019 | -0.056 |
| ENSMUSG00000026479.13 | Lamc2 | chr1 | - | 153129963 | 153130150 | 153130035 | 153130150 | 0.00214613267752 | 0.057 |
| ENSMUSG00000045411.16 | 2410002F23Rik | chr7 | + | 44246807 | 44247047 | 44246807 | 44246823 | 0.0119988469525 | 0.057 |
| ENSMUSG00000053070.5 | 9230110C19Rik | chr9 | - | 8042628 | 8042813 | 8042669 | 8042813 | 0.0062169711463 | 0.059 |
| ENSMUSG00000024937.14 | Ehbp1l1 | chr19 | - | 5717661 | 5720407 | 5720184 | 5720407 | 0.00447022578475 | -0.06 |
| ENSMUSG00000022307.15 | Oxr1 | chr15 | + | 41830957 | 41831129 | 41830957 | 41831077 | 0.00935212672276 | -0.062 |
| ENSMUSG00000031109.16 | Enox2 | chrX | - | 49286842 | 49286904 | 49286844 | 49286904 | 0.0272326363912 | 0.064 |
| ENSMUSG00000073563.2 | Csnk1g3 | chr18 | + | 53936852 | 53936948 | 53936852 | 53936942 | 0.00688007475848 | -0.064 |
| ENSMUSG00000037570.15 | Mcrs1 | chr15 | - | 99251721 | 99251961 | 99251861 | 99251961 | 0.020558500783 | 0.065 |
| ENSMUSG00000068284.14 | Usf3 | chr16 | + | 44198903 | 44199167 | 44198903 | 44199021 | 0.000624088893201 | -0.065 |
| ENSMUSG00000061607.14 | Mdc1 | chr17 | + | 35850294 | 35851005 | 35850294 | 35850513 | 0.0127838201487 | 0.066 |
| ENSMUSG00000030057.15 | Cnbp | chr6 | - | 87845655 | 87845781 | 87845676 | 87845781 | 0.00148779629937 | -0.067 |
| ENSMUSG00000025968.16 | Ndufs1 | chr1 | - | 63176524 | 63176667 | 63176585 | 63176667 | 0.0233667141172 | -0.068 |
| ENSMUSG00000022992.16 | Kansl2 | chr15 | - | 98524400 | 98524590 | 98524560 | 98524590 | 0.0330392811749 | -0.069 |
| ENSMUSG00000042772.15 | Smg7 | chr1 | - | 152848595 | 152849016 | 152848733 | 152849016 | 0.0338284893696 | -0.069 |
| ENSMUSG00000024799.14 | Tm7sf2 | chr19 | - | 6067710 | 6067867 | 6067777 | 6067867 | 0.0458058347801 | 0.071 |
| ENSMUSG00000036678.7 | Aaas | chr15 | - | 102350543 | 102350759 | 102350565 | 102350759 | 0.000719212995039 | 0.072 |
| ENSMUSG00000033032.15 | Afap1l1 | chr18 | - | 61756812 | 61756934 | 61756839 | 61756934 | 0.0367875691 68 | 0.074 |
| ENSMUSG00000045128.9 | Rpl18a | chr8 | - | 70897128 | 70897283 | 70897279 | 70897283 | 0.0330392811749 | 0.076 |
| ENSMUSG00000038900.17 | Rpl12 | chr2 | + | 32962074 | 32962122 | 32962074 | 32962108 | 0.0296601095735 | 0.077 |
| ENSMUSG00000034908.16 | Sidt2 | chr9 | - | 45947423 | 45947552 | 45947540 | 45947552 | 0.0151410791413 | 0.078 |
| ENSMUSG00000024327.15 | Slc39a7 | chr17 | - | 34031540 | 34031656 | 34031547 | 34031656 | 0.03721582376 | 0.078 |
| ENSMUSG00000023961.16 | Enpp4 | chr17 | - | 44105521 | 44105764 | 44105613 | 44105764 | 0.0337482297191 | -0.078 |
| ENSMUSG00000031833.10 | Mast3 | chr8 | - | 70789537 | 70789703 | 70789682 | 70789703 | 0.000339240400523 | 0.079 |
| ENSMUSG00000045128.9 | Rpl18a | chr8 | - | 70897130 | 70897283 | 70897279 | 70897283 | 0.0274336736997 | 0.08 |
| ENSMUSG00000049106.6 | Dcaf5 | chr12 | - | 80436184 | 80436601 | 80436363 | 80436601 | 0.000346806906005 | -0.08 |
| ENSMUSG00000078974.10 | Sec61g | chr11 | - | 16508074 | 16508166 | 16508099 | 16508166 | 0.0472738627036 | 0.085 |
| ENSMUSG00000027797.15 | Dclk1 | chr3 | + | 55477804 | 55477937 | 55477804 | 55477889 | 0.00485049809489 | 0.086 |
| ENSMUSG00000074182.8 | Znhit6 | chr3 | + | 145576335 | 145576407 | 145576335 | 145576359 | 0.00323054545681 | 0.087 |
| ENSMUSG00000052369.7 | Tmem106c | chr15 | + | 97964228 | 97964378 | 97964228 | 97964335 | 0.0246332202846 | 0.087 |
| ENSMUSG00000028173.10 | Wls | chr3 | + | 159934250 | 159934473 | 159934250 | 159934369 | 0.000997810955654 | 0.089 |
| ENSMUSG00000032294.17 | Pkm | chr9 | + | 59658242 | 59658549 | 59658242 | 59658302 | 0.0458058347801 | -0.091 |
| ENSMUSG00000042557.14 | Sin3a | chr9 | + | 57072039 | 57072216 | 57072039 | 57072063 | 0.0378219738726 | 0.092 |
| ENSMUSG00000109324.1 | Prmt1 | chr7 | - | 44983734 | 44983973 | 44983770 | 44983973 | 0.0282964413054 | 0.093 |
| ENSMUSG00000021957.6 | Tkt | chr14 | + | 30557645 | 30557693 | 30557645 | 30557688 | 0.0200596189979 | 0.095 |
| ENSMUSG00000045128.9 | Rpl18a | chr8 | - | 70897168 | 70897412 | 70897279 | 70897412 | 0.0195727420136 | 0.097 |
| ENSMUSG00000023030.16 | Slc11a2 | chr15 | - | 100395216 | 100395323 | 100395269 | 100395323 | 0.000339240400523 | 0.098 |
| ENSMUSG00000029177.9 | Cenpa | chr5 | + | 30667580 | 30667687 | 30667580 | 30667607 | 0.00274981876519 | 0.098 |
| ENSMUSG00000028797.20 | Tmem234 | chr4 | + | 129600982 | 129601243 | 129600982 | 129601134 | 0.000786629409985 | 0.101 |
| ENSMUSG00000002031.12 | Ift46 | chr9 | + | 44773262 | 44773477 | 44773262 | 44773287 | 0.0384379552297 | 0.102 |
| ENSMUSG00000046027.17 | Stard5 | chr7 | + | 83631999 | 83632096 | 83631999 | 83632088 | 0.046381865126 | 0.102 |
| ENSMUSG00000052369.7 | Tmem106c | chr15 | + | 97964228 | 97964503 | 97964228 | 97964335 | 0.00729438711004 | 0.103 |
| ENSMUSG00000029622.16 | Arpc1b | chr5 | + | 145123973 | 145124290 | 145123973 | 145124053 | 0.0272326363912 | 0.103 |
| ENSMUSG00000036026.15 | Tmem63b | chr17 | - | 45685781 | 45685973 | 45685915 | 45685973 | 0.002859907793 | 0.113 |
| ENSMUSG00000057113.13 | Npm1 | chr11 | - | 33160823 | 33160921 | 33160827 | 33160921 | 0.02319659571 | -0.119 |
| ENSMUSG00000034165.16 | Ccnd3 | chr17 | + | 47594321 | 47594483 | 47594321 | 47594405 | 0.0100734442453 | 0.121 |
| ENSMUSG00000096916.7 | Zfp850 | chr7 | - | 28014015 | 28014115 | 28014020 | 28014115 | 0.0188839616164 | -0.121 |
| ENSMUSG00000037772.13 | Mrpl23 | chr7 | + | 142533161 | 142533191 | 142533161 | 142533166 | 0.0367903299154 | -0.121 |
| ENSMUSG00000023235.14 | Ccl25 | chr8 | + | 4347713 | 4348340 | 4347713 | 4347913 | 0.0121173571915 | -0.123 |
| ENSMUSG00000013822.7 | Elof1 | chr9 | - | 22116654 | 22117086 | 22116844 | 22117086 | 0.0294746483524 | -0.123 |
| ENSMUSG00000042042.14 | Csgalnact2 | chr6 | - | 118124311 | 118126338 | 118126121 | 118126338 | 0.0395321775973 | -0.125 |
| ENSMUSG00000031583.13 | Wrn | chr8 | - | 33342976 | 33343692 | 33343546 | 33343692 | 0.0309502809243 | 0.126 |
| ENSMUSG00000037608.16 | Bclaf1 | chr10 | + | 20332505 | 20332569 | 20332505 | 20332518 | 0.0243076279142 | 0.127 |
| ENSMUSG00000089917.7 | Uckl1 | chr2 | - | 181571168 | 181571257 | 181571237 | 181571257 | 0.0477690751211 | -0.128 |
| ENSMUSG00000027777.15 | Schip1 | chr3 | + | 68493490 | 68493658 | 68493490 | 68493605 | 0.0115027889629 | 0.131 |
| ENSMUSG00000032258.15 | Lca5 | chr9 | - | 83440923 | 83441116 | 83441036 | 83441116 | 0.0384379552297 | 0.133 |
| ENSMUSG00000038260.10 | Trpm4 | chr7 | - | 45305335 | 45305494 | 45305373 | 45305494 | 0.00734838496271 | -0.134 |
| ENSMUSG00000022974.12 | Paxbp1 | chr16 | - | 91030415 | 91030556 | 91030432 | 91030556 | 0.020558500783 | 0.135 |
| ENSMUSG00000055401.14 | Fbxo6 | chr4 | - | 148151768 | 148151925 | 148151829 | 148151925 | 0.0227792217016 | 0.135 |
| ENSMUSG00000019461.12 | Plscr3 | chr11 | + | 69846375 | 69846515 | 69846375 | 69846497 | 0.037159415225 | 0.136 |
| ENSMUSG00000063052.9 | Lrrc40 | chr3 | + | 158048400 | 158048559 | 158048400 | 158048546 | 0.0458058347801 | 0.137 |
| ENSMUSG00000105617.4 | Gm43809 | chr5 | + | 31613971 | 31614156 | 31613971 | 31614022 | 0.00214613267752 | -0.137 |
| ENSMUSG00000037772.13 | Mrpl23 | chr7 | + | 142533011 | 142533189 | 142533011 | 142533166 | 0.0288722395909 | -0.137 |
| ENSMUSG00000059743.12 | Fdps | chr3 | - | 89101654 | 89101888 | 89101861 | 89101888 | 0.036397552023 | 0.138 |
| ENSMUSG00000034445.7 | Cyb561a3 | chr19 | + | 10577453 | 10577692 | 10577453 | 10577563 | 0.0264007627707 | 0.142 |
| ENSMUSG00000002908.17 | Kcnn1 | chr8 | - | 70848174 | 70848302 | 70848199 | 70848302 | 0.00935212672276 | 0.147 |
| ENSMUSG00000056821.13 | 1700028I16Rik | chr10 | + | 82812296 | 82812403 | 82812296 | 82812359 | 0.048798840157 | -0.147 |
| ENSMUSG00000030397.10 | Mark4 | chr7 | - | 19456956 | 19457099 | 19457065 | 19457099 | 0.022877164925 | -0.149 |
| ENSMUSG00000037816.10 | Fbxw17 | chr13 | + | 50419839 | 50420028 | 50419839 | 50419877 | 0.00511049630358 | 0.15 |
| ENSMUSG00000091228.1 | Gm20390 | chr11 | - | 93968173 | 93968293 | 93968186 | 93968293 | 0.0458058347801 | 0.153 |
| ENSMUSG00000038260.10 | Trpm4 | chr7 | - | 45305344 | 45305494 | 45305373 | 45305494 | 0.00991896417098 | -0.153 |
| ENSMUSG00000037772.13 | Mrpl23 | chr7 | + | 142533160 | 142533241 | 142533160 | 142533166 | 0.020558500783 | -0.154 |
| ENSMUSG00000044968.16 | Napepld | chr5 | - | 21701204 | 21701396 | 21701356 | 21701396 | 0.0472738627036 | 0.155 |
| ENSMUSG00000027637.3 | 1110008F13Rik | chr2 | + | 156866651 | 156866877 | 156866651 | 156866736 | 0.00905729492155 | -0.155 |
| ENSMUSG00000034820.3 | Cpsf7 | chr19 | + | 10532130 | 10532334 | 10532130 | 10532196 | 0.00671188495912 | 0.157 |
| ENSMUSG00000042766.17 | Trim46 | chr3 | - | 89245704 | 89245897 | 89245714 | 89245897 | 0.0380660730859 | -0.157 |
| ENSMUSG00000002249.18 | Tead3 | chr17 | - | 28336172 | 28336344 | 28336273 | 28336344 | 0.0440468256868 | 0.159 |
| ENSMUSG00000097867.1 | Lppos | chr16 | - | 24393353 | 24393588 | 24393481 | 24393588 | 0.0144859804085 | -0.16 |
| ENSMUSG00000029430.13 | Ran | chr5 | + | 129020911 | 129021117 | 129020911 | 129021071 | 0.037159415225 | 0.162 |
| ENSMUSG00000020776.18 | Fbf1 | chr11 | - | 116167516 | 116167697 | 116167551 | 116167697 | 0.00400303807729 | -0.162 |
| ENSMUSG00000037772.13 | Mrpl23 | chr7 | + | 142533160 | 142533236 | 142533160 | 142533166 | 0.0216314612278 | -0.162 |
| ENSMUSG00000097867.1 | Lppos | chr16 | - | 24393358 | 24393588 | 24393481 | 24393588 | 0.0110951024435 | -0.164 |
| ENSMUSG00000001847.14 | Rac1 | chr5 | - | 143517157 | 143517229 | 143517159 | 143517229 | 0.037159415225 | 0.165 |
| ENSMUSG00000025135.12 | Anapc11 | chr11 | + | 120603943 | 120604185 | 120603943 | 120603970 | 0.0101030606726 | -0.168 |
| ENSMUSG00000020074.6 | Ccar1 | chr10 | - | 62792141 | 62792368 | 62792204 | 62792368 | 0.02319659571 | 0.173 |
| ENSMUSG00000086877.7 | A230072C01Rik | chrX | + | 20962113 | 20962271 | 20962113 | 20962218 | 0.0487815772734 | -0.175 |
| ENSMUSG00000028936.15 | Rpl22 | chr4 | + | 152329259 | 152329503 | 152329259 | 152329399 | 0.000719212995039 | 0.177 |
| ENSMUSG00000061859.16 | Patj | chr4 | + | 98395854 | 98395924 | 98395854 | 98395865 | 0.0222677847323 | 0.177 |

| | | | | | | | | | |
|---|---|---|---|---|---|---|---|---|---|
| ENSMUSG00000027509.11 | Rae1 | chr2 | + | 173009369 | 173009506 | 173009369 | 173009477 | 0.037159415225 | -0.179 |
| ENSMUSG00000087365.7 | C430049B03Rik | chrX | - | 53055696 | 53055905 | 53055855 | 53055905 | 0.00147785089173 | 0.181 |
| ENSMUSG00000050157.5 | Gm867 | chr10 | - | 75939161 | 75939331 | 75939172 | 75939331 | 0.0382146420645 | 0.182 |
| ENSMUSG00000064138.13 | Fam172a | chr13 | - | 78066055 | 78066168 | 78066055 | 78066129 | 0.0471273169129 | 0.182 |
| ENSMUSG00000029822.15 | Osbpl3 | chr6 | - | 50370117 | 50370215 | 50370122 | 50370215 | 0.0380561895063 | 0.184 |
| ENSMUSG00000024287.7 | Thoc1 | chr18 | + | 9959097 | 9959234 | 9959097 | 9959171 | 0.0269095053606 | -0.184 |
| ENSMUSG00000020074.6 | Ccar1 | chr10 | - | 62792129 | 62792368 | 62792204 | 62792368 | 0.0396970955227 | 0.185 |
| ENSMUSG00000047909.11 | Ankrd16 | chr2 | + | 11779884 | 11780128 | 11779884 | 11779897 | 0.000878348578856 | 0.186 |
| ENSMUSG00000073795.11 | 6430531B16Rik | chr7 | - | 139978033 | 139978723 | 139978546 | 139978723 | 0.0127253249255 | 0.186 |
| ENSMUSG00000047909.11 | Ankrd16 | chr2 | + | 11779884 | 11780084 | 11779884 | 11779897 | 0.00236261382259 | 0.187 |
| ENSMUSG00000049436.4 | Upk1b | chr16 | - | 38800147 | 38800203 | 38800149 | 38800203 | 0.0472738627036 | -0.189 |
| ENSMUSG00000110105.1 | Vmn2r29 | chr7 | - | 7278096 | 7278210 | 7278116 | 7278210 | 0.0447719346276 | 0.193 |
| ENSMUSG00000063316.13 | Rpl27 | chr11 | + | 101443954 | 101444086 | 101443954 | 101443999 | 0.0395321775973 | 0.196 |
| ENSMUSG00000036768.6 | Kif15 | chr9 | + | 122993041 | 122993219 | 122993041 | 122993087 | 0.0120066521436 | 0.197 |
| ENSMUSG00000060002.14 | Chpt1 | chr10 | - | 88475230 | 88475477 | 88475332 | 88475477 | 0.0012954297491 | 0.205 |
| ENSMUSG00000090841.1 | Myl6 | chr10 | - | 128491283 | 128491304 | 128491285 | 128491304 | 0.00315566102115 | 0.209 |
| ENSMUSG00000046792.8 | Zfp787 | chr7 | - | 6134536 | 6134708 | 6134641 | 6134708 | 0.0332891843922 | 0.211 |
| ENSMUSG00000042203.7 | Tbc1d22b | chr17 | + | 29571664 | 29571896 | 29571664 | 29571794 | 0.0254685385472 | -0.212 |
| ENSMUSG00000057842.13 | Zfp595 | chr13 | - | 67320791 | 67320954 | 67320827 | 67320954 | 0.00485049809489 | 0.217 |
| ENSMUSG00000042104.18 | Uggt2 | chr14 | - | 119032628 | 119032787 | 119032741 | 119032787 | 0.00181045960678 | -0.217 |
| ENSMUSG00000049295.16 | Zfp219 | chr14 | - | 52014850 | 52015095 | 52014947 | 52015095 | 0.0066742990511 | 0.219 |
| ENSMUSG00000107478.1 | Gm45234 | chr6 | - | 124755864 | 124756021 | 124755983 | 124756021 | 0.0254685385472 | 0.22 |
| ENSMUSG00000037563.15 | Rps16 | chr7 | + | 28350651 | 28350877 | 28350651 | 28350853 | 0.00823868959558 | -0.222 |
| ENSMUSG00000029823.16 | Luc7l2 | chr6 | + | 38554979 | 38555094 | 38554979 | 38555063 | 0.00513063614941 | 0.225 |
| ENSMUSG00000031708.16 | Tecr | chr8 | - | 83572518 | 83572753 | 83572680 | 83572753 | 0.00190442123858 | -0.225 |
| ENSMUSG00000002458.13 | Rgs19 | chr2 | - | 181689295 | 181689500 | 181689376 | 181689500 | 0.00113232645411 | 0.226 |
| ENSMUSG00000110105.1 | Vmn2r29 | chr7 | - | 7278093 | 7278232 | 7278116 | 7278232 | 0.0472738627036 | 0.226 |
| ENSMUSG00000044968.16 | Napepld | chr5 | - | 21700974 | 21701394 | 21701356 | 21701394 | 0.0490121700912 | 0.226 |
| ENSMUSG00000031708.16 | Tecr | chr8 | - | 83572575 | 83572753 | 83572680 | 83572753 | 0.00214613267752 | -0.236 |
| ENSMUSG00000057342.15 | Sphk2 | chr7 | - | 45717175 | 45717408 | 45717242 | 45717408 | 0.000576242502148 | -0.238 |
| ENSMUSG00000031708.16 | Tecr | chr8 | - | 83572551 | 83572753 | 83572680 | 83572753 | 0.0016498379739 | -0.241 |
| ENSMUSG00000067629.11 | Syngap1 | chr17 | + | 26952396 | 26952440 | 26952396 | 26952436 | 0.000902128668462 | 0.243 |
| ENSMUSG00000020473.13 | Aebp1 | chr11 | + | 5862744 | 5862972 | 5862744 | 5862765 | 0.0330392811749 | -0.243 |
| ENSMUSG00000022587.14 | Ly6e | chr15 | + | 74955652 | 74955818 | 74955652 | 74955742 | 0.000878348578856 | 0.247 |
| ENSMUSG00000061859.16 | Patj | chr4 | + | 98395784 | 98395865 | 98395784 | 98395837 | 0.0458058347801 | -0.252 |
| ENSMUSG00000001348.15 | Acp5 | chr9 | - | 22131500 | 22131653 | 22131589 | 22131653 | 0.00698917367671 | 0.254 |
| ENSMUSG00000068246.5 | Apol9b | chr15 | + | 77729120 | 77729246 | 77729120 | 77729148 | 0.00191893565418 | 0.255 |
| ENSMUSG00000057738.13 | Sptan1 | chr2 | + | 29968775 | 29968966 | 29968775 | 29968826 | 0.0395175303373 | -0.256 |
| ENSMUSG00000087433.1 | Gm14167 | chr2 | - | 151542252 | 151542348 | 151542306 | 151542348 | 0.00306452147954 | 0.257 |
| ENSMUSG00000021266.16 | Wars | chr12 | - | 108888311 | 108888548 | 108888371 | 108888548 | 0.00214613267752 | 0.258 |
| ENSMUSG00000019969.13 | Psen1 | chr12 | + | 83688562 | 83688933 | 83688562 | 83688701 | 0.045445704469 | 0.258 |
| ENSMUSG00000092176.1 | Gm20460 | chr17 | - | 34627246 | 34627349 | 34627324 | 34627349 | 0.00244851652229 | 0.264 |
| ENSMUSG00000033047.4 | Eif3l | chr15 | + | 79075222 | 79075457 | 79075222 | 79075303 | 0.00309511327355 | 0.265 |
| ENSMUSG00000063052.9 | Lrrc40 | chr3 | + | 158048400 | 158048615 | 158048400 | 158048546 | 0.0042355950113 | 0.267 |
| ENSMUSG00000001999.15 | Blvra | chr2 | + | 127070692 | 127070932 | 127070692 | 127070725 | 0.00239175090049 | 0.269 |
| ENSMUSG00000022307.15 | Oxr1 | chr15 | + | 41789448 | 41789618 | 41789448 | 41789506 | 0.0110951024435 | -0.274 |
| ENSMUSG00000030846.15 | Tial1 | chr7 | - | 128447038 | 128447107 | 128447041 | 128447107 | 0.00886679529264 | -0.276 |
| ENSMUSG00000002846.9 | Timmdc1 | chr16 | - | 38522638 | 38522663 | 38522643 | 38522663 | 0.0347994848683 | 0.281 |
| ENSMUSG00000056758.14 | Hmga2 | chr10 | - | 120475948 | 120476064 | 120475952 | 120476064 | 0.00671188495912 | -0.282 |
| ENSMUSG00000036980.13 | Taf6 | chr5 | - | 138182354 | 138182487 | 138182413 | 138182487 | 0.0336275226959 | 0.286 |
| ENSMUSG00000033047.4 | Eif3l | chr15 | + | 79075222 | 79075360 | 79075222 | 79075303 | 0.0396970955227 | 0.286 |
| ENSMUSG00000022890.13 | Atp5j | chr16 | - | 84834637 | 84834832 | 84834796 | 84834832 | 0.0032447775349 | 0.289 |
| ENSMUSG00000043716.13 | Rpl7 | chr1 | - | 16103989 | 16104227 | 16104010 | 16104227 | 0.00835581183514 | 0.29 |
| ENSMUSG00000036246.14 | Gmip | chr8 | + | 69813264 | 69813435 | 69813264 | 69813291 | 0.0133460748734 | -0.29 |
| ENSMUSG00000025142.17 | Aspscr1 | chr11 | + | 120707904 | 120707996 | 120707904 | 120707957 | 0.000198179637697 | -0.292 |
| ENSMUSG00000022475.18 | Hdac7 | chr15 | - | 97795583 | 97795831 | 97795606 | 97795831 | 0.00030955550446 | -0.293 |
| ENSMUSG00000022890.13 | Atp5j | chr16 | - | 84834656 | 84834832 | 84834796 | 84834832 | 0.00381492617223 | 0.301 |
| ENSMUSG00000037239.8 | Spred3 | chr7 | - | 29167351 | 29167447 | 29167372 | 29167447 | 0.00308472241183 | -0.302 |
| ENSMUSG00000024530.7 | Slmo1 | chr18 | + | 67476963 | 67477103 | 67476963 | 67477066 | 0.0115092784086 | 0.307 |
| ENSMUSG00000058600.13 | Rpl30 | chr15 | - | 34443020 | 34443132 | 34443024 | 34443132 | 0.0422303164895 | 0.307 |
| ENSMUSG00000089872.10 | Rps6kc1 | chr1 | - | 190871272 | 190871491 | 190871391 | 190871491 | 0.00147785089173 | 0.312 |
| ENSMUSG00000110631.1 | RP24-286J21.7 | chr8 | + | 121833246 | 121833346 | 121833246 | 121833332 | 0.0264007627707 | -0.313 |
| ENSMUSG00000031930.11 | Wwp2 | chr8 | + | 107552276 | 107552318 | 107552276 | 107552304 | 0.0487815772734 | -0.316 |
| ENSMUSG00000086035.1 | Gm12610 | chr4 | + | 89460928 | 89461035 | 89460928 | 89460964 | 0.0418562250708 | 0.318 |
| ENSMUSG00000035086.13 | Becn1 | chr11 | - | 101296193 | 101296313 | 101296257 | 101296313 | 0.000120469198123 | 0.319 |
| ENSMUSG00000017404.12 | Rpl19 | chr11 | + | 98026956 | 98027113 | 98026956 | 98027026 | 0.01969232527 | 0.319 |
| ENSMUSG00000040044.11 | Orc3 | chr4 | - | 34614748 | 34614904 | 34614864 | 34614904 | 0.0339032019101 | 0.321 |
| ENSMUSG00000063652.10 | Slc22a21 | chr11 | - | 53960517 | 53960621 | 53960552 | 53960621 | 0.0482448818063 | 0.321 |
| ENSMUSG00000030189.15 | Ybx3 | chr6 | - | 131379353 | 131379480 | 131379357 | 131379480 | 0.000441613978911 | 0.322 |
| ENSMUSG00000029033.16 | Acap3 | chr4 | + | 155895491 | 155895695 | 155895491 | 155895601 | 0.047594582311 | 0.322 |
| ENSMUSG00000031533.4 | Mrps31 | chr8 | + | 22424346 | 22424509 | 22424346 | 22424420 | 0.000878348578856 | -0.326 |
| ENSMUSG00000039105.6 | Atp6v1g1 | chr4 | + | 63548584 | 63548745 | 63548584 | 63548699 | 0.0492679354108 | 0.332 |
| ENSMUSG00000097113.7 | Gm19705 | chr1 | + | 136677122 | 136677266 | 136677122 | 136677261 | 0.0062169711463 | -0.333 |
| ENSMUSG00000006931.15 | P3h4 | chr11 | - | 100411976 | 100412177 | 100412063 | 100412177 | 0.022877164925 | 0.335 |
| ENSMUSG00000030061.16 | Uba3 | chr6 | - | 97185331 | 97185546 | 97185482 | 97185546 | 0.0264007627707 | -0.336 |
| ENSMUSG00000007817.14 | Zmiz1 | chr14 | + | 25607356 | 25607470 | 25607356 | 25607398 | 0.0336275226959 | -0.336 |
| ENSMUSG00000039680.9 | Mrps6 | chr16 | + | 92059009 | 92059252 | 92059009 | 92059116 | 0.00150205555531 | -0.34 |
| ENSMUSG00000032411.15 | Tfdp2 | chr9 | + | 96231730 | 96231873 | 96231730 | 96231799 | 0.00604091248445 | 0.341 |
| ENSMUSG00000042104.18 | Uggt2 | chr14 | - | 119032690 | 119032787 | 119032741 | 119032787 | 0.00023860191048 | -0.346 |
| ENSMUSG00000067203.7 | H2-K2 | chr17 | - | 33977770 | 33978046 | 33977870 | 33978046 | 0.0471273169129 | -0.346 |
| ENSMUSG00000026197.12 | Zfand2b | chr1 | + | 75168927 | 75169071 | 75168927 | 75168962 | 0.00209107825423 | 0.348 |
| ENSMUSG00000032294.17 | Pkm | chr9 | + | 59657239 | 59657312 | 59657239 | 59657297 | 0.00340989825786 | 0.353 |
| ENSMUSG00000028832.11 | Stmn1 | chr4 | + | 134470102 | 134470179 | 134470102 | 134470175 | 0.0272326363912 | 0.354 |
| ENSMUSG00000034088.15 | Hdlbp | chr1 | - | 93442649 | 93442791 | 93442710 | 93442791 | 0.000256938474121 | 0.355 |
| ENSMUSG00000054717.7 | Hmgb2 | chr8 | + | 57511906 | 57512010 | 57511906 | 57512005 | 0.022073986907 | -0.355 |
| ENSMUSG00000002733.8 | Plekha3 | chr2 | + | 76692878 | 76693014 | 76692878 | 76692894 | 0.00228490513995 | -0.357 |
| ENSMUSG00000030727.12 | Rabep2 | chr7 | + | 126440347 | 126440595 | 126440347 | 126440446 | 0.00871618777259 | 0.358 |
| ENSMUSG00000024082.4 | Ndufaf7 | chr17 | + | 78937134 | 78937240 | 78937134 | 78937176 | 0.00519629319493 | -0.361 |
| ENSMUSG00000039382.12 | Wdr45 | chrX | + | 7722926 | 7723161 | 7722926 | 7722981 | 0.0317146477942 | 0.367 |
| ENSMUSG00000033991.8 | Ttc37 | chr13 | + | 76098733 | 76098886 | 76098733 | 76098833 | 0.011225155746 | -0.367 |
| ENSMUSG00000042298.18 | Ttc19 | chr11 | + | 62283223 | 62283377 | 62283223 | 62283255 | 0.00671188495912 | 0.368 |
| ENSMUSG00000099703.6 | Gm28285 | chr18 | + | 35557336 | 35557391 | 35557336 | 35557369 | 0.0337482297191 | 0.368 |

| | | | | | | | | | |
|---|---|---|---|---|---|---|---|---|---|
| ENSMUSG00000039128.13 | Cdc123 | chr2 | - | 5795531 | 5795710 | 5795543 | 5795710 | 0.0337482297191 | 0.37 |
| ENSMUSG00000019804.12 | Snx3 | chr10 | + | 42528310 | 42528402 | 42528310 | 42528362 | 0.00238835701455 | 0.377 |
| ENSMUSG00000048175.13 | Asb8 | chr15 | - | 98142040 | 98142111 | 98142044 | 98142111 | 0.0485268298537 | 0.379 |
| ENSMUSG00000022407.9 | Adsl | chr15 | + | 80967624 | 80967770 | 80967624 | 80967765 | 0.00646414516144 | 0.385 |
| ENSMUSG00000031782.15 | Coq9 | chr8 | + | 94838414 | 94838608 | 94838414 | 94838502 | 0.00991896417098 | 0.393 |
| ENSMUSG00000006494.11 | Pdk1 | chr2 | + | 71887685 | 71887824 | 71887685 | 71887817 | 0.0282964413054 | 0.394 |
| ENSMUSG00000002416.13 | Ndufb2 | chr6 | + | 39596450 | 39596514 | 39596450 | 39596480 | 0.0307132695742 | 0.402 |
| ENSMUSG00000026743.16 | Mllt10 | chr2 | + | 18082510 | 18082748 | 18082510 | 18082725 | 0.00633861367527 | 0.419 |
| ENSMUSG00000068739.13 | Sars | chr3 | - | 108427852 | 108427902 | 108427855 | 108427902 | 0.0351344122009 | -0.42 |
| ENSMUSG00000026519.16 | Tmem63a | chr1 | + | 180946365 | 180946476 | 180946365 | 180946439 | 0.0143104466281 | -0.429 |
| ENSMUSG00000087433.1 | Gm14167 | chr2 | - | 151542291 | 151542348 | 151542306 | 151542348 | 0.000429783304955 | 0.431 |
| ENSMUSG00000021713.8 | Ppwd1 | chr13 | - | 104219620 | 104219837 | 104219691 | 104219837 | 0.0130519035238 | -0.433 |
| ENSMUSG00000037772.13 | Mrpl23 | chr7 | + | 142533160 | 142533241 | 142533160 | 142533191 | 0.000256938474121 | -0.44 |
| ENSMUSG00000024346.5 | Pfdn1 | chr18 | - | 36416874 | 36417030 | 36416946 | 36417030 | 0.000281280956745 | -0.443 |
| ENSMUSG00000039168.15 | Dap | chr15 | + | 31224559 | 31224710 | 31224559 | 31224689 | 0.0251840297252 | 0.446 |
| ENSMUSG00000027357.16 | Crls1 | chr2 | + | 132847733 | 132847904 | 132847733 | 132847880 | 0.0203756172258 | 0.451 |
| ENSMUSG00000003865.16 | Gys1 | chr7 | + | 45447916 | 45448028 | 45447916 | 45448011 | 0.0120636538856 | 0.452 |
| ENSMUSG00000028453.10 | Fancg | chr4 | - | 43005206 | 43005312 | 43005210 | 43005312 | 0.0201074569362 | -0.455 |
| ENSMUSG00000050373.13 | Snx21 | chr2 | + | 164791557 | 164791578 | 164791557 | 164791573 | 0.000685836297499 | 0.457 |
| ENSMUSG00000006931.15 | P3h4 | chr11 | - | 100412014 | 100412177 | 100412063 | 100412177 | 0.00485049809489 | 0.462 |
| ENSMUSG00000096054.3 | Syne1 | chr10 | - | 5426212 | 5426275 | 5426254 | 5426275 | 0.00384213368502 | 0.482 |
| ENSMUSG00000048495.16 | Tyw5 | chr1 | - | 57393570 | 57393691 | 57393584 | 57393691 | 0.00028579545779 | -0.483 |
| ENSMUSG00000024392.17 | Bag6 | chr17 | + | 35146025 | 35146206 | 35146025 | 35146169 | 0.00357650588192 | 0.489 |
| ENSMUSG00000042700.15 | Sipa1l1 | chr12 | + | 82268032 | 82268196 | 82268032 | 82268157 | 0.031167549974 | 0.493 |
| ENSMUSG00000050373.13 | Snx21 | chr2 | + | 164791557 | 164791617 | 164791557 | 164791573 | 0.000181107636209 | 0.498 |
| ENSMUSG00000005774.12 | Rfx5 | chr3 | + | 94954762 | 94954931 | 94954762 | 94954877 | 0.000579069961823 | 0.507 |
| ENSMUSG00000051550.10 | Zfp579 | chr7 | - | 4985177 | 4986486 | 4986400 | 4986486 | 0.0144859804085 | -0.519 |
| ENSMUSG00000027678.17 | Ncoa3 | chr2 | + | 166049139 | 166049317 | 166049139 | 166049236 | 0.0133733831538 | 0.537 |
| ENSMUSG00000024392.17 | Bag6 | chr17 | + | 35146025 | 35146206 | 35146025 | 35146169 | 0.00381492617223 | 0.544 |
| ENSMUSG00000057133.14 | Chd6 | chr2 | - | 161059699 | 161059806 | 161059790 | 161059806 | 0.0110951024435 | 0.565 |
| ENSMUSG00000053617.11 | Sh3pxd2a | chr19 | - | 47272578 | 47272803 | 47272706 | 47272803 | 0.00907882403216 | -0.565 |
| ENSMUSG00000090112.8 | Shprh | chr10 | + | 11205846 | 11205980 | 11205846 | 11205918 | 0.000878348578856 | 0.606 |
| ENSMUSG00000019539.11 | Rcn3 | chr7 | - | 45091888 | 45092123 | 45092093 | 45092123 | 0.000267797353554 | -0.621 |
| ENSMUSG00000024900.4 | Cpt1a | chr19 | + | 3374681 | 3374783 | 3374681 | 3374762 | 0.0125448537766 | 0.64 |
| ENSMUSG00000034674.18 | Tdg | chr10 | + | 82636968 | 82637138 | 82636968 | 82637104 | 0.0272326363912 | 0.687 |
| ENSMUSG00000034269.12 | Setd5 | chr6 | + | 113111116 | 113111272 | 113111116 | 113111243 | 0.0032447775349 | 0.712 |
| ENSMUSG00000078441.10 | Scamp4 | chr10 | + | 80608802 | 80608854 | 80608802 | 80608850 | 0.009762501811 | 0.73 |
